# EZH2 alters aggressive variant prostate cancer subtype evolution, but is not required for neuroendocrine prostate cancer

**DOI:** 10.1101/2025.05.13.653549

**Authors:** Justine J. Jacobi, Kristine M. Wadosky, Neha Jaiswal, Xiaojing Zhang, Yanqing Wang, Prashant Singh, Jie Wang, Eduardo Cortes Gomez, Jianmin Wang, Bo Xu, Paloma Cejas, Shweta Kukreja, Henry W. Long, David W. Goodrich

**Author notes:** These authors contributed equally as first authors (Justine J. Jacobi, Kristine M. Wadosky).

## Abstract

Advanced prostate cancer (PrCa) remains a leading cause of cancer-related death among men because nearly all patients progress on standard therapy that targets androgen receptor (AR) signaling. An important mechanism driving therapeutic resistance is lineage plasticity, the ability of PrCa cells to reprogram into lineage variants like neuroendocrine (NE) PrCa that no longer depend on AR signaling. The histone methyltransferase EZH2 has been implicated in PrCa lineage plasticity and EZH2 inhibitors are being evaluated clinically for the treatment of PrCa. Here we test how *Ezh2* impacts the evolution of PrCa lineage plasticity using genetically engineered mice and human patient data. Reduced *Ezh2* expression diversifies the evolution of PrCa lineage variants, including a newly discovered variant with high expression of neurofilament genes, in both mice and humans. Lineage variant diversification is associated with activation of Klf transcription factor programs and AR cistrome reprogramming that is normally repressed by Ezh2. These findings advance the understanding of PrCa lineage plasticity and have implications for using EZH2 as a target for PrCa therapy.

## Introduction

Prostate cancer (PrCa) is a leading cause of cancer associated mortality in men^1-3^. Standard of care for patients with advanced disease include therapies designed to block androgen receptor (AR) signaling. While effective initially, virtually all patients progress on these therapies to a disease state commonly referred to as castrate resistance prostate cancer (CRPC)^4,5^. Genetic mechanisms underlying CRPC include alterations in AR that restore signaling, or activation of signaling pathways that compensate for reduced AR signaling^6^. A more recently appreciated mechanism driving acquired therapeutic resistance is lineage plasticity, transcriptional reprogramming of PrCa cells to aggressive variant prostate cancers (AVPC) like neuroendocrine PrCa (NEPC) that are no longer dependent on AR signaling^7-11^. Lineage plasticity is observed currently in about 15-25% of CRPC patients, but this fraction is increasing as more potent AR targeted therapies are deployed in the clinic^12-14^. Mechanisms underlying PrCa lineage plasticity are incompletely defined and therapies effective in treating or preventing AVPC are not available. Lineage plasticity confounds the clinical management of PrCa because it is unclear if different AVPC subtypes exhibit similar or unique therapeutic vulnerabilities. Patients diagnosed with AVPC thus have very poor prognosis^15,16^.

Multi-omic analyses of clinical specimens from CRPC patients have discovered a range of distinct AVPC variants classified using various criteria^17-23^. In the criteria of Labreque et al.^21^, CRPC lineage variants include amphicrine that express both AR and neuroendocrine markers (AR+/NE+), AR-low (reduced AR, NE−), double negative (AR−/NE−), and NEPC (AR−/NE+)^18,21^. The NEPC subtype exhibits additional heterogeneity defined by expression of pro-neuronal transcription factors ASCL1 and/or NEUROD1 that act as key oncogenic drivers in aggressive neuroendocrine cancers^19,24,25^. Given the dearth of experimental models that recapitulate this phenotypic diversity, the evolutionary relationships between these PrCa lineage variants and the molecular factors governing their development remain largely unknown.

Genetic alterations in *RB1*, *TP53*, *PTEN*, and MYCN are highly recurrent in human NEPC and have been functionally linked to PrCa lineage plasticity in experimental models*^11,12,15,16,26-35^*. Not all PrCa containing these genetic alterations convert to NEPC, however, and NEPC conversion is rare among similarly mutated PrCa cells^20,32,36^. Accumulating evidence thus highlights the importance epigenetic mechanisms, including DNA methylation and histone modifications, in driving PrCa lineage plasticity^10,11,25,37-40^. Enhancer of zeste homolog 2 (EZH2), for example, has been implicated in PrCa lineage plasticity and acquired therapeutic resistance^27,32,41-44^. EZH2 inhibitors are thus being evaluated clinically for the treatment of CRPC (e.g. NCT04846478, NCT04179864, NCT03460977). *EZH2* encodes the catalytic subunit of the Polycomb Repressive Complex 2 (PRC2) that catalyzes histone 3 lysine 27 trimethylation (H3K27me3) to generate transcriptionally repressive chromatin. However, EZH2 inhibitors only modestly affected tumor growth in preclinical NEPC models^42^. How EZH2 impacts lineage plasticity and the evolution of different AVPC variants is unknown.

Leveraging genetically engineered mouse models (GEMM) and publicly available genomic data from patient specimens, we investigated how *Ezh2* deficiency impacts the evolution of PrCa lineage plasticity in vivo. Multi-omics analysis revealed that *Ezh2* constrains the evolution of PrCa lineage variants, favoring development of the *Ascl1* expressing NEPC subtype. *Ezh2* deficiency thus increases the diversity of AVPC lineage variants detected, but is not required for NEPC development. Notably, evolutionary relationships between PrCa lineage variants and transcription factor networks have been identified that likely dictate their unique phenotypic states. PrCa lineage variants observed in mice and humans are similar at the transcriptional level, demonstrating clinical relevance. These results and the mouse models described advance the understanding of PrCa lineage plasticity, will be useful in discovering therapeutic approaches effective in treating AVPC variants, and inform the use of EZH2 inhibitors for treating PrCa.

## Results

### Effects of Ezh2 deficiency on PrCa progression in vivo

We and others have described genetically engineered mice in which deletion of tumor suppressor genes specifically in prostate epithelium initiates development of PrCa that recapitulates lineage plasticity and NEPC development analogous to that in human patients^32^. *Pten* deletion alone initiates prostate adenocarcinoma that does not progress to NEPC^45^. Combined deletion of *Pten* and *Rb1* (DKO) initially presents as adenocarcinoma, but progresses to metastatic NEPC after long latency. Deletion of *Pten*, *Rb1*, and *Trp53* (TKO) accelerates progression of adenocarcinoma to NEPC. In contrast, combined deletion of *Pten* and *Trp53* causes development of sarcomatoid PrCa, rarely observed in human patients, with some foci of NE differentiation^46,47^. *Rb1* loss is thus a rate limiting event in the development of NEPC, potentially because Rb1 controls the activity of key transcription factors and epigenetic regulators like Ezh2^48-51^. To investigate the impact of *Ezh2* deficiency, floxed alleles of the mouse Ezh2 gene^52^ were bred into the DKO model to generate DKO_EP and DKO_EE mice with one or two *Ezh2* floxed alleles, respectively **(Figure 1A)**. PCR genotyping of tail or prostate DNA confirmed expected prostate specific Cre mediated gene deletion (**Figure S1A**). As expected, DKO_EE prostate tissue had reduced *Ezh2* RNA (**Figure S1B**) and protein **(Figure 1B-C)** compared to tissue from DKO mice. H3K27me3 levels were also reduced (**Figure 1D**), but remained detectable, likely because Ezh1 can catalyze the H3K27me3 modification with reduced specific activity^52,53^.

**Figure 1:**
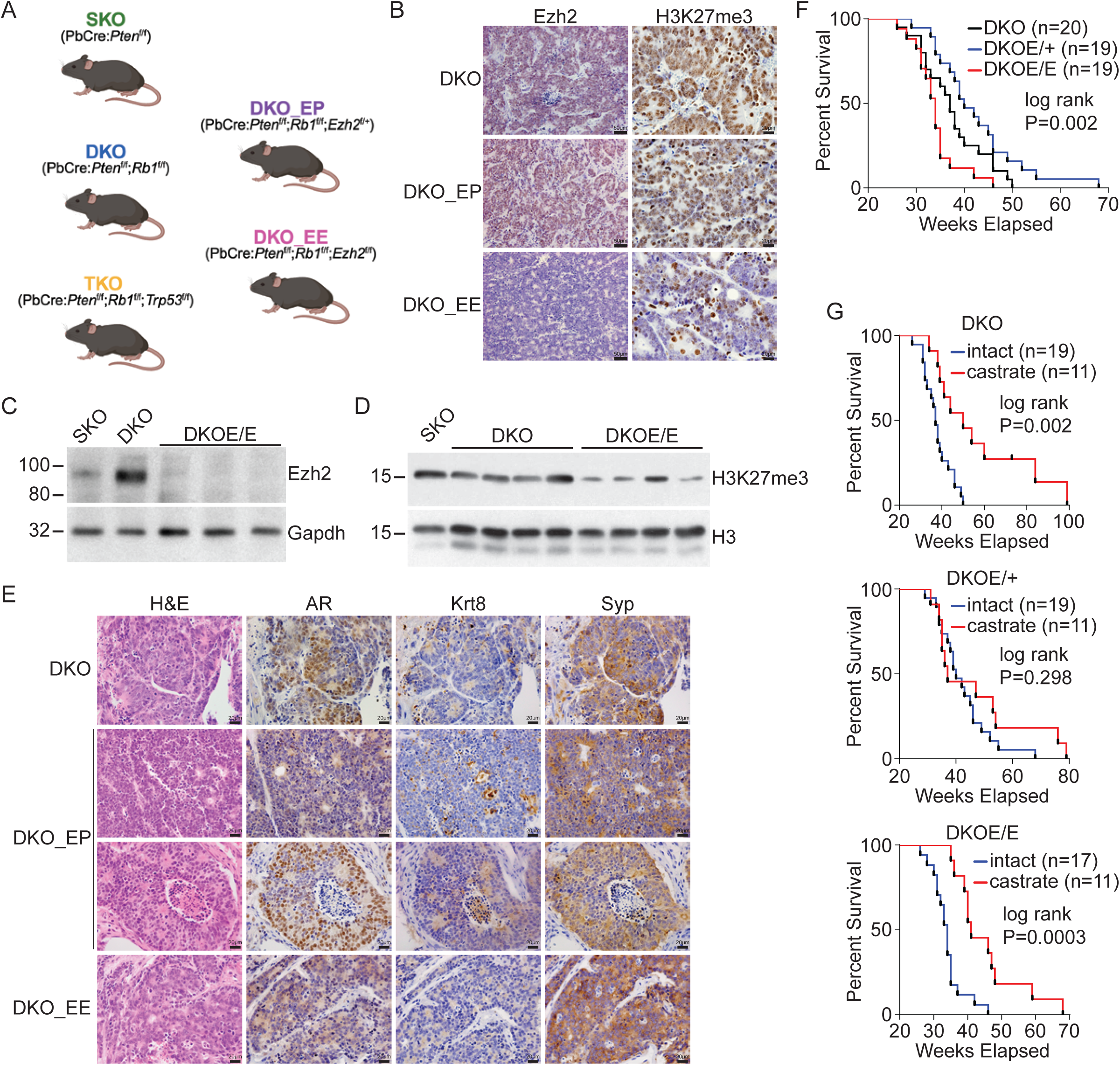
*Ezh2* deficiency alters PrCa phenotype in vivo. A) Schematic summarizing the genetically engineered mice used in the present study. B) Immunostaining of PrCa tissue sections from mice of the indicated genotypes showing marked decreases in Ezh2 protein and in the H3K27me3 mark catalyzed by Ezh2. Scale bars represent 50µm or 20 µm as indicated. C) and D) Western blot analysis of PrCa tissue from mice of the indicated genotypes for Ezh2 or H3K27me3. Gapdh or histone H3 (H3) serve as protein loading controls. E) Immunostaining of PrCa tissue sections from mice of the indicated genotypes for Ar, the epithelial marker Krt8, and the neuroendocrine marker Syp. Scale bars represent 20 µm. F) Survival of mice of the indicated genotypes is plotted in a Kaplan-Meier curve with the indicated sample sizes (n). Survival is significantly different between genotypes by log-rank analysis. G) Mice of the indicated genotypes were castrated at 12 weeks of age and survival monitored as in F). Castration significantly lengthened survival for DKO and DKO_EE mice, but not DKO_EP mice.

End stage mice were necropsied and tissues collected to assess primary tumors and distant metastases. DKO mice developed heterogeneous end stage tumors with mixed AR, KRT8 (luminal marker), and SYP (NE marker) expression as described previously (**Figure 1E**)^32^. DKO_EP mice developed end stage PrCa with two phenotypes distinguishable by immunostaining, AR-/NE+ NEPC and AR+/NE+ amphicrine PrCa whose relative proportions varied per mouse. End stage tumors in DKO_EE mice were heterogeneous but primarily AR-/NE+ NEPC. NEPC was detectable in DKO_EE mice as young as 12 weeks of age (**Figure S1C**), sooner than in DKO mice where NEPC is typically detectable in mice older than 20 weeks of age^32^. *Ezh2* was thus not required for, and likely accelerated, NEPC development in this genetic context. Tumors from all three genotypes metastasize to clinically relevant sites like the lung, liver, and bone (**Figure S1D**). Metastatic lesions developing in DKO and DKO_EP mice had a mix of amphicrine and NEPC phenotypes, demonstrating both variants had metastatic potential (**Figure S1E**). The survival of DKO, DKO_EP, and DKO_EE mice were significantly different (log-rank P=0.002) with median survival of 37, 40, and 34 weeks respectively (**Figure 1F**), with marked variation in survival among mice within each genotype (**Figure S1F**). We note that *Ezh2* deletion in prostate epithelial cells in vivo, alone or in combination with *Rb1* loss, did not cause detectable PrCa development (**Figure S1G**). *Ezh2* loss in SKO mice did not detectably affect PrCa phenotype, and these tumors did not progress to NEPC (**Figure S1H**).

Separate cohorts of mice were surgically castrated to test if *Ezh2* deficiency altered responses of developing PrCa to androgen deprivation therapy. Castration of DKO mice significantly extended median survival from 37 to 50 weeks, consistent with our prior published work (**Figure 1G**)^32^. Castration of DKO_EE mice extended median survival from 34 to 41 weeks while castration of DKO_EP mice did not alter survival. The survival curve for DKO_EP mice appeared biphasic, however, suggesting distinct tumor phenotypes detected in these mice, NEPC and amphicrine, might have different responses to castration. Genetic *Ezh2* deficiency, therefore, did not improve outcomes for androgen deprivation therapy in vivo.

### Ezh2 deficiency induces lineage-specific transcriptional reprogramming

To investigate the impact of *Ezh2* loss on transcription, normal and PrCa tissue was analyzed by mRNA sequencing (RNA-seq). Principal component analysis revealed that transcriptional patterns in DKO_EP and DKO_EE tumors were distinct from other tumor genotypes, excepting overlap with some DKO tumors **(Figure 2A)**. Consistent with immunostaining analysis, gene expression patterns in DKO, DKO_EP, DKO_EE, and TKO tumors all showed increased expression of genes associated with NEPC and reduced expression of genes associated with AR signaling and adenocarcinoma (**Figure 2B**)^13,19^. Morel et al.^54^ reported that treating PrCa cells with EZH2 inhibitors induced expression of inflammation and interferon response genes. Expression of these genes was also elevated in DKO_EE PrCa relative to DKO PrCa (**Figure S2A**). Across most samples, an inverse relationship between *Ezh2* and *Ezh1* expression was observed, suggesting possible compensation or lineage-specific regulation of *Ezh1/2* expression **(Figure 2C)**. Notable gene expression features distinguishing DKO_EE tumors included increased expression of *Neurod1* and Neurod1 regulated genes like *Syt1* (**Figure 2C**). Immunostaining of DKO_EE tumor sections confirmed Neurod1 protein expression (**Figure 2D**). Cells staining for nuclear Neurod1 were intermixed with Neurod1 negative cells within the same tumor foci, however, suggesting the transcription factors driving NE differentiation in these tumors were heterogeneous. Consistent with this data, master regulator analysis indicated that DKO_EE tumors exhibited increased Neurod1 transcription factor activity as well as increased activity for transcription factors like Ascl1, Foxa2, and Prox1 (**Figure 2E, Table S3)**. DKO_EE tumors also exhibited activation of developmental and neurogenesis transcription factors like Germ Cell Nuclear Factor (Nr6a1) as well as the Pou2f and Klf transcription factor families. Interestingly, *Ezh2* loss decreased the activity of transcription factors like Mycn, Onecut2, and Sox2 that have been implicated in NEPC. DKO_EP tumors, in contrast, exhibited unique up regulation of *Ar* and *Onecut2* expression. Master regulator analysis demonstrated increased activity for Ar, Onecut2, Ascl1, and Klf transcription factors (**Figure 2E, Table S3**). Both DKO_EE and DKO_EP tumors showed negative enrichment of Rest activity, a transcriptional repressor of NE differentiation. DKO_EE and DKO_EP tumors also exhibited decreased expression of genes associated with oxidative phosphorylation (**Figure S2B**). Castration reduced expression of these oxidative phosphorylation genes further (**Figure S2C**). Reduced expression of oxidative phosphorylation genes correlated with reduced activity of Myc family transcription factors known to directly regulate these metabolic genes (**Figure 2E**)^55^.

**Figure 2:**
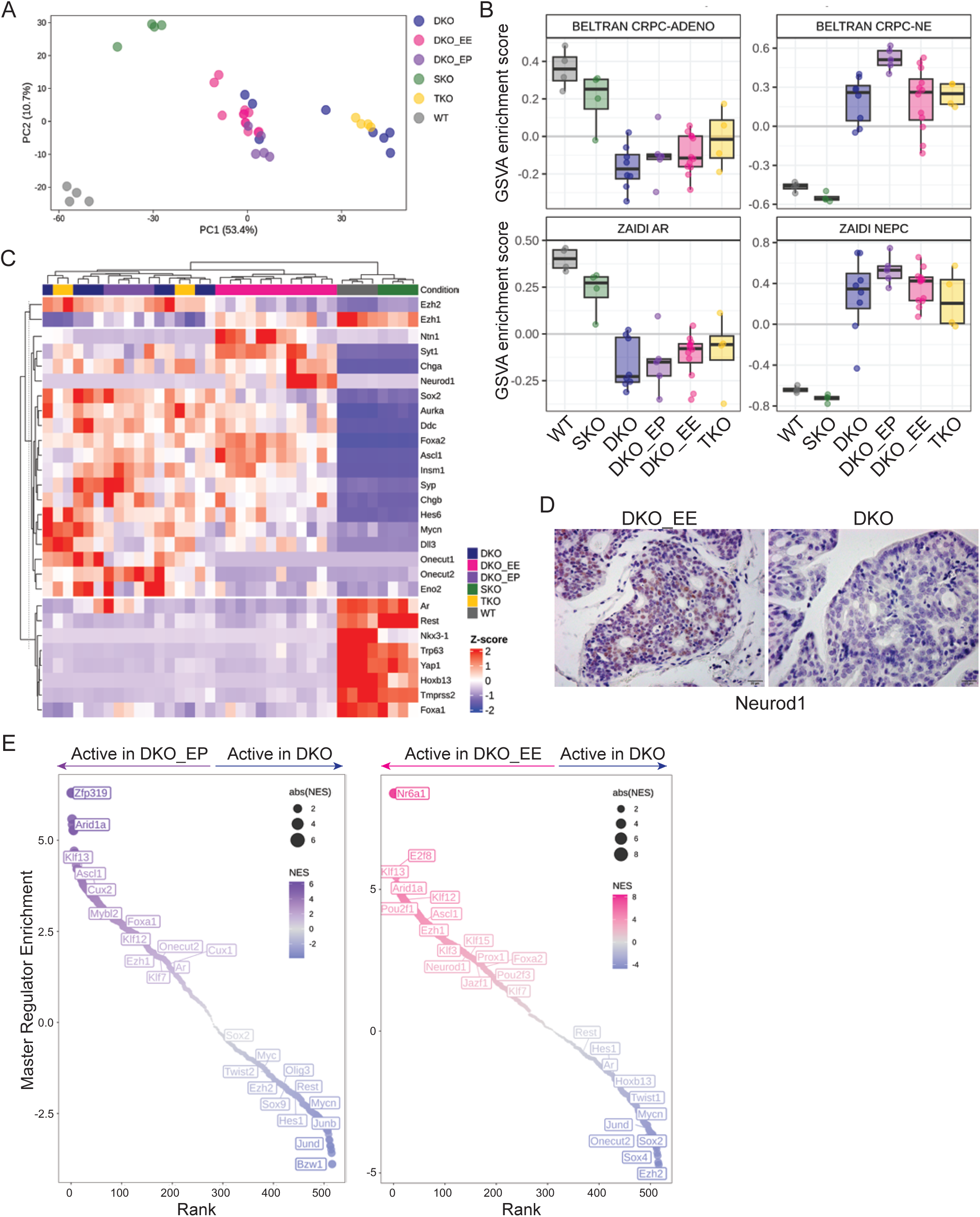
*Ezh2* deficiency alters lineage specific transcriptional reprogramming during PrCa progression. A) RNA-seq data was generated from prostate tissue of the indicated genotypes, the data was analyzed by principal component analysis, and samples color coded by genotype plotted based the first two principal components. B) RNA-seq data was analyzed by gene set variation analysis using the indicated published gene sets^13,19^, and scores for each sample plotted by genotype. C) RNA-seq data, coded by genotype, was hierarchically clustered and presented as a heat map using genes that distinguish prostate adenocarcinoma from NEPC. D) Immunostaining of PrCa tissues sections with the indicated genotypes confirms increased Neurod1 protein expression in some tumor foci upon *Ezh2* loss. E) Master regulator analysis of the RNA-seq data comparing the indicated genotypes indicates active transcription factors in the relevant samples.

Tumors developing in TKO mice were treated by castration plus administration of CPI1688 to test effects of pharmacological Ezh2 inhibition on PrCa progression in vivo. CPI1688 treatment did not significantly alter the overall survival of castrated TKO mice (**Figure S2D**). While PrCa gene expression was heterogeneous between mice, effects of CPI1688 mirrored some effects caused by genetic *Ezh2* deficiency. For example, *Neurod1* expression was elevated in PrCa arising in three of four CPI1688 treated mice, relative to control mice (**Figure S2E-F, S4D**). Overall, CPI1688 increased expression of genes associated with adenocarcinoma and Ar signaling, but had mixed effects on genes associated with NEPC (**Figure S2G**).

### Ezh2 loss alters the evolution of aggressive variant prostate cancer subtypes

To quantitate the impact of *Ezh2* deficiency on AVPC subtype evolution at the level of individual cells, single-cell RNA sequencing (scRNA-seq) analysis was performed on GEMM PrCa tissue. UMAP projections of over 210,000 individual cell transcriptomes from SKO, DKO, TKO, DKO_EP, DKO_EE and wild type prostate tissue revealed clear genotype-specific differences in gene expression **(Figure 3A)**. Broad cell type labeling (**Figure 3B**) was validated using anchor-based reference mapping **(Figure S3A)^56^** and well-established marker genes from the published literature including *Epcam/Krt8* (luminal epithelial), *Krt5* (basal epithelial), *Ascl1* (NE), Col1a1 (mesenchyme), Pecam1 (endothelial), Svs1 (seminal vesicle), and Ptprc/Cd45 (lymphoid) **(Figure 3C)**.

**Figure 3:**
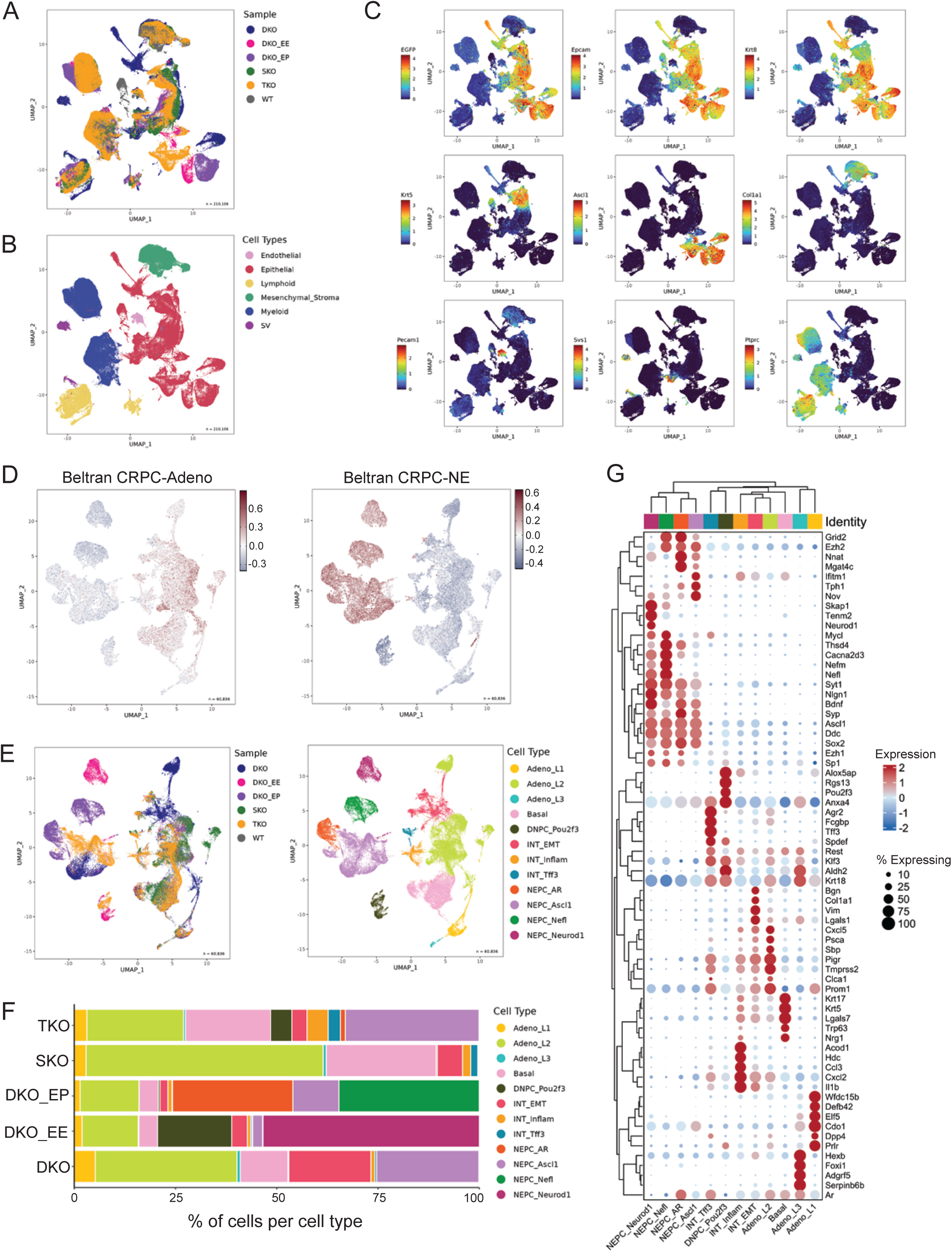
*Ezh2* deficiency alters PrCa lineage variant evolution. A) Prostate tissue from mice of the indicated genotypes was analyzed by scRNA-seq, clustered based on transcriptional profiles, and plotted as a uniform manifold approximation and projection (UMAP). B) The broad cell type of each analyzed cell was assigned using anchor-based reference mapping and displayed in color coded fashion on the UMAP. C) Cells on the UMAP are color coded based on the expression of well-established marker genes characteristic of different cell types. D) Cre recombined cells identified by EGFP expression were analyzed separately and those cells exhibiting expression of either CRPC Adenocarcinoma or CRPC Neuroendocrine gene signatures were projected on a UMAP. E) The UMAP of EGFP expressing PrCa cells color coded by genotype (left panel) or PrCa cell type (right panel). F) The relative proportion of PrCa cell types making up the total PrCa cells analyzed for each genotype. G) Genes whose expression characterize different PrCa cell types based on hierarchical clustering, with expression level coded by color and the fraction of expressing cells coded by size.

All GEMMs incorporated an enhanced green fluorescent protein (EGFP) based Cre recombination reporter gene^57^, thus EGFP expression was used to identify Cre recombined prostate epithelial cells. EGFP expression correlated with expression of epithelial markers (**Figure 3C, S3B**), confirming the validity of this approach. Focusing on EGFP+/Epcam+ cells, approximately 60,000 cells were identified that displayed enrichment for either CRPC Adenocarcinoma or CRPC Neuroendocrine gene signatures **(Figure 3D)**, identifying them as PrCa cells. PrCa cells in this dataset included those with gene expression analogous to each of the four normal epithelial subtypes identified previously in the mouse prostate (**Figure S3C**) (Adeno_L1, Adeno_L2, Adeno_L3, Basal)^58^. The Adeno_L2 population was predominant in SKO PrCa and was the most prevalent adenocarcinoma phenotype across genotypes (**Figure 3E-G**). Basal epithelial-like PrCa cells were notably diminished among *Ezh2* deficient PrCa suggesting Ezh2 activity supports this lineage variant. Previously discovered intermediate PrCa cell types such as inflammatory (INT_Inflam), EMT-like (INT_EMT), and intestinal lineage-like (INT_Tff3) were also detected^19,56^.

All PrCa cells exhibiting NE differentiation expressed the neuronal lineage driver *Ascl1* and its target gene, *Ddc* (**Figure 3G, S3B)**. NE PrCa cells from DKO and TKO tumors primarily aligned with the well described Ascl1+/Sox2+/Ezh2+ NEPC subtype (NEPC_Ascl1). *Ezh2* deficient PrCa revealed additional lineage variants. DKO_EP PrCa had an increased proportion of NEPC_AR cells resembling the human amphicrine PrCa variant (Ar+, NE+). These cells did not express canonical Ar regulated genes like *Tmprss2* **(Figure 3G, S3B)**, however, suggesting Ar activity may be reprogrammed to regulate different genes in this context (see below). *Ezh2* deficient PrCa cells also revealed a novel NEPC subtype characterized by high expression of neurofilament genes like *Nefl* and *Nefm* (NEPC_Nefl) **(Figure 3G, S3B)**. Nefl protein expression was confirmed by immunostaining of both DKO_EP and DKO_EE PrCa (**Figure S3D**). Nefl immunostaining was patchy but relatively homogeneous within tumor and metastatic foci where it was expressed. In line with bulk RNA-seq data, DKO_EE tumors gave rise to an NEPC subtype expressing *Neurod1* (NEPC_Neurod1) **(Figure 3E-G, S3B)** that has also been observed in human patient specimens^25^. DKO_EE also exhibited an increased fraction of PrCa cells with negative enrichment for both CRPC Adenocarcinoma and Neuroendocrine gene expression signatures **(Figure 3C)** while uniquely expressing tuft cell lineage markers Pou2f3 and the Pou2f3 regulated gene *Rgs13* (DNPC_Pou2f3) **(Figure 3G, S3B)**. It is notable that DNPC_Pou2f3 cells were also detected in TKO mice that retain wild type *Ezh2* alleles, but TKO DNPC_Pou2f3 cells also exhibited low *Ezh2* expression (**Figure S3E**). DNPC_Pou2f3 cells resembled the AR-/NE- double negative mCRPC lineage variant detected in human patients that is associated with the worst clinical prognosis in the West Coast Dream Team patient cohort^18^. A Pou2f3+/Rgs13+ subtype has also been detected in human small cell lung cancers^59,60^. These findings highlight the critical role of Ezh2 in shaping the evolution of AVPC lineage variants.

Hallmark gene set enrichment analysis was performed to compare the phenotypes of different PrCa lineage variants detected (**Figure S4A**), and subtype specific differences were noted. The NEPC subtypes varied in their expression of genes related to metabolic processes, suggesting different lineage variants may exhibit distinct metabolism. *Ezh2* loss was associated with increased inflammatory gene expression in bulk RNA-seq data, but scRNA-seq data indicated this increase was largely confined to adenocarcinoma, basal, Int_EMT, and Int_Inflam subtypes as both DNPC_Pou2f3 and NEPC variants showed reduced expression of these inflammatory genes (**Figure S4A-B**). Consistent with prior reports^56,61^, this suggests an inflammatory plastic state serves as an important intermediate during reprogramming to variant AVPC lineage states.

To assess the clinical relevance of *EZH2* expression for human PrCa, RNA-seq data from the SU2C West Coast Dream Team metastatic CRPC patient dataset was analyzed^17^. *EZH2* expression was high in the NEPC subtype expressing *ASCL1* as expected. However, *EZH2* expression was significantly lower in other PrCa lineage variants including DNPC (AR-/NE-) and amphicrine (AR+/NE+)(**Figure 4A-B**). Labrecque et al.^21^ demonstrated that human amphicrine and AR-/NE+ PrCa exhibited comparable expression of REST transcription factor repressed genes (Neuro I), but amphicrine PrCa expressed lower levels of core NE transcription factors like *SOX2* (Neuro II). Reduced *EZH2* expression in human PrCa samples was correlated with reduced Neuro II gene expression (**Figure 4B**), consistent with mouse data showing *Ezh2* loss reduced expression of Neuro II genes like *Sox2*, *MycN*, and *Onecut2* (**Figure 2C**). Additionally, Labrecque et al. identified DNPC (AR-/NE-; Arlow/NE-) tumors with squamous differentiation. As with the mouse DNPC_Pou2f3 lineage variant, these human DNPC expressed lower levels of *EZH2* and uniquely expressed *POU2F3* and KLF family transcription factors. We extracted the top 50 differentially expressed genes from mouse lineage variants and assessed their gene activity in each human specimen using GSVA (**Figure 4C, Table S4**). Human DNPC gave the highest expression for the mouse DNPC_Pou2f3 signature genes. Human AR-/NE+ showed the highest activity for all the mouse NE variants, including NEPC_Nefl, suggesting this new AVPC lineage variant might also be detected in human PrCa.

**Figure 4:**
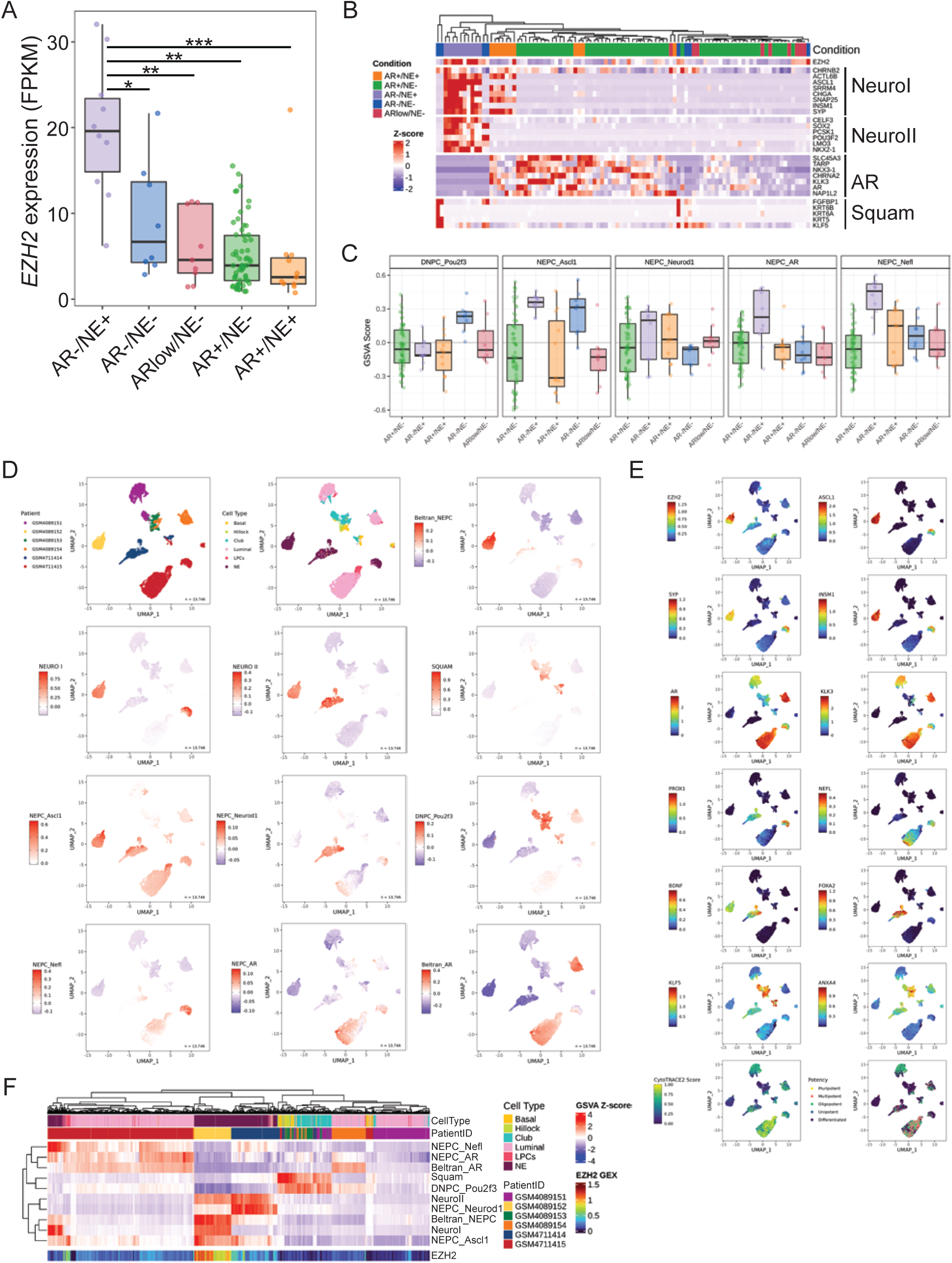
*EZH2* expression is low in most human AVPC lineage variants. A) RNA-seq data derived from human metastatic CRPC specimens classified by AVPC subtype were analyzed for *EZH2* expression. Asterisks indicate significant differences with P< 0.05, 0.01, or 0.001, respectively. B) The heatmap shows the human samples hierarchically clustered with AVPC subtypes distinguished by gene expression signatures described in Labrecque et al^21^. C) The top 50 differentially expressed genes in the mouse AVPC lineage variants were used to score the human metastatic CRPC samples by GSVA, with scores segregated based on their assigned human AVPC subtype. D) scRNA-seq data generated from CRPC patient biopsies^62^ was analyzed and presented as UMAPs color coded based on sample identity, PrCa cell subtype, and the activity of either established human gene expression signatures or gene expression signatures dervied from mice in this study. E) UMAPs of the human scRNA-seq data color coded for expression of genes characterizing different PrCa subtypes. F) A heatmap showing the activity of human and mouse gene expression signatures within human CRPC specimens highlighting the similarity between human and mouse AVPC subtypes.

Single cell RNA-seq data collected by Dong et al.^62^ from CRPC patient biopsies was thus analyzed to dissect AVPC heterogeneity (**Figure 4D-F**). This dataset contains six patients with varying populations of luminal, basal, hillock/club, luminal progenitor (LPC), and neuroendocrine (NE) PrCa subtypes. Of the six patients, three had evidence of NEPC. Clear intra- and inter-tumoral heterogeneity existed between the NEPC samples. *EZH2* expression was largely confined to NEPC from one patient (GSM4089152) that also had the highest expression of the Beltran NE signature, *ASCL1,* as well as the NEUROI and NEUROII signatures. The mouse NEPC_Ascl1 gene signature was most enriched in this patient sample, defining it as an ASCL1 NEPC subtype. NEPC in two other patients exhibited low *EZH2* expression. In one of these patients (GSM4711414), expression of the human NEURO II and mouse NEPC_Neurod1 gene signatures was high but expression of the NEURO I genes was low. This patient specimen expressed genes like *BDNF* that define it as the NEUROD1 NEPC subtype. Interestingly, the remaining NEPC patient sample (GSM4711415) was enriched for the mouse NEPC_Nefl gene expression program, including high expression of genes like *PROX1* and *NEFL,* but low expression of *EZH2.* This patient sample also included PrCa cells with an amphicrine AR+/NE+ phenotype as well as luminal adenocarcinoma PrCa cells expressing NEFL. This PrCa sample thus exhibited considerable plasticity. Consistent with this notion, PrCa cells in this specimen exhibited the highest plasticity scores as measured by CytoTRACE2 (**Figure 4E**), an scRNA-seq analysis algorithm designed to measure developmental potency, plasticity, and differentiation states^63^. Cells expressing the mouse DNPC_Pou2f3 or human Squam DNPC gene expression signatures were detected in 5 of six patients, although typically as a minority population. The exception was PrCa GSM4089153 that was composed primarily of cells enriched for mouse DNPC_Pou2f3, defining it as a DNPC subtype. Analogous AVPC variants were thus detected in both mice and humans, and increased AVPC subtype diversity was associated with lower EZH2 expression in both species.

### Ezh2 deficiency diversifies transcription factor programs active during AVPC evolution

To measure lineage plasticity in mouse PrCa samples, scRNA-seq data was also analyzed by CytoTRACE2. DKO_EP and DKO_EE PrCa cells exhibited higher plasticity than DKO PrCa cells (**Figure 5A**), indicating Ezh2 constrained overall PrCa lineage plasticity. Among PrCa variants, NEPC_Nefl and NEPC_Ascl1 had higher plasticity (less differentiated) while NEPC_Neurod1 and NEPC_Ar had lower plasticity (more differentiated) **(Figure 5B)**. The DNPC_Pou2f3 subtype exhibited intermediate plasticity. Gene set enrichment analysis (GSEA) was performed on the top 300 genes most significantly correlated with high and low plasticity cells (**Table S5**). Gene expression that correlated with high plasticity was enriched for pathways related to stemness, chromatin remodeling, cell cycle, and progenitor cell functions (**Figure 5C**). Gene expression that correlated with low plasticity was enriched for neuroendocrine or tuft cell lineages, neuronal signaling, and brain developmental programs. Transcriptional plasticity scores thus align with expected biological programs in plastic stem/progenitor like PrCa cells versus those differentiating towards terminal AVPC lineage states.

**Figure 5:**
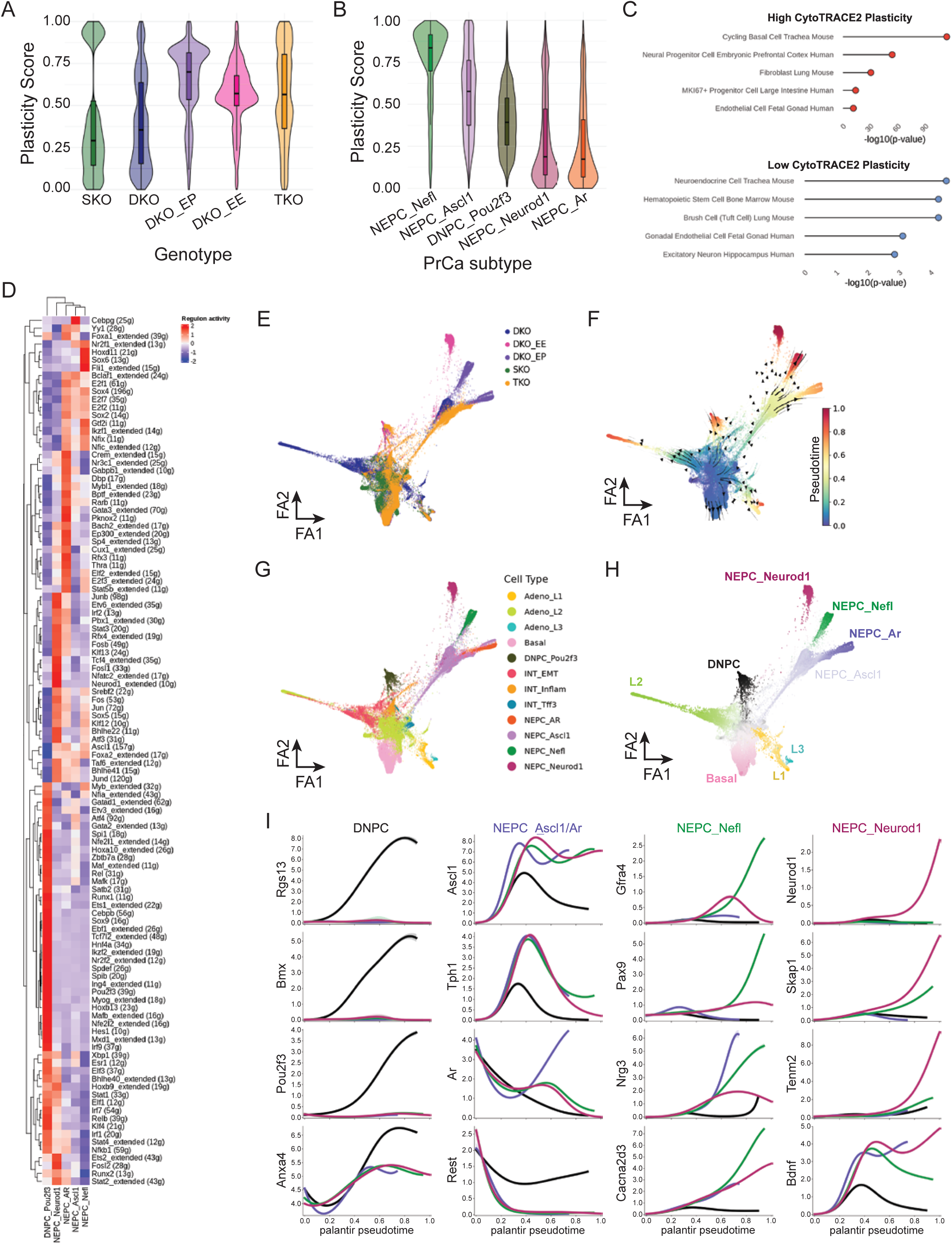
*Ezh2* deficiency diversifies transcriptional programs active during AVPC subtype evolution. A) scRNA-seq data generated from PrCa cells segregated by genotype were analyzed by CytoTRACE2, and CytoTRACE2 scores for each cell graphed in violiln plots. B) scRNA-seq data from PrCa cells segregated by lineage variant were analyzed and graphed as in A). C) GSEA was performed on the top 300 genes most significantly correlated with high and low CytoTRACE2 scoring cells. D) SCENIC analysis of scRNA-seq data was performed to identify transcription factor regulon activity in the different PrCa lineage variants with results displayed as a hierarchically clustered heat map. E) scRNA-seq data from PrCa cells of the different genotypes was displayed as a force-directed graph projection. F) Pseudotime analysis was performed and plotted on a force-directed graph projection to explore cell state transitions during AVPC evolution. The Adeno-L2 cell state was assumed to be the initial cell state based on prior work^56^. G) The force-directed projection of scRNA-seq data color coded based on PrCa subtype assignment. H) The force-directed projection identifying the terminal lineage states based on pseudotime and palantir entropy. I) Relative expression of the indicated genes are graphed individually over pseudotime in each of the terminal PrCa lineage states.

To assess the transcription factor networks regulating the evolution of PrCa lineage variants, SCENIC (Single-Cell rEgulatory Network Inference and Clustering)^64^ was used to integrate scRNA-seq data with cis-regulatory motif analysis. NEPC-associated transcription factors like Ascl1 and Foxa2 were active across all NE differentiated lineage variants (**Figure 5D**). Notably, variants enriched in DKO_EE PrCa (NEPC_Neurod1, DNPC_Pou2f3) exhibited loss of Sox2 transcriptional activity, consistent with loss of Neuro II gene expression in patient samples that exhibit low *EZH2* expression (**Figure 4B**). This suggested Ezh2 may support a SOX2-mediated transcriptional program during AVPC development. In agreement with SCENIC and master regulator analysis, transcription factor Klf12 was highly expressed and active in *Ezh2* deficient NE variants (**Figure S5A**). DNPC_Pou2f3 PrCa cells exhibited Pou2f3 regulon activity, lacked activity for NE lineage driving transcription factors (e.g. Ascl1, Neurod1), and showed increased expression of *Klf3* and activation of the Klf regulon (**Figure 5D, S5A**). This mirrored activation of KLF transcription factors in the human DNPC variant (**Figure 4B,E**)^17,65^ and suggested a Klf family transcriptional programs were activated in *Ezh2* deficient PrCa cells thus influencing subsequent evolution of terminal AVPC lineage states. Consistent with this possibility, Klf transcription factor gene expression was detectably elevated in bulk prostate tissue from most *Ezh2* deficient mice as young as 12 weeks of age, well before development of NE lineage variants (**Figure S4C-D**). Interestingly, the Ar regulon was not enriched in NEPC_AR cells despite robust nuclear Ar expression (**Figure 1E**), but other nuclear receptors and AR co-factors including Retinoic Acid Receptor Beta (Rarb), Thyroid Hormone Receptor Alpha (Thra), Glucocorticoid Receptor (Nr3c1), and Foxa1 were active. This suggested engagement of an alternative Ar regulatory program in NEPC_AR cells (see below). The NEPC_Neurod1 population uniquely exhibited Neurod1 regulon activity as expected.

Force-directed graph projection was used to explore cell state transitions within the EGFP positive PrCa cells (**Figure 5E**). Estimating the initial lineage state as Adeno_L2^56^, cell lineage transitions were inferred using transcriptional similarities across pseudotime **(Figure 5F)**. Estimated terminal lineage states aligned with the Ezh2-dependent lineage variants detected and corresponded to our initial EGFP⁺ cell-type annotations **(Figure 5G-H, S5B-D)**. This analysis revealed that NEPC_Neurod1, NEPC_Nefl, and NEPC_AR lineage variant cells evolved from NEPC_Ascl1 cells. Notably, DNPC_Pou2f3 cells were an early branching and distinct lineage trajectory that emerged from the intermediate states exhibiting the highest terminal lineage potential (palantir entropy) (INT_EMT, INT_Inflam) (**Figure S5C**). Thus DNPC_Pou2f3 was not an intermediate during NEPC development, but arose from an evolutionary trajectory distinct from terminal NE differentiated lineage states. Lineage fate probabilities, calculated per cell and grouped by genotype, showed that SKO, DKO, and TKO cells preferentially transitioned to the NEPC_Ascl1 variant, with some TKO cells progressing to NEPC_AR (**Figure S5D**). Loss of a single *Ezh2* allele (DKO_EP) skewed DKO cells towards the NEPC_AR or NEPC_Nefl lineages. Loss of both *Ezh2* alleles favored evolution toward the NEPC_Neurod1 or DNPC_Pou2f3 lineage states. Wild type Ezh2 activity thus caused convergence towards the NEPC_Ascl1 lineage state.

Terminal lineage states exhibited distinct, temporally regulated gene expression patterns **(Figure 5H)**. An early notable change in pseudotime was reduced expression of *Ar* and Ar regulated genes like *Tmprss2* within the intermediate states that exhibit high Palantir entropy (**Figure 5C, S5C-D**). These intermediate states also showed reduced expression of luminal epithelial markers and the *Rest* transcription factor that suppresses NE gene expression. *Ascl1* expression then increased in all lineage variants, with DNPC_Pou2f3 cells exhibiting the lowest *Ascl1* expression. Ascl1 expression was maintained in terminal NEPC lineage variants, but declined in DNPC_Pou2f3. *Pou2f3* expression expression was coincident with loss of *Ascl1* expression, potentially indicating commitment to the tuft-cell like DNPC lineage state. As cells progressed through pseudotime, *Ar* expression re-emerged in some cells to generate the NEPC_AR variant. Expression of genes distinguishing the NEPC_Nefl and NEPC_Neurod1 lineage variants occurs late in pseudotime, consistent with their evolution from NEPC_Ascl1 cells. NEPC_Nefl cells exhibited high expression of Gfra4, a receptor for the neurotrophic factor persephin involved in neural crest and enteric nervous system development. GFRA4 overexpression was recently linked to activation of the Hedgehog pathway^66^ which was also activated in NEPC_Nefl cells **(Figure S4A).**

### Ezh2 deficiency causes chromatin remodeling that facilitates elaboration of alternative transcriptional programs

To examine how *Ezh2* deficiency alters AVPC evolution, the chromatin state of PrCa developing in GEMMs was analyzed by both histone mark chromatin immunoprecipitation and DNA methylation arrays. H3K27me3 marks were reduced at transcription start sites (TSS) in DKO_EE PrCa (**Figure 6A**) in agreement with decreased total H3K27me3 (**Figure 1E**). H3K4me3 and H3K27ac marks associated with active transcription were increased in DKO_EE and, to a lesser extent, DKO_EP PrCa. The H3K4me3 mark was elevated in Ezh2-deficient PrCa at the promoters of AVPC-driving genes including *Ascl1*, *Neurod1*, and *Onecut2*, as well as Klf gene family members and *Ezh1* (**Figure S6A**). DKO_EP tumors exhibited genomic regions of both increased or decreased H3K27ac (988 gain, 426 loss) whereas DKO_EE tumors displayed a very strong bias toward H3K27ac gains (7,148 gain, 17 loss) **(Figure 6B, Table S6)**. Using Binding and Expression Target Analysis (BETA)^67^, genomic regions of increased H3K27ac in DKO_EE tumors were significantly associated with transcriptional activation **(Figure 6C)**. Querying 1,1667 genes that exhibited both increased H3K27ac and increased gene expression within the same DKO_EE tumors, there was strong enrichment for Klf transcription factor family member motifs as well as motifs for other AVPC lineage specifying transcription factors like Pou2f3, Hes1, Neurod1, and Ascl1 **(Figure 6D)**. Overall, these 1,1667 genes were enriched for pathways associated with chromatin organization and GTPase signaling **(Figure 6E)**, both of which have been implicated in lineage plasticity^68,69^.

**Figure 6:**
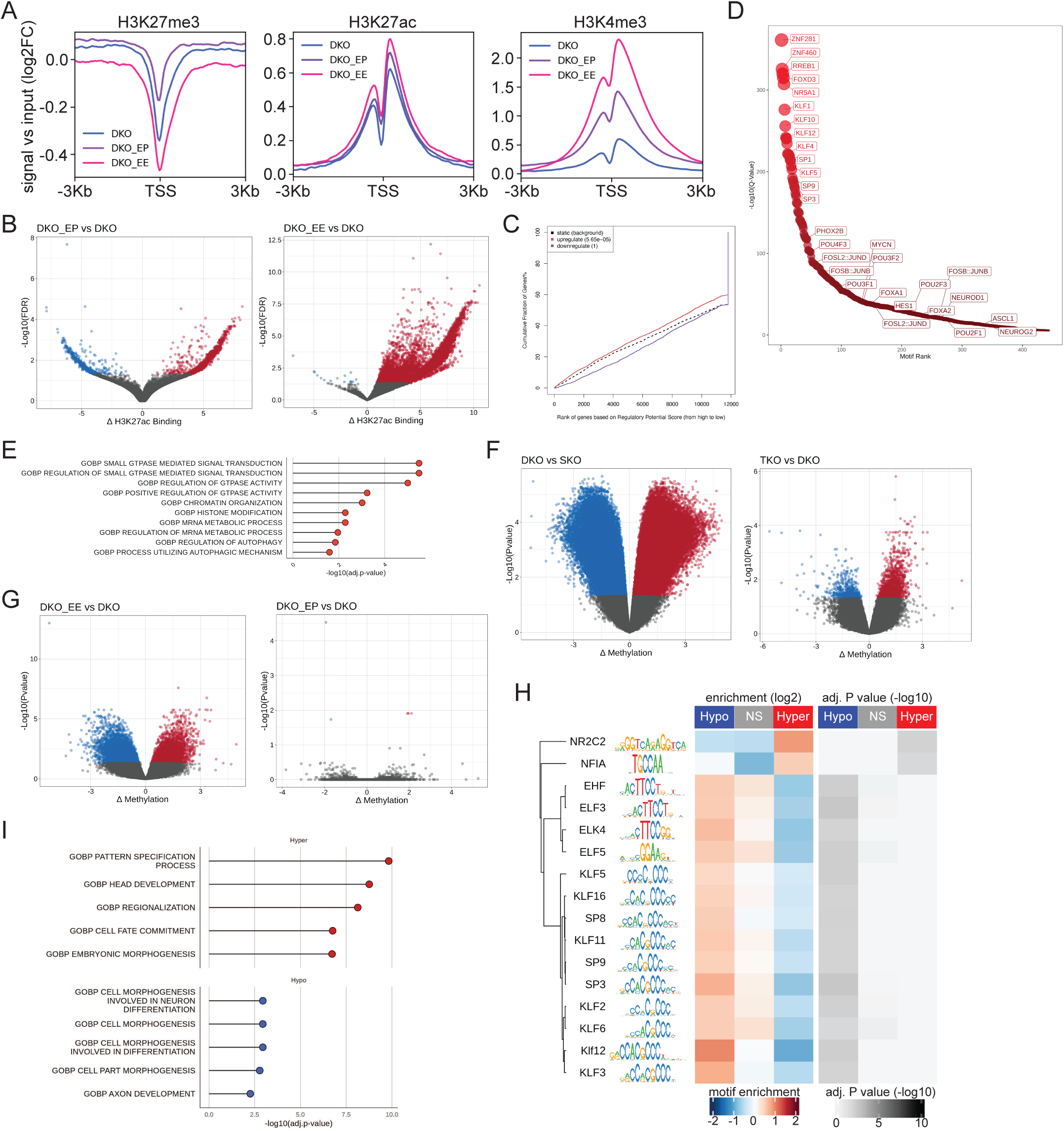
*Ezh2* deficiency alters chromatin to facilitate alternative transcriptional programs. A) Histone H3 K27me3, K27ac, or K4me3 ChIP-seq was used to analyze prostate tissue from mice of the indicated genotypes. ChIP-seq signal versus input is graphed relative to transcription start sites (TSS) genome wide. B) Genome sites of differential histone H3K27ac between DKO_EP or DKO_EE and DKO prostate tissue were graphed, demonstrating a larger number H3K27ac sites upon *Ezh2* loss. C) BETA analysis indicates genomic regions of increased H3K27ac were significantly associated with transcriptional activation. D) SCENIC analysis of genes exhibiting both increased H3K27ac and transcription in DKO_EE prostate tissue demonstrates significant enrichment of KLF transcription factor family motifs, Pou2f3, and Neurod1. E) GSEA of genes exhibiting increased H3K27ac and transcription in DKO_EE prostate tissue. F) Genomic regions exhibiting differential DNA methylation between DKO and SKO prostate tissue or TKO and DKO prostate tissue demonstrating a marked effect of Rb1 loss. G) Genomic regions exhibiting differential DNA methylation between DKO_EP or DKO_EE and DKO prostate tissue demonstrating only complete *Ezh2* loss significantly affects DNA methylation. H) Transcription factor motif analysys of sites with significant differential DNA methylation in DKO_EE prostate tissue, hypomethylated (hypo) or hypermethylated (hyper) as indicated. KLF family transcription factor DNA binding motifs are enriched in hypomethylated regions. I) Gene set enrichment analysis of genes affected by differential DNA methylation in DKO_EE prostate tissue.

*Rb1* loss had a profound effect on DNA methylation, driving both dramatic hyper- and hypomethylation changes in DKO relative to SKO PrCa (65,466 hypo; 71,692 hyper) (**Figure 6F, Table S7**). Additional loss of *Trp53* caused more modest DNA methylation changes (487 hypo; 1,728 hyper). DKO hypomethylated regions were enriched for transcription factor binding sites related to lineage plasticity including Sox2, Zeb1, and Ascl1 while hypermethylated regions enriched for Jun/Fos and ETS-family members (**Figure S6B**). Complete loss of *Ezh2* caused significant further DNA methylation changes (DKO_EE versus DKO) skewed towards hypomethylation (11,374 hypo; 7,233 hyper), but loss of a single *Ezh2* allele had minimal effects on DNA methylation (**Figure 6G, Table S7**). Motif analysis of differentially hypomethylated regions in DKO_EE PrCa revealed strong enrichment for Klf transcription factor family binding sites **(Figure 6H)**. Loss of *Ezh2* in DKO_EE PrCa cells thus reversed hypermethylation of Klf motif containing genomic sites that occurred upon *Rb1* loss, congruent with increased expression and activity of these transcription factors in mouse DNPC_Pou2f3 and NEPC_Neurod1 cells. Consistent with mouse data, KLF5 was previously identified as a regulator of the DNPC AR-/NE- and ARlow/NE- DNA methylation profile in human AVPC specimens, with its activity linked to *RB1* loss^65^. Genes affected by differential hypomethylation in DKO_EE PrCa enriched for neuronal differentiation while hypermethylated genes enriched for developmental and morphogenesis pathways (**Figure 6I**). Hypomethylated genes exhibited increased gene expression in the Ezh2 deficient AVPC lineage variants relative to the NEPC_Ascl1 variant (**Figure S6C-D)**. These findings highlight how chromatin changes driven by *Ezh2* deficiency facilitate activation additional transcription factor programs, like Klf transcription factors, to diversify AVPC lineage variant evolution.

### The AR cistrome is reprogrammed in Ezh2 deficient PrCa cells

NEPC_AR PrCa cells prevalent in DKO_EP PrCa samples expressed robust levels of nuclear localized Ar (**Figure 1E**), yet failed to express genes whose transcription is canonically stimulated by Ar signaling (**Figure 2C**). Ar chromatin immunoprecipitation analysis was thus performed to assess Ar activity. DKO_EP PrCa exhibited a marked increase in global Ar DNA binding compared to DKO or DKO_EE PrCa **(Figure 7A)**. The pattern of Ar binding across these samples was k-means clustered into four groups (**Figure 7B, Table S8**). The overall pattern of Ar binding in DKO_EP PrCa was most similar to that in SKO PrCa while DKO_EE PrCa had low Ar binding signal in all clusters. Ar binding sites in clusters 1 and 3 mapped mostly to gene promoters (**Figure 7C**). DKO_EP PrCa exhibited the highest Ar binding at cluster 1 sites with 13,490 of these sites showing significantly increased binding signal. BETA was utilized to integrate RNA-seq data and predict if Ar binding is associated with changes in transcription. Genes bound by Ar in cluster 1 were strongly associated with transcriptional up regulation in DKO_EP relative to DKO (**Figure 7D**). Ar binding at cluster 3 sites showed comparable signals in SKO and DKO_EP PrCa with lower signal in DKO and DKO_EE PrCa. Cluster 3 Ar binding was significantly associated with transcriptional repression in DKO_EP relative to DKO. Ar-binding sites in clusters 2 and 4 showed relatively low Ar-binding signal, mapped more frequently to intergenic or intronic regions, exhibited smaller differences between genotypes, and were not associated with marked changes in transcription (**Figure S7A**).

**Figure 7:**
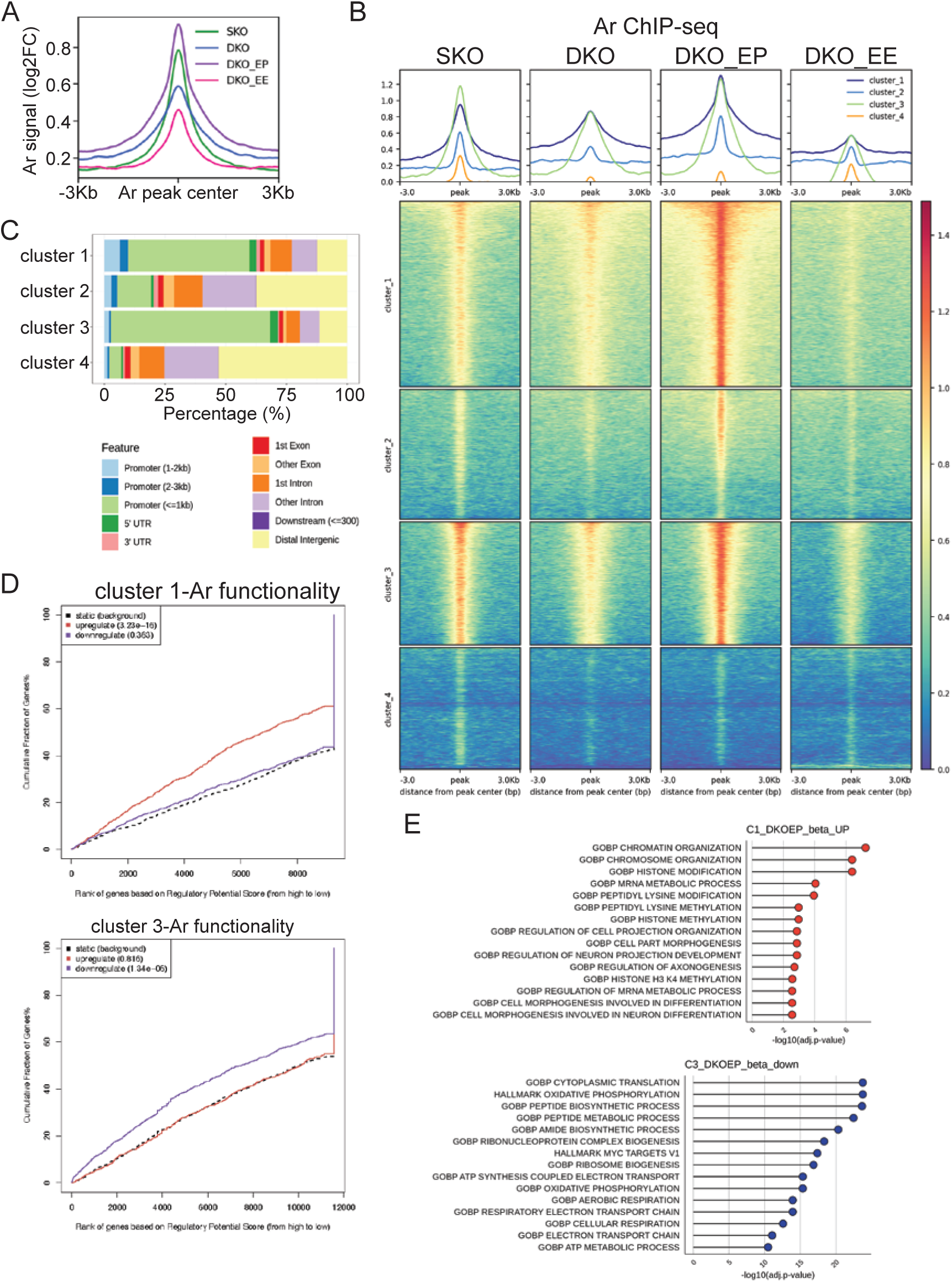
The AR cistrome is reprogrammed in *Ezh2* deficient PrCa. A) Prostate tissue with each of the indicated genotypes was analyzed by Ar ChIP-seq and Ar binding signal plotted versus distance from peak center, demonstrating highest Ar binding in the DKO_EP genotype. B) K-means clustering of Ar binding patterns in the four genotypes aligned by peak center across the genome identifies four clusters of Ar binding sites. C) The relative distribution of genomic features associated with Ar binding sites in each of the four clusters is presented in stacked bar graphs. D) BETA analysis indicates cluster 1 binding sites are significantly associated with transcriptional activation while cluster 3 Ar binding sites are associated with transcriptional repression. E) Gene set enrichment analysis of genes associated with cluster 1 and cluster 3 Ar binding sites in DKO_EP prostate tissue.

The biological programs and transcription factor DNA binding motifs associated with Ar-binding sites were examined to assess functional consequences. Cluster 1 Ar-bound gene promoters with increased transcription were significantly enriched for chromatin organization and neurogenesis-related pathways rather than canonical Ar regulated genes **(Figure 7E, Table S9)**. Interestingly, these cluster 1 Ar-binding sites were also significantly enriched for Klf transcription factor DNA binding motifs (**Figure S7B**). Ar-bound gene promoters in cluster 3 associated with reduced transcription were enriched for aerobic respiration and other energy-related processes biological programs, consistent with decreased expression of such genes in *Ezh2* deficient PrCa cells (**Figure S2B**). Cluster 3 sites were also significantly enriched for Klf transcription factor DNA binding motifs (**Figure S7B**). These observations suggested increased Klf transcription factor expression and activity may facilitate reprogramming of the Ar cistrome to facilitate transcriptional changes driving evolution of AVPC lineage states. Ar-bound sites in cluster 2 enriched for gene sets related to metastasis and cell-cell signaling with the top enriched transcription factor DNA binding motifs being nuclear hormone receptors and FOX transcription factor family members (**Figure S7B**). The Cluster 4 Ar-binding sites with strongest binding signal in SKO PrCa showed strong enrichment for the canonical Ar binding DNA sequence motif (**Figure S7B**) and enriched for genes related to Notch signaling (**Table S9**). This observation was consistent with our prior report demonstrating Notch signaling is suppressed during NEPC transdifferentiation^70^.

## Discussion

With increasing analysis of advanced PrCa patient specimens, it is becoming increasingly clear that CRPC is a heterogeneous disease state that can evolve into multiple lineage variants that diverge from the luminal epithelial-like state of primary tumor cells^7-11^. Lineage variants currently classified as AVPC are clinically aggressive and no longer rely on AR signaling for growth and survival. Given that AVPC are resistant to AR targeted therapies and don’t respond durably to chemotherapy, prognosis for such patients is poor^15,16^. The incidence of AVPC is increasing with more widespread deployment increasingly potent AR targeted therapies, thus PrCa lineage plasticity represents a growing clinical challenge^14,21^. The evolutionary relationships between AVPC variants, if any, and the molecular pathways that drive their development are poorly understood. These knowledge gaps have hampered the discovery of effective approaches for predicting, diagnosing, and treating AVPC.

Here we characterize a series of GEMMs that recapitulate the diversity of AVPC lineage variants observed in human patients. These mice develop AVPC variants analogous to well established human PrCa lineage variants including NEPC (NEPC_Ascl1), an NEPC variant expressing *NEUROD1* (NEPC_Neurod1), amphicrine (NEPC_AR), and DNPC (DNPC_Pou2f3). A novel lineage variant detected in mice distinguished by elevated expression of neurofilament genes has also been discovered in human patient data. These GEMMs thus represent clinically relevant experimental models for studying PrCa lineage plasticity and AVPC/CRPC evolution. Based on trajectory analysis of scRNA-seq data from these mice, terminal AVPC lineage variants arise from highly transcriptionally heterogeneous intermediate states characterized by inflammatory and epithelial to mesenchymal gene expression (INT_Inflam, INT_EMT), consistent with prior studies^56^. The NEPC-Ascl1 variant arising from these intermediate cell states can evolve further giving rise to the NEPC_Neurod1, NEPC_Nefl, and NEPC_AR variants, particularly in the context of *Ezh2* deficiency. In contrast, DNPC_Pou2f3 and Basal variants represent separate, distinct evolutionary trajectories rather than intermediate lineage states during evolution to NEPC lineage variants as has been proposed^18^. This interpretation is consistent with prior studies where DNPC experimental models have failed to evolve to NEPC^18^.

Inactivating mutations in the *RB1*, *PTEN*, and *TP53* tumor suppressor genes are highly recurrent in AVPC^29^ and enable PrCa lineage plasticity in experimental models like the GEMMs described here^31,32^, but these mutations are not sufficient. Epigenetic factors like EZH2 have been proposed to drive phenotypic transitions to CRPC and AVPC in the context of PrCa genetic backgrounds with sufficient transcriptional plasticity^27,44,71^. This has inspired clinical trials testing the utility of newly developed EZH2 inhibitors for the treatment of advanced PrCa. Contrary to expectation, data presented here demonstrate *Ezh2* is not required for NEPC development. Instead *Ezh2* influences AVPC evolution, favoring or stabilizing the NEPC_Ascl1 and Basal lineage states. Mouse DKO PrCa genetically deficient for *Ezh2* evolve a greater diversity of AVPC variants and show a diminished incidence of the Basal lineage state. PrCa lacking one *Ezh2* allele have increased probability of developing NEPC_AR (amphicrine) and NEPC_Nefl AVPC, while PrCa lacking both *Ezh2* alleles are more likely to develop NEPC_Neurod1 and DNPC_Pou2f3. Some of these variants can also be detected in TKO mice that retain both *Ezh2* alleles, and these variant PrCa cells also show reduced Ezh2 expression. Congruent with these findings in mice, our analysis indicates human AVPC NEPC_Ascl1 exhibits high *EZH2* expression while other AVPC variants show relatively low *EZH2* expression. Complete *Ezh2* loss thus does not prevent NEPC in mice and can actually accelerate it. This may be related to earlier development of the highly plastic intermediate INT_Inflam and INT_EMT lineage states (**Figure S4D**)^54^.

The mechanisms underlying observed effects of *Ezh2* deficiency involve chromatin changes, including DNA methylation and histone modifications, that enable the activation of alternative lineage specifying transcription factor mediated gene expression programs. In particular, Klf transcription factor expression and activity is increased early in the development of *Ezh2* deficient PrCa (12 weeks), before the appearance of NE lineage variant cells (**Figure S4D**). Changes in histone marks, DNA methylation, and the Ar cistrome observed in *Ezh2* deficient lineage variants converge to highlight the activity of Klf transcription factors. In the NEPC_AR variant, for example, binding of Ar is reprogrammed from canonical Ar target genes to genes whose expression supports the variant NE lineage states, and these genes often exhibit Klf DNA binding motifs. As Klf transcription factors both regulate *Ar* expression and bind Ar^72,73^, expression and reprogramming of the Ar cistrome may be mediated directly by Klf transcription factors. KLF5 have been implicated in the human DNPC lineage variant^18^.

Ezh2 inhibition has been reported to sensitize CRPC cells to AR targeted therapies, potentially by reversing epigenetic changes driving acquired therapeutic resistance and PrCa lineage plasticity. Based on results reported here, effects of EZH2 inhibition are likely to be complex. Ezh2 deficiency increases expression of genes associated with the epithelial adenocarcinoma lineage states, but alters rather suppresses expression of genes associated with NE differentiation. *Ezh2* deficiency also suppresses the Basal lineage variant, a variant observed in more than 40% of CRPC cases^74^. DKO_EE PrCa completely lacking functional Ezh2 retains some sensitivity to castration, but DKO_EP PrCa lacking one functional *Ezh2* allele is castration resistant. This may also be related to Klf transcription factors. Cluster 1 Ar-bound genes include Klf transcription factors (e.g. *Klf3/7/13*) and their expression is increased upon castration in both DKO_EP and DKO_EE mice. DKO_EP PrCa may be resistant to castration due to its higher baseline nuclear Ar expression, established positive feedback between AR and KLF transcription factors, and reprogramming of the AR cistrome to support Ar-independent NE lineage variants.

A limitation in this study is that all experimental data is generated in mice that may not accurately model AVPC evolution in humans. This is mitigated by retrospective analysis of human advanced PrCa RNA-seq and scRNA-seq data, demonstrating remarkable transcriptional similarities between human and mouse AVPC variants. Indeed a new NE variant discovered in mice successfully predicted its detection in human sample data. There is heterogeneity in AVPC development in mice of the same genotype. This coupled with random sampling of tissue likely introduces variability in the molecular data obtained. This potential limitation could be addressed by analysis of larger sample sizes. Nonetheless, the use of bulk tissue and single cell experimental approaches analyzing different cohorts of mice yields convergent results. Genetic *Ezh2* deficiency at the time of PrCa initiation may have different effects than pharmacological inhibition of Ezh2 in pre-existing PrCa. However, some effects like increased Neurod1 expression and activity are convergent in both contexts.

The main findings reported have implications for clinical efforts to treat PrCa by targeting EZH2. *Ezh2* loss does not prevent AVPC development but clearly affects the evolution of PrCa lineage variants. *Ezh2* deficiency suppresses the relative incidence of some variants (NEPC_Ascl1, Basal) while augmenting others (DNPC_Pou2f3, NEPC_AR, NEPC_Nefl). This indicates EZH2 inhibition may be more effective in treating some PrCa lineage variants than others and potentially predicts mechanisms by which PrCa cells can adapt to this inhibition. More nuanced approaches may thus be required to optimize treatment outcomes with EZH2 inibitors, and the data and experimental models reported here will be useful in discovering these approaches.

## Materials and Methods

### Genetically engineered mouse models (GEMMs) of prostate cancer

All procedures involving animals were approved by Roswell Park Comprehensive Cancer Center’s Institutional Animal Care and Use Committee (IACUC). The SKO, DKO, and TKO prostate cancer GEMMs and the floxed *Ezh2* allele bred into the DKO mice were described previously^32,45,52,57,75^. All experimental mice were maintained on a mixed C57BL/6:129/Sv:FVB genetic background. Animals were visually monitored daily with weekly palpation of the abdomen to assess prostate tumors. Moribund mice were euthanized and considered to have “endstage” disease. Necropsy was performed and tissues harvested to characterize prostate cancer phenotype and assess distant metastases. The Kaplan-Meier method was used to assess overall survival and was performed using GraphPad Prism software (version 10.4.1).

Mice were genotyped by analyzing tail DNA using established PCR assays specific for each allele. Mouse tails clipped at the time of weaning were extracted by boiling in 100 µL 50 mM NaOH for 30 minutes. The reaction was buffered by adding 30 µL 1 M Tris-HCl, pH 7.4 and crude DNA collected by centrifuging. DNA extract was diluted 1:20 for use as template in PCR reactions using primers at 0.5 µM final concentration (**Table S1**), and the KAPA Taq PCR Kit (Roche BK1000) used to perform reactions according to the manufacturer’s instructions. Annealing temperature for all PCR reactions was 61.2°C. PCR reactions were resolved on 1.5 % agarose gels along with molecular weight standards to determine PCR fragment sizes.

To genotype prostate tissue specimens, DNA was extracted from flash frozen tissue samples using 50 mM Tris, pH 7, 5 mM EDTA, 1% SDS, 0.2 M NaCl, and 0.2 mg/mL proteinase K) incubated overnight at 55°C with shaking. Extracts were centrifuged at 16,100g for 5 minutes and the supernatant transferred to a new tube. DNA was precipitated using isopropanol, centrifuged at 12,000g for 10 minutes, and the DNA pellet dissolved in DNase/RNase-free H_2_O (between 40 to 60 µL) by shaking at 55°C for 1-2 hours. PCR reactions with prostate tissue DNA was carried out as above.

### Histological and immunostaining analysis

Soft tissues were fixed in phosphate-buffered 4% paraformaldehyde overnight at 4°C, washed with phospho-buffered saline (PBS) three times for 20 minutes with shaking per wash, then washed with 65% ethanol for 30 minutes with shaking, and stored in 70% ethanol. Harvested tibias were fixed in 10% neutral buffered formalin for one week. Following fixation, tissue was washed with PBS, as described above, and decalcified in neutral EDTA (143 mg/mL) for three weeks. All processed tissue was paraffin-embedded and serially sectioned at 5 µm thickness and mounted on slides. Slides were stained with H&E for histological analysis.

For immunostaining, tissues sections were deparaffinized by incubating at 55°C for 30-60 minutes, washed three times in xylene for 5 minutes per wash, incubated in 100% ethanol for 10 minutes, washed two times in 95% ethanol for 10 minutes per wash, and finally incubated in 70% ethanol for 10 minutes. Tissue sections were rehydrated by washing two times in distilled H_2_O for 5 minutes per wash. Antigen retrieval was conducted using either citrate buffer containing 1.8 mM citric acid, 8.2 mM sodium citrate at pH 6 or Tris-EDTA Buffer, pH 9.0 (Antigen Retrieval Buffer Abcam ab93684)(Supplementary Table 2), in conjunction with heat induced epitope retrieval. Tissue slides were immersed in antigen retrieval buffer and microwaved on high until buffer was approximately 95°C. When the buffer reached 95°C, the microwave power was reduced to the lowest setting and tissue slides were heated 10-20 minutes. Tissue slides cooled to room temperature were washed in distilled H_2_O three times for 5 minutes per wash, incubated in 3% hydrogen peroxide (v/v in H_2_O) for 10 minutes, washed again in distilled H_2_O three times for 5 minutes per wash, and finally washed one time in 1x PBS, plus Tween-20 at 1% (v/v) (PBS-T).

Blocking buffer (ImmPRESS^®^ HRP Goat Anti-Mouse IgG Polymer Detection Kit, Peroxidase-Vector Laboratories MP-7452 or Goat Anti-Rabbit IgG Polymer Detection Kit, Peroxidase-Vector Laboratories-MP-7451) was added to each section and incubated for 1 hour. Primary antibodies were prepared at the indicated dilutions (**Table S2**) in PBS-T and antibody incubated with tissue sections overnight at 4°C. Slides were washed three times in PBS-T for 5 minutes per wash and incubated with Goat Anti-Mouse or Goat Anti-Rabbit secondary antibodies from the kits described above for 1 hour at room temperature. After secondary antibody incubation, sections were washed three times with PBS-T for 5 minutes per wash. Peroxidase detection was carried out using 3,3’-diaminobenzidine (DAB) stain via the DAB Substrate Kit, Brown (Vector Laboratories, SK-4100) according to the manufacturer’s instructions. Positive control slides for each antibody were used to determine optimal exposure time to DAB – approximately 30 seconds to 2 minutes, based on the primary antibody used. After DAB exposure, the reaction was terminated by immersing slides in H_2_O. Tissues sections were counterstained by incubating with hematoxylin nuclear staining solution (1.044 g/mL, Sigma 51275) for 2 minutes. Stained sections were washed under continuously running water for 3 minutes, washed twice with 1% acid alcohol (1% [v/v] hydrochloric acid in 70% ethanol), rinsed under running water for 4 minutes, washed in 95% ethanol for 3 minutes, 100% ethanol for 3 minutes, and then in xylene three times for 3 minutes per wash. Slides were mounted using Cytoseal™ mounting solution (Fisher 22-050-262).

### Western blotting analysis

Protein lysis buffer containing 20 mM PIPES, 150 mM NaCl, 1 mM EGTA, 1.5 mM MgCl_2_, 1% Triton (v/v) supplemented with protease (Pierce^TM^ Protease Inhibitor Tablets, ThermoFisher 88265) and phosphatase inhibitors (Pierce^TM^ Phosphatase Inhibitor Mini Tablets, ThermoFisher 88667) was used to prepare primary prostate tissue lysates using a dounce homogenizer. Protein concentration was measured using the DC protein assay (BioRad 5000111) and the indicated protein concentrations of tumor lysates (Supplementary Table 2) prepared in loading buffer containing 0.4 M Tris, pH 6.8, 8% SDS (m/v), 39% glycerol (v/v), and 0.04% bromophenol blue (m/v) with reducing agent beta-mercaptoethanol at 5% (v/v). Proteins were resolved by SDS-PAGE using Tris-Glycine gels at indicated percentages (Supplementary Table 2) in Tris-Glycine buffer at 100V for 2 hours. All SDS-PAGE was performed with stacking gels at 5% Tris-Glycine. Protein was transferred to nitrocellulose membranes in Tris-glycine buffer with 20% methanol overnight at 30V at 4°C. After overnight transfer, membranes were blocked with 5% Bovine Serum Albumin (BSA) in Tris-Buffered Saline (TBS) with 1 % (v/v) of Tween-20 (TBS-T). Indicated dilutions of primary antibodies (Supplementary Table 2) were made in TBS-T plus 5% BSA and membranes incubated for 1 hour at RT. After washing three times for 10 minutes each with TBS-T, membranes were incubated with rabbit-horseradish peroxidase (HRP) secondary antibody at 1:5000 dilution in TBS-T plus 5% BSA for an additional hour. Membranes were washed three times for 10 minutes each with TBS-T and developed using either SuperSignal™ West Pico PLUS Chemiluminescent Substrate (Thermo 34580) or Pierce™ ECL Western Blotting Substrate (Thermo 32106) based on protein abundance (Supplementary Table 2). Protein bands were detected using X-ray film.

### Quantitative real-time PCR analysis

Total RNA was extracted from tissue using the TRIzol RNA isolation reagent and a dounce homogenizer. RNA concentration was analyzed using a Nanodrop spectrophotometer and cDNA was generated using the BioRad iScript cDNA synthesis kit (BioRad 1708890) from 2 µg RNA. Gene expression of *Ezh2* was measured with *L32* as a reference gene using iTaq™ Universal SYBR® Green Supermix (BioRad 1725120) and a BioRad CFX Connect Real-Time PCR Detection System. Quantitative RT-PCR primers used are listed in Supplementary Table 2.

### Bulk RNA sequencing analysis

Total RNA was extracted using the miRNeasy Mini Kit (Qiagen). Lysis was performed with 700 µL QIAzol Lysis Reagent followed by vortexing. After chloroform addition, phase separation was achieved by centrifugation. Ethanol was added to optimize RNA binding and the sample purified an RNeasy Mini spin column. Membrane-bound RNA was washed, on-column DNase digestion performed, and RNA eluted in RNase-free water. RNA yield and purity were assessed using a Qubit Broad Range RNA Kit (Thermo Fisher), and integrity was evaluated via the TapeStation 4200 (Agilent Technologies).

Total RNA (500 ng) was used to prepare sequencing libraries with the RNA HyperPrep Kit with RiboErase (HMR)(Roche Sequencing Solutions), following the manufacturer’s protocol. Ribosomal RNA (rRNA) was first depleted, and the remaining RNA was treated with DNase to remove genomic DNA contamination. First-strand cDNA synthesis was performed using random primers, followed by second-strand synthesis, during which dUTP was incorporated in place of dTTP to enable strand-specificity. The resulting double-stranded (ds) cDNA was purified using Pure Beads (KAPA Biosystems). To facilitate adapter ligation, an adenine (A) overhang was added to the 3’ ends of the blunt fragments and indexing adapters with a complementary thymine (T) overhang were ligated. Adapter-ligated libraries were amplified by PCR, purified using Pure Beads, and assessed for fragment size distribution using the Agilent 4200 TapeStation D1000 ScreenTape (Agilent Technologies). Library quantification was performed using the KAPA qPCR kit (KAPA Biosystems), and libraries were pooled equimolarly based on the experimental design. Pooled libraries were denatured, diluted to 350 pM, and spiked with 1% PhiX control. Sequencing was conducted on an Illumina NovaSeq 6000 system using 100 bp paired-end reads, following the manufacturer’s protocol. On average, 50 million paired-end reads were generated per sample.

Bulk RNA-seq reads were first assessed for quality using fastqc and then mapped to the mouse GRCm38-mm10 genome using STAR and quantified with RSEM. Batch effect removal was performed using limma. Features not expressed in at least three samples were filtered out. Normalized counts were calculated within DESeq2 with variance stabilization. Differential gene expression was performed using DESeq2 using Wald statistic and alpha = 0.05. Genes were deemed significantly differentially expressed if log2FC was greater or less than 1.2 and adjusted p.value less than 0.05. Gene set variation analysis using published gene signatures was performed on normalized counts using the GSVA package. Gene network inference and master regulator analysis was performed on normalized counts using corto. Gene set enrichment analysis of differentially expressed genes ranked by Wald statistic was performed using clusterProfiler (fgsea) against the mouse collection MSigDB gene sets (Hallmark and Gene Ontology: Biological Processes) with minGSSize = 15 and maxGSSize = 1000, and gene sets were deemed significantly enriched if the adjusted p.value < 0.05.

### Single cell RNA sequencing analysis

Mouse prostates were dissected, and half the prostate was put on ice in Advanced Dulbecco’s Modified Eagle Medium/Ham’s F-12 (Thermo 12634010) in a 6-well dish and the other half of the prostate was fixed as described above for histological analysis. Prostate tissue was minced into 1 mm^3^ pieces and dissociated for 1-1.5 hours at 37° C in AdDMEM/F12 containing 5 mg/mL collagenase II, 10 µM Rock inhibitor (Y-27632 dihydrochloride), and 1 nM synthetic androgen R1881. Tissue dissociation was terminated by addition of excess cold AdDMEM/F12, and cells collected by centrifugation at 1000 rpm at 4°C. Cells were resuspended in warm 0.25% Trypsin, 2.21 mM EDTA containing 10 µM Rock inhibitor and 1 nM R1881 and incubated at 37°C for 15 minutes with occasional agitation by pipetting. Trypsin digestion reaction was terminated by excess cold AdDMEM/F12 and cells were pelleted by centrifugation at 1000 rpm at 4°C. The cell pellet was resuspended in 3 mL AdDMEM/F12 and the cell suspension strained through a sterile 40 µm filter. The strained cells suspension was washed once with 1 mL AdDMEM/F12 centrifuged at 1000 rpm at 4°C to collect cells and assess red blood cells within the sample. If contaminated with red blood cells, the cell pellet was resuspended in 1x Ammonium-Chloride-Potassium (ACK) buffer (Thermo A1049201) for 10 minutes at room temperature to lyse red blood. The remaining cells were collected by centrifugation at 1000 rpm at 4°C. Cell suspensions were then used to create single cell libraries. Cells were either used for scRNA-seq directly or after flow sorting for viable cells; flow sorting had no detectable impact on the data based on PCA analysis.

Single-cell transcriptomic libraries were generated using the 10x Genomics Chromium Next GEM Single Cell 3’ v3.1 chemistry. Prior to library construction, single-cell suspensions were assessed for quality using the ViaStain™ AOPI Staining Solution in combination with a Cellometer K2 automated cell counter (Nexcelom Bioscience) to determine cell concentration, viability, and to confirm the absence of aggregates or debris that might interfere with droplet encapsulation. Cells were loaded onto the Chromium X Controller (10x Genomics), where they were partitioned into nanoliter-scale Gel Beads-in-Emulsion (GEMs), each containing a single cell and a barcoded bead. Within each GEM, reverse transcription was performed to produce uniquely barcoded, full-length cDNA. Following emulsion breakage, the barcoded cDNA was purified and amplified via PCR. Amplified cDNA was enzymatically fragmented, followed by end-repair, A-tailing, adaptor ligation, and sample indexing via PCR to generate Illumina-compatible libraries. The final libraries were assessed for size distribution and integrity using the D1000 ScreenTape assay on a TapeStation 4200 (Agilent Technologies) and quantified using the KAPA Library Quantification Kit for Illumina platforms (Roche). Sequencing libraries were pooled at equimolar concentrations, denatured, and diluted to a final loading concentration of 300 pM with 1% PhiX control library (Illumina). Pooled libraries were sequenced on an Illumina NovaSeq 6000 system, following the manufacturer’s protocol. An average sequencing depth of ∼20,000 paired-end reads per cell was achieved for downstream analysis.

Raw sequence data was demultiplexed, aligned, and processed for feature calling using the 10x Genomics Cell Ranger pipeline (v8.0.1) using a modified mm10 genome to include enhanced GFP (“EGFP”) and tdTomato reporter gene sequences. High-quality cells that passed quality thresholds (percent mt-reads < 10%; nFeature_RNA >300 or <3,000; nCount_RNA <80,000; feature detected in min 3 cells) were further analyzed with Seurat (v4). Additional doublet detection and filtering was performed with Scrublet. A final count of 210,106 high quality cells were used for downstream analysis. Cell cycle phase prediction was performed using CellCycleScoring within Seurat. Gene expression counts were normalized using SCTransform with differences in percent.mt regressed out. Following normalization, the top 2,000 highly variable features were identified using VariableFeatures. Principal component analysis (PCA) was performed to reduce dimensionality, and the first 30 principal components (PCs) were selected for downstream clustering and visualization based on elbow plot and variance explained. All cells were initially clustered using the Louvain algorithm with a shared nearest neighbor (SNN) graph (resolution=0.3). Uniform Manifold Approximation and Projection (UMAP) was used for visualization of the merged datasets. Differential expression analysis was conducted using FindMarkers from Seurat using MAST (min.pct = 0.25, logfc.threshold = 0.25). Cell annotation mapping on all cells was performed using Seurat’s MapQuery function using ‘coarse’ and ‘fine’ mouse prostate cell type labels from Chan et al., 2022 (GSE210358) as a reference. Broad cell type annotations were then implemented based on established gene markers from the literature and reference mapping predictions. MAGIC (magic-impute) was used to impute marker gene counts for UMAP visualizations.

Cells which underwent Cre-mediated recombination of the reporter gene were bioinformatically identified by positive EGFP expression. Initially, cells with RNA counts of >1 for the epithelial cell marker Epcam were identified yielding 79, 420 cells. Further subsetting of these cells based on EGFP counts > 1 yielded 60,836 cells. These cells were re-analyzed and re-projected in UMAP space and clustered (resolution = 0.2) using PCA with the first 30 principal components. Again, MAGIC (magic-impute) was used to impute marker gene counts for low-dimensional visualizations. Custom module scoring was performed with AUCell using curated gene sets from the literature. EGFP+ cell type labels were then implemented based on established gene markers from the literature (gene set scoring) and top differential markers (cluster vs rest). Pathway analysis of Hallmark gene sets (mouse) was performed using SeuratExtend via AUCell. To quantify cellular plasticity at single-cell resolution, we applied CytoTRACE2 and visualized these scores per cell, grouped by either cell type or genotype. Identifying genes most significantly correlated (Pearson) with high or low plasticity scoring cells was achieved using ccaPP. The top 300 most positive and negatively correlated genes with plasticity were used for pathway enrichment analysis using EnrichR. Transcription factor regulatory networks were determined at single-cell resolution with SCENIC and grouped by EGFP+ cell type.

EGFP+ cells were used for trajectory analysis using CellRank2. First, ForceAtlas2 was used to visualize the continuous state of lineage transitions and bifurcations in a force-directed graphical layout. The cell-cell transition matrix was computed on imputed expression within the CellRank.PseudotimeKernel using pseudotime calculated by Palantir as the time key, with the root being set within the “Adeno_L2” cell population. To predict the initial, intermediate, and terminal cellular macrostates, the Generalized Perron Cluster Cluster Analysis (GPCCA) estimator was fitted and converged on eight optimal macrostates which agreed with our annotated EGFP+ cell types, so were labeled in that regard. Within those eight macrostates, the GPCCA estimator further predicted the single top initial state (“Adeno_L2”). Fate probabilities were computed by absorption probabilities on the Markov chain using random walks from each query cell towards one of the terminal states. Entropy was calculated within CellRank2 as a differentiation potential score based on the fate (branching) probabilities. To determine putative driver genes for each lineage, we correlated the fate probabilities with gene expression from cells in the Adeno_L2 population to each of the terminal cell populations and significance of the correlation was determined by multiple tested-corrected q-values. To compute and visualize gene expression trends along the different lineage trajectories, cells were first ordered in pseudotime, and gene expression data was used to fit the Generalized Additive Model (GAM) within CellRank2. Gene expression trends of selected top correlated drivers were visualized by stream plots across Palantir pseudotime.

### DNA methylation analysis

Genomic DNA was extracted from mouse tissue using the QIAamp DNA Mini kit (Qiagen) following the manufacturer’s protocol. High-quality DNA was eluted in nuclease-free water (Corning, Cat# 46000CI). Quantitative assessment of the purified DNA was performed by using a Qubit Broad Range DNA kit (Thermofisher). DNA methylation analysis was performed using Illumina Mouse Methylation BeadChips following the manufacturer’s protocol. 500 ng of high-quality genomic DNA was bisulfite-converted using the EZ DNA Methylation-Gold Kit (Zymo Research) per sample. Bisulfite-converted DNA was hybridized to the Infinium Mouse Methylation BeadChip (Illumina). Array processing—including amplification, fragmentation, hybridization, single-base extension, and staining—was carried out according to the manufacturer’s protocol and arrays were scanned using the Illumina iScan system.

Data processing and quantification was accomplished using the ChAMP package revised for 285K Mouse array. Detectible beta values for all probed CpG sites were initially compiled and filtered to remove those associated with multiple alignments, detected in <5% of samples, and known SNPs. To adjust for probe design bias (Infinium Type-I, Type-II), the beta-mixture quantile normalization method (BMIQ) was employed. Differentially methylated positions (DMP) were determined using normalized beta values and deemed significant if FDR < 0.05 and 10% change in methylation for each comparison. Differentially methylated regions (DMR) were similarly calculated using FDR < 0.05 and change in regional methylation > 5%. DMRs were determined using DMRcate within ChAMP. CpG probes were annotated to genes based on the Infinium Mouse Methylation BeadChip Manifest probe id annotations for mm10 and queried against GO: Biological Processes gene sets using clusterProfiler with minGSSize = 15 and maxGSSize = 1000. Motif enrichment within DMR’s was performed using monaLisa on binned genomic regions and querying JASPAR 2020 vertebrates’ collection. Significantly enriched motifs were visualized using hierarchical clustering of the pearson residuals and motif similarity. Significantly hypomethylated DMRs in DKO_EE relative to DKO were annotated and compiled into a signature which was scored for activity within our scRNA-seq dataset using AUCell.

### ChIP-seq analysis

Chromain immunoprecipitation library preparation and sequencing was as described previously^32^. Briefly, **c**hopped tissue (∼50mg) was pulverized with plastic pestles in 1.5 ml tubes, pulverized tissue was directly resuspended in 2 mM of disuccinimidyl glutarate (DSG, Pierce), incubated for 45 minutes at room temperature, pelleted and the pellet resuspended in 1 ml of 1% formaldehyde. After incubation for 10 minutes at room temperature, crosslinked tissue was quenched with 0.125 M glycine for 5 minutes, washed with PBS, and pellets resuspended in 500 μl of 1% SDS (50 mM Tris-HCl pH 8, 10 mM EDTA). Fixed samples were sonicated for 10 minutes using Covaris E220 sonicator at 140 peak incident power, 5% duty factor, and 200 cycles per burst in 1 ml AFA fiber millitubes. Chromatin (40 ug) was immunoprecipitated with 5 μg of antibody, immunoprecipitated using Protein A/G beads (Invitrogen Dynabeads 1000-2D and 1000-4D), washed 6X with RIPA buffer, reverse crosslinked with Proteinase K, and column purified. ChIP-seq libraries were made using Swift DNA Library Prep Kit (Swift Biosciences 10024). 75bp paired end reads were sequenced on a NextSeq instrument (Illumina).

Raw fastq files were assessed for quality following adapter trimming using fastqc and MultiQC and then aligned to the mouse genome (mm10) from UCSC using bwa. Outputs were sorted with samtools and assessed for mapping alignment quality. Duplicated reads that occur due to PCR artifacts were marked and removed using Picard (Broad Institute). Peaks were called using macs2 using the ‘narrow peak’ parameter for H3K27ac and H3K4me3 or ‘broad peak’ for H3K27me3 and filtered for standard chromosomes and non-blacklist regions. Peaks were deemed significant relative to input DNA if FDR < 0.05. Peaks were annotated to UCSC-mm10 genomic regions and gene transcripts (TxDb.Mmusculus.UCSC.mm10.knownGene) using ChIPseeker. Differential peak calling was performed with DiffBind. Briefly, peaks between groups were merged, normalized based on library size, and the significantly differentially bound regions (FDR <0.05) were determined between sample groups using the DESeq2 algorithm. deepTools was used to normalize reads (RPKM) and plot signal across candidate genomic loci using karyoploteR with the ENCODE candidate cis-Regulatory Elements track from UCSC for visualizing functional genomic regions. Normalized signal at peak centers of TSS (UCSC-mm10) was plotted using deepTools bigwigCompare (log2 ratio of signal vs input), computeMatrix (+/- 3kb), plotProfile, and plotHeatmap (kmeans=4) functions. Functional enrichment of peak-genes was performed using clusterProfiler against the mouse collection MSigDB gene sets (Hallmark and Gene Ontology: Biological Processes) with minGSSize = 15 and maxGSSize = 1000, and gene sets were deemed significantly enriched if the adjusted p.value < 0.05. Binding and Expression Target Analysis (BETA) predictions were performed on regions of significantly gained H3K27ac or AR-bound clusters and integrating our bulk RNA-seq data from the noted conditions. The HOmo sapiens COmprehensive MOdel COllection (HOCOMOCO) v12 database (mouse orthologs) and JASPAR 2020 vertebrates’ collection were queried to identify transcription factor binding motifs in peaks, and enrichment was calculated relative to shuffled background DNA sequences performed with MEME Suite using a Fisher’s exact test. Enrichment is based on the best match to the motif in each sequence and ranked by ‘E-value’ which is calculated by multiplying by each motif hit p-value by the total motifs queried.

## Acknowledgements

This work was supported by grants from the NCI (R01CA234162 and U24CA274159 to D.W.G.; R50CA283805 to P.S.) and the Prostate Cancer Foundation (22CHAL13 to D.W.G.). This work was supported by the Roswell Park Comprehensive Cancer Center and NCI grant P30CA016056 that supports the Genomics and Comparative Oncology Shared Resources.

