## Supplementary Figures and Tables for "EZH2 alters aggressive variant prostate cancer subtype evolution, but is not required for neuroendocrine prostate cancer"

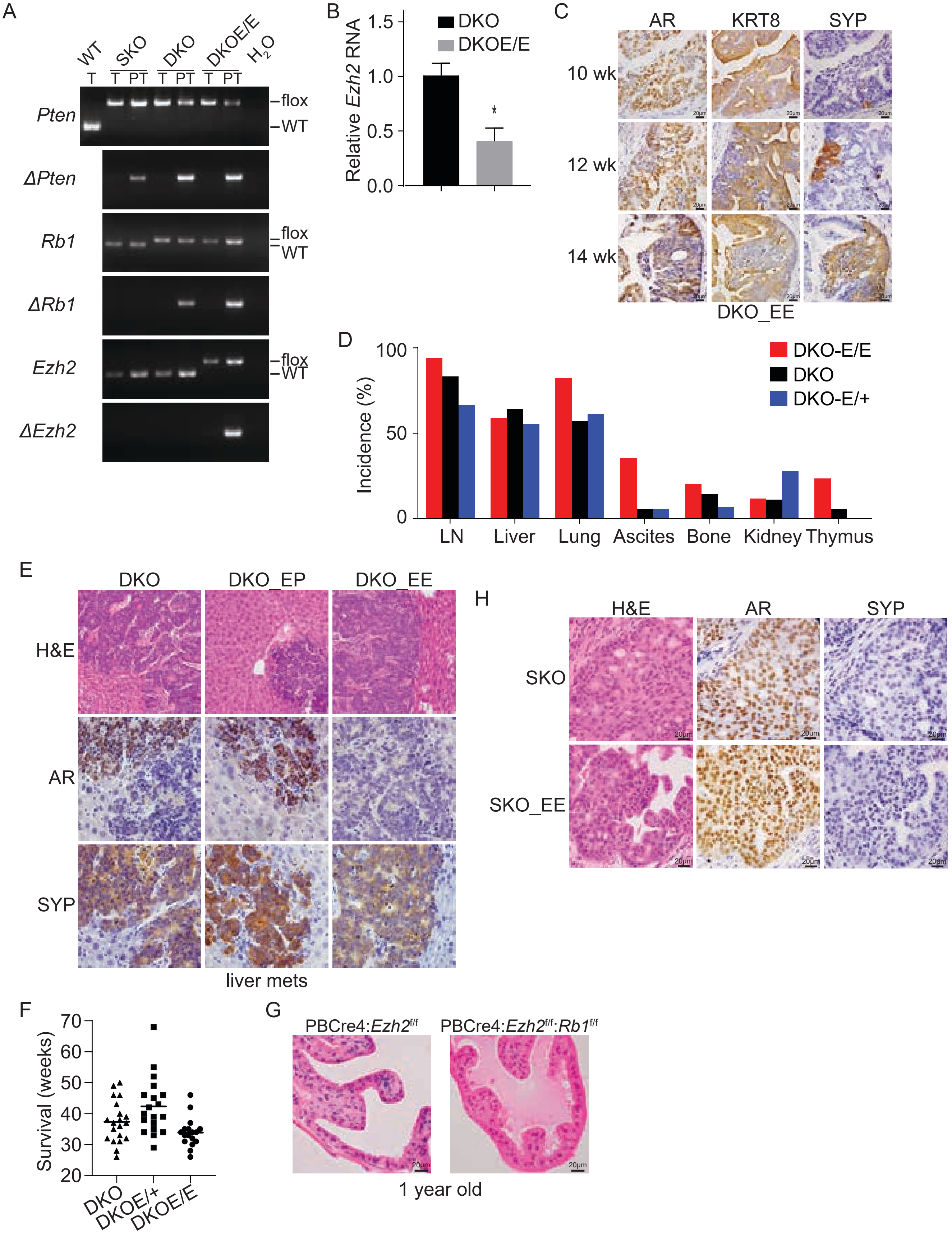

Supplementary Figure S1: Charadterization of *Ezh2* deficient GEMMs.

*Supplementary Figure 1: Characterization of Ezh2 deficient GEMMs.* A) PCR assays designed to detect floxed, wild type, or Cre-deleted gene alleles were used to analyze tail or prostate tissue DNA from mice of the indicated genotypes. Representative images of EtBr-stained agarose gels resolving PCR DNA fragments are shown, indicating the size of expected PCR fragments. B) *Ezh2* RNA in prostate tissue of the indicated genotypes was quantitated by real time reverse transcription PCR with the mean and standard deviation shown (n=3). C) Prostate tissue from DKO_EE mice of the noted ages was sectioned and immunostained for the proteins listed. Representative images of stained sections are shown at the indicated scale. D) Mice of the indicated genotype were necropsied at endstage and the listed tissue sites examined for PrCa metastases. The incidence of metastasis at each tissue site is shown for each mouse genotype. E) Liver tissue from mice of the indicated genotypes was sectioned and stained for Hematoxylin & Eosin (H&E) or immunostained for the proteins listed. Representative images of liver metastases are shown. F) Survival time of mice with the listed genotypes is plotted with each point representing an individual mouse. The bar indicates median survival time. G) Prostate tissue from mice of the indicated genotypes was sectioned and stained for H&E. Representative images are shown at the indicated scale. H) Prostate tissue from mice of the indicated genotypes was sectioned and stained and is E).

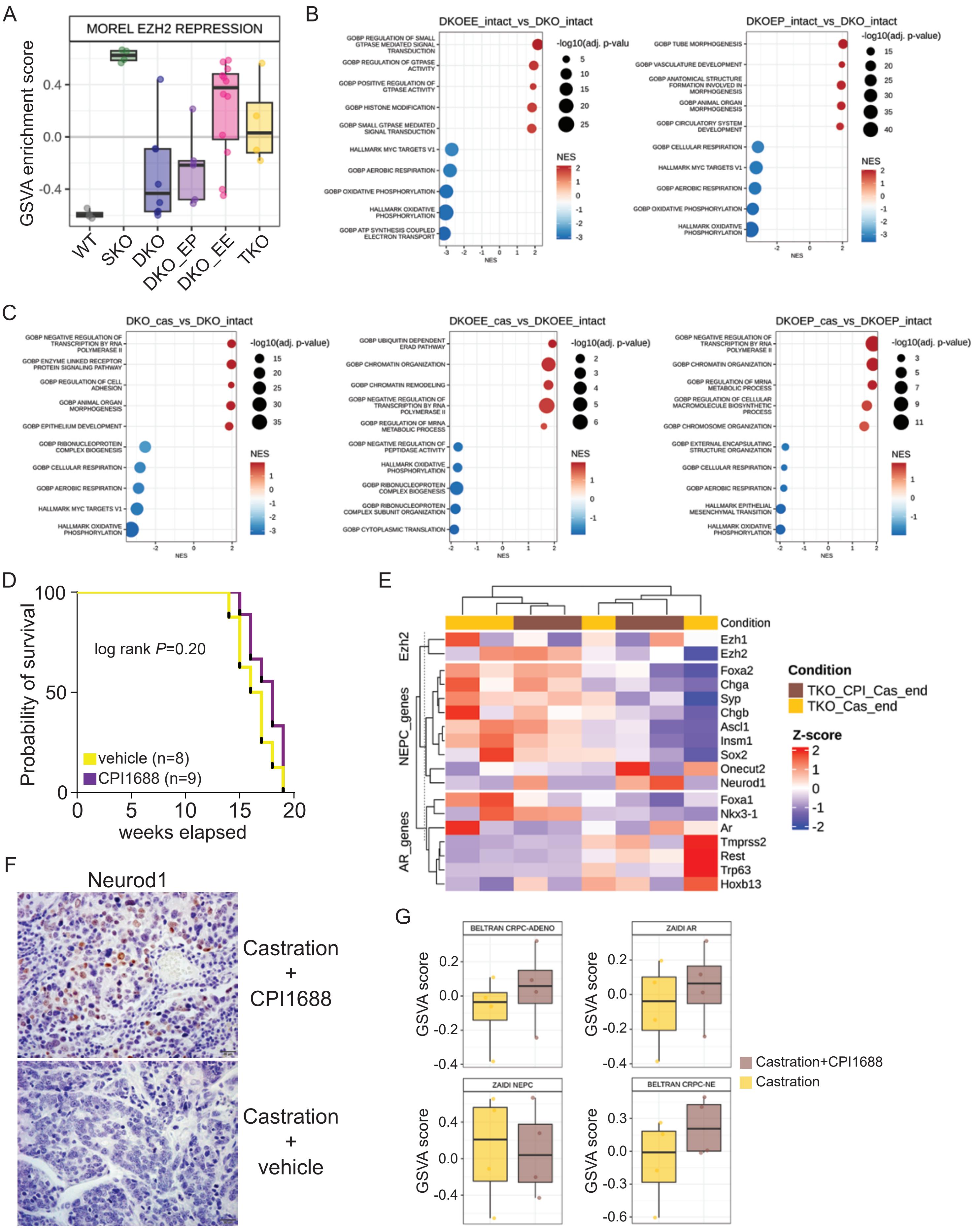

Figure S2: *Ezh2* deficiency alters transcriptional reprogramming during AVPC development

*Supplementary Figure 2: Ezh2 deficiency alters transcriptional reprogramming during AVPC development.* A) GSVA was used to analyzed PrCa RNA-seq data from mice of the indicated genotypes for gene expression induced by pharmacological EZH2 inhibition as determined by Morel et al.[54] Each point is the GSVA score for an individual sample with cumulative data displayed as a box and whisker plot. B) Differentially expressed genes in RNA-seq data from the indicated genotypes was analyzed by GSEA. The five most up- and down-regulated gene sets are shown. C) Prostate tissue from intact or castrated mice of the listed genotypes was analyzed by RNA-seq, and GSEA performed as in B). D) TKO mice castrated at 12 weeks of age were treated with vehicle or the EZH2 inhibitor CPI1688 daily (75 mg/kg), and survival of mice assessed by Kaplan-Meier plot. Survival was not significantly different between groups. E) Prostate tissue from end stage mice treated in D) was analyzed by RNA-seq analysis. The heat map shows differentially expressed genes relevant to NEPC, including *Neurod1*, and AR signaling. F) Prostate tissue from end stage mice treated in D) was sectioned and immunostained for Neurod1. Images at the indicated scale show detectable nuclear Neurod1 immunostaining in CPI1688 treated samples. G) RNA-seq from TKO mice treated in D) was analyzed by GSVA for the indicated gene signatures[13,19] showing evidence of increased AR activity in CPI1688 treated samples.

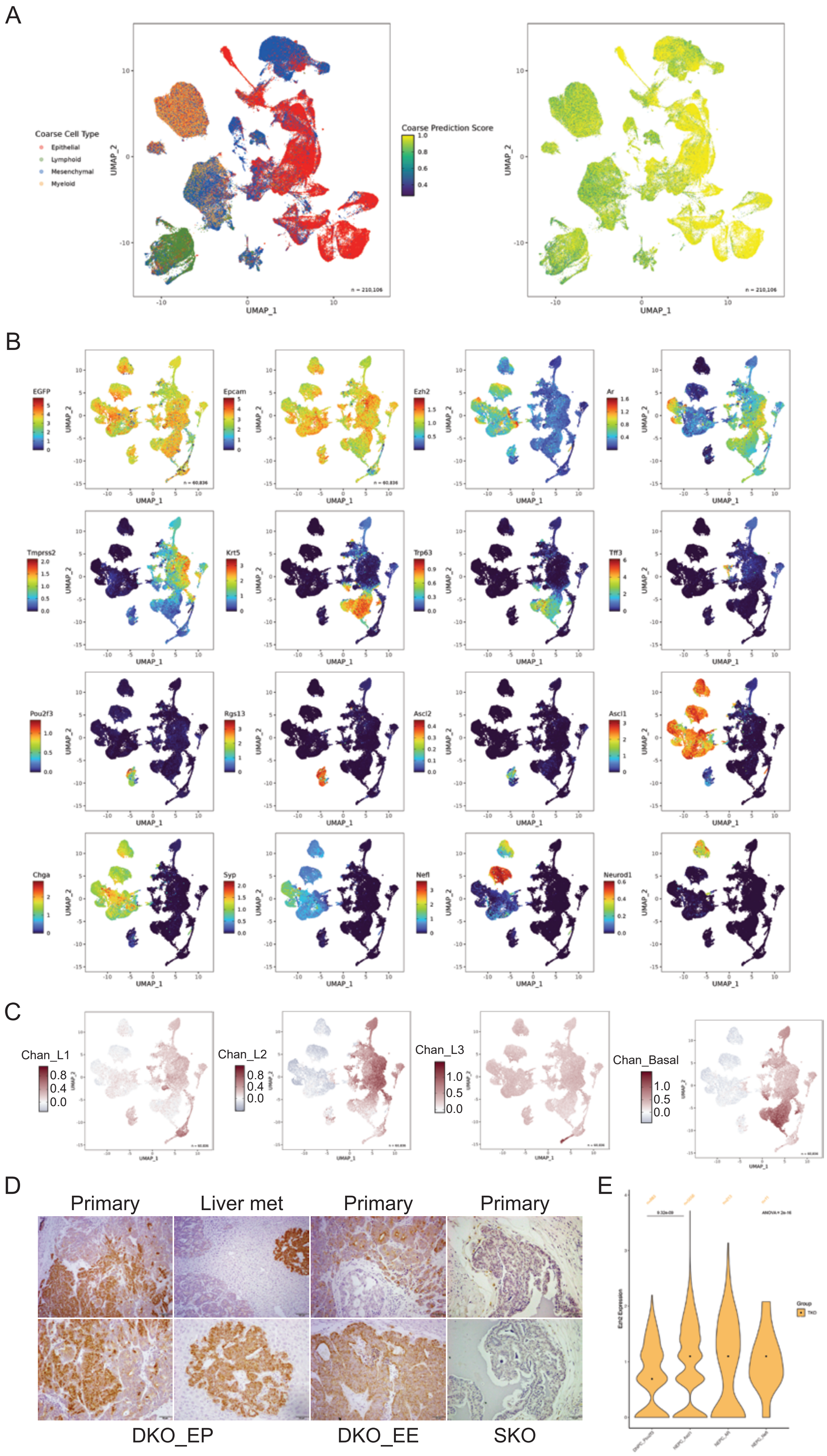

Supplementary Figure 3: Detection of PrCa lineage variants in vivo

*Supplementary Figure 3: Detection of PrCa lineage variants in vivo.* A) PrCa tissue cells collected from mice of all genotypes examined were analyzed by scRNA-seq and plotted as a UMAP projection of cells colored by cell type (left) or prediction score (right) from anchor-based reference mapping of mouse scRNA-seq data from Chan et al[56]. B) EGFP+/Epcam+ PrCa cells were analyzed separately and plotted as a UMAP color coded by the relative expression of the indicated genes. C) The PrCa UMAP is color coded based on gene expression similarity to the indicated and previously established signatures for normal prostate epithelial cells in the mouse[56]. D) Prostate or liver tissue sections from mice of the indicated genotypes was immunostained for Nefl. Representative images are shown at the indicated scale. E) PrCa lineage variants developing in TKO mice were identified by scRNA-seq and *Ezh2* expression plotted for each cell with the dot representing the median relative expression level (n=#cells in each subtype analyzed). *Ezh2* expression in DNPC_Pou2f3 cells was signficantly lower than NEPC_Ascl1 cells with the p values listed.

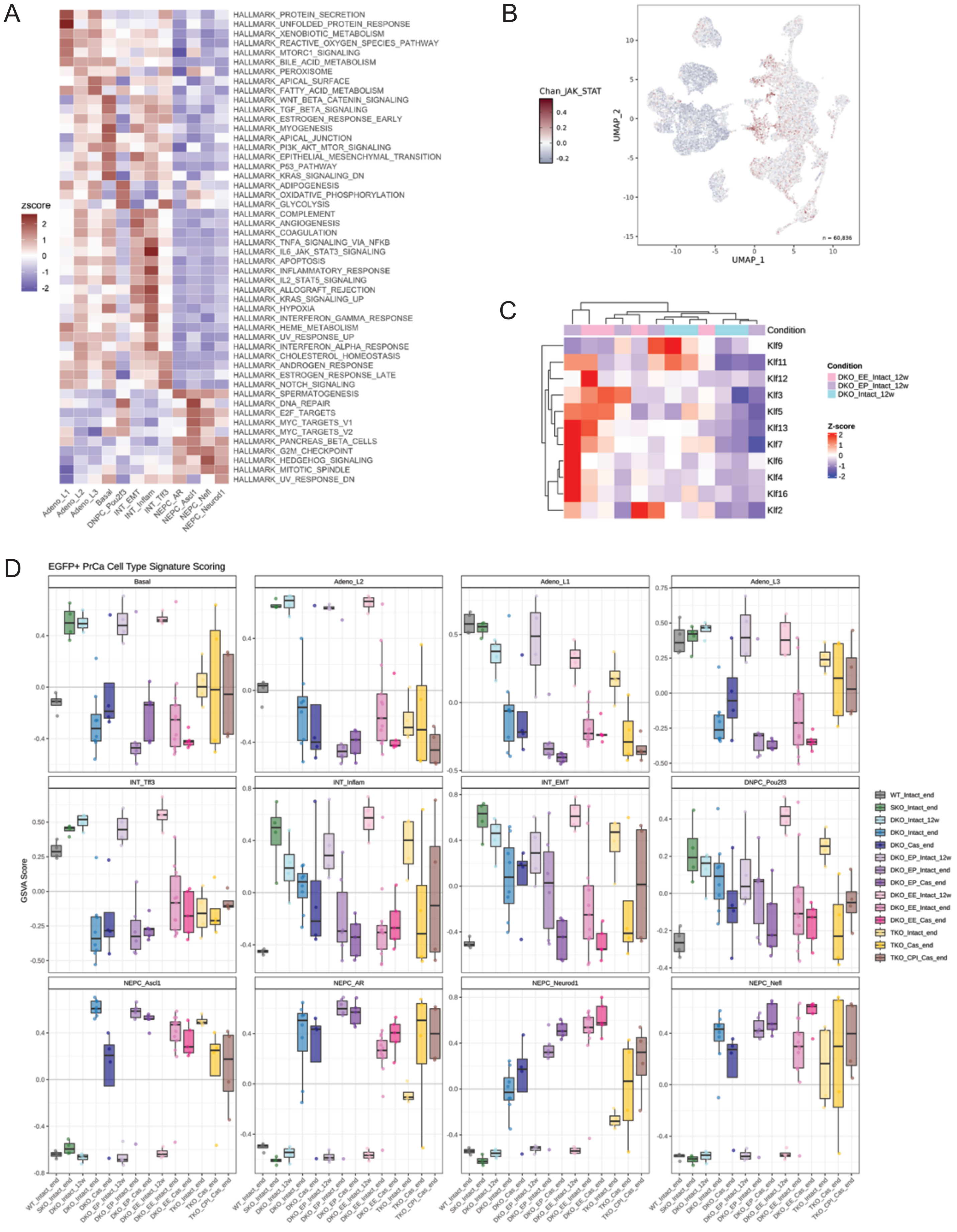

Supplementary Figure 4: Phentype of PrCa lineage variants developing in mice

*Supplementary Figure 4: Phenotype of PrCa lineage variants developing in mice.* A) scRNA-seq data from each mouse PrCa lineage variant was analyzed by GSEA using the Hallmark gene signatures set. The Z scores are displayed as a heatmap. B) The UMAP of mouse PrCa cells is color coded for similarity to the indicated inflammatory gene set that correlates with plasticity[56]. C) Bulk RNA-seq data from prostate tissue of 12 week old mice with the indicated genotypes was analyzed for Klf transcription factor family gene expression. The Z scores are displayed as a heatmap. D) Bulk RNA-seq data from prostate tissue samples of the indicated genotype, age (12W=12 weeks, end=end stage), and treatment (Cas=castrated, intact=not castrated) was analyzed for similarity to the indicated gene signatures by GSVA. Note Adeno_L2 at 12 weeks precedes development of NE lineage variants, DNPC_Pou2f3 is enriched in DKO_EE 12 weeks samples, and NEPC_Neurod1 is enriched in end stage Ezh2 deficient samples consistent with scRNA-seq data.

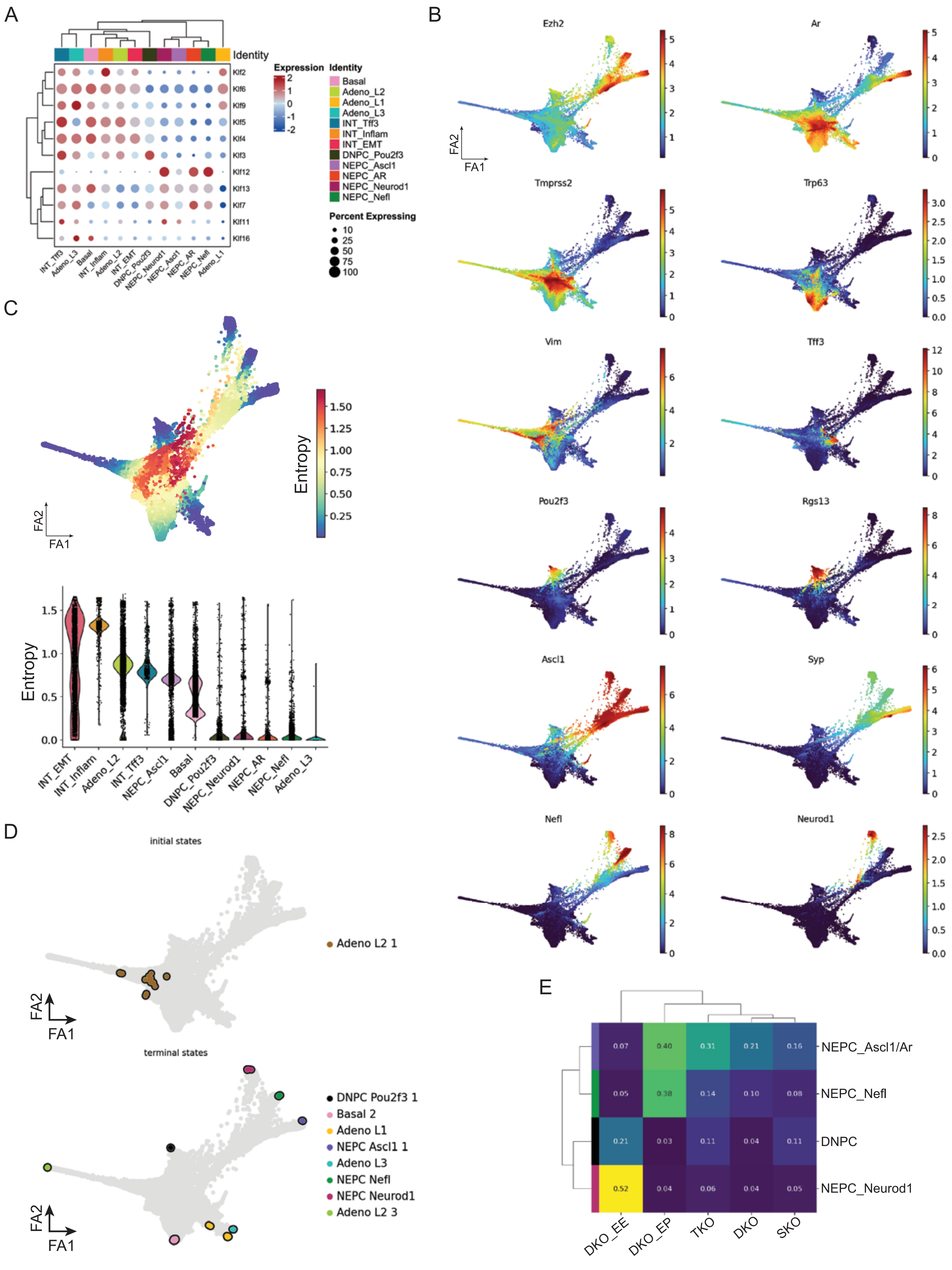

Supplementary Figure 5: *Ezh2* deficiency alters transcription factor activity and AVPC subtype

*Supplementary Figure 5: Ezh2 deficiency alters transcription factor activity and AVPC subtype evolution.* A) PrCa subtypes identified by scRNA-seq data were analyzed for Klf transcription factor family expression. Data is presented as a dot plot for each gene and subtype, color coded for expression level and size coded for percent of expressing cells. B) Force directed projection of PrCa scRNA-seq showing genotype specific differences in terminal AVPC lineage states color coded by expression of the indicated genes. C) Force directed PrCa cell projection color coded by palantir entropy that correlates with developmental potential (upper panel). The lower panel displays the range of palantir entropy in cells of the indicated PrCa subtypes. D) Terminal lineage states calculated from pseudotime analysis of scRNA-seq data using Adeno_L2 as the initial state[56]. E) The probability matrix of assuming the indicated AVPC terminal lineage states for PrCa cells of the listed genotypes.

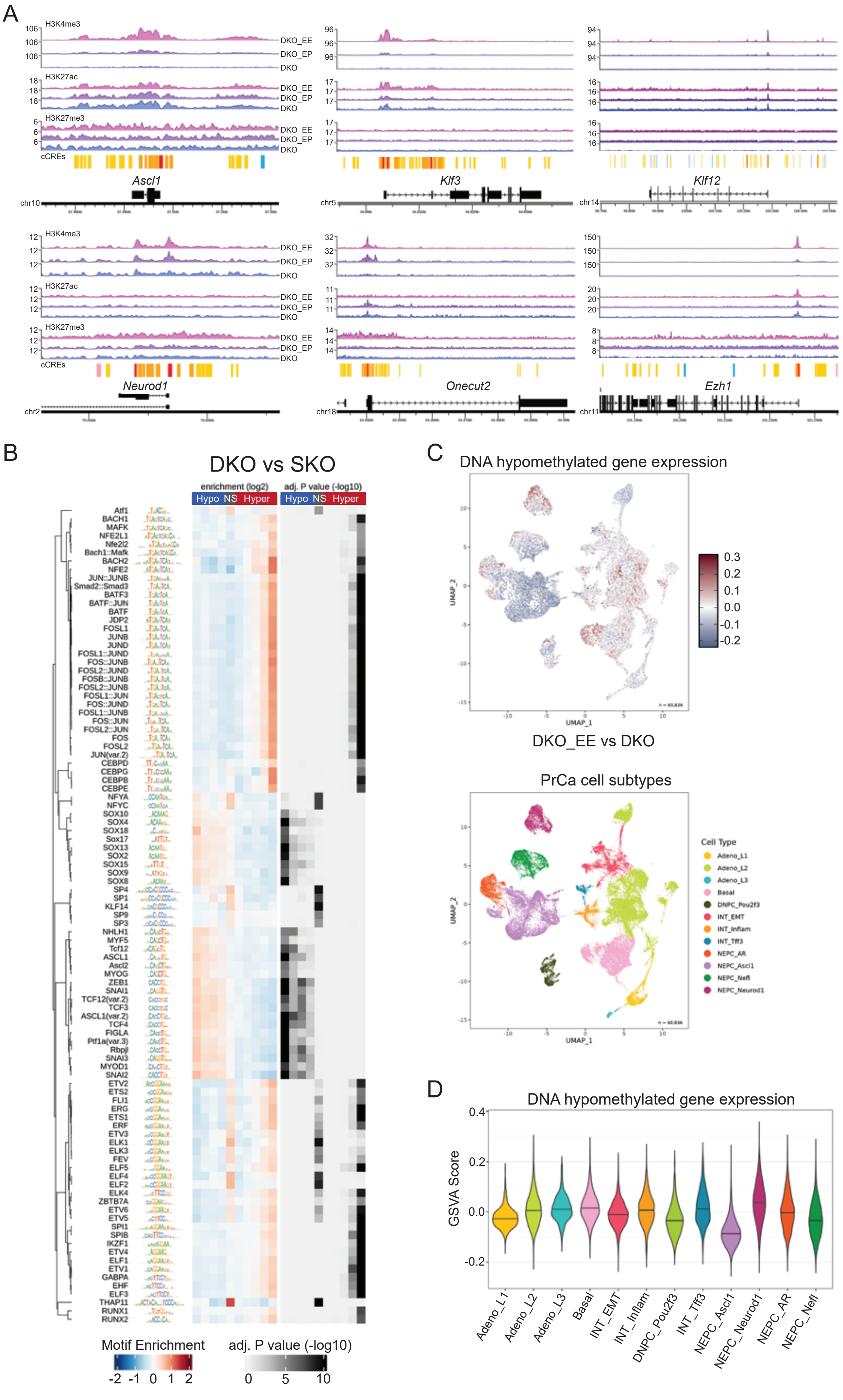

Supplementary Figure 6: Chromatin remodeling in PrCa AVPC lineage variants

*Figure S6: Chromatin remodeling in PrCa AVPC lineage variants.* A) ChIP-seq reads (upper panels) for the indicated histone marks and PrCa genotype are mapped on the listed genes (lower panels) and surrounding genomic region. The position of Encode determined candidate cis-regulatory elements (cCRE) are shown relative to these genes. B) Transcription factor motifs enriched in differentially DNA hypermethylated or hypomethylated regions comparing DKO versus SKO PrCa. Hypomethylated regions are enriched for motifs bound by transcription factors implicated in PrCa lineage plasticity. C) UMAP representation of single cell RNA-seq data from EGFP positive cancer cells color coded by expression levels of differentially DNA hypomethylated genes in DKO_EE versus DKO PrCa (top panel). UMAP representation color coded by PrCa cell subtype (bottom panel). Expression of DNA hypomethylated genes is higher in non-NEPC_Ascl1 subtype NEPC lineage variants enriched in *Ezh2* deficient samples. D) GSVA analysis of scRNA-seq data showing relative expression of DNA hypomethylated gene set in the DKO_EE versus DKO compmarison with the value of each cell displayed in a violin plot. The bar represents the median score highlighting increased expression in PrCa lineage variants enriched by *Ezh2* deficiency.

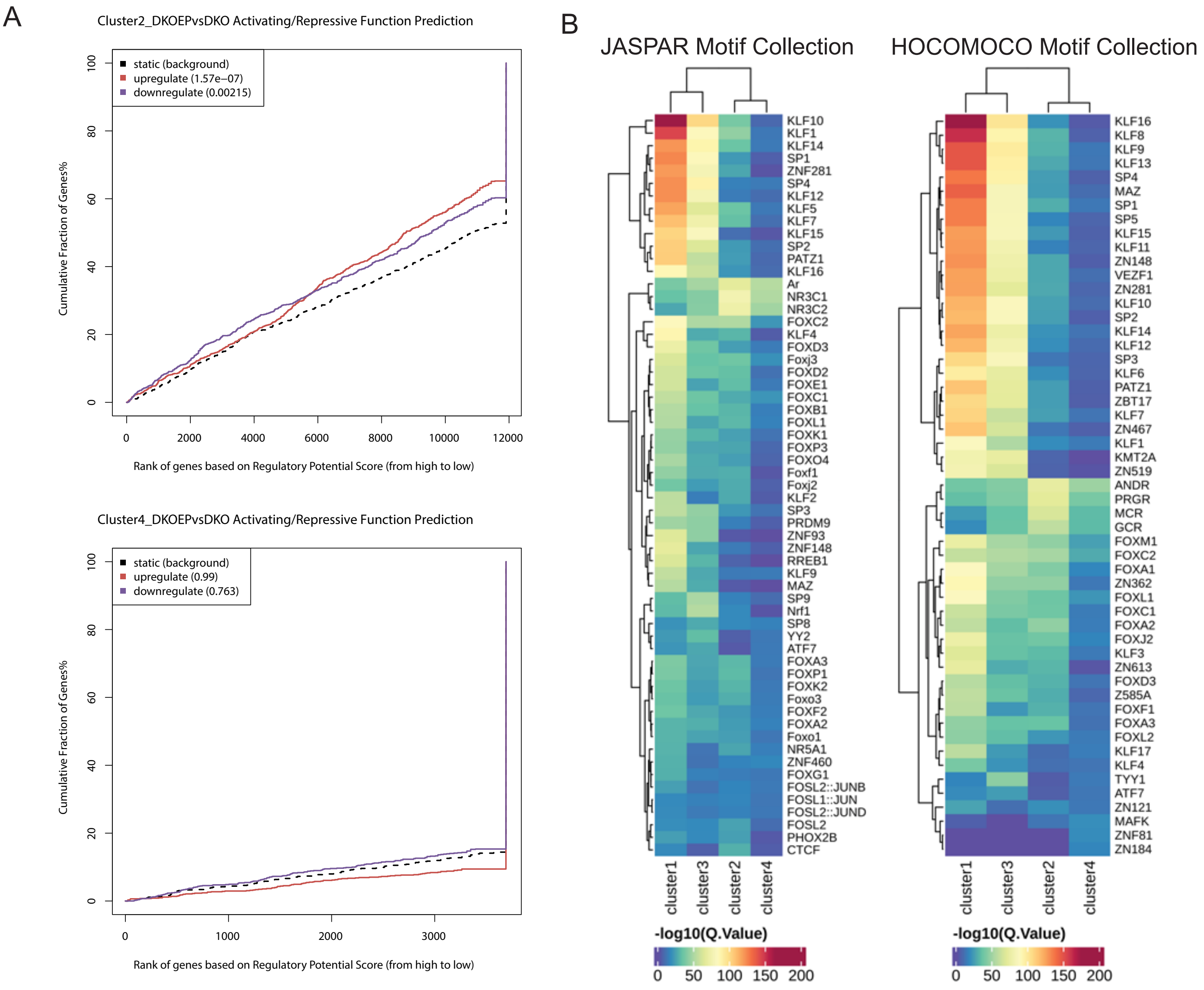

Supplementary Figure 7: AR genome binding patterns and transcription are altered in *Ezh2* deficient PrCa

*Figure S7: AR genome binding patterns and transcriptional effects are altered in Ezh2 deficient PrCa.* A) BETA analysis indicates that Ar binding sites in clusters 2 (upper panel) and 4 (lower panel) are not associated with marked changes in transcription, consistent with the relatively low fraction of these genes mapping near gene promoters. B) Transcription factor motif enrichment near Ar binding sites from each cluster of Ar binding highlighting enrichment of Klf transcription factor motifs in cluster 1 that are associated with increased transcription.

**Table S1 PCR primers and use**

| Template | Sequence | Band Size (bp) | Purpose/Source | Notes |
|---|---|---|---|---|
| *CRE* gene | 5'- GGA CAT GTT CAG GGA TCG CCA GGC G -3'<br>5'- GCA TAA CCA GTG AAA CAG CAT TGC TG -3' | 296 | PCR Genotyping/Tail DNA | |
| *Rb1* floxed | 5'- GGA ATT CCG GCG TGT GCC ATC AAT G -3'<br>5'- AGC TCT CAA GAG CTC AGA CTC ATG G -3' | WT: 247<br>FL: 295 | PCR Genotyping/Tail DNA | 2.5 - 3% agarose gel for separation |
| *Rb1* recombined | 5'- GGA ATT CCG GCG TGT GCC ATC AAT G -3'<br>5'- CGA AAG GAA AGT CAG GGA CAT TGG G -3' | 150 | PCR Gene deletion/Tumor DNA | |
| *Pten* floxed | 5'- CAA GCA CTC TGC GAA CTG AG -3'<br>5'- AAG TTT TTG AAG GCA AGA TGC -3' | WT: 156<br>FL: 328 | PCR Genotyping/Tail DNA | |
| *Pten* recombined | 5'- CAA GCA CTC TGC GAA CTG AG -3'<br>5'- GCT TGA TAT CGA ATT CCT GCA GC -3’ | 300 | PCR Gene deletion/Tumor | |
| *Ezh2* floxed | 5’ - CAT GTG CAG CTT TCT GTT CA – 3’<br>5’ - CAC AGC CTT TCT GCT CAC TG - 3’ | FL: 300<br>WT: 200 | PCR Genotyping/Tail | 2.5 - 3% agarose gel for separation |
| *Ezh2* recombined | 5’ - CCCATGTTTAAGGGCATAGTGACATG – 3’<br>5’ TCGAGGGACCTAATAACTCGTATAGCA -3’ | 200 | PCR Gene deletion/Tumor | |
| *Trp53* floxed | 5’-GTT AAG GGG TAT GAG GGA CAA GGT A-3’<br>5’-CCA TGA GAC AGG GTC TTG CTA TTG T-3’<br>5’-GTG CCC TCC GTC CTT TTT CGC AAT C-3’ | WT: 169<br>FL: 316 | PCR Genotyping/Tail | 2.5 - 3% agarose gel for separation |
| *Trp53* recombined | 5’-GTT AAG GGG TAT GAG GGA CAA GGT A-3’<br>5’-CCA TGA GAC AGG GTC TTG CTA TTG T-3’<br>5’-GTG CCC TCC GTC CTT TTT CGC AAT C-3’ | 198 | PCR Gene deletion/Tumor DNA | |
| *mT/mG* reporter | 5’-CTC TGC TGC CTC CTG GCT TCT -3’<br>5’-CGA GGC GGA TCA CAA GCA ATA-3’<br>5’-TCA ATG GGC GGG GGT CGT T-3’ | WT: 330<br>Tom: 250 | PCR Genotyping/Tail | |
| *Ezh2* mRNA | 5’- TCC CGT TAA AGA CCC TGA ATG-3’<br>5’- TGA AAG TGC CAT CCT GAT CC-3’ | NA | SYBR Green q-RT-PCR/ Tumor cDNA | Melt curve done with each run showing one amplicon |
| *L32* mRNA | 5’-TTC CTG GTC CAC AAC GTC AAG-3’<br>5’-TGT GAG CGA TCT CGG CAC-3’ | NA | SYBR Green q-RT-PCR/ Tumor cDNA | Melt curve done with each run showing one amplicon |

All genotyping reactions run on 1.5% agarose gel, unless noted otherwise

**Table S2: Antibodies and conditions used**

| Antibody | Purpose | Dilution/Protocol | Source |
|---|---|---|---|
| Ezh2 | IB/IHC | IB: 1:2000, 15 µg, 10 % gel, Pico ECL<br>IHC: 1:800, pH 6 | Cell Sig 5246 |
| GAPDH | IB | 1:10,000, 15 µg, 10 % gel, Pico ECL | Cell Sig 2118 |
| H3K27me3 | IB/IHC | IB: 1:2000, 3 µg, 12% gel, Pico ECL<br>IHC: 1:2000, pH 6 | Cell Sig 9733 |
| H3K27me3 | ChIP-seq | 5 µg | Active Motif 39055 |
| H3K27ac | ChIP-seq | 5 µg | Diagenode C15410196 |
| H3K4Me3 | ChIP-seq | 5 µg | Abcam 8580 |
| H3 | IB | 1:4000, 3 µg, 12% gel, ECL | Cell Sig 9715 |
| AR | IHC | 1:400, pH 6 | Santa Cruz sc-816 |
| AR | ChIP-seq | 5 µg | Abcam 108341 |
| Krt8 (CK8) | IHC | 1:400, pH 6 | Research Diagnostics RDI-PRO61038 |
| Synaptophysin | IHC | 1:500, pH 6 | Roche MRQ-40 |
| NeuroD1 | IHC | 1:60, pH 6 | Abcam ab205300 |
| Nefl | IHC | 1:100, pH 9 | ThermoFisher 3L9Y10 |

Table S3: Bulk RNA-seq DKO_EP vs DKO regulons

| TF | NES | rank |
|---|---|---|
| Zfp319 | 6.29882428 | 1 |
| Nfic | 5.57999774 | 2 |
| Arid1a | 5.42744085 | 3 |
| Cic | 5.28050205 | 4 |
| Prr12 | 5.26048481 | 5 |
| Casz1 | 4.70516208 | 6 |
| Hinfp | 4.65190815 | 7 |
| Dot1l | 4.6476028 | 8 |
| Foxp4 | 4.58107495 | 9 |
| Wiz | 4.32036292 | 10 |
| Zfp609 | 4.21155539 | 11 |
| Zfp628 | 4.20525379 | 12 |
| Terf2 | 4.20161087 | 13 |
| Baz2a | 4.1950958 | 14 |
| Sox1 | 4.19372498 | 15 |
| Trerf1 | 4.09788422 | 16 |
| Zfp395 | 4.0820575 | 17 |
| Rere | 4.06360509 | 18 |
| Foxj2 | 4.04722202 | 19 |
| Gmeb2 | 3.91798013 | 20 |
| Zfhx3 | 3.911362 | 21 |
| Zbtb7a | 3.85985864 | 22 |
| Klf13 | 3.79800169 | 23 |
| Foxk1 | 3.77304555 | 24 |
| Zfp687 | 3.75098007 | 25 |
| Zfp362 | 3.73792931 | 26 |
| Myrf | 3.71314371 | 27 |
| Mnt | 3.69031542 | 28 |
| Rela | 3.67202297 | 29 |
| Tead1 | 3.65713158 | 30 |
| Zfp646 | 3.64187449 | 31 |
| Mlxipl | 3.62596748 | 32 |
| Zfp335 | 3.61279743 | 33 |
| Rfx1 | 3.60377053 | 34 |
| Zfp579 | 3.58526615 | 35 |
| Deaf1 | 3.54386075 | 36 |
| Gon4l | 3.54108829 | 37 |
| Hivep3 | 3.49802668 | 38 |
| Mta1 | 3.49632371 | 39 |

| | | |
|---|---|---|
| Znf512b | 3.49602319 | 40 |
| Rfx2 | 3.48306503 | 41 |
| Pou5f2 | 3.4809088 | 42 |
| Cramp1l | 3.44086248 | 43 |
| Cux2 | 3.41131288 | 44 |
| E2f1 | 3.39885197 | 45 |
| Zfp692 | 3.396553 | 46 |
| Chd6 | 3.38267258 | 47 |
| Zfp827 | 3.37897353 | 48 |
| Ep400 | 3.37459142 | 49 |
| Zfp414 | 3.37121593 | 50 |
| Elf4 | 3.31517614 | 51 |
| Smarcc2 | 3.31083957 | 52 |
| Tada2b | 3.27242326 | 53 |
| Ascl1 | 3.26905787 | 54 |
| E2f7 | 3.26815132 | 55 |
| Mybl2 | 3.23240696 | 56 |
| Zfp111 | 3.22351199 | 57 |
| Foxm1 | 3.20428307 | 58 |
| Hivep1 | 3.16707157 | 59 |
| Prox1 | 3.16676921 | 60 |
| Bptf | 3.16343231 | 61 |
| Zbtb34 | 3.14452957 | 62 |
| Zfp592 | 3.14221989 | 63 |
| Irf2 | 3.1087433 | 64 |
| Arid1b | 3.10139867 | 65 |
| Sp2 | 3.08944337 | 66 |
| E2f8 | 3.07655211 | 67 |
| Mlxip | 3.0460955 | 68 |
| Zfp169 | 3.02320796 | 69 |
| Pou2f1 | 3.01717613 | 70 |
| Maz | 3.01638741 | 71 |
| Ppard | 3.00875003 | 72 |
| Mef2d | 2.99340633 | 73 |
| Foxn3 | 2.98477878 | 74 |
| E2f2 | 2.96662901 | 75 |
| Tbx3 | 2.9620925 | 76 |
| Zfp446 | 2.95999798 | 77 |
| Nfrkb | 2.95006618 | 78 |
| Etv6 | 2.9206022 | 79 |
| Gmeb1 | 2.9089719 | 80 |

| | | |
|---|---|---|
| Creb3l2 | 2.8929296 | 81 |
| Sox5 | 2.87502416 | 82 |
| Zhx3 | 2.83377121 | 83 |
| Zfp607 | 2.83112437 | 84 |
| Mafg | 2.82424023 | 85 |
| Arntl2 | 2.81776054 | 86 |
| Gatad2b | 2.81731975 | 87 |
| Nr2c2 | 2.8035412 | 88 |
| Ikzf4 | 2.8014771 | 89 |
| Tshz1 | 2.78572492 | 90 |
| Zfp652 | 2.78162326 | 91 |
| Zbtb40 | 2.78098433 | 92 |
| Zfp451 | 2.78085725 | 93 |
| Nfatc3 | 2.76729224 | 94 |
| Lmx1b | 2.75965113 | 95 |
| Zfp518b | 2.75023925 | 96 |
| Hivep2 | 2.741048 | 97 |
| Foxa1 | 2.73936635 | 98 |
| Hnf1b | 2.7184886 | 99 |
| Nr6a1 | 2.70284738 | 100 |
| Smarcc1 | 2.70276527 | 101 |
| Ski | 2.63119322 | 102 |
| Prdm10 | 2.62873791 | 103 |
| Baz2b | 2.624757 | 104 |
| Hmbox1 | 2.61919226 | 105 |
| Plagl2 | 2.61477745 | 106 |
| Nfat5 | 2.59320322 | 107 |
| Zfp418 | 2.58999563 | 108 |
| Yy2 | 2.57306589 | 109 |
| Zfp608 | 2.57092769 | 110 |
| Sox13 | 2.56328917 | 111 |
| Dnajc1 | 2.56067536 | 112 |
| Atf7 | 2.53588008 | 113 |
| Klf12 | 2.52658744 | 114 |
| Zfp276 | 2.51060048 | 115 |
| Gata3 | 2.50485116 | 116 |
| Zfp236 | 2.50434816 | 117 |
| Ybx2 | 2.4938718 | 118 |
| Rreb1 | 2.48956829 | 119 |
| Rbl1 | 2.47883368 | 120 |
| Foxo6 | 2.47479151 | 121 |

| | | |
|---|---|---|
| E2f4 | 2.45989699 | 122 |
| Zbtb38 | 2.45524326 | 123 |
| Hsf1 | 2.45149385 | 124 |
| Zfp580 | 2.43593819 | 125 |
| Rfx3 | 2.43231654 | 126 |
| Jazf1 | 2.43073773 | 127 |
| Elk4 | 2.43029782 | 128 |
| Foxj3 | 2.4269694 | 129 |
| Zbtb16 | 2.41331934 | 130 |
| Zfp612 | 2.37678203 | 131 |
| Rfxank | 2.36806746 | 132 |
| Zfhx4 | 2.32661717 | 133 |
| Smad2 | 2.31496164 | 134 |
| Stat5b | 2.30934623 | 135 |
| Zfp704 | 2.29432343 | 136 |
| Elk1 | 2.28084408 | 137 |
| Pds5b | 2.28020893 | 138 |
| Arnt | 2.26239945 | 139 |
| Rfx7 | 2.24991177 | 140 |
| Mga | 2.24917721 | 141 |
| Arid2 | 2.21343924 | 142 |
| Zbtb44 | 2.20895562 | 143 |
| Pknox2 | 2.19607693 | 144 |
| Setbp1 | 2.18105149 | 145 |
| Tfap2a | 2.17384904 | 146 |
| Mef2a | 2.15996727 | 147 |
| Sp1 | 2.14739776 | 148 |
| Zfp287 | 2.10336521 | 149 |
| Sim2 | 2.09221239 | 150 |
| Arnt2 | 2.07383982 | 151 |
| Tsc22d2 | 2.04638799 | 152 |
| Myt1 | 2.01292625 | 153 |
| Zfp329 | 1.99287629 | 154 |
| Mysm1 | 1.98198403 | 155 |
| Nr1d1 | 1.97309432 | 156 |
| Zbtb37 | 1.96186915 | 157 |
| Zfp280b | 1.95441513 | 158 |
| Stat2 | 1.94378121 | 159 |
| Ahctf1 | 1.92202426 | 160 |
| Hlf | 1.92092373 | 161 |
| Wdhd1 | 1.92073755 | 162 |

| | | |
|---|---|---|
| Foxo1 | 1.91894152 | 163 |
| Pbx4 | 1.91877827 | 164 |
| Zfp292 | 1.88097862 | 165 |
| Pou6f2 | 1.84341289 | 166 |
| Zfp445 | 1.8281874 | 167 |
| Bcl11a | 1.82683504 | 168 |
| Hnf4g | 1.81777643 | 169 |
| Sox10 | 1.81238648 | 170 |
| Tcf7l1 | 1.80892769 | 171 |
| Meis2 | 1.80849413 | 172 |
| Etv3 | 1.79821497 | 173 |
| Tfdp2 | 1.79505315 | 174 |
| Ezh1 | 1.78666669 | 175 |
| Klf7 | 1.78035391 | 176 |
| Zfp280d | 1.77907227 | 177 |
| Zfp280c | 1.77763301 | 178 |
| Onecut2 | 1.76917121 | 179 |
| Zfp41 | 1.76644297 | 180 |
| Rfx5 | 1.76090198 | 181 |
| E4f1 | 1.75352323 | 182 |
| Ctcfl | 1.74635816 | 183 |
| Zbed4 | 1.7462196 | 184 |
| Lin28a | 1.71772518 | 185 |
| Rlf | 1.70070546 | 186 |
| Pou2f3 | 1.69478851 | 187 |
| Ubp1 | 1.66755573 | 188 |
| Fosl2 | 1.66650207 | 189 |
| Lin28b | 1.64805525 | 190 |
| Notch4 | 1.6432335 | 191 |
| Nrf1 | 1.61118347 | 192 |
| Cux1 | 1.59787246 | 193 |
| Pbx3 | 1.52716399 | 194 |
| Foxc1 | 1.51191265 | 195 |
| Gata2 | 1.487613 | 196 |
| Ar | 1.47672426 | 197 |
| Tcf7l2 | 1.47103694 | 198 |
| Tfcp2 | 1.46296185 | 199 |
| Alx4 | 1.43038436 | 200 |
| Ttf1 | 1.40661383 | 201 |
| Zfp7 | 1.39856058 | 202 |
| Nr2c1 | 1.39708303 | 203 |

| | | |
|---|---|---|
| Nr4a2 | 1.3773116 | 204 |
| Phox2a | 1.36864668 | 205 |
| Zfp9 | 1.33032429 | 206 |
| Rel | 1.29278116 | 207 |
| Zeb1 | 1.29004071 | 208 |
| Arid3c | 1.28661262 | 209 |
| Chd7 | 1.25376777 | 210 |
| Bcl6 | 1.24388159 | 211 |
| Tub | 1.23112802 | 212 |
| Tbx6 | 1.22759166 | 213 |
| Sp8 | 1.19425957 | 214 |
| Alx3 | 1.1807285 | 215 |
| Zfpm1 | 1.16227538 | 216 |
| E2f6 | 1.11544411 | 217 |
| Atf2 | 1.11055229 | 218 |
| Nkx2-1 | 1.10542833 | 219 |
| Ctcf | 1.09981496 | 220 |
| Vezf1 | 1.07794406 | 221 |
| Runx3 | 1.06803395 | 222 |
| Smad9 | 0.99629732 | 223 |
| Tead3 | 0.96457979 | 224 |
| Nkx2-6 | 0.92445027 | 225 |
| Nfia | 0.91706465 | 226 |
| Pbx2 | 0.9165204 | 227 |
| Hoxb5 | 0.90758688 | 228 |
| Zfp2 | 0.90071077 | 229 |
| Tox | 0.89462697 | 230 |
| Mybl1 | 0.89314538 | 231 |
| St18 | 0.88881309 | 232 |
| Foxo4 | 0.86642966 | 233 |
| Tcf7 | 0.84605212 | 234 |
| Zfp595 | 0.84140918 | 235 |
| Phox2b | 0.78959116 | 236 |
| Six1 | 0.76378059 | 237 |
| Batf | 0.7168713 | 238 |
| Tgif1 | 0.70950683 | 239 |
| Zfp605 | 0.7058037 | 240 |
| Sall1 | 0.70272029 | 241 |
| Hmgb3 | 0.68748432 | 242 |
| Gata5 | 0.67520181 | 243 |
| Foxa2 | 0.66251192 | 244 |

| | | |
|---|---|---|
| Zfp157 | 0.66062586 | 245 |
| Nkx3-1 | 0.64869758 | 246 |
| Lef1 | 0.64305334 | 247 |
| Epas1 | 0.62528798 | 248 |
| 1700049G17I | 0.62292708 | 249 |
| Nrk | 0.5912409 | 250 |
| Six5 | 0.58894989 | 251 |
| Zscan21 | 0.56074927 | 252 |
| Notch2 | 0.55496499 | 253 |
| Hoxb7 | 0.54605889 | 254 |
| Atf6 | 0.53998123 | 255 |
| Gfi1 | 0.53755729 | 256 |
| Zbtb26 | 0.52001148 | 257 |
| Fosb | 0.48502564 | 258 |
| Esrrb | 0.4828758 | 259 |
| Gli1 | 0.46880945 | 260 |
| Zfp710 | 0.46122801 | 261 |
| Neurod1 | 0.46013625 | 262 |
| Tead4 | 0.42156188 | 263 |
| Egr1 | 0.42142056 | 264 |
| Notch1 | 0.4083096 | 265 |
| Zfp512 | 0.34114582 | 266 |
| Rfx4 | 0.31082759 | 267 |
| Foxk2 | 0.31059944 | 268 |
| Hsf4 | 0.30525861 | 269 |
| Zfp618 | 0.282834 | 270 |
| Klf9 | 0.2578059 | 271 |
| Rora | 0.21751817 | 272 |
| Nfe2l1 | 0.21371542 | 273 |
| Zfp821 | 0.18498337 | 274 |
| Mafk | 0.17326829 | 275 |
| Six4 | 0.1700507 | 276 |
| Irf5 | 0.10471842 | 277 |
| Thrb | 0.07022373 | 278 |
| Zfp354c | 0.02934481 | 279 |
| Nr3c2 | 0.01363947 | 280 |
| Smad4 | -0.042484 | 281 |
| Pparg | -0.0566222 | 282 |
| Nr4a1 | -0.1175292 | 283 |
| Rxra | -0.121192 | 284 |
| Elf2 | -0.1323123 | 285 |

| | | |
|---|---|---|
| Sox3 | -0.1370747 | 286 |
| Nr2f2 | -0.1388307 | 287 |
| Mafb | -0.146971 | 288 |
| Bhlhe40 | -0.1886839 | 289 |
| Sp4 | -0.1980934 | 290 |
| Grhl1 | -0.2084688 | 291 |
| Bhlhe22 | -0.2621374 | 292 |
| Hoxa10 | -0.2871883 | 293 |
| Nfatc4 | -0.2946331 | 294 |
| Stat1 | -0.2998138 | 295 |
| Mef2c | -0.3115761 | 296 |
| Bcl11b | -0.3234032 | 297 |
| Hoxc5 | -0.3409158 | 298 |
| Klf2 | -0.3515648 | 299 |
| Sox2 | -0.3551737 | 300 |
| Klf15 | -0.3635993 | 301 |
| Nfkb1 | -0.4008876 | 302 |
| Ebf3 | -0.4088076 | 303 |
| Mis18bp1 | -0.4578667 | 304 |
| Zfp219 | -0.4681368 | 305 |
| Irf7 | -0.4750377 | 306 |
| Atoh8 | -0.4951159 | 307 |
| Elk3 | -0.508889 | 308 |
| Heyl | -0.5331233 | 309 |
| Ebf1 | -0.5406235 | 310 |
| Bhlha15 | -0.54065 | 311 |
| Trp73 | -0.5519336 | 312 |
| Gata6 | -0.570255 | 313 |
| Dach1 | -0.574571 | 314 |
| Egr3 | -0.5943393 | 315 |
| Nfib | -0.6032867 | 316 |
| Isx | -0.6040726 | 317 |
| Ets1 | -0.6273997 | 318 |
| Rbl2 | -0.6308756 | 319 |
| Zfp521 | -0.6344956 | 320 |
| Tsc22d4 | -0.6362985 | 321 |
| Ikzf3 | -0.6645232 | 322 |
| Etv1 | -0.6945236 | 323 |
| Arntl | -0.7207549 | 324 |
| Runx1 | -0.721599 | 325 |
| Smad3 | -0.7590942 | 326 |

| | | |
|---|---|---|
| Hsf2 | -0.7607701 | 327 |
| Nr0b2 | -0.7674201 | 328 |
| Zfp691 | -0.7714187 | 329 |
| Tsc22d3 | -0.7797118 | 330 |
| Pknox1 | -0.786509 | 331 |
| Cbx2 | -0.7880757 | 332 |
| Sox7 | -0.7937937 | 333 |
| Noc3l | -0.8118584 | 334 |
| Hoxa11 | -0.8200929 | 335 |
| Lyl1 | -0.8406271 | 336 |
| Stat4 | -0.84076 | 337 |
| Tcf23 | -0.8423518 | 338 |
| Tulp2 | -0.8452244 | 339 |
| Maf | -0.8495362 | 340 |
| Zbtb41 | -0.855204 | 341 |
| Phtf1 | -0.8557879 | 342 |
| Esrrg | -0.8629003 | 343 |
| Lhx2 | -0.9058357 | 344 |
| Foxp3 | -0.9120187 | 345 |
| Klf4 | -0.9219001 | 346 |
| Foxp1 | -0.9338409 | 347 |
| Hand2 | -0.9545525 | 348 |
| Isl1 | -0.9839422 | 349 |
| Trp63 | -0.988078 | 350 |
| Bnc2 | -0.990503 | 351 |
| Mecom | -0.9954228 | 352 |
| Rarg | -0.9984531 | 353 |
| Nab2 | -1.0002949 | 354 |
| Tbx2 | -1.0085056 | 355 |
| Hmgxb4 | -1.0101191 | 356 |
| Rbpj | -1.0124939 | 357 |
| Dzip1l | -1.0125384 | 358 |
| Klf3 | -1.0290339 | 359 |
| Phtf2 | -1.049178 | 360 |
| Creb1 | -1.0507714 | 361 |
| Six2 | -1.0520271 | 362 |
| Ovol1 | -1.0830096 | 363 |
| Myb | -1.0832771 | 364 |
| Egr2 | -1.0864514 | 365 |
| Eomes | -1.0990493 | 366 |
| Arid3a | -1.108297 | 367 |

| | | |
|---|---|---|
| Prrx1 | -1.1157616 | 368 |
| Stat6 | -1.118489 | 369 |
| Tulp1 | -1.122397 | 370 |
| Zfp644 | -1.1394237 | 371 |
| Six3 | -1.1583407 | 372 |
| Ebf2 | -1.1720404 | 373 |
| Sox6 | -1.1722972 | 374 |
| Pou2af1 | -1.1819452 | 375 |
| Sox4 | -1.1972337 | 376 |
| Elf1 | -1.1999458 | 377 |
| Osr1 | -1.1999524 | 378 |
| Irf4 | -1.2120542 | 379 |
| Tcf3 | -1.2405047 | 380 |
| Myc | -1.2559829 | 381 |
| Tfcp2l1 | -1.2701926 | 382 |
| Tlx3 | -1.2775537 | 383 |
| Ahr | -1.278984 | 384 |
| Nfkbia | -1.3335311 | 385 |
| Satb2 | -1.3442197 | 386 |
| Irx3 | -1.3481907 | 387 |
| Emx2 | -1.3516087 | 388 |
| Twist2 | -1.3521265 | 389 |
| Runx2 | -1.4258243 | 390 |
| Tead2 | -1.4352159 | 391 |
| Nfkb2 | -1.4501812 | 392 |
| Pax9 | -1.4591615 | 393 |
| Satb1 | -1.4611203 | 394 |
| Zbtb7b | -1.4612984 | 395 |
| Batf3 | -1.4680812 | 396 |
| Hoxc8 | -1.4843041 | 397 |
| Pou3f3 | -1.5035336 | 398 |
| Hnf4a | -1.5129501 | 399 |
| Esr1 | -1.5404834 | 400 |
| Ubtf | -1.5654371 | 401 |
| Aebp1 | -1.5819316 | 402 |
| Rara | -1.5910718 | 403 |
| Sox17 | -1.5929267 | 404 |
| Foxp2 | -1.6037068 | 405 |
| Zfp715 | -1.6113932 | 406 |
| Nr5a2 | -1.6290933 | 407 |
| Tfec | -1.6588089 | 408 |

| | | |
|---|---|---|
| Max | -1.6590523 | 409 |
| Stat5a | -1.6590792 | 410 |
| Sox18 | -1.6606609 | 411 |
| Tfdp1 | -1.6636678 | 412 |
| Fli1 | -1.6873176 | 413 |
| Bach1 | -1.6878519 | 414 |
| Tcfl5 | -1.6879151 | 415 |
| Nfyc | -1.715686 | 416 |
| Tfap2c | -1.7322936 | 417 |
| Ezh2 | -1.7340053 | 418 |
| Ssrp1 | -1.7412267 | 419 |
| Olig3 | -1.7421871 | 420 |
| Zfp358 | -1.749817 | 421 |
| Nfkbie | -1.750767 | 422 |
| Sox9 | -1.7937025 | 423 |
| Pax8 | -1.7965754 | 424 |
| Atoh1 | -1.797356 | 425 |
| Tcf21 | -1.8058976 | 426 |
| Ikzf2 | -1.8414915 | 427 |
| Hes6 | -1.8694839 | 428 |
| Wt1 | -1.8836772 | 429 |
| Zfp518a | -1.8852532 | 430 |
| Relb | -1.8870402 | 431 |
| Bnc1 | -1.8890133 | 432 |
| Irf8 | -1.9095041 | 433 |
| Npas2 | -1.9193502 | 434 |
| Tal1 | -1.9246345 | 435 |
| Twist1 | -1.9405658 | 436 |
| Tbr1 | -1.9405745 | 437 |
| Rxrg | -1.9412039 | 438 |
| Mxd3 | -1.9547841 | 439 |
| Erg | -1.9615672 | 440 |
| Fos | -1.978243 | 441 |
| Foxq1 | -2.0024263 | 442 |
| Hes1 | -2.0085029 | 443 |
| Tbx5 | -2.0118096 | 444 |
| Rest | -2.0130713 | 445 |
| Nab1 | -2.0176722 | 446 |
| Nfkbib | -2.0200923 | 447 |
| Zfp593 | -2.0202155 | 448 |
| Nr1h4 | -2.0319082 | 449 |

| | | |
|---|---|---|
| Carhsp1 | -2.0657153 | 450 |
| Gatad1 | -2.0732285 | 451 |
| Zfp712 | -2.0800483 | 452 |
| Zfp791 | -2.1089264 | 453 |
| Zfp422 | -2.120809 | 454 |
| Irf3 | -2.1247744 | 455 |
| Cebpd | -2.1491076 | 456 |
| Mta2 | -2.2319653 | 457 |
| Hoxd12 | -2.2387771 | 458 |
| Atf4 | -2.2480419 | 459 |
| Tcf12 | -2.2496999 | 460 |
| Atf3 | -2.2514584 | 461 |
| Zfp110 | -2.2593246 | 462 |
| Scx | -2.2727907 | 463 |
| Cebpa | -2.2941894 | 464 |
| Vdr | -2.2968295 | 465 |
| Nkx6-2 | -2.297094 | 466 |
| Zfp91 | -2.3085538 | 467 |
| Mrpl28 | -2.316872 | 468 |
| Zfp51 | -2.3214758 | 469 |
| Hoxb13 | -2.3780543 | 470 |
| Hmgb2 | -2.3884169 | 471 |
| Elf5 | -2.424881 | 472 |
| Terf2ip | -2.4257573 | 473 |
| Zfp768 | -2.4365165 | 474 |
| Mycn | -2.4514786 | 475 |
| Fosl1 | -2.4590385 | 476 |
| Mitf | -2.4696693 | 477 |
| Nfyb | -2.4792302 | 478 |
| Hmg20b | -2.4863219 | 479 |
| Spdef | -2.4975272 | 480 |
| Ehf | -2.5105937 | 481 |
| Esrra | -2.5116182 | 482 |
| Nr1h2 | -2.5139648 | 483 |
| Junb | -2.5236477 | 484 |
| Smarca5 | -2.5316775 | 485 |
| Zfp260 | -2.5433886 | 486 |
| Cbfb | -2.5754363 | 487 |
| Zfp191 | -2.5916732 | 488 |
| Crem | -2.5938974 | 489 |
| Hif1a | -2.59428 | 490 |

| | | |
|---|---|---|
| Drap1 | -2.6213206 | 491 |
| Ybx1 | -2.6315225 | 492 |
| Nr4a3 | -2.6478342 | 493 |
| Ppara | -2.6927579 | 494 |
| Hoxa13 | -2.7114787 | 495 |
| Hoxd13 | -2.7565095 | 496 |
| Jun | -2.7889759 | 497 |
| Creb3l4 | -2.7990426 | 498 |
| Nr2f6 | -2.8119109 | 499 |
| Nfe2l2 | -2.8231601 | 500 |
| Zfp771 | -2.8276286 | 501 |
| Zfp207 | -2.8297568 | 502 |
| Creb3 | -2.8480811 | 503 |
| Sp3 | -2.8726702 | 504 |
| Tfam | -2.9427608 | 505 |
| Mxd4 | -2.9433998 | 506 |
| Cdc5l | -2.9761396 | 507 |
| Nfil3 | -2.9781493 | 508 |
| Zdhhc13 | -3.0733958 | 509 |
| Jund | -3.2152459 | 510 |
| Smarce1 | -3.2370383 | 511 |
| Csde1 | -3.3875856 | 512 |
| Cebpz | -3.3973312 | 513 |
| Dr1 | -3.4943632 | 514 |
| Cebpb | -3.5884239 | 515 |
| Bzw1 | -3.8918979 | 516 |

Table S3: Bulk RNA-seq DKO_EE vs DKO regulons

| TF | NES | rank |
|---|---|---|
| Nr6a1 | 8.38262196 | 1 |
| E2f8 | 5.70344262 | 2 |
| Zbtb38 | 5.67386229 | 3 |
| Nr3c2 | 5.61258066 | 4 |
| Gmeb2 | 5.45639262 | 5 |
| Zfp609 | 5.17881998 | 6 |
| Tshz1 | 5.16163981 | 7 |
| Rfx2 | 5.11773099 | 8 |
| Prdm10 | 5.08920607 | 9 |
| E2f4 | 5.07911167 | 10 |
| Foxj2 | 5.05023642 | 11 |
| Lin28b | 4.9834209 | 12 |
| Rora | 4.93003689 | 13 |
| Arntl | 4.81710076 | 14 |
| Hinfp | 4.81049424 | 15 |
| Dnajc1 | 4.79183168 | 16 |
| Arid1a | 4.73256283 | 17 |
| Klf13 | 4.73106139 | 18 |
| Foxm1 | 4.65860248 | 19 |
| Rere | 4.65183664 | 20 |
| Klf12 | 4.63339559 | 21 |
| Nfrkb | 4.61666041 | 22 |
| Znf512b | 4.59459421 | 23 |
| Gmeb1 | 4.5777952 | 24 |
| Wiz | 4.51732243 | 25 |
| Nr2c2 | 4.51223523 | 26 |
| Nr4a2 | 4.48108679 | 27 |
| Smarcc2 | 4.47264014 | 28 |
| Zfp280b | 4.46398013 | 29 |
| Foxk1 | 4.44086041 | 30 |
| Terf2 | 4.42696486 | 31 |
| Gatad2b | 4.40851296 | 32 |
| Plagl2 | 4.40765595 | 33 |
| E2f7 | 4.40581741 | 34 |
| Sp8 | 4.38867421 | 35 |
| Zhx3 | 4.35084792 | 36 |
| Zfp618 | 4.3322635 | 37 |
| Baz2a | 4.30990594 | 38 |
| Hivep2 | 4.29738879 | 39 |

| | | |
|---|---|---|
| Irf2 | 4.29717232 | 40 |
| Pou2f1 | 4.27840722 | 41 |
| Zfp287 | 4.2502972 | 42 |
| Baz2b | 4.24785209 | 43 |
| Casz1 | 4.21131723 | 44 |
| Elk4 | 4.19150505 | 45 |
| Nab2 | 4.18898284 | 46 |
| Hivep3 | 4.1331772 | 47 |
| Esrrb | 4.12471896 | 48 |
| Dot1l | 4.11028325 | 49 |
| Fosb | 4.07496898 | 50 |
| Zbtb34 | 4.0449867 | 51 |
| Zfp827 | 4.03990984 | 52 |
| Meis2 | 4.01288824 | 53 |
| Rlf | 3.98990057 | 54 |
| Bptf | 3.95525491 | 55 |
| Tada2b | 3.92479945 | 56 |
| Ezh1 | 3.89330086 | 57 |
| E2f2 | 3.89145014 | 58 |
| Arid2 | 3.89119872 | 59 |
| Ascl1 | 3.8746144 | 60 |
| Arid1b | 3.87287051 | 61 |
| Zfp646 | 3.85309752 | 62 |
| Creb3l2 | 3.83621747 | 63 |
| Zfp169 | 3.80941018 | 64 |
| Etv6 | 3.79650611 | 65 |
| E2f1 | 3.76600648 | 66 |
| Hmbox1 | 3.75935406 | 67 |
| Ep400 | 3.75851081 | 68 |
| Ubp1 | 3.7437452 | 69 |
| Trerf1 | 3.74136593 | 70 |
| Nfat5 | 3.70303874 | 71 |
| Zfhx3 | 3.69298791 | 72 |
| Bcl11a | 3.65548476 | 73 |
| Mlxipl | 3.63114233 | 74 |
| Zfp652 | 3.62739167 | 75 |
| Nfib | 3.62404719 | 76 |
| Atf7 | 3.57071179 | 77 |
| Pbx4 | 3.54473351 | 78 |
| Pbx2 | 3.53179813 | 79 |
| Atf6 | 3.51682605 | 80 |

| | | |
|---|---|---|
| Foxo1 | 3.4575483 | 81 |
| Mnt | 3.45612496 | 82 |
| Foxn3 | 3.43676203 | 83 |
| Tead3 | 3.43153014 | 84 |
| Rbl1 | 3.41744901 | 85 |
| Foxc1 | 3.40633794 | 86 |
| Rfx5 | 3.37711621 | 87 |
| Zfp111 | 3.37668621 | 88 |
| Bhlhe40 | 3.3634702 | 89 |
| Rfx4 | 3.36023585 | 90 |
| Pbx3 | 3.33736339 | 91 |
| Bcl6 | 3.33683927 | 92 |
| Tfdp2 | 3.31324802 | 93 |
| Chd6 | 3.31033132 | 94 |
| Zfp362 | 3.30563898 | 95 |
| Foxp4 | 3.29025625 | 96 |
| Foxj3 | 3.2573235 | 97 |
| Nfatc3 | 3.21590928 | 98 |
| Mlxip | 3.20386036 | 99 |
| Sp1 | 3.18508792 | 100 |
| Foxo6 | 3.18078399 | 101 |
| Smarcc1 | 3.17186386 | 102 |
| Klf3 | 3.1701111 | 103 |
| Zfp445 | 3.15974766 | 104 |
| Zfp592 | 3.13845384 | 105 |
| Cux2 | 3.13637006 | 106 |
| Nr1d1 | 3.12990262 | 107 |
| Hivep1 | 3.10425797 | 108 |
| Cramp1l | 3.09880024 | 109 |
| Arnt | 3.0980113 | 110 |
| Zfp446 | 3.09777713 | 111 |
| Zfp292 | 3.09664444 | 112 |
| Deaf1 | 3.08400068 | 113 |
| Rfx7 | 3.05942204 | 114 |
| Zfp687 | 3.04957966 | 115 |
| Mysm1 | 3.04420053 | 116 |
| Zfp335 | 3.03545471 | 117 |
| Prr12 | 3.03173088 | 118 |
| Arntl2 | 3.01117465 | 119 |
| Hnf1b | 3.00284597 | 120 |
| Tfap2a | 2.99309982 | 121 |

| | | |
|---|---|---|
| Tead1 | 2.97462254 | 122 |
| Smad2 | 2.96781991 | 123 |
| Mef2a | 2.95188191 | 124 |
| Zfp451 | 2.93468964 | 125 |
| Ovol1 | 2.90531882 | 126 |
| Nfic | 2.88501162 | 127 |
| Elf4 | 2.88399925 | 128 |
| Zbtb7a | 2.88214646 | 129 |
| Ppard | 2.8757557 | 130 |
| Runx2 | 2.87175136 | 131 |
| Sp2 | 2.84742488 | 132 |
| Zfp280d | 2.82420324 | 133 |
| Egr1 | 2.81819579 | 134 |
| Gata2 | 2.81782107 | 135 |
| Zfp395 | 2.81249278 | 136 |
| Rel | 2.80453008 | 137 |
| Gon4l | 2.76738821 | 138 |
| Ikzf2 | 2.73500306 | 139 |
| Zbtb16 | 2.72710054 | 140 |
| Stat2 | 2.72611026 | 141 |
| Pds5b | 2.72010948 | 142 |
| Neurod1 | 2.71561987 | 143 |
| Grhl1 | 2.67013662 | 144 |
| Setbp1 | 2.66423235 | 145 |
| Mga | 2.64840027 | 146 |
| Zfp512 | 2.64562993 | 147 |
| Klf15 | 2.64493546 | 148 |
| Mta1 | 2.63260614 | 149 |
| Maz | 2.61775418 | 150 |
| Zfp579 | 2.59755448 | 151 |
| Gfi1 | 2.59556341 | 152 |
| Rfx1 | 2.59276717 | 153 |
| Ctcf | 2.5794973 | 154 |
| Zfp692 | 2.57528294 | 155 |
| Hlf | 2.53072432 | 156 |
| Rreb1 | 2.52716911 | 157 |
| Ttf1 | 2.50759355 | 158 |
| Zbtb44 | 2.50652662 | 159 |
| Jazf1 | 2.48383384 | 160 |
| Yy2 | 2.4765977 | 161 |
| Prox1 | 2.47146389 | 162 |

| | | |
|---|---|---|
| Myb | 2.43520606 | 163 |
| Chd7 | 2.42751602 | 164 |
| Elk1 | 2.3982487 | 165 |
| Foxp1 | 2.37345966 | 166 |
| Fos | 2.37046757 | 167 |
| Zfp628 | 2.36649156 | 168 |
| Cic | 2.36440307 | 169 |
| Tfcp2l1 | 2.36203525 | 170 |
| Ahctf1 | 2.35241065 | 171 |
| Zfp236 | 2.32882228 | 172 |
| Nkx2-6 | 2.31841408 | 173 |
| Foxa2 | 2.24962712 | 174 |
| Pou5f2 | 2.24696633 | 175 |
| Irf7 | 2.23609124 | 176 |
| Zfp580 | 2.22159559 | 177 |
| Tgif1 | 2.19510829 | 178 |
| Pou6f2 | 2.15632389 | 179 |
| Zfp319 | 2.13417517 | 180 |
| Zfp9 | 2.12666406 | 181 |
| Tox | 2.10126729 | 182 |
| Elf1 | 2.09869013 | 183 |
| Zfhx4 | 2.09416377 | 184 |
| Zfp518b | 2.08241572 | 185 |
| Mef2d | 2.06591545 | 186 |
| Zfp280c | 2.05720304 | 187 |
| Pou2f3 | 2.01594682 | 188 |
| Tcf7l1 | 2.00680585 | 189 |
| Olig3 | 1.99416568 | 190 |
| Nrf1 | 1.94630708 | 191 |
| Zbtb37 | 1.93090333 | 192 |
| Stat1 | 1.92018159 | 193 |
| Nfia | 1.88752791 | 194 |
| Nfil3 | 1.88718278 | 195 |
| Tsc22d2 | 1.84118939 | 196 |
| Nr2c1 | 1.8165536 | 197 |
| Notch2 | 1.76973836 | 198 |
| Phox2a | 1.75953577 | 199 |
| Sox5 | 1.74775511 | 200 |
| Elf5 | 1.71715912 | 201 |
| Pknox1 | 1.69040897 | 202 |
| Mafg | 1.67464296 | 203 |

| | | |
|---|---|---|
| Mef2c | 1.66822089 | 204 |
| Vezf1 | 1.65914458 | 205 |
| Zfp691 | 1.64306637 | 206 |
| Npas2 | 1.64270728 | 207 |
| Zfp607 | 1.64109153 | 208 |
| Wdhd1 | 1.64012427 | 209 |
| Sp4 | 1.57572766 | 210 |
| Ctcfl | 1.55443261 | 211 |
| Trp63 | 1.55112776 | 212 |
| Zfp414 | 1.54416613 | 213 |
| Klf7 | 1.52443435 | 214 |
| Foxa1 | 1.52056589 | 215 |
| Stat6 | 1.51364349 | 216 |
| Foxp2 | 1.51305719 | 217 |
| Mybl2 | 1.49760045 | 218 |
| Mybl1 | 1.49483419 | 219 |
| Hoxb5 | 1.46802655 | 220 |
| Zbtb40 | 1.44220881 | 221 |
| Runx3 | 1.41783698 | 222 |
| Ikzf4 | 1.41485757 | 223 |
| Ski | 1.36696502 | 224 |
| Epas1 | 1.36073544 | 225 |
| Sox10 | 1.34080013 | 226 |
| Nfkb1 | 1.33014823 | 227 |
| Trp73 | 1.28263762 | 228 |
| Tcf7 | 1.2697299 | 229 |
| Klf9 | 1.23372333 | 230 |
| Notch1 | 1.22415045 | 231 |
| Nfe2l1 | 1.22306984 | 232 |
| Ybx2 | 1.19446142 | 233 |
| Tbx6 | 1.19171089 | 234 |
| Esrrg | 1.18296917 | 235 |
| Isx | 1.16658081 | 236 |
| Sox1 | 1.15694426 | 237 |
| Nrk | 1.10854097 | 238 |
| Ehf | 1.10389977 | 239 |
| Tead4 | 1.1013694 | 240 |
| Etv3 | 1.09616317 | 241 |
| Runx1 | 1.07647494 | 242 |
| Zfp612 | 1.07142966 | 243 |
| Lef1 | 1.06601783 | 244 |

| | | |
|---|---|---|
| Gli1 | 0.99521124 | 245 |
| Smad4 | 0.99516977 | 246 |
| Mafk | 0.984485 | 247 |
| E4f1 | 0.95507448 | 248 |
| Atf2 | 0.95152856 | 249 |
| Zfp329 | 0.93893801 | 250 |
| Arid3c | 0.92938692 | 251 |
| Tcf7l2 | 0.91564529 | 252 |
| Rarg | 0.90945457 | 253 |
| Phox2b | 0.9029137 | 254 |
| Nr4a1 | 0.88665646 | 255 |
| Hoxb7 | 0.86290866 | 256 |
| E2f6 | 0.84988984 | 257 |
| Rela | 0.84884607 | 258 |
| Notch4 | 0.80760005 | 259 |
| Zfp276 | 0.78572179 | 260 |
| Hand2 | 0.7797892 | 261 |
| Bcl11b | 0.74910719 | 262 |
| Hmgxb4 | 0.73023769 | 263 |
| Bach1 | 0.7146869 | 264 |
| Fosl2 | 0.57887039 | 265 |
| Rxra | 0.5710442 | 266 |
| Bnc1 | 0.55916289 | 267 |
| Foxk2 | 0.5589256 | 268 |
| Irx3 | 0.55042162 | 269 |
| Cbx2 | 0.52914894 | 270 |
| Zbed4 | 0.52234309 | 271 |
| Rfx3 | 0.51864309 | 272 |
| Pparg | 0.51479667 | 273 |
| Dzip1l | 0.50704212 | 274 |
| Zfp418 | 0.49587816 | 275 |
| Hoxd13 | 0.47662847 | 276 |
| Gata5 | 0.47245192 | 277 |
| Zfp7 | 0.43950898 | 278 |
| Nr1h4 | 0.41941942 | 279 |
| Zfp605 | 0.41279315 | 280 |
| Zfp608 | 0.41270199 | 281 |
| Ahr | 0.38492051 | 282 |
| Klf2 | 0.37376652 | 283 |
| Nab1 | 0.36988854 | 284 |
| Ikzf3 | 0.35142743 | 285 |

| | | |
|---|---|---|
| Tcf23 | 0.34884283 | 286 |
| Rbl2 | 0.34731474 | 287 |
| Foxq1 | 0.34540407 | 288 |
| Hnf4a | 0.31921027 | 289 |
| Nkx2-1 | 0.31342109 | 290 |
| Egr3 | 0.29206464 | 291 |
| Tsc22d3 | 0.24727456 | 292 |
| 1700049G17I | 0.24023131 | 293 |
| Lin28a | 0.18121649 | 294 |
| Atf3 | 0.16405174 | 295 |
| Tub | 0.15715925 | 296 |
| Smad9 | 0.14516268 | 297 |
| Tulp2 | 0.13181413 | 298 |
| Zfp91 | 0.1315994 | 299 |
| Tal1 | 0.12485842 | 300 |
| Vdr | 0.12374668 | 301 |
| Tulp1 | 0.1024862 | 302 |
| Zeb1 | 0.0884082 | 303 |
| Sall1 | 0.08762427 | 304 |
| Gata3 | 0.08641213 | 305 |
| Ebf3 | 0.08092203 | 306 |
| Klf4 | 0.06052392 | 307 |
| Stat5b | 0.05779414 | 308 |
| Isl1 | 0.0491897 | 309 |
| Sox17 | 0.02373659 | 310 |
| Etv1 | 0.0126639 | 311 |
| Zbtb7b | -0.0215984 | 312 |
| St18 | -0.0229784 | 313 |
| Hsf1 | -0.0263483 | 314 |
| Mafb | -0.0571176 | 315 |
| Satb1 | -0.0625558 | 316 |
| Pou2af1 | -0.0900106 | 317 |
| Nfkbie | -0.0909386 | 318 |
| Foxp3 | -0.1071219 | 319 |
| Bhlha15 | -0.1112567 | 320 |
| Mecom | -0.1297327 | 321 |
| Ets1 | -0.159157 | 322 |
| Nfkbia | -0.1597979 | 323 |
| Max | -0.1660682 | 324 |
| Nkx3-1 | -0.1767961 | 325 |
| Ebf1 | -0.1887836 | 326 |

| | | |
|---|---|---|
| Stat5a | -0.2034139 | 327 |
| Rxrg | -0.2179489 | 328 |
| Smad3 | -0.2198276 | 329 |
| Zfp704 | -0.2204332 | 330 |
| Hmgb2 | -0.2209861 | 331 |
| Sox13 | -0.2431762 | 332 |
| Jun | -0.260936 | 333 |
| Tcfl5 | -0.2794903 | 334 |
| Nfatc4 | -0.3153137 | 335 |
| Tsc22d4 | -0.3190012 | 336 |
| Erg | -0.329661 | 337 |
| Zfp521 | -0.3414332 | 338 |
| Sox9 | -0.3677513 | 339 |
| Tbr1 | -0.3678613 | 340 |
| Rfxank | -0.3686575 | 341 |
| Zfp821 | -0.3724735 | 342 |
| Myt1 | -0.3822673 | 343 |
| Zfp2 | -0.3850353 | 344 |
| Wt1 | -0.3868942 | 345 |
| Zfp422 | -0.3979833 | 346 |
| Nr4a3 | -0.4438542 | 347 |
| Elf2 | -0.4518824 | 348 |
| Nfyc | -0.4530833 | 349 |
| Zfp791 | -0.4763086 | 350 |
| Six3 | -0.4817484 | 351 |
| Tbx2 | -0.4829946 | 352 |
| Lyl1 | -0.5655823 | 353 |
| Mis18bp1 | -0.5712321 | 354 |
| Tfap2c | -0.5769615 | 355 |
| Satb2 | -0.5822072 | 356 |
| Ssrp1 | -0.5862129 | 357 |
| Zfp710 | -0.5924456 | 358 |
| Tfdp1 | -0.6023339 | 359 |
| Pax9 | -0.6163845 | 360 |
| Osr1 | -0.6199385 | 361 |
| Bnc2 | -0.6200854 | 362 |
| Cux1 | -0.6359466 | 363 |
| Rbpj | -0.6857736 | 364 |
| Stat4 | -0.686603 | 365 |
| Hoxd12 | -0.7145262 | 366 |
| Hoxc8 | -0.742046 | 367 |

| | | |
|---|---|---|
| Rest | -0.7582096 | 368 |
| Foxo4 | -0.775343 | 369 |
| Hmgb3 | -0.7867451 | 370 |
| Six2 | -0.8209715 | 371 |
| Tcf12 | -0.8258592 | 372 |
| Esr1 | -0.8291634 | 373 |
| Atoh1 | -0.8367633 | 374 |
| Hoxa11 | -0.8754801 | 375 |
| Nfkb2 | -0.8976546 | 376 |
| Zfpm1 | -0.8982374 | 377 |
| Tfcp2 | -0.90211 | 378 |
| Sox6 | -0.9029951 | 379 |
| Eomes | -0.9074085 | 380 |
| Dach1 | -0.9420441 | 381 |
| Relb | -0.9456402 | 382 |
| Egr2 | -0.9503156 | 383 |
| Emx2 | -0.9518883 | 384 |
| Irf4 | -0.9580841 | 385 |
| Tfec | -0.9672491 | 386 |
| Tlx3 | -0.967683 | 387 |
| Zscan21 | -0.9732268 | 388 |
| Prrx1 | -0.9733233 | 389 |
| Irf5 | -0.9746937 | 390 |
| Cebpd | -0.9866339 | 391 |
| Nr2f2 | -0.9941917 | 392 |
| Hoxc5 | -1.0482125 | 393 |
| Cdc5l | -1.0599327 | 394 |
| Zfp644 | -1.082421 | 395 |
| Zfp157 | -1.1096453 | 396 |
| Twist2 | -1.1266403 | 397 |
| Heyl | -1.1493948 | 398 |
| Ebf2 | -1.1527782 | 399 |
| Atoh8 | -1.1739552 | 400 |
| Maf | -1.1851684 | 401 |
| Pknox2 | -1.2187639 | 402 |
| Noc3l | -1.2350061 | 403 |
| Aebp1 | -1.2462929 | 404 |
| Hsf2 | -1.2694417 | 405 |
| Batf | -1.2824011 | 406 |
| Pou3f3 | -1.2844552 | 407 |
| Alx4 | -1.2899919 | 408 |

| | | |
|---|---|---|
| Hoxa10 | -1.2900994 | 409 |
| Nfkbib | -1.2978015 | 410 |
| Six5 | -1.3345912 | 411 |
| Sox7 | -1.3819496 | 412 |
| Bhlhe22 | -1.4158359 | 413 |
| Six4 | -1.4221391 | 414 |
| Nr5a2 | -1.4241443 | 415 |
| Ppara | -1.4256099 | 416 |
| Rara | -1.4317434 | 417 |
| Myc | -1.4320189 | 418 |
| Batf3 | -1.4846774 | 419 |
| Nr0b2 | -1.486447 | 420 |
| Gata6 | -1.4883266 | 421 |
| Hes1 | -1.5056107 | 422 |
| Fosl1 | -1.5161286 | 423 |
| Hoxa13 | -1.523802 | 424 |
| Tcf21 | -1.5268732 | 425 |
| Elk3 | -1.5343583 | 426 |
| Tbx3 | -1.547514 | 427 |
| Tbx5 | -1.6085132 | 428 |
| Ar | -1.6951931 | 429 |
| Fli1 | -1.7117283 | 430 |
| Spdef | -1.7139298 | 431 |
| Irf8 | -1.7142895 | 432 |
| Nfyb | -1.7418189 | 433 |
| Sox3 | -1.7476178 | 434 |
| Pax8 | -1.7504751 | 435 |
| Mitf | -1.7936321 | 436 |
| Cebpz | -1.8143077 | 437 |
| Sox18 | -1.8426565 | 438 |
| Smarce1 | -1.8690445 | 439 |
| Zdhhc13 | -1.9549728 | 440 |
| Hsf4 | -2.0319051 | 441 |
| Ubtf | -2.033617 | 442 |
| Hoxb13 | -2.0728522 | 443 |
| Arnt2 | -2.0747489 | 444 |
| Hes6 | -2.1329551 | 445 |
| Junb | -2.1521173 | 446 |
| Thrb | -2.1571535 | 447 |
| Scx | -2.1778945 | 448 |
| Zfp207 | -2.1782828 | 449 |

| | | |
|---|---|---|
| Zfp715 | -2.2208602 | 450 |
| Zfp593 | -2.2218819 | 451 |
| Tcf3 | -2.2326808 | 452 |
| Nfe2l2 | -2.2410378 | 453 |
| Mta2 | -2.2453723 | 454 |
| Drap1 | -2.2561489 | 455 |
| Zfp354c | -2.2670721 | 456 |
| Atf4 | -2.2819268 | 457 |
| Zfp191 | -2.294565 | 458 |
| Creb3 | -2.3886267 | 459 |
| Sp3 | -2.4054775 | 460 |
| Ybx1 | -2.4403507 | 461 |
| Mxd4 | -2.4497172 | 462 |
| Nr1h2 | -2.4662394 | 463 |
| Zfp768 | -2.4737496 | 464 |
| Alx3 | -2.5210917 | 465 |
| Mrpl28 | -2.5262647 | 466 |
| Creb3l4 | -2.5390209 | 467 |
| Six1 | -2.5406726 | 468 |
| Phtf2 | -2.5462092 | 469 |
| Esrra | -2.5995307 | 470 |
| Zbtb26 | -2.6095923 | 471 |
| Smarca5 | -2.6242582 | 472 |
| Mxd3 | -2.6350607 | 473 |
| Twist1 | -2.6504352 | 474 |
| Hnf4g | -2.7542802 | 475 |
| Csde1 | -2.8246284 | 476 |
| Phtf1 | -2.8249389 | 477 |
| Carhsp1 | -2.8583346 | 478 |
| Tfam | -2.8837571 | 479 |
| Zfp41 | -2.8961852 | 480 |
| Zfp51 | -2.9054217 | 481 |
| Zbtb41 | -2.9423397 | 482 |
| Nkx6-2 | -3.0035213 | 483 |
| Crem | -3.0163969 | 484 |
| Gatad1 | -3.1019425 | 485 |
| Nr2f6 | -3.1729179 | 486 |
| Creb1 | -3.1887851 | 487 |
| Cebpa | -3.2062193 | 488 |
| Terf2ip | -3.2472527 | 489 |
| Tead2 | -3.272056 | 490 |

| | | |
|---|---|---|
| Mycn | -3.2754482 | 491 |
| Bzw1 | -3.2760382 | 492 |
| Hmg20b | -3.2915481 | 493 |
| Zfp771 | -3.2963236 | 494 |
| Lhx2 | -3.3114984 | 495 |
| Jund | -3.4413714 | 496 |
| Myrf | -3.490221 | 497 |
| Cbfb | -3.5173919 | 498 |
| Onecut2 | -3.5433323 | 499 |
| Zfp110 | -3.5769438 | 500 |
| Hif1a | -3.5937663 | 501 |
| Sim2 | -3.593867 | 502 |
| Zfp219 | -3.6209283 | 503 |
| Zfp358 | -3.629482 | 504 |
| Zfp712 | -3.6428813 | 505 |
| Arid3a | -3.8403365 | 506 |
| Zfp260 | -3.8688176 | 507 |
| Cebpb | -3.8914202 | 508 |
| Irf3 | -3.9785609 | 509 |
| Zfp595 | -4.0317096 | 510 |
| Lmx1b | -4.071999 | 511 |
| Dr1 | -4.1413022 | 512 |
| Sox2 | -4.2976622 | 513 |
| Sox4 | -4.3907557 | 514 |
| Zfp518a | -4.6258228 | 515 |
| Ezh2 | -4.7069321 | 516 |

Table S4: EGFP Cell Type Gene Signatures

| Basal | Adeno_L2 | Adeno_L1 | Adeno_L3 | INT_Tff3 | INT_Inflam | INT_EMT | DNPC_Pou2f3 |
|---|---|---|---|---|---|---|---|
| ANXA2 | AW112010 | ADH1 | ACKR4 | AGR2 | ACOD1 | AY036118 | AGT |
| APOE | AY036118 | AKR1B3 | ADCY5 | ATOH1 | ANKRD33B | APOD | ALDH2 |
| BCAM | ANXA1 | B2M | ADGRF5 | ATP2C2 | ANTXR2 | APOE | ALOX5AP |
| CYR61 | ANXA3 | BMPR1B | ADORA1 | ATP4A | ARG2 | BCL2A1B | MCUB |
| COL17A1 | ARL14 | CD52 | AFAP1L1 | BC016579 | BCL2A1B | BGN | ATP2A3 |
| CP | CAR2 | CDO1 | ATP6V0A4 | BEST2 | CCL3 | C1QA | AVIL |
| DAPL1 | CBR2 | CEACAM10 | ATP6V0D2 | CAPN9 | CCL4 | C1QB | BCL2L14 |
| DCN | CEACAM1 | CES1D | ATP6V1B1 | CHODL | CCR1 | C1QC | BMX |
| DST | CLU | CHAC1 | ATP6V1C2 | LMAN1L | CCRL2 | CCL2 | BPGM |
| ECM1 | CTSE | CLU | ATP6V1G3 | CPM | CD14 | CCL5 | CADM2 |
| EXT1 | CXCL15 | CORO2A | BMPR1B | CREB3L4 | CD53 | CCL7 | CALML4 |
| FN1 | CXCL17 | CWH43 | CD200 | CRYM | CEBPB | CCL8 | CD24A |
| FOSL1 | CXCL5 | CYP4A12A | CDR2 | DPYSL3 | CLEC4D | CD14 | CDHR5 |
| FST | CYBA | DEFB1 | CLCNKB | ERN2 | CLEC4E | CD74 | CDKN1C |
| HBEGF | GM26917 | DEFB42 | CLDN10 | FCGBP | CLEC4N | COL1A1 | CLDN14 |
| HSPB1 | GSDMC2 | ELF5 | CLIC6 | FFAR4 | GM5483 | COL1A2 | CRIP1 |
| IFI202B | GSTA4 | FOLR1 | CPXM2 | GAP43 | CXCL2 | COL3A1 | CRISP3 |
| IGFBP5 | GSTO1 | FYB2 | CWH43 | GNE | CYP4F18 | CST3 | DCLK1 |
| IGFBP7 | KRT19 | GSTM1 | DAB2 | GPR155 | ENTPD1 | CTLA2A | ETHE1 |
| IL33 | KRT4 | HERPUD1 | DEFB1 | HEPACAM2 | FCER1G | CTSS | FOLR1 |
| INHBA | LCN2 | ITM2B | DKK3 | HSD11B2 | FTH1 | CXCL14 | FYB |
| ITGA2 | LGALS3 | LCN2 | DMRT2 | INSRR | FTL1 | DCN | GALK1 |
| ITGB1 | LGALS4 | NDUFA1 | EDN3 | KCNMA1 | FXYD5 | FCER1G | GNAT3 |
| ITGB4 | LRG1 | NEFH | FOXI1 | KLK1B11 | G0S2 | FXYD5 | GNG13 |
| KRT14 | LTF | NUCB2 | GPM6B | KLK1B9 | GADD45B | H2-AA | HCK |
| KRT15 | LY6A | ODAM | GSTM1 | TMEM159 | HCAR2 | H2-AB1 | HEPACAM2 |
| KRT16 | LY6D | A630095E13RIK | GSTM7 | LRRC31 | HDC | H2-EB1 | HPGDS |
| KRT17 | LYPD8 | PATE4 | KCNMA1 | MGAT5 | IL1A | IGFBP4 | LRMP |
| KRT5 | MMP7 | D730048I06RIK | LDHB | MICAL1 | IL1B | IGFBP6 | ITPR2 |
| KRT6A | NUPR1 | PDZK1IP1 | LEF1 | MUC13 | IL1R2 | IGFBP7 | KCTD12 |
| LAMB3 | PGLYRP1 | GGT1 | MCF2 | NR3C2 | IL1RN | IL1B | LTC4S |
| LGALS7 | PIGR | PHYH | MGLL | PDE7B | LILR4B | LAPTM5 | LY6G6F |
| LTBP1 | PLAT | PLAC8 | MUC20 | PIEZO2 | LMNB1 | LARS2 | MCTP1 |
| MFGE8 | PRDX6 | PRLR | PDE1A | RAMP1 | MXD1 | LGALS1 | MGST3 |
| MT2 | PSCA | RCN1 | PDLIM3 | SLC12A2 | NFKBIA | LTF | NRGN |
| NRG1 | REG1 | SVS2 | PLA2R1 | SLC12A8 | NINJ1 | LYZ2 | PGM2L1 |
| PERP | REG3B | SERP1 | SERPINB6B | SLC22A1 | NLRP3 | MFAP5 | PLAC8 |
| PHLDA1 | REG3G | SERPINE2 | SERPINB9 | SMIM14 | PIM1 | MGP | PTGS1 |
| RBMS3 | S100A11 | SH3BGRL3 | SLC26A3 | SPDEF | PLEK | C3 | PTPN18 |
| RBP1 | S100A6 | SHISA4 | SLCO1A5 | SVAL1 | PTGS2 | REG3B | PYGL |
| SFN | SBP | SLC31A2 | SPINK8 | SYTL2 | RETNLG | RGS1 | REEP5 |
| S100A10 | SLC12A2 | SPINK8 | SULT1A1 | TESC | S100A8 | GM42418 | RGS13 |
| SDC1 | SLC39A4 | SSR4 | TDRP | TFF3 | S100A9 | SELENOP | RGS2 |

| | | | | | | | |
|---|---|---|---|---|---|---|---|
| SERPINB5 | SPINK1 | SUCO | TMEM61 | TMEM56 | SAMSN1 | SERPING1 | SELENOM |
| SGK1 | SPINK5 | SVS4 | TMOD1 | TRIM31 | SLC7A11 | SFRP1 | SGCZ |
| TPM1 | SPRR2A3 | SVS5 | TTN | TTC8 | SLFN4 | SPARC | SH2D6 |
| TIMP3 | TSPAN8 | SVS6 | VSIG1 | ZC3H12D | SLPI | SRGN | SKAP2 |
| TM4SF1 | WFDC18 | TMED3 | VWC2 | BC030870 | SRGN | TYROBP | FAM19A1 |
| TNFRSF12A | WFDC2 | WFDC15B | | | TREM1 | VIM | TRPM5 |
| TPM2 | | | | | TYROBP | WFDC17 | TSPAN6 |

| NEPC_Ascl1 | NEPC_AR | NEPC_Neurod1 | NEPC_Nefl |
|---|---|---|---|
| ALDH1A2 | DCC | A2ML1 | ACVR1 |
| ARL6IP1 | AGBL4 | AFF3 | ADGRB3 |
| ASCL1 | AUTS2 | ATP8A1 | ANKS1B |
| BEX1 | CACNA1C | AUTS2 | ASCL1 |
| BEX2 | CACNA2D1 | C130073E24RIK | CACNA2D3 |
| CDK1 | CALCA | CACNA1C | CARMIL1 |
| CENPA | CALN1 | CAMK4 | CDH13 |
| CENPF | CCSER1 | CDH13 | CDH20 |
| CHGA | CDH13 | CDH20 | CDK6 |
| CKB | CIB3 | CEP112 | CHD7 |
| DDC | CNTN4 | COL8A1 | CNTN4 |
| ESPN | CNTNAP5A | DENND4A | CPLX2 |
| FABP5 | CTNNA2 | ETV6 | DACH2 |
| H1F0 | EFNA5 | EXOC4 | EFNA5 |
| H1FX | FAT3 | FARP1 | FGF14 |
| H2AFZ | FBXO38 | FAT3 | FRMD5 |
| HES6 | FGF14 | FMNL2 | FXYD6 |
| HMGB2 | FRMD5 | HCN1 | GRID2 |
| HMGN2 | GAD1 | KCND2 | HCN1 |
| IFITM1 | GALNT13 | MAP2 | HES6 |
| IGFBP2 | GALNT17 | MDGA2 | HS6ST3 |
| INSM1 | GRID2 | MEIS2 | ID4 |
| KRT7 | HCN1 | MPZL1 | KCNB2 |
| LMO4 | HS6ST3 | FAM155A | KCNIP4 |
| LOCKD | IMMP2L | NEUROD1 | GRIP1 |
| MAP1B | KCNB2 | NKAIN2 | LMO4 |
| MKI67 | KCNIP4 | NLGN1 | MAGI2 |
| GM1673 | MGAT4C | NRXN1 | FAM155A |
| PCLAF | NEGR1 | PDE11A | NECAB1 |
| PCSK1 | NNAT | PDE4D | NEFL |
| PKIB | NRG3 | PIP5K1B | NEFM |
| NOV | NRXN1 | PITPNC1 | NRG3 |
| PRC1 | OXR1 | PTPRN2 | NRXN1 |
| RESP18 | PCDH17 | PTPRT | NXPH1 |
| RTN1 | PHACTR1 | TOX | PAX9 |
| RUNDC3A | RAPGEF4 | RIMS2 | PDE4D |
| SMC4 | TOX | SHROOM3 | PEX5L |
| SNCA | RUNX1T1 | SKAP1 | PIP5K1B |
| SNHG11 | SDK1 | SOX5 | PKD1L3 |
| STMN1 | SLC8A1 | SPOCK1 | TOX |
| STMN3 | SNAP25 | SYT1 | RTN1 |
| TMEM158 | SNHG11 | TENM1 | RUNX1T1 |
| TOP2A | TCERG1L | TENM2 | SGIP1 |

| | | | |
|---|---|---|---|
| TPH1 | TENM4 | TENM3 | SPP1 |
| TUBA1A | TMEM117 | TMEM158 | SLC25A21 |
| TUBA1B | TMEM163 | TUBB2B | SYT1 |
| TUBB5 | UBE2E2 | UPP1 | TCERG1L |
| UBE2C | ZBTB20 | VWF | TENM1 |
| UBE2S | ZFP704 | XKR4 | THSD4 |
| UCHL1 | | | TMEM163 |

Table S5: CytoTrace Correlated Gene Expression

| Gene | Correlation | Correlation with Plasticity |
|---|---|---|
| Tubb5 | 0.506289254 | Positive |
| H2afz | 0.499034613 | Positive |
| Ptma | 0.486818916 | Positive |
| Hmgn2 | 0.477639297 | Positive |
| Pmf1 | 0.471828581 | Positive |
| Tmem14c | 0.465451358 | Positive |
| Pclaf | 0.457951627 | Positive |
| Calm2 | 0.450601387 | Positive |
| Ran | 0.449593285 | Positive |
| Hmgb2 | 0.448084553 | Positive |
| Hmgb1 | 0.43991841 | Positive |
| Stmn1 | 0.435996336 | Positive |
| Tmpo | 0.435123492 | Positive |
| Ppp1ca | 0.43451523 | Positive |
| H2afv | 0.432763925 | Positive |
| Acat1 | 0.431266097 | Positive |
| Tyms | 0.425925684 | Positive |
| H2afy | 0.423694473 | Positive |
| Selenoh | 0.423540335 | Positive |
| Dut | 0.423165809 | Positive |
| Snrpd1 | 0.422153466 | Positive |
| Fabp5 | 0.420116104 | Positive |
| Ppia | 0.418691898 | Positive |
| Snrpa1 | 0.416079091 | Positive |
| Nudc | 0.415701176 | Positive |
| Tuba1b | 0.412652968 | Positive |
| Hnrnpa1 | 0.412110676 | Positive |
| Vdac3 | 0.411334711 | Positive |
| Hnrnpab | 0.410477227 | Positive |
| Hmgn1 | 0.410349816 | Positive |
| Sumo2 | 0.406551523 | Positive |
| Nsmce4a | 0.405893313 | Positive |
| Ilf2 | 0.403853189 | Positive |
| Anp32b | 0.397888333 | Positive |
| Srsf3 | 0.396666127 | Positive |
| Gmnn | 0.396269276 | Positive |
| Psip1 | 0.396145298 | Positive |
| Rbm3 | 0.395365744 | Positive |
| Ranbp1 | 0.391973015 | Positive |
| Lmnb1 | 0.391081161 | Positive |
| Dnajc9 | 0.390701067 | Positive |
| Cks1b | 0.38956122 | Positive |
| Lsm3 | 0.389056067 | Positive |

| | | |
|---|---|---|
| Csrp2 | 0.388447684 | Positive |
| Ccdc34 | 0.388100165 | Positive |
| Rrm1 | 0.387033953 | Positive |
| Smc2 | 0.386652413 | Positive |
| Cdk1 | 0.383940898 | Positive |
| Npm1 | 0.38294494 | Positive |
| Anapc5 | 0.382494717 | Positive |
| Lsm5 | 0.381390181 | Positive |
| Rfc5 | 0.381026022 | Positive |
| Rtraf | 0.380990389 | Positive |
| Rfc4 | 0.379992699 | Positive |
| Cenpm | 0.379939373 | Positive |
| Anp32e | 0.379510682 | Positive |
| Sae1 | 0.379390034 | Positive |
| Jpt1 | 0.379256871 | Positive |
| Lsm2 | 0.378195798 | Positive |
| Fkbp3 | 0.377636498 | Positive |
| Pbk | 0.375289382 | Positive |
| Rangap1 | 0.373944882 | Positive |
| Ube2t | 0.373745805 | Positive |
| Smc4 | 0.373490807 | Positive |
| Thoc7 | 0.371913711 | Positive |
| Cks2 | 0.37184791 | Positive |
| Exosc8 | 0.371577304 | Positive |
| Ube2s | 0.37121018 | Positive |
| Dtymk | 0.37063462 | Positive |
| Ezh2 | 0.370319342 | Positive |
| Alyref | 0.370022234 | Positive |
| Ncl | 0.369422129 | Positive |
| Dek | 0.368887703 | Positive |
| Ccna2 | 0.36812932 | Positive |
| Hint1 | 0.367146626 | Positive |
| Hsp90aa1 | 0.3670038 | Positive |
| Cenpk | 0.366329615 | Positive |
| Dctpp1 | 0.365787349 | Positive |
| Rdx | 0.365766758 | Positive |
| Spc24 | 0.365736327 | Positive |
| Impdh2 | 0.364365978 | Positive |
| H3f3a | 0.363659774 | Positive |
| Top2a | 0.363482245 | Positive |
| Racgap1 | 0.362164281 | Positive |
| Usp1 | 0.362062664 | Positive |
| Slc25a4 | 0.36054502 | Positive |
| Dazap1 | 0.360439167 | Positive |

| | | |
|---|---|---|
| Ckap2l | 0.359653922 | Positive |
| Ift27 | 0.357548539 | Positive |
| Birc5 | 0.356730667 | Positive |
| Hmgb3 | 0.354900189 | Positive |
| Cdca8 | 0.354677525 | Positive |
| Gm1673 | 0.354394687 | Positive |
| Spc25 | 0.354188002 | Positive |
| Snrpe | 0.353973105 | Positive |
| Fam216a | 0.353831963 | Positive |
| Atp5f1 | 0.353797995 | Positive |
| Nsg1 | 0.353699449 | Positive |
| Elof1 | 0.352865791 | Positive |
| Tcp1 | 0.352676786 | Positive |
| Tsen34 | 0.352670192 | Positive |
| Sub1 | 0.352553572 | Positive |
| Nucks1 | 0.352389073 | Positive |
| Rpsa | 0.352356083 | Positive |
| Lmo4 | 0.352003443 | Positive |
| Atad2 | 0.35108555 | Positive |
| Cdkn2d | 0.350572784 | Positive |
| Eif4a1 | 0.350148127 | Positive |
| Prmt1 | 0.35005277 | Positive |
| Nt5dc2 | 0.349683517 | Positive |
| Asf1b | 0.349346569 | Positive |
| Hnrnpa2b1 | 0.347682996 | Positive |
| Oxct1 | 0.347663951 | Positive |
| Cox4i1 | 0.347532775 | Positive |
| Banf1 | 0.346956206 | Positive |
| Tubg1 | 0.346640515 | Positive |
| Ubald2 | 0.346238744 | Positive |
| Tubb4b | 0.346235523 | Positive |
| Nefl | 0.346012418 | Positive |
| Ybx1 | 0.345754911 | Positive |
| Cox5a | 0.345749249 | Positive |
| Timm50 | 0.344960049 | Positive |
| H3f3b | 0.344371133 | Positive |
| Batf3 | 0.344267095 | Positive |
| Smc3 | 0.344035983 | Positive |
| Nfib | 0.34227536 | Positive |
| Hes6 | 0.34216276 | Positive |
| Atp5o.1 | 0.341947389 | Positive |
| Lsm8 | 0.341728687 | Positive |
| Ndufa4 | 0.341591497 | Positive |
| Uhrf1 | 0.341541938 | Positive |

| | | |
|---|---|---|
| Nusap1 | 0.341458586 | Positive |
| Nap1l1 | 0.341281085 | Positive |
| Pdcd5 | 0.34124304 | Positive |
| Dnph1 | 0.341204095 | Positive |
| Cetn3 | 0.340631741 | Positive |
| Rbm8a | 0.340615543 | Positive |
| Tk1 | 0.339935635 | Positive |
| Nasp | 0.339790267 | Positive |
| Hnrnpk | 0.339630861 | Positive |
| Nono | 0.339420326 | Positive |
| Raly | 0.338994406 | Positive |
| Serbp1 | 0.338152584 | Positive |
| Nudt21 | 0.338086596 | Positive |
| Rrm2 | 0.337328742 | Positive |
| Aif1l | 0.336784201 | Positive |
| H2afx | 0.336179436 | Positive |
| Mrpl28 | 0.335620206 | Positive |
| Hirip3 | 0.335022122 | Positive |
| Eif4g2 | 0.334948159 | Positive |
| Incenp | 0.334906051 | Positive |
| Mns1 | 0.334788585 | Positive |
| Fen1 | 0.33433755 | Positive |
| Smc6 | 0.334296479 | Positive |
| Tlx3 | 0.333930049 | Positive |
| 1110004F10Rik | 0.332140771 | Positive |
| Ppp1cc | 0.331137178 | Positive |
| Hmmr | 0.330660776 | Positive |
| Prc1 | 0.330557802 | Positive |
| Pdap1 | 0.330179795 | Positive |
| Fbxo5 | 0.33013278 | Positive |
| Kif22 | 0.329884292 | Positive |
| Prelid3b | 0.329812146 | Positive |
| Supt16 | 0.329607227 | Positive |
| Kif23 | 0.329124016 | Positive |
| Rad21 | 0.329009473 | Positive |
| Cbx5 | 0.328145613 | Positive |
| Mbd3 | 0.326879186 | Positive |
| Prim1 | 0.326446067 | Positive |
| Ncaph | 0.326428762 | Positive |
| Rad51 | 0.325910076 | Positive |
| Gnb2 | 0.325562687 | Positive |
| Mki67 | 0.32540432 | Positive |
| Fxyd6 | 0.325112423 | Positive |
| Nol7 | 0.324916678 | Positive |

| | | |
|---|---|---|
| Slbp | 0.324793604 | Positive |
| Nt5c3b | 0.324573643 | Positive |
| Pax9 | 0.3242453 | Positive |
| Sept7 | 0.323757768 | Positive |
| Cyth2 | 0.323498685 | Positive |
| Lsm6 | 0.323084922 | Positive |
| Sgo2a | 0.32292573 | Positive |
| Gemin6 | 0.322262033 | Positive |
| Eif5a | 0.322223506 | Positive |
| Hdgf | 0.322045727 | Positive |
| Ssna1 | 0.321456817 | Positive |
| D030056L22Rik | 0.321427979 | Positive |
| Ndc80 | 0.321361846 | Positive |
| Lmo1 | 0.321225384 | Positive |
| Hnrnpf | 0.320490774 | Positive |
| Nefm | 0.320153543 | Positive |
| Srsf2 | 0.319472667 | Positive |
| Hnrnpa3 | 0.319406809 | Positive |
| Smarcc1 | 0.319132643 | Positive |
| Dbf4 | 0.319043834 | Positive |
| Cdc45 | 0.31891646 | Positive |
| Pcm1 | 0.318574318 | Positive |
| Actl6a | 0.318110998 | Positive |
| Maz | 0.31811082 | Positive |
| Krt7 | 0.317997667 | Positive |
| Gm11837 | 0.317975162 | Positive |
| Ppm1g | 0.317873136 | Positive |
| Cnbp | 0.317637129 | Positive |
| Phf5a | 0.317627771 | Positive |
| Ndufa5 | 0.317289423 | Positive |
| Melk | 0.316958107 | Positive |
| Hjurp | 0.316756633 | Positive |
| Sf3b1 | 0.31661478 | Positive |
| Pcbp4 | 0.316239809 | Positive |
| Cdca3 | 0.31622073 | Positive |
| Snrpc | 0.316084008 | Positive |
| Tacc3 | 0.315994985 | Positive |
| Plk1 | 0.315843808 | Positive |
| Mrpl18 | 0.315802582 | Positive |
| Bex2 | 0.315452425 | Positive |
| Mapk13 | 0.314966311 | Positive |
| Eif4h | 0.314371078 | Positive |
| Nsmce1 | 0.314229305 | Positive |
| Ssx2ip | 0.313970617 | Positive |

| | | |
|---|---|---|
| Nmral1 | 0.313819463 | Positive |
| Ddx39b | 0.313745569 | Positive |
| Aurkb | 0.313462056 | Positive |
| Ncapg2 | 0.312997581 | Positive |
| Clspn | 0.312561542 | Positive |
| Aurka | 0.312153363 | Positive |
| Cenpe | 0.311826018 | Positive |
| Ccnb1 | 0.311768074 | Positive |
| Ssrp1 | 0.311217721 | Positive |
| Cnih2 | 0.310792679 | Positive |
| Ywhae | 0.310393291 | Positive |
| Idh3a | 0.31026936 | Positive |
| Etfa | 0.310106861 | Positive |
| Ncapd2 | 0.310086167 | Positive |
| Ssb | 0.310045746 | Positive |
| Nfix | 0.31003068 | Positive |
| Plk4 | 0.309817297 | Positive |
| Sgo1 | 0.3097719 | Positive |
| Hnrnpu | 0.309169871 | Positive |
| Psmc3 | 0.309074839 | Positive |
| Fkbp4 | 0.309070345 | Positive |
| Tcf19 | 0.309069061 | Positive |
| Shcbp1 | 0.30862893 | Positive |
| Cirbp | 0.308430807 | Positive |
| Rfc2 | 0.308354981 | Positive |
| Gnas | 0.308095092 | Positive |
| Id4 | 0.307759082 | Positive |
| Rad51ap1 | 0.30774186 | Positive |
| Gltp | 0.307686847 | Positive |
| Commd1 | 0.307519622 | Positive |
| Snu13 | 0.307161209 | Positive |
| Dck | 0.307102484 | Positive |
| Rfc3 | 0.306904767 | Positive |
| Ncapd3 | 0.306861054 | Positive |
| Atp5b | 0.306603708 | Positive |
| Pbrm1 | 0.306593289 | Positive |
| Pafah1b3 | 0.306261537 | Positive |
| Snrpb | 0.304840493 | Positive |
| Psmc6 | 0.304590735 | Positive |
| Cenpa | 0.304527645 | Positive |
| Matr3 | 0.304455284 | Positive |
| Nap1l4 | 0.303926903 | Positive |
| Crlf1 | 0.303812862 | Positive |
| Mcm7 | 0.303562144 | Positive |

| | | |
|---|---|---|
| Il17b | 0.303022223 | Positive |
| Cdc20 | 0.302820884 | Positive |
| Cdk6 | 0.302764613 | Positive |
| Srrm1 | 0.302633936 | Positive |
| Cdk5rap2 | 0.302630465 | Positive |
| Psmc4 | 0.302440656 | Positive |
| Cmc2 | 0.302261954 | Positive |
| Rnf187 | 0.30222734 | Positive |
| Hdac2 | 0.301997815 | Positive |
| Cenpw | 0.301904306 | Positive |
| Rnf126 | 0.301590768 | Positive |
| Mtch1 | 0.301454134 | Positive |
| Pcbp2 | 0.301393082 | Positive |
| Nme1 | 0.301391439 | Positive |
| Uba2 | 0.301244765 | Positive |
| Tpx2 | 0.301230684 | Positive |
| Ywhah | 0.301156204 | Positive |
| Mxd3 | 0.300887778 | Positive |
| Timm8b | 0.300783668 | Positive |
| Mis18bp1 | 0.300740319 | Positive |
| Prdx5 | 0.300394273 | Positive |
| Atp5a1 | 0.300366558 | Positive |
| Mlf1 | 0.300093136 | Positive |
| Sult2b1 | 0.299967781 | Positive |
| Adh5 | 0.299808623 | Positive |
| Elob | 0.29972031 | Positive |
| Aebp1 | 0.299665915 | Positive |
| Necab1 | 0.299649002 | Positive |
| Kif20a | 0.299544333 | Positive |
| Bex1 | 0.299464489 | Positive |
| Cltb | 0.299248766 | Positive |
| Nuf2 | 0.299231443 | Positive |
| Rrp1 | 0.299229882 | Positive |
| Fam92a | 0.299183222 | Positive |
| Tpr | 0.299095174 | Positive |
| Rab2a | 0.298801966 | Positive |
| Smarca5 | 0.29879481 | Positive |
| Ppard | -0.101748639 | Negative |
| Parp8 | -0.101893625 | Negative |
| Arhgef3 | -0.102022597 | Negative |
| Zfp408 | -0.102719424 | Negative |
| Xbp1 | -0.102912893 | Negative |
| Dock9 | -0.10291551 | Negative |
| Blnk | -0.102923999 | Negative |

| | | |
|---|---|---|
| Fbxl17 | -0.10312957 | Negative |
| Cpq | -0.103548575 | Negative |
| Pla2g16 | -0.103569678 | Negative |
| Nfasc | -0.103689651 | Negative |
| Tgtp2 | -0.103780944 | Negative |
| Apof | -0.103788174 | Negative |
| Tmem173 | -0.103817473 | Negative |
| Liph | -0.10381877 | Negative |
| Unc93b1 | -0.10383639 | Negative |
| Man2a1 | -0.10418334 | Negative |
| Itpr2 | -0.104223837 | Negative |
| Pawr | -0.10441696 | Negative |
| Maml3 | -0.104531881 | Negative |
| Gpr146 | -0.104558899 | Negative |
| Efhb | -0.104653774 | Negative |
| Kcnq1 | -0.104679379 | Negative |
| Tmem176a | -0.10469737 | Negative |
| Hivep3 | -0.105014014 | Negative |
| Mkl2 | -0.105128964 | Negative |
| Ano6 | -0.105142833 | Negative |
| Dnah9 | -0.105203458 | Negative |
| Afap1l2 | -0.105218047 | Negative |
| Smg1 | -0.105261446 | Negative |
| Cables1 | -0.105401652 | Negative |
| Ssfa2 | -0.105458081 | Negative |
| Kcnma1 | -0.105582358 | Negative |
| Exoc6b | -0.10558266 | Negative |
| Pygl | -0.105623384 | Negative |
| Spock1 | -0.106086892 | Negative |
| Fam46a | -0.106341942 | Negative |
| Aff2 | -0.106686236 | Negative |
| Tenm3 | -0.106874243 | Negative |
| Gm44148 | -0.106885111 | Negative |
| Epsti1 | -0.107086242 | Negative |
| Kcnj6 | -0.107133222 | Negative |
| Nalcn | -0.107316659 | Negative |
| Adcy2 | -0.107361467 | Negative |
| Stk32c | -0.107418375 | Negative |
| Mast4 | -0.10796837 | Negative |
| Capn13 | -0.107969702 | Negative |
| Naip2 | -0.108002136 | Negative |
| Thrb | -0.108086701 | Negative |
| Plscr1 | -0.108377771 | Negative |
| Trpv3 | -0.108424241 | Negative |

| | | |
|---|---|---|
| Galnt7 | -0.108529233 | Negative |
| Fbxo38 | -0.108630285 | Negative |
| Tubb2b | -0.108747526 | Negative |
| Rbm47 | -0.108901184 | Negative |
| Ttc39aos1 | -0.109189449 | Negative |
| Gm38505 | -0.109246611 | Negative |
| Stab2 | -0.109577934 | Negative |
| Gm15283 | -0.110466776 | Negative |
| H2-T23 | -0.110714029 | Negative |
| Crebrf | -0.11096078 | Negative |
| 4930539E08Rik | -0.110994504 | Negative |
| Kcnq4 | -0.111008132 | Negative |
| Cd200r2 | -0.111510775 | Negative |
| Gpatch2l | -0.111527725 | Negative |
| Cacnb4 | -0.111686267 | Negative |
| Baiap2l1 | -0.111786474 | Negative |
| Rac3 | -0.111886245 | Negative |
| Rbms3 | -0.111959095 | Negative |
| Myo3b | -0.111980112 | Negative |
| Lpcat2 | -0.112131521 | Negative |
| Oxr1 | -0.112326673 | Negative |
| Dennd4a | -0.112566168 | Negative |
| Lrch1 | -0.112591252 | Negative |
| Gbe1 | -0.112921644 | Negative |
| Hoxb7 | -0.113432959 | Negative |
| Gm37240 | -0.113651829 | Negative |
| Cacna1e | -0.113712176 | Negative |
| Pdgfc | -0.11378753 | Negative |
| Arhgap29 | -0.114165049 | Negative |
| Ets2 | -0.114209653 | Negative |
| Unc79 | -0.114232444 | Negative |
| Gm26848 | -0.115247228 | Negative |
| B3gat2 | -0.115264486 | Negative |
| Kank1 | -0.115368115 | Negative |
| Ppfibp2 | -0.115469835 | Negative |
| Gm26835 | -0.115541463 | Negative |
| Smoc2 | -0.115800984 | Negative |
| Mfap3l | -0.115882728 | Negative |
| Gm15810 | -0.115936203 | Negative |
| Ankrd28 | -0.115937592 | Negative |
| Clstn2 | -0.116314338 | Negative |
| Camk2n1 | -0.116774093 | Negative |
| Cdon | -0.116822205 | Negative |
| Ern1 | -0.116955563 | Negative |

| | | |
|---|---|---|
| Socs3 | -0.116991131 | Negative |
| Pacrg | -0.117073764 | Negative |
| Zeb1 | -0.11708866 | Negative |
| Gcnt2 | -0.117674848 | Negative |
| Vmp1 | -0.1177502 | Negative |
| Nek10 | -0.117761765 | Negative |
| Cntn3 | -0.117940121 | Negative |
| Lypd6b | -0.118002705 | Negative |
| Kcnh5 | -0.118221481 | Negative |
| Plcl2 | -0.118253095 | Negative |
| Setbp1 | -0.118860256 | Negative |
| Neurod1 | -0.119256536 | Negative |
| Ppm1e | -0.119392431 | Negative |
| Ell2 | -0.11954318 | Negative |
| Barx2 | -0.119661396 | Negative |
| Zfhx4 | -0.1201226 | Negative |
| 1810034E14Rik | -0.120129457 | Negative |
| Slc37a1 | -0.120553788 | Negative |
| Ccdc162 | -0.120896075 | Negative |
| Camk2a | -0.121348016 | Negative |
| Ccnd3 | -0.121553452 | Negative |
| Arid5b | -0.121835894 | Negative |
| Arhgap22 | -0.121874375 | Negative |
| Reps2 | -0.122206394 | Negative |
| Xkr4 | -0.122214778 | Negative |
| Egln3 | -0.12226257 | Negative |
| Nlrc5 | -0.122547417 | Negative |
| Pde11a | -0.122818136 | Negative |
| Scg2 | -0.123003627 | Negative |
| Mgat4c | -0.123064462 | Negative |
| Cnn3 | -0.123226045 | Negative |
| Naip5 | -0.123405513 | Negative |
| Auts2 | -0.12351696 | Negative |
| Gch1 | -0.12378368 | Negative |
| Kalrn | -0.124061429 | Negative |
| Crim1 | -0.124816772 | Negative |
| Aff3 | -0.124877981 | Negative |
| Gm572 | -0.125123763 | Negative |
| Ahnak | -0.125343315 | Negative |
| Ptprn2 | -0.126172623 | Negative |
| Ctxn3 | -0.126458273 | Negative |
| Sidt1 | -0.126663821 | Negative |
| Ypel2 | -0.126865428 | Negative |
| Atxn1 | -0.126978539 | Negative |

| | | |
|---|---|---|
| Il13ra1 | -0.127030554 | Negative |
| Pde1c | -0.127532883 | Negative |
| Btbd11 | -0.127624096 | Negative |
| Sec24d | -0.127941698 | Negative |
| Smoc1 | -0.128097466 | Negative |
| St14 | -0.128150296 | Negative |
| Irf1 | -0.128432594 | Negative |
| Pappa | -0.128697431 | Negative |
| Tmtc1 | -0.128733223 | Negative |
| Pgpep1l | -0.129345047 | Negative |
| Bdnf | -0.129437086 | Negative |
| Nnat | -0.129699976 | Negative |
| Usp6nl | -0.13019783 | Negative |
| Kcnn3 | -0.130580746 | Negative |
| Slc24a3 | -0.131439009 | Negative |
| Map6 | -0.131553521 | Negative |
| Ubxn10 | -0.131770958 | Negative |
| Stat3 | -0.131926241 | Negative |
| Gm15478 | -0.13200959 | Negative |
| Rrbp1 | -0.132453297 | Negative |
| Fgf12 | -0.132730833 | Negative |
| C130073E24Rik | -0.132778838 | Negative |
| Rgs22 | -0.133030207 | Negative |
| Myo1b | -0.133319916 | Negative |
| Tanc1 | -0.133483512 | Negative |
| Rab27a | -0.134233223 | Negative |
| Bace2 | -0.134659227 | Negative |
| Atp10a | -0.134765151 | Negative |
| Tshz3 | -0.13476679 | Negative |
| Jak2 | -0.13495131 | Negative |
| Ube2e2 | -0.135077074 | Negative |
| Eps8 | -0.135727877 | Negative |
| Utp20 | -0.135891593 | Negative |
| Cacnb2 | -0.136207617 | Negative |
| Clmn | -0.136334365 | Negative |
| Psd3 | -0.136698579 | Negative |
| Slc22a23 | -0.137163002 | Negative |
| Ppm1l | -0.137286742 | Negative |
| Corin | -0.137374267 | Negative |
| Rab3c | -0.137750206 | Negative |
| Cntnap5a | -0.137873023 | Negative |
| Lpp | -0.13795102 | Negative |
| R3hdml | -0.13826284 | Negative |
| Smad1 | -0.138304972 | Negative |

| | | |
|---|---|---|
| Ank | -0.138478191 | Negative |
| Naip6 | -0.13886294 | Negative |
| Plekhg3 | -0.139026492 | Negative |
| Dcc | -0.139218651 | Negative |
| Ank3 | -0.139265727 | Negative |
| Rab3b | -0.14010419 | Negative |
| Wipf1 | -0.140282728 | Negative |
| 5330417C22Rik | -0.141039114 | Negative |
| Pdcd4 | -0.141383195 | Negative |
| Dock1 | -0.141477568 | Negative |
| Rnf130 | -0.141724319 | Negative |
| 2010111I01Rik | -0.142312119 | Negative |
| Fam189a2 | -0.142379527 | Negative |
| Cdc14b | -0.142695972 | Negative |
| Mdga2 | -0.143112429 | Negative |
| B2m | -0.143715794 | Negative |
| Phldb2 | -0.143751779 | Negative |
| Cdh1 | -0.144781292 | Negative |
| Nr3c2 | -0.145140747 | Negative |
| Lamc2 | -0.145800755 | Negative |
| Lmtk2 | -0.146005198 | Negative |
| Arntl2 | -0.146210143 | Negative |
| Myo1d | -0.146408767 | Negative |
| Pdlim5 | -0.146869554 | Negative |
| Sgce | -0.146989751 | Negative |
| Pam | -0.147574283 | Negative |
| Rin2 | -0.148272501 | Negative |
| Asap2 | -0.148392723 | Negative |
| Ankrd34c | -0.149058478 | Negative |
| Slc12a8 | -0.149177921 | Negative |
| Nhs | -0.149632148 | Negative |
| App | -0.149668054 | Negative |
| Sh3d19 | -0.150345755 | Negative |
| Tmtc2 | -0.151083377 | Negative |
| Heg1 | -0.152408067 | Negative |
| Rabgap1l | -0.152882348 | Negative |
| Gsdma | -0.154566081 | Negative |
| Camk2d | -0.154809907 | Negative |
| Skap1 | -0.154837518 | Negative |
| 0610040J01Rik | -0.155219844 | Negative |
| Zfp618 | -0.155279121 | Negative |
| Ldlrad4 | -0.1553694 | Negative |
| Nr6a1 | -0.155790171 | Negative |
| Fat3 | -0.156026129 | Negative |

| | | |
|---|---|---|
| Ptprt | -0.156027914 | Negative |
| Rassf3 | -0.156219079 | Negative |
| Zfhx3 | -0.156757856 | Negative |
| Rgs7 | -0.157215928 | Negative |
| Mbnl2 | -0.157646735 | Negative |
| Rps6ka3 | -0.157870088 | Negative |
| Nlgn1 | -0.158898462 | Negative |
| Epb41l4a | -0.160092484 | Negative |
| Sema5a | -0.16095076 | Negative |
| Sgms2 | -0.161963369 | Negative |
| Resp18 | -0.162549672 | Negative |
| Rora | -0.162701599 | Negative |
| Fndc3b | -0.163534525 | Negative |
| Mybpc1 | -0.163838642 | Negative |
| Qsox1 | -0.165489498 | Negative |
| Hivep2 | -0.165735908 | Negative |
| Sh3rf1 | -0.166213467 | Negative |
| Cdk14 | -0.166443553 | Negative |
| Ptk2b | -0.166573079 | Negative |
| Ntn4 | -0.166693197 | Negative |
| Gm15915 | -0.167280187 | Negative |
| Fndc1 | -0.16998904 | Negative |
| Sgms1 | -0.170110729 | Negative |
| Bcl3 | -0.170196094 | Negative |
| Atp8a1 | -0.170734779 | Negative |
| St8sia5 | -0.170975683 | Negative |
| Zdhhc14 | -0.171320521 | Negative |
| Car8 | -0.172081658 | Negative |
| Lncpint | -0.172668169 | Negative |
| Gm26883 | -0.174171598 | Negative |
| Tenm2 | -0.175119318 | Negative |
| Cacna2d1 | -0.175289216 | Negative |
| Trpc4 | -0.177173711 | Negative |
| Adamts17 | -0.177659725 | Negative |
| Pdzrn3 | -0.177881213 | Negative |
| 9030622O22Rik | -0.17916401 | Negative |
| Cap2 | -0.1793041 | Negative |
| Lifr | -0.179784171 | Negative |
| Sorcs2 | -0.181828135 | Negative |
| Col8a1 | -0.182963895 | Negative |
| Tox2 | -0.183227957 | Negative |
| Hoxb8 | -0.184439221 | Negative |
| Slc35f3 | -0.184596454 | Negative |
| Srgap1 | -0.185241418 | Negative |

| | | |
|---|---|---|
| Ptprj | -0.186029552 | Negative |
| Ror2 | -0.18709605 | Negative |
| Igfbp5 | -0.18798707 | Negative |
| Tshz2 | -0.189973926 | Negative |
| Cacna1c | -0.191832841 | Negative |
| Shroom3 | -0.192180598 | Negative |
| Cers6 | -0.193304854 | Negative |
| Nebl | -0.193668055 | Negative |
| Ralgapa2 | -0.199329607 | Negative |
| Mpzl1 | -0.199481323 | Negative |
| Entpd3 | -0.203150987 | Negative |
| Itga9 | -0.204999679 | Negative |
| Upp1 | -0.205286481 | Negative |
| 1700113H08Rik | -0.206465852 | Negative |
| Nxpe4 | -0.206569836 | Negative |
| Meis2 | -0.208457898 | Negative |
| Pitpnc1 | -0.209338726 | Negative |
| 2610035D17Rik | -0.209451785 | Negative |
| Arhgap26 | -0.214470076 | Negative |
| Nptx1 | -0.217682902 | Negative |
| Tmem45b | -0.220540023 | Negative |
| Glis3 | -0.221365071 | Negative |
| Fmnl2 | -0.223386245 | Negative |
| Phactr1 | -0.225117229 | Negative |
| Tmem117 | -0.230435883 | Negative |
| Mgat5 | -0.23107272 | Negative |
| Camk4 | -0.233584162 | Negative |
| Etv6 | -0.234894746 | Negative |
| Mctp2 | -0.260228553 | Negative |

### Table S6: Differential H3K27ac DKO_EP vs DKO Top500

| Symbol | Fold | p.value | FDR | annotation |
|---|---|---|---|---|
| Hoxd3 | -6.194419697 | 6.32E-13 | 2.08E-08 | Promoter (<=1kb) |
| Fhod1 | 8.192230744 | 1.67E-09 | 2.35E-05 | Promoter (<=1kb) |
| Hoxd8 | -4.708149591 | 2.14E-09 | 2.35E-05 | Promoter (<=1kb) |
| Lelp1 | -7.878108385 | 3.15E-09 | 2.59E-05 | Exon (ENSMUST00000200051.1/ENSMUST00000200051.1_ exon 1 of 4) |
| Mmp14 | 3.161928333 | 4.57E-09 | 3.01E-05 | Promoter (<=1kb) |
| Bbox1 | -7.867639742 | 7.14E-09 | 3.92E-05 | 3' UTR |
| Fhdc1 | 7.915475755 | 1.70E-08 | 8.03E-05 | Distal Intergenic |
| 2610035D17Rik | 7.585285486 | 2.25E-08 | 8.65E-05 | Intron (ENSMUST00000136438.7/72386_ intron 1 of 2) |
| 1110002J07Rik | 5.169041502 | 2.36E-08 | 8.65E-05 | Promoter (<=1kb) |
| 0610040F04Rik | 7.359214636 | 2.93E-08 | 9.66E-05 | Distal Intergenic |
| Pmepa1 | 8.007181666 | 3.56E-08 | 0.000106669 | Promoter (<=1kb) |
| Fut4 | 7.50793166 | 4.67E-08 | 0.000126983 | Promoter (<=1kb) |
| Ap5z1 | 7.466777237 | 5.01E-08 | 0.000126983 | Intron (ENSMUST00000072837.6/17425_ intron 2 of 8) |
| Wwc1 | 7.526877722 | 8.26E-08 | 0.000194533 | Exon (ENSMUST00000018993.6/211652_ exon 2 of 23) |
| Nxn | 4.782204659 | 9.08E-08 | 0.000199573 | Intron (ENSMUST00000021204.3/18230_ intron 1 of 7) |
| Sspn | 7.236964656 | 1.05E-07 | 0.000214668 | Exon (ENSMUST00000203815.1/ENSMUST00000203815.1_ exon 1 of 1) |
| Adgrl2 | 7.525937767 | 1.19E-07 | 0.000214668 | Promoter (<=1kb) |
| Gm45521 | 7.217904269 | 1.21E-07 | 0.000214668 | Promoter (<=1kb) |
| Rnf38 | 7.384147031 | 1.24E-07 | 0.000214668 | Promoter (<=1kb) |
| Mir3971 | 7.579343401 | 1.39E-07 | 0.000221582 | 3' UTR |
| Morf4l1-ps1 | 7.046195381 | 1.41E-07 | 0.000221582 | Intron (ENSMUST00000231352.1/210126_ intron 3 of 6) |
| Rac1 | 7.214888044 | 1.58E-07 | 0.000232619 | Exon (ENSMUST00000080537.13/19353_ exon 3 of 6) |
| Cebpd | 5.020330345 | 1.62E-07 | 0.000232619 | Intron (ENSMUST00000040248.8/224008_ intron 10 of 19) |
| Gm12992 | 7.213102047 | 1.78E-07 | 0.000243061 | Promoter (<=1kb) |
| Gm19461 | 7.290113288 | 1.89E-07 | 0.000243061 | Intron (ENSMUST00000212252.1/240753_ intron 3 of 9) |
| B230217C12Rik | 7.114782615 | 1.93E-07 | 0.000243061 | Intron (ENSMUST00000043843.11/16796_ intron 4 of 6) |
| Sprtn | 7.155674484 | 1.99E-07 | 0.000243061 | Intron (ENSMUST00000034469.6/112405_ intron 1 of 4) |
| Runx1 | 4.34544234 | 2.07E-07 | 0.000243543 | 5' UTR |
| Mir6995 | 6.935914231 | 2.67E-07 | 0.00029348 | 5' UTR |
| Dhrs3 | 7.17342236 | 2.69E-07 | 0.00029348 | 3' UTR |
| Arhgef10l | 7.09487245 | 2.76E-07 | 0.00029348 | Intron (ENSMUST00000039204.9/72754_ intron 1 of 27) |
| Rassf2 | 7.220163593 | 2.89E-07 | 0.000297713 | Exon (ENSMUST00000155829.1/215653_ exon 1 of 6) |
| 4930511A08Rik | -6.982054827 | 3.02E-07 | 0.000302227 | Intron (ENSMUST00000055224.14/14924_ intron 1 of 21) |
| Pip5k1c | 7.133348216 | 3.12E-07 | 0.000302381 | Intron (ENSMUST00000163075.7/18717_ intron 1 of 16) |
| Artn | 6.904992871 | 3.97E-07 | 0.000368384 | Exon (ENSMUST00000146982.1/ENSMUST00000146982.1_ exon 1 of 3) |
| P2ry2 | 4.845634027 | 4.02E-07 | 0.000368384 | Promoter (<=1kb) |
| 4930554I06Rik | -4.178302379 | 4.40E-07 | 0.000385366 | Distal Intergenic |
| Bak1 | 6.757504767 | 4.61E-07 | 0.000385366 | 3' UTR |
| Lrp1 | 6.96334476 | 4.66E-07 | 0.000385366 | 3' UTR |
| Plin3 | 4.931507162 | 4.70E-07 | 0.000385366 | Promoter (<=1kb) |
| Ly6d | 6.909827288 | 4.91E-07 | 0.000385366 | Promoter (<=1kb) |
| Ceacam1 | 6.687151398 | 4.91E-07 | 0.000385366 | 3' UTR |
| Epas1 | 6.74752614 | 5.32E-07 | 0.000406044 | Promoter (<=1kb) |
| Gm13648 | 7.079010058 | 5.45E-07 | 0.000406044 | Distal Intergenic |
| Spns2 | 6.786454112 | 5.60E-07 | 0.000406044 | Promoter (<=1kb) |
| Ppp1r14b | 6.988051063 | 5.66E-07 | 0.000406044 | 5' UTR |
| Tex2 | 6.843517318 | 5.91E-07 | 0.000414595 | Exon (ENSMUST00000184424.1/ENSMUST00000184424.1_ exon 1 of 1) |
| Slc39a4 | 6.9114835 | 6.22E-07 | 0.000427342 | Promoter (<=1kb) |
| Gm5431 | -6.933810612 | 6.51E-07 | 0.000438051 | Promoter (<=1kb) |
| Fzd2 | 6.741307461 | 7.01E-07 | 0.000462329 | Promoter (<=1kb) |
| Mir721 | 6.798494681 | 7.63E-07 | 0.000493012 | Intron (ENSMUST00000176216.8/13047_ intron 2 of 22) |
| Adam19 | 7.094469345 | 7.95E-07 | 0.000503943 | Promoter (<=1kb) |
| Baiap2 | 5.255037408 | 8.54E-07 | 0.000522902 | Intron (ENSMUST00000075180.11/108100_ intron 3 of 13) |
| Gm10451 | 6.824301474 | 8.59E-07 | 0.000522902 | Distal Intergenic |
| Adamts1 | 6.646108027 | 8.72E-07 | 0.000522902 | Promoter (<=1kb) |

| | | | | |
|---|---|---|---|---|
| Akap13 | 6.804405752 | 9.53E-07 | 0.000561379 | Exon (ENSMUST00000207998.1/75547_ exon 1 of 9) |
| Ak1 | 6.802823824 | 9.71E-07 | 0.000561949 | 5' UTR |
| Apobec1 | 6.879046218 | 9.98E-07 | 0.000567071 | Promoter (<=1kb) |
| Mir3097 | 6.850470242 | 1.07E-06 | 0.000595382 | 5' UTR |
| Ehf | 6.697903672 | 1.08E-06 | 0.000595382 | Exon (ENSMUST00000090475.9/13661_ exon 4 of 9) |
| Lif | 6.572890553 | 1.11E-06 | 0.000597327 | Promoter (1-2kb) |
| Unc5b | 6.805730699 | 1.12E-06 | 0.000597327 | Promoter (<=1kb) |
| Tinagl1 | 6.731869355 | 1.14E-06 | 0.000597817 | Promoter (<=1kb) |
| Dapl1 | 6.697061072 | 1.23E-06 | 0.00063579 | Distal Intergenic |
| Edn1 | 5.445183134 | 1.41E-06 | 0.000717443 | Promoter (<=1kb) |
| Slc8b1 | 6.501678855 | 1.46E-06 | 0.000731563 | Promoter (1-2kb) |
| Tagap1 | 3.38458191 | 1.49E-06 | 0.000733545 | 5' UTR |
| Rgs3 | 6.71922448 | 1.52E-06 | 0.000737306 | Intron (ENSMUST00000065870.13/50780_ intron 16 of 19) |
| Zbtb40 | 6.719933651 | 1.60E-06 | 0.00076528 | Promoter (<=1kb) |
| Clic6 | 6.853778339 | 1.64E-06 | 0.000774256 | Exon (ENSMUST00000180989.3/ENSMUST00000180989.3_ exon 1 of 5) |
| Cox4i2 | 6.479444188 | 1.73E-06 | 0.000802992 | 3' UTR |
| Gm15880 | 6.689049071 | 1.76E-06 | 0.000805467 | Promoter (<=1kb) |
| Tnk2os | 6.554718322 | 1.79E-06 | 0.000810537 | Promoter (2-3kb) |
| Klf5 | 6.804198959 | 1.82E-06 | 0.000810537 | Distal Intergenic |
| Zfp36l1 | -6.456952669 | 1.86E-06 | 0.000818406 | 3' UTR |
| 1700012D01Rik | -6.534926168 | 1.96E-06 | 0.00083962 | Promoter (2-3kb) |
| Txn2 | 6.623353753 | 1.97E-06 | 0.00083962 | 3' UTR |
| Sirt2 | 4.850411247 | 1.99E-06 | 0.00083962 | Promoter (<=1kb) |
| Kcnn4 | 6.444706407 | 2.13E-06 | 0.000887063 | Promoter (<=1kb) |
| Pde2a | 6.577386352 | 2.19E-06 | 0.000902839 | 5' UTR |
| Soat2 | 6.690200029 | 2.23E-06 | 0.000908058 | Distal Intergenic |
| Gstt3 | 6.672653515 | 2.36E-06 | 0.000939313 | Promoter (<=1kb) |
| Tob2 | 6.937263891 | 2.36E-06 | 0.000939313 | Distal Intergenic |
| Sphk1 | 6.486243347 | 2.43E-06 | 0.000955381 | Promoter (<=1kb) |
| Niban2 | 6.570565817 | 2.47E-06 | 0.000957026 | Intron (ENSMUST00000028135.14/227737_ intron 1 of 13) |
| Eml1 | 6.648701456 | 2.50E-06 | 0.000957065 | Promoter (<=1kb) |
| Tpm2 | 6.336565962 | 2.59E-06 | 0.000981596 | Promoter (<=1kb) |
| Slc22a16 | 6.310053645 | 2.81E-06 | 0.001032213 | Distal Intergenic |
| Get4 | 6.453620368 | 2.84E-06 | 0.001032213 | Promoter (2-3kb) |
| Fgfr1 | 4.039932774 | 2.87E-06 | 0.001032213 | 5' UTR |
| Basp1 | 6.399916559 | 2.87E-06 | 0.001032213 | Distal Intergenic |
| Krt31 | -6.44078514 | 2.90E-06 | 0.001032213 | Exon (ENSMUST00000238944.1/ENSMUST00000238944.1_ exon 1 of 1) |
| Gm10248 | -6.378836154 | 2.91E-06 | 0.001032213 | Distal Intergenic |
| BB123696 | 6.41221738 | 3.15E-06 | 0.001096523 | Distal Intergenic |
| Fasn | 6.644752298 | 3.17E-06 | 0.001096523 | Intron (ENSMUST00000056781.4/71276_ intron 18 of 18) |
| Ndfip2 | -6.565534259 | 3.19E-06 | 0.001096523 | Distal Intergenic |
| Ddah2 | 4.526769556 | 3.25E-06 | 0.001096523 | Promoter (<=1kb) |
| Htra3 | 6.509010305 | 3.38E-06 | 0.001096523 | 5' UTR |
| Gdpd5 | 6.324893033 | 3.39E-06 | 0.001096523 | 5' UTR |
| Pxn | 6.35187019 | 3.40E-06 | 0.001096523 | Exon (ENSMUST00000125007.7/19303_ exon 1 of 11) |
| Niban2 | 6.575286148 | 3.45E-06 | 0.001096523 | Exon (ENSMUST00000154514.7/227737_ exon 1 of 6) |
| Anpep | 6.372472485 | 3.52E-06 | 0.001096523 | Intron (ENSMUST00000075657.7/11778_ intron 4 of 5) |
| Gm21057 | 6.585915218 | 3.53E-06 | 0.001096523 | Promoter (<=1kb) |
| Plekha6 | 6.323385841 | 3.53E-06 | 0.001096523 | Intron (ENSMUST00000212252.1/240753_ intron 3 of 9) |
| A530013C23Rik | 6.71247221 | 3.54E-06 | 0.001096523 | Distal Intergenic |
| Sh2d4b | 6.482582083 | 3.55E-06 | 0.001096523 | Distal Intergenic |
| A930015D03Rik | 6.537566685 | 3.56E-06 | 0.001096523 | Promoter (2-3kb) |
| Zbtb16 | 6.340910313 | 3.66E-06 | 0.001099142 | Intron (ENSMUST00000093852.4/235320_ intron 2 of 6) |
| Itgb1 | 6.460738426 | 3.66E-06 | 0.001099142 | Exon (ENSMUST00000212623.1/ENSMUST00000212623.1_ exon 1 of 1) |
| Dusp16 | -5.018669707 | 3.68E-06 | 0.001099142 | Distal Intergenic |
| Sulf1 | -6.471326476 | 3.70E-06 | 0.001099142 | Intron (ENSMUST00000177608.7/240725_ intron 2 of 21) |

| | | | | |
|---|---|---|---|---|
| Abcc3 | 6.30089846 | 3.77E-06 | 0.001109551 | Promoter (<=1kb) |
| Zc2hc1a | 6.601544117 | 3.84E-06 | 0.001121529 | Promoter (<=1kb) |
| Hdac7 | 6.534710599 | 4.05E-06 | 0.00116827 | Intron (ENSMUST00000079838.13/56233_ intron 1 of 24) |
| Tbc1d1 | -6.375209096 | 4.07E-06 | 0.00116827 | Intron (ENSMUST00000043893.12/57915_ intron 2 of 21) |
| Cemip2 | 4.598192673 | 4.11E-06 | 0.00116894 | Exon (ENSMUST00000096194.8/83921_ exon 12 of 25) |
| Arrb1 | 3.245242762 | 4.36E-06 | 0.001229747 | Promoter (<=1kb) |
| Ppt1 | -6.58391089 | 4.51E-06 | 0.001260402 | Intron (ENSMUST00000119988.1/ENSMUST00000119988.1_ intron 1 of 3) |
| Sh2b2 | 6.405581777 | 4.73E-06 | 0.001301954 | Promoter (<=1kb) |
| Chaserr | 6.549910914 | 4.74E-06 | 0.001301954 | Distal Intergenic |
| Pik3r1 | 6.388154006 | 4.78E-06 | 0.001303298 | Intron (ENSMUST00000055518.12/18708_ intron 2 of 15) |
| Rgs3 | 6.584276079 | 4.92E-06 | 0.00132937 | Exon (ENSMUST00000084521.10/50780_ exon 11 of 16) |
| Map3k13 | 6.504059541 | 5.02E-06 | 0.001338841 | 5' UTR |
| Gm15417 | 6.251419177 | 5.04E-06 | 0.001338841 | 5' UTR |
| Ino80c | 6.348476433 | 5.12E-06 | 0.001350044 | Exon (ENSMUST00000153360.7/225280_ exon 3 of 5) |
| Acer3 | 6.208783548 | 5.26E-06 | 0.001377631 | Distal Intergenic |
| Gm2176 | 6.431386411 | 5.41E-06 | 0.001401672 | Distal Intergenic |
| Prr13 | 2.495041756 | 5.44E-06 | 0.001401672 | Promoter (<=1kb) |
| Fry | -6.535949716 | 5.48E-06 | 0.001401672 | Intron (ENSMUST00000200960.3/320365_ intron 2 of 4) |
| E130114P18Rik | -6.30933759 | 5.53E-06 | 0.001403326 | Intron (ENSMUST00000075448.12/18027_ intron 4 of 10) |
| 2610035D17Rik | 6.217550609 | 5.61E-06 | 0.00140427 | Intron (ENSMUST00000136438.7/72386_ intron 1 of 2) |
| Dgat2 | 6.427225096 | 5.62E-06 | 0.00140427 | Intron (ENSMUST00000033001.5/67800_ intron 1 of 7) |
| Syt12 | 6.441698203 | 5.66E-06 | 0.00140427 | Promoter (<=1kb) |
| Tcp11 | 6.921141885 | 5.72E-06 | 0.001406749 | Intron (ENSMUST00000114842.8/224650_ intron 12 of 24) |
| Gm6525 | -6.599553828 | 5.79E-06 | 0.001413237 | Intron (ENSMUST00000107692.7/80890_ intron 1 of 11) |
| Naglu | 6.343567979 | 6.25E-06 | 0.001515702 | Exon (ENSMUST00000103110.9/11975_ exon 19 of 21) |
| Hsp90ab1 | 6.472982733 | 6.43E-06 | 0.001547102 | 3' UTR |
| Sh3pxd2a | 6.358390363 | 6.74E-06 | 0.001610279 | Intron (ENSMUST00000081619.9/14218_ intron 1 of 14) |
| Tab1 | 6.480651465 | 6.82E-06 | 0.001611941 | Intron (ENSMUST00000230479.1/66513_ intron 1 of 1) |
| Atl2 | 6.397041538 | 6.84E-06 | 0.001611941 | Distal Intergenic |
| 1110028F18Rik | 6.592137577 | 7.10E-06 | 0.00166066 | Promoter (<=1kb) |
| Hoxd9 | -5.084254318 | 7.25E-06 | 0.001684307 | Promoter (<=1kb) |
| A330093E20Rik | 6.328397976 | 7.53E-06 | 0.001736419 | Distal Intergenic |
| Atp2b1 | -6.323829635 | 8.02E-06 | 0.00183716 | Distal Intergenic |
| Shmt2 | 6.401949973 | 8.32E-06 | 0.001893028 | Promoter (<=1kb) |
| Agmat | 3.998450913 | 8.52E-06 | 0.001923755 | Exon (ENSMUST00000038014.10/214063_ exon 10 of 15) |
| Tmem176a | 6.433566043 | 8.61E-06 | 0.001932207 | Distal Intergenic |
| Serpinh1 | 6.127143197 | 8.69E-06 | 0.001935064 | Promoter (<=1kb) |
| 9130015G15Rik | 6.366537838 | 8.83E-06 | 0.001954524 | Intron (ENSMUST00000183659.7/78286_ intron 2 of 39) |
| 8030442B05Rik | -6.239009102 | 8.89E-06 | 0.001954736 | Promoter (<=1kb) |
| Podxl | 3.890769848 | 9.38E-06 | 0.002047674 | 3' UTR |
| Synpo | 5.178951423 | 9.48E-06 | 0.002057379 | Exon (ENSMUST00000237783.1/15531_ exon 8 of 15) |
| Or13c7 | 6.246061714 | 9.66E-06 | 0.002074989 | Distal Intergenic |
| Fxyd4 | 6.032706799 | 9.69E-06 | 0.002074989 | Promoter (<=1kb) |
| Hic1 | 6.260021383 | 9.82E-06 | 0.002080651 | Promoter (<=1kb) |
| Cd47 | -6.331253909 | 9.84E-06 | 0.002080651 | Distal Intergenic |
| Asb7 | 4.739624263 | 1.02E-05 | 0.002131934 | 3' UTR |
| Mink1 | 6.068978081 | 1.02E-05 | 0.002131934 | Intron (ENSMUST00000072237.12/50932_ intron 1 of 32) |
| Etv6 | -6.208503905 | 1.04E-05 | 0.002153074 | Intron (ENSMUST00000081028.12/14011_ intron 2 of 7) |
| Nrp2 | 6.166821579 | 1.05E-05 | 0.002164909 | Promoter (2-3kb) |
| Magi3 | -6.269784984 | 1.06E-05 | 0.002165563 | Distal Intergenic |
| Fam118b | 6.323673308 | 1.07E-05 | 0.002176467 | Promoter (2-3kb) |
| Fam3b | 6.283754768 | 1.08E-05 | 0.002176467 | Intron (ENSMUST00000049721.8/52793_ intron 2 of 7) |
| C330024C12Rik | 6.161363638 | 1.10E-05 | 0.002189731 | Distal Intergenic |
| Max | -4.291361987 | 1.10E-05 | 0.002189731 | Distal Intergenic |
| 1700096K18Rik | 6.197576564 | 1.11E-05 | 0.002189731 | Distal Intergenic |
| Tbkbp1 | 6.281783182 | 1.12E-05 | 0.002189731 | Promoter (<=1kb) |

| | | | | |
|---|---|---|---|---|
| Ier5l | 5.91996131 | 1.12E-05 | 0.002189731 | Distal Intergenic |
| B4galt2 | 6.385143735 | 1.13E-05 | 0.002189731 | Promoter (<=1kb) |
| Chd9 | 4.890876766 | 1.13E-05 | 0.002189731 | Exon (ENSMUST00000211215.1/109151_ exon 3 of 3) |
| Ly6g6c | 4.671941832 | 1.16E-05 | 0.002222586 | Promoter (<=1kb) |
| Abi3 | 6.307173884 | 1.16E-05 | 0.002222586 | Promoter (<=1kb) |
| Fndc3b | 3.866630842 | 1.17E-05 | 0.002229513 | Intron (ENSMUST00000195008.5/72007_ intron 1 of 26) |
| Plcd1 | 6.045262354 | 1.20E-05 | 0.002267011 | Promoter (<=1kb) |
| Ucp3 | -6.160569228 | 1.21E-05 | 0.00227704 | Exon (ENSMUST00000098259.10/277939_ exon 21 of 32) |
| 6530402F18Rik | 6.408503059 | 1.23E-05 | 0.002286321 | Promoter (<=1kb) |
| Cipc | 6.398368157 | 1.23E-05 | 0.002286321 | Distal Intergenic |
| Mnt | 6.071710225 | 1.23E-05 | 0.002286321 | Promoter (2-3kb) |
| Tril | 6.40644872 | 1.27E-05 | 0.002345337 | Promoter (<=1kb) |
| Gm32865 | -6.26304195 | 1.29E-05 | 0.002349269 | Distal Intergenic |
| Mrps27 | 6.385508541 | 1.29E-05 | 0.002349269 | Intron (ENSMUST00000052249.6/218506_ intron 2 of 10) |
| Cxcl16 | 6.303400819 | 1.30E-05 | 0.002359393 | 3' UTR |
| Spef1l | 5.952427745 | 1.36E-05 | 0.002410958 | 3' UTR |
| Btg1 | -6.31937218 | 1.36E-05 | 0.002410958 | Distal Intergenic |
| Mir701 | 6.262379645 | 1.36E-05 | 0.002410958 | Exon (ENSMUST00000199644.1/ENSMUST00000199644.1_ exon 1 of 1) |
| Foxp1 | 6.311539748 | 1.37E-05 | 0.002410958 | Exon (ENSMUST00000158074.1/ENSMUST00000158074.1_ exon 1 of 1) |
| Lad1 | 2.906796651 | 1.37E-05 | 0.002410958 | Promoter (<=1kb) |
| Mgst1 | 5.957953109 | 1.37E-05 | 0.002410958 | Promoter (<=1kb) |
| Man1c1 | 6.143950421 | 1.39E-05 | 0.002429818 | Promoter (<=1kb) |
| Tm4sf4 | -6.205781169 | 1.42E-05 | 0.002472872 | Intron (ENSMUST00000029380.13/97064_ intron 4 of 6) |
| Daam1 | 6.163987257 | 1.44E-05 | 0.002490877 | Intron (ENSMUST00000223272.1/208846_ intron 1 of 25) |
| Ell2 | -5.942196625 | 1.45E-05 | 0.002492498 | Intron (ENSMUST00000001583.7/192657_ intron 4 of 11) |
| Nynrin | 6.093271309 | 1.47E-05 | 0.002519725 | Promoter (<=1kb) |
| 2810414N06Rik | 5.934973037 | 1.50E-05 | 0.002542038 | Intron (ENSMUST00000148810.7/63913_ intron 5 of 13) |
| Gbp4 | -6.433270114 | 1.50E-05 | 0.002542038 | Distal Intergenic |
| Gm10532 | 6.465094921 | 1.52E-05 | 0.002554225 | Intron (ENSMUST00000165559.2/269037_ intron 9 of 11) |
| Tnfrsf1a | 6.125688789 | 1.54E-05 | 0.002574588 | Intron (ENSMUST00000176655.7/20276_ intron 8 of 12) |
| Ccnd1 | 5.874964962 | 1.55E-05 | 0.002574588 | Promoter (<=1kb) |
| Frmd6 | -5.939588829 | 1.58E-05 | 0.002614663 | Intron (ENSMUST00000220504.1/ENSMUST00000220504.1_ intron 2 of 2) |
| Soat2 | 6.0244493 | 1.59E-05 | 0.002614663 | Promoter (<=1kb) |
| Ikzf2 | -6.057313014 | 1.60E-05 | 0.002618422 | Promoter (<=1kb) |
| Mb | 6.128717213 | 1.62E-05 | 0.002633603 | 3' UTR |
| NA | 6.106033243 | 1.63E-05 | 0.002633603 | Intron (ENSMUST00000098066.8/677884_ intron 7 of 10) |
| Fgd6 | 6.303409497 | 1.64E-05 | 0.002633603 | Intron (ENSMUST00000020208.4/13998_ intron 3 of 20) |
| Ano7 | 6.450295647 | 1.64E-05 | 0.002633603 | Promoter (<=1kb) |
| Slc22a23 | -6.172047728 | 1.65E-05 | 0.002633603 | Distal Intergenic |
| Kat6b | 3.82895868 | 1.66E-05 | 0.002648279 | Intron (ENSMUST00000183201.7/54169_ intron 2 of 5) |
| Ptgfrn | 6.363118616 | 1.72E-05 | 0.002720759 | Intron (ENSMUST00000102694.3/19221_ intron 4 of 8) |
| Mllt1 | 4.863453021 | 1.73E-05 | 0.002734307 | Exon (ENSMUST00000025053.9/64144_ exon 4 of 12) |
| Ccdc97 | 2.350461437 | 1.78E-05 | 0.002781956 | 3' UTR |
| Gm7550 | 5.914713214 | 1.78E-05 | 0.002781956 | Distal Intergenic |
| Psma1 | -4.947278169 | 1.80E-05 | 0.002793209 | Intron (ENSMUST00000163996.1/71045_ intron 3 of 3) |
| Nradd | 6.287175834 | 1.82E-05 | 0.002801148 | Promoter (<=1kb) |
| Col18a1 | 6.089983032 | 1.82E-05 | 0.002801148 | Distal Intergenic |
| A730020M07Rik | 6.013027976 | 1.85E-05 | 0.002834896 | Distal Intergenic |
| Perp | -6.113050899 | 1.86E-05 | 0.00284048 | Distal Intergenic |
| Tpm1 | -5.92986988 | 1.93E-05 | 0.002924693 | Distal Intergenic |
| Map1lc3a | 6.127978695 | 1.93E-05 | 0.002924693 | Exon (ENSMUST00000139854.1/228812_ exon 2 of 4) |
| Smco4 | 6.382349652 | 1.97E-05 | 0.002965456 | Intron (ENSMUST00000215907.1/170748_ intron 1 of 2) |
| Rdh10 | -5.848104872 | 2.03E-05 | 0.003044628 | 3' UTR |
| Gab1 | -5.971296654 | 2.07E-05 | 0.003090566 | Intron (ENSMUST00000210676.1/14388_ intron 1 of 10) |
| 2210408F21Rik | -6.046872569 | 2.13E-05 | 0.003156533 | Intron (ENSMUST00000144612.7/73652_ intron 2 of 5) |
| 9430078K24Rik | 5.984018063 | 2.16E-05 | 0.003185919 | Distal Intergenic |

| | | | | |
|---|---|---|---|---|
| Umps | 6.353130127 | 2.17E-05 | 0.003185919 | Intron (ENSMUST00000115028.10/16419_ intron 10 of 14) |
| Zfp618 | 6.104234295 | 2.17E-05 | 0.003185919 | Promoter (<=1kb) |
| Slc19a1 | 6.118223845 | 2.20E-05 | 0.003205666 | 5' UTR |
| Gnai3 | 6.040609845 | 2.21E-05 | 0.003208514 | Intron (ENSMUST00000000001.4/14679_ intron 1 of 8) |
| Slc38a1 | 6.080450304 | 2.23E-05 | 0.0032215 | Intron (ENSMUST00000088454.12/105727_ intron 4 of 15) |
| Snx10 | 5.905815695 | 2.26E-05 | 0.003247866 | 5' UTR |
| Synj2 | 6.376146672 | 2.27E-05 | 0.003247866 | Distal Intergenic |
| Tnfaip2 | 5.954295222 | 2.30E-05 | 0.003286962 | Promoter (<=1kb) |
| Hibadh | 5.960444225 | 2.34E-05 | 0.003301247 | Promoter (<=1kb) |
| 9430078K24Rik | -4.685110164 | 2.34E-05 | 0.003301247 | Intron (ENSMUST00000171210.2/19363_ intron 7 of 10) |
| Tns1 | 6.290745858 | 2.34E-05 | 0.003301247 | Exon (ENSMUST00000187281.6/21961_ exon 1 of 7) |
| Vsig8 | 6.120068931 | 2.36E-05 | 0.003312436 | 3' UTR |
| Mirlet7a-1 | -6.021514671 | 2.40E-05 | 0.003351146 | Distal Intergenic |
| Hemk1 | 6.155142772 | 2.42E-05 | 0.003364174 | Distal Intergenic |
| Rrp1b | 5.745501839 | 2.46E-05 | 0.003412645 | Intron (ENSMUST00000002145.11/74377_ intron 7 of 7) |
| 2310001H17Rik | -6.279965331 | 2.51E-05 | 0.003457497 | Intron (ENSMUST00000181238.1/76432_ intron 1 of 2) |
| Agr3 | -5.834938433 | 2.56E-05 | 0.003516658 | Distal Intergenic |
| Tspan15 | 6.238921494 | 2.57E-05 | 0.003521606 | Intron (ENSMUST00000047883.10/70423_ intron 1 of 7) |
| Gm4668 | 5.823539085 | 2.60E-05 | 0.003527433 | Promoter (<=1kb) |
| Tagln2 | 5.989016415 | 2.60E-05 | 0.003527433 | Exon (ENSMUST00000135267.7/93842_ exon 4 of 6) |
| Gm4668 | 5.850530244 | 2.62E-05 | 0.003537626 | Distal Intergenic |
| Rras2 | 5.975229857 | 2.63E-05 | 0.003542354 | Exon (ENSMUST00000046687.15/233744_ exon 7 of 16) |
| Sdc3 | 6.079576622 | 2.65E-05 | 0.003550408 | Promoter (2-3kb) |
| Septin9 | 6.143373008 | 2.70E-05 | 0.003599353 | 3' UTR |
| Sipa1l3 | 5.822468807 | 2.72E-05 | 0.003599353 | Intron (ENSMUST00000085809.10/74206_ intron 3 of 22) |
| 2310001H17Rik | -6.021448644 | 2.72E-05 | 0.003599353 | Exon (ENSMUST00000181238.1/76432_ exon 3 of 3) |
| Itga5 | 5.691922928 | 2.73E-05 | 0.00360037 | Promoter (<=1kb) |
| Mgat5 | -6.095587048 | 2.74E-05 | 0.003601811 | Intron (ENSMUST00000189427.6/107895_ intron 1 of 7) |
| Tm4sf1 | 6.121227736 | 2.77E-05 | 0.003608956 | Promoter (<=1kb) |
| Ccdc85c | 5.746456398 | 2.77E-05 | 0.003608956 | Intron (ENSMUST00000222310.1/668158_ intron 1 of 5) |
| Gm4651 | 5.875217043 | 2.81E-05 | 0.003648525 | Exon (ENSMUST00000212208.1/194401_ exon 6 of 11) |
| Pxk | -4.906818324 | 2.86E-05 | 0.003684264 | Promoter (1-2kb) |
| 1700001K23Rik | 5.808014524 | 2.88E-05 | 0.003684264 | Distal Intergenic |
| Trp63 | 5.941456796 | 2.90E-05 | 0.003684264 | 5' UTR |
| Ibtk | 5.853885203 | 2.92E-05 | 0.003684264 | Exon (ENSMUST00000083772.1/ENSMUST00000083772.1_ exon 1 of 1) |
| Prr5l | 6.009148877 | 2.92E-05 | 0.003684264 | 5' UTR |
| Mir29a | 5.605969766 | 2.93E-05 | 0.003684264 | Promoter (<=1kb) |
| Zbtb16 | 6.256510654 | 2.94E-05 | 0.003684264 | Intron (ENSMUST00000093852.4/235320_ intron 3 of 6) |
| Esyt3 | 5.941321917 | 2.94E-05 | 0.003684264 | Exon (ENSMUST00000035045.14/17532_ exon 5 of 6) |
| Lamc2 | 5.849741086 | 2.95E-05 | 0.003684264 | Intron (ENSMUST00000186368.6/226518_ intron 7 of 11) |
| 9430078K24Rik | -6.067165363 | 2.95E-05 | 0.003684264 | Intron (ENSMUST00000171210.2/19363_ intron 9 of 10) |
| Entrep1 | 6.01320648 | 3.05E-05 | 0.003771833 | Intron (ENSMUST00000096164.5/381217_ intron 2 of 10) |
| Fgf1 | 5.866283199 | 3.07E-05 | 0.003771833 | Intron (ENSMUST00000148850.7/71302_ intron 1 of 5) |
| Ensa | -6.095554664 | 3.08E-05 | 0.003771833 | 3' UTR |
| Itm2b | 5.980603123 | 3.09E-05 | 0.003771833 | Distal Intergenic |
| Tox3 | -6.124047299 | 3.09E-05 | 0.003771833 | Distal Intergenic |
| Kctd10 | 5.706391938 | 3.10E-05 | 0.003771833 | Exon (ENSMUST00000102581.10/330171_ exon 2 of 7) |
| 9430078K24Rik | 3.799923715 | 3.10E-05 | 0.003771833 | Intron (ENSMUST00000171210.2/19363_ intron 8 of 10) |
| Gstm5 | 5.963567006 | 3.13E-05 | 0.003796216 | Exon (ENSMUST00000198085.1/ENSMUST00000198085.1_ exon 1 of 1) |
| 7530416G11Rik | 5.937710236 | 3.14E-05 | 0.003796343 | Intron (ENSMUST00000163242.2/54138_ intron 9 of 11) |
| Coro2a | 5.635337871 | 3.20E-05 | 0.003839344 | Intron (ENSMUST00000107756.3/107684_ intron 1 of 11) |
| Arl4d | 3.811496052 | 3.22E-05 | 0.003839344 | Distal Intergenic |
| Srcin1 | 6.115271912 | 3.23E-05 | 0.003839344 | Promoter (<=1kb) |
| Hspg2 | 5.883499368 | 3.24E-05 | 0.003839344 | Promoter (<=1kb) |
| Plekho1 | 5.876918951 | 3.24E-05 | 0.003839344 | Exon (ENSMUST00000015889.9/67220_ exon 4 of 6) |
| Cables1 | 6.096938274 | 3.28E-05 | 0.0038818 | Distal Intergenic |

| | | | | |
|---|---|---|---|---|
| Ippk | -5.778980115 | 3.31E-05 | 0.003902475 | Distal Intergenic |
| Ltb4r2 | 6.05099832 | 3.33E-05 | 0.003909337 | Promoter (<=1kb) |
| Smurf2 | 5.919918711 | 3.35E-05 | 0.00391267 | Distal Intergenic |
| Ifnlr1 | 5.743081865 | 3.44E-05 | 0.004002939 | Distal Intergenic |
| Mboat2 | -6.056421818 | 3.49E-05 | 0.004038152 | Intron (ENSMUST00000222198.1/67216_ intron 1 of 4) |
| Ak3 | -4.273291236 | 3.49E-05 | 0.004038152 | Promoter (1-2kb) |
| Mir6403 | 5.875514945 | 3.51E-05 | 0.004049077 | Intron (ENSMUST00000038628.3/230815_ intron 3 of 12) |
| Pdlim2 | 5.550076285 | 3.55E-05 | 0.004082092 | Promoter (<=1kb) |
| Csf3 | 5.557992486 | 3.57E-05 | 0.004082092 | Promoter (2-3kb) |
| Pik3cb | 5.839373058 | 3.58E-05 | 0.004082092 | Distal Intergenic |
| Snx13 | -5.842528195 | 3.64E-05 | 0.004123913 | Distal Intergenic |
| Prx | -4.742848978 | 3.64E-05 | 0.004123913 | 5' UTR |
| Cgnl1 | 5.810698169 | 3.68E-05 | 0.004141909 | Promoter (<=1kb) |
| Foxp1 | -5.915601182 | 3.68E-05 | 0.004141909 | Intron (ENSMUST00000113326.8/108655_ intron 3 of 17) |
| Mir5122 | 5.661138308 | 3.73E-05 | 0.004171773 | Intron (ENSMUST00000030669.7/20544_ intron 1 of 11) |
| Ikzf1 | 5.991410161 | 3.78E-05 | 0.004171773 | Promoter (2-3kb) |
| Ate1 | 5.974778405 | 3.78E-05 | 0.004171773 | 5' UTR |
| Znrf2 | 5.865953276 | 3.79E-05 | 0.004171773 | Intron (ENSMUST00000079869.12/387524_ intron 1 of 4) |
| Rilpl2 | 6.218863378 | 3.79E-05 | 0.004171773 | Promoter (1-2kb) |
| Gm6260 | -6.280876929 | 3.80E-05 | 0.004171773 | Exon (ENSMUST00000192421.1/ENSMUST00000192421.1_ exon 1 of 2) |
| Ptgs1 | 5.869111168 | 3.81E-05 | 0.004171773 | Promoter (<=1kb) |
| D030018L15Rik | 5.925055065 | 3.82E-05 | 0.004171773 | Distal Intergenic |
| Qsox1 | -5.762763415 | 3.83E-05 | 0.004171773 | 3' UTR |
| Mx1 | -6.003267936 | 3.83E-05 | 0.004171773 | Exon (ENSMUST00000231393.1/ENSMUST00000231393.1_ exon 1 of 1) |
| 5830428M24Rik | 5.530659078 | 3.88E-05 | 0.004211228 | Distal Intergenic |
| Kcnk6 | 5.981041947 | 3.93E-05 | 0.004249191 | Promoter (<=1kb) |
| Nrip1 | -6.060262825 | 3.99E-05 | 0.004294373 | 5' UTR |
| Epha2 | 4.510623337 | 4.00E-05 | 0.004295709 | Promoter (<=1kb) |
| D030018L15Rik | -5.795812082 | 4.02E-05 | 0.004306282 | Distal Intergenic |
| Mtarc2 | 5.946115693 | 4.08E-05 | 0.004345448 | Distal Intergenic |
| Epm2aip1 | -5.78983307 | 4.09E-05 | 0.004345448 | Distal Intergenic |
| Slco1a6 | 5.743830028 | 4.10E-05 | 0.004345448 | Intron (ENSMUST00000174455.1/28254_ intron 1 of 2) |
| 1700056E22Rik | 5.862946166 | 4.11E-05 | 0.004345448 | Distal Intergenic |
| Gm19519 | -5.768490053 | 4.14E-05 | 0.00435842 | Intron (ENSMUST00000236816.1/100503033_ intron 1 of 5) |
| Sema4b | 5.662092204 | 4.16E-05 | 0.004364571 | Distal Intergenic |
| Lpin2 | 6.084918129 | 4.17E-05 | 0.004364571 | Promoter (2-3kb) |
| Fkbp5 | 5.805928946 | 4.20E-05 | 0.004378967 | 5' UTR |
| Rgmb | 5.723748439 | 4.21E-05 | 0.004378967 | Intron (ENSMUST00000170578.2/68799_ intron 2 of 2) |
| Adcy6 | 5.910770291 | 4.24E-05 | 0.004396012 | 5' UTR |
| Mir7057 | 5.838135066 | 4.27E-05 | 0.00441185 | Exon (ENSMUST00000015277.13/233328_ exon 3 of 34) |
| Rassf3 | -5.459949601 | 4.28E-05 | 0.00441185 | Exon (ENSMUST00000219500.1/192678_ exon 1 of 4) |
| Robo4 | 5.825302763 | 4.33E-05 | 0.004446655 | 3' UTR |
| Nod2 | 5.754369541 | 4.41E-05 | 0.004515585 | Promoter (<=1kb) |
| Ncoa3 | 5.736842279 | 4.42E-05 | 0.004516497 | Distal Intergenic |
| Cux1 | -5.652336265 | 4.47E-05 | 0.004540019 | Exon (ENSMUST00000192800.1/ENSMUST00000192800.1_ exon 1 of 1) |
| Plekhf2 | 6.163936073 | 4.48E-05 | 0.004540019 | Distal Intergenic |
| Adap2 | 6.220415171 | 4.51E-05 | 0.004560864 | Promoter (<=1kb) |
| Galnt3 | -5.797767887 | 4.53E-05 | 0.004565502 | Intron (ENSMUST00000028378.3/14425_ intron 1 of 10) |
| Laptm5 | -5.669553231 | 4.58E-05 | 0.004605354 | Distal Intergenic |
| 5830408C22Rik | 5.969030813 | 4.62E-05 | 0.004615944 | Intron (ENSMUST00000064883.13/83436_ intron 2 of 11) |
| Car9 | 5.795229768 | 4.62E-05 | 0.004615944 | Promoter (<=1kb) |
| Cd82 | -5.660149727 | 4.65E-05 | 0.004615944 | Intron (ENSMUST00000099693.8/100038525_ intron 2 of 4) |
| Tspan32 | 6.112676976 | 4.66E-05 | 0.004615944 | Promoter (2-3kb) |
| Tmem37 | 5.858623517 | 4.67E-05 | 0.004615944 | Promoter (<=1kb) |
| Lrrc71 | 5.510282547 | 4.68E-05 | 0.004615944 | 5' UTR |
| Fam83h | 5.67517184 | 4.71E-05 | 0.004635971 | 3' UTR |

| | | | | |
|---|---|---|---|---|
| Stk39 | 6.053423149 | 4.75E-05 | 0.004661788 | Intron (ENSMUST00000102715.3/53416_ intron 1 of 17) |
| Rsu1 | 5.73516595 | 4.81E-05 | 0.004706354 | Intron (ENSMUST00000091436.6/65969_ intron 31 of 66) |
| Ccdc68 | -2.175675949 | 4.88E-05 | 0.004765262 | Promoter (<=1kb) |
| Clic6 | 5.965725866 | 4.99E-05 | 0.004846911 | Intron (ENSMUST00000023673.13/12394_ intron 5 of 7) |
| Fermt1 | -5.753499992 | 5.00E-05 | 0.004846911 | Promoter (<=1kb) |
| Mbp | 5.473969648 | 5.02E-05 | 0.004849991 | Intron (ENSMUST00000091789.10/17196_ intron 2 of 3) |
| Nfe2l3 | 5.814491507 | 5.03E-05 | 0.004849991 | Promoter (<=1kb) |
| Tfeb | 5.785854142 | 5.06E-05 | 0.004861185 | 5' UTR |
| Arap3 | -5.495031085 | 5.12E-05 | 0.004908973 | Distal Intergenic |
| Pcdh1 | 5.628109641 | 5.16E-05 | 0.004908973 | Exon (ENSMUST00000161701.2/75599_ exon 2 of 4) |
| Glrp1 | 5.642799075 | 5.18E-05 | 0.004908973 | Promoter (<=1kb) |
| Trim8 | 5.465798509 | 5.18E-05 | 0.004908973 | Exon (ENSMUST00000123252.1/24069_ exon 1 of 3) |
| Arhgap17 | 5.692098173 | 5.18E-05 | 0.004908973 | Intron (ENSMUST00000206721.1/30949_ intron 1 of 2) |
| Dbx1 | 5.642592309 | 5.29E-05 | 0.004986828 | Intron (ENSMUST00000183659.7/78286_ intron 38 of 39) |
| C77080 | 5.744752833 | 5.29E-05 | 0.004986828 | Promoter (<=1kb) |
| Susd6 | 5.693122429 | 5.36E-05 | 0.00503412 | Intron (ENSMUST00000220238.1/217684_ intron 2 of 6) |
| Vgll4 | 5.755514995 | 5.38E-05 | 0.005041073 | Intron (ENSMUST00000032459.13/232334_ intron 2 of 4) |
| Sesn3 | -5.696200539 | 5.44E-05 | 0.005077504 | Distal Intergenic |
| Arhgap18 | -5.784808964 | 5.48E-05 | 0.005077504 | Intron (ENSMUST00000176060.7/73910_ intron 1 of 4) |
| 4930543E12Rik | 5.553389649 | 5.48E-05 | 0.005077504 | Distal Intergenic |
| Cyth3 | 5.879154476 | 5.48E-05 | 0.005077504 | Intron (ENSMUST00000116456.9/19159_ intron 1 of 12) |
| Tnfrsf21 | -5.449904515 | 5.50E-05 | 0.005079939 | Intron (ENSMUST00000024708.5/94185_ intron 4 of 5) |
| Nabp1 | 5.431566765 | 5.52E-05 | 0.005080113 | Exon (ENSMUST00000194442.1/ENSMUST00000194442.1_ exon 1 of 1) |
| Slc7a11 | 2.975870602 | 5.53E-05 | 0.005080113 | Promoter (<=1kb) |
| Chst9 | -5.877717289 | 5.55E-05 | 0.005082316 | Intron (ENSMUST00000053017.12/71367_ intron 1 of 3) |
| Ube2n | 5.69483564 | 5.57E-05 | 0.005084108 | Intron (ENSMUST00000099329.4/93765_ intron 1 of 3) |
| Gda | 5.92486063 | 5.64E-05 | 0.005113198 | Intron (ENSMUST00000087600.9/14544_ intron 1 of 13) |
| Mcc | 5.717843612 | 5.65E-05 | 0.005113198 | Promoter (<=1kb) |
| Smu1 | 5.567559867 | 5.65E-05 | 0.005113198 | Exon (ENSMUST00000121600.1/ENSMUST00000121600.1_ exon 1 of 1) |
| Ccdc122 | 5.646569209 | 5.66E-05 | 0.005113198 | Distal Intergenic |
| Usp31 | 2.688175721 | 5.73E-05 | 0.005164397 | Promoter (<=1kb) |
| Nsd3 | -5.846175463 | 5.82E-05 | 0.005209944 | Intron (ENSMUST00000155861.7/234135_ intron 4 of 9) |
| Smco4 | 5.246103504 | 5.82E-05 | 0.005209944 | 5' UTR |
| Pygb | 5.921593273 | 5.83E-05 | 0.005209944 | Intron (ENSMUST00000056149.14/76192_ intron 4 of 12) |
| Slc25a51 | 5.64177193 | 5.88E-05 | 0.005241522 | 3' UTR |
| Il12a | 5.306294167 | 5.93E-05 | 0.0052507 | 5' UTR |
| H2-Ab1 | 3.854302161 | 5.94E-05 | 0.0052507 | 5' UTR |
| Mir1961 | 5.429420814 | 5.94E-05 | 0.0052507 | Intron (ENSMUST00000113055.8/27428_ intron 2 of 10) |
| Cnot8 | 5.725815571 | 6.04E-05 | 0.005317176 | 5' UTR |
| Polr2b | 5.503864123 | 6.05E-05 | 0.005317176 | Intron (ENSMUST00000046746.9/29817_ intron 1 of 4) |
| Cab39l | 5.690831375 | 6.09E-05 | 0.005322884 | 5' UTR |
| Hebp2 | 5.841022377 | 6.10E-05 | 0.005322884 | Intron (ENSMUST00000207038.1/215819_ intron 2 of 6) |
| Rgs3 | 5.491829766 | 6.10E-05 | 0.005322884 | 5' UTR |
| Hoxa1 | 5.707331241 | 6.14E-05 | 0.005338115 | Promoter (<=1kb) |
| Zc3hc1 | 5.903215488 | 6.24E-05 | 0.005410314 | Promoter (<=1kb) |
| Mst1r | 5.777976302 | 6.30E-05 | 0.005454285 | Promoter (<=1kb) |
| Sorbs2os | 5.683373303 | 6.32E-05 | 0.005459307 | Intron (ENSMUST00000067065.13/234214_ intron 2 of 25) |
| Susd1 | 5.502300805 | 6.35E-05 | 0.005463274 | Intron (ENSMUST00000040166.13/634731_ intron 8 of 16) |
| Prlr | -3.955976309 | 6.38E-05 | 0.005480156 | Promoter (<=1kb) |
| Apoe | 5.941656795 | 6.43E-05 | 0.005510224 | Promoter (<=1kb) |
| Enpp6 | 5.375080456 | 6.49E-05 | 0.005536198 | Distal Intergenic |
| Epas1 | 5.707539036 | 6.50E-05 | 0.005536198 | Intron (ENSMUST00000024954.10/13819_ intron 1 of 15) |
| Ski | 5.639963871 | 6.52E-05 | 0.00553961 | Distal Intergenic |
| Itgb5 | 5.61682308 | 6.58E-05 | 0.005579668 | Intron (ENSMUST00000115028.10/16419_ intron 3 of 14) |
| Ccdc68 | 5.82353485 | 6.80E-05 | 0.00575005 | Intron (ENSMUST00000235804.1/ENSMUST00000235804.1_ intron 1 of 1) |
| Nod1 | -5.73155938 | 6.87E-05 | 0.005796949 | Intron (ENSMUST00000168172.3/107607_ intron 1 of 11) |

| | | | | |
|---|---|---|---|---|
| Tgif1 | -5.353884814 | 6.92E-05 | 0.005812199 | Intron (ENSMUST00000133983.7/224997_ intron 8 of 13) |
| Mief1 | 5.36425874 | 6.94E-05 | 0.005812199 | Distal Intergenic |
| Bcas3 | 5.627493721 | 6.96E-05 | 0.005812199 | Promoter (<=1kb) |
| Platr4 | 2.986513726 | 6.96E-05 | 0.005812199 | Distal Intergenic |
| Ap3b1 | 5.639581999 | 7.04E-05 | 0.00584519 | Exon (ENSMUST00000022196.4/11774_ exon 8 of 27) |
| Slc41a2 | -5.742224125 | 7.05E-05 | 0.00584519 | Distal Intergenic |
| Klf9 | 3.439261644 | 7.06E-05 | 0.00584519 | Intron (ENSMUST00000236056.1/ENSMUST00000236056.1_ intron 1 of 1) |
| Sema3f | 5.309302057 | 7.09E-05 | 0.00585931 | Intron (ENSMUST00000080560.8/20350_ intron 2 of 17) |
| Stard5 | 4.876826479 | 7.19E-05 | 0.005924034 | 5' UTR |
| Unc13d | 5.411478042 | 7.24E-05 | 0.005956042 | Promoter (<=1kb) |
| Snap25 | -3.872656403 | 7.26E-05 | 0.005956313 | Distal Intergenic |
| Gm17746 | 5.994575365 | 7.32E-05 | 0.005977806 | Exon (ENSMUST00000222804.1/ENSMUST00000222804.1_ exon 2 of 2) |
| Tcim | -5.63231326 | 7.32E-05 | 0.005977806 | Distal Intergenic |
| Lamc2 | -5.354994172 | 7.35E-05 | 0.005983045 | 3' UTR |
| Chd9 | 5.480307692 | 7.41E-05 | 0.006016812 | Intron (ENSMUST00000048665.7/109151_ intron 2 of 39) |
| Mir7034 | -5.687039057 | 7.52E-05 | 0.00609108 | Intron (ENSMUST00000153500.7/18984_ intron 2 of 7) |
| Ggta1 | 5.55215849 | 7.56E-05 | 0.006093298 | Promoter (<=1kb) |
| Znrf2 | 5.887547073 | 7.56E-05 | 0.006093298 | Distal Intergenic |
| Tmem17 | 5.872647023 | 7.58E-05 | 0.006093298 | Distal Intergenic |
| Ccdc158 | -5.659783531 | 7.65E-05 | 0.006135564 | Promoter (<=1kb) |
| Tns1 | 5.726427765 | 7.77E-05 | 0.00621528 | Exon (ENSMUST00000191264.6/21961_ exon 1 of 7) |
| AU019990 | 5.877530621 | 7.84E-05 | 0.006249875 | Promoter (<=1kb) |
| Notch1 | 5.615637293 | 7.85E-05 | 0.006249875 | Intron (ENSMUST00000132820.1/18128_ intron 2 of 3) |
| Dtnbp1 | 5.552333172 | 7.87E-05 | 0.006249875 | Distal Intergenic |
| 1810013A23Rik | 5.850989607 | 8.16E-05 | 0.00646634 | Intron (ENSMUST00000002320.15/19015_ intron 2 of 7) |
| Tbl1xr1 | -5.607976755 | 8.26E-05 | 0.006533947 | Distal Intergenic |
| Apobec3 | 3.610785328 | 8.30E-05 | 0.006549185 | Exon (ENSMUST00000229056.1/ENSMUST00000229056.1_ exon 1 of 1) |
| Tubb4b | -4.620673204 | 8.32E-05 | 0.006549185 | 3' UTR |
| S100a4 | -5.172774998 | 8.39E-05 | 0.006587626 | Promoter (1-2kb) |
| Slc7a8 | 5.734755152 | 8.45E-05 | 0.006618323 | Promoter (<=1kb) |
| Gm45521 | 5.875958506 | 8.50E-05 | 0.006641723 | Exon (ENSMUST00000052678.8/286940_ exon 19 of 46) |
| Lgals1 | 4.008131883 | 8.55E-05 | 0.006658433 | Promoter (<=1kb) |
| Myof | 5.284019353 | 8.56E-05 | 0.006658433 | Exon (ENSMUST00000172095.2/226101_ exon 19 of 53) |
| Axl | 5.633147486 | 8.65E-05 | 0.00670861 | Intron (ENSMUST00000002677.10/26362_ intron 4 of 19) |
| 1700020N01Rik | -5.812529361 | 8.81E-05 | 0.006821087 | Intron (ENSMUST00000057341.5/67692_ intron 2 of 2) |
| Thumpd3 | -5.520325931 | 8.86E-05 | 0.006842785 | Distal Intergenic |
| Gm19276 | 5.560184306 | 8.92E-05 | 0.006874924 | Intron (ENSMUST00000180604.1/75259_ intron 5 of 6) |
| Rfx4 | 5.493598096 | 9.24E-05 | 0.00707513 | Intron (ENSMUST00000077175.6/70428_ intron 23 of 27) |
| Timp3 | 5.245703705 | 9.25E-05 | 0.00707513 | Promoter (<=1kb) |
| 9430078K24Rik | 5.473959831 | 9.25E-05 | 0.00707513 | Exon (ENSMUST00000171210.2/19363_ exon 9 of 11) |
| Arrdc4 | 3.262337674 | 9.29E-05 | 0.00707513 | Promoter (<=1kb) |
| Zeb2os | 2.295337981 | 9.32E-05 | 0.00707513 | Promoter (<=1kb) |
| Smim38 | 5.711608523 | 9.32E-05 | 0.00707513 | Distal Intergenic |
| Ptpn1 | 3.263697256 | 9.33E-05 | 0.00707513 | Exon (ENSMUST00000147210.7/19246_ exon 4 of 7) |
| Ncoa3 | 4.108820172 | 9.41E-05 | 0.007112783 | Intron (ENSMUST00000146963.1/ENSMUST00000146963.1_ intron 2 of 2) |
| AI463170 | 5.818211255 | 9.53E-05 | 0.007187544 | Intron (ENSMUST00000021458.12/20741_ intron 32 of 35) |
| Bmp8b | 5.481171815 | 9.56E-05 | 0.007195198 | Distal Intergenic |
| Zdhhc7 | 5.46427386 | 9.61E-05 | 0.007218838 | Intron (ENSMUST00000127664.1/108148_ intron 1 of 15) |
| 9430078K24Rik | -5.394732332 | 9.69E-05 | 0.007242923 | Promoter (<=1kb) |
| 2610035D17Rik | 5.46260581 | 9.69E-05 | 0.007242923 | Intron (ENSMUST00000145206.1/ENSMUST00000145206.1_ intron 1 of 1) |
| Golt1a | 5.784219603 | 9.74E-05 | 0.007266196 | Promoter (<=1kb) |
| Rlbp1 | 5.293887407 | 9.79E-05 | 0.007285068 | Intron (ENSMUST00000037315.12/54608_ intron 5 of 10) |
| Apol7a | 5.806131371 | 9.95E-05 | 0.007385819 | Promoter (<=1kb) |
| Prkci | -5.419625804 | 0.000100024 | 0.0073953 | Intron (ENSMUST00000108249.8/18759_ intron 1 of 17) |
| Thbd | 5.425179302 | 0.00010016 | 0.0073953 | Promoter (<=1kb) |
| Rbpj | 5.510611827 | 0.000100258 | 0.0073953 | Intron (ENSMUST00000201991.3/19664_ intron 1 of 6) |

| | | | | |
|---|---|---|---|---|
| Surf6 | 5.446735467 | 0.000100934 | 0.007428528 | 3' UTR |
| Gask1b | 5.53900338 | 0.00010139 | 0.007445486 | Promoter (<=1kb) |
| Zfp532 | 5.553879685 | 0.000102093 | 0.007480469 | Intron (ENSMUST00000235822.1/328977_ intron 4 of 10) |
| Pbx1 | 5.595302871 | 0.000102515 | 0.007484519 | Intron (ENSMUST00000188912.6/18514_ intron 2 of 7) |
| Sash1 | 5.563426713 | 0.000102602 | 0.007484519 | Intron (ENSMUST00000015449.5/70097_ intron 8 of 19) |
| Cep68 | -5.592378419 | 0.000103643 | 0.007522668 | Intron (ENSMUST00000050611.13/216543_ intron 4 of 6) |
| Hesx1 | 5.680453709 | 0.000104048 | 0.007522668 | Promoter (<=1kb) |
| Uox | 5.77610085 | 0.000104195 | 0.007522668 | Distal Intergenic |
| Gm15713 | -5.774674893 | 0.000104593 | 0.007522668 | 5' UTR |
| Csrp2 | 5.704855309 | 0.00010461 | 0.007522668 | Intron (ENSMUST00000020403.6/13008_ intron 1 of 5) |
| Pdxk | 5.426493847 | 0.000104708 | 0.007522668 | Exon (ENSMUST00000217736.1/ENSMUST00000217736.1_ exon 1 of 1) |
| Hdac4 | -5.712365418 | 0.000104888 | 0.007522668 | Intron (ENSMUST00000097644.8/208727_ intron 3 of 26) |
| Aebp1 | 5.62552726 | 0.00010495 | 0.007522668 | Promoter (<=1kb) |
| Smim1 | 5.343162111 | 0.000106474 | 0.007615331 | Promoter (<=1kb) |
| Gm13205 | -5.550658144 | 0.000107103 | 0.007633593 | Intron (ENSMUST00000103217.10/56273_ intron 3 of 8) |
| Actn4 | 3.238614636 | 0.000107193 | 0.007633593 | Exon (ENSMUST00000208229.1/ENSMUST00000208229.1_ exon 2 of 2) |
| Pzp | -5.756426222 | 0.000107639 | 0.007648856 | Exon (ENSMUST00000060574.8/232400_ exon 25 of 36) |
| Hdac4 | 5.497676842 | 0.000108174 | 0.007650789 | Intron (ENSMUST00000097644.8/208727_ intron 3 of 26) |
| Mir1961 | 5.4899272 | 0.00010828 | 0.007650789 | Intron (ENSMUST00000113055.8/27428_ intron 2 of 10) |
| Barx2 | -5.571869096 | 0.000108362 | 0.007650789 | Intron (ENSMUST00000116615.4/12023_ intron 2 of 3) |
| Smurf2 | -5.502580057 | 0.000108924 | 0.007674052 | Intron (ENSMUST00000103067.9/66313_ intron 1 of 17) |
| Cadm1 | 5.515925968 | 0.000109732 | 0.007714465 | Intron (ENSMUST00000114548.7/54725_ intron 1 of 9) |
| 4930554I06Rik | 5.607936489 | 0.000111948 | 0.007841966 | Intron (ENSMUST00000237231.1/ENSMUST00000237231.1_ intron 2 of 4) |
| Fabp12 | 5.49665265 | 0.000112021 | 0.007841966 | Intron (ENSMUST00000029043.12/75497_ intron 1 of 5) |
| Mboat2 | -5.518096014 | 0.000112454 | 0.007842109 | Intron (ENSMUST00000222198.1/67216_ intron 3 of 4) |
| Dram1 | 5.555850055 | 0.000112499 | 0.007842109 | Intron (ENSMUST00000156097.1/71712_ intron 1 of 6) |
| Speer4f1 | 5.587238598 | 0.000113379 | 0.00788681 | Distal Intergenic |
| Emg1 | -5.550032987 | 0.000113644 | 0.007888538 | 3' UTR |
| Grhl2 | 5.224618526 | 0.000114128 | 0.007905494 | Distal Intergenic |
| Car8 | 5.57334633 | 0.00011473 | 0.007924685 | Intron (ENSMUST00000066674.7/12319_ intron 3 of 8) |
| Sdc3 | 3.084968308 | 0.000114885 | 0.007924685 | Promoter (<=1kb) |
| Pgbd5 | 5.422524862 | 0.00011543 | 0.007945643 | Exon (ENSMUST00000127664.1/108148_ exon 15 of 16) |
| Gm30790 | 5.652567849 | 0.000116327 | 0.007990678 | Intron (ENSMUST00000208702.1/ENSMUST00000208702.1_ intron 1 of 1) |
| Shb | 4.072023901 | 0.000116953 | 0.008003713 | Intron (ENSMUST00000147926.7/230126_ intron 1 of 4) |
| Fkbp5 | -5.224838724 | 0.000117168 | 0.008003713 | Exon (ENSMUST00000232983.1/ENSMUST00000232983.1_ exon 1 of 3) |
| Slc27a2 | -5.6129506 | 0.000117245 | 0.008003713 | Intron (ENSMUST00000061491.13/26458_ intron 8 of 9) |
| Snora3 | -5.48973658 | 0.000119062 | 0.008110985 | Intron (ENSMUST00000079282.7/76954_ intron 2 of 19) |
| Sp1 | -5.503822347 | 0.000120883 | 0.008212224 | Distal Intergenic |
| Dnah7a | -5.65880439 | 0.000121046 | 0.008212224 | Distal Intergenic |
| Gm16325 | -5.542787678 | 0.000123462 | 0.008358911 | 5' UTR |
| Hs3st1 | 5.366130107 | 0.000124155 | 0.008372387 | Intron (ENSMUST00000117944.1/15476_ intron 2 of 2) |
| Lrrc74b | 5.635010896 | 0.000124169 | 0.008372387 | Promoter (<=1kb) |
| A530013C23Rik | 5.625327151 | 0.000124519 | 0.008378834 | Distal Intergenic |
| Tgfb3 | 5.621919424 | 0.000124942 | 0.008390181 | Intron (ENSMUST00000040179.13/320244_ intron 30 of 33) |
| Slc45a3 | 5.194025925 | 0.000125858 | 0.008433469 | Distal Intergenic |
| Map3k14 | 5.344046641 | 0.000126098 | 0.008433469 | Distal Intergenic |
| Hdgfl3 | -6.198924129 | 0.000127578 | 0.008515192 | Promoter (<=1kb) |
| Micos10 | 5.551402286 | 0.000128546 | 0.008560642 | Distal Intergenic |
| Hdac7 | 5.544084717 | 0.000128837 | 0.008560642 | Intron (ENSMUST00000116408.8/56233_ intron 1 of 23) |
| C1qtnf9 | 5.580567903 | 0.000129038 | 0.008560642 | Intron (ENSMUST00000160973.7/219140_ intron 4 of 12) |
| Snap23 | -5.494957771 | 0.000130257 | 0.008624167 | Intron (ENSMUST00000110711.8/20619_ intron 1 of 7) |
| Rnh1 | 5.499204833 | 0.000131397 | 0.008648468 | Exon (ENSMUST00000210314.1/107702_ exon 3 of 10) |

### Table S6: Differential H3K27ac DKO_EE vs DKO Top500

| Symbol | Fold | p.value | FDR | annotation |
|---|---|---|---|---|
| Arrb1 | 5.97490255 | 2.03E-17 | 6.70E-13 | Promoter (<=1kb) |
| Runx1 | 6.887607649 | 2.27E-16 | 3.74E-12 | 5' UTR |
| Gsn | 4.33292507 | 7.64E-16 | 8.40E-12 | 5' UTR |
| Plin3 | 7.52421921 | 3.50E-14 | 2.88E-10 | Promoter (<=1kb) |
| Fhod1 | 10.55725389 | 1.68E-13 | 1.11E-09 | Promoter (<=1kb) |
| Baiap2 | 7.913412833 | 3.75E-13 | 2.06E-09 | Intron (ENSMUST00000075180.11/108100_ intron 3 of 13) |
| Lad1 | 4.845917481 | 4.72E-13 | 2.22E-09 | Promoter (<=1kb) |
| Gm49890 | 3.436922038 | 5.46E-13 | 2.25E-09 | Promoter (<=1kb) |
| Rbm47 | 4.477047255 | 6.36E-13 | 2.33E-09 | 5' UTR |
| Fam3d | 4.218994506 | 7.48E-13 | 2.47E-09 | Promoter (<=1kb) |
| Slc22a16 | 10.03936344 | 1.11E-12 | 3.19E-09 | Distal Intergenic |
| 2610035D17Rik | 10.14046965 | 1.16E-12 | 3.19E-09 | Intron (ENSMUST00000136438.7/72386_ intron 1 of 2) |
| Mmp15 | 5.002505782 | 1.40E-12 | 3.55E-09 | Promoter (<=1kb) |
| Mbp | 3.391960334 | 4.52E-12 | 1.06E-08 | 3' UTR |
| Nxn | 6.238384983 | 5.30E-12 | 1.17E-08 | Intron (ENSMUST00000021204.3/18230_ intron 1 of 7) |
| Gm45521 | 9.785770308 | 8.54E-12 | 1.76E-08 | Promoter (<=1kb) |
| Ccdc85c | 9.832807193 | 1.05E-11 | 2.04E-08 | Intron (ENSMUST00000222310.1/668158_ intron 1 of 5) |
| Tacstd2 | 7.313318472 | 1.44E-11 | 2.42E-08 | Promoter (<=1kb) |
| Mlph | 4.248920281 | 1.46E-11 | 2.42E-08 | 5' UTR |
| Fut4 | 9.751428394 | 1.53E-11 | 2.42E-08 | Promoter (<=1kb) |
| Arrdc4 | 5.724601419 | 1.54E-11 | 2.42E-08 | Promoter (<=1kb) |
| Adgrl2 | 10.12100158 | 1.73E-11 | 2.46E-08 | Promoter (<=1kb) |
| Lif | 9.497877794 | 1.75E-11 | 2.46E-08 | Promoter (1-2kb) |
| Pmepa1 | 10.37310074 | 1.89E-11 | 2.46E-08 | Promoter (<=1kb) |
| Tinagl1 | 9.765276066 | 1.95E-11 | 2.46E-08 | Promoter (<=1kb) |
| Artn | 9.581673438 | 1.98E-11 | 2.46E-08 | Exon (ENSMUST00000146982.1/ENSMUST00000146982.1_ exon 1 of 3) |
| Rhov | 5.843486532 | 2.08E-11 | 2.46E-08 | Promoter (<=1kb) |
| Actn4 | 5.753999942 | 2.24E-11 | 2.46E-08 | Exon (ENSMUST00000208229.1/ENSMUST00000208229.1_ exon 2 of 2) |
| Mir3971 | 10.21008297 | 2.28E-11 | 2.46E-08 | 3' UTR |
| Ltf | 6.605462783 | 2.30E-11 | 2.46E-08 | Distal Intergenic |
| Trim14 | 9.877662051 | 2.36E-11 | 2.46E-08 | Intron (ENSMUST00000030021.13/107684_ intron 3 of 12) |
| Shb | 9.406566119 | 2.39E-11 | 2.46E-08 | Exon (ENSMUST00000158267.1/ENSMUST00000158267.1_ exon 1 of 1) |
| Kcnn4 | 9.506159036 | 2.47E-11 | 2.47E-08 | Promoter (<=1kb) |
| 0610040F04Rik | 9.222817941 | 2.98E-11 | 2.87E-08 | Distal Intergenic |
| Zfp648 | 9.64582689 | 3.05E-11 | 2.87E-08 | Exon (ENSMUST00000185465.6/ENSMUST00000185465.6_ exon 1 of 3) |
| Agmat | 6.0218493 | 3.21E-11 | 2.94E-08 | Exon (ENSMUST00000038014.10/214063_ exon 10 of 15) |
| Edn1 | 7.661636412 | 3.64E-11 | 3.16E-08 | Promoter (<=1kb) |
| Kcnk16 | 9.523761012 | 3.64E-11 | 3.16E-08 | Distal Intergenic |
| 4933424L21Rik | 5.747579182 | 4.16E-11 | 3.52E-08 | Intron (ENSMUST00000066987.13/171212_ intron 1 of 11) |
| Gm19461 | 9.718716722 | 4.48E-11 | 3.70E-08 | Intron (ENSMUST00000212252.1/240753_ intron 3 of 9) |
| Hacd2 | 9.562015952 | 5.06E-11 | 4.07E-08 | Intron (ENSMUST00000061156.9/70757_ intron 2 of 6) |
| Synpo | 7.903700626 | 5.37E-11 | 4.21E-08 | Exon (ENSMUST00000237783.1/15531_ exon 8 of 15) |
| Cblc | 9.280048351 | 6.56E-11 | 5.03E-08 | Promoter (<=1kb) |
| Csf3 | 9.212281773 | 6.81E-11 | 5.11E-08 | Promoter (2-3kb) |
| Cpd | 9.469905836 | 7.08E-11 | 5.18E-08 | Intron (ENSMUST00000021201.5/12874_ intron 2 of 20) |
| Mroh4 | 6.152380186 | 7.80E-11 | 5.59E-08 | Exon (ENSMUST00000137963.7/69439_ exon 8 of 24) |
| Adamts1 | 9.175771324 | 8.51E-11 | 5.97E-08 | Promoter (<=1kb) |
| 1110002J07Rik | 6.087390435 | 8.70E-11 | 5.98E-08 | Promoter (<=1kb) |
| Ceacam1 | 8.988260513 | 9.16E-11 | 6.17E-08 | 3' UTR |
| 2610035D17Rik | 9.273837783 | 1.06E-10 | 6.91E-08 | Intron (ENSMUST00000136438.7/72386_ intron 1 of 2) |
| Tatdn2 | 9.201099575 | 1.07E-10 | 6.91E-08 | 3' UTR |

| | | | | |
|---|---|---|---|---|
| Cebpd | 6.254611722 | 1.19E-10 | 7.47E-08 | Intron (ENSMUST00000040248.8/224008_ intron 10 of 19) |
| Myo1e | 9.179718676 | 1.20E-10 | 7.47E-08 | Intron (ENSMUST00000213863.1/71602_ intron 1 of 3) |
| Spns2 | 9.132501555 | 1.25E-10 | 7.62E-08 | Promoter (<=1kb) |
| Gm28856 | 9.167054821 | 1.36E-10 | 8.13E-08 | Promoter (<=1kb) |
| Bak1 | 8.972300863 | 1.47E-10 | 8.66E-08 | 3' UTR |
| Epas1 | 9.003932669 | 1.54E-10 | 8.91E-08 | Promoter (<=1kb) |
| Rab4a | 9.150069876 | 1.59E-10 | 8.97E-08 | Intron (ENSMUST00000127664.1/108148_ intron 1 of 15) |
| Pik3ip1 | 5.079441415 | 1.60E-10 | 8.97E-08 | Exon (ENSMUST00000138907.7/216505_ exon 4 of 5) |
| Ly6d | 9.203205804 | 1.72E-10 | 9.32E-08 | Promoter (<=1kb) |
| Rab11fip1 | 3.764740029 | 1.74E-10 | 9.32E-08 | Exon (ENSMUST00000211028.1/ENSMUST00000211028.1_ exon 3 of 3) |
| Szrd1 | 9.456347892 | 1.75E-10 | 9.32E-08 | Intron (ENSMUST00000148204.7/213491_ intron 1 of 3) |
| Fhdc1 | 9.427087433 | 1.91E-10 | 9.99E-08 | Distal Intergenic |
| Mir6995 | 8.96656123 | 2.02E-10 | 1.04E-07 | 5' UTR |
| Cox4i2 | 8.99330417 | 2.18E-10 | 1.11E-07 | 3' UTR |
| Rgs3 | 3.892010568 | 2.32E-10 | 1.16E-07 | Exon (ENSMUST00000084521.10/50780_ exon 11 of 16) |
| Ikzf2 | 8.88811545 | 2.56E-10 | 1.25E-07 | Intron (ENSMUST00000027146.8/22779_ intron 3 of 7) |
| Sspn | 9.015531921 | 2.58E-10 | 1.25E-07 | Exon (ENSMUST00000203815.1/ENSMUST00000203815.1_ exon 1 of 1) |
| Tnk2os | 9.083087614 | 2.75E-10 | 1.31E-07 | Promoter (2-3kb) |
| Baiap2 | 9.077293492 | 2.89E-10 | 1.35E-07 | Exon (ENSMUST00000075180.11/108100_ exon 2 of 14) |
| Hoxb6 | 9.69861135 | 2.90E-10 | 1.35E-07 | Promoter (<=1kb) |
| Sprtn | 9.102057854 | 2.94E-10 | 1.35E-07 | Intron (ENSMUST00000034469.6/112405_ intron 1 of 4) |
| Anxa9 | 9.040840322 | 3.03E-10 | 1.36E-07 | 3' UTR |
| Pxn | 9.017448686 | 3.05E-10 | 1.36E-07 | Exon (ENSMUST00000125007.7/19303_ exon 1 of 11) |
| Lrp1 | 9.118224864 | 3.27E-10 | 1.44E-07 | 3' UTR |
| Coro2a | 8.927932809 | 3.56E-10 | 1.53E-07 | Intron (ENSMUST00000107756.3/107684_ intron 1 of 11) |
| Isg20 | 4.068784811 | 3.58E-10 | 1.53E-07 | Promoter (<=1kb) |
| Morf4l1-ps1 | 8.766088225 | 3.64E-10 | 1.54E-07 | Intron (ENSMUST00000231352.1/210126_ intron 3 of 6) |
| Nabp1 | 8.886419558 | 3.84E-10 | 1.60E-07 | Exon (ENSMUST00000194442.1/ENSMUST00000194442.1_ exon 1 of 1) |
| 5830428M24Rik | 8.814025711 | 4.20E-10 | 1.72E-07 | Distal Intergenic |
| Fasn | 9.409023198 | 4.23E-10 | 1.72E-07 | Intron (ENSMUST00000056781.4/71276_ intron 18 of 18) |
| Ankrd55 | 8.980266633 | 4.32E-10 | 1.73E-07 | Distal Intergenic |
| Inava | 9.135818714 | 4.36E-10 | 1.73E-07 | Promoter (<=1kb) |
| Smad3 | 8.744466984 | 4.90E-10 | 1.92E-07 | Intron (ENSMUST00000034973.9/17127_ intron 1 of 8) |
| Cemip2 | 6.297377574 | 5.01E-10 | 1.94E-07 | Exon (ENSMUST00000096194.8/83921_ exon 12 of 25) |
| Rara | 2.88691778 | 5.16E-10 | 1.98E-07 | Promoter (2-3kb) |
| Mllt1 | 7.213923831 | 5.35E-10 | 2.01E-07 | Exon (ENSMUST00000025053.9/64144_ exon 4 of 12) |
| 2210408F21Rik | 8.923680579 | 5.37E-10 | 2.01E-07 | Exon (ENSMUST00000123149.7/73652_ exon 3 of 7) |
| Grhl2 | 8.972237415 | 5.49E-10 | 2.02E-07 | Distal Intergenic |
| Gm12992 | 8.999785694 | 5.51E-10 | 2.02E-07 | Promoter (<=1kb) |
| Ap5z1 | 8.90975474 | 5.63E-10 | 2.04E-07 | Intron (ENSMUST00000072837.6/17425_ intron 2 of 8) |
| Pde2a | 9.035953456 | 5.88E-10 | 2.11E-07 | 5' UTR |
| Plekhs1 | 9.158488524 | 6.03E-10 | 2.14E-07 | Promoter (<=1kb) |
| Efnb2 | 8.903455332 | 6.34E-10 | 2.22E-07 | Distal Intergenic |
| Akr1d1 | 8.968331191 | 6.38E-10 | 2.22E-07 | Distal Intergenic |
| Tnfsf10 | 8.90221258 | 6.94E-10 | 2.38E-07 | Promoter (2-3kb) |
| Ccnd1 | 8.778732941 | 7.00E-10 | 2.38E-07 | Promoter (<=1kb) |
| Fam83h | 9.124059748 | 7.09E-10 | 2.39E-07 | 3' UTR |
| Abcc3 | 8.769093187 | 7.67E-10 | 2.55E-07 | Promoter (<=1kb) |
| Wwc1 | 9.082460894 | 8.10E-10 | 2.67E-07 | Exon (ENSMUST00000018993.6/211652_ exon 2 of 23) |
| Basp1 | 8.791426493 | 8.18E-10 | 2.67E-07 | Distal Intergenic |
| Ak1 | 8.948700462 | 8.53E-10 | 2.76E-07 | 5' UTR |
| Slc25a51 | 9.154718016 | 9.67E-10 | 3.10E-07 | 3' UTR |

| | | | | |
|---|---|---|---|---|
| Rac1 | 8.807551449 | 1.05E-09 | 3.33E-07 | Exon (ENSMUST00000080537.13/19353_ exon 3 of 6) |
| Arl4d | 5.663297329 | 1.10E-09 | 3.46E-07 | Distal Intergenic |
| Ikzf2 | 8.739106415 | 1.11E-09 | 3.47E-07 | Distal Intergenic |
| Klhl8 | 8.977958325 | 1.13E-09 | 3.47E-07 | Intron (ENSMUST00000054979.9/17355_ intron 7 of 19) |
| Cryaa | 8.716649616 | 1.14E-09 | 3.47E-07 | Distal Intergenic |
| Gm11681 | 6.550869017 | 1.15E-09 | 3.47E-07 | Distal Intergenic |
| 9430078K24Rik | 8.789161592 | 1.18E-09 | 3.55E-07 | Distal Intergenic |
| Niban2 | 9.078094093 | 1.20E-09 | 3.56E-07 | Exon (ENSMUST00000154514.7/227737_ exon 1 of 6) |
| Tent5c | 9.136270283 | 1.24E-09 | 3.64E-07 | Distal Intergenic |
| Slc45a3 | 8.801372484 | 1.26E-09 | 3.66E-07 | Distal Intergenic |
| Enpp6 | 8.619895683 | 1.27E-09 | 3.66E-07 | Distal Intergenic |
| Snx10 | 8.919329792 | 1.30E-09 | 3.72E-07 | 5' UTR |
| Adh6a | 9.083078543 | 1.33E-09 | 3.79E-07 | Exon (ENSMUST00000160080.1/ENSMUST00000160080.1_ exon 1 of 3) |
| Gsta1 | 8.901460573 | 1.45E-09 | 4.09E-07 | Exon (ENSMUST00000216513.1/ENSMUST00000216513.1_ exon 1 of 1) |
| Ezr | 8.784114778 | 1.46E-09 | 4.09E-07 | Exon (ENSMUST00000064234.6/22350_ exon 7 of 13) |
| Tob2 | 9.43499208 | 1.52E-09 | 4.19E-07 | Distal Intergenic |
| Sik1 | 8.74417352 | 1.53E-09 | 4.19E-07 | Distal Intergenic |
| Mb | 9.079877143 | 1.55E-09 | 4.20E-07 | 3' UTR |
| Il12a | 8.366894158 | 1.55E-09 | 4.20E-07 | 5' UTR |
| Srp9 | 8.921217046 | 1.57E-09 | 4.21E-07 | Distal Intergenic |
| 2210408F21Rik | 8.889783612 | 1.58E-09 | 4.21E-07 | Intron (ENSMUST00000144612.7/73652_ intron 1 of 5) |
| Thumpd3 | 8.543351378 | 1.62E-09 | 4.27E-07 | Distal Intergenic |
| Mlph | 8.790799224 | 1.66E-09 | 4.35E-07 | Promoter (<=1kb) |
| Trim14 | 9.177658047 | 1.72E-09 | 4.46E-07 | 5' UTR |
| Tnks1bp1 | 3.479321135 | 1.74E-09 | 4.46E-07 | Promoter (<=1kb) |
| Dap | 8.476218119 | 1.74E-09 | 4.46E-07 | Distal Intergenic |
| Or2w25 | 8.579454167 | 1.76E-09 | 4.46E-07 | Distal Intergenic |
| Plekhg6 | 8.532488092 | 1.83E-09 | 4.61E-07 | Distal Intergenic |
| Ehf | 8.64505198 | 1.87E-09 | 4.63E-07 | Exon (ENSMUST00000090475.9/13661_ exon 4 of 9) |
| Prr13 | 3.240385042 | 1.87E-09 | 4.63E-07 | Promoter (<=1kb) |
| Rgs3 | 8.696536236 | 2.03E-09 | 4.99E-07 | 5' UTR |
| Mir3076 | 8.700387734 | 2.06E-09 | 4.99E-07 | 3' UTR |
| 1110028F18Rik | 9.353574804 | 2.06E-09 | 4.99E-07 | Promoter (<=1kb) |
| Gm15417 | 8.581090412 | 2.16E-09 | 5.21E-07 | 5' UTR |
| Ccdc97 | 3.226571803 | 2.18E-09 | 5.21E-07 | 3' UTR |
| Synj2 | 5.60494429 | 2.28E-09 | 5.40E-07 | Intron (ENSMUST00000115791.9/20975_ intron 1 of 26) |
| Pdlim2 | 8.397035936 | 2.36E-09 | 5.55E-07 | Promoter (<=1kb) |
| Slc8b1 | 8.400627436 | 2.49E-09 | 5.78E-07 | Promoter (1-2kb) |
| Zbtb16 | 8.543565434 | 2.49E-09 | 5.78E-07 | Intron (ENSMUST00000093852.4/235320_ intron 2 of 6) |
| Gmppb | 6.19357723 | 2.57E-09 | 5.92E-07 | Intron (ENSMUST00000176454.1/27399_ intron 8 of 10) |
| Unc5b | 8.744218146 | 2.61E-09 | 5.97E-07 | Promoter (<=1kb) |
| Lmod3 | 3.180171906 | 2.71E-09 | 6.16E-07 | 3' UTR |
| Foxq1 | 8.420271613 | 2.76E-09 | 6.23E-07 | Promoter (<=1kb) |
| Ppp1r14b | 8.717747115 | 2.84E-09 | 6.35E-07 | 5' UTR |
| Rrp1b | 8.484582022 | 2.87E-09 | 6.35E-07 | Intron (ENSMUST00000002145.11/74377_ intron 7 of 7) |
| Gadd45g | 3.947897871 | 2.91E-09 | 6.35E-07 | Distal Intergenic |
| Eml1 | 8.822354579 | 2.92E-09 | 6.35E-07 | Promoter (<=1kb) |
| Denn2b | 8.384501696 | 2.93E-09 | 6.35E-07 | Intron (ENSMUST00000207745.1/76954_ intron 1 of 3) |
| Myof | 8.704450683 | 2.96E-09 | 6.35E-07 | Intron (ENSMUST00000172095.2/226101_ intron 3 of 52) |
| Podxl | 5.236869838 | 2.96E-09 | 6.35E-07 | 3' UTR |
| Gstt3 | 8.831362423 | 2.97E-09 | 6.35E-07 | Promoter (<=1kb) |
| Ecscr | 3.630121878 | 3.08E-09 | 6.55E-07 | Promoter (<=1kb) |

| | | | | |
|---|---|---|---|---|
| Dhrs3 | 8.675763395 | 3.16E-09 | 6.69E-07 | 3' UTR |
| Arl14ep | 8.802269316 | 3.21E-09 | 6.73E-07 | Intron (ENSMUST00000111063.7/77015_ intron 4 of 6) |
| Klf9 | 5.218051357 | 3.24E-09 | 6.77E-07 | Intron (ENSMUST00000236056.1/ENSMUST00000236056.1_ intron 1 of 1) |
| A530013C23Rik | 8.659169524 | 3.32E-09 | 6.87E-07 | Distal Intergenic |
| 1700020M21Rik | 8.700924142 | 3.34E-09 | 6.87E-07 | Distal Intergenic |
| Lyzl4 | 9.006776401 | 3.37E-09 | 6.87E-07 | Intron (ENSMUST00000077706.9/69032_ intron 1 of 5) |
| Slc39a4 | 8.599480895 | 3.38E-09 | 6.87E-07 | Promoter (<=1kb) |
| Ywhaz | 4.916977525 | 3.49E-09 | 6.99E-07 | Exon (ENSMUST00000226620.1/ENSMUST00000226620.1_ exon 1 of 2) |
| Slc5a9 | 8.651562066 | 3.49E-09 | 6.99E-07 | Promoter (<=1kb) |
| Capza1b | 5.253019274 | 3.50E-09 | 6.99E-07 | 5' UTR |
| Anpep | 8.492094048 | 3.54E-09 | 7.03E-07 | Intron (ENSMUST00000075657.7/11778_ intron 4 of 5) |
| E130008D07Rik | 8.391944711 | 3.56E-09 | 7.03E-07 | Distal Intergenic |
| Actn4 | 8.59257373 | 3.64E-09 | 7.15E-07 | 3' UTR |
| Myo5c | 8.750105463 | 3.70E-09 | 7.23E-07 | 3' UTR |
| Mir5122 | 8.520187924 | 3.99E-09 | 7.74E-07 | Intron (ENSMUST00000030669.7/20544_ intron 1 of 11) |
| Ddah2 | 5.797528041 | 4.05E-09 | 7.77E-07 | Promoter (<=1kb) |
| Mir1901 | 3.727487023 | 4.05E-09 | 7.77E-07 | Intron (ENSMUST00000171109.8/63955_ intron 1 of 9) |
| Runx1 | 8.265114002 | 4.17E-09 | 7.94E-07 | 3' UTR |
| Pan3 | 8.796389419 | 4.38E-09 | 8.31E-07 | Distal Intergenic |
| Ccdc25 | 8.565289294 | 4.45E-09 | 8.38E-07 | Distal Intergenic |
| Stk39 | 8.61517135 | 4.80E-09 | 8.95E-07 | Intron (ENSMUST00000102715.3/53416_ intron 1 of 17) |
| Mir29a | 8.213305905 | 4.81E-09 | 8.95E-07 | Promoter (<=1kb) |
| Myo5c | 8.88730993 | 4.97E-09 | 9.21E-07 | 3' UTR |
| Slc45a1 | 3.25601976 | 5.17E-09 | 9.52E-07 | Distal Intergenic |
| 9430078K24Rik | 8.414186595 | 5.35E-09 | 9.81E-07 | Intron (ENSMUST00000171210.2/19363_ intron 7 of 10) |
| A930015D03Rik | 8.65515273 | 5.41E-09 | 9.85E-07 | Promoter (2-3kb) |
| Mecom | 9.044015174 | 5.71E-09 | 1.03E-06 | 5' UTR |
| Mir1195 | 5.75387208 | 5.74E-09 | 1.03E-06 | Intron (ENSMUST00000180468.1/ENSMUST00000180468.1_ intron 1 of 2) |
| Acer3 | 8.295293524 | 5.80E-09 | 1.04E-06 | Distal Intergenic |
| Surf6 | 8.755775255 | 5.89E-09 | 1.05E-06 | 3' UTR |
| Gm13648 | 8.648584389 | 6.00E-09 | 1.06E-06 | Distal Intergenic |
| Dapl1 | 8.412865096 | 6.01E-09 | 1.06E-06 | Distal Intergenic |
| Arrdc1 | 8.84039791 | 6.03E-09 | 1.06E-06 | 3' UTR |
| Mir3097 | 8.581218323 | 6.08E-09 | 1.06E-06 | 5' UTR |
| Map1lc3a | 8.833818708 | 6.22E-09 | 1.08E-06 | Exon (ENSMUST00000139854.1/228812_ exon 2 of 4) |
| Get4 | 8.384868065 | 6.26E-09 | 1.08E-06 | Promoter (2-3kb) |
| Cab39l | 8.80597948 | 6.32E-09 | 1.09E-06 | 5' UTR |
| Apobec3 | 5.465548818 | 6.42E-09 | 1.10E-06 | Exon (ENSMUST00000229056.1/ENSMUST00000229056.1_ exon 1 of 1) |
| Nek7 | 8.37110415 | 6.47E-09 | 1.10E-06 | Intron (ENSMUST00000186017.6/59125_ intron 1 of 9) |
| Cx3cl1 | 8.509683389 | 6.53E-09 | 1.10E-06 | Distal Intergenic |
| Ampd3 | 8.611988959 | 6.65E-09 | 1.11E-06 | Promoter (<=1kb) |
| Isg20 | 8.478028481 | 6.66E-09 | 1.11E-06 | Exon (ENSMUST00000206812.1/ENSMUST00000206812.1_ exon 1 of 1) |
| B230217C12Rik | 8.287654159 | 6.80E-09 | 1.13E-06 | Intron (ENSMUST00000043843.11/16796_ intron 4 of 6) |
| Tnnt2 | 8.482581583 | 6.93E-09 | 1.15E-06 | Exon (ENSMUST00000188028.6/21956_ exon 5 of 16) |
| Arhgef10l | 8.37632242 | 7.05E-09 | 1.16E-06 | Intron (ENSMUST00000039204.9/72754_ intron 1 of 27) |
| Ier5l | 8.142656263 | 7.05E-09 | 1.16E-06 | Distal Intergenic |
| Alox12 | 6.261165103 | 7.08E-09 | 1.16E-06 | Promoter (<=1kb) |
| Mir205 | 8.733044496 | 7.19E-09 | 1.17E-06 | Distal Intergenic |
| Ahnak | 6.002824714 | 7.31E-09 | 1.18E-06 | Promoter (<=1kb) |
| Tjp1 | 8.198618358 | 7.39E-09 | 1.19E-06 | Intron (ENSMUST00000206228.1/21872_ intron 2 of 4) |
| 4833427F10Rik | 8.306807879 | 7.59E-09 | 1.21E-06 | Promoter (<=1kb) |
| 9430078K24Rik | 8.495857339 | 7.60E-09 | 1.21E-06 | Distal Intergenic |

| | | | | |
|---|---|---|---|---|
| Gm15880 | 8.487754 | 7.64E-09 | 1.21E-06 | Promoter (<=1kb) |
| Myof | 6.636958168 | 7.89E-09 | 1.25E-06 | Promoter (<=1kb) |
| Scamp1 | 5.760160026 | 7.97E-09 | 1.25E-06 | 5' UTR |
| Il15 | 5.685976651 | 8.14E-09 | 1.27E-06 | Promoter (<=1kb) |
| Plce1 | 8.375466368 | 8.16E-09 | 1.27E-06 | Distal Intergenic |
| Soat2 | 8.574838728 | 8.19E-09 | 1.27E-06 | Distal Intergenic |
| Gm21057 | 8.620167963 | 8.23E-09 | 1.27E-06 | Promoter (<=1kb) |
| Mir7041 | 8.724644069 | 8.27E-09 | 1.27E-06 | Intron (ENSMUST00000204645.2/16206_ intron 3 of 19) |
| Spred2 | 5.361356505 | 8.32E-09 | 1.27E-06 | Intron (ENSMUST00000093299.12/114716_ intron 1 of 6) |
| Coro2a | 8.535723908 | 8.71E-09 | 1.32E-06 | Intron (ENSMUST00000107756.3/107684_ intron 1 of 11) |
| Pip5k1c | 8.418265399 | 8.78E-09 | 1.33E-06 | Intron (ENSMUST00000163075.7/18717_ intron 1 of 16) |
| Gm7550 | 8.321828538 | 8.81E-09 | 1.33E-06 | Distal Intergenic |
| 4930578N18Rik | 2.960111431 | 8.88E-09 | 1.33E-06 | Distal Intergenic |
| Ido2 | 8.023905134 | 8.89E-09 | 1.33E-06 | Promoter (<=1kb) |
| Ccnd3 | 3.268165313 | 9.18E-09 | 1.36E-06 | Promoter (<=1kb) |
| Map3k13 | 8.625724886 | 9.35E-09 | 1.38E-06 | 5' UTR |
| 2200002D01Rik | 8.359784555 | 9.38E-09 | 1.38E-06 | Promoter (<=1kb) |
| Tomm20 | 8.300720262 | 9.40E-09 | 1.38E-06 | Distal Intergenic |
| Pstpip1 | 8.293564874 | 9.49E-09 | 1.38E-06 | Promoter (<=1kb) |
| Herpud1 | 8.489101788 | 9.57E-09 | 1.39E-06 | 3' UTR |
| Spdef | 8.683420258 | 9.74E-09 | 1.41E-06 | Promoter (<=1kb) |
| Mink1 | 8.256355811 | 9.94E-09 | 1.43E-06 | Intron (ENSMUST00000072237.12/50932_ intron 1 of 32) |
| Paxbp1 | 8.232219823 | 1.02E-08 | 1.46E-06 | 5' UTR |
| Vcpkmt | 2.421663799 | 1.03E-08 | 1.48E-06 | Distal Intergenic |
| Krt78 | 8.429517119 | 1.06E-08 | 1.50E-06 | Distal Intergenic |
| Naglu | 8.442548907 | 1.06E-08 | 1.50E-06 | Exon (ENSMUST00000103110.9/11975_ exon 19 of 21) |
| Tagap1 | 4.009403647 | 1.08E-08 | 1.52E-06 | 5' UTR |
| Fxyd4 | 8.145735024 | 1.10E-08 | 1.54E-06 | Promoter (<=1kb) |
| Etfbkmt | 8.841489372 | 1.11E-08 | 1.55E-06 | 5' UTR |
| Fkbp5 | 8.312219859 | 1.12E-08 | 1.56E-06 | Intron (ENSMUST00000079413.10/14229_ intron 1 of 10) |
| Gm12992 | 5.033973834 | 1.13E-08 | 1.56E-06 | Exon (ENSMUST00000124738.1/545681_ exon 2 of 5) |
| Ppp1r3b | 5.136770399 | 1.13E-08 | 1.56E-06 | Distal Intergenic |
| Gnai2 | 8.345022552 | 1.14E-08 | 1.56E-06 | Intron (ENSMUST00000192615.5/14678_ intron 1 of 8) |
| Mgst1 | 8.190357718 | 1.14E-08 | 1.56E-06 | Promoter (<=1kb) |
| Sema4b | 8.311868652 | 1.19E-08 | 1.63E-06 | Distal Intergenic |
| Plekha6 | 8.114086383 | 1.22E-08 | 1.65E-06 | Intron (ENSMUST00000212252.1/240753_ intron 3 of 9) |
| Shb | 6.24386818 | 1.23E-08 | 1.66E-06 | Intron (ENSMUST00000147926.7/230126_ intron 1 of 4) |
| Pdzrn3 | 8.390793375 | 1.23E-08 | 1.66E-06 | Distal Intergenic |
| Tcim | 8.572986535 | 1.26E-08 | 1.68E-06 | Exon (ENSMUST00000209183.1/ENSMUST00000209183.1_ exon 1 of 1) |
| Pdlim5 | 5.336101901 | 1.26E-08 | 1.68E-06 | Intron (ENSMUST00000029941.15/56376_ intron 3 of 12) |
| Fam3b | 8.57641834 | 1.27E-08 | 1.68E-06 | Intron (ENSMUST00000049721.8/52793_ intron 2 of 7) |
| Capzb | 8.111808783 | 1.27E-08 | 1.68E-06 | Intron (ENSMUST00000102508.9/12345_ intron 1 of 8) |
| Wwc2 | 4.958429933 | 1.30E-08 | 1.72E-06 | Promoter (<=1kb) |
| Bace2 | 8.419544167 | 1.31E-08 | 1.73E-06 | Intron (ENSMUST00000047275.7/56175_ intron 2 of 8) |
| Epha2 | 6.403320882 | 1.32E-08 | 1.73E-06 | Promoter (<=1kb) |
| Fndc3b | 5.054753633 | 1.37E-08 | 1.78E-06 | Intron (ENSMUST00000195008.5/72007_ intron 1 of 26) |
| Supt3 | 8.462345283 | 1.38E-08 | 1.78E-06 | Intron (ENSMUST00000162816.7/12393_ intron 5 of 5) |
| Chd9 | 6.468146193 | 1.38E-08 | 1.78E-06 | Exon (ENSMUST00000211215.1/109151_ exon 3 of 3) |
| Gm12596 | 2.868762446 | 1.39E-08 | 1.79E-06 | Distal Intergenic |
| Far1os | 8.414826307 | 1.40E-08 | 1.79E-06 | Exon (ENSMUST00000211320.1/ENSMUST00000211320.1_ exon 1 of 1) |
| Nfkbia | 3.786597414 | 1.42E-08 | 1.82E-06 | Distal Intergenic |
| Etv6 | 6.190452841 | 1.44E-08 | 1.83E-06 | Intron (ENSMUST00000081028.12/14011_ intron 1 of 7) |

| | | | | |
|---|---|---|---|---|
| Klf5 | 8.510806469 | 1.44E-08 | 1.83E-06 | Distal Intergenic |
| Furin | 8.469900792 | 1.49E-08 | 1.88E-06 | Exon (ENSMUST00000206518.1/12144_ exon 1 of 2) |
| Rnf38 | 8.299004534 | 1.50E-08 | 1.88E-06 | Promoter (<=1kb) |
| Fzd2 | 8.032869362 | 1.50E-08 | 1.88E-06 | Promoter (<=1kb) |
| Dtnbp1 | 8.508410319 | 1.51E-08 | 1.89E-06 | Distal Intergenic |
| Plcd3 | 8.487492026 | 1.58E-08 | 1.96E-06 | Intron (ENSMUST00000103077.1/72469_ intron 1 of 14) |
| Ly6g6c | 6.141977925 | 1.59E-08 | 1.97E-06 | Promoter (<=1kb) |
| Rassf3 | 8.25468344 | 1.60E-08 | 1.97E-06 | Intron (ENSMUST00000026902.8/192678_ intron 1 of 4) |
| Rxra | 8.481643059 | 1.64E-08 | 2.01E-06 | Intron (ENSMUST00000129514.7/20181_ intron 1 of 3) |
| 2210409D07Rik | 8.75603769 | 1.68E-08 | 2.06E-06 | Intron (ENSMUST00000079738.9/75444_ intron 6 of 6) |
| Psd4 | 8.351492181 | 1.69E-08 | 2.06E-06 | 5' UTR |
| Trmt5 | 4.776663884 | 1.70E-08 | 2.07E-06 | Intron (ENSMUST00000187549.6/17420_ intron 6 of 6) |
| Get1 | 8.176935799 | 1.72E-08 | 2.08E-06 | Promoter (2-3kb) |
| Cd2ap | 3.407738894 | 1.72E-08 | 2.08E-06 | Distal Intergenic |
| Naa20 | 3.562815955 | 1.75E-08 | 2.10E-06 | 3' UTR |
| Trit1 | 8.285005955 | 1.79E-08 | 2.14E-06 | Exon (ENSMUST00000102649.3/66966_ exon 4 of 11) |
| Plcd3 | 8.437003383 | 1.79E-08 | 2.14E-06 | Promoter (<=1kb) |
| Niban2 | 8.230700945 | 1.83E-08 | 2.17E-06 | Intron (ENSMUST00000028135.14/227737_ intron 1 of 13) |
| 0610040F04Rik | 4.586820785 | 1.83E-08 | 2.17E-06 | Distal Intergenic |
| 2810414N06Rik | 8.077593506 | 1.83E-08 | 2.17E-06 | Intron (ENSMUST00000148810.7/63913_ intron 5 of 13) |
| 9130015A21Rik | 8.008818725 | 1.85E-08 | 2.17E-06 | Exon (ENSMUST00000221491.1/71581_ exon 2 of 2) |
| Pcdh1 | 8.282190152 | 1.86E-08 | 2.18E-06 | Exon (ENSMUST00000161701.2/75599_ exon 2 of 4) |
| Anxa2 | 8.242025595 | 1.87E-08 | 2.18E-06 | Distal Intergenic |
| Gm17546 | 7.90532654 | 1.90E-08 | 2.21E-06 | Exon (ENSMUST00000168960.2/102634389_ exon 2 of 2) |
| Kctd10 | 8.083907767 | 1.93E-08 | 2.24E-06 | Exon (ENSMUST00000102581.10/330171_ exon 2 of 7) |
| Rgs3 | 8.548091468 | 1.95E-08 | 2.25E-06 | Exon (ENSMUST00000084521.10/50780_ exon 11 of 16) |
| Abcc4 | 8.233652101 | 1.95E-08 | 2.25E-06 | Intron (ENSMUST00000036554.13/239273_ intron 3 of 30) |
| 9430078K24Rik | 5.204691915 | 1.99E-08 | 2.29E-06 | Intron (ENSMUST00000171210.2/19363_ intron 8 of 10) |
| Mir1941 | 8.003309339 | 2.08E-08 | 2.39E-06 | Intron (ENSMUST00000229584.1/ENSMUST00000229584.1_ intron 1 of 4) |
| Fbxw7 | 8.237661232 | 2.14E-08 | 2.44E-06 | Intron (ENSMUST00000154148.7/99889_ intron 3 of 9) |
| Wbp1l | 4.209786853 | 2.20E-08 | 2.50E-06 | Intron (ENSMUST00000138302.8/226178_ intron 1 of 3) |
| E230032D23Rik | 4.49684453 | 2.23E-08 | 2.52E-06 | 5' UTR |
| Car9 | 8.464088924 | 2.23E-08 | 2.52E-06 | Promoter (<=1kb) |
| Zzef1 | 8.369357702 | 2.25E-08 | 2.53E-06 | Exon (ENSMUST00000150551.7/195018_ exon 10 of 13) |
| Yy1 | 8.172191093 | 2.27E-08 | 2.54E-06 | Distal Intergenic |
| Ccl6 | 8.183106139 | 2.27E-08 | 2.54E-06 | Promoter (2-3kb) |
| Cryaa | 6.943629183 | 2.30E-08 | 2.56E-06 | Distal Intergenic |
| Platr4 | 4.226563788 | 2.31E-08 | 2.57E-06 | Distal Intergenic |
| Mir5122 | 8.069333101 | 2.33E-08 | 2.58E-06 | Intron (ENSMUST00000030669.7/20544_ intron 1 of 11) |
| 9130015G15Rik | 8.589275161 | 2.34E-08 | 2.58E-06 | Intron (ENSMUST00000183659.7/78286_ intron 2 of 39) |
| Prkch | 5.482815868 | 2.36E-08 | 2.60E-06 | Intron (ENSMUST00000221153.1/18755_ intron 2 of 14) |
| Tpm2 | 7.840414442 | 2.37E-08 | 2.60E-06 | Promoter (<=1kb) |
| Pola2 | 8.270113308 | 2.40E-08 | 2.62E-06 | Exon (ENSMUST00000025752.14/18969_ exon 5 of 18) |
| Pbp2 | 8.287727138 | 2.45E-08 | 2.67E-06 | Intron (ENSMUST00000203686.1/ENSMUST00000203686.1_ intron 2 of 3) |
| Wnt4 | 8.256671086 | 2.59E-08 | 2.81E-06 | Promoter (<=1kb) |
| Sphk1 | 8.009304713 | 2.61E-08 | 2.82E-06 | Promoter (<=1kb) |
| Mpzl3 | 5.110749116 | 2.67E-08 | 2.87E-06 | Promoter (2-3kb) |
| Ate1 | 8.602730034 | 2.67E-08 | 2.87E-06 | 5' UTR |
| Sgk1 | 8.040488025 | 2.71E-08 | 2.90E-06 | Distal Intergenic |
| Pik3cb | 8.316037681 | 2.73E-08 | 2.92E-06 | Distal Intergenic |
| Mir1961 | 7.952676794 | 2.80E-08 | 2.98E-06 | Intron (ENSMUST00000113055.8/27428_ intron 2 of 10) |
| Capzb | 7.859036976 | 2.81E-08 | 2.98E-06 | Intron (ENSMUST00000102508.9/12345_ intron 2 of 8) |

| | | | | |
|---|---|---|---|---|
| Gm2176 | 8.249269091 | 2.89E-08 | 3.05E-06 | Distal Intergenic |
| 5033406O09Rik | 5.7586269 | 2.93E-08 | 3.08E-06 | Distal Intergenic |
| Fam220a | 7.903208998 | 2.94E-08 | 3.09E-06 | Promoter (1-2kb) |
| Ppp2ca | 8.912136232 | 2.99E-08 | 3.13E-06 | Intron (ENSMUST00000121591.7/213084_ intron 13 of 13) |
| Clic6 | 8.33510913 | 3.11E-08 | 3.24E-06 | Exon (ENSMUST00000180989.3/ENSMUST00000180989.3_ exon 1 of 5) |
| Sh3pxd2a | 8.218970472 | 3.13E-08 | 3.25E-06 | Intron (ENSMUST00000081619.9/14218_ intron 1 of 14) |
| 9030204H09Rik | 8.0110307 | 3.17E-08 | 3.29E-06 | Intron (ENSMUST00000112986.8/69601_ intron 3 of 16) |
| Cilk1 | 8.146486859 | 3.22E-08 | 3.33E-06 | Intron (ENSMUST00000118869.7/56542_ intron 1 of 13) |
| Ifnlr1 | 8.076241374 | 3.24E-08 | 3.34E-06 | Distal Intergenic |
| Fkbp5 | 8.309812326 | 3.32E-08 | 3.41E-06 | 5' UTR |
| A730020M07Rik | 8.165531112 | 3.34E-08 | 3.41E-06 | Distal Intergenic |
| Mrpl33 | 8.053457723 | 3.35E-08 | 3.41E-06 | Promoter (1-2kb) |
| Tsen2 | 8.537197585 | 3.35E-08 | 3.41E-06 | Intron (ENSMUST00000171644.7/19016_ intron 7 of 7) |
| Mnt | 8.043246297 | 3.39E-08 | 3.44E-06 | Promoter (2-3kb) |
| Plb1 | 7.446550589 | 3.40E-08 | 3.44E-06 | Distal Intergenic |
| Tril | 8.598556129 | 3.47E-08 | 3.50E-06 | Promoter (<=1kb) |
| Src | 7.996736971 | 3.54E-08 | 3.56E-06 | 3' UTR |
| Pyroxd1 | 8.285814342 | 3.59E-08 | 3.60E-06 | Distal Intergenic |
| Rab4a | 8.24193288 | 3.62E-08 | 3.61E-06 | Intron (ENSMUST00000127664.1/108148_ intron 1 of 15) |
| Igfbp7 | 7.858810728 | 3.62E-08 | 3.61E-06 | Intron (ENSMUST00000046746.9/29817_ intron 1 of 4) |
| Sh2d4b | 8.07212517 | 3.68E-08 | 3.65E-06 | Distal Intergenic |
| Gm4651 | 8.133875834 | 3.80E-08 | 3.74E-06 | Intron (ENSMUST00000207889.1/194401_ intron 1 of 31) |
| Inpp5k | 8.415308015 | 3.80E-08 | 3.74E-06 | 3' UTR |
| Rras2 | 8.348949131 | 3.80E-08 | 3.74E-06 | 3' UTR |
| Laptm4a | 7.746207952 | 3.83E-08 | 3.76E-06 | Distal Intergenic |
| Mir5104 | 8.162189584 | 3.88E-08 | 3.80E-06 | Promoter (1-2kb) |
| Cgnl1 | 8.183244875 | 3.97E-08 | 3.86E-06 | Promoter (<=1kb) |
| Slc39a14 | 7.880966384 | 3.97E-08 | 3.86E-06 | 5' UTR |
| Barx2 | 8.373345329 | 3.98E-08 | 3.86E-06 | Distal Intergenic |
| E330017L17Rik | 8.218705495 | 4.03E-08 | 3.90E-06 | Distal Intergenic |
| Kat6b | 4.92812203 | 4.13E-08 | 3.98E-06 | Intron (ENSMUST00000183201.7/54169_ intron 2 of 5) |
| Hibadh | 8.150561607 | 4.17E-08 | 4.00E-06 | Promoter (<=1kb) |
| Atosa | 8.293647578 | 4.19E-08 | 4.02E-06 | Distal Intergenic |
| Kif21b | 8.162664518 | 4.21E-08 | 4.02E-06 | Promoter (1-2kb) |
| Txn2 | 7.993292674 | 4.25E-08 | 4.02E-06 | 3' UTR |
| Trp63 | 8.241171266 | 4.25E-08 | 4.02E-06 | 5' UTR |
| Egln3 | 2.156365986 | 4.26E-08 | 4.02E-06 | Intron (ENSMUST00000039516.3/112407_ intron 1 of 4) |
| Spef1l | 7.839726241 | 4.27E-08 | 4.02E-06 | 3' UTR |
| Mir721 | 7.876382065 | 4.27E-08 | 4.02E-06 | Intron (ENSMUST00000176216.8/13047_ intron 2 of 22) |
| D030018L15Rik | 8.378711416 | 4.43E-08 | 4.16E-06 | Distal Intergenic |
| Pgbd5 | 8.114345497 | 4.44E-08 | 4.16E-06 | 3' UTR |
| Rgmb | 8.082521181 | 4.52E-08 | 4.22E-06 | Intron (ENSMUST00000170578.2/68799_ intron 2 of 2) |
| Zmiz1 | 8.031353904 | 4.58E-08 | 4.27E-06 | Intron (ENSMUST00000225188.1/ENSMUST00000225188.1_ intron 2 of 3) |
| Bcl6 | 2.601533829 | 4.61E-08 | 4.28E-06 | Promoter (<=1kb) |
| Chd9 | 8.117590577 | 4.65E-08 | 4.31E-06 | Intron (ENSMUST00000048665.7/109151_ intron 2 of 39) |
| Cgnl1 | 7.906544633 | 4.69E-08 | 4.33E-06 | Intron (ENSMUST00000072899.8/68178_ intron 8 of 18) |
| Akap13 | 7.940572189 | 4.70E-08 | 4.33E-06 | Exon (ENSMUST00000207998.1/75547_ exon 1 of 9) |
| Gpx2 | 7.81252809 | 4.73E-08 | 4.34E-06 | 5' UTR |
| Pgbd5 | 8.265066671 | 4.82E-08 | 4.41E-06 | Exon (ENSMUST00000127664.1/108148_ exon 15 of 16) |
| Tbc1d16 | 8.43077553 | 4.84E-08 | 4.42E-06 | Intron (ENSMUST00000036113.3/207592_ intron 3 of 11) |
| Tgfa | 3.297131426 | 4.96E-08 | 4.52E-06 | Intron (ENSMUST00000032066.12/21802_ intron 1 of 5) |
| Tagln2 | 8.213684618 | 4.97E-08 | 4.52E-06 | Exon (ENSMUST00000135267.7/93842_ exon 4 of 6) |

| | | | | |
|---|---|---|---|---|
| Slc29a2 | 8.162382416 | 5.06E-08 | 4.58E-06 | Distal Intergenic |
| Ifitm10 | 7.670318106 | 5.13E-08 | 4.63E-06 | Promoter (<=1kb) |
| Dtnbp1 | 8.136419025 | 5.14E-08 | 4.63E-06 | Exon (ENSMUST00000221315.1/ENSMUST00000221315.1_ exon 1 of 2) |
| Mief1 | 7.739739245 | 5.16E-08 | 4.64E-06 | Distal Intergenic |
| Fam53b | 6.676977416 | 5.18E-08 | 4.64E-06 | Intron (ENSMUST00000124096.7/14183_ intron 2 of 6) |
| Nln | 8.077593524 | 5.20E-08 | 4.65E-06 | Intron (ENSMUST00000109315.4/75805_ intron 12 of 12) |
| Sorbs2 | 5.184832426 | 5.30E-08 | 4.72E-06 | 5' UTR |
| Slc25a10 | 5.000864676 | 5.31E-08 | 4.72E-06 | Distal Intergenic |
| Htra3 | 7.986011206 | 5.45E-08 | 4.83E-06 | 5' UTR |
| Gm36229 | 5.585305078 | 5.55E-08 | 4.91E-06 | Distal Intergenic |
| Cryaa | 7.780172808 | 5.58E-08 | 4.92E-06 | Distal Intergenic |
| Zc3h4 | 7.945567336 | 5.73E-08 | 5.02E-06 | Exon (ENSMUST00000214735.1/330474_ exon 2 of 2) |
| Niban2 | 8.109348568 | 5.74E-08 | 5.02E-06 | Distal Intergenic |
| Mir7661 | 7.923715024 | 5.75E-08 | 5.02E-06 | Intron (ENSMUST00000032192.8/16438_ intron 46 of 61) |
| 4933411K16Rik | 8.053692574 | 5.75E-08 | 5.02E-06 | Intron (ENSMUST00000172244.7/67432_ intron 1 of 2) |
| Fchsd2 | 8.238823536 | 5.99E-08 | 5.21E-06 | Exon (ENSMUST00000207053.1/ENSMUST00000207053.1_ exon 1 of 1) |
| 2310001H17Rik | 3.149834786 | 6.10E-08 | 5.29E-06 | Exon (ENSMUST00000181238.1/76432_ exon 2 of 3) |
| Soat2 | 7.939294761 | 6.13E-08 | 5.30E-06 | Promoter (<=1kb) |
| Slc28a3 | 8.026393433 | 6.14E-08 | 5.30E-06 | Intron (ENSMUST00000022036.13/114304_ intron 1 of 17) |
| 4930556N09Rik | 8.496849038 | 6.15E-08 | 5.30E-06 | Intron (ENSMUST00000219918.1/ENSMUST00000219918.1_ intron 4 of 4) |
| Mir29b-1 | 8.703584346 | 6.18E-08 | 5.30E-06 | Intron (ENSMUST00000229711.1/232685_ intron 2 of 3) |
| Mroh3 | 8.193703677 | 6.24E-08 | 5.35E-06 | Exon (ENSMUST00000180496.1/ENSMUST00000180496.1_ exon 2 of 2) |
| Gm5820 | 6.676658826 | 6.26E-08 | 5.35E-06 | 3' UTR |
| AU022754 | 8.154303699 | 6.51E-08 | 5.55E-06 | Promoter (<=1kb) |
| Tent5c | 3.382376214 | 6.54E-08 | 5.56E-06 | Promoter (<=1kb) |
| Zfp691 | 7.804129089 | 6.68E-08 | 5.66E-06 | Distal Intergenic |
| Marf1 | 7.86000215 | 6.77E-08 | 5.72E-06 | Intron (ENSMUST00000023359.12/67203_ intron 4 of 8) |
| 6330415G19Rik | 8.359264284 | 6.91E-08 | 5.83E-06 | Distal Intergenic |
| Mbp | 7.639519048 | 7.11E-08 | 5.96E-06 | Intron (ENSMUST00000091789.10/17196_ intron 2 of 3) |
| Apobec1 | 7.947935735 | 7.12E-08 | 5.96E-06 | Promoter (<=1kb) |
| Ncoa3 | 5.860481602 | 7.12E-08 | 5.96E-06 | Intron (ENSMUST00000146963.1/ENSMUST00000146963.1_ intron 2 of 2) |
| S1pr2 | 7.714966331 | 7.42E-08 | 6.20E-06 | Promoter (2-3kb) |
| Stxbp6 | 7.780446703 | 7.45E-08 | 6.21E-06 | Promoter (<=1kb) |
| Sirt2 | 5.571369956 | 7.50E-08 | 6.23E-06 | Promoter (<=1kb) |
| Ehd4 | 7.810951091 | 7.56E-08 | 6.26E-06 | Exon (ENSMUST00000028755.7/98878_ exon 3 of 6) |
| Mrgpre | 3.811073722 | 7.74E-08 | 6.39E-06 | 3' UTR |
| Tlr6 | 6.191902892 | 7.89E-08 | 6.51E-06 | Promoter (<=1kb) |
| Mir7661 | 8.011152376 | 7.95E-08 | 6.54E-06 | Intron (ENSMUST00000032192.8/16438_ intron 60 of 61) |
| Efna5 | 8.14207025 | 8.03E-08 | 6.59E-06 | Intron (ENSMUST00000232942.1/ENSMUST00000232942.1_ intron 3 of 4) |
| 1700021F02Rik | 4.105218855 | 8.17E-08 | 6.68E-06 | Distal Intergenic |
| Gm45521 | 7.743756075 | 8.20E-08 | 6.69E-06 | Distal Intergenic |
| Sipa1l3 | 7.822036277 | 8.29E-08 | 6.75E-06 | Intron (ENSMUST00000085809.10/74206_ intron 3 of 22) |
| Tex2 | 7.669432982 | 8.42E-08 | 6.84E-06 | Exon (ENSMUST00000184424.1/ENSMUST00000184424.1_ exon 1 of 1) |
| Susd1 | 7.792455365 | 8.57E-08 | 6.94E-06 | Intron (ENSMUST00000040166.13/634731_ intron 8 of 16) |
| Ephb4 | 7.915558818 | 8.60E-08 | 6.94E-06 | Promoter (<=1kb) |
| P2ry2 | 5.180654962 | 8.61E-08 | 6.94E-06 | Promoter (<=1kb) |
| Rassf5 | 8.146245638 | 8.75E-08 | 7.03E-06 | Intron (ENSMUST00000027688.14/54354_ intron 1 of 5) |
| Rxra | 8.172624095 | 8.82E-08 | 7.08E-06 | Intron (ENSMUST00000129514.7/20181_ intron 1 of 3) |
| Pik3r1 | 7.824016976 | 8.97E-08 | 7.18E-06 | Intron (ENSMUST00000055518.12/18708_ intron 2 of 15) |
| Krt78 | 3.618961145 | 9.07E-08 | 7.24E-06 | Distal Intergenic |
| Mvp | 3.856887893 | 9.16E-08 | 7.28E-06 | Promoter (<=1kb) |
| Gm30173 | 7.907139351 | 9.16E-08 | 7.28E-06 | Distal Intergenic |

| | | | | |
|---|---|---|---|---|
| Glod4 | 7.603174238 | 9.21E-08 | 7.30E-06 | Intron (ENSMUST00000021204.3/18230_ intron 1 of 7) |
| A930002C04Rik | 3.444385968 | 9.24E-08 | 7.31E-06 | Intron (ENSMUST00000068891.11/218215_ intron 2 of 7) |
| Rlbp1 | 7.706440577 | 9.27E-08 | 7.31E-06 | Intron (ENSMUST00000037315.12/54608_ intron 5 of 10) |
| Zbtb16 | 8.569774909 | 9.30E-08 | 7.31E-06 | Intron (ENSMUST00000093852.4/235320_ intron 3 of 6) |
| Heatr6 | 7.912158133 | 9.43E-08 | 7.40E-06 | Distal Intergenic |
| Evi2b | 8.212444155 | 9.47E-08 | 7.42E-06 | Promoter (<=1kb) |
| Cpeb4 | 8.090310793 | 9.59E-08 | 7.49E-06 | Distal Intergenic |
| Suco | 8.354752237 | 9.63E-08 | 7.50E-06 | Distal Intergenic |
| Cluh | 3.974656938 | 9.69E-08 | 7.53E-06 | 3' UTR |
| Kbtbd11 | 8.453718745 | 9.72E-08 | 7.54E-06 | Intron (ENSMUST00000084207.11/234094_ intron 26 of 28) |
| Tcp11 | 8.683576373 | 9.86E-08 | 7.61E-06 | Intron (ENSMUST00000114842.8/224650_ intron 12 of 24) |
| Tmem238l | 7.987899611 | 9.87E-08 | 7.61E-06 | Promoter (<=1kb) |
| Gm28856 | 7.958770797 | 9.88E-08 | 7.61E-06 | Intron (ENSMUST00000189150.1/433347_ intron 2 of 3) |
| Apobec1 | 8.016486884 | 9.91E-08 | 7.62E-06 | 5' UTR |
| Chmp3 | 7.949158763 | 1.00E-07 | 7.68E-06 | Intron (ENSMUST00000059462.11/66700_ intron 2 of 5) |
| Pde9a | 7.909701038 | 1.01E-07 | 7.76E-06 | 5' UTR |
| 1110002J07Rik | 8.092959447 | 1.03E-07 | 7.87E-06 | Distal Intergenic |
| Agtpbp1 | 8.270677933 | 1.06E-07 | 8.08E-06 | 3' UTR |
| Tent5c | 8.194340511 | 1.07E-07 | 8.12E-06 | Distal Intergenic |
| Dcaf4 | 8.367489854 | 1.08E-07 | 8.15E-06 | 5' UTR |
| Cp | 3.662489116 | 1.08E-07 | 8.17E-06 | Promoter (<=1kb) |
| Mapkapk3 | 8.014717244 | 1.08E-07 | 8.17E-06 | 5' UTR |
| Pou2af3 | 8.00326668 | 1.09E-07 | 8.21E-06 | Distal Intergenic |
| Bcas3 | 8.013838561 | 1.09E-07 | 8.22E-06 | Promoter (<=1kb) |
| Pvt1 | 7.759680941 | 1.10E-07 | 8.24E-06 | Intron (ENSMUST00000182956.7/19296_ intron 1 of 6) |
| Mir7661 | 4.844967718 | 1.10E-07 | 8.25E-06 | Intron (ENSMUST00000032192.8/16438_ intron 60 of 61) |
| Agr2 | 8.268863194 | 1.13E-07 | 8.39E-06 | Distal Intergenic |
| Ppif | 7.695232636 | 1.13E-07 | 8.39E-06 | Intron (ENSMUST00000162645.7/328365_ intron 6 of 23) |
| Mir6404 | 8.344402655 | 1.13E-07 | 8.39E-06 | Distal Intergenic |
| Fgfr1 | 4.597085821 | 1.13E-07 | 8.40E-06 | 5' UTR |
| Smu1 | 7.750835569 | 1.14E-07 | 8.46E-06 | Exon (ENSMUST00000121600.1/ENSMUST00000121600.1_ exon 1 of 1) |
| Gm45521 | 8.626512282 | 1.15E-07 | 8.46E-06 | Exon (ENSMUST00000052678.8/286940_ exon 19 of 46) |
| Irf2 | 8.280855484 | 1.17E-07 | 8.63E-06 | 5' UTR |
| 9430078K24Rik | 8.065267026 | 1.18E-07 | 8.65E-06 | Intron (ENSMUST00000171210.2/19363_ intron 8 of 10) |
| Vsig8 | 8.129857978 | 1.19E-07 | 8.69E-06 | 3' UTR |
| A230108P19Rik | 8.467653704 | 1.19E-07 | 8.71E-06 | Distal Intergenic |
| Naglu | 7.81101183 | 1.23E-07 | 8.96E-06 | Intron (ENSMUST00000103110.9/11975_ intron 18 of 20) |
| Rbms3 | 7.808290908 | 1.23E-07 | 8.98E-06 | Intron (ENSMUST00000111773.9/207181_ intron 3 of 14) |
| Snx29 | 7.983123965 | 1.24E-07 | 9.01E-06 | Intron (ENSMUST00000180792.7/74478_ intron 13 of 20) |
| Vopp1 | 6.265184929 | 1.25E-07 | 9.07E-06 | Intron (ENSMUST00000127485.7/232023_ intron 1 of 4) |
| Tc2n | 8.01296433 | 1.26E-07 | 9.10E-06 | Exon (ENSMUST00000021603.8/23876_ exon 9 of 11) |
| BB123696 | 7.592318761 | 1.27E-07 | 9.15E-06 | Distal Intergenic |
| Clu | 8.398122787 | 1.27E-07 | 9.17E-06 | Promoter (<=1kb) |
| Zbtb16 | 8.373955851 | 1.29E-07 | 9.29E-06 | Intron (ENSMUST00000093852.4/235320_ intron 2 of 6) |
| Ino80c | 7.693467647 | 1.31E-07 | 9.36E-06 | Exon (ENSMUST00000153360.7/225280_ exon 3 of 5) |
| 4933401H06Rik | 7.7967112 | 1.32E-07 | 9.44E-06 | Intron (ENSMUST00000081978.9/67547_ intron 3 of 8) |
| Adamtsl4 | 4.747869087 | 1.34E-07 | 9.59E-06 | Distal Intergenic |
| Tnfrsf1a | 7.86653052 | 1.36E-07 | 9.65E-06 | Intron (ENSMUST00000176655.7/20276_ intron 8 of 12) |
| Clstn1 | 3.955691145 | 1.39E-07 | 9.86E-06 | Exon (ENSMUST00000145263.1/ENSMUST00000145263.1_ exon 1 of 2) |
| Tnfrsf1a | 8.038279188 | 1.40E-07 | 9.91E-06 | Promoter (2-3kb) |
| Adcy6 | 8.095037455 | 1.40E-07 | 9.91E-06 | 5' UTR |
| Smco4 | 7.230551918 | 1.40E-07 | 9.91E-06 | 5' UTR |

| | | | | |
|---|---|---|---|---|
| Pik3ip1 | 7.789484621 | 1.43E-07 | 1.01E-05 | Exon (ENSMUST00000110029.8/16886_ exon 8 of 12) |
| 1700001K23Rik | 7.675445734 | 1.47E-07 | 1.03E-05 | Distal Intergenic |
| Srebf2 | 7.875100682 | 1.47E-07 | 1.03E-05 | Intron (ENSMUST00000023100.7/20788_ intron 1 of 18) |
| Mcc | 7.932739085 | 1.51E-07 | 1.06E-05 | Promoter (<=1kb) |
| Nrarp | 7.875696721 | 1.52E-07 | 1.06E-05 | Intron (ENSMUST00000238850.1/ENSMUST00000238850.1_ intron 13 of 13) |
| Sik1 | 8.276017996 | 1.55E-07 | 1.07E-05 | Exon (ENSMUST00000237005.1/ENSMUST00000237005.1_ exon 1 of 1) |
| Shisa5 | 2.481905601 | 1.55E-07 | 1.07E-05 | Promoter (<=1kb) |
| Cpd | 7.763669913 | 1.56E-07 | 1.08E-05 | Distal Intergenic |
| Itpr3 | 7.702115865 | 1.58E-07 | 1.09E-05 | Exon (ENSMUST00000236359.1/ENSMUST00000236359.1_ exon 1 of 2) |
| Cyp17a1 | 7.95394664 | 1.58E-07 | 1.09E-05 | 3' UTR |
| Baiap2 | 4.169169325 | 1.59E-07 | 1.09E-05 | Exon (ENSMUST00000143428.1/ENSMUST00000143428.1_ exon 1 of 2) |
| Ikzf1 | 8.118899658 | 1.59E-07 | 1.09E-05 | Promoter (2-3kb) |
| Hdac7 | 7.804732708 | 1.59E-07 | 1.09E-05 | Intron (ENSMUST00000079838.13/56233_ intron 1 of 24) |
| Qsox1 | 7.675660072 | 1.60E-07 | 1.09E-05 | Exon (ENSMUST00000140809.1/104009_ exon 2 of 2) |
| F630206G17Rik | 8.169003715 | 1.60E-07 | 1.09E-05 | Intron (ENSMUST00000137002.1/100009664_ intron 1 of 3) |
| Eya2 | 8.66136004 | 1.60E-07 | 1.09E-05 | Intron (ENSMUST00000088132.12/14049_ intron 1 of 15) |
| Gm4651 | 7.745922715 | 1.60E-07 | 1.09E-05 | Exon (ENSMUST00000212208.1/194401_ exon 6 of 11) |
| Eva1a | 7.60695752 | 1.62E-07 | 1.10E-05 | Promoter (<=1kb) |
| Polr2b | 7.587530795 | 1.64E-07 | 1.11E-05 | Intron (ENSMUST00000046746.9/29817_ intron 1 of 4) |
| Snx11 | 8.276765288 | 1.64E-07 | 1.11E-05 | 3' UTR |
| Timp3 | 7.424747925 | 1.64E-07 | 1.11E-05 | Promoter (<=1kb) |
| Itgb1 | 7.648129261 | 1.65E-07 | 1.11E-05 | Exon (ENSMUST00000212623.1/ENSMUST00000212623.1_ exon 1 of 1) |
| E230013L22Rik | 4.997768687 | 1.66E-07 | 1.12E-05 | Intron (ENSMUST00000033900.6/19332_ intron 1 of 1) |
| Iqgap1 | 7.512398847 | 1.66E-07 | 1.12E-05 | Promoter (<=1kb) |
| Card14 | 7.743696109 | 1.68E-07 | 1.12E-05 | 5' UTR |
| C330024C12Rik | 7.689786097 | 1.70E-07 | 1.13E-05 | Distal Intergenic |
| Sectm1a | 7.681622996 | 1.74E-07 | 1.16E-05 | Distal Intergenic |
| Smim1 | 7.683541123 | 1.78E-07 | 1.19E-05 | Promoter (<=1kb) |
| Tmem45b | 8.238794405 | 1.80E-07 | 1.20E-05 | Distal Intergenic |
| Nrp2 | 7.653522242 | 1.85E-07 | 1.22E-05 | Promoter (2-3kb) |
| Piezo1 | 2.850915069 | 1.85E-07 | 1.23E-05 | Promoter (<=1kb) |
| Tgfa | 7.643660494 | 1.86E-07 | 1.23E-05 | Distal Intergenic |

Table S7: DNA Methylation DMP DKO vs SKO Top500

| Gene | Feature | logFC | P.Value | adj.P.Val |
|---|---|---|---|---|
| LOC102635707 | tss_body | -1.47 | 7.58E-12 | 2.16E-06 |
| Fbln1 | tss_body | 2.07 | 1.78E-11 | 2.53E-06 |
| Fam207a | tss_body | 1.81 | 2.78E-11 | 2.63E-06 |
| NA | NA | -1.95 | 3.80E-11 | 2.64E-06 |
| Mup20 | tss_1500 | -2.2 | 5.14E-11 | 2.64E-06 |
| Gm14634 | tss_body | 1.92 | 5.57E-11 | 2.64E-06 |
| NA | NA | -2.41 | 6.90E-11 | 2.80E-06 |
| Ust | tss_body | 2.42 | 9.43E-11 | 3.30E-06 |
| Gm34998 | tss_body | -5.17 | 1.04E-10 | 3.30E-06 |
| Cblb | tss_body | 3.57 | 1.26E-10 | 3.31E-06 |
| Olfr166 | tss_body | -2.52 | 1.43E-10 | 3.31E-06 |
| Syna | tss_1500 | 2.17 | 1.59E-10 | 3.31E-06 |
| Vsig10 | tss_body | 1.72 | 1.69E-10 | 3.31E-06 |
| Ube2cbp | tss_body | 1.39 | 1.72E-10 | 3.31E-06 |
| LOC102635707 | tss_body | -1.39 | 1.74E-10 | 3.31E-06 |
| Tbc1d10a | tss_body | 1.25 | 1.92E-10 | 3.42E-06 |
| Mylk | tss_body | 1.87 | 2.07E-10 | 3.46E-06 |
| Gm34541 | tss_body | 1.78 | 2.30E-10 | 3.64E-06 |
| Limk2 | tss_body | 1.78 | 2.56E-10 | 3.72E-06 |
| Akr1b7 | tss_200 | 2.27 | 2.64E-10 | 3.72E-06 |
| NA | NA | 1.44 | 3.03E-10 | 3.72E-06 |
| Gm5084 | tss_200 | 1.78 | 3.06E-10 | 3.72E-06 |
| Ccl27a | tss_body | -1.47 | 3.08E-10 | 3.72E-06 |
| NA | NA | 2.21 | 3.14E-10 | 3.72E-06 |
| Gm5084 | tss_1500 | 2.61 | 3.53E-10 | 3.93E-06 |
| NA | NA | -2.65 | 3.85E-10 | 3.93E-06 |
| NA | NA | 1.47 | 3.96E-10 | 3.93E-06 |
| P2rx7 | tss_body | 1.8 | 4.01E-10 | 3.93E-06 |
| Nav1 | tss_body | 1.77 | 4.14E-10 | 3.93E-06 |
| NA | NA | 1.75 | 4.29E-10 | 3.93E-06 |
| Gm34719 | tss_body | -2.48 | 4.38E-10 | 3.93E-06 |
| NA | NA | 1.39 | 4.43E-10 | 3.93E-06 |
| NA | NA | -2.27 | 4.56E-10 | 3.93E-06 |
| NA | NA | 1.89 | 5.06E-10 | 4.08E-06 |
| NA | NA | 2.36 | 5.12E-10 | 4.08E-06 |
| Ccl27a | tss_body | -1.15 | 5.40E-10 | 4.08E-06 |
| Olfr1089 | tss_body | -1.89 | 5.44E-10 | 4.08E-06 |
| Prkd2 | tss_body | 1.66 | 5.44E-10 | 4.08E-06 |
| Mettl8 | tss_body | 1.69 | 5.73E-10 | 4.10E-06 |
| NA | tss_1500 | 1.8 | 5.76E-10 | 4.10E-06 |

| | | | | |
|---|---|---|---|---|
| NA | NA | 2.5 | 6.10E-10 | 4.22E-06 |
| Shisa6 | tss_body | 2.68 | 6.23E-10 | 4.22E-06 |
| Phlpp1 | tss_body | 1.33 | 6.85E-10 | 4.53E-06 |
| Slc25a13 | tss_body | 1.41 | 7.27E-10 | 4.70E-06 |
| NA | NA | -1.83 | 8.11E-10 | 5.04E-06 |
| NA | tss_1500 | -1.84 | 8.26E-10 | 5.04E-06 |
| AY761185 | tss_200 | -2.24 | 8.32E-10 | 5.04E-06 |
| Mpp5 | tss_body | 1.5 | 8.82E-10 | 5.23E-06 |
| Sipa1l3 | tss_body | 1.94 | 9.10E-10 | 5.28E-06 |
| Olfr1424 | tss_200 | -2.03 | 9.58E-10 | 5.31E-06 |
| Astn2 | tss_body | 1.44 | 9.73E-10 | 5.31E-06 |
| Nlrp9a | tss_body | -2.61 | 1.01E-09 | 5.31E-06 |
| NA | NA | -1.7 | 1.04E-09 | 5.31E-06 |
| LOC108168946 | tss_body | -2.07 | 1.05E-09 | 5.31E-06 |
| Rnf112 | tss_body | 1.67 | 1.12E-09 | 5.31E-06 |
| NA | NA | 1.08 | 1.13E-09 | 5.31E-06 |
| Tnfaip8 | tss_body | 2.32 | 1.14E-09 | 5.31E-06 |
| NA | NA | -1.51 | 1.16E-09 | 5.31E-06 |
| NA | tss_body | 2.09 | 1.17E-09 | 5.31E-06 |
| Cdon | tss_body | 1.79 | 1.17E-09 | 5.31E-06 |
| Cmss1 | tss_body | 2.13 | 1.18E-09 | 5.31E-06 |
| A430048G15Rik | tss_200 | 1.39 | 1.20E-09 | 5.31E-06 |
| Ociad2 | tss_body | 1.85 | 1.20E-09 | 5.31E-06 |
| Adh6a | tss_body | 1.58 | 1.21E-09 | 5.31E-06 |
| Dsc3 | tss_body | 3.39 | 1.24E-09 | 5.31E-06 |
| NA | NA | 1.52 | 1.25E-09 | 5.31E-06 |
| NA | tss_body | -2.58 | 1.26E-09 | 5.31E-06 |
| Mucl2 | tss_body | -2.57 | 1.28E-09 | 5.31E-06 |
| A0A494BAU3 | tss_1500 | 2.31 | 1.29E-09 | 5.31E-06 |
| Phyhip | tss_body | 2.87 | 1.33E-09 | 5.40E-06 |
| Bdnf | tss_body | 1.82 | 1.43E-09 | 5.60E-06 |
| Cecr2 | tss_body | -1.73 | 1.46E-09 | 5.60E-06 |
| NA | NA | -2.61 | 1.46E-09 | 5.60E-06 |
| Gm33594 | tss_body | 1.76 | 1.47E-09 | 5.60E-06 |
| Akr1b7 | tss_200 | 2.5 | 1.48E-09 | 5.60E-06 |
| Gm2163 | tss_body | -1.9 | 1.51E-09 | 5.66E-06 |
| NA | NA | 1.82 | 1.59E-09 | 5.84E-06 |
| Rian | tss_body | -2.39 | 1.60E-09 | 5.84E-06 |
| Mmrn2 | tss_body | 1.4 | 1.63E-09 | 5.87E-06 |
| Sptlc1 | tss_body | 1.72 | 1.72E-09 | 6.06E-06 |
| Dym | tss_body | 1.4 | 1.74E-09 | 6.06E-06 |

| | | | | |
|---|---|---|---|---|
| Gm31813 | tss_body | 1.88 | 1.76E-09 | 6.06E-06 |
| LOC105242408 | tss_body | 1.47 | 1.81E-09 | 6.06E-06 |
| NA | NA | -1.99 | 1.87E-09 | 6.06E-06 |
| Olfr703 | tss_1500 | -2.92 | 1.87E-09 | 6.06E-06 |
| Ksr1 | tss_body | 1.88 | 1.88E-09 | 6.06E-06 |
| Usp46 | tss_body | 1.95 | 1.88E-09 | 6.06E-06 |
| Fam205a4 | tss_body | -1.56 | 1.94E-09 | 6.06E-06 |
| Fndc3a | tss_body | -2.29 | 1.94E-09 | 6.06E-06 |
| Sez6l | tss_body | -2.1 | 1.96E-09 | 6.06E-06 |
| Nlrp4f | tss_body | -2.17 | 1.97E-09 | 6.06E-06 |
| LOC108169009 | tss_body | -2.26 | 2.04E-09 | 6.06E-06 |
| Rreb1 | tss_body | 1.48 | 2.17E-09 | 6.06E-06 |
| Vmn1r30 | tss_body | -1.47 | 2.17E-09 | 6.06E-06 |
| Camta1 | tss_body | 1.74 | 2.17E-09 | 6.06E-06 |
| NA | tss_body | -2.48 | 2.18E-09 | 6.06E-06 |
| Ext1 | tss_body | 1.94 | 2.23E-09 | 6.06E-06 |
| LOC102637912 | tss_body | -1.09 | 2.23E-09 | 6.06E-06 |
| Gm11677 | tss_body | -1.81 | 2.26E-09 | 6.06E-06 |
| Pcca | tss_body | 2.53 | 2.26E-09 | 6.06E-06 |
| Rian | tss_body | -2.53 | 2.27E-09 | 6.06E-06 |
| Olfr1513 | tss_body | -2.34 | 2.30E-09 | 6.06E-06 |
| LOC105242408 | tss_body | 2.54 | 2.34E-09 | 6.06E-06 |
| Olfr299 | tss_1500 | -2.31 | 2.39E-09 | 6.06E-06 |
| Msi2 | tss_body | 2.01 | 2.39E-09 | 6.06E-06 |
| Plekha6 | tss_body | 1.72 | 2.41E-09 | 6.06E-06 |
| Slc1a7 | tss_body | 1.54 | 2.42E-09 | 6.06E-06 |
| Gm34458 | tss_body | 1.68 | 2.42E-09 | 6.06E-06 |
| Lrrc32 | tss_1500 | 1.99 | 2.43E-09 | 6.06E-06 |
| Mir329 | tss_1500 | -1.51 | 2.44E-09 | 6.06E-06 |
| Gm12407 | tss_1500 | -1.61 | 2.46E-09 | 6.06E-06 |
| Ostm1 | tss_body | 1.54 | 2.46E-09 | 6.06E-06 |
| NA | NA | 1.65 | 2.47E-09 | 6.06E-06 |
| NA | NA | 1.83 | 2.53E-09 | 6.06E-06 |
| Gm32328 | tss_body | -1.57 | 2.53E-09 | 6.06E-06 |
| Kcnma1 | tss_body | -2.49 | 2.59E-09 | 6.06E-06 |
| Vti1a | tss_body | 1.2 | 2.59E-09 | 6.06E-06 |
| Creb5 | tss_body | -2.48 | 2.60E-09 | 6.06E-06 |
| Gria3 | tss_body | 1.8 | 2.60E-09 | 6.06E-06 |
| Skint7 | tss_body | -2.6 | 2.62E-09 | 6.06E-06 |
| Gm12407 | tss_1500 | -1.78 | 2.64E-09 | 6.06E-06 |
| Ccdc58 | tss_body | 1.3 | 2.66E-09 | 6.06E-06 |

| | | | | |
|---|---|---|---|---|
| Magi1 | tss_body | 2.35 | 2.67E-09 | 6.06E-06 |
| Olfr461 | tss_200 | -2.09 | 2.69E-09 | 6.06E-06 |
| Gm12407 | tss_1500 | -1.65 | 2.69E-09 | 6.06E-06 |
| NA | NA | -3.05 | 2.70E-09 | 6.06E-06 |
| Uaca | tss_body | 1.84 | 2.71E-09 | 6.06E-06 |
| Elmo1 | tss_body | -2.34 | 2.79E-09 | 6.20E-06 |
| Snap25 | tss_body | -1.95 | 2.92E-09 | 6.45E-06 |
| NA | NA | -1.33 | 3.01E-09 | 6.50E-06 |
| A2m | tss_1500 | -1.99 | 3.02E-09 | 6.50E-06 |
| LOC108169009 | tss_body | -2.28 | 3.08E-09 | 6.50E-06 |
| Gm36600 | tss_body | 1.47 | 3.10E-09 | 6.50E-06 |
| Dsc3 | tss_body | 3.4 | 3.10E-09 | 6.50E-06 |
| NA | tss_200 | -1.4 | 3.10E-09 | 6.50E-06 |
| NA | tss_body | -2.2 | 3.12E-09 | 6.50E-06 |
| NA | tss_body | -2.1 | 3.13E-09 | 6.50E-06 |
| Scn10a | tss_body | -1.49 | 3.24E-09 | 6.60E-06 |
| Tmem246 | tss_body | 2.09 | 3.27E-09 | 6.60E-06 |
| Gpt | tss_1500 | 1.13 | 3.27E-09 | 6.60E-06 |
| NA | NA | -1.94 | 3.28E-09 | 6.60E-06 |
| Dscam | tss_body | 1.81 | 3.30E-09 | 6.60E-06 |
| NA | NA | 2.66 | 3.32E-09 | 6.60E-06 |
| Atp13a5 | tss_body | 1.31 | 3.37E-09 | 6.66E-06 |
| A430048G15Rik | tss_200 | 1.49 | 3.41E-09 | 6.68E-06 |
| LOC108169009 | tss_body | -1.41 | 3.53E-09 | 6.88E-06 |
| Hnrnpll | tss_body | 1.65 | 3.56E-09 | 6.90E-06 |
| Rbm47 | tss_body | 1.78 | 3.61E-09 | 6.94E-06 |
| NA | NA | -2.46 | 3.66E-09 | 6.99E-06 |
| Fes | tss_body | 2.5 | 3.78E-09 | 7.06E-06 |
| NA | NA | 2.17 | 3.83E-09 | 7.06E-06 |
| Tmco5 | tss_body | -1.92 | 3.86E-09 | 7.06E-06 |
| Magi1 | tss_body | 1.53 | 3.89E-09 | 7.06E-06 |
| Adgrv1 | tss_body | 1.69 | 3.89E-09 | 7.06E-06 |
| Rabl2 | tss_body | 1.77 | 3.91E-09 | 7.06E-06 |
| Myo10 | tss_body | 1.76 | 3.93E-09 | 7.06E-06 |
| Rara | tss_body | 1.71 | 3.95E-09 | 7.06E-06 |
| NA | NA | 1.19 | 4.01E-09 | 7.06E-06 |
| Creb3l2 | tss_body | 1.78 | 4.01E-09 | 7.06E-06 |
| Cyp4a14 | tss_body | -2.21 | 4.03E-09 | 7.06E-06 |
| NA | NA | -1.86 | 4.04E-09 | 7.06E-06 |
| Cfhr2 | tss_1500 | -2.04 | 4.05E-09 | 7.06E-06 |
| Gsk3b | tss_body | 1.68 | 4.07E-09 | 7.06E-06 |

| | | | | |
|---|---|---|---|---|
| Mid2 | tss_body | 1.81 | 4.09E-09 | 7.06E-06 |
| LOC108169009 | tss_body | -1.42 | 4.11E-09 | 7.06E-06 |
| Fam65c | tss_body | 1.78 | 4.12E-09 | 7.06E-06 |
| NA | NA | 1.45 | 4.14E-09 | 7.06E-06 |
| Tg | tss_body | 2.46 | 4.17E-09 | 7.06E-06 |
| Gm2163 | tss_body | -1.37 | 4.21E-09 | 7.06E-06 |
| Gm2163 | tss_body | -1.66 | 4.22E-09 | 7.06E-06 |
| Disp1 | tss_body | 2.01 | 4.30E-09 | 7.10E-06 |
| Daam2 | tss_body | -1.78 | 4.35E-09 | 7.10E-06 |
| Rora | tss_body | -2.72 | 4.35E-09 | 7.10E-06 |
| Q7TRS3 | tss_1500 | -2.21 | 4.36E-09 | 7.10E-06 |
| Lrrtm4 | tss_body | -1.74 | 4.37E-09 | 7.10E-06 |
| NA | tss_1500 | 2.35 | 4.39E-09 | 7.10E-06 |
| NA | tss_1500 | -1.54 | 4.46E-09 | 7.16E-06 |
| Med12 | tss_body | 1.48 | 4.49E-09 | 7.17E-06 |
| Platr6 | tss_body | 1.74 | 4.53E-09 | 7.17E-06 |
| Adh6a | tss_1500 | 1.54 | 4.54E-09 | 7.17E-06 |
| 4930539M17Rik | tss_body | 1.76 | 4.57E-09 | 7.18E-06 |
| Tspan5 | tss_body | 1.85 | 4.60E-09 | 7.18E-06 |
| Tnni1 | tss_body | 1.8 | 4.64E-09 | 7.18E-06 |
| Kcnf1 | tss_body | -1.31 | 4.64E-09 | 7.18E-06 |
| NA | tss_200 | -2.55 | 4.70E-09 | 7.22E-06 |
| NA | tss_body | -2.06 | 4.76E-09 | 7.25E-06 |
| Osgin1 | tss_body | 1.76 | 4.79E-09 | 7.25E-06 |
| NA | tss_body | 2.5 | 4.79E-09 | 7.25E-06 |
| Gm40032 | tss_body | -1.36 | 4.91E-09 | 7.33E-06 |
| Glp2r | tss_body | -2.08 | 4.92E-09 | 7.33E-06 |
| LOC108169009 | tss_body | -1.97 | 4.93E-09 | 7.33E-06 |
| NA | tss_body | -2.44 | 4.97E-09 | 7.33E-06 |
| Sdr9c7 | tss_body | 1.1 | 4.98E-09 | 7.33E-06 |
| 1500009L16Rik | tss_body | 1.85 | 5.02E-09 | 7.36E-06 |
| Kcnma1 | tss_body | -2.14 | 5.13E-09 | 7.39E-06 |
| Arhgef6 | tss_body | 4.66 | 5.14E-09 | 7.39E-06 |
| NA | NA | 2.48 | 5.15E-09 | 7.39E-06 |
| Gm41492 | tss_body | -2.17 | 5.18E-09 | 7.39E-06 |
| Ccl27a | tss_body | -1.2 | 5.18E-09 | 7.39E-06 |
| NA | tss_body | -1.53 | 5.20E-09 | 7.39E-06 |
| Fam69c | tss_body | -1.58 | 5.22E-09 | 7.39E-06 |
| Rfx4 | tss_body | 1.77 | 5.25E-09 | 7.39E-06 |
| Tg | tss_body | 2.52 | 5.29E-09 | 7.42E-06 |
| Sox2ot | tss_body | -2.34 | 5.33E-09 | 7.43E-06 |

| | | | | |
|---|---|---|---|---|
| Gm2163 | tss_body | -1.51 | 5.39E-09 | 7.44E-06 |
| Asap1 | tss_body | 1.71 | 5.40E-09 | 7.44E-06 |
| NA | NA | 1.16 | 5.42E-09 | 7.44E-06 |
| Epb41l4a | tss_body | 1.95 | 5.46E-09 | 7.46E-06 |
| Trim35 | tss_body | -1.39 | 5.57E-09 | 7.55E-06 |
| Csn1s1 | tss_body | -1.5 | 5.57E-09 | 7.55E-06 |
| NA | tss_body | -1.98 | 5.61E-09 | 7.55E-06 |
| NA | NA | 1.21 | 5.67E-09 | 7.55E-06 |
| CN725425 | tss_body | -1.79 | 5.69E-09 | 7.55E-06 |
| NA | NA | -2.06 | 5.69E-09 | 7.55E-06 |
| Gm31813 | tss_body | 1.77 | 5.71E-09 | 7.55E-06 |
| NA | tss_body | -1.64 | 5.76E-09 | 7.59E-06 |
| NA | NA | -2.47 | 5.86E-09 | 7.60E-06 |
| Sytl3 | tss_body | 1.66 | 5.87E-09 | 7.60E-06 |
| Mak | tss_body | 1.74 | 5.88E-09 | 7.60E-06 |
| 1700094M24Rik | tss_body | -3.03 | 5.91E-09 | 7.60E-06 |
| Ctnnbl1 | tss_body | 1.2 | 5.93E-09 | 7.60E-06 |
| Dock2 | tss_body | -2.14 | 5.95E-09 | 7.60E-06 |
| Ppm1b | tss_body | 1.84 | 6.04E-09 | 7.60E-06 |
| Cd300ld | tss_1500 | -1.65 | 6.07E-09 | 7.60E-06 |
| NA | tss_body | -1.47 | 6.07E-09 | 7.60E-06 |
| Pdgfa | tss_body | 1.25 | 6.08E-09 | 7.60E-06 |
| Ift88 | tss_body | 1.57 | 6.10E-09 | 7.60E-06 |
| Arl6ip5 | tss_body | 2.18 | 6.11E-09 | 7.60E-06 |
| Tbc1d1 | tss_body | 1.72 | 6.15E-09 | 7.60E-06 |
| Etnppl | tss_body | 2.82 | 6.16E-09 | 7.60E-06 |
| NA | NA | -2.08 | 6.17E-09 | 7.60E-06 |
| NA | NA | 1.4 | 6.28E-09 | 7.67E-06 |
| LOC108169009 | tss_body | -1.22 | 6.34E-09 | 7.67E-06 |
| Gm4858 | tss_1500 | -2.04 | 6.35E-09 | 7.67E-06 |
| Pecam1 | tss_body | 1.68 | 6.37E-09 | 7.67E-06 |
| Gm2163 | tss_body | -2.02 | 6.39E-09 | 7.67E-06 |
| Gm21075 | tss_1500 | -2.02 | 6.40E-09 | 7.67E-06 |
| 2310043O21Rik | tss_body | 1.56 | 6.46E-09 | 7.67E-06 |
| Vac14 | tss_body | 1.46 | 6.47E-09 | 7.67E-06 |
| A8Y5L4 | tss_body | 1.64 | 6.48E-09 | 7.67E-06 |
| NA | NA | 1.87 | 6.50E-09 | 7.67E-06 |
| Olfr399 | tss_body | -1.17 | 6.54E-09 | 7.68E-06 |
| Pigr | tss_body | 1.66 | 6.62E-09 | 7.68E-06 |
| 4-Mar | tss_body | 1.5 | 6.67E-09 | 7.68E-06 |
| NA | NA | 1.18 | 6.67E-09 | 7.68E-06 |

| | | | | |
|---|---|---|---|---|
| Mucl2 | tss_body | -2.17 | 6.68E-09 | 7.68E-06 |
| Fgf2 | tss_body | 1.99 | 6.70E-09 | 7.68E-06 |
| Olfr536 | tss_body | -1.74 | 6.71E-09 | 7.68E-06 |
| NA | NA | 1.99 | 6.72E-09 | 7.68E-06 |
| Psg18 | tss_body | -2.17 | 6.78E-09 | 7.72E-06 |
| Atg4a | tss_body | 2.28 | 6.84E-09 | 7.75E-06 |
| NA | NA | -1.85 | 6.88E-09 | 7.76E-06 |
| Dcx | tss_body | 2.79 | 6.90E-09 | 7.76E-06 |
| F8WI90 | tss_200 | 1.73 | 6.94E-09 | 7.77E-06 |
| Vwce | tss_body | 1.46 | 7.12E-09 | 7.87E-06 |
| NA | NA | -1.96 | 7.13E-09 | 7.87E-06 |
| Dcbld2 | tss_body | 1.13 | 7.16E-09 | 7.87E-06 |
| Nav1 | tss_body | 1.2 | 7.16E-09 | 7.87E-06 |
| NA | NA | -2.74 | 7.20E-09 | 7.87E-06 |
| Platr6 | tss_body | 1.86 | 7.21E-09 | 7.87E-06 |
| Tmem108 | tss_body | 2.84 | 7.22E-09 | 7.87E-06 |
| Stk39 | tss_body | 2.21 | 7.25E-09 | 7.87E-06 |
| Cyp4a32 | tss_body | -2.3 | 7.30E-09 | 7.89E-06 |
| Fam205a4 | tss_body | -1.62 | 7.43E-09 | 8.01E-06 |
| Ccdc58 | tss_body | 1.34 | 7.55E-09 | 8.01E-06 |
| Gm21075 | tss_1500 | -1.76 | 7.55E-09 | 8.01E-06 |
| 1500009L16Rik | tss_body | 1.61 | 7.56E-09 | 8.01E-06 |
| Wdr7 | tss_body | 2.34 | 7.57E-09 | 8.01E-06 |
| Zfp385b | tss_body | 1.44 | 7.62E-09 | 8.01E-06 |
| A0A140LI27 | tss_body | -1.23 | 7.64E-09 | 8.01E-06 |
| Pfkfb1 | tss_body | 1.54 | 7.64E-09 | 8.01E-06 |
| LOC108169009 | tss_body | -2.34 | 7.65E-09 | 8.01E-06 |
| Smc5 | tss_body | -2.19 | 7.76E-09 | 8.09E-06 |
| Cttnbp2 | tss_body | 1.94 | 7.79E-09 | 8.09E-06 |
| NA | tss_1500 | -2.31 | 7.86E-09 | 8.13E-06 |
| LOC108169009 | tss_body | -1.31 | 7.92E-09 | 8.16E-06 |
| Gm34388 | tss_body | 1.83 | 7.95E-09 | 8.16E-06 |
| Rad51b | tss_body | 3.66 | 8.02E-09 | 8.16E-06 |
| Olfr666 | tss_200 | -2.07 | 8.03E-09 | 8.16E-06 |
| Gm21936 | tss_1500 | -2.12 | 8.12E-09 | 8.16E-06 |
| Ccl27a | tss_body | -1.15 | 8.13E-09 | 8.16E-06 |
| NA | NA | -2.34 | 8.17E-09 | 8.16E-06 |
| M0QWH4 | tss_body | 2.39 | 8.17E-09 | 8.16E-06 |
| chr4-145640284-145641297 | SShore | -2.04 | 8.20E-09 | 8.16E-06 |
| Pml | tss_body | 1.71 | 8.20E-09 | 8.16E-06 |
| Olfr695 | tss_1500 | -2.06 | 8.23E-09 | 8.16E-06 |

| | | | | |
|---|---|---|---|---|
| Ptprc | tss_body | -1.62 | 8.23E-09 | 8.16E-06 |
| Rian | tss_body | -2.53 | 8.29E-09 | 8.18E-06 |
| Ankrd44 | tss_body | 1.39 | 8.35E-09 | 8.18E-06 |
| Esp31 | tss_body | -1.77 | 8.37E-09 | 8.18E-06 |
| Gm39089 | tss_1500 | 3.17 | 8.39E-09 | 8.18E-06 |
| Gm15325 | tss_200 | 1.43 | 8.41E-09 | 8.18E-06 |
| Sik1 | tss_body | 1.32 | 8.43E-09 | 8.18E-06 |
| NA | NA | 1.49 | 8.50E-09 | 8.22E-06 |
| LOC108169009 | tss_body | -1.17 | 8.58E-09 | 8.26E-06 |
| Fam205a4 | tss_body | -1.34 | 8.60E-09 | 8.26E-06 |
| Cttnbp2 | tss_body | 1.71 | 8.79E-09 | 8.42E-06 |
| NA | NA | -2.17 | 8.92E-09 | 8.49E-06 |
| NA | tss_200 | -2.43 | 8.94E-09 | 8.49E-06 |
| Akirin1 | tss_1500 | 1.35 | 8.95E-09 | 8.49E-06 |
| Ica1l | tss_body | 1.07 | 8.98E-09 | 8.49E-06 |
| Zfp608 | tss_body | -2.44 | 9.12E-09 | 8.57E-06 |
| NA | NA | -1.69 | 9.16E-09 | 8.57E-06 |
| Eno2 | tss_body | 1.63 | 9.16E-09 | 8.57E-06 |
| S100a16 | tss_body | 2.49 | 9.20E-09 | 8.57E-06 |
| NA | NA | -2.25 | 9.23E-09 | 8.57E-06 |
| Stpg1 | tss_body | 1.93 | 9.28E-09 | 8.57E-06 |
| Tnfsfm13 | tss_body | 2.48 | 9.28E-09 | 8.57E-06 |
| Cacna1c | tss_body | 2.57 | 9.37E-09 | 8.61E-06 |
| Ube2o | tss_body | 1.4 | 9.39E-09 | 8.61E-06 |
| Gm38794 | tss_body | 1.31 | 9.47E-09 | 8.65E-06 |
| NA | tss_body | 1.1 | 9.49E-09 | 8.65E-06 |
| A730004F24Rik | tss_body | -2.69 | 9.66E-09 | 8.78E-06 |
| NA | tss_body | 1.69 | 9.79E-09 | 8.87E-06 |
| NA | tss_body | 1.48 | 9.89E-09 | 8.93E-06 |
| Dst | tss_body | 1.69 | 9.96E-09 | 8.97E-06 |
| 4833422C13Rik | tss_body | 1.83 | 1.00E-08 | 9.00E-06 |
| Osbpl1a | tss_body | 1.7 | 1.01E-08 | 9.00E-06 |
| NA | NA | 1.99 | 1.02E-08 | 9.07E-06 |
| Lrp6 | tss_body | 1.68 | 1.05E-08 | 9.27E-06 |
| Gm33415 | tss_body | 1.78 | 1.05E-08 | 9.27E-06 |
| Insr | tss_body | 1.3 | 1.06E-08 | 9.27E-06 |
| NA | tss_body | 2.07 | 1.06E-08 | 9.27E-06 |
| Sspn | tss_body | 1.12 | 1.06E-08 | 9.27E-06 |
| Tspan9 | tss_body | 1.62 | 1.06E-08 | 9.27E-06 |
| NA | NA | -2.27 | 1.06E-08 | 9.27E-06 |
| Olfr1499 | tss_1500 | -1.74 | 1.07E-08 | 9.31E-06 |

| | | | | |
|---|---|---|---|---|
| Ccdc57 | tss_body | 1.43 | 1.07E-08 | 9.31E-06 |
| Mrgprb8 | tss_1500 | -2.06 | 1.08E-08 | 9.37E-06 |
| NA | tss_body | -1.93 | 1.09E-08 | 9.41E-06 |
| NA | NA | 2.15 | 1.11E-08 | 9.44E-06 |
| Rapgef5 | tss_body | 1.35 | 1.11E-08 | 9.44E-06 |
| Pisd | tss_body | -1.49 | 1.11E-08 | 9.44E-06 |
| Mroh6 | tss_body | 1.2 | 1.11E-08 | 9.44E-06 |
| Col16a1 | tss_body | 1.36 | 1.12E-08 | 9.44E-06 |
| Prkag2 | tss_body | 1.53 | 1.12E-08 | 9.44E-06 |
| Cfh | tss_body | -1.94 | 1.12E-08 | 9.44E-06 |
| Ccdc70 | tss_body | -1.9 | 1.13E-08 | 9.44E-06 |
| Jade1 | tss_body | 1.29 | 1.13E-08 | 9.44E-06 |
| NA | NA | -1.5 | 1.13E-08 | 9.44E-06 |
| Kat6a | tss_body | 1.74 | 1.13E-08 | 9.44E-06 |
| Myo10 | tss_body | 1.97 | 1.14E-08 | 9.44E-06 |
| Vdr | tss_1500 | -1.86 | 1.14E-08 | 9.44E-06 |
| LOC108168771 | tss_body | -1.51 | 1.14E-08 | 9.44E-06 |
| LOC108168771 | tss_body | -2.12 | 1.14E-08 | 9.44E-06 |
| Fgf12 | tss_body | 2.13 | 1.15E-08 | 9.49E-06 |
| Vmn2r78 | tss_1500 | -1.88 | 1.16E-08 | 9.50E-06 |
| Tmem132d | tss_body | -1.89 | 1.16E-08 | 9.50E-06 |
| NA | tss_1500 | -1.87 | 1.18E-08 | 9.60E-06 |
| Cep350 | tss_body | -2.62 | 1.18E-08 | 9.62E-06 |
| Atxn10 | tss_body | 2.26 | 1.19E-08 | 9.62E-06 |
| Oxr1 | tss_body | 1.89 | 1.19E-08 | 9.62E-06 |
| Spata13 | tss_body | 1.47 | 1.19E-08 | 9.62E-06 |
| Sema4a | tss_body | 1.77 | 1.21E-08 | 9.71E-06 |
| Rlbp1 | tss_body | -2.26 | 1.22E-08 | 9.71E-06 |
| Bdnf | tss_body | 2.23 | 1.22E-08 | 9.71E-06 |
| Pdzd2 | tss_body | 2.59 | 1.22E-08 | 9.71E-06 |
| Q8VGV6 | tss_200 | -2.47 | 1.22E-08 | 9.71E-06 |
| Exog | tss_body | 1.25 | 1.23E-08 | 9.72E-06 |
| Phyhip | tss_body | 2.17 | 1.24E-08 | 9.80E-06 |
| Tbcd | tss_body | 1.14 | 1.25E-08 | 9.80E-06 |
| Ablim1 | tss_body | 2.03 | 1.25E-08 | 9.80E-06 |
| Gml2 | tss_body | 1.59 | 1.25E-08 | 9.80E-06 |
| chr2-150470084-150470288 | NShore | -2.3 | 1.25E-08 | 9.80E-06 |
| NA | NA | 1.87 | 1.27E-08 | 9.91E-06 |
| Grap2 | tss_body | 1.22 | 1.28E-08 | 9.94E-06 |
| NA | NA | 1.25 | 1.28E-08 | 9.94E-06 |
| Srl | tss_body | 1.05 | 1.31E-08 | 1.01E-05 |

| | | | | |
|---|---|---|---|---|
| Gm34726 | tss_body | -1.75 | 1.31E-08 | 1.01E-05 |
| NA | NA | 1.06 | 1.32E-08 | 1.02E-05 |
| Samd5 | tss_body | -0.95 | 1.33E-08 | 1.02E-05 |
| NA | NA | 1.55 | 1.33E-08 | 1.02E-05 |
| Gzmc | tss_body | -2.17 | 1.33E-08 | 1.02E-05 |
| LOC108169009 | tss_body | -1.78 | 1.35E-08 | 1.02E-05 |
| Gm36071 | tss_body | -1.92 | 1.35E-08 | 1.02E-05 |
| Gm12295 | tss_body | -2.95 | 1.36E-08 | 1.03E-05 |
| NA | NA | 2.81 | 1.37E-08 | 1.03E-05 |
| Vmn1r39 | tss_body | -2.37 | 1.38E-08 | 1.03E-05 |
| Efl1 | tss_body | 1.9 | 1.38E-08 | 1.03E-05 |
| NA | tss_body | -1.86 | 1.38E-08 | 1.03E-05 |
| St3gal3 | tss_body | 2.81 | 1.39E-08 | 1.03E-05 |
| Gm2163 | tss_body | -2.07 | 1.39E-08 | 1.03E-05 |
| Zmiz1 | tss_body | 1.59 | 1.39E-08 | 1.03E-05 |
| Ppm1h | tss_body | 1.23 | 1.41E-08 | 1.05E-05 |
| Aspscr1 | tss_body | 2.34 | 1.42E-08 | 1.05E-05 |
| Gm36268 | tss_body | 1.59 | 1.42E-08 | 1.05E-05 |
| NA | NA | -1.9 | 1.44E-08 | 1.05E-05 |
| Olfr1507 | tss_body | -2.29 | 1.44E-08 | 1.05E-05 |
| Npepps | tss_body | 1.48 | 1.44E-08 | 1.05E-05 |
| LOC108169009 | tss_body | -1.34 | 1.44E-08 | 1.05E-05 |
| Trib3 | tss_body | 1.29 | 1.44E-08 | 1.05E-05 |
| Agpat4 | tss_body | 1.57 | 1.45E-08 | 1.05E-05 |
| Chd2 | tss_body | 1.62 | 1.45E-08 | 1.05E-05 |
| Stat5a | tss_1500 | 3.1 | 1.46E-08 | 1.05E-05 |
| Pgap2 | tss_body | 1.21 | 1.46E-08 | 1.05E-05 |
| Creb3l1 | tss_body | 1.69 | 1.47E-08 | 1.06E-05 |
| NA | tss_body | -1.65 | 1.48E-08 | 1.06E-05 |
| NA | NA | -0.99 | 1.50E-08 | 1.07E-05 |
| NA | NA | 2.18 | 1.50E-08 | 1.07E-05 |
| Gm2163 | tss_body | -2.25 | 1.51E-08 | 1.07E-05 |
| Fam205a4Gm2163 | tss_body | -2.21 | 1.51E-08 | 1.07E-05 |
| NA | tss_1500 | 2.99 | 1.52E-08 | 1.07E-05 |
| NA | NA | 1.51 | 1.52E-08 | 1.07E-05 |
| Gm31143 | tss_body | 2.22 | 1.53E-08 | 1.07E-05 |
| Fam205a4 | tss_body | -2.14 | 1.53E-08 | 1.07E-05 |
| Etnppl | tss_body | 2.6 | 1.53E-08 | 1.07E-05 |
| C87414 | tss_body | -1.51 | 1.53E-08 | 1.07E-05 |
| NA | NA | -2.45 | 1.54E-08 | 1.07E-05 |
| 4833447I15Rik | tss_body | 2.63 | 1.54E-08 | 1.07E-05 |

| | | | | |
|---|---|---|---|---|
| Macrod2 | tss_body | 1.92 | 1.54E-08 | 1.07E-05 |
| Fam205a4 | tss_body | -1.64 | 1.54E-08 | 1.07E-05 |
| NA | NA | -1.3 | 1.55E-08 | 1.07E-05 |
| NA | tss_body | -2.35 | 1.57E-08 | 1.08E-05 |
| NA | tss_body | -1.61 | 1.59E-08 | 1.10E-05 |
| Fam205a4 | tss_body | -2.67 | 1.60E-08 | 1.10E-05 |
| Gramd2 | tss_body | 1.28 | 1.61E-08 | 1.10E-05 |
| Gm2163 | tss_body | -1.87 | 1.62E-08 | 1.10E-05 |
| Tmem110 | tss_body | 1.64 | 1.62E-08 | 1.10E-05 |
| Gm2163 | tss_body | -1.49 | 1.63E-08 | 1.10E-05 |
| Limk2 | tss_body | 1.29 | 1.64E-08 | 1.10E-05 |
| NA | NA | 1.98 | 1.64E-08 | 1.10E-05 |
| NA | NA | 3.53 | 1.64E-08 | 1.10E-05 |
| A2A517 | tss_body | -1.19 | 1.64E-08 | 1.10E-05 |
| Gm30246 | tss_body | 1.96 | 1.66E-08 | 1.11E-05 |
| Ppm1b | tss_body | 1.15 | 1.66E-08 | 1.11E-05 |
| Frrs1l | tss_body | 1.33 | 1.68E-08 | 1.12E-05 |
| Ophn1 | tss_body | 2.12 | 1.70E-08 | 1.13E-05 |
| Gm34276 | tss_body | 1.14 | 1.70E-08 | 1.13E-05 |
| Fam205a4 | tss_body | -1.47 | 1.71E-08 | 1.13E-05 |
| NA | tss_body | -2.3 | 1.71E-08 | 1.13E-05 |
| Ccr6 | tss_body | -2.33 | 1.71E-08 | 1.13E-05 |
| Abca16 | tss_body | -2.11 | 1.71E-08 | 1.13E-05 |
| NA | tss_200 | -1.93 | 1.74E-08 | 1.14E-05 |
| Pml | tss_body | 2.98 | 1.75E-08 | 1.14E-05 |
| Psd4 | tss_body | 1.46 | 1.75E-08 | 1.14E-05 |
| Itpr3 | tss_body | 2.21 | 1.75E-08 | 1.14E-05 |
| Fpr-rs6 | tss_body | -1.94 | 1.76E-08 | 1.14E-05 |
| NA | tss_body | -1.65 | 1.76E-08 | 1.14E-05 |
| Tas2r116 | tss_body | -2.19 | 1.77E-08 | 1.14E-05 |
| 2210408F21Rik | tss_body | -1.79 | 1.81E-08 | 1.17E-05 |
| Efcab6 | tss_body | 1.83 | 1.82E-08 | 1.17E-05 |
| Krt75 | tss_body | -1.56 | 1.83E-08 | 1.17E-05 |
| Hrnr | tss_body | -1.63 | 1.83E-08 | 1.17E-05 |
| Gm20751 | tss_body | -1.63 | 1.83E-08 | 1.17E-05 |
| Folh1 | tss_body | -2.52 | 1.83E-08 | 1.17E-05 |
| Sfn | tss_1500 | 1.1 | 1.83E-08 | 1.17E-05 |
| Sat1 | tss_body | 2.46 | 1.84E-08 | 1.17E-05 |
| Ube2v2 | tss_body | 1.75 | 1.84E-08 | 1.17E-05 |
| NA | tss_body | -2.19 | 1.85E-08 | 1.17E-05 |
| NA | NA | 1.81 | 1.86E-08 | 1.17E-05 |

| | | | | |
|---|---|---|---|---|
| NA | NA | 2.37 | 1.86E-08 | 1.17E-05 |
| Calr4 | tss_body | 1.64 | 1.87E-08 | 1.17E-05 |
| Unc13d | tss_1500 | 2.66 | 1.87E-08 | 1.17E-05 |
| NA | tss_body | 1.87 | 1.87E-08 | 1.17E-05 |
| NA | NA | -2.08 | 1.88E-08 | 1.17E-05 |
| NA | NA | 2.2 | 1.88E-08 | 1.17E-05 |
| Pramel7 | tss_body | -1.91 | 1.89E-08 | 1.18E-05 |
| Reg3d | tss_1500 | -2.03 | 1.90E-08 | 1.18E-05 |
| Ezr | tss_body | 1.92 | 1.90E-08 | 1.18E-05 |
| Sult2a4 | tss_body | -1.74 | 1.90E-08 | 1.18E-05 |
| Lilr4b | tss_body | -1.87 | 1.92E-08 | 1.18E-05 |
| Olfr729 | tss_body | -1.41 | 1.92E-08 | 1.18E-05 |
| NA | NA | 1.57 | 1.93E-08 | 1.18E-05 |
| Lockd | tss_body | -2.7 | 1.94E-08 | 1.18E-05 |
| Gm2163 | tss_body | -1.72 | 1.94E-08 | 1.18E-05 |
| NA | NA | 1.14 | 1.94E-08 | 1.18E-05 |
| P2rx7 | tss_body | 1.11 | 1.94E-08 | 1.18E-05 |
| NA | NA | 1.88 | 1.95E-08 | 1.18E-05 |
| A730046J19Rik | tss_200 | 1.91 | 1.95E-08 | 1.18E-05 |
| Alad | tss_body | 2.29 | 1.95E-08 | 1.18E-05 |
| NA | tss_body | -2.51 | 1.96E-08 | 1.19E-05 |
| Usp9y | tss_body | -1.95 | 1.98E-08 | 1.19E-05 |
| Palm2 | tss_body | -2.38 | 1.98E-08 | 1.19E-05 |
| Hrnr | tss_body | -1.7 | 1.99E-08 | 1.19E-05 |
| Mthfsl | tss_1500 | -1.94 | 1.99E-08 | 1.19E-05 |
| NA | tss_body | 2.06 | 2.00E-08 | 1.19E-05 |
| Ezr | tss_body | 1.94 | 2.00E-08 | 1.19E-05 |
| NA | NA | 1.23 | 2.01E-08 | 1.19E-05 |
| chr6-51168827-51169063 | SShelf | 2.77 | 2.01E-08 | 1.19E-05 |
| NA | NA | 1.35 | 2.01E-08 | 1.19E-05 |
| NA | NA | -1.8 | 2.01E-08 | 1.19E-05 |
| U2surp | tss_body | -2.66 | 2.03E-08 | 1.20E-05 |
| NA | NA | -2.6 | 2.03E-08 | 1.20E-05 |
| NA | tss_1500 | 1.79 | 2.04E-08 | 1.20E-05 |
| Gm36684 | tss_body | -2.22 | 2.06E-08 | 1.21E-05 |
| NA | NA | 1.63 | 2.07E-08 | 1.21E-05 |
| Rbm20 | tss_body | 1.51 | 2.08E-08 | 1.21E-05 |
| Aptx | tss_body | 1.67 | 2.09E-08 | 1.22E-05 |
| NA | NA | 1.58 | 2.09E-08 | 1.22E-05 |
| Sfxn1 | tss_body | 1.13 | 2.11E-08 | 1.22E-05 |
| Cngb3 | tss_body | -2.19 | 2.11E-08 | 1.22E-05 |

| | | | | |
|---|---|---|---|---|
| Tnrc6b | tss_body | -2.04 | 2.12E-08 | 1.22E-05 |
| NA | NA | -1.63 | 2.13E-08 | 1.23E-05 |
| BC048546 | tss_body | 1.43 | 2.13E-08 | 1.23E-05 |
| Plec | tss_body | 1.23 | 2.14E-08 | 1.23E-05 |
| NA | NA | 1.76 | 2.15E-08 | 1.23E-05 |
| NA | NA | -1.58 | 2.17E-08 | 1.24E-05 |
| Rbms3 | tss_body | 3.42 | 2.17E-08 | 1.24E-05 |
| D17Wsu92e | tss_body | 1.42 | 2.18E-08 | 1.24E-05 |

Table S7: DNA Methylation DMP TKO vs DKO Top500

| Gene | Feature | logFC | P.Value | adj.P.Val |
|---|---|---|---|---|
| LOC102635707 | tss_body | 1.5 | 5.46E-12 | 1.55E-06 |
| LOC102635707 | tss_body | 1.45 | 7.83E-11 | 1.11E-05 |
| LOC102635707 | tss_body | 2.56 | 5.11E-10 | 4.85E-05 |
| LOC108168771 | tss_body | 1.76 | 7.86E-10 | 5.59E-05 |
| LOC102635707 | tss_body | 2.33 | 1.33E-09 | 7.01E-05 |
| Tekt1 | tss_body | 2 | 1.48E-09 | 7.01E-05 |
| LOC108168771 | tss_body | 2.17 | 2.11E-09 | 8.57E-05 |
| Gm36071 | tss_body | 2.12 | 2.61E-09 | 9.09E-05 |
| LOC102635707 | tss_body | 2.22 | 2.88E-09 | 9.09E-05 |
| NA | NA | 2.43 | 3.80E-09 | 0.00010804 |
| Cyb5d1 | tss_body | -3.9 | 6.10E-09 | 0.00015773 |
| NA | NA | 1.72 | 6.91E-09 | 0.00016375 |
| Myl1 | tss_body | -5.62 | 7.61E-09 | 0.00016661 |
| Acadl | tss_body | 1.45 | 1.04E-08 | 0.00018005 |
| NA | NA | 1.33 | 1.04E-08 | 0.00018005 |
| Ccl27a | tss_body | 0.96 | 1.22E-08 | 0.00018005 |
| LOC108168771 | tss_body | 2.38 | 1.28E-08 | 0.00018005 |
| Tekt1 | tss_body | 1.92 | 1.30E-08 | 0.00018005 |
| Gm34719 | tss_body | 2.04 | 1.30E-08 | 0.00018005 |
| Vamp2 | tss_body | 3.28 | 1.48E-08 | 0.00018005 |
| Ccl27a | tss_body | 1.17 | 1.50E-08 | 0.00018005 |
| Gm36011 | tss_body | 2.15 | 1.53E-08 | 0.00018005 |
| Gm11677 | tss_body | 1.61 | 1.55E-08 | 0.00018005 |
| NA | tss_body | 3.03 | 1.58E-08 | 0.00018005 |
| Palm2 | tss_body | 1.51 | 1.58E-08 | 0.00018005 |
| Specc1 | tss_body | -4.94 | 1.76E-08 | 0.000187437 |
| LOC102635707 | tss_body | 2.09 | 1.78E-08 | 0.000187437 |
| Ccl27a | tss_body | 1.09 | 2.30E-08 | 0.00022503 |
| NA | NA | 2.08 | 2.32E-08 | 0.00022503 |
| chr6-51168827-51169063 | SShelf | -2.75 | 2.37E-08 | 0.00022503 |
| NA | NA | 1.71 | 2.56E-08 | 0.000235149 |
| Gm32328 | tss_body | 1.36 | 3.01E-08 | 0.000267875 |
| Gm4872 | tss_body | 2.28 | 3.24E-08 | 0.000279167 |
| Gm30219 | tss_body | 1.38 | 3.87E-08 | 0.000323753 |
| NA | NA | 1.04 | 4.14E-08 | 0.000336823 |
| NA | NA | 1.29 | 4.65E-08 | 0.000367508 |
| Eif5a | tss_body | 1.69 | 5.56E-08 | 0.000427686 |
| Ces1a | tss_body | 2.61 | 6.14E-08 | 0.000459056 |
| Abca16 | tss_body | 1.94 | 6.33E-08 | 0.000459056 |
| NA | tss_body | 2.59 | 6.45E-08 | 0.000459056 |

| | | | | |
|---|---|---|---|---|
| NA | tss_1500 | 2.02 | 7.07E-08 | 0.000483305 |
| NA | NA | 1.69 | 7.13E-08 | 0.000483305 |
| NA | tss_body | 1.39 | 8.53E-08 | 0.000564493 |
| NA | NA | 1.38 | 1.00E-07 | 0.000643694 |
| NA | NA | 1.43 | 1.02E-07 | 0.000643694 |
| Kcnma1 | tss_body | 1.88 | 1.11E-07 | 0.000678976 |
| Kcnma1 | tss_body | 1.98 | 1.12E-07 | 0.000678976 |
| Palm2 | tss_body | 1.43 | 1.21E-07 | 0.000717739 |
| NA | tss_1500 | 1.25 | 1.35E-07 | 0.000757804 |
| NA | tss_body | -1.83 | 1.35E-07 | 0.000757804 |
| Cdkn2a | tss_body | 3.02 | 1.36E-07 | 0.000757804 |
| NA | NA | 2.31 | 1.39E-07 | 0.00075802 |
| Mup20 | tss_1500 | 1.38 | 1.47E-07 | 0.00076817 |
| Q924H8 | tss_1500 | -1.32 | 1.48E-07 | 0.00076817 |
| Gm31143 | tss_body | -1.93 | 1.48E-07 | 0.00076817 |
| NA | tss_body | 2.14 | 1.58E-07 | 0.0008004 |
| Lpar1 | tss_body | 1.63 | 1.61E-07 | 0.00080389 |
| Rab38 | tss_body | 2.18 | 1.65E-07 | 0.000810022 |
| Creb5 | tss_body | 1.92 | 1.69E-07 | 0.000816984 |
| Olfr224 | tss_body | 2.03 | 1.79E-07 | 0.000850655 |
| NA | NA | 1.4 | 1.90E-07 | 0.00088718 |
| NA | tss_body | 1.7 | 2.05E-07 | 0.000916659 |
| LOC102635707 | tss_body | 1.89 | 2.06E-07 | 0.000916659 |
| Slit3 | tss_body | 1.7 | 2.06E-07 | 0.000916659 |
| 9430019J16Rik | tss_body | 1.54 | 2.11E-07 | 0.000925582 |
| NA | tss_body | 1.38 | 2.18E-07 | 0.000938503 |
| Gm39851 | tss_body | 1.58 | 2.24E-07 | 0.000953133 |
| Rora | tss_body | 1.37 | 2.36E-07 | 0.00098881 |
| Gm36011 | tss_body | 2.21 | 2.40E-07 | 0.00098881 |
| NA | NA | 1.65 | 2.44E-07 | 0.00098973 |
| NA | NA | 1.45 | 2.47E-07 | 0.000991505 |
| NA | tss_1500 | 2.04 | 2.66E-07 | 0.001034868 |
| NA | NA | 1.18 | 2.66E-07 | 0.001034868 |
| Kirrel3 | tss_body | 1.88 | 2.74E-07 | 0.001049718 |
| Ces1a | tss_body | 1.94 | 2.77E-07 | 0.001049718 |
| NA | NA | 1.81 | 2.89E-07 | 0.001071017 |
| Lpar1 | tss_body | 1.98 | 2.90E-07 | 0.001071017 |
| A630050E04Rik | tss_1500 | 2.01 | 2.96E-07 | 0.001071017 |
| NA | NA | 1.76 | 2.97E-07 | 0.001071017 |
| 4930441J16Rik | tss_body | 2.25 | 3.05E-07 | 0.001082966 |
| 1700025K24Rik | tss_body | 0.77 | 3.09E-07 | 0.001086692 |

| | | | | |
|---|---|---|---|---|
| NA | NA | 1.22 | 3.21E-07 | 0.001114066 |
| NA | NA | 2.24 | 3.54E-07 | 0.001211824 |
| Hdgfrp3 | tss_body | -1.86 | 3.67E-07 | 0.00124217 |
| Lpar1 | tss_body | 2.19 | 3.94E-07 | 0.001320056 |
| Gpatch2l | tss_body | 1.79 | 4.17E-07 | 0.001379872 |
| NA | tss_1500 | 1.4 | 4.22E-07 | 0.001379872 |
| Sh3bp4 | tss_body | -3.27 | 4.51E-07 | 0.001429196 |
| NA | tss_body | 1.17 | 4.52E-07 | 0.001429196 |
| Gm36011 | tss_body | 1.63 | 4.59E-07 | 0.001429196 |
| Gm36011 | tss_body | 1.57 | 4.60E-07 | 0.001429196 |
| NA | NA | 2.58 | 4.62E-07 | 0.001429196 |
| Nek11 | tss_body | 2.04 | 4.79E-07 | 0.001465521 |
| Cecr2 | tss_body | 1.21 | 5.02E-07 | 0.001519082 |
| LOC108168771 | tss_body | 1.67 | 5.14E-07 | 0.001529852 |
| Kcnma1 | tss_body | 1.55 | 5.16E-07 | 0.001529852 |
| Acadl | tss_1500 | 0.94 | 5.30E-07 | 0.001555951 |
| LOC108168769 | tss_body | 2.6 | 5.59E-07 | 0.001623071 |
| Gm30132 | tss_body | 1.23 | 5.68E-07 | 0.001630937 |
| Commd1 | tss_body | -0.99 | 5.81E-07 | 0.001652886 |
| LOC108168946 | tss_body | 1.39 | 6.34E-07 | 0.001786504 |
| Ccl27a | tss_body | 0.87 | 6.45E-07 | 0.001798144 |
| NA | tss_1500 | 1.51 | 6.62E-07 | 0.001828381 |
| Slit3 | tss_body | 1.33 | 6.82E-07 | 0.001865393 |
| Txndc8 | tss_200 | 1.03 | 6.92E-07 | 0.001875786 |
| NA | NA | 1.27 | 7.14E-07 | 0.00191528 |
| NA | tss_1500 | 2.37 | 7.39E-07 | 0.001964916 |
| Fmo4 | tss_1500 | 1.21 | 7.54E-07 | 0.001975297 |
| Lpar1 | tss_body | 1.88 | 7.57E-07 | 0.001975297 |
| Shisa6 | tss_body | -1.73 | 7.70E-07 | 0.001992247 |
| Cdr2 | tss_body | -1.31 | 7.93E-07 | 0.002021427 |
| Lpar1 | tss_body | 1.61 | 7.96E-07 | 0.002021427 |
| Lpar1 | tss_body | 1.45 | 8.13E-07 | 0.002047492 |
| NA | NA | 1.65 | 8.29E-07 | 0.002050042 |
| NA | NA | 1.92 | 8.35E-07 | 0.002050042 |
| Gm35426 | tss_body | 1.85 | 8.36E-07 | 0.002050042 |
| Lhfpl3 | tss_body | 1.93 | 8.51E-07 | 0.002068378 |
| NA | NA | -1.38 | 8.80E-07 | 0.002122426 |
| NA | tss_200 | 1.74 | 8.91E-07 | 0.00212928 |
| Kcnma1 | tss_body | 1.78 | 9.11E-07 | 0.002153071 |
| LOC108168771 | tss_body | 1.33 | 9.23E-07 | 0.002153071 |
| Gm38809 | tss_body | 1.29 | 9.26E-07 | 0.002153071 |

| | | | | |
|---|---|---|---|---|
| Myocd | tss_body | 0.98 | 9.31E-07 | 0.002153071 |
| NA | NA | 1.59 | 9.95E-07 | 0.002283154 |
| Gm29683 | tss_body | 1.72 | 1.01E-06 | 0.002307078 |
| NA | tss_body | 1.59 | 1.06E-06 | 0.002377929 |
| Kirrel3 | tss_body | 1.66 | 1.06E-06 | 0.002377929 |
| Ngef | tss_body | 1.21 | 1.07E-06 | 0.002378685 |
| Olfr1151 | tss_body | 2.12 | 1.12E-06 | 0.002459107 |
| Elmo1 | tss_body | 1.61 | 1.12E-06 | 0.002459107 |
| NA | NA | 1.08 | 1.13E-06 | 0.002459683 |
| Rimklb | tss_body | 1.78 | 1.14E-06 | 0.002462851 |
| Ugt1a10 | tss_body | 3.25 | 1.20E-06 | 0.002562767 |
| LOC102635707 | tss_body | 2.37 | 1.22E-06 | 0.002584818 |
| Gm31265 | tss_body | 1.22 | 1.23E-06 | 0.00258541 |
| Rian | tss_body | 1.7 | 1.30E-06 | 0.002708814 |
| NA | NA | 1.54 | 1.31E-06 | 0.002708814 |
| NA | NA | 1.26 | 1.31E-06 | 0.002708814 |
| NA | NA | 1.86 | 1.35E-06 | 0.002763471 |
| Olfr316 | tss_body | -3.78 | 1.37E-06 | 0.002780646 |
| LOC108168771 | tss_body | 2.21 | 1.44E-06 | 0.00291359 |
| Bnc2 | tss_body | 1.29 | 1.46E-06 | 0.00291359 |
| NA | tss_200 | 1.49 | 1.47E-06 | 0.00291359 |
| NA | NA | 1.73 | 1.47E-06 | 0.00291359 |
| LOC108168771 | tss_body | 2.7 | 1.52E-06 | 0.002979837 |
| Sytl2 | tss_body | 1.83 | 1.55E-06 | 0.00300978 |
| 4930441J16Rik | tss_body | 2.3 | 1.56E-06 | 0.00300978 |
| NA | NA | 1.1 | 1.61E-06 | 0.003050412 |
| Yae1d1 | tss_body | -1.52 | 1.61E-06 | 0.003050412 |
| NA | NA | -2.37 | 1.61E-06 | 0.003050412 |
| NA | tss_body | 1.88 | 1.62E-06 | 0.003050412 |
| Palm2 | tss_body | 1.78 | 1.70E-06 | 0.003188312 |
| NA | NA | 2.22 | 1.72E-06 | 0.003195494 |
| Myocd | tss_body | 0.89 | 1.74E-06 | 0.003204118 |
| NA | NA | 1.8 | 1.76E-06 | 0.003204118 |
| Usp9y | tss_body | 1.46 | 1.77E-06 | 0.003204118 |
| A730004F24Rik | tss_body | 1.26 | 1.77E-06 | 0.003204118 |
| Gucy1a2 | tss_body | 1.74 | 1.84E-06 | 0.003315361 |
| NA | NA | 1.3 | 1.88E-06 | 0.003363378 |
| Q8VEZ1 | tss_body | 1.47 | 1.90E-06 | 0.003373721 |
| Kcnma1 | tss_body | 1.47 | 1.93E-06 | 0.003416441 |
| Lhfpl3 | tss_body | 1.8 | 2.03E-06 | 0.003556505 |
| LOC108167525 | tss_body | 1.09 | 2.08E-06 | 0.00361646 |

| | | | | |
|---|---|---|---|---|
| Gm35727 | tss_body | 1.24 | 2.08E-06 | 0.00361646 |
| NA | NA | -1.22 | 2.13E-06 | 0.003672807 |
| Rora | tss_body | 1.83 | 2.20E-06 | 0.003731835 |
| NA | NA | 1.83 | 2.21E-06 | 0.003731835 |
| NA | NA | 1.94 | 2.21E-06 | 0.003731835 |
| NA | NA | -1.95 | 2.22E-06 | 0.003731835 |
| Kcnma1 | tss_body | 1.7 | 2.26E-06 | 0.003775593 |
| NA | NA | 1.64 | 2.31E-06 | 0.003819567 |
| Olfr702 | tss_body | 1.68 | 2.33E-06 | 0.003819567 |
| NA | NA | 1.3 | 2.34E-06 | 0.003819567 |
| Kcnma1 | tss_body | 1.87 | 2.35E-06 | 0.003819567 |
| NA | NA | 1.69 | 2.36E-06 | 0.003819567 |
| NA | tss_body | 1.49 | 2.36E-06 | 0.003819567 |
| NA | tss_1500 | 1.51 | 2.44E-06 | 0.003906811 |
| Commd1 | tss_body | -0.63 | 2.46E-06 | 0.003906811 |
| Kcnma1 | tss_body | 2.03 | 2.47E-06 | 0.003906811 |
| Gm34081 | tss_body | 1.33 | 2.47E-06 | 0.003906811 |
| Ccl27a | tss_body | 1.08 | 2.50E-06 | 0.003927444 |
| A2RST7 | tss_1500 | 1.56 | 2.52E-06 | 0.003934355 |
| Sparc | tss_body | 1.54 | 2.54E-06 | 0.003936006 |
| LOC108169141 | tss_body | 1.21 | 2.55E-06 | 0.003936006 |
| NA | tss_200 | 1.43 | 2.56E-06 | 0.003936006 |
| NA | NA | 1.34 | 2.61E-06 | 0.003984778 |
| NA | tss_1500 | 1.58 | 2.73E-06 | 0.004148815 |
| Gm3331 | tss_body | 0.97 | 2.79E-06 | 0.00421507 |
| Commd1 | tss_body | -0.72 | 2.87E-06 | 0.004326371 |
| NA | NA | 1.7 | 2.90E-06 | 0.00433665 |
| NA | tss_body | 1.52 | 3.02E-06 | 0.004468877 |
| Rimklb | tss_body | 1.87 | 3.03E-06 | 0.004468877 |
| NA | tss_body | 2.14 | 3.03E-06 | 0.004468877 |
| Ikzf2 | tss_body | 1.63 | 3.09E-06 | 0.004502422 |
| Tvp23b | tss_body | 1.03 | 3.10E-06 | 0.004502422 |
| Commd1 | tss_body | -0.8 | 3.10E-06 | 0.004502422 |
| Gm30219 | tss_body | 1.4 | 3.13E-06 | 0.004525646 |
| NA | NA | 1.74 | 3.22E-06 | 0.004624339 |
| NA | tss_body | 2.34 | 3.27E-06 | 0.004650715 |
| NA | tss_body | 1.19 | 3.28E-06 | 0.004650715 |
| NA | NA | 1.6 | 3.29E-06 | 0.004650715 |
| Commd1 | tss_body | -1.01 | 3.32E-06 | 0.004660587 |
| Slc38a4 | tss_body | 1.16 | 3.33E-06 | 0.004660587 |
| Hand2 | tss_body | -1.42 | 3.34E-06 | 0.004660587 |

| | | | | |
|---|---|---|---|---|
| Gm31614 | tss_body | 1.45 | 3.38E-06 | 0.004683359 |
| Fmo1 | tss_body | 0.95 | 3.39E-06 | 0.004683359 |
| Fam205a4 | tss_body | 1.09 | 3.41E-06 | 0.00468367 |
| Kirrel3 | tss_body | 1.3 | 3.44E-06 | 0.004699384 |
| Ccl27a | tss_body | 1.15 | 3.48E-06 | 0.004740925 |
| B130024G19Rik | tss_body | 1 | 3.57E-06 | 0.004837859 |
| NA | NA | 1.4 | 3.62E-06 | 0.004885763 |
| 1700003I22Rik | tss_200 | 1.04 | 3.67E-06 | 0.004911635 |
| NA | tss_200 | 1.32 | 3.68E-06 | 0.004911635 |
| LOC102635707 | tss_body | 1.37 | 3.70E-06 | 0.004915005 |
| Elovl6 | tss_body | 1.94 | 3.74E-06 | 0.004944702 |
| Olfr847 | tss_body | 2.03 | 3.76E-06 | 0.004944702 |
| NA | NA | 1.23 | 3.84E-06 | 0.004944702 |
| NA | tss_body | 1.33 | 3.86E-06 | 0.004944702 |
| 2610002M06Rik | tss_body | -1.22 | 3.87E-06 | 0.004944702 |
| Fpr-rs6 | tss_body | 1.36 | 3.90E-06 | 0.004944702 |
| NA | NA | -1.28 | 3.91E-06 | 0.004944702 |
| NA | NA | -0.71 | 3.91E-06 | 0.004944702 |
| Tbce | tss_1500 | -3.1 | 3.92E-06 | 0.004944702 |
| Zfp692 | tss_body | -1 | 3.93E-06 | 0.004944702 |
| LOC102635707 | tss_body | 1.61 | 3.93E-06 | 0.004944702 |
| Ces1e | tss_body | 1.5 | 3.93E-06 | 0.004944702 |
| Olfr648 | tss_200 | 1.4 | 4.01E-06 | 0.005031113 |
| NA | NA | 2.18 | 4.11E-06 | 0.005112771 |
| Kcnma1 | tss_body | 1.4 | 4.12E-06 | 0.005112771 |
| NA | NA | 1.54 | 4.17E-06 | 0.005130325 |
| Efcab6 | tss_body | -1.28 | 4.19E-06 | 0.005130325 |
| NA | NA | 1.73 | 4.20E-06 | 0.005130325 |
| Fam205a4 | tss_body | 0.95 | 4.20E-06 | 0.005130325 |
| Olfr224 | tss_body | -2.09 | 4.24E-06 | 0.005130325 |
| NA | tss_body | 1.72 | 4.24E-06 | 0.005130325 |
| Ntrk2 | tss_body | 1.55 | 4.26E-06 | 0.005131107 |
| Cyp4a14 | tss_body | 1.41 | 4.28E-06 | 0.005141891 |
| C130060K24Rik | tss_body | 1.47 | 4.36E-06 | 0.005212225 |
| Olfr30 | tss_body | 2.03 | 4.38E-06 | 0.005212225 |
| NA | tss_body | 2.06 | 4.43E-06 | 0.005238216 |
| NA | NA | 1.53 | 4.45E-06 | 0.005238216 |
| Ldhb | tss_body | 1.64 | 4.46E-06 | 0.005238216 |
| Utrn | tss_body | 1.27 | 4.76E-06 | 0.005560387 |
| NA | tss_body | 1.41 | 4.79E-06 | 0.005560387 |
| Svep1 | tss_body | 0.84 | 4.79E-06 | 0.005560387 |

| | | | | |
|---|---|---|---|---|
| Ces1a | tss_body | 1.8 | 4.84E-06 | 0.005595299 |
| Olfr420 | tss_body | 1.09 | 4.90E-06 | 0.005596092 |
| Gm31143 | tss_1500 | -1.56 | 4.91E-06 | 0.005596092 |
| Prdm5 | tss_body | 1.81 | 4.93E-06 | 0.005596092 |
| NA | tss_body | 1.21 | 4.94E-06 | 0.005596092 |
| Vmn1r43 | tss_200 | 1.76 | 4.94E-06 | 0.005596092 |
| NA | NA | 1.53 | 4.96E-06 | 0.005598753 |
| Pygo1 | tss_body | 1.24 | 5.07E-06 | 0.005672914 |
| Gm12381 | tss_body | 1.24 | 5.08E-06 | 0.005672914 |
| NA | tss_body | 1.53 | 5.10E-06 | 0.005672914 |
| Gm30392 | tss_body | 1.59 | 5.11E-06 | 0.005672914 |
| Dzip1l | tss_1500 | -1.28 | 5.12E-06 | 0.005672914 |
| Vmn1r50 | tss_body | 1.57 | 5.27E-06 | 0.005807583 |
| Olfr224 | tss_body | -2.65 | 5.38E-06 | 0.005912147 |
| NA | tss_body | 0.89 | 5.46E-06 | 0.005968342 |
| Islr2 | tss_body | -1.42 | 5.48E-06 | 0.005968342 |
| Ces1c | tss_body | 0.74 | 5.52E-06 | 0.005979081 |
| Dkk2 | tss_body | 1.37 | 5.56E-06 | 0.005979081 |
| NA | tss_body | 1.29 | 5.56E-06 | 0.005979081 |
| Hs3st3a1 | tss_body | -1.09 | 5.58E-06 | 0.005979081 |
| Map2 | tss_body | -1.24 | 5.62E-06 | 0.005979081 |
| Kirrel3 | tss_body | 1.5 | 5.63E-06 | 0.005979081 |
| Glp2r | tss_body | 1.28 | 5.63E-06 | 0.005979081 |
| Olfr224 | tss_body | -3.68 | 5.67E-06 | 0.005994746 |
| Vax1 | tss_body | -1.41 | 5.71E-06 | 0.006020308 |
| Rian | tss_body | 1.41 | 5.94E-06 | 0.006233696 |
| AY761185 | tss_200 | 1.27 | 6.01E-06 | 0.006278589 |
| Ctsc | tss_body | 1.86 | 6.04E-06 | 0.006278589 |
| Gm35332 | tss_body | 1.95 | 6.05E-06 | 0.006278589 |
| Kirrel3 | tss_body | 1.34 | 6.09E-06 | 0.00630021 |
| Mtus2 | tss_body | 1.61 | 6.17E-06 | 0.006360605 |
| Plekhs1 | tss_body | -1.76 | 6.29E-06 | 0.006442675 |
| Pramel7 | tss_body | 1.3 | 6.31E-06 | 0.006442675 |
| Sfrp5 | tss_body | -2.59 | 6.32E-06 | 0.006442675 |
| Lhfpl3 | tss_body | 1.34 | 6.44E-06 | 0.006528828 |
| Gm33642 | tss_body | -1.56 | 6.45E-06 | 0.006528828 |
| NA | tss_body | 1.55 | 6.50E-06 | 0.006555995 |
| Utrn | tss_body | 1.73 | 6.55E-06 | 0.006555995 |
| AU023762 | tss_body | 1.75 | 6.55E-06 | 0.006555995 |
| Mir3969 | tss_1500 | 1.64 | 6.58E-06 | 0.006555995 |
| Gm35988 | tss_body | 1.35 | 6.59E-06 | 0.006555995 |

| | | | | |
|---|---|---|---|---|
| Kcnj12 | tss_body | 1.3 | 6.62E-06 | 0.006562369 |
| NA | tss_1500 | 1.16 | 6.75E-06 | 0.006648239 |
| Gata3 | tss_body | 1.32 | 6.77E-06 | 0.006648239 |
| NA | NA | 1.03 | 6.78E-06 | 0.006648239 |
| Wwox | tss_body | -2 | 6.81E-06 | 0.006660788 |
| Kcnma1 | tss_body | 1.35 | 6.88E-06 | 0.00667939 |
| Ebf1 | tss_body | 1.55 | 6.88E-06 | 0.00667939 |
| Ankrd36 | tss_body | -0.92 | 6.94E-06 | 0.006711661 |
| NA | NA | 1.8 | 7.12E-06 | 0.00685437 |
| NA | NA | -1.77 | 7.14E-06 | 0.00685437 |
| NA | NA | 0.98 | 7.16E-06 | 0.00685437 |
| Gm13067 | tss_body | 1.32 | 7.20E-06 | 0.006875687 |
| Kcnma1 | tss_body | 1.58 | 7.33E-06 | 0.006957226 |
| NA | tss_200 | -4.05 | 7.35E-06 | 0.006957226 |
| Q7TRG4 | tss_body | 1.99 | 7.42E-06 | 0.006957226 |
| Olfr279 | tss_200 | 1.69 | 7.44E-06 | 0.006957226 |
| NA | tss_body | 1.37 | 7.48E-06 | 0.006957226 |
| Kirrel3 | tss_body | 1.27 | 7.49E-06 | 0.006957226 |
| Mgat5b | tss_body | -1.3 | 7.51E-06 | 0.006957226 |
| Lpar1 | tss_body | 1.4 | 7.58E-06 | 0.006957226 |
| Tnfsfm13 | tss_body | -1.59 | 7.59E-06 | 0.006957226 |
| NA | tss_body | 1.36 | 7.61E-06 | 0.006957226 |
| Gm41162 | tss_body | 1.96 | 7.65E-06 | 0.006957226 |
| Lhfpl3 | tss_body | 1.34 | 7.65E-06 | 0.006957226 |
| Scn10a | tss_body | 0.9 | 7.69E-06 | 0.006957226 |
| Larp1 | tss_body | 1.65 | 7.71E-06 | 0.006957226 |
| NA | tss_body | 1.64 | 7.71E-06 | 0.006957226 |
| Ccl27a | tss_body | 0.8 | 7.72E-06 | 0.006957226 |
| Gm35727 | tss_body | 1.84 | 7.72E-06 | 0.006957226 |
| NA | NA | 1.4 | 7.73E-06 | 0.006957226 |
| 4921504E06Rik | tss_body | -1.36 | 7.77E-06 | 0.006959771 |
| NA | tss_200 | 2 | 7.80E-06 | 0.006959771 |
| NA | tss_body | 0.92 | 7.80E-06 | 0.006959771 |
| NA | NA | 1.84 | 7.85E-06 | 0.006973915 |
| Ano2 | tss_body | 1.14 | 7.88E-06 | 0.006973915 |
| NA | tss_body | 0.71 | 7.89E-06 | 0.006973915 |
| Guk1 | tss_body | 1.72 | 7.95E-06 | 0.007000608 |
| Fmo4 | tss_body | 0.93 | 7.99E-06 | 0.007018456 |
| NA | NA | 1.9 | 8.02E-06 | 0.007018456 |
| Gm33149 | tss_body | 1.3 | 8.13E-06 | 0.007097493 |
| chr18-81916242-81916539 | SShore | 1.85 | 8.28E-06 | 0.007204702 |

| | | | | |
|---|---|---|---|---|
| NA | tss_body | 1.1 | 8.43E-06 | 0.007292778 |
| NA | NA | 1.31 | 8.43E-06 | 0.007292778 |
| Vmn1r30 | tss_body | 0.86 | 8.57E-06 | 0.007386453 |
| Rgs6 | tss_body | 1.24 | 8.62E-06 | 0.007406602 |
| NA | tss_body | 1.12 | 8.75E-06 | 0.007485636 |
| NA | NA | 1.13 | 8.76E-06 | 0.007485636 |
| NA | NA | 1.01 | 8.80E-06 | 0.00749845 |
| NA | NA | 1 | 8.83E-06 | 0.0075024 |
| AU023762 | tss_body | 1.9 | 8.90E-06 | 0.007538053 |
| Gzmc | tss_body | 1.4 | 9.07E-06 | 0.007657395 |
| Mecom | tss_body | 1.36 | 9.14E-06 | 0.007685891 |
| NA | NA | 0.95 | 9.16E-06 | 0.007685891 |
| Gm38809 | tss_body | 1.06 | 9.22E-06 | 0.007713132 |
| NA | tss_body | 1.55 | 9.27E-06 | 0.007713132 |
| NA | tss_body | -0.74 | 9.28E-06 | 0.007713132 |
| Ugt1a10 | tss_body | 1.98 | 9.31E-06 | 0.007713132 |
| Gm41492 | tss_body | 1.32 | 9.34E-06 | 0.007713132 |
| Gm4070 | tss_body | 1.12 | 9.40E-06 | 0.007713132 |
| NA | tss_body | 1.49 | 9.40E-06 | 0.007713132 |
| Foxp2 | tss_body | 1.47 | 9.41E-06 | 0.007713132 |
| Creb5 | tss_body | 1.47 | 9.55E-06 | 0.007774787 |
| Svep1 | tss_body | -1.57 | 9.57E-06 | 0.007774787 |
| Hs6st2 | tss_body | 1.75 | 9.59E-06 | 0.007774787 |
| NA | NA | 1.89 | 9.61E-06 | 0.007774787 |
| Fam110b | tss_body | 1.93 | 9.62E-06 | 0.007774787 |
| E330023G01Rik | tss_200 | 1.1 | 9.68E-06 | 0.007788782 |
| Gm42324 | tss_body | -1.15 | 9.69E-06 | 0.007788782 |
| NA | NA | 1.5 | 9.85E-06 | 0.007893555 |
| NA | NA | 1.22 | 1.01E-05 | 0.008041407 |
| Gm31731 | tss_1500 | 1.4 | 1.01E-05 | 0.008041407 |
| A2m | tss_1500 | 1.17 | 1.01E-05 | 0.008041407 |
| Oosp1 | tss_body | 1.4 | 1.01E-05 | 0.008041407 |
| Daam2 | tss_body | 1.07 | 1.03E-05 | 0.00812784 |
| Lpar1 | tss_body | 2.01 | 1.03E-05 | 0.008140434 |
| NA | NA | -1.8 | 1.04E-05 | 0.008179877 |
| Rora | tss_body | 1.62 | 1.04E-05 | 0.008180075 |
| Kiz | tss_body | -1.36 | 1.05E-05 | 0.008183769 |
| Reg3d | tss_1500 | 1.33 | 1.05E-05 | 0.008183769 |
| NA | NA | 1.97 | 1.05E-05 | 0.008183769 |
| NA | NA | 0.92 | 1.07E-05 | 0.008308238 |
| NA | NA | 1.87 | 1.08E-05 | 0.008357898 |

| | | | | |
|---|---|---|---|---|
| Rimklb | tss_body | 1.51 | 1.09E-05 | 0.008402846 |
| NA | NA | 1.95 | 1.09E-05 | 0.008408363 |
| Tmem132d | tss_body | 1.19 | 1.12E-05 | 0.008623206 |
| Sacs | tss_1500 | -1.53 | 1.13E-05 | 0.008630187 |
| Kcnb1 | tss_body | 1.97 | 1.13E-05 | 0.008630187 |
| NA | NA | 1.43 | 1.13E-05 | 0.008630187 |
| Palm2 | tss_body | 0.81 | 1.14E-05 | 0.008676961 |
| Fam205a4 | tss_body | 1.05 | 1.15E-05 | 0.008683152 |
| Gm34719 | tss_body | 1.46 | 1.15E-05 | 0.008683152 |
| Vmn2r118 | tss_body | 1.35 | 1.16E-05 | 0.008683152 |
| NA | tss_body | 1.7 | 1.16E-05 | 0.008683152 |
| Col25a1 | tss_body | 1.46 | 1.16E-05 | 0.008683152 |
| Gm32771 | tss_body | -2.26 | 1.16E-05 | 0.008683152 |
| LOC102635707 | tss_body | 1.69 | 1.17E-05 | 0.008683152 |
| Slc1a3 | tss_body | 0.87 | 1.17E-05 | 0.008687788 |
| NA | tss_1500 | 1.89 | 1.19E-05 | 0.0088015 |
| Olfr703 | tss_1500 | 1.65 | 1.19E-05 | 0.0088015 |
| NA | NA | 1.25 | 1.20E-05 | 0.008814346 |
| NA | tss_1500 | 1.58 | 1.22E-05 | 0.008970672 |
| Tas2r116 | tss_body | 1.41 | 1.23E-05 | 0.009025127 |
| Phactr1 | tss_body | 1.59 | 1.24E-05 | 0.009040695 |
| Q8VGU9 | tss_body | 1.51 | 1.24E-05 | 0.009045832 |
| A930003O13Rik | tss_body | 1.91 | 1.25E-05 | 0.009106607 |
| Pparg | tss_body | -0.97 | 1.26E-05 | 0.009111291 |
| Dhrs7c | tss_body | 1.49 | 1.27E-05 | 0.009167566 |
| Gm29478 | tss_body | 1.34 | 1.27E-05 | 0.009185821 |
| NA | NA | -2.2 | 1.28E-05 | 0.009219652 |
| Pmfbp1 | tss_body | 1.75 | 1.29E-05 | 0.00924297 |
| NA | NA | -1.31 | 1.29E-05 | 0.009277041 |
| NA | tss_1500 | 0.79 | 1.32E-05 | 0.009428809 |
| Cyp4b1 | tss_body | 1.43 | 1.32E-05 | 0.009428809 |
| NA | NA | 1.76 | 1.34E-05 | 0.009530785 |
| Atp2a1 | tss_body | -1.13 | 1.36E-05 | 0.009623676 |
| Uaca | tss_body | -1.49 | 1.37E-05 | 0.009712733 |
| NA | tss_body | 1.37 | 1.39E-05 | 0.009778007 |
| Wscd2 | tss_body | 1.26 | 1.39E-05 | 0.009778007 |
| NA | NA | 1.59 | 1.40E-05 | 0.009797163 |
| Csmd2 | tss_body | 1.38 | 1.40E-05 | 0.009797163 |
| NA | NA | -1.36 | 1.42E-05 | 0.009910504 |
| NA | NA | 1.67 | 1.43E-05 | 0.009944531 |
| Cps1 | tss_body | 1.23 | 1.44E-05 | 0.010006292 |

| | | | | |
|---|---|---|---|---|
| Ldhb | tss_body | 1.31 | 1.44E-05 | 0.010006292 |
| Hoxc6 | tss_body | -1 | 1.45E-05 | 0.010026121 |
| NA | NA | 1.79 | 1.45E-05 | 0.010026121 |
| Gm20878 | tss_1500 | 0.82 | 1.46E-05 | 0.010029134 |
| NA | NA | 1.31 | 1.47E-05 | 0.010117097 |
| NA | NA | 1.47 | 1.48E-05 | 0.010117097 |
| Acadl | tss_body | -1.58 | 1.48E-05 | 0.010117097 |
| Nr2f2 | tss_body | -1.48 | 1.50E-05 | 0.010239807 |
| 2900060L22Rik | tss_body | -0.99 | 1.51E-05 | 0.010257414 |
| NA | NA | 1.15 | 1.51E-05 | 0.010272397 |
| NA | NA | 1.33 | 1.52E-05 | 0.01027243 |
| Slco1a5 | tss_body | 1.19 | 1.52E-05 | 0.01027243 |
| Ces1c | tss_1500 | 1.42 | 1.52E-05 | 0.01027243 |
| Gm30192 | tss_body | 1.27 | 1.53E-05 | 0.01027243 |
| Gm31711 | tss_body | 1.39 | 1.54E-05 | 0.010321222 |
| Rora | tss_body | 1.69 | 1.55E-05 | 0.010350318 |
| Fam205a4 | tss_body | 1.03 | 1.55E-05 | 0.010350318 |
| Gm5065 | tss_body | 1.19 | 1.56E-05 | 0.010350318 |
| Gm36139 | tss_body | 0.84 | 1.56E-05 | 0.010350318 |
| C130060K24Rik | tss_body | 1.69 | 1.56E-05 | 0.010362461 |
| NA | NA | 1.96 | 1.57E-05 | 0.010362461 |
| NA | NA | 1.58 | 1.57E-05 | 0.010376943 |
| Ctnnbip1 | tss_body | -1.13 | 1.61E-05 | 0.010576644 |
| Kcnma1 | tss_body | 1.89 | 1.61E-05 | 0.010576644 |
| NA | NA | 1.59 | 1.62E-05 | 0.010593252 |
| Ebf2 | tss_body | 1.29 | 1.64E-05 | 0.010724635 |
| Myh13 | tss_body | 0.75 | 1.65E-05 | 0.010752409 |
| NA | tss_200 | 1.59 | 1.66E-05 | 0.010775517 |
| Ces1a | tss_body | 1.47 | 1.67E-05 | 0.01080396 |
| Olfr1513 | tss_body | 1.3 | 1.67E-05 | 0.01080396 |
| Elovl6 | tss_body | 2.33 | 1.69E-05 | 0.010902437 |
| NA | NA | 1.55 | 1.70E-05 | 0.010979934 |
| Rnf182 | tss_body | 1.26 | 1.71E-05 | 0.011018539 |
| NA | tss_1500 | 1.72 | 1.73E-05 | 0.011082843 |
| Bax | tss_body | 1.27 | 1.73E-05 | 0.011082843 |
| Elmo1 | tss_body | -1.16 | 1.74E-05 | 0.011092899 |
| Gng4 | tss_body | 5.22 | 1.74E-05 | 0.011092899 |
| Gm34501 | tss_body | 1.24 | 1.75E-05 | 0.011092899 |
| LOC108169141 | tss_body | 1.13 | 1.75E-05 | 0.011092899 |
| Lpar1 | tss_body | 0.89 | 1.75E-05 | 0.011092899 |
| NA | tss_body | 1.09 | 1.76E-05 | 0.011116066 |

| | | | | |
|---|---|---|---|---|
| 4930504O13Rik | tss_body | 0.73 | 1.77E-05 | 0.011166116 |
| Sez6l | tss_body | 1.15 | 1.78E-05 | 0.011182808 |
| Dnajc4 | tss_1500 | 1.34 | 1.81E-05 | 0.011379027 |
| NA | NA | 1.32 | 1.82E-05 | 0.011385679 |
| Gm36139 | tss_body | 1.01 | 1.82E-05 | 0.011385679 |
| NA | tss_body | 1.87 | 1.83E-05 | 0.011385679 |
| Hspa12b | tss_body | 1.02 | 1.83E-05 | 0.011385679 |
| 4930557F10Rik | tss_body | 1.85 | 1.83E-05 | 0.011385679 |
| Spire2 | tss_body | 0.84 | 1.86E-05 | 0.011471162 |
| C1qtnf2 | tss_body | 1.66 | 1.86E-05 | 0.011471162 |
| Celsr1 | tss_body | 1.65 | 1.86E-05 | 0.011471162 |
| Gm4858 | tss_1500 | 1.19 | 1.86E-05 | 0.011471162 |
| Lrrtm4 | tss_body | 1.48 | 1.87E-05 | 0.011488117 |
| Gm13306 | tss_body | 0.67 | 1.88E-05 | 0.011498957 |
| NA | NA | 1.2 | 1.88E-05 | 0.011527847 |
| Rab31 | tss_body | 1.48 | 1.89E-05 | 0.011539943 |
| NA | tss_body | 1.28 | 1.90E-05 | 0.011586186 |
| Slc2a5 | tss_1500 | -1.19 | 1.92E-05 | 0.01164773 |
| Gm12253 | tss_body | 1.91 | 1.94E-05 | 0.011736522 |
| NA | NA | -1.78 | 1.94E-05 | 0.011736522 |
| NA | NA | 1.63 | 1.95E-05 | 0.011736522 |
| Gm29536 | tss_body | -1.16 | 1.95E-05 | 0.011736522 |
| chr5-127241442-127242848 | NShore | 1.65 | 1.95E-05 | 0.011736522 |
| NA | NA | 1.26 | 1.97E-05 | 0.011826223 |
| Slc1a3 | tss_body | 1.24 | 1.98E-05 | 0.011881453 |
| Gm42266 | tss_body | 1.7 | 2.01E-05 | 0.011917826 |
| NA | tss_1500 | 1.53 | 2.01E-05 | 0.011917826 |
| chr4-99650896-99651569 | SShelf | -1.47 | 2.01E-05 | 0.011917826 |
| Gm29683 | tss_body | 1.39 | 2.02E-05 | 0.011917826 |
| Ntm | tss_body | 1.8 | 2.02E-05 | 0.011917826 |
| NA | NA | 1.09 | 2.02E-05 | 0.011917826 |
| Scly | tss_body | 1.31 | 2.02E-05 | 0.011917826 |
| NA | NA | -1.56 | 2.03E-05 | 0.011917826 |
| Cyp4a10 | tss_body | 1.09 | 2.03E-05 | 0.011917826 |
| NA | tss_1500 | 1.23 | 2.04E-05 | 0.011982964 |
| Rora | tss_body | 1.46 | 2.06E-05 | 0.012041247 |
| NA | NA | 1.43 | 2.06E-05 | 0.012041247 |
| NA | NA | -0.9 | 2.07E-05 | 0.012066012 |
| 5730559C18Rik | tss_body | 1.53 | 2.07E-05 | 0.012066012 |
| NA | tss_body | 0.92 | 2.08E-05 | 0.012070306 |
| Rerg | tss_body | -1.31 | 2.10E-05 | 0.012159983 |

| | | | | |
|---|---|---|---|---|
| Commd1 | tss_body | -0.82 | 2.11E-05 | 0.012206547 |
| Gm30288 | tss_body | 1.02 | 2.12E-05 | 0.012242298 |
| NA | tss_body | 1.35 | 2.13E-05 | 0.012262418 |
| chr1-69376995-69377218 | SShore | 1.56 | 2.16E-05 | 0.0123881 |
| Zscan10 | tss_body | 1.9 | 2.16E-05 | 0.012397141 |
| Ccl27a | tss_body | 1.35 | 2.17E-05 | 0.012397141 |
| NA | NA | 1.39 | 2.18E-05 | 0.012482137 |
| Grin2b | tss_body | -1.76 | 2.20E-05 | 0.012517444 |

TAble S7: DNA Methylation DMP DKO_EP vs DKO Top500

| Gene | Feature | logFC | P.Value | adj.P.Val |
|---|---|---|---|---|
| NA | NA | -1.94 | 1.03E-10 | 2.92E-05 |
| Gm37013 | tss_body | 1.96 | 1.07E-07 | 0.012350197 |
| Chil5 | tss_body | 2.11 | 1.61E-07 | 0.012350197 |
| Asb4 | tss_200 | 1.92 | 1.74E-07 | 0.012350197 |
| Yae1d1 | tss_body | -1.7 | 3.24E-07 | 0.018422839 |
| Palm2 | tss_body | 1.07 | 3.05E-06 | 0.124898602 |
| Fndc3c1 | tss_body | 1.58 | 3.07E-06 | 0.124898602 |
| Itln1 | tss_body | 2.71 | 5.43E-06 | 0.193151747 |
| NA | NA | -2 | 1.15E-05 | 0.331156643 |
| NA | NA | -1.64 | 1.23E-05 | 0.331156643 |
| NA | NA | -2.18 | 1.28E-05 | 0.331156643 |
| Pcdh11x | tss_body | -2.21 | 1.55E-05 | 0.348944572 |
| Ptpdc1 | tss_body | -3.1 | 1.69E-05 | 0.348944572 |
| Tssc1 | tss_body | 4.52 | 1.72E-05 | 0.348944572 |
| Rtkn | tss_body | 2.21 | 2.06E-05 | 0.368896518 |
| Chil3 | tss_body | 1.25 | 2.07E-05 | 0.368896518 |
| Olfr418 | tss_body | -1.09 | 2.28E-05 | 0.379713608 |
| Cd244 | tss_body | 2.14 | 2.40E-05 | 0.379713608 |
| Tbce | tss_1500 | -2.69 | 2.55E-05 | 0.381831981 |
| Mob3b | tss_body | -1.74 | 2.70E-05 | 0.384091985 |
| Chil3 | tss_200 | 1.78 | 3.39E-05 | 0.432620657 |
| NA | tss_body | 1.54 | 3.50E-05 | 0.432620657 |
| NA | NA | 1.06 | 3.58E-05 | 0.432620657 |
| LOC102632256 | tss_body | -1.2 | 3.79E-05 | 0.432620657 |
| Oxr1 | tss_body | -1.38 | 3.80E-05 | 0.432620657 |
| NA | NA | -1.46 | 4.06E-05 | 0.444791416 |
| Ttll11 | tss_body | 1.4 | 4.67E-05 | 0.47315208 |
| Haglr | tss_body | -1.79 | 4.73E-05 | 0.47315208 |
| Oxr1 | tss_body | -1.59 | 4.88E-05 | 0.47315208 |
| NA | NA | -0.57 | 5.05E-05 | 0.47315208 |
| Gm33190 | tss_body | 1.21 | 5.16E-05 | 0.47315208 |
| NA | NA | 2.78 | 5.51E-05 | 0.474898844 |
| NA | NA | 1.61 | 5.59E-05 | 0.474898844 |
| Runx3 | tss_body | -0.68 | 5.68E-05 | 0.474898844 |
| Gm39696 | tss_body | 1.26 | 6.53E-05 | 0.519792785 |
| Man1c1 | tss_body | -1.61 | 6.58E-05 | 0.519792785 |
| E0CZ38 | tss_body | -1.89 | 7.15E-05 | 0.521184635 |
| Hand2 | tss_body | -1.11 | 7.24E-05 | 0.521184635 |
| Mob3b | tss_body | -3.83 | 7.33E-05 | 0.521184635 |
| Tfpi | tss_body | -1.96 | 7.33E-05 | 0.521184635 |

| | | | | |
|---|---|---|---|---|
| Myo3b | tss_body | -1.71 | 7.51E-05 | 0.521184635 |
| chr11-46056019-46056350 | NShelf | -1.02 | 7.90E-05 | 0.522666045 |
| Itln1 | tss_body | -1.62 | 7.90E-05 | 0.522666045 |
| Pifo | tss_body | 1.84 | 8.89E-05 | 0.575014933 |
| Q5NCC8 | tss_body | 0.93 | 9.50E-05 | 0.575564952 |
| NA | tss_body | 1.12 | 9.59E-05 | 0.575564952 |
| Gm31333 | tss_body | 1.36 | 9.69E-05 | 0.575564952 |
| Rnf220 | tss_body | -1.23 | 9.71E-05 | 0.575564952 |
| NA | NA | -0.81 | 0.000101871 | 0.581424533 |
| NA | NA | -0.96 | 0.000102181 | 0.581424533 |
| NA | NA | -0.89 | 0.000117411 | 0.634451602 |
| NA | NA | -1.12 | 0.000119388 | 0.634451602 |
| Mybpc2 | tss_body | 1.46 | 0.000119904 | 0.634451602 |
| Gm30094 | tss_body | -1.11 | 0.00012042 | 0.634451602 |
| Haglr | tss_body | -1.53 | 0.000128014 | 0.649003946 |
| NA | NA | -0.94 | 0.000131473 | 0.649003946 |
| Gm36857 | tss_1500 | 0.95 | 0.000131729 | 0.649003946 |
| Sox10 | tss_body | 1.03 | 0.000132306 | 0.649003946 |
| Myt1l | tss_body | 2.22 | 0.000139493 | 0.669484593 |
| Myt1l | tss_body | 2.17 | 0.000143093 | 0.669484593 |
| Dcdc2c | tss_body | 2.26 | 0.000145708 | 0.669484593 |
| Mks1 | tss_body | 0.59 | 0.000145894 | 0.669484593 |
| Pcdh11x | tss_body | -1.44 | 0.000148294 | 0.669694296 |
| Ldlrap1 | tss_body | -1.84 | 0.000151683 | 0.674295524 |
| Zfy1 | tss_body | 1.17 | 0.000154054 | 0.674300557 |
| Lyve1 | tss_body | -1.18 | 0.000157084 | 0.677145234 |
| Gm33763 | tss_body | 1.34 | 0.000159545 | 0.67748926 |
| Rtkn | tss_body | 1.86 | 0.000169998 | 0.695981823 |
| Tmem56 | tss_body | -1.03 | 0.000171032 | 0.695981823 |
| NA | tss_body | 1.46 | 0.000176832 | 0.695981823 |
| Cdkn2a | tss_body | 1.76 | 0.00017914 | 0.695981823 |
| Myt1l | tss_body | 3.07 | 0.000180537 | 0.695981823 |
| Rnf220 | tss_body | 0.86 | 0.000181771 | 0.695981823 |
| Gm35507 | tss_body | 1.13 | 0.000182226 | 0.695981823 |
| Ralyl | tss_body | 1.55 | 0.000184129 | 0.695981823 |
| NA | tss_body | -2.85 | 0.000185956 | 0.695981823 |
| Zfp423 | tss_body | -0.77 | 0.000189524 | 0.695981823 |
| NA | NA | 1.02 | 0.000190809 | 0.695981823 |
| NA | NA | -1.91 | 0.00019439 | 0.696466669 |
| NA | NA | -1.15 | 0.000196201 | 0.696466669 |
| Pcdh11x | tss_body | -1.52 | 0.000198285 | 0.696466669 |

| | | | | |
|---|---|---|---|---|
| Tmem233 | tss_body | 1.97 | 0.000207511 | 0.705425595 |
| Pcdh11x | tss_body | -1.62 | 0.000211066 | 0.705425595 |
| Myo3b | tss_body | -2.09 | 0.000211628 | 0.705425595 |
| NA | NA | -0.94 | 0.000212935 | 0.705425595 |
| Q8VGL4 | tss_body | 1.4 | 0.000213233 | 0.705425595 |
| Leo1 | tss_body | -1.36 | 0.000220482 | 0.721020427 |
| NA | NA | -1.45 | 0.00022844 | 0.727206799 |
| LOC108169009 | tss_body | 1.39 | 0.000229508 | 0.727206799 |
| Meikin | tss_body | 1.63 | 0.000230041 | 0.727206799 |
| Pcdh11x | tss_body | -1.79 | 0.000235554 | 0.73149946 |
| Gdf15 | tss_body | 0.78 | 0.000242727 | 0.73149946 |
| Map3k7 | tss_body | -1.72 | 0.000242856 | 0.73149946 |
| Tenm4 | tss_body | 1.89 | 0.000244643 | 0.73149946 |
| Gm37013 | tss_body | 1.48 | 0.0002465 | 0.73149946 |
| Gad1os | tss_body | 1.23 | 0.000249062 | 0.73149946 |
| NA | NA | -0.98 | 0.00025163 | 0.73149946 |
| Chil3 | tss_body | 1.21 | 0.000251968 | 0.73149946 |
| Itln1 | tss_1500 | -1.63 | 0.000260722 | 0.74582123 |
| Nek11 | tss_body | 0.6 | 0.000262544 | 0.74582123 |
| NA | NA | -1.05 | 0.000266757 | 0.74582123 |
| Chrm3 | tss_body | -1.71 | 0.000267387 | 0.74582123 |
| 2810459M11Rik | tss_body | -0.62 | 0.000271753 | 0.746791846 |
| Epb41l4a | tss_body | -1.69 | 0.000272985 | 0.746791846 |
| Man1c1 | tss_body | -1.1 | 0.000281723 | 0.762960176 |
| Agbl1 | tss_body | 1.28 | 0.000284632 | 0.762960176 |
| chr4-125729578-125729806 | Island | 2.49 | 0.00028694 | 0.762960176 |
| Tnfaip8l3 | tss_body | 0.75 | 0.000298938 | 0.779500199 |
| NA | NA | -1.1 | 0.000304286 | 0.779500199 |
| Rbm7 | tss_1500 | -1.6 | 0.000308626 | 0.779500199 |
| Gpr158 | tss_body | 1.56 | 0.000310623 | 0.779500199 |
| Gm33763 | tss_body | 1.21 | 0.000310752 | 0.779500199 |
| Mapk4 | tss_body | 2.98 | 0.000311911 | 0.779500199 |
| Foxl2os | tss_body | 1.37 | 0.000312339 | 0.779500199 |
| Sec23a | tss_body | 0.6 | 0.000320601 | 0.789079348 |
| Sgce | tss_1500 | -0.94 | 0.000321889 | 0.789079348 |
| Prrt4 | tss_1500 | 1.62 | 0.000324498 | 0.789079348 |
| NA | tss_1500 | -0.96 | 0.000334545 | 0.797721324 |
| NA | tss_body | -1 | 0.000338117 | 0.797721324 |
| Fam13c | tss_body | 0.72 | 0.000340615 | 0.797721324 |
| Lpin1 | tss_body | -1.07 | 0.000341303 | 0.797721324 |
| Gm5617 | tss_body | -1.34 | 0.000345778 | 0.797721324 |

| | | | | |
|---|---|---|---|---|
| Aurkc | tss_body | 1.22 | 0.00034775 | 0.797721324 |
| NA | tss_1500 | -1.27 | 0.000347991 | 0.797721324 |
| Cys1 | tss_body | 2.19 | 0.00035336 | 0.797721324 |
| Tmem150c | tss_body | -1.26 | 0.000355062 | 0.797721324 |
| Arhgap23 | tss_body | 1.18 | 0.000365987 | 0.797721324 |
| Chil4 | tss_body | 1.57 | 0.000368404 | 0.797721324 |
| Wdr64 | tss_body | 1.37 | 0.000369625 | 0.797721324 |
| NA | NA | -1.07 | 0.000371347 | 0.797721324 |
| Pard3b | tss_body | 0.65 | 0.00037154 | 0.797721324 |
| Mcc | tss_body | -0.84 | 0.000372085 | 0.797721324 |
| Ppp1r12a | tss_body | 0.85 | 0.000372914 | 0.797721324 |
| 3110039M20Rik | tss_body | -0.93 | 0.000384763 | 0.816926267 |
| NA | tss_200 | -1.04 | 0.000389021 | 0.819848329 |
| Metap1d | tss_1500 | 2.19 | 0.000398755 | 0.83418415 |
| NA | NA | 1.07 | 0.000408282 | 0.847878543 |
| Tenm4 | tss_body | 2.08 | 0.000412096 | 0.849399225 |
| Pbx1 | tss_body | -1.2 | 0.000414985 | 0.849399225 |
| Tchhl1 | tss_body | 1.28 | 0.00043556 | 0.854362083 |
| Sall3 | tss_body | -2.41 | 0.000436044 | 0.854362083 |
| NA | tss_200 | -1.98 | 0.000438641 | 0.854362083 |
| Tmem178 | tss_body | -0.73 | 0.00044264 | 0.854362083 |
| NA | tss_body | 1.04 | 0.000451469 | 0.854362083 |
| NA | NA | 1.23 | 0.000454817 | 0.854362083 |
| Satb1 | tss_body | -1.39 | 0.000464292 | 0.854362083 |
| Adamts20 | tss_body | 1.1 | 0.000465975 | 0.854362083 |
| Nrde2 | tss_body | -0.5 | 0.000471015 | 0.854362083 |
| NA | NA | 1.08 | 0.000471047 | 0.854362083 |
| Olfr1320 | tss_200 | -0.87 | 0.000472848 | 0.854362083 |
| Gm33763 | tss_body | 1.37 | 0.000476011 | 0.854362083 |
| Oxr1 | tss_body | -1.51 | 0.000478584 | 0.854362083 |
| Chil3 | tss_body | 1.79 | 0.000485812 | 0.854362083 |
| B230323A14Rik | tss_body | 0.77 | 0.000493697 | 0.854362083 |
| Runx1 | tss_body | -1.28 | 0.00049604 | 0.854362083 |
| Dnajc15 | tss_body | 0.6 | 0.000498719 | 0.854362083 |
| Chil4 | tss_body | 1.5 | 0.000500026 | 0.854362083 |
| Htr5a | tss_body | 1.03 | 0.000501435 | 0.854362083 |
| Oosp1 | tss_body | 1 | 0.000502141 | 0.854362083 |
| NA | NA | 1.85 | 0.000506867 | 0.854362083 |
| Gm30505 | tss_body | -0.93 | 0.000508056 | 0.854362083 |
| NA | tss_200 | -1.01 | 0.00050884 | 0.854362083 |
| NA | NA | 1.12 | 0.000509402 | 0.854362083 |

| | | | | |
|---|---|---|---|---|
| Atxn7l1 | tss_body | 0.73 | 0.000515206 | 0.854362083 |
| NA | NA | -1.07 | 0.000518158 | 0.854362083 |
| NA | tss_body | 0.79 | 0.000526002 | 0.854362083 |
| NA | NA | 0.74 | 0.000527183 | 0.854362083 |
| Wapl | tss_body | -0.93 | 0.000527892 | 0.854362083 |
| Gm40149 | tss_1500 | 1 | 0.000536501 | 0.854362083 |
| Olfr1228 | tss_body | 1.06 | 0.000538417 | 0.854362083 |
| Prrt4 | tss_1500 | 1.45 | 0.000538566 | 0.854362083 |
| NA | NA | 1.52 | 0.000547408 | 0.854362083 |
| Rnf43 | tss_body | -1.08 | 0.000547526 | 0.854362083 |
| chr4-125729578-125729806 | Island | 2.57 | 0.000548226 | 0.854362083 |
| 1700057G04Rik | tss_body | -0.72 | 0.000549184 | 0.854362083 |
| Cpb1 | tss_body | -1.12 | 0.000553679 | 0.854362083 |
| Msrb3 | tss_body | 0.76 | 0.000558192 | 0.854362083 |
| NA | tss_body | 1.18 | 0.000558626 | 0.854362083 |
| 6430628N08Rik | tss_body | -1.68 | 0.00056243 | 0.854362083 |
| Hnrnpa1 | tss_body | 0.72 | 0.000562589 | 0.854362083 |
| Fcer1g | tss_body | 0.61 | 0.000563291 | 0.854362083 |
| NA | NA | -1.22 | 0.000564032 | 0.854362083 |
| Gad1 | tss_body | 0.94 | 0.000564999 | 0.854362083 |
| Atf6 | tss_body | 1.01 | 0.000565587 | 0.854362083 |
| Dnaaf3 | tss_body | 1.13 | 0.000565943 | 0.854362083 |
| NA | NA | 0.65 | 0.000568481 | 0.854362083 |
| Brinp3 | tss_body | 0.87 | 0.000568689 | 0.854362083 |
| Pcdha1 | tss_body | -2.16 | 0.000573499 | 0.854362083 |
| Plppr2 | tss_body | 1.45 | 0.000581413 | 0.854362083 |
| LOC108169009 | tss_body | 1.17 | 0.000581963 | 0.854362083 |
| Osbpl3 | tss_body | -0.68 | 0.000584293 | 0.854362083 |
| NA | NA | 0.9 | 0.000585985 | 0.854362083 |
| NA | tss_body | -0.9 | 0.000588416 | 0.854362083 |
| NA | NA | -0.52 | 0.000588671 | 0.854362083 |
| Cald1 | tss_body | 0.83 | 0.000589618 | 0.854362083 |
| NA | NA | 0.73 | 0.00059015 | 0.854362083 |
| Mtus2 | tss_body | -1.78 | 0.00059158 | 0.854362083 |
| NA | tss_body | 0.97 | 0.000602081 | 0.857451056 |
| Arhgef10l | tss_body | -1.58 | 0.000603092 | 0.857451056 |
| Fibcd1 | tss_body | 1.55 | 0.00060402 | 0.857451056 |
| Ppa2 | tss_1500 | 0.71 | 0.000607223 | 0.857451056 |
| Tenm4 | tss_body | 1.73 | 0.000612548 | 0.857451056 |
| NA | NA | -0.86 | 0.00061368 | 0.857451056 |
| Oxr1 | tss_body | -1.47 | 0.000614816 | 0.857451056 |

| | | | | | |
|---|---|---|---|---|---|
| Ctr9 | | tss_body | -0.6 | 0.000620286 | 0.859050421 |
| Q7TRI0 | | tss_1500 | 1.1 | 0.000624488 | 0.859050421 |
| Pik3r1 | | tss_body | 0.96 | 0.000625021 | 0.859050421 |
| Tyr | | tss_body | -0.93 | 0.000631555 | 0.861996884 |
| Myof | | tss_body | -0.68 | 0.000635701 | 0.861996884 |
| | 1-Mar | tss_body | -0.54 | 0.000637454 | 0.861996884 |
| NA | | NA | -0.77 | 0.000639284 | 0.861996884 |
| NA | | tss_1500 | -1.64 | 0.000651831 | 0.871873371 |
| NA | | NA | -0.83 | 0.000652737 | 0.871873371 |
| NA | | NA | -2.36 | 0.000663031 | 0.875810798 |
| NA | | tss_1500 | 0.95 | 0.000665119 | 0.875810798 |
| Scn2a1 | | tss_body | 1.29 | 0.000668983 | 0.875810798 |
| NA | | NA | 0.74 | 0.000670613 | 0.875810798 |
| NA | | NA | -1 | 0.000671785 | 0.875810798 |
| Lpin1 | | tss_body | -2.07 | 0.000674155 | 0.875810798 |
| Gm37013 | | tss_body | 1.2 | 0.000681615 | 0.878926179 |
| Osbpl3 | | tss_body | -0.57 | 0.000684853 | 0.878926179 |
| Gm37013 | | tss_body | 0.86 | 0.000686048 | 0.878926179 |
| Mir7660 | | tss_1500 | 0.39 | 0.00068891 | 0.878926179 |
| Bcat1 | | tss_body | 1.49 | 0.000694215 | 0.881739827 |
| Myh13 | | tss_body | 0.54 | 0.000701427 | 0.886940524 |
| Prdm5 | | tss_body | 0.91 | 0.000706011 | 0.888786028 |
| NA | | NA | 0.88 | 0.000712322 | 0.892781127 |
| Ssrp1 | | tss_body | -0.7 | 0.000717108 | 0.892786189 |
| Chil5 | | tss_body | 1.55 | 0.00072005 | 0.892786189 |
| Fshr | | tss_200 | -0.78 | 0.000726742 | 0.892786189 |
| NA | | NA | -1.41 | 0.000727754 | 0.892786189 |
| NA | | NA | 0.94 | 0.000728016 | 0.892786189 |
| 1700111N16Rik | | tss_body | -0.75 | 0.0007331 | 0.895161841 |
| Kcnn3 | | tss_body | -0.77 | 0.00074321 | 0.903296869 |
| Atp1a4 | | tss_body | 0.56 | 0.000748873 | 0.903296869 |
| Abcc9 | | tss_body | -0.75 | 0.00075026 | 0.903296869 |
| Tfpi | | tss_body | -1.53 | 0.000752462 | 0.903296869 |
| Q8VEZ1 | | tss_body | 0.89 | 0.000760138 | 0.904705473 |
| LOC108169009 | | tss_body | 1.35 | 0.000762607 | 0.904705473 |
| Plppr2 | | tss_body | 1.4 | 0.000763175 | 0.904705473 |
| Olfr727 | | tss_1500 | -1.67 | 0.000776015 | 0.915593929 |
| Cfap99 | | tss_body | -0.95 | 0.000778796 | 0.915593929 |
| Gdf15 | | tss_body | 1.14 | 0.000790606 | 0.925653058 |
| Txndc8 | | tss_body | 0.82 | 0.000801668 | 0.930966115 |
| BC067074 | | tss_body | 0.81 | 0.000805565 | 0.930966115 |

| | | | | |
|---|---|---|---|---|
| chr8-84197390-84197864 | SShelf | 1.91 | 0.000810095 | 0.930966115 |
| LOC108169009 | tss_body | 1.22 | 0.000812489 | 0.930966115 |
| E030019B13Rik | tss_200 | 0.93 | 0.000813416 | 0.930966115 |
| Mtl5 | tss_1500 | 0.56 | 0.000815425 | 0.930966115 |
| NA | tss_body | 1.32 | 0.000824081 | 0.930966115 |
| Zfp423 | tss_body | -0.86 | 0.000824801 | 0.930966115 |
| En1 | tss_body | 1.17 | 0.000829317 | 0.930966115 |
| Pstpip2 | tss_body | 1.38 | 0.000831844 | 0.930966115 |
| NA | NA | -0.61 | 0.000836836 | 0.930966115 |
| chr6-51168827-51169063 | SShelf | 1.14 | 0.000837942 | 0.930966115 |
| A0A140T8K7 | tss_body | -1.56 | 0.000841601 | 0.930966115 |
| Gm29968 | tss_1500 | 0.75 | 0.000843743 | 0.930966115 |
| LOC108169009 | tss_body | 1.14 | 0.000848149 | 0.930966115 |
| Slc10a2 | tss_1500 | -1.03 | 0.000854545 | 0.930966115 |
| Dync1i2 | tss_body | -1.07 | 0.000855196 | 0.930966115 |
| NA | NA | -1.27 | 0.00085651 | 0.930966115 |
| Olfr73 | tss_body | 0.91 | 0.000864993 | 0.930966115 |
| Nek11 | tss_body | 1.25 | 0.000867304 | 0.930966115 |
| Iqcg | tss_body | -1.68 | 0.000868006 | 0.930966115 |
| Pstpip2 | tss_body | 1.48 | 0.000869677 | 0.930966115 |
| Sulf2 | tss_body | -0.66 | 0.000880333 | 0.930966115 |
| LOC108168457 | tss_body | 1.32 | 0.000881943 | 0.930966115 |
| Gm31724 | tss_body | 1.31 | 0.000882704 | 0.930966115 |
| Gdf15 | tss_body | 0.79 | 0.000886129 | 0.930966115 |
| Tenm4 | tss_body | 1.59 | 0.000890335 | 0.930966115 |
| NA | NA | -1.63 | 0.000892401 | 0.930966115 |
| Lrrtm4 | tss_body | 1.12 | 0.000899615 | 0.930966115 |
| Ush2a | tss_body | 1.32 | 0.000901679 | 0.930966115 |
| Tdh | tss_body | 1.04 | 0.00090218 | 0.930966115 |
| NA | NA | -1 | 0.000908504 | 0.930966115 |
| Sox6 | tss_body | 0.85 | 0.000916322 | 0.930966115 |
| Homer1 | tss_body | -0.75 | 0.000916774 | 0.930966115 |
| Rxfp2 | tss_body | 5.26 | 0.000922283 | 0.930966115 |
| NA | NA | -1.15 | 0.00092472 | 0.930966115 |
| Lin28a | tss_body | 1.2 | 0.000926754 | 0.930966115 |
| chr4-125729578-125729806 | Island | 2.17 | 0.000927476 | 0.930966115 |
| Tbx20 | tss_body | 1.01 | 0.000929625 | 0.930966115 |
| Tenm4 | tss_body | 1.33 | 0.000929661 | 0.930966115 |
| Ralyl | tss_body | 1.72 | 0.000932357 | 0.930966115 |
| Msra | tss_body | 0.51 | 0.000932576 | 0.930966115 |
| Olfr331 | tss_1500 | 0.88 | 0.000942042 | 0.932949595 |

| | | | | |
|---|---|---|---|---|
| NA | NA | 2.08 | 0.000942077 | 0.932949595 |
| Otp | tss_body | 0.82 | 0.000944435 | 0.932949595 |
| Tnrc6a | tss_body | -1.25 | 0.00094768 | 0.932949595 |
| Gm31406 | tss_body | -0.79 | 0.000956966 | 0.936798802 |
| Gpr158 | tss_body | 1.77 | 0.000965254 | 0.936798802 |
| Olfr533 | tss_200 | 0.9 | 0.000965551 | 0.936798802 |
| Epsti1 | tss_body | 1.06 | 0.000968505 | 0.936798802 |
| Gm31175 | tss_body | 1.44 | 0.000973641 | 0.936798802 |
| NA | NA | -1.99 | 0.000975772 | 0.936798802 |
| Pitpnc1 | tss_body | -1.1 | 0.000978961 | 0.936798802 |
| Gm42321 | tss_body | -0.88 | 0.00097969 | 0.936798802 |
| Q7TRT9 | tss_1500 | 1.06 | 0.000981224 | 0.936798802 |
| NA | NA | -1.24 | 0.000987429 | 0.939569601 |
| Dzip1 | tss_body | -1.4 | 0.000992875 | 0.941113026 |
| Dcdc5 | tss_body | 1.19 | 0.000997994 | 0.941113026 |
| Olfr648 | tss_body | 0.97 | 0.000998974 | 0.941113026 |
| Cnot8 | tss_body | -0.82 | 0.001005128 | 0.94250072 |
| Pakap | tss_body | 1.63 | 0.001009155 | 0.94250072 |
| Olfr1061 | tss_body | -0.87 | 0.001015223 | 0.94250072 |
| Q7TRV7 | tss_body | 1.01 | 0.001016506 | 0.94250072 |
| NA | tss_body | 1.19 | 0.00101806 | 0.94250072 |
| Flt1 | tss_body | -1.47 | 0.001022845 | 0.94250072 |
| Ppara | tss_body | -0.69 | 0.001023636 | 0.94250072 |
| Nt5c1b | tss_body | -1.01 | 0.001031791 | 0.943637854 |
| NA | tss_body | 1.49 | 0.001031843 | 0.943637854 |
| E9QMD3 | tss_body | -0.92 | 0.00103761 | 0.943637854 |
| Gm595 | tss_body | 0.98 | 0.001038138 | 0.943637854 |
| 6330403K07Rik | tss_body | -1.59 | 0.001054053 | 0.951734418 |
| Fam172a | tss_body | 0.92 | 0.00105435 | 0.951734418 |
| Sulf2 | tss_body | -0.95 | 0.001057081 | 0.951734418 |
| NA | NA | -0.57 | 0.001061523 | 0.95271892 |
| NA | NA | -0.71 | 0.001075892 | 0.957148574 |
| Olfr541 | tss_200 | 1.13 | 0.001078912 | 0.957148574 |
| Hpse2 | tss_body | 0.79 | 0.001085004 | 0.957148574 |
| Gpr89 | tss_body | 0.43 | 0.00108832 | 0.957148574 |
| Rftn1 | tss_body | -0.82 | 0.001091667 | 0.957148574 |
| NA | NA | -1.15 | 0.001099939 | 0.957148574 |
| Nuggc | tss_body | 0.97 | 0.001100767 | 0.957148574 |
| Ccdc170 | tss_body | -1.71 | 0.001104266 | 0.957148574 |
| Gm16845 | tss_body | -0.59 | 0.001104436 | 0.957148574 |
| Actrt3 | tss_200 | 2.35 | 0.001108573 | 0.957148574 |

| | | | | |
|---|---|---|---|---|
| NA | NA | 1.71 | 0.001110775 | 0.957148574 |
| Lmo3 | tss_body | 0.93 | 0.001112542 | 0.957148574 |
| Fxyd6 | tss_body | 1.12 | 0.001113404 | 0.957148574 |
| Gm42411 | tss_body | 0.77 | 0.001113558 | 0.957148574 |
| NA | NA | -1.66 | 0.001121921 | 0.959163605 |
| Slc30a9 | tss_body | -0.73 | 0.001122645 | 0.959163605 |
| Cyp21a1 | tss_1500 | 1.25 | 0.001143995 | 0.967382546 |
| NA | NA | -1.1 | 0.001145057 | 0.967382546 |
| Fam71f2 | tss_body | 0.97 | 0.001145924 | 0.967382546 |
| Pebp4 | tss_body | 0.89 | 0.001147783 | 0.967382546 |
| Azin2 | tss_body | -0.51 | 0.001151243 | 0.967382546 |
| Mrps35 | tss_body | 2.03 | 0.001152666 | 0.967382546 |
| Six3os1 | tss_body | -1.25 | 0.001156379 | 0.967607333 |
| Hoxc13 | tss_body | 1.4 | 0.00116497 | 0.967607333 |
| Q7TRW1 | tss_1500 | 1.5 | 0.001172136 | 0.967607333 |
| Chchd5 | tss_1500 | 1.37 | 0.001173181 | 0.967607333 |
| Dctn6 | tss_body | -0.43 | 0.001177118 | 0.967607333 |
| Exoc2 | tss_body | -0.98 | 0.001177129 | 0.967607333 |
| Epb41l4a | tss_body | -1.31 | 0.001190337 | 0.967607333 |
| Dnajc14 | tss_body | -0.75 | 0.001197338 | 0.967607333 |
| Nudt5 | tss_body | -1.22 | 0.00120148 | 0.967607333 |
| Grik3 | tss_body | 1.38 | 0.001201683 | 0.967607333 |
| Macc1 | tss_body | 0.67 | 0.001203162 | 0.967607333 |
| NA | tss_body | 0.65 | 0.001203357 | 0.967607333 |
| Gpr3 | tss_body | 0.7 | 0.00120347 | 0.967607333 |
| Sema5b | tss_body | -0.53 | 0.001203867 | 0.967607333 |
| Saa2 | tss_1500 | 0.64 | 0.001208571 | 0.967607333 |
| chr2-71544134-71546923 | NShelf | 0.57 | 0.001211902 | 0.967607333 |
| Sim1 | tss_1500 | 0.56 | 0.001217121 | 0.967607333 |
| Lyrm5 | tss_1500 | 0.45 | 0.001217779 | 0.967607333 |
| Tcf4 | tss_body | 1 | 0.001217905 | 0.967607333 |
| NA | tss_body | -0.84 | 0.001225616 | 0.967607333 |
| 9630013D21Rik | tss_1500 | -1.36 | 0.001225964 | 0.967607333 |
| Nell1 | tss_body | -1.64 | 0.001227755 | 0.967607333 |
| Nbas | tss_body | -0.43 | 0.001243799 | 0.969038206 |
| Lpin1 | tss_body | 1.59 | 0.001244694 | 0.969038206 |
| Gm37013 | tss_body | 1.4 | 0.00124553 | 0.969038206 |
| Hoxd12 | tss_body | -1.28 | 0.001250426 | 0.969038206 |
| NA | NA | 1.34 | 0.001251791 | 0.969038206 |
| Mansc4 | tss_200 | -0.85 | 0.001255311 | 0.969038206 |
| Gm30515 | tss_body | 0.82 | 0.001255537 | 0.969038206 |

| | | | | |
|---|---|---|---|---|
| NA | NA | -0.56 | 0.001256819 | 0.969038206 |
| Lhfpl3 | tss_body | 1.01 | 0.001270506 | 0.976670657 |
| Gm35742 | tss_1500 | -1.27 | 0.001274267 | 0.976670657 |
| NA | tss_1500 | -0.89 | 0.001277017 | 0.976670657 |
| Asph | tss_body | -1.36 | 0.001282713 | 0.978397437 |
| Ptprr | tss_body | 0.88 | 0.00128638 | 0.978570254 |
| Fcrlb | tss_body | 1.91 | 0.001309322 | 0.986439211 |
| NA | tss_1500 | -1.25 | 0.001312864 | 0.986439211 |
| NA | NA | -0.79 | 0.001314459 | 0.986439211 |
| Nfatc4 | tss_body | 1.08 | 0.001317986 | 0.986439211 |
| Grik2 | tss_body | -1.14 | 0.001319409 | 0.986439211 |
| Adcy6 | tss_body | -1.26 | 0.001319769 | 0.986439211 |
| Lrrc4c | tss_body | 1.16 | 0.001324776 | 0.986439211 |
| Gdf15 | tss_body | 1.13 | 0.001326766 | 0.986439211 |
| Gm16223 | tss_1500 | -0.84 | 0.00133148 | 0.986439211 |
| Glp2r | tss_body | 0.57 | 0.001340028 | 0.986439211 |
| NA | NA | 1.22 | 0.001340461 | 0.986439211 |
| 4930557F10Rik | tss_body | 0.87 | 0.00134069 | 0.986439211 |
| Il1rn | tss_body | -1.35 | 0.001348781 | 0.986439211 |
| NA | NA | 1.01 | 0.001351179 | 0.986439211 |
| Impa1 | tss_body | 0.76 | 0.001358544 | 0.986439211 |
| Arhgef10l | tss_body | -1.64 | 0.001368271 | 0.986439211 |
| Pip4k2b | tss_body | 0.78 | 0.001369007 | 0.986439211 |
| Ntn1 | tss_body | -0.46 | 0.001369292 | 0.986439211 |
| Nhsl2 | tss_body | -0.88 | 0.001369648 | 0.986439211 |
| NA | NA | -0.95 | 0.001370736 | 0.986439211 |
| NA | NA | 1.76 | 0.00137634 | 0.986439211 |
| Grhl2 | tss_body | -0.66 | 0.00137851 | 0.986439211 |
| Olfr46 | tss_body | 1.15 | 0.001378902 | 0.986439211 |
| Zbtb20 | tss_body | -0.42 | 0.001381429 | 0.986439211 |
| NA | NA | -0.66 | 0.001383403 | 0.986439211 |
| NA | NA | 0.81 | 0.001402461 | 0.99626128 |
| Gal | tss_body | -0.4 | 0.001405524 | 0.99626128 |
| Shroom4 | tss_body | 1.73 | 0.001411684 | 0.99626128 |
| Ccdc170 | tss_body | -1.34 | 0.001415396 | 0.99626128 |
| Gm36287 | tss_body | -1.08 | 0.001419492 | 0.99626128 |
| Tnrc6a | tss_body | -0.53 | 0.001421674 | 0.99626128 |
| 1700112E06Rik | tss_body | -0.64 | 0.001423702 | 0.99626128 |
| Pakap | tss_body | 0.73 | 0.001426878 | 0.99626128 |
| Zswim6 | tss_body | 0.62 | 0.001428693 | 0.99626128 |
| NA | NA | -0.97 | 0.001442143 | 0.998817423 |

| | | | | |
|---|---|---|---|---|
| Msi1 | tss_body | 0.97 | 0.001446005 | 0.998817423 |
| Zfy1 | tss_body | 1.04 | 0.001455424 | 0.998817423 |
| 9630013D21Rik | tss_body | -1.56 | 0.00145858 | 0.998817423 |
| NA | NA | -0.93 | 0.001461992 | 0.998817423 |
| Olfr119 | tss_200 | -0.84 | 0.001464515 | 0.998817423 |
| Ndst4 | tss_body | 0.74 | 0.001467747 | 0.998817423 |
| Dbhos | tss_body | -1.63 | 0.001471467 | 0.998817423 |
| Kcnd3 | tss_body | 0.57 | 0.00147331 | 0.998817423 |
| Akt1s1 | tss_1500 | -0.37 | 0.001474123 | 0.998817423 |
| NA | NA | 1.2 | 0.001475868 | 0.998817423 |
| Haglr | tss_body | -0.85 | 0.001477872 | 0.998817423 |
| Q8VFE8 | tss_body | 0.96 | 0.001480797 | 0.998817423 |
| NA | NA | -0.77 | 0.001481508 | 0.998817423 |
| Etl4 | tss_body | -0.9 | 0.001485159 | 0.99891151 |
| chr17-27415155-27415387 | SShore | -0.62 | 0.001488861 | 0.998989443 |
| Chia1 | tss_body | 1.68 | 0.001497243 | 0.998989443 |
| NA | NA | 0.78 | 0.001499845 | 0.998989443 |
| Peg3 | tss_body | -0.57 | 0.001504769 | 0.998989443 |
| Nuf2 | tss_body | -0.56 | 0.001506636 | 0.998989443 |
| Mageb16 | tss_1500 | -1.07 | 0.001509337 | 0.998989443 |
| Lims2 | tss_body | -0.85 | 0.001515299 | 0.998989443 |
| Dll1 | tss_body | -0.48 | 0.001526697 | 0.998989443 |
| D1Ertd622e | tss_body | -0.62 | 0.001527145 | 0.998989443 |
| Nrn1 | tss_1500 | 0.87 | 0.001527681 | 0.998989443 |
| Lpar1 | tss_body | 0.77 | 0.001529083 | 0.998989443 |
| NA | NA | 1.08 | 0.001533295 | 0.998989443 |
| NA | NA | 1.13 | 0.001533651 | 0.998989443 |
| Slc2a4 | tss_body | 0.85 | 0.001542454 | 0.998989443 |
| Gm20878 | tss_body | 1.08 | 0.001542595 | 0.998989443 |
| Gjb6 | tss_body | 1.51 | 0.001546776 | 0.998989443 |
| NA | NA | 1.27 | 0.001548226 | 0.998989443 |
| Dnmt3a | tss_body | -0.78 | 0.001551328 | 0.998989443 |
| Dph6 | tss_body | -0.49 | 0.001553601 | 0.998989443 |
| Wdr64 | tss_body | -1.37 | 0.0015555 | 0.998989443 |
| NA | NA | 0.73 | 0.001562121 | 0.999982763 |
| NA | NA | 0.64 | 0.001565321 | 0.999982763 |
| NA | NA | -0.91 | 0.001574613 | 0.999982763 |
| Tnfaip8l3 | tss_body | 0.59 | 0.001578227 | 0.999982763 |
| chr19-43524343-43524983 | SShore | -0.49 | 0.001579697 | 0.999982763 |
| Chil5 | tss_body | 1.42 | 0.001587685 | 0.999982763 |
| NA | NA | 0.75 | 0.001593841 | 0.999982763 |

| | | | | |
|---|---|---|---|---|
| Trpv4 | tss_body | -0.7 | 0.001600856 | 0.999982763 |
| Myo7b | tss_body | 1.4 | 0.001602837 | 0.999982763 |
| Asb4 | tss_body | 0.9 | 0.001605968 | 0.999982763 |
| Ipmk | tss_body | 0.99 | 0.001623482 | 0.999982763 |
| Gm38636 | tss_body | 0.69 | 0.001627294 | 0.999982763 |
| 4921508D12Rik | tss_1500 | -0.46 | 0.001630514 | 0.999982763 |
| 2210408F21Rik | tss_body | 1.95 | 0.001635582 | 0.999982763 |
| NA | NA | -1.03 | 0.001640391 | 0.999982763 |
| Gm30192 | tss_body | 1.15 | 0.001640958 | 0.999982763 |
| NA | NA | -0.91 | 0.001644573 | 0.999982763 |
| Olfr279 | tss_200 | 1.03 | 0.00164491 | 0.999982763 |
| chr4-129585157-129585410 | Island | -0.47 | 0.001647903 | 0.999982763 |
| Vmn2r100 | tss_body | -0.66 | 0.00165532 | 0.999982763 |
| NA | NA | 0.88 | 0.001675014 | 0.999982763 |
| Acvr1 | tss_body | 1.35 | 0.001680939 | 0.999982763 |
| LOC102639543 | tss_body | 3.34 | 0.001689058 | 0.999982763 |
| NA | NA | -1.46 | 0.001692202 | 0.999982763 |
| Palm2 | tss_body | 0.66 | 0.001697466 | 0.999982763 |
| St7 | tss_body | -0.37 | 0.001697558 | 0.999982763 |
| Dlg1 | tss_body | 0.81 | 0.001698303 | 0.999982763 |
| LOC108167682 | tss_200 | 1.38 | 0.001699278 | 0.999982763 |
| Wdr64 | tss_body | -1.66 | 0.001709816 | 0.999982763 |
| NA | NA | 0.97 | 0.001715558 | 0.999982763 |
| NA | NA | 0.84 | 0.001715663 | 0.999982763 |
| 6430628N08Rik | tss_body | -1.57 | 0.001716264 | 0.999982763 |
| 1700030N03Rik | tss_body | 2.05 | 0.001721286 | 0.999982763 |
| Wee2 | tss_body | -2.14 | 0.001723866 | 0.999982763 |
| Cobll1 | tss_body | -0.88 | 0.001728409 | 0.999982763 |
| Ptprr | tss_body | 1.31 | 0.001730545 | 0.999982763 |
| Dzip1l | tss_body | 1 | 0.00173094 | 0.999982763 |
| NA | tss_body | 0.99 | 0.001733889 | 0.999982763 |
| Tenm4 | tss_body | 1.68 | 0.001736806 | 0.999982763 |
| Podn | tss_body | 1.4 | 0.00174215 | 0.999982763 |
| LOC108167675 | tss_body | 1.22 | 0.001747373 | 0.999982763 |
| NA | NA | 0.69 | 0.001749451 | 0.999982763 |
| E9PY33 | tss_body | -0.5 | 0.001754128 | 0.999982763 |
| Col6a2 | tss_body | -0.74 | 0.00175432 | 0.999982763 |
| Ccdc170 | tss_body | -1.44 | 0.001754866 | 0.999982763 |
| NA | tss_200 | -0.77 | 0.001755579 | 0.999982763 |
| Nsa2 | tss_body | 0.93 | 0.001763112 | 0.999982763 |
| Bcl6 | tss_body | -0.76 | 0.001767048 | 0.999982763 |

| | | | | |
|---|---|---|---|---|
| Dnhd1 | tss_body | 0.87 | 0.001770132 | 0.999982763 |
| Cpe | tss_body | -0.92 | 0.001771706 | 0.999982763 |
| Dnase1l2 | tss_body | -1.09 | 0.001771945 | 0.999982763 |
| Dnhd1 | tss_body | 0.92 | 0.001783663 | 0.999982763 |
| Fras1 | tss_body | -0.86 | 0.001785921 | 0.999982763 |
| Gpr182 | tss_body | 1.64 | 0.001796255 | 0.999982763 |
| A0A1B0GRG8 | tss_body | -0.97 | 0.00179837 | 0.999982763 |
| Snora16a | tss_1500 | -2.06 | 0.001806831 | 0.999982763 |

Table S7: DNA Methylation DMP DKO_EE vs DKO Top500

| Gene | Feature | logFC | P.Value | adj.P.Val |
|---|---|---|---|---|
| NA | NA | -5.27 | 3.66E-19 | 1.04E-13 |
| LOC102635707 | tss_body | 1.79 | 1.82E-13 | 2.58E-08 |
| NA | NA | 3.35 | 1.87E-12 | 1.77E-07 |
| Fam135b | tss_body | -2.57 | 3.43E-11 | 1.71E-06 |
| LOC102635707 | tss_body | 2.95 | 3.94E-11 | 1.71E-06 |
| Rictor | tss_body | -1.87 | 3.99E-11 | 1.71E-06 |
| Rimklb | tss_body | 3.29 | 4.21E-11 | 1.71E-06 |
| LOC102635707 | tss_body | 1.48 | 5.26E-11 | 1.82E-06 |
| Pitx2 | tss_body | 1.88 | 6.17E-11 | 1.82E-06 |
| Csmd1 | tss_body | -2.28 | 7.64E-11 | 1.82E-06 |
| Rimklb | tss_body | 3.58 | 7.76E-11 | 1.82E-06 |
| chr13-98502955-98503214 | SShore | 2.36 | 8.61E-11 | 1.82E-06 |
| Ano2 | tss_body | 2.21 | 8.75E-11 | 1.82E-06 |
| Rab38 | tss_body | 3.39 | 9.06E-11 | 1.82E-06 |
| Mnx1 | tss_body | 1.55 | 9.61E-11 | 1.82E-06 |
| Foxd2os | tss_body | -2.8 | 1.56E-10 | 2.77E-06 |
| D3Z790 | tss_body | 1.62 | 1.76E-10 | 2.87E-06 |
| Vax1 | tss_body | 1.4 | 1.82E-10 | 2.87E-06 |
| NA | NA | 2.48 | 2.05E-10 | 2.88E-06 |
| Uncx | tss_1500 | 1.46 | 2.10E-10 | 2.88E-06 |
| 3110099E03Rik | tss_body | -2.8 | 2.13E-10 | 2.88E-06 |
| Gm41931 | tss_body | 2.07 | 2.53E-10 | 3.27E-06 |
| Dlg2 | tss_body | 2.42 | 3.42E-10 | 4.10E-06 |
| Ccdc85c | tss_body | -2.01 | 3.46E-10 | 4.10E-06 |
| Pitx1 | tss_body | 2.79 | 3.91E-10 | 4.45E-06 |
| Mcc | tss_body | -1.94 | 4.58E-10 | 5.01E-06 |
| Pitx2 | tss_body | 1.65 | 4.83E-10 | 5.09E-06 |
| Lhx4 | tss_body | 2 | 5.02E-10 | 5.10E-06 |
| 3110099E03Rik | tss_body | -2.45 | 5.83E-10 | 5.72E-06 |
| Plet1os | tss_body | -1.7 | 6.55E-10 | 5.85E-06 |
| Gm41492 | tss_body | 2.08 | 6.73E-10 | 5.85E-06 |
| NA | NA | 1.93 | 6.88E-10 | 5.85E-06 |
| Pitx2 | tss_body | 0.99 | 7.10E-10 | 5.85E-06 |
| Fam135b | tss_body | -2.46 | 7.53E-10 | 5.85E-06 |
| 3110099E03Rik | tss_body | -2.05 | 7.59E-10 | 5.85E-06 |
| Dmrt2 | tss_1500 | 2.19 | 7.94E-10 | 5.85E-06 |
| Igsf11 | tss_body | -2.58 | 8.16E-10 | 5.85E-06 |
| NA | NA | -2.57 | 8.18E-10 | 5.85E-06 |
| Nup93 | tss_body | -3.29 | 8.20E-10 | 5.85E-06 |
| LOC108168771 | tss_body | 1.76 | 8.55E-10 | 5.85E-06 |

| | | | | |
|---|---|---|---|---|
| Lrriq1 | tss_body | 1.95 | 8.62E-10 | 5.85E-06 |
| 3110099E03Rik | tss_200 | -3.4 | 8.70E-10 | 5.85E-06 |
| Nedd4l | tss_body | -2.68 | 8.84E-10 | 5.85E-06 |
| Rimklb | tss_body | 2.76 | 9.07E-10 | 5.86E-06 |
| NA | tss_body | 1.74 | 1.12E-09 | 7.11E-06 |
| Rbfox3 | tss_body | -1.51 | 1.18E-09 | 7.15E-06 |
| Tshz1 | tss_body | -2.38 | 1.19E-09 | 7.15E-06 |
| Alpl | tss_body | -1.74 | 1.23E-09 | 7.15E-06 |
| Prss16 | tss_body | 1.59 | 1.23E-09 | 7.15E-06 |
| Intu | tss_body | -2.22 | 1.27E-09 | 7.22E-06 |
| Pitx2 | tss_body | 1.01 | 1.30E-09 | 7.26E-06 |
| Mnx1 | tss_body | 2.11 | 1.39E-09 | 7.62E-06 |
| Ccdc24 | tss_body | -2.4 | 1.46E-09 | 7.64E-06 |
| Zic5 | tss_body | 1.16 | 1.47E-09 | 7.64E-06 |
| Pygl | tss_body | -2.51 | 1.48E-09 | 7.64E-06 |
| Igsf11 | tss_body | -2.24 | 1.59E-09 | 8.08E-06 |
| Plet1os | tss_body | -2.25 | 1.62E-09 | 8.10E-06 |
| A730004F24Rik | tss_body | 1.94 | 1.70E-09 | 8.35E-06 |
| Dlg2 | tss_body | 2.38 | 2.00E-09 | 9.35E-06 |
| NA | NA | -3.09 | 2.01E-09 | 9.35E-06 |
| Camk2d | tss_body | -2.02 | 2.01E-09 | 9.35E-06 |
| Kat6b | tss_body | -1.44 | 2.04E-09 | 9.35E-06 |
| LOC108168771 | tss_body | 2.18 | 2.09E-09 | 9.44E-06 |
| Mboat2 | tss_body | -3.13 | 2.22E-09 | 9.87E-06 |
| Med27 | tss_body | -1.47 | 2.27E-09 | 9.94E-06 |
| Nedd4l | tss_body | -2.18 | 2.55E-09 | 1.10E-05 |
| Gm15559 | tss_body | -1.46 | 2.94E-09 | 1.25E-05 |
| Otx1 | tss_body | 3.25 | 3.01E-09 | 1.26E-05 |
| Neurog2 | tss_body | 1.28 | 3.08E-09 | 1.27E-05 |
| chr4-99650896-99651569 | Island | 1.93 | 3.24E-09 | 1.32E-05 |
| Tmem110 | tss_body | -1.25 | 3.29E-09 | 1.32E-05 |
| LOC102635707 | tss_body | 2.2 | 3.37E-09 | 1.32E-05 |
| Cdc14a | tss_body | -2.19 | 3.40E-09 | 1.32E-05 |
| Olfr536 | tss_body | 1.81 | 3.44E-09 | 1.32E-05 |
| Lrig1 | tss_body | -2.38 | 3.48E-09 | 1.32E-05 |
| Dpp10 | tss_body | -3.13 | 3.54E-09 | 1.33E-05 |
| Mir124a-1hg | tss_1500 | 1.74 | 4.27E-09 | 1.58E-05 |
| LOC108168769 | tss_body | 3.52 | 4.42E-09 | 1.61E-05 |
| NA | tss_body | -1.64 | 4.53E-09 | 1.63E-05 |
| NA | NA | -2.46 | 4.69E-09 | 1.65E-05 |
| Ano2 | tss_body | 1.86 | 4.70E-09 | 1.65E-05 |

| | | | | |
|---|---|---|---|---|
| Rora | tss_body | 1.75 | 4.85E-09 | 1.68E-05 |
| LOC102635707 | tss_body | 2.15 | 5.08E-09 | 1.74E-05 |
| Gm34726 | tss_body | 1.85 | 5.24E-09 | 1.78E-05 |
| En2 | tss_1500 | 1.91 | 5.33E-09 | 1.78E-05 |
| NA | tss_1500 | 2.36 | 5.57E-09 | 1.83E-05 |
| NA | NA | 2.16 | 5.63E-09 | 1.83E-05 |
| Fuk | tss_body | -2.38 | 5.66E-09 | 1.83E-05 |
| Nup93 | tss_body | -2.79 | 5.74E-09 | 1.83E-05 |
| 9430019J16Rik | tss_body | 1.93 | 5.95E-09 | 1.88E-05 |
| Park2 | tss_body | -2.01 | 6.35E-09 | 1.97E-05 |
| Fam174b | tss_body | -2.22 | 6.37E-09 | 1.97E-05 |
| Tex2 | tss_body | -2.16 | 6.50E-09 | 1.99E-05 |
| NA | NA | 1.79 | 6.59E-09 | 2.00E-05 |
| Commd1 | tss_body | -1.2 | 6.76E-09 | 2.02E-05 |
| NA | NA | 1.94 | 6.92E-09 | 2.05E-05 |
| Fat1 | tss_body | -1.35 | 7.17E-09 | 2.10E-05 |
| NA | tss_1500 | 1.9 | 7.41E-09 | 2.15E-05 |
| Rora | tss_body | 2.64 | 7.50E-09 | 2.16E-05 |
| Ropn1 | tss_body | -2.97 | 7.64E-09 | 2.17E-05 |
| Ccdc125 | tss_body | -2.44 | 7.80E-09 | 2.20E-05 |
| Wipf3 | tss_body | -1.73 | 7.90E-09 | 2.20E-05 |
| NA | tss_200 | 1.63 | 7.98E-09 | 2.20E-05 |
| Gm8439 | tss_body | -1.1 | 8.03E-09 | 2.20E-05 |
| Nup93 | tss_body | -3.25 | 8.20E-09 | 2.22E-05 |
| Dlgap1 | tss_body | 2.43 | 8.43E-09 | 2.25E-05 |
| Q8VFE8 | tss_body | 2.42 | 8.48E-09 | 2.25E-05 |
| Gm32351 | tss_body | 1.5 | 8.98E-09 | 2.36E-05 |
| Intu | tss_body | -1.96 | 9.81E-09 | 2.56E-05 |
| Abca16 | tss_body | 2.17 | 1.02E-08 | 2.63E-05 |
| Fscn1 | tss_body | 2.45 | 1.04E-08 | 2.64E-05 |
| NA | NA | 1.71 | 1.05E-08 | 2.64E-05 |
| Myrip | tss_body | 1.5 | 1.05E-08 | 2.64E-05 |
| Srgap1 | tss_body | -2.34 | 1.07E-08 | 2.67E-05 |
| NA | NA | -1.44 | 1.09E-08 | 2.70E-05 |
| Gm2694 | tss_body | -1.38 | 1.11E-08 | 2.70E-05 |
| A730004F24Rik | tss_body | 2.66 | 1.11E-08 | 2.70E-05 |
| Pde6h | tss_body | -2.16 | 1.12E-08 | 2.70E-05 |
| Lhx1os | tss_body | 2.04 | 1.16E-08 | 2.78E-05 |
| Gm35727 | tss_body | 1.73 | 1.20E-08 | 2.85E-05 |
| Srm | tss_body | -1.48 | 1.26E-08 | 2.96E-05 |
| Gm41958 | tss_body | -2.08 | 1.30E-08 | 3.03E-05 |

| | | | | |
|---|---|---|---|---|
| Cldn18 | tss_body | 1.47 | 1.32E-08 | 3.04E-05 |
| Anks1 | tss_body | -1.72 | 1.32E-08 | 3.04E-05 |
| NA | NA | 2.05 | 1.33E-08 | 3.04E-05 |
| Sash1 | tss_body | -1.76 | 1.37E-08 | 3.07E-05 |
| Gm35223 | tss_body | 1.68 | 1.37E-08 | 3.07E-05 |
| Sorbs2 | tss_body | -1.85 | 1.45E-08 | 3.19E-05 |
| Hs6st1 | tss_body | 1.39 | 1.46E-08 | 3.19E-05 |
| Osbpl3 | tss_body | -1.29 | 1.47E-08 | 3.19E-05 |
| Dmbx1 | tss_body | -1.93 | 1.47E-08 | 3.19E-05 |
| Smad7 | tss_body | 1.06 | 1.50E-08 | 3.24E-05 |
| Isg20 | tss_body | -1.93 | 1.56E-08 | 3.33E-05 |
| LOC108168771 | tss_body | 2.35 | 1.57E-08 | 3.33E-05 |
| Fzd5 | tss_body | -1.43 | 1.58E-08 | 3.33E-05 |
| Fam135b | tss_body | -2.84 | 1.60E-08 | 3.34E-05 |
| Hmx2 | tss_body | 1.24 | 1.66E-08 | 3.46E-05 |
| chr13-98504157-98504424 | NShore | 1.85 | 1.69E-08 | 3.48E-05 |
| NA | NA | -1.72 | 1.71E-08 | 3.50E-05 |
| NA | NA | -1.98 | 1.83E-08 | 3.72E-05 |
| Intu | tss_body | -1.58 | 1.87E-08 | 3.78E-05 |
| NA | NA | -2.23 | 1.91E-08 | 3.82E-05 |
| Gm33293 | tss_body | 2.25 | 2.05E-08 | 4.08E-05 |
| Scn2a1 | tss_body | -2.14 | 2.07E-08 | 4.08E-05 |
| Slc18b1 | tss_body | -1.55 | 2.12E-08 | 4.13E-05 |
| LOC108169141 | tss_body | 1.79 | 2.14E-08 | 4.13E-05 |
| Gm30796 | tss_body | -1.62 | 2.14E-08 | 4.13E-05 |
| Mboat2 | tss_body | -1.58 | 2.16E-08 | 4.13E-05 |
| Nedd4l | tss_body | -1.28 | 2.16E-08 | 4.13E-05 |
| Kalrn | tss_body | -2.06 | 2.23E-08 | 4.22E-05 |
| chr1-23256018-23257054 | SShore | -2.33 | 2.36E-08 | 4.45E-05 |
| Dlx1 | tss_body | 0.79 | 2.45E-08 | 4.59E-05 |
| NA | tss_body | -1.82 | 2.48E-08 | 4.62E-05 |
| Gm41958 | tss_body | -2.45 | 2.50E-08 | 4.62E-05 |
| Utrn | tss_body | -1.52 | 2.55E-08 | 4.67E-05 |
| 4930578E11Rik | tss_body | -1.85 | 2.56E-08 | 4.67E-05 |
| Bdh1 | tss_body | -1.02 | 2.61E-08 | 4.73E-05 |
| Foxn3 | tss_body | -1.46 | 2.63E-08 | 4.74E-05 |
| 3110099E03Rik | tss_1500 | -2.3 | 2.66E-08 | 4.76E-05 |
| Gad1os | tss_body | 1.13 | 2.71E-08 | 4.81E-05 |
| chr4-124277515-124277995 | SShore | 2.09 | 2.76E-08 | 4.88E-05 |
| Mall | tss_body | -1.43 | 2.89E-08 | 5.08E-05 |
| NA | NA | 1.94 | 2.92E-08 | 5.10E-05 |

| | | | | |
|---|---|---|---|---|
| Foxb2 | tss_body | 0.86 | 2.95E-08 | 5.12E-05 |
| NA | NA | -1.77 | 2.98E-08 | 5.13E-05 |
| NA | NA | -1.71 | 3.00E-08 | 5.13E-05 |
| Pxylp1 | tss_body | -1.39 | 3.01E-08 | 5.13E-05 |
| chr9-98973077-98973428 | Island | 1.73 | 3.09E-08 | 5.24E-05 |
| NA | tss_200 | -2.9 | 3.14E-08 | 5.29E-05 |
| Prelp | tss_body | -1.55 | 3.20E-08 | 5.32E-05 |
| NA | NA | -2.67 | 3.24E-08 | 5.32E-05 |
| Commd1 | tss_body | -1.19 | 3.25E-08 | 5.32E-05 |
| NA | tss_body | 1.55 | 3.26E-08 | 5.32E-05 |
| Lrrc8e | tss_body | -1.24 | 3.27E-08 | 5.32E-05 |
| Camk2d | tss_body | -2.5 | 3.27E-08 | 5.32E-05 |
| Gm41492 | tss_body | 1.93 | 3.44E-08 | 5.56E-05 |
| Lrig1 | tss_body | -1.9 | 3.48E-08 | 5.60E-05 |
| Onecut2 | tss_1500 | 1.33 | 3.52E-08 | 5.62E-05 |
| Scn2a1 | tss_body | -2.05 | 3.71E-08 | 5.87E-05 |
| Rfx3 | tss_body | -2.64 | 3.76E-08 | 5.87E-05 |
| NA | NA | 1.55 | 3.77E-08 | 5.87E-05 |
| Zfp326 | tss_body | -1.69 | 3.78E-08 | 5.87E-05 |
| St3gal6 | tss_1500 | -2.58 | 3.78E-08 | 5.87E-05 |
| NA | tss_body | 2.5 | 3.86E-08 | 5.97E-05 |
| C430049B03Rik | tss_body | 2.54 | 3.89E-08 | 5.97E-05 |
| Tle4 | tss_body | -2.3 | 3.99E-08 | 6.10E-05 |
| Cdkl3 | tss_body | -2.38 | 4.04E-08 | 6.15E-05 |
| Slitrk6 | tss_body | -1.94 | 4.08E-08 | 6.18E-05 |
| Pou2f1 | tss_body | -1.92 | 4.13E-08 | 6.21E-05 |
| Tec | tss_body | -2.62 | 4.18E-08 | 6.26E-05 |
| Myh13 | tss_body | 1.13 | 4.21E-08 | 6.27E-05 |
| Fam170b | tss_200 | 1.93 | 4.25E-08 | 6.28E-05 |
| Tbx15 | tss_body | 2.05 | 4.26E-08 | 6.28E-05 |
| Csmd1 | tss_body | -2.34 | 4.43E-08 | 6.49E-05 |
| Cnnm1 | tss_body | 1.8 | 4.77E-08 | 6.91E-05 |
| Trpv3 | tss_1500 | -1.42 | 4.81E-08 | 6.91E-05 |
| NA | tss_body | 1.94 | 4.81E-08 | 6.91E-05 |
| Plcb1 | tss_body | -1.21 | 4.81E-08 | 6.91E-05 |
| LOC108168771 | tss_body | 2.76 | 4.87E-08 | 6.96E-05 |
| Cers6 | tss_body | -1.37 | 5.02E-08 | 7.15E-05 |
| LOC102635707 | tss_body | 1.96 | 5.20E-08 | 7.35E-05 |
| Dmrt2 | tss_body | 2.42 | 5.31E-08 | 7.45E-05 |
| Dnajc13 | tss_body | -1.4 | 5.33E-08 | 7.45E-05 |
| Q8BGX4 | tss_body | -1.51 | 5.34E-08 | 7.45E-05 |

| | | | | |
|---|---|---|---|---|
| Catsper3 | tss_body | -1.96 | 5.41E-08 | 7.51E-05 |
| D3Z790 | tss_body | 2.65 | 5.56E-08 | 7.67E-05 |
| Rora | tss_body | 2.2 | 5.58E-08 | 7.67E-05 |
| Commd1 | tss_body | -0.94 | 5.67E-08 | 7.75E-05 |
| Lhfpl3 | tss_body | 2.3 | 5.69E-08 | 7.75E-05 |
| NA | NA | -1.89 | 5.80E-08 | 7.84E-05 |
| NA | tss_1500 | -1.19 | 5.83E-08 | 7.84E-05 |
| Adgrd1 | tss_body | 1.61 | 5.84E-08 | 7.84E-05 |
| C430049B03Rik | tss_body | 1.66 | 6.00E-08 | 8.02E-05 |
| NA | tss_body | -1.16 | 6.09E-08 | 8.10E-05 |
| Osbpl3 | tss_body | -1.37 | 6.19E-08 | 8.20E-05 |
| chr8-90828686-90828982 | NShelf | -1.49 | 6.26E-08 | 8.25E-05 |
| Rictor | tss_body | -1.27 | 6.30E-08 | 8.26E-05 |
| Gas7 | tss_body | -1.93 | 6.39E-08 | 8.34E-05 |
| NA | NA | 1.5 | 6.48E-08 | 8.42E-05 |
| Msrb3 | tss_body | -2.24 | 6.57E-08 | 8.46E-05 |
| E9QMD3 | tss_body | 1.79 | 6.57E-08 | 8.46E-05 |
| NA | tss_body | -1.44 | 6.60E-08 | 8.46E-05 |
| Tle4 | tss_body | -1.61 | 6.64E-08 | 8.47E-05 |
| Hs2st1 | tss_body | -1.43 | 6.76E-08 | 8.59E-05 |
| NA | NA | -2.61 | 6.89E-08 | 8.72E-05 |
| Sfrp5 | tss_body | -3.51 | 6.99E-08 | 8.80E-05 |
| Ntn4 | tss_1500 | -1.62 | 7.08E-08 | 8.87E-05 |
| Wscd2 | tss_body | 1.47 | 7.45E-08 | 9.30E-05 |
| NA | NA | -1.51 | 7.52E-08 | 9.34E-05 |
| Gm30132 | tss_body | 1.41 | 7.67E-08 | 9.47E-05 |
| Gm31816 | tss_body | -1.79 | 7.69E-08 | 9.47E-05 |
| Q8VGW6 | tss_1500 | -1.92 | 7.72E-08 | 9.47E-05 |
| NA | NA | 2.62 | 7.87E-08 | 9.61E-05 |
| NA | NA | -1.48 | 7.94E-08 | 9.62E-05 |
| Tsc1 | tss_body | -1.33 | 7.96E-08 | 9.62E-05 |
| Thrb | tss_body | -1.05 | 7.98E-08 | 9.62E-05 |
| NA | NA | -1.76 | 8.08E-08 | 9.67E-05 |
| Bnc2 | tss_body | 1.91 | 8.09E-08 | 9.67E-05 |
| Anks1 | tss_body | -1.57 | 8.16E-08 | 9.71E-05 |
| Gng2 | tss_body | -3.07 | 8.21E-08 | 9.73E-05 |
| NA | NA | -1.22 | 8.29E-08 | 9.79E-05 |
| Gm30296 | tss_body | -1.58 | 8.45E-08 | 9.94E-05 |
| Peak1 | tss_body | -3.1 | 8.56E-08 | 0.00010018 |
| Scn2a1 | tss_body | -2.14 | 8.62E-08 | 0.000100463 |
| Foxd2os | tss_body | -2.66 | 8.69E-08 | 0.000100859 |

| | | | | |
|---|---|---|---|---|
| Gm41492 | tss_body | 2.07 | 8.74E-08 | 0.000101133 |
| NA | NA | -1.57 | 8.82E-08 | 0.000101642 |
| Sim1 | tss_1500 | 1.19 | 9.16E-08 | 0.000104394 |
| Rfx3 | tss_body | -2.75 | 9.19E-08 | 0.000104394 |
| Nebl | tss_body | -2.04 | 9.19E-08 | 0.000104394 |
| Otp | tss_body | 1.16 | 9.21E-08 | 0.000104394 |
| Kalrn | tss_body | -1.96 | 9.39E-08 | 0.000105987 |
| Slc7a2 | tss_body | -2.45 | 9.59E-08 | 0.000107769 |
| NA | tss_200 | 2.01 | 9.62E-08 | 0.000107769 |
| NA | NA | 2.61 | 9.76E-08 | 0.000108784 |
| Phc1 | tss_body | -1.18 | 9.81E-08 | 0.000108784 |
| En2 | tss_1500 | 1.42 | 9.83E-08 | 0.000108784 |
| Tmem245 | tss_body | -2 | 9.88E-08 | 0.000108784 |
| NA | NA | 2.27 | 9.90E-08 | 0.000108784 |
| NA | tss_body | 2 | 1.01E-07 | 0.000110315 |
| NA | NA | -2.5 | 1.02E-07 | 0.000110774 |
| Ncald | tss_body | -2.1 | 1.03E-07 | 0.000110774 |
| Klk4 | tss_body | 1.44 | 1.03E-07 | 0.000110774 |
| NA | NA | 2.91 | 1.03E-07 | 0.000110774 |
| NA | NA | -2.8 | 1.08E-07 | 0.000115026 |
| NA | NA | 2.21 | 1.08E-07 | 0.000115026 |
| NA | NA | 1.72 | 1.08E-07 | 0.000115026 |
| Cntn1 | tss_body | 2.5 | 1.08E-07 | 0.000115026 |
| Commd1 | tss_body | -1.28 | 1.09E-07 | 0.000115646 |
| Ccdc39 | tss_body | -2.92 | 1.10E-07 | 0.000115704 |
| Peak1 | tss_body | -2.74 | 1.17E-07 | 0.000122316 |
| Sorbs1 | tss_body | -1.84 | 1.18E-07 | 0.000123372 |
| Gtsf1 | tss_body | 1.08 | 1.23E-07 | 0.000127475 |
| NA | NA | 1.61 | 1.23E-07 | 0.000127475 |
| Col22a1 | tss_body | -1.98 | 1.26E-07 | 0.000130458 |
| Rictor | tss_body | -2.12 | 1.28E-07 | 0.000131467 |
| NA | tss_body | -1.77 | 1.29E-07 | 0.000131467 |
| Gspt2 | tss_body | -2.13 | 1.29E-07 | 0.000131467 |
| Gad1os | tss_body | 1.52 | 1.29E-07 | 0.000131467 |
| Lpar1 | tss_body | 1.65 | 1.30E-07 | 0.000132598 |
| Q8VFQ1 | tss_1500 | -1.41 | 1.32E-07 | 0.000133485 |
| Gm36568 | tss_body | -1.79 | 1.32E-07 | 0.000133485 |
| B4galt2 | tss_1500 | -0.85 | 1.35E-07 | 0.000134872 |
| Bcl11a | tss_body | 2.3 | 1.36E-07 | 0.000134872 |
| Arntl | tss_body | -1.97 | 1.36E-07 | 0.000134872 |
| Gm35988 | tss_body | 1.83 | 1.36E-07 | 0.000134872 |

| | | | | |
|---|---|---|---|---|
| Stx8 | tss_body | -2.07 | 1.36E-07 | 0.000134872 |
| Dlx2 | tss_body | 1.23 | 1.38E-07 | 0.000136125 |
| Ano2 | tss_body | 1.6 | 1.39E-07 | 0.000136125 |
| Gm35062 | tss_body | 1.73 | 1.39E-07 | 0.000136125 |
| Ano2 | tss_body | 1.84 | 1.48E-07 | 0.000145107 |
| NA | NA | -1.83 | 1.52E-07 | 0.000148035 |
| NA | tss_body | 1.45 | 1.53E-07 | 0.000148097 |
| Commd1 | tss_body | -0.76 | 1.53E-07 | 0.000148467 |
| Dpp10 | tss_body | -2.52 | 1.56E-07 | 0.000150396 |
| Wscd2 | tss_body | 1.73 | 1.57E-07 | 0.000151177 |
| Gm41492 | tss_body | 1.17 | 1.58E-07 | 0.000151177 |
| Lhx2 | tss_body | 0.97 | 1.58E-07 | 0.000151177 |
| Gm28651 | tss_body | 2 | 1.62E-07 | 0.000154389 |
| Pax6os1 | tss_body | 1.15 | 1.63E-07 | 0.000154399 |
| NA | NA | 1.93 | 1.64E-07 | 0.000154575 |
| NA | tss_body | 1.55 | 1.67E-07 | 0.000156651 |
| NA | NA | -1.52 | 1.67E-07 | 0.000156651 |
| NA | tss_body | 1.97 | 1.72E-07 | 0.00016099 |
| Gm38585 | tss_body | -1.46 | 1.77E-07 | 0.000164773 |
| E9QAY5 | tss_body | -1.27 | 1.77E-07 | 0.000164773 |
| Tle4 | tss_body | -1.7 | 1.78E-07 | 0.000164914 |
| Kcnma1 | tss_body | 1.98 | 1.79E-07 | 0.000164969 |
| Dmrt2 | tss_200 | 1.27 | 1.81E-07 | 0.000166165 |
| Abcd2 | tss_body | 2.13 | 1.82E-07 | 0.000166165 |
| 4930474G06Rik | tss_body | -1.5 | 1.82E-07 | 0.000166165 |
| Gm884 | tss_body | 1.18 | 1.83E-07 | 0.000166165 |
| LOC102635707 | tss_body | 2.85 | 1.83E-07 | 0.000166165 |
| Lrig1 | tss_body | -1.71 | 1.83E-07 | 0.000166165 |
| NA | NA | -2.13 | 1.85E-07 | 0.000166981 |
| Capn5 | tss_body | -0.82 | 1.87E-07 | 0.000168368 |
| Pde4d | tss_body | -1.75 | 1.90E-07 | 0.000169745 |
| Sdcbp2 | tss_body | -1.62 | 1.90E-07 | 0.000169745 |
| Zic5 | tss_body | -2.27 | 1.90E-07 | 0.000169798 |
| NA | NA | 1.52 | 1.92E-07 | 0.000170076 |
| NA | NA | 1.87 | 1.92E-07 | 0.000170076 |
| NA | NA | -2.58 | 1.93E-07 | 0.000170284 |
| NA | NA | 2.76 | 1.94E-07 | 0.000170769 |
| NA | NA | 2.21 | 1.96E-07 | 0.000171548 |
| Cntn1 | tss_body | 2.4 | 1.96E-07 | 0.000171548 |
| Lhx2 | tss_body | 1.39 | 1.97E-07 | 0.000171604 |
| Prkaa2 | tss_body | -1.52 | 1.99E-07 | 0.000172787 |

| | | | | |
|---|---|---|---|---|
| Nedd4l | tss_body | -2.07 | 1.99E-07 | 0.000173027 |
| Vax2os | tss_body | 1.1 | 2.00E-07 | 0.000173253 |
| NA | tss_200 | -2.02 | 2.01E-07 | 0.000173493 |
| Cdh7 | tss_body | 1.94 | 2.03E-07 | 0.000174517 |
| NA | tss_1500 | 2.48 | 2.04E-07 | 0.000174517 |
| NA | NA | 1.62 | 2.05E-07 | 0.000174868 |
| Osr2 | tss_body | 1.07 | 2.07E-07 | 0.000176067 |
| Onecut2 | tss_1500 | 1.31 | 2.12E-07 | 0.000180405 |
| Cog5 | tss_body | -3.23 | 2.16E-07 | 0.000182948 |
| St3gal3 | tss_1500 | 2.86 | 2.20E-07 | 0.000185256 |
| Pebp4 | tss_body | -1.82 | 2.20E-07 | 0.000185256 |
| Pygl | tss_body | -2.13 | 2.24E-07 | 0.000188332 |
| Rap1gap2 | tss_body | -1.56 | 2.26E-07 | 0.000188614 |
| Otx1 | tss_body | 2.2 | 2.26E-07 | 0.000188614 |
| Q7TRJ3 | tss_body | -1.58 | 2.27E-07 | 0.00018905 |
| NA | NA | 1.97 | 2.29E-07 | 0.000189955 |
| chr13-98502955-98503214 | SShore | 1.35 | 2.31E-07 | 0.000190817 |
| C430049B03Rik | tss_body | 1.9 | 2.35E-07 | 0.000193736 |
| Tmed8 | tss_body | -1.53 | 2.37E-07 | 0.000194474 |
| Aff3 | tss_body | -1.26 | 2.38E-07 | 0.000194629 |
| Ccl27a | tss_body | 0.94 | 2.38E-07 | 0.000194629 |
| Gm33293 | tss_body | 1.31 | 2.44E-07 | 0.000198443 |
| NA | NA | -1.25 | 2.44E-07 | 0.000198443 |
| Olfr122 | tss_body | -1.72 | 2.46E-07 | 0.000199378 |
| Ptpdc1 | tss_body | -4.19 | 2.47E-07 | 0.000199589 |
| NA | NA | -1.08 | 2.49E-07 | 0.000199836 |
| LOC102635707 | tss_body | 1.95 | 2.49E-07 | 0.000199836 |
| NA | tss_200 | -1.01 | 2.49E-07 | 0.000199836 |
| NA | tss_body | 2.26 | 2.53E-07 | 0.000201918 |
| R3hdml | tss_body | -1.79 | 2.53E-07 | 0.000201918 |
| Ces1c | tss_body | 0.91 | 2.55E-07 | 0.000202468 |
| Fam46c | tss_body | -1.16 | 2.57E-07 | 0.000203511 |
| Stx8 | tss_body | -1.5 | 2.64E-07 | 0.000208687 |
| NA | NA | -1.81 | 2.66E-07 | 0.000208721 |
| NA | tss_1500 | -1.1 | 2.66E-07 | 0.000208721 |
| Prdm12 | tss_body | 1.79 | 2.67E-07 | 0.000208721 |
| NA | NA | -2.29 | 2.68E-07 | 0.000208721 |
| Ly6g6d | tss_body | -1.89 | 2.69E-07 | 0.000208721 |
| Dpp10 | tss_body | -2.74 | 2.69E-07 | 0.000208721 |
| Dcaf4 | tss_body | -2.33 | 2.69E-07 | 0.000208721 |
| Gm35623 | tss_body | 1.76 | 2.73E-07 | 0.00021143 |

| | | | | |
|---|---|---|---|---|
| Plpp3 | tss_body | -1.35 | 2.81E-07 | 0.000215738 |
| NA | NA | -1.39 | 2.81E-07 | 0.000215738 |
| NA | tss_body | 1.49 | 2.82E-07 | 0.000215738 |
| Cftr | tss_body | -2.87 | 2.82E-07 | 0.000215738 |
| Kcnma1 | tss_body | 1.68 | 2.83E-07 | 0.000215738 |
| NA | NA | -1.61 | 2.84E-07 | 0.000215738 |
| Gsg1l2 | tss_body | 0.89 | 2.85E-07 | 0.000216428 |
| chr5-139546513-139547812 | SShore | 1.58 | 2.86E-07 | 0.000216428 |
| Dlx4os | tss_body | -0.89 | 2.88E-07 | 0.000216637 |
| Acss3 | tss_body | -1.4 | 2.88E-07 | 0.000216637 |
| Cacnb2 | tss_body | -1.36 | 2.89E-07 | 0.000216637 |
| Vax1 | tss_body | 1.06 | 2.90E-07 | 0.000216862 |
| Arhgef28 | tss_body | -1.55 | 2.92E-07 | 0.000217349 |
| Cdh13 | tss_body | -2.09 | 2.92E-07 | 0.000217349 |
| Tmem154 | tss_1500 | -1.69 | 2.93E-07 | 0.000217349 |
| Gm32385 | tss_body | 1.16 | 2.94E-07 | 0.000217615 |
| Bcar3 | tss_body | -1.92 | 2.96E-07 | 0.000218394 |
| NA | NA | -1.55 | 2.96E-07 | 0.000218394 |
| chr13-98365821-98366560 | Island | 1.54 | 3.04E-07 | 0.000223187 |
| A930003O13Rik | tss_body | 2.49 | 3.07E-07 | 0.000224825 |
| Gm35585 | tss_body | -1.94 | 3.09E-07 | 0.000225407 |
| Klhl3 | tss_body | -1.58 | 3.09E-07 | 0.000225407 |
| Myocd | tss_body | 1 | 3.10E-07 | 0.000225493 |
| Q64378 | tss_body | -1.33 | 3.11E-07 | 0.000225493 |
| NA | NA | -1.61 | 3.14E-07 | 0.000226862 |
| Cog5 | tss_body | -1.66 | 3.14E-07 | 0.000226862 |
| NA | tss_1500 | 1.5 | 3.27E-07 | 0.000235487 |
| Foxd2 | tss_body | -1.91 | 3.28E-07 | 0.000235666 |
| LOC102635707 | tss_body | 1.83 | 3.30E-07 | 0.000236481 |
| Plb1 | tss_body | -0.9 | 3.31E-07 | 0.000236596 |
| Nfam1 | tss_body | -1.21 | 3.34E-07 | 0.000237479 |
| 2900026A02Rik | tss_body | 2.21 | 3.34E-07 | 0.000237479 |
| Creb5 | tss_body | 1.84 | 3.37E-07 | 0.000238632 |
| Gm34055 | tss_body | 2.01 | 3.37E-07 | 0.000238632 |
| Prkce | tss_body | -1.51 | 3.39E-07 | 0.000239394 |
| Dlx2 | tss_body | 1.47 | 3.42E-07 | 0.00024094 |
| Ccr6 | tss_body | 1.98 | 3.44E-07 | 0.000241981 |
| NA | NA | -2.03 | 3.47E-07 | 0.000243388 |
| NA | NA | -2.1 | 3.50E-07 | 0.000244497 |
| NA | NA | 1.63 | 3.51E-07 | 0.000244497 |
| Klhl2 | tss_body | -1.08 | 3.52E-07 | 0.000244647 |

| | | | | |
|---|---|---|---|---|
| Hnf4aos | tss_body | -1.5 | 3.55E-07 | 0.000245929 |
| NA | NA | -2.23 | 3.56E-07 | 0.000245929 |
| Nkx2-1 | tss_body | 1.27 | 3.56E-07 | 0.000245929 |
| D430041D05Rik | tss_body | -1.3 | 3.60E-07 | 0.00024783 |
| Prkci | tss_body | -1.67 | 3.63E-07 | 0.000249192 |
| Prkch | tss_body | -1.43 | 3.64E-07 | 0.000249192 |
| Tet3 | tss_body | -1.73 | 3.65E-07 | 0.000249192 |
| Serp2 | tss_body | -1.38 | 3.65E-07 | 0.000249192 |
| Hs6st2 | tss_body | 1.38 | 3.72E-07 | 0.000253006 |
| Ago4 | tss_body | -1.37 | 3.73E-07 | 0.000253006 |
| Lrig1 | tss_body | -1.71 | 3.75E-07 | 0.000254176 |
| 4930578E11Rik | tss_body | -1.84 | 3.79E-07 | 0.000256209 |
| Auh | tss_body | -1.41 | 3.87E-07 | 0.00026072 |
| Nr0b1 | tss_200 | 2.25 | 3.88E-07 | 0.000261274 |
| Hcar2 | tss_body | -1.74 | 3.94E-07 | 0.000263744 |
| Acly | tss_body | -1.18 | 3.94E-07 | 0.000263744 |
| NA | NA | -1.25 | 3.95E-07 | 0.000264079 |
| Rap1a | tss_body | -1.45 | 3.99E-07 | 0.000265483 |
| Vstm4 | tss_body | 2.07 | 4.00E-07 | 0.000265483 |
| Il1rn | tss_200 | -1.64 | 4.01E-07 | 0.000265483 |
| Smim14 | tss_body | -1.88 | 4.01E-07 | 0.000265483 |
| Immp2l | tss_body | -1.29 | 4.06E-07 | 0.00026783 |
| Nedd4l | tss_body | -1.91 | 4.07E-07 | 0.00026783 |
| Gm15559 | tss_body | -1.25 | 4.08E-07 | 0.00026783 |
| Myo1c | tss_body | -1.22 | 4.09E-07 | 0.00026783 |
| NA | NA | 2.2 | 4.11E-07 | 0.000268626 |
| Ly6d | tss_1500 | -1.55 | 4.12E-07 | 0.000268901 |
| NA | NA | -2.22 | 4.18E-07 | 0.000272437 |
| NA | tss_body | -1.61 | 4.22E-07 | 0.000273591 |
| Nxf7 | tss_body | 2.57 | 4.22E-07 | 0.000273591 |
| Commd1 | tss_body | -0.94 | 4.28E-07 | 0.000276778 |
| NA | tss_body | 1.94 | 4.30E-07 | 0.000277575 |
| NA | NA | -1.41 | 4.35E-07 | 0.000280237 |
| Dpp10 | tss_body | -1.64 | 4.39E-07 | 0.000282173 |
| Zfp385b | tss_body | -1.37 | 4.41E-07 | 0.000282348 |
| NA | NA | 2.17 | 4.47E-07 | 0.000286093 |
| NA | tss_200 | 1.75 | 4.49E-07 | 0.000286141 |
| Gm2694 | tss_body | -1.72 | 4.54E-07 | 0.000288939 |
| Ablim1 | tss_body | -1.5 | 4.55E-07 | 0.000289203 |
| NA | NA | -1.51 | 4.61E-07 | 0.000292159 |
| 2900026A02Rik | tss_body | 2.87 | 4.64E-07 | 0.000293672 |

| | | | | |
|---|---|---|---|---|
| Ccl27a | tss_body | 0.76 | 4.70E-07 | 0.000296518 |
| Otop1 | tss_body | 1.8 | 4.73E-07 | 0.00029722 |
| Dbndd1 | tss_body | -1.33 | 4.76E-07 | 0.00029722 |
| Lhx1os | tss_body | 2.16 | 4.77E-07 | 0.00029722 |
| Fam170b | tss_200 | 1.81 | 4.77E-07 | 0.00029722 |
| Gm17576 | tss_1500 | 1.3 | 4.78E-07 | 0.00029722 |
| NA | NA | 1.7 | 4.79E-07 | 0.00029722 |
| Gsx1 | tss_1500 | 2.63 | 4.79E-07 | 0.00029722 |
| NA | NA | 2.18 | 4.80E-07 | 0.00029722 |
| Prkca | tss_body | -1.33 | 4.81E-07 | 0.00029722 |
| NA | NA | -2.1 | 4.82E-07 | 0.00029722 |
| Lhx4 | tss_body | 0.81 | 4.90E-07 | 0.000301453 |
| Ccr6 | tss_body | 1.71 | 4.92E-07 | 0.000302628 |
| Klk8 | tss_body | 1.47 | 4.99E-07 | 0.000305774 |
| Ccl27a | tss_body | 0.94 | 5.00E-07 | 0.000305774 |
| Ablim1 | tss_body | -1.97 | 5.08E-07 | 0.00031036 |
| Pard3 | tss_body | -0.86 | 5.12E-07 | 0.000312175 |
| Gm35201 | tss_body | -1.62 | 5.17E-07 | 0.000314087 |
| Pomp | tss_body | -1.39 | 5.23E-07 | 0.000317003 |
| Gng2 | tss_1500 | -1.55 | 5.27E-07 | 0.00031923 |
| Arl15 | tss_body | -1.04 | 5.48E-07 | 0.000331073 |
| Nlgn1 | tss_body | -2 | 5.50E-07 | 0.000331468 |
| Asb4 | tss_body | 1.77 | 5.51E-07 | 0.000331468 |
| Kcnma1 | tss_body | 1.88 | 5.52E-07 | 0.000331472 |
| NA | NA | 1.99 | 5.54E-07 | 0.000332086 |
| Mcc | tss_body | -2.97 | 5.62E-07 | 0.000336202 |
| Myrip | tss_body | 1.34 | 5.67E-07 | 0.000336538 |
| Ptpra | tss_body | -0.78 | 5.68E-07 | 0.000336538 |
| Commd1 | tss_body | -1.07 | 5.68E-07 | 0.000336538 |
| NA | tss_body | 2.02 | 5.69E-07 | 0.000336538 |
| Gm35426 | tss_body | 1.9 | 5.69E-07 | 0.000336538 |
| Nsf | tss_body | -1.22 | 5.73E-07 | 0.000338014 |
| Ikzf1 | tss_body | 0.91 | 5.75E-07 | 0.000338014 |
| NA | NA | -1.47 | 5.75E-07 | 0.000338014 |
| Yae1d1 | tss_body | -1.63 | 5.85E-07 | 0.000342794 |
| LOC102632594 | tss_body | -2.38 | 5.86E-07 | 0.000342794 |
| Gm5089 | tss_body | 1.11 | 5.87E-07 | 0.000342794 |
| Tspan12 | tss_body | -1.4 | 5.88E-07 | 0.000342794 |
| Pdgfra | tss_body | 1 | 5.98E-07 | 0.000347461 |
| Gm26728 | tss_body | -1.2 | 5.98E-07 | 0.000347461 |
| Zbtb38 | tss_body | -1.88 | 6.01E-07 | 0.000348216 |

| | | | | |
|---|---|---|---|---|
| Mipep | tss_body | -1.09 | 6.08E-07 | 0.00035113 |
| Gpc5 | tss_body | -1.72 | 6.09E-07 | 0.00035113 |
| Gm35223 | tss_1500 | 1.71 | 6.10E-07 | 0.00035113 |
| Bsx | tss_body | 1 | 6.14E-07 | 0.000352173 |
| Gm34529 | tss_1500 | -1.16 | 6.14E-07 | 0.000352173 |
| Rock2 | tss_body | -1.02 | 6.15E-07 | 0.000352173 |
| Olfr530 | tss_body | 1.69 | 6.23E-07 | 0.000355768 |
| NA | NA | -2.63 | 6.24E-07 | 0.000355768 |

Table S7: DNA Methylation DMR DKO vs SKO Top500

| Gene Name | meanbetafc | minfdr | Stouffer | Annotation | Distance to TSS |
|---|---|---|---|---|---|
| Gm13298 | -0.156825841 | 0 | 0 | exon (NM_001085530, exon 4 of 4).2 | 3508 |
| Gm13298 | -0.177604672 | 1.50E-164 | 4.90E-74 | intron (NR_033506, intron 1 of 3) | -10023 |
| Ccdc58 | 0.155198744 | 2.73E-42 | 4.09E-10 | intron (NM_001159421, intron 2 of 3) | 7809 |
| Zfp629 | 0.235557529 | 1.16E-21 | 0.008062051 | Intergenic | -38304 |
| Scgb3a2 | -0.143717351 | 2.64E-09 | 7.04E-06 | promoter-TSS (NM_001289644) | -45 |
| Hrasls | 0.15282732 | 2.64E-09 | 3.74E-05 | promoter-TSS (NM_013751) | -236 |
| Mir190b | -0.141463869 | 2.64E-09 | 0.000346965 | promoter-TSS (NR_030543) | -164 |
| Tuba1a | -0.341443839 | 2.65E-09 | 3.77E-06 | Intergenic | -1056 |
| Cped1 | 0.291846547 | 2.67E-09 | 1.37E-05 | intron (NM_001081351, intron 8 of 21) | 136044 |
| Nkx6-2 | -0.279259824 | 2.67E-09 | 1.39E-05 | intron (NM_001127363, intron 8 of 15) | 54039 |
| Gm20362 | -0.224015208 | 2.68E-09 | 4.58E-06 | Intergenic | 356656 |
| Gm525 | 0.216624699 | 2.68E-09 | 1.45E-05 | exon (NM_001033266, exon 2 of 4) | 1210 |
| Arl10 | 0.126404035 | 2.68E-09 | 1.94E-05 | intron (NM_019968, intron 2 of 3) | 3117 |
| Ror2 | 0.204244602 | 1.17E-21 | 4.10E-13 | intron (NM_013846, intron 1 of 8) | 107745 |
| Bmp7 | -0.203468064 | 2.69E-09 | 6.46E-06 | Intergenic | 183932 |
| Cyp26c1 | 0.129363176 | 2.70E-09 | 1.28E-05 | intron (NM_175353, intron 20 of 21) | -28727 |
| Col6a1 | -0.104375881 | 2.71E-09 | 1.95E-05 | exon (NM_009933, exon 32 of 35) | 14710 |
| Isca1 | 0.121682771 | 2.71E-09 | 4.95E-05 | intron (NM_026921, intron 1 of 3) | 3196 |
| Slc19a3 | -0.13736601 | 2.71E-09 | 0.105758007 | intron (NM_030556, intron 1 of 6) | 1428 |
| Arhgap36 | -0.124499485 | 2.72E-09 | 0.001071337 | intron (NM_001081123, intron 1 of 12) | 7524 |
| 3300005D01Rik | 0.188696811 | 2.73E-09 | 3.94E-09 | promoter-TSS (NR_045079) | 414 |
| Mir6340 | -0.433359956 | 2.73E-09 | 1.32E-06 | promoter-TSS (NR_105758) | 39 |
| Mir192 | 0.135522137 | 1.20E-21 | 1.91E-10 | promoter-TSS (NR_029720) | -20 |
| Zmiz1 | 0.169337599 | 2.73E-09 | 3.98E-06 | Intergenic | -80028 |
| Sfpq | 0.155406586 | 2.73E-09 | 4.56E-06 | Intergenic | -20537 |
| Max | -0.225079841 | 2.73E-09 | 5.63E-06 | intron (NM_008558, intron 3 of 4) | 17930 |
| Tbl1xr1 | -0.225927557 | 2.73E-09 | 1.27E-05 | intron (NM_030732, intron 1 of 15) | 37824 |
| 5430427O19Rik | -0.140066822 | 2.74E-09 | 2.82E-06 | Intergenic | -5838 |
| Ksr2 | 0.196050294 | 2.74E-09 | 2.55E-05 | intron (NM_001114545, intron 4 of 19) | 200564 |
| Thsd4 | 0.184230978 | 2.74E-09 | 2.74E-05 | intron (NM_001040426, intron 7 of 17) | 1388 |
| Tbrg4 | -0.143421221 | 2.75E-09 | 0.003415952 | promoter-TSS (NM_134011) | -497 |
| Snord1a | 0.218072318 | 1.20E-21 | 1.32E-08 | promoter-TSS (NR_028570) | -158 |
| Cnga1 | -0.167224644 | 2.76E-09 | 1.95E-07 | promoter-TSS (NM_007723) | -460 |
| Eng | 0.153367452 | 2.76E-09 | 0.038746695 | promoter-TSS (NM_007932) | -160 |
| Lyplal1 | -0.296058706 | 2.77E-09 | 7.73E-06 | Intergenic | -136996 |
| Acsm2 | -0.206214774 | 2.77E-09 | 2.30E-05 | promoter-TSS (NM_001177977) | -257 |
| Arhgef10l | -0.191921323 | 2.78E-09 | 1.31E-06 | intron (NM_001112722, intron 1 of 27) | 389 |
| Stat5b | 0.255517345 | 2.78E-09 | 3.38E-06 | intron (NM_001113563, intron 1 of 18) | -3880 |
| Bcl3 | 0.253205298 | 2.78E-09 | 8.06E-06 | Intergenic | -4164 |
| Orc6 | -0.120322391 | 2.78E-09 | 7.77E-05 | promoter-TSS (NM_022997) | 731 |
| 1700020N15Rik | 0.251719537 | 1.23E-21 | 1.33E-08 | intron (NR_033213, intron 3 of 8) | 62082 |
| Cdh10 | -0.262170597 | 2.79E-09 | 3.99E-05 | Intergenic | 574956 |
| Smarca1 | 0.177902467 | 2.80E-09 | 0.000538749 | promoter-TSS (NM_053123) | -195 |
| Rhbdd3 | 0.103097722 | 2.81E-09 | 1.79E-05 | intron (NM_001290492, intron 3 of 8) | 1305 |
| Med7 | -0.177540052 | 2.81E-09 | 2.80E-05 | 5' UTR (NM_001104530, exon 2 of 3) | 2858 |

| | | | | | |
|---|---|---|---|---|---|
| Scgb2b27 | -0.265416382 | 2.81E-09 | 0.000138856 | Intergenic | 16928 |
| Kdelr2 | -0.16589936 | 2.81E-09 | 0.000185053 | intron (NM_001159321, intron 16 of 21) | -16071 |
| Ccm2 | 0.214872812 | 2.82E-09 | 1.51E-06 | promoter-TSS (NM_001190344) | -707 |
| Mir7242 | 0.128323968 | 2.82E-09 | 0.000159498 | promoter-TSS (NR_106101) | -271 |
| a | 0.313170018 | 1.26E-21 | 1.87E-08 | intron (NM_015770, intron 1 of 3) | 5187 |
| Ranbp3 | -0.199349992 | 2.83E-09 | 5.71E-06 | intron (NR_045519, intron 2 of 16) | 18766 |
| Mgat4e | 0.101409948 | 2.83E-09 | 7.40E-06 | promoter-TSS (NM_001145300) | -546 |
| Lrrtm1 | 0.346842845 | 2.83E-09 | 1.30E-05 | intron (NM_001109764, intron 12 of 19) | -263977 |
| Eef1e1 | -0.156981109 | 2.84E-09 | 3.56E-06 | Intergenic | -134665 |
| Gm6260 | 0.174788439 | 2.84E-09 | 8.67E-06 | Intergenic | 61984 |
| 1700030F04Rik | -0.352294188 | 2.85E-09 | 6.47E-06 | Intergenic | 139068 |
| Lypd2 | 0.271908296 | 2.85E-09 | 9.86E-06 | Intergenic | -1555 |
| Cpa4 | 0.108114062 | 2.85E-09 | 5.44E-05 | intron (NM_027926, intron 1 of 10) | 652 |
| Ube2v1 | -0.240372574 | 2.86E-09 | 1.24E-06 | intron (NM_023230, intron 1 of 3) | 2797 |
| Tspan5 | -0.377838193 | 2.86E-09 | 1.56E-06 | intron (NR_121195, intron 1 of 7) | 67486 |
| Vrtn | 0.179531334 | 2.86E-09 | 6.97E-06 | promoter-TSS (NM_001168589) | -268 |
| Prss44 | -0.156901401 | 2.86E-09 | 1.57E-05 | exon (NM_148940, exon 1 of 5) | 223 |
| Zfp488 | 0.117392166 | 2.87E-09 | 5.18E-05 | exon (NM_001013777, exon 2 of 2) | 8176 |
| Klf11 | -0.228277521 | 2.88E-09 | 6.50E-06 | Intergenic | -9397 |
| Fam78a | 0.166932133 | 1.29E-21 | 3.60E-08 | exon (NM_175511, exon 2 of 2) | 14275 |
| Ssx2ip | -0.346622456 | 2.90E-09 | 6.54E-06 | intron (NM_001253768, intron 5 of 14) | 17692 |
| Txk | 0.270596795 | 2.91E-09 | 6.52E-06 | promoter-TSS (NM_001122754) | 1 |
| Hmcn1 | -0.116333196 | 2.93E-09 | 1.44E-05 | Intergenic | -8067 |
| Egfl8 | 0.132043462 | 2.94E-09 | 4.05E-07 | promoter-TSS (NM_152922) | -483 |
| Mbnl2 | -0.230942857 | 2.94E-09 | 2.20E-05 | Intergenic | -80341 |
| Ccdc73 | -0.201617494 | 2.94E-09 | 0.000445101 | 5' UTR (NM_177600, exon 1 of 18) | 144 |
| LOC102634101 | 0.187202047 | 1.31E-21 | 2.42E-11 | Intergenic | -82451 |
| Optc | 0.202476817 | 2.95E-09 | 6.47E-07 | Intergenic | 16378 |
| Ldlrad4 | 0.216677812 | 2.96E-09 | 6.89E-06 | intron (NM_172631, intron 1 of 5) | 102094 |
| Olfr1110 | -0.249484495 | 2.96E-09 | 8.04E-06 | Intergenic | 11998 |
| Phka2 | 0.114322169 | 2.97E-09 | 2.26E-07 | Intergenic | 99434 |
| Tas2r130 | -0.191106789 | 2.97E-09 | 1.28E-06 | exon (NM_199156, exon 1 of 1) | 116 |
| Olfr1364 | -0.270092044 | 2.97E-09 | 6.12E-06 | exon (NM_146540, exon 1 of 1) | 192 |
| Lrfn4 | 0.151794234 | 2.97E-09 | 0.000206143 | intron (NM_001162946, intron 14 of 20) | -3073 |
| Mir667 | -0.309820074 | 1.32E-21 | 6.69E-08 | TTS (NR_030426) | 194 |
| 3110009F21Rik | -0.202708344 | 2.98E-09 | 3.64E-06 | Intergenic | 185221 |
| Gm2176 | 0.206155705 | 2.99E-09 | 1.90E-07 | intron (NM_198429, intron 9 of 9) | -67113 |
| Snora81 | -0.154679307 | 3.00E-09 | 1.28E-08 | TTS (NR_034048) | 453 |
| Agmat | 0.119447642 | 3.00E-09 | 4.06E-06 | exon (NM_001081408, exon 1 of 7) | 208 |
| Mink1 | 0.355480253 | 3.01E-09 | 6.95E-06 | intron (NM_176893, intron 1 of 31) | 24196 |
| Upp1 | 0.119062216 | 3.01E-09 | 0.000339759 | TTS (NM_001159402) | 17813 |
| Alpl | 0.113518416 | 3.02E-09 | 7.85E-06 | Intergenic | -4369 |
| Tbc1d2 | 0.273569416 | 3.02E-09 | 9.75E-06 | intron (NM_198664, intron 2 of 12) | 6520 |
| Cyp2a22 | -0.228616822 | 3.58E-42 | 2.12E-15 | promoter-TSS (NM_001101467) | 11 |
| Ror1 | 0.127487027 | 3.02E-09 | 6.94E-05 | intron (NM_013845, intron 1 of 8) | 30892 |
| Plscr1 | 0.179757581 | 3.02E-09 | 0.002179091 | promoter-TSS (NM_011636) | -173 |

| | | | | | |
|---|---|---|---|---|---|
| Stk38 | 0.121723123 | 3.03E-09 | 2.25E-05 | intron (NM_134115, intron 1 of 13) | 7630 |
| Mbd2 | 0.228728016 | 3.04E-09 | 2.07E-05 | exon (NM_007831, exon 22 of 29) | 774451 |
| Mir5625 | -0.156470778 | 3.04E-09 | 3.57E-05 | Intergenic | -10468 |
| Chl1 | -0.115759695 | 3.06E-09 | 4.88E-07 | promoter-TSS (NM_007697) | 7 |
| D130058E03 | -0.39711085 | 1.34E-21 | 3.56E-12 | promoter-TSS (NR_073373) | -384 |
| Cks1brt | -0.253459535 | 3.06E-09 | 5.56E-06 | promoter-TSS (NM_001037922) | -55 |
| Mir6350 | -0.214415732 | 3.06E-09 | 7.07E-06 | Intergenic | -1627916 |
| Cdh26 | 0.263590555 | 3.07E-09 | 6.49E-06 | Intergenic | 481776 |
| Grik3 | -0.265023576 | 3.08E-09 | 5.93E-06 | Intergenic | -171438 |
| Sulf1 | -0.183616906 | 3.08E-09 | 7.08E-06 | exon (NM_001198565, exon 18 of 22) | 124208 |
| Ikbip | 0.132511452 | 3.08E-09 | 1.48E-05 | intron (NM_027078, intron 2 of 2) | 11963 |
| Gm4265 | 0.131025967 | 3.09E-09 | 0.001540544 | promoter-TSS (NR_046077) | -13 |
| Cyb5b | 0.239418512 | 1.35E-21 | 1.98E-08 | Intergenic | -14039 |
| Lce1h | -0.263200608 | 3.11E-09 | 1.41E-05 | intron (NM_026335, intron 1 of 1) | 308 |
| Wdfy3 | 0.138028821 | 3.11E-09 | 0.000388776 | intron (NM_172882, intron 1 of 66) | 13850 |
| Cilp2 | -0.300641183 | 3.12E-09 | 3.88E-06 | intron (NM_001024954, intron 2 of 7) | 23382 |
| Nde1 | 0.149873534 | 3.12E-09 | 6.76E-06 | intron (NM_001114085, intron 5 of 8) | 22134 |
| Galnt7 | -0.382144312 | 3.12E-09 | 6.77E-06 | Intergenic | -46603 |
| Gmds | 0.110654352 | 3.12E-09 | 9.90E-06 | Intergenic | -72915 |
| 2310015B20Rik | -0.199984407 | 3.14E-09 | 3.12E-07 | promoter-TSS (NR_038041) | 87 |
| Rtbdn | 0.103724285 | 3.14E-09 | 1.52E-06 | 5' UTR (NM_144929, exon 2 of 6) | 3498 |
| Pip4k2a | 0.197851599 | 1.37E-21 | 1.74E-08 | intron (NM_008845, intron 3 of 9) | 102200 |
| C1qtnf9 | -0.321841211 | 3.15E-09 | 1.54E-06 | exon (NM_001033272, exon 3 of 13) | -61371 |
| 2310008N11Rik | 0.168794654 | 3.15E-09 | 1.01E-05 | Intergenic | -17692 |
| Zfp648 | 0.383443232 | 3.16E-09 | 7.45E-06 | Intergenic | -83332 |
| Olfr809 | -0.233319404 | 3.16E-09 | 8.36E-06 | TTS (NM_146324) | 996 |
| Lce1l | -0.281827302 | 3.16E-09 | 1.17E-05 | promoter-TSS (NM_028628) | -14 |
| Kcnq2 | 0.157370547 | 3.17E-09 | 6.71E-06 | exon (NM_010611, exon 6 of 17) | 25456 |
| Gm10440 | -0.22148574 | 3.18E-09 | 6.65E-06 | Intergenic | 981050 |
| Gm5086 | -0.406381822 | 3.18E-09 | 7.78E-06 | Intergenic | 41422 |
| 4933438B17Rik | -0.125710351 | 3.18E-09 | 0.009254655 | Intergenic | -695157 |
| Gm8994 | -0.206591618 | 1.40E-21 | 4.87E-10 | intron (NM_001142734, intron 1 of 1) | 696 |
| Hsf4 | 0.106134635 | 3.20E-09 | 1.73E-06 | TTS (NM_015821) | 423 |
| Smim18 | -0.344642398 | 3.20E-09 | 6.74E-06 | promoter-TSS (NM_001206849) | -160 |
| Ifna13 | -0.181567847 | 3.20E-09 | 0.000821648 | promoter-TSS (NM_177347) | -435 |
| Sik1 | 0.116631305 | 3.20E-09 | 0.004092965 | Intergenic | -85964 |
| Mir2136 | -0.202226278 | 3.21E-09 | 2.10E-06 | promoter-TSS (NR_035516) | 10 |
| Syngr4 | 0.166228802 | 3.21E-09 | 2.68E-05 | intron (NM_001291064, intron 3 of 3) | 9553 |
| Abcg3 | -0.205879106 | 3.21E-09 | 3.49E-05 | intron (NM_030239, intron 1 of 15) | 4437 |
| Vmn2r15 | -0.263940224 | 1.41E-21 | 9.87E-11 | intron (NM_001104626, intron 1 of 5) | 582 |
| Olfr1391 | -0.222449299 | 3.22E-09 | 6.71E-06 | exon (NM_146468, exon 1 of 1) | 411 |
| Zfp648 | 0.235337278 | 3.22E-09 | 9.03E-06 | intron (NM_001204908, intron 1 of 1) | 418 |
| Wbscr17 | 0.103089772 | 3.22E-09 | 0.016434267 | intron (NM_177047, intron 5 of 18) | -250497 |
| Dennd2a | 0.236464454 | 3.23E-09 | 3.39E-07 | intron (NM_172477, intron 1 of 18) | 30221 |
| Acot1 | -0.249606367 | 3.23E-09 | 1.34E-06 | intron (NM_012006, intron 1 of 2) | 2195 |
| Bpifb1 | -0.210065228 | 3.23E-09 | 1.59E-06 | intron (NM_001012392, intron 1 of 15) | 143 |

| | | | | | |
|---|---|---|---|---|---|
| Olfr1361 | -0.241425046 | 3.23E-09 | 9.62E-06 | exon (NM_146541, exon 1 of 1) | 270 |
| Ythdf2 | 0.197197717 | 3.24E-09 | 1.85E-06 | Intergenic | 30220 |
| Tsku | 0.24764344 | 3.24E-09 | 7.62E-06 | Intergenic | -30304 |
| 3830408C21Rik | 0.368642839 | 1.41E-21 | 1.40E-08 | Intergenic | 12387 |
| Mir5127 | -0.192919918 | 3.25E-09 | 1.50E-05 | Intergenic | 142148 |
| Olfr907 | -0.243565226 | 3.25E-09 | 5.29E-05 | Intergenic | -2824 |
| Lepre1 | -0.13673755 | 3.25E-09 | 0.000312207 | intron (NM_001286148, intron 5 of 14) | 5139 |
| Dok5 | -0.180742769 | 3.26E-09 | 1.88E-05 | Intergenic | 483728 |
| Rai1 | -0.138505451 | 3.27E-09 | 1.11E-07 | intron (NM_009021, intron 2 of 5) | 680 |
| Ldhb | 0.114057077 | 3.27E-09 | 3.49E-06 | promoter-TSS (NM_008492) | 9 |
| Ttc18 | 0.247349364 | 3.27E-09 | 8.74E-06 | exon (NM_001163638, exon 3 of 29) | 4697 |
| Defb22 | -0.323077293 | 3.28E-09 | 7.38E-06 | exon (NM_001002791, exon 2 of 2) | 4262 |
| Ifnk | -0.316518961 | 3.28E-09 | 9.98E-06 | intron (NM_178061, intron 2 of 3) | -88668 |
| Mir7018 | -0.13140306 | 3.29E-09 | 8.67E-08 | TTS (NR_105985) | 589 |
| Pou2f1 | 0.129654909 | 3.29E-09 | 8.22E-06 | 3' UTR (NM_198933, exon 16 of 16) | 60053 |
| Sumo3 | 0.153252987 | 3.29E-09 | 2.89E-05 | intron (NM_019929, intron 1 of 3) | 3024 |
| Itgb1bp2 | 0.109550485 | 3.29E-09 | 0.019909786 | promoter-TSS (NM_013712) | -120 |
| Soat2 | 0.31028264 | 3.30E-09 | 6.79E-06 | promoter-TSS (NM_146064) | -13 |
| Ino80c | -0.428658748 | 3.31E-09 | 6.95E-06 | intron (NM_172625, intron 1 of 4) | 6962 |
| Cds1 | 0.110104247 | 3.31E-09 | 1.05E-05 | 3' UTR (NM_172882, exon 67 of 67) | 68978 |
| 1700029N11Rik | 0.275758425 | 3.32E-09 | 7.67E-07 | Intergenic | 64526 |
| Pvrl1 | 0.186481762 | 1.48E-21 | 2.17E-08 | Intergenic | -23376 |
| Ankfn1 | -0.251515157 | 3.33E-09 | 1.04E-05 | promoter-TSS (NM_001080933) | -48 |
| Nanp | -0.326331029 | 3.34E-09 | 9.76E-06 | intron (NM_026086, intron 1 of 1) | 1446 |
| Slc39a14 | 0.155390468 | 3.36E-09 | 8.78E-06 | intron (NM_144808, intron 1 of 9) | 16962 |
| Spaca4 | 0.200079082 | 3.37E-09 | 2.41E-08 | intron (NM_017465, intron 2 of 6) | -16217 |
| Rnf138rt1 | -0.164249038 | 3.37E-09 | 6.77E-06 | Intergenic | 37988 |
| Gli3 | 0.121029059 | 3.37E-09 | 9.06E-06 | intron (NM_008130, intron 2 of 14) | 80033 |
| Mgp | 0.272752733 | 1.52E-21 | 2.58E-08 | promoter-TSS (NM_008597) | -62 |
| Grip1 | 0.165560544 | 3.39E-09 | 3.22E-06 | promoter-TSS (NM_001277295) | 59 |
| Efhc2 | -0.110850381 | 3.39E-09 | 1.19E-05 | intron (NM_028916, intron 3 of 14) | 81801 |
| Uty | 0.122393405 | 3.39E-09 | 3.49E-05 | intron (NM_009484, intron 2 of 26) | 4980 |
| Usp9y | -0.177410652 | 3.40E-09 | 7.27E-10 | Intergenic | -287705 |
| Grm4 | 0.131317599 | 3.40E-09 | 2.10E-05 | Intergenic | -10463 |
| Vrk1 | 0.237657142 | 3.41E-09 | 3.55E-08 | intron (NM_001029843, intron 1 of 13) | 20994 |
| Cic | -0.144582418 | 3.42E-09 | 8.54E-05 | intron (NM_001110132, intron 10 of 19) | 7197 |
| Dynlt1a | -0.100792877 | 4.12E-42 | 1.95E-28 | intron (NM_001166629, intron 1 of 4) | 1307 |
| Ext1 | 0.283048733 | 1.54E-21 | 1.97E-08 | intron (NM_010162, intron 1 of 10) | 88516 |
| Rbl1 | 0.144770886 | 3.44E-09 | 1.54E-08 | intron (NM_011249, intron 14 of 21) | 33935 |
| Trak1 | 0.36079158 | 3.44E-09 | 9.44E-06 | 5' UTR (NM_175114, exon 1 of 16) | 129 |
| Olfr1273-ps | -0.188981341 | 3.44E-09 | 4.83E-05 | Intergenic | 3881 |
| Slc39a4 | 0.114094648 | 3.44E-09 | 0.000244369 | promoter-TSS (NM_028064) | -158 |
| Epb4.1l4a | 0.119004101 | 3.44E-09 | 0.008647598 | Intergenic | -39298 |
| Gm6639 | 0.304069191 | 3.45E-09 | 1.31E-06 | Intergenic | 325133 |
| Zfp819 | -0.271640731 | 3.46E-09 | 9.06E-08 | intron (NM_028913, intron 1 of 4) | 2269 |
| Mir190a | 0.170818372 | 1.58E-21 | 4.32E-11 | intron (NM_001081242, intron 47 of 57) | -24287 |

| | | | | | |
|---|---|---|---|---|---|
| Zfp592 | 0.150379107 | 3.46E-09 | 2.00E-05 | Intergenic | -6418 |
| BC055402 | -0.266883557 | 3.46E-09 | 3.67E-05 | Intergenic | 87667 |
| 0610011F06Rik | -0.112047686 | 3.46E-09 | 0.000132733 | exon (NM_026686, exon 4 of 6) | 1017 |
| Mapkapk3 | 0.301634135 | 3.47E-09 | 3.02E-05 | intron (NM_178907, intron 2 of 10) | 9132 |
| Aatk | 0.288835503 | 3.49E-09 | 7.94E-07 | intron (NM_001198785, intron 1 of 13) | 639 |
| H2-Oa | -0.149530606 | 3.50E-09 | 8.17E-06 | TTS (NR_046183) | -8535 |
| Grin2b | -0.30401739 | 3.51E-09 | 6.74E-06 | intron (NM_008171, intron 3 of 14) | 19066 |
| Fpr2 | -0.168172328 | 3.51E-09 | 0.001946004 | Intergenic | 24751 |
| Gimap6 | 0.316956322 | 3.52E-09 | 8.00E-06 | promoter-TSS (NM_153175) | -42 |
| Ugt1a5 | -0.154271446 | 3.53E-09 | 5.50E-05 | promoter-TSS (NM_201643) | -5 |
| Arhgef4 | 0.140652883 | 3.53E-09 | 0.00012787 | Intergenic | -77134 |
| S1pr4 | -0.19157204 | 3.53E-09 | 0.000469623 | intron (NM_010304, intron 5 of 6) | -9079 |
| Ifnk | 0.125915197 | 3.54E-09 | 0.000100487 | intron (NM_178061, intron 1 of 3) | -23938 |
| Elf4 | 0.187220295 | 3.55E-09 | 7.19E-06 | intron (NM_019680, intron 1 of 8) | 27408 |
| Ddx1 | -0.14001165 | 3.55E-09 | 0.000129091 | promoter-TSS (NM_134040) | -381 |
| Ovol1 | -0.247305202 | 3.56E-09 | 5.81E-08 | intron (NM_019935, intron 1 of 3) | 6533 |
| Thsd7a | 0.172224476 | 3.56E-09 | 5.36E-06 | Intergenic | -96000 |
| Bcor | 0.215240264 | 3.56E-09 | 0.000495041 | Intergenic | 250658 |
| Cntnap5c | -0.273116137 | 1.65E-21 | 2.80E-08 | intron (NM_001081653, intron 3 of 23) | 131691 |
| Pantr1 | -0.234211054 | 3.57E-09 | 7.14E-06 | Intergenic | 339021 |
| Olfr1094 | -0.210518632 | 3.57E-09 | 9.78E-06 | Intergenic | -3624 |
| H2afy2 | 0.174255148 | 3.57E-09 | 0.000222808 | Intergenic | -42640 |
| Olfr362 | -0.210558203 | 3.58E-09 | 6.42E-06 | Intergenic | -12033 |
| Skap2 | -0.390591015 | 3.58E-09 | 6.94E-06 | Intergenic | 221949 |
| Bmp2k | 0.187082235 | 3.59E-09 | 1.80E-06 | intron (NM_080708, intron 1 of 15) | 4516 |
| 9930014A18Rik | -0.111625776 | 3.59E-09 | 7.29E-06 | Intergenic | -152123 |
| Zfy2 | -0.133378854 | 1.66E-21 | 5.33E-21 | Intergenic | 70178 |
| St3gal6 | -0.219329072 | 3.60E-09 | 2.83E-05 | intron (NM_018784, intron 7 of 10) | 48592 |
| Kdm4b | 0.192763936 | 3.61E-09 | 7.09E-06 | intron (NM_172132, intron 5 of 22) | 34602 |
| Homer2 | -0.212042749 | 3.61E-09 | 8.43E-06 | Intergenic | -19286 |
| Ammecr1 | 0.14478769 | 3.62E-09 | 0.00104029 | intron (NM_001199357, intron 6 of 6) | 124641 |
| Spock1 | 0.173328608 | 3.63E-09 | 1.42E-06 | intron (NM_001166464, intron 2 of 10) | 28337 |
| Pdcd1 | 0.166336808 | 3.63E-09 | 1.24E-05 | exon (NM_008798, exon 1 of 5) | 138 |
| 6330410L21Rik | -0.139771953 | 3.63E-09 | 0.057617399 | intron (NM_010113, intron 6 of 23) | -28872 |
| Fermt2 | 0.261785343 | 1.72E-21 | 1.59E-08 | intron (NM_146054, intron 4 of 14) | 52193 |
| 2810032G03Rik | 0.194522628 | 3.64E-09 | 1.91E-06 | Intergenic | 498743 |
| Olfr1280 | -0.181712133 | 3.64E-09 | 8.31E-06 | Intergenic | -1567 |
| Glis1 | 0.258992126 | 3.64E-09 | 2.07E-05 | intron (NM_147221, intron 2 of 10) | 88078 |
| S100g | 0.133600237 | 3.64E-09 | 0.000198688 | intron (NR_038088, intron 1 of 5) | -193377 |
| Eml2 | 0.13125627 | 3.65E-09 | 0.000902704 | Intergenic | -3340 |
| Bid | 0.221739257 | 3.66E-09 | 7.17E-06 | intron (NM_001270475, intron 20 of 31) | -61443 |
| Grrp1 | 0.173593345 | 3.66E-09 | 7.36E-06 | Intergenic | -3002 |
| Kcnd1 | -0.384531494 | 1.73E-21 | 2.55E-10 | intron (NM_001290455, intron 21 of 25) | -10616 |
| Pef1 | -0.256056225 | 3.66E-09 | 1.24E-05 | Intergenic | -4785 |
| Olfr1457 | -0.174973596 | 3.66E-09 | 2.04E-05 | Intergenic | -1945 |
| Olfr231 | -0.312707367 | 3.68E-09 | 6.82E-06 | Intergenic | -2732 |

| | | | | | |
|---|---|---|---|---|---|
| Cpm | -0.116253305 | 3.68E-09 | 0.012071713 | intron (NM_027468, intron 2 of 8) | 12419 |
| Mir669j | -0.116704299 | 3.69E-09 | 3.32E-06 | promoter-TSS (NR_035416) | -243 |
| Mir211 | 0.121266861 | 3.69E-09 | 7.76E-06 | exon (NM_018752, exon 5 of 14) | -2770 |
| Th | -0.173833689 | 3.69E-09 | 8.03E-06 | promoter-TSS (NM_009377) | -10 |
| Smc5 | 0.300950452 | 3.70E-09 | 7.44E-06 | intron (NM_174857, intron 11 of 13) | -49118 |
| Zfp750 | 0.195094472 | 3.71E-09 | 1.10E-05 | intron (NM_029878, intron 14 of 38) | -28210 |
| Ptpn6 | 0.250018097 | 1.79E-21 | 2.65E-10 | promoter-TSS (NM_013545) | -20 |
| Itga7 | -0.191584699 | 3.71E-09 | 5.50E-05 | intron (NM_008398, intron 6 of 24) | 9284 |
| Xrcc6bp1 | 0.143641296 | 3.71E-09 | 5.54E-05 | Intergenic | 191771 |
| Vmn2r52 | -0.216851708 | 3.73E-09 | 5.85E-06 | intron (NM_001105191, intron 1 of 5) | 407 |
| Tln2 | 0.124492209 | 3.74E-09 | 1.36E-05 | Intergenic | -64731 |
| Gtf2f2 | -0.196508896 | 3.74E-09 | 0.006785004 | Intergenic | -10457 |
| Flnb | -0.255646355 | 3.75E-09 | 8.03E-06 | intron (NM_001081427, intron 1 of 45) | 3583 |
| Timp4 | 0.232839848 | 1.79E-21 | 1.32E-09 | promoter-TSS (NM_080639) | -230 |
| Slc35f1 | 0.126718436 | 3.77E-09 | 3.51E-06 | intron (NM_178675, intron 1 of 7) | 187772 |
| Cacna1c | 0.227821336 | 3.77E-09 | 6.80E-06 | intron (NM_001159534, intron 3 of 46) | 70456 |
| Vmn2r88 | -0.248315292 | 3.77E-09 | 9.94E-05 | Intergenic | -30846 |
| Mpped2 | -0.219126754 | 3.78E-09 | 7.66E-06 | intron (NM_029837, intron 3 of 6) | 58368 |
| Eno1 | -0.196331549 | 3.78E-09 | 0.000611265 | Intergenic | -6878 |
| Gm6040 | -0.133787276 | 3.78E-09 | 0.007267379 | promoter-TSS (NM_001025353) | -412 |
| Map7 | -0.374957253 | 3.80E-09 | 7.22E-06 | intron (NM_001198635, intron 1 of 17) | 22277 |
| Gpx4 | 0.415254289 | 3.81E-09 | 1.62E-06 | intron (NM_183426, intron 4 of 31) | 21302 |
| Asb11 | 0.169092897 | 3.81E-09 | 6.67E-06 | promoter-TSS (NM_026853) | -16 |
| Olfr1087 | -0.22918374 | 3.81E-09 | 7.46E-06 | Intergenic | -14687 |
| Rprl1 | -0.13608389 | 3.81E-09 | 0.00075686 | Intergenic | -501848 |
| 2810055G20Rik | 0.117357614 | 3.82E-09 | 2.00E-06 | promoter-TSS (NR_015543) | -38 |
| Snx10 | 0.121486359 | 3.82E-09 | 7.22E-06 | promoter-TSS (NM_001127348) | -182 |
| Nrros | 0.307607434 | 5.23E-42 | 3.12E-20 | intron (NM_146069, intron 1 of 3) | 538 |
| 4632428C04Rik | 0.124205743 | 1.81E-21 | 5.49E-06 | intron (NR_033631, intron 1 of 2) | 983 |
| Pcgf5 | 0.113290472 | 3.82E-09 | 9.15E-06 | promoter-TSS (NM_029508) | -3 |
| Pea15b | 0.151011284 | 3.82E-09 | 1.46E-05 | Intergenic | 26124 |
| Olfr629 | -0.244258594 | 3.82E-09 | 1.94E-05 | exon (NM_146821, exon 1 of 1) | 673 |
| Glp2r | -0.132277207 | 3.82E-09 | 2.43E-05 | Intergenic | -3520 |
| Mycn | 0.114064878 | 3.83E-09 | 7.45E-06 | Intergenic | 32104 |
| Snx30 | 0.132314357 | 3.83E-09 | 8.21E-06 | Intergenic | -4220 |
| Npas3 | -0.272438201 | 3.84E-09 | 8.16E-06 | intron (NM_013780, intron 4 of 11) | 415522 |
| Cyp2b19 | -0.185226906 | 3.84E-09 | 4.01E-05 | promoter-TSS (NM_007814) | -37 |
| Podnl1 | 0.173475764 | 3.88E-09 | 7.92E-06 | promoter-TSS (NM_001013384) | -189 |
| Dclk2 | -0.148179001 | 3.88E-09 | 0.000948795 | intron (NM_001195497, intron 1 of 15) | 1393 |
| Sdc1 | 0.123645807 | 3.88E-09 | 0.04911229 | intron (NM_011519, intron 1 of 4) | 2081 |
| Ikbkb | 0.146853047 | 3.89E-09 | 1.15E-05 | intron (NM_010546, intron 5 of 20) | 17703 |
| Nhlh2 | 0.160819239 | 3.89E-09 | 2.04E-05 | promoter-TSS (NM_178777) | -164 |
| Phykpl | 0.205628951 | 3.90E-09 | 2.23E-06 | 3' UTR (NM_153393, exon 29 of 29) | -1876 |
| H60b | -0.114670573 | 3.90E-09 | 0.029525444 | intron (NM_009018, intron 5 of 5) | -6898 |
| Ccnd3 | 0.270667472 | 1.99E-21 | 1.25E-15 | intron (NM_001081636, intron 4 of 6) | 3400 |
| Gapvd1 | 0.239811216 | 3.92E-09 | 9.29E-06 | Intergenic | 126786 |

| | | | | | |
|---|---|---|---|---|---|
| Cd79b | 0.168161922 | 3.92E-09 | 7.14E-05 | intron (NM_008339, intron 1 of 5) | 186 |
| Card11 | 0.247387189 | 3.94E-09 | 7.07E-06 | promoter-TSS (NM_175362) | -179 |
| Socs3 | 0.344023714 | 3.94E-09 | 1.35E-05 | Intergenic | 19570 |
| Mdfi | -0.185930165 | 3.95E-09 | 3.56E-07 | intron (NM_001276391, intron 4 of 5) | 9242 |
| Limk2 | 0.2694723 | 3.97E-09 | 7.57E-06 | intron (NM_010718, intron 2 of 15) | -3612 |
| Agrp | 0.195481274 | 3.97E-09 | 8.97E-06 | promoter-TSS (NM_001271806) | -98 |
| Tmem229b | -0.161712738 | 2.09E-21 | 9.41E-11 | promoter-TSS (NM_178745) | 79 |
| Tlr13 | 0.306116722 | 3.98E-09 | 7.22E-06 | promoter-TSS (NM_205820) | -77 |
| Pla2g2d | 0.189839947 | 3.99E-09 | 8.39E-06 | exon (NM_011109, exon 1 of 4) | 101 |
| Inafm1 | 0.184920123 | 3.99E-09 | 0.003516019 | promoter-TSS (NM_001136270) | -326 |
| Hbb-y | -0.254589408 | 4.00E-09 | 7.92E-06 | promoter-TSS (NM_008221) | -234 |
| Ifna4 | -0.222177418 | 4.00E-09 | 1.44E-05 | promoter-TSS (NM_010504) | -170 |
| Dnah17 | 0.104308697 | 4.00E-09 | 6.07E-05 | Intergenic | -1414 |
| Gse1 | 0.122569441 | 4.01E-09 | 1.91E-06 | Intergenic | -116733 |
| Ppp1r26 | -0.172023228 | 4.01E-09 | 1.36E-05 | promoter-TSS (NM_001005420) | -216 |
| Aldh7a1 | 0.27981445 | 2.12E-21 | 6.46E-13 | promoter-TSS (NM_138600) | 30 |
| Mirlet7j | -0.272073493 | 4.03E-09 | 2.42E-06 | Intergenic | -166496 |
| Tbx10 | 0.138989017 | 4.03E-09 | 1.08E-05 | promoter-TSS (NM_011533) | -135 |
| Olfr97 | -0.214367057 | 4.03E-09 | 3.34E-05 | Intergenic | -5642 |
| Olfr1448 | -0.208469019 | 4.04E-09 | 1.39E-05 | exon (NM_146701, exon 1 of 1) | 171 |
| Ctnnd2 | 0.276093038 | 4.05E-09 | 8.23E-06 | intron (NM_008729, intron 2 of 22) | 211174 |
| Olfr95 | -0.294362188 | 4.05E-09 | 9.85E-06 | Intergenic | -1504 |
| Basp1 | 0.206114663 | 4.07E-09 | 0.000222855 | Intergenic | 110701 |
| Map3k5 | 0.141544315 | 4.07E-09 | 0.00676535 | intron (NM_008580, intron 1 of 29) | 23536 |
| Gm1966 | -0.28417972 | 2.17E-21 | 1.90E-08 | intron (NM_001277182, intron 1 of 3) | 7734 |
| Gm10436 | -0.139866383 | 4.08E-09 | 7.21E-06 | Intergenic | 97762 |
| Retn | -0.249964413 | 4.08E-09 | 7.36E-06 | 3' UTR (NM_022984, exon 4 of 4) | 2040 |
| Cyth4 | 0.263659162 | 4.11E-09 | 1.50E-06 | promoter-TSS (NM_028195) | 79 |
| Olfr583 | -0.258646325 | 4.12E-09 | 7.35E-06 | promoter-TSS (NM_146757) | -590 |
| Tmprss11c | -0.236199877 | 4.13E-09 | 1.14E-06 | promoter-TSS (NM_001030297) | 34 |
| Olfr1116-ps | -0.164005352 | 4.13E-09 | 8.99E-06 | promoter-TSS (NM_001011734) | -158 |
| Sim1 | 0.127127206 | 4.13E-09 | 1.27E-05 | intron (NM_011376, intron 1 of 10) | 7546 |
| Higd2a | -0.132709409 | 4.13E-09 | 1.39E-05 | TTS (NM_025933) | 1410 |
| Angptl2 | 0.19383725 | 2.19E-21 | 3.15E-13 | intron (NM_011923, intron 1 of 4) | 892 |
| D630013N20Rik | 0.150081837 | 4.13E-09 | 3.80E-05 | Intergenic | -32758 |
| Znrf3 | -0.171424886 | 4.14E-09 | 9.84E-06 | intron (NM_001080924, intron 1 of 8) | 27898 |
| Rpl38 | -0.191420314 | 4.14E-09 | 4.31E-05 | intron (NR_037269, intron 2 of 4) | -10877 |
| Olfr1449 | -0.168007992 | 4.15E-09 | 4.49E-05 | Intergenic | -3006 |
| Wrn | -0.17737092 | 4.16E-09 | 0.007791523 | intron (NM_001122822, intron 1 of 33) | 684 |
| Gm6602 | -0.136135169 | 4.17E-09 | 1.94E-06 | Intergenic | -307520 |
| Hey1 | -0.36323075 | 4.17E-09 | 7.28E-06 | Intergenic | 38893 |
| B3gnt2 | 0.164125186 | 4.17E-09 | 1.67E-05 | Intergenic | 29558 |
| Apof | -0.380973333 | 4.17E-09 | 2.13E-05 | 5' UTR (NM_133997, exon 1 of 2) | 607 |
| Myh4 | -0.149964327 | 4.17E-09 | 2.90E-05 | exon (NM_010855, exon 3 of 40) | 2858 |
| Mir29b-1 | 0.140835178 | 2.22E-21 | 2.73E-14 | promoter-TSS (NR_029744) | 98 |
| Olfr1026 | -0.24399472 | 4.18E-09 | 1.80E-06 | Intergenic | -4948 |

| | | | | | |
|---|---|---|---|---|---|
| Prdm1 | 0.235021166 | 4.19E-09 | 1.75E-05 | intron (NM_007548, intron 1 of 6) | 1483 |
| Gm4265 | -0.223740137 | 4.19E-09 | 0.000109593 | Intergenic | 96411 |
| Gja1 | -0.26873874 | 4.20E-09 | 8.16E-06 | Intergenic | 511132 |
| Csf3 | 0.161671949 | 4.20E-09 | 1.01E-05 | exon (NM_011869, exon 20 of 26) | 6413 |
| Mir30d | -0.155177776 | 4.21E-09 | 5.02E-06 | promoter-TSS (NR_029718) | 7 |
| Lhx2 | 0.30706234 | 4.22E-09 | 1.65E-06 | exon (NM_010710, exon 3 of 5) | 3414 |
| Bpnt1 | -0.269960907 | 4.22E-09 | 1.51E-05 | intron (NM_011794, intron 1 of 8) | 1452 |
| Olfr801 | -0.263486405 | 4.23E-09 | 1.57E-08 | Intergenic | -1069 |
| B4galt6 | 0.212923971 | 4.23E-09 | 3.39E-06 | intron (NM_019737, intron 3 of 8) | 27857 |
| Podxl | 0.144057075 | 4.23E-09 | 1.15E-05 | Intergenic | -222692 |
| Tfeb | 0.290954042 | 4.24E-09 | 1.61E-06 | intron (NM_001161723, intron 1 of 8) | -16795 |
| Olfr1381 | -0.182920424 | 4.24E-09 | 1.04E-05 | Intergenic | 5722 |
| Col7a1 | 0.309152555 | 2.26E-21 | 1.36E-10 | intron (NM_007738, intron 1 of 117) | 249 |
| Olfr1454 | -0.195850673 | 4.26E-09 | 4.63E-06 | Intergenic | -3088 |
| Pacs1 | 0.113188457 | 4.26E-09 | 2.97E-05 | promoter-TSS (NM_153129) | -381 |
| Rhag | -0.28008994 | 4.27E-09 | 7.35E-06 | promoter-TSS (NM_011269) | -987 |
| Gm10790 | 0.213938137 | 4.27E-09 | 1.41E-05 | intron (NR_033545, intron 1 of 6) | 6405 |
| D730050B12Rik | 0.34356873 | 4.28E-09 | 1.32E-06 | promoter-TSS (NR_046196) | -218 |
| Gm20611 | -0.100265314 | 4.28E-09 | 0.003198498 | Intergenic | 20590 |
| Olfr663 | -0.169509217 | 4.29E-09 | 8.26E-06 | Intergenic | 30333 |
| Tln2 | 0.109763304 | 4.29E-09 | 8.61E-06 | Intergenic | -47035 |
| C87414 | -0.163458752 | 5.41E-42 | 1.24E-18 | Intergenic | 183957 |
| Slc24a1 | 0.214773958 | 2.33E-21 | 2.29E-08 | 3' UTR (NM_001162917, exon 35 of 35) | 32936 |
| Nox3 | 0.184235422 | 4.29E-09 | 1.85E-05 | intron (NM_198958, intron 8 of 12) | 24635 |
| Tmem33 | -0.19368481 | 4.30E-09 | 0.001104248 | Intergenic | -26006 |
| Treml4 | -0.184270935 | 4.31E-09 | 7.41E-06 | intron (NM_001033922, intron 1 of 5) | 273 |
| Frmd4a | 0.129767924 | 4.31E-09 | 7.79E-06 | intron (NM_172475, intron 1 of 22) | -68420 |
| Fam92b | 0.155813451 | 4.31E-09 | 8.64E-06 | exon (NM_001033980, exon 2 of 8) | 2665 |
| Max | 0.106900685 | 4.31E-09 | 0.001668278 | Intergenic | -10428 |
| Phf11d | -0.262391486 | 4.32E-09 | 1.73E-05 | intron (NM_199015, intron 4 of 10) | 9242 |
| Zmiz1 | -0.146125691 | 4.32E-09 | 0.010083192 | Intergenic | -140284 |
| Vmn2r120 | -0.333825252 | 2.37E-21 | 1.30E-08 | promoter-TSS (NM_001104591) | -52 |
| Champ1 | -0.359965912 | 4.33E-09 | 7.31E-06 | intron (NM_181854, intron 1 of 1) | 6465 |
| Tec | 0.346016299 | 4.33E-09 | 7.76E-06 | intron (NM_013689, intron 1 of 17) | 19385 |
| Insrr | 0.13450932 | 4.34E-09 | 0.003228682 | exon (NM_011832, exon 5 of 22) | 6259 |
| Olfr1364 | -0.243917817 | 4.35E-09 | 1.18E-06 | Intergenic | -7125 |
| Calml3 | -0.266450639 | 4.35E-09 | 7.74E-06 | Intergenic | 44461 |
| Nxt2 | 0.136589804 | 4.35E-09 | 6.28E-05 | 3' UTR (NM_001290532, exon 5 of 5) | 9388 |
| Sel1l2 | 0.110314772 | 4.35E-09 | 0.002364335 | promoter-TSS (NM_001033296) | -130 |
| Nkd1 | 0.312067733 | 4.36E-09 | 9.16E-06 | intron (NM_001163660, intron 1 of 7) | 43357 |
| Gm1943 | -0.247218386 | 4.36E-09 | 2.67E-05 | Intergenic | 91311 |
| Gm5069 | 0.47455121 | 2.53E-21 | 4.03E-10 | intron (NM_001081175, intron 2 of 7) | -73414 |
| Prrg1 | -0.25179739 | 4.36E-09 | 9.96E-05 | Intergenic | -72319 |
| Ly6g6e | 0.194752163 | 4.37E-09 | 1.22E-05 | TTS (NM_001163192) | 3349 |
| Cldn24 | -0.178426742 | 4.37E-09 | 0.000717303 | Intergenic | -48974 |
| Tmem33 | 0.136530402 | 4.38E-09 | 8.00E-05 | Intergenic | -32484 |

| | | | | | |
|---|---|---|---|---|---|
| Apoo | 0.170773785 | 4.39E-09 | 1.14E-05 | Intergenic | -45588 |
| Tbx1 | 0.12471871 | 4.40E-09 | 4.67E-06 | intron (NM_001285493, intron 8 of 8) | 31609 |
| Tmtc2 | 0.166072265 | 4.41E-09 | 5.51E-05 | intron (NM_177368, intron 1 of 11) | 51591 |
| Nr1d1 | 0.123259435 | 4.42E-09 | 1.05E-07 | intron (NM_145434, intron 7 of 7) | 6810 |
| Tmeff2 | -0.220690197 | 4.42E-09 | 7.61E-06 | Intergenic | -530422 |
| Gpatch2l | 0.362607945 | 2.53E-21 | 6.96E-10 | Intergenic | -27401 |
| Slc22a18 | 0.224825937 | 4.42E-09 | 8.51E-06 | promoter-TSS (NM_001042760) | -30 |
| Il1rl1 | -0.175626849 | 4.43E-09 | 1.38E-05 | intron (NM_001294171, intron 1 of 7) | -2744 |
| Mast4 | 0.242785884 | 4.44E-09 | 1.50E-06 | intron (NM_175171, intron 1 of 28) | 108311 |
| Steap4 | 0.141250481 | 4.44E-09 | 1.64E-06 | promoter-TSS (NM_054098) | -101 |
| Ywhaz | -0.282950966 | 4.44E-09 | 9.82E-06 | intron (NM_001253806, intron 1 of 5) | 342 |
| 1700001D01Rik | 0.248190521 | 4.44E-09 | 1.14E-05 | intron (NR_045475, intron 1 of 4) | 2574 |
| Ghrl | -0.195102259 | 4.44E-09 | 0.000342534 | intron (NM_001286406, intron 1 of 2) | 1845 |
| Rpl27 | -0.112914311 | 4.44E-09 | 0.000639137 | intron (NM_011289, intron 2 of 4) | 936 |
| Avpi1 | 0.142907346 | 2.53E-21 | 1.08E-09 | intron (NM_027106, intron 1 of 2) | 1374 |
| Foxk1 | 0.140767774 | 4.46E-09 | 7.01E-06 | intron (NM_177879, intron 5 of 44) | -533264 |
| Chn1os3 | 0.271550878 | 4.46E-09 | 8.06E-06 | Intergenic | -6395 |
| Lef1 | 0.184400269 | 4.49E-09 | 8.48E-06 | intron (NM_001276402, intron 3 of 9) | 4547 |
| Il10 | 0.240477766 | 4.51E-09 | 6.11E-06 | promoter-TSS (NM_010548) | -80 |
| Foxk1 | -0.308331115 | 4.51E-09 | 0.00014553 | Intergenic | -146869 |
| E130006D01Rik | -0.168102928 | 4.52E-09 | 1.41E-06 | Intergenic | -30253 |
| Olfr745 | -0.297051464 | 2.55E-21 | 2.19E-08 | Intergenic | -2726 |
| 5730507C01Rik | -0.143985625 | 4.52E-09 | 0.000860655 | Intergenic | -188564 |
| Zfp583 | -0.145991765 | 4.53E-09 | 1.82E-06 | promoter-TSS (NM_001033249) | -288 |
| Zscan20 | 0.239723605 | 4.54E-09 | 9.57E-06 | Intergenic | 30735 |
| Erbb2ip | 0.207823055 | 4.55E-09 | 8.82E-06 | Intergenic | -57694 |
| Raver1-fdx1l | 0.233953588 | 4.55E-09 | 1.49E-05 | Intergenic | -6132 |
| Cxcl11 | 0.308393874 | 4.56E-09 | 7.63E-06 | promoter-TSS (NM_019494) | 1 |
| Srsf10 | -0.191772005 | 4.56E-09 | 0.000152212 | Intergenic | -1746 |
| 1600014K23Rik | 0.123361271 | 4.58E-09 | 1.06E-05 | exon (NM_007868, exon 73 of 79) | -99419 |
| Gm6760 | -0.197339314 | 4.58E-09 | 1.97E-05 | exon (NM_001177377, exon 1 of 2) | 178 |
| Hip1r | 0.149210701 | 4.59E-09 | 3.76E-06 | intron (NM_145070, intron 1 of 31) | 7036 |
| Nfil3 | 0.415728085 | 4.59E-09 | 8.21E-06 | Intergenic | -4742 |
| 4933438B17Rik | -0.151151147 | 4.59E-09 | 2.29E-05 | Intergenic | -635246 |
| Tmem168 | -0.100842884 | 4.60E-09 | 3.63E-06 | intron (NM_028990, intron 1 of 4) | 301 |
| Sema6a | -0.131602552 | 4.61E-09 | 0.00092156 | intron (NM_018744, intron 1 of 18) | 15353 |
| Gm10640 | -0.199286754 | 4.62E-09 | 7.77E-06 | Intergenic | 26848 |
| Ckmt2 | -0.14116842 | 2.63E-21 | 2.14E-08 | promoter-TSS (NM_198415) | -245 |
| Tmem38b | 0.123523913 | 4.62E-09 | 0.006276191 | Intergenic | 195613 |
| Micall2 | 0.212436915 | 4.63E-09 | 7.77E-05 | Intergenic | 95205 |
| Hcn3 | 0.179464598 | 4.65E-09 | 1.46E-05 | intron (NM_008227, intron 2 of 7) | 7612 |
| Cldn14 | 0.109351789 | 4.65E-09 | 0.000259744 | intron (NM_019500, intron 2 of 2) | 5944 |
| Gm41 | -0.275392147 | 4.65E-09 | 0.000493792 | Intergenic | 26344 |
| Add3 | 0.224792502 | 4.66E-09 | 2.33E-05 | Intergenic | -11775 |
| Olfr821 | -0.25891255 | 4.66E-09 | 0.000109516 | promoter-TSS (NM_146776) | -217 |
| Clec4n | -0.19460053 | 4.68E-09 | 3.05E-05 | intron (NM_001190320, intron 3 of 4) | 14533 |

| | | | | | |
|---|---|---|---|---|---|
| Mir653 | -0.131350103 | 4.68E-09 | 0.002649692 | promoter-TSS (NR_030612) | -225 |
| Bcl11a | -0.2945792 | 2.68E-21 | 4.00E-08 | Intergenic | -32055 |
| Akp3 | -0.140601209 | 4.69E-09 | 1.90E-08 | exon (NM_007432, exon 5 of 11) | 954 |
| Supt20 | 0.295599014 | 4.70E-09 | 8.07E-06 | Intergenic | -99939 |
| Lingo1 | 0.168641582 | 4.72E-09 | 9.02E-06 | intron (NM_181074, intron 3 of 3) | 46209 |
| Aplp2 | 0.2076082 | 4.73E-09 | 1.02E-05 | intron (NM_001102456, intron 1 of 16) | 7840 |
| N4bp2l1 | -0.200543996 | 4.73E-09 | 1.19E-05 | Intergenic | -2714 |
| Gtsf1l | -0.128611491 | 4.73E-09 | 0.000635857 | exon (NM_026630, exon 2 of 2) | 2072 |
| Mir1897 | -0.114322905 | 2.72E-21 | 7.61E-10 | intron (NR_015580, intron 3 of 4) | 3991 |
| Trp53cor1 | 0.175781104 | 4.74E-09 | 7.62E-06 | promoter-TSS (NR_036469) | 30 |
| Ttn | -0.33774402 | 4.74E-09 | 8.61E-06 | 5' UTR (NM_011652, exon 2 of 313) | 2353 |
| Ccer1 | 0.22130129 | 4.74E-09 | 9.38E-06 | intron (NR_033537, intron 4 of 4) | 32863 |
| Pkp4 | -0.366220891 | 4.74E-09 | 2.95E-05 | intron (NM_026361, intron 1 of 21) | 17997 |
| Mir325 | 0.282347665 | 4.74E-09 | 3.91E-05 | intron (NM_145456, intron 2 of 14) | -49361 |
| Minpp1 | -0.163405993 | 4.74E-09 | 5.37E-05 | intron (NM_010799, intron 1 of 4) | 2287 |
| Zfp521 | -0.190280307 | 4.76E-09 | 0.001695835 | intron (NM_145492, intron 2 of 7) | 30291 |
| Ubxn6 | -0.149432395 | 4.76E-09 | 0.001703435 | promoter-TSS (NM_024432) | -282 |
| Gm3376 | -0.11978484 | 4.76E-09 | 0.046325413 | Intergenic | 196080 |
| She | 0.262698486 | 4.77E-09 | 3.06E-06 | Intergenic | -9881 |
| Adh6a | 0.251114856 | 6.72E-42 | 3.16E-10 | promoter-TSS (NM_026945) | -136 |
| Gm13124 | -0.346825665 | 2.72E-21 | 1.28E-08 | promoter-TSS (NM_001085542) | -264 |
| Calhm2 | 0.277045091 | 4.77E-09 | 5.13E-06 | promoter-TSS (NM_133746) | -216 |
| Mir125b-1 | -0.229199837 | 4.77E-09 | 8.21E-06 | Intergenic | 100769 |
| Speer2 | 0.165869394 | 4.77E-09 | 4.92E-05 | Intergenic | 314165 |
| 3830408C21Rik | 0.282689232 | 4.79E-09 | 9.59E-06 | Intergenic | 72529 |
| Plagl2 | 0.138164753 | 4.79E-09 | 4.11E-05 | intron (NM_018807, intron 1 of 2) | 3436 |
| Cep112 | -0.264333444 | 4.79E-09 | 0.006195111 | intron (NM_029606, intron 20 of 27) | -76721 |
| Gbp3 | 0.111715162 | 4.81E-09 | 1.01E-06 | 5' UTR (NM_001289492, exon 2 of 12) | 270 |
| Plekhh3 | 0.247311155 | 4.82E-09 | 1.11E-06 | intron (NM_146030, intron 10 of 12) | 6710 |
| Mag | -0.154560731 | 2.85E-21 | 2.24E-06 | intron (NM_010758, intron 1 of 12) | 337 |
| Olfr220 | -0.169641784 | 4.82E-09 | 7.90E-06 | Intergenic | -6490 |
| Sh3bp2 | 0.235553172 | 4.84E-09 | 1.66E-06 | intron (NM_011893, intron 1 of 12) | -8457 |
| Gm10791 | 0.175527443 | 4.84E-09 | 0.001779482 | Intergenic | -52344 |
| Slc16a9 | -0.226911997 | 4.85E-09 | 6.38E-06 | Intergenic | 42509 |
| Slc17a9 | 0.140186846 | 4.85E-09 | 2.14E-05 | promoter-TSS (NM_183161) | -232 |
| Nlrp4f | -0.236453931 | 4.85E-09 | 2.47E-05 | intron (NM_175290, intron 5 of 9) | 15619 |
| Sh2d4a | 0.266204231 | 2.94E-21 | 1.91E-08 | exon (NM_028182, exon 2 of 9) | 5798 |
| Mrgpra3 | -0.199510051 | 4.85E-09 | 2.89E-05 | promoter-TSS (NM_153067) | -100 |
| Zer1 | -0.144655176 | 4.86E-09 | 2.47E-06 | Intergenic | -4173 |
| Scn8a | 0.162928682 | 4.86E-09 | 0.000232819 | intron (NM_001077499, intron 1 of 26) | -2790 |
| Usp9y | -0.184009982 | 4.86E-09 | 0.001913916 | intron (NM_148943, intron 12 of 46) | 28831 |
| 4930413M19Rik | -0.141653127 | 4.88E-09 | 1.09E-05 | promoter-TSS (NR_045759) | -119 |
| Mfap4 | 0.160501897 | 4.89E-09 | 5.50E-07 | promoter-TSS (NM_029568) | -212 |
| Rhog | 0.235683389 | 4.89E-09 | 5.39E-06 | intron (NM_019566, intron 1 of 1) | 2429 |
| 1700017N19Rik | -0.227598485 | 4.91E-09 | 6.19E-07 | Intergenic | -1539 |
| Tff1 | 0.115081214 | 4.91E-09 | 4.25E-06 | promoter-TSS (NM_009362) | -98 |

| | | | | | |
|---|---|---|---|---|---|
| Btnl2 | -0.356495056 | 2.96E-21 | 2.94E-10 | exon (NM_079835, exon 6 of 9) | 8557 |
| Gm4788 | -0.279102643 | 4.92E-09 | 7.85E-06 | intron (NM_001160304, intron 12 of 13) | 50653 |
| Samd14 | 0.117211856 | 4.93E-09 | 1.24E-07 | 3' UTR (NM_146025, exon 10 of 10) | 15000 |
| Proz | 0.167145678 | 4.93E-09 | 4.46E-06 | intron (NM_025834, intron 5 of 7) | 6494 |
| Thbs4 | -0.203242982 | 4.94E-09 | 7.97E-06 | exon (NM_011582, exon 1 of 22) | 234 |
| Mir29b-1 | 0.144889311 | 2.98E-21 | 7.89E-15 | Intergenic | -25269 |
| Exoc3l4 | -0.153610673 | 4.95E-09 | 4.15E-07 | exon (NM_028807, exon 4 of 13) | 6562 |
| Mir8104 | -0.420273981 | 4.96E-09 | 7.79E-06 | intron (NM_176919, intron 2 of 8) | 110597 |
| Olfr164 | -0.176220135 | 4.96E-09 | 1.91E-05 | Intergenic | 50175 |
| Ankib1 | 0.103102157 | 4.98E-09 | 1.36E-05 | intron (NM_001289527, intron 1 of 19) | 7938 |
| Gipc2 | 0.155118205 | 5.00E-09 | 9.32E-06 | intron (NM_016867, intron 2 of 5) | 32294 |
| Plcb4 | -0.300942179 | 5.01E-09 | 7.74E-06 | intron (NM_013829, intron 1 of 36) | 63693 |
| Plekhf2 | 0.159024201 | 5.01E-09 | 7.95E-06 | intron (NM_175175, intron 1 of 1) | 8454 |
| Hccs | 0.165641578 | 5.03E-09 | 1.09E-05 | Intergenic | -127317 |
| Plb1 | 0.149040188 | 5.03E-09 | 0.000854292 | Intergenic | 44340 |
| Galnt4 | 0.168377994 | 5.06E-09 | 8.03E-06 | intron (NM_027740, intron 10 of 11) | 57261 |
| Awat2 | 0.127761735 | 5.06E-09 | 9.77E-06 | promoter-TSS (NM_001290395) | 38 |
| Olfr1229 | -0.176914012 | 5.06E-09 | 0.000112744 | exon (NM_001011761, exon 1 of 1) | 428 |
| Klk1b16 | 0.139376973 | 5.07E-09 | 1.17E-05 | promoter-TSS (NM_008454) | -365 |
| Gjc3 | -0.239910826 | 5.08E-09 | 1.39E-05 | promoter-TSS (NM_080450) | -13 |
| Prok2 | -0.289174881 | 5.08E-09 | 6.77E-05 | Intergenic | -16542 |
| Ano3 | -0.248877168 | 5.09E-09 | 8.10E-06 | Intergenic | -107314 |
| Rac1 | 0.250545194 | 2.99E-21 | 1.31E-09 | intron (NM_009007, intron 1 of 5) | 9727 |
| Esp8 | -0.183496201 | 5.09E-09 | 8.49E-06 | promoter-TSS (NM_001177584) | -37 |
| Gm15679 | 0.216791117 | 5.09E-09 | 8.64E-06 | Intergenic | -274287 |
| Coa3 | 0.115464801 | 5.09E-09 | 3.88E-05 | exon (NM_175638, exon 17 of 19) | 2442 |
| Gnal | -0.179868505 | 5.10E-09 | 7.87E-06 | Intergenic | -21721 |
| Timp1 | 0.28143115 | 5.10E-09 | 7.89E-06 | exon (NM_001044384, exon 4 of 6) | 2100 |
| Tns3 | 0.188934124 | 5.10E-09 | 1.15E-05 | intron (NM_001083587, intron 9 of 30) | 124526 |
| Ddx26b | 0.167991188 | 5.11E-09 | 4.40E-06 | exon (NM_172779, exon 7 of 17) | 28972 |
| Fam181a | 0.126121947 | 5.11E-09 | 1.09E-05 | Intergenic | -3926 |
| Fam167b | 0.15787841 | 5.12E-09 | 5.95E-06 | exon (NM_182783, exon 1 of 2) | 213 |
| Rprl1 | -0.257708568 | 5.12E-09 | 8.22E-06 | Intergenic | -594627 |
| Olfr735 | -0.350665412 | 3.02E-21 | 1.58E-08 | promoter-TSS (NM_001011754) | 89 |
| 2610018G03Rik | 0.117384198 | 5.13E-09 | 0.000109873 | intron (NM_133729, intron 2 of 11) | 23299 |
| Dmtn | 0.255382228 | 5.14E-09 | 1.21E-06 | promoter-TSS (NM_001252663) | -148 |
| Olfr1457 | -0.207800719 | 5.14E-09 | 5.60E-05 | exon (NM_146575, exon 1 of 1) | 681 |
| Marveld2 | -0.27211258 | 5.15E-09 | 2.64E-06 | promoter-TSS (NM_178410) | -120 |
| Cdkn2c | -0.261389834 | 5.15E-09 | 8.28E-06 | 3' UTR (NM_007671, exon 2 of 2) | 4170 |
| Ccz1 | -0.175389783 | 5.16E-09 | 9.94E-06 | intron (NM_033039, intron 1 of 4) | -11319 |

Table S7: DNA Methylation DMR TKO vs DKO Top500

| gene_name | meanbetafc | minfdr | Stouffer | Annotation | Distance to TSS |
|---|---|---|---|---|---|
| 1700129C05Rik | 0.224618596 | 2.49E-13 | 0.000341163 | Intergenic | 516485 |
| Pced1b | 0.14379217 | 2.54E-13 | 0.000387566 | Intergenic | -79529 |
| Mir3095 | 0.217231728 | 2.93E-13 | 0.000362665 | TTS (NR_037278) | 131 |
| Tbr1 | -0.159093855 | 3.31E-13 | 0.000501782 | intron (NM_009322, intron 1 of 5) | 1122 |
| Txndc8 | 0.133627933 | 3.47E-13 | 0.00065113 | promoter-TSS (NM_026132) | -46 |
| Nr2f2 | 0.184193336 | 4.51E-13 | 0.001473282 | Intergenic | -730163 |
| 4933427E13Rik | 0.288398814 | 4.59E-13 | 0.000464017 | Intergenic | -93364 |
| Sacs | -0.152031587 | 6.12E-13 | 0.000759275 | promoter-TSS (NM_172809) | -97 |
| C530044C16Rik | 0.149988058 | 6.75E-13 | 0.00923521 | Intergenic | 206021 |
| Slc38a4 | 0.152843987 | 8.27E-13 | 0.000468502 | intron (NM_027052, intron 14 of 15) | 56767 |
| Mir503 | 0.143748371 | 1.03E-12 | 0.005991316 | promoter-TSS (NR_029776) | 280 |
| C530044C16Rik | 0.193191791 | 1.49E-12 | 0.001436804 | Intergenic | 71635 |
| Mgst2 | -0.13459557 | 1.89E-12 | 0.000455477 | intron (NM_001004176, intron 2 of 4) | 78242 |
| 2410088K16Rik | -0.12730319 | 1.94E-12 | 0.001574037 | Intergenic | -2348 |
| 4930438E09Rik | 0.226455398 | 2.13E-12 | 0.000640992 | intron (NR_045961, intron 3 of 5) | 26309 |
| Myocd | 0.142421904 | 2.13E-12 | 0.002912428 | intron (NM_146386, intron 4 of 12) | 51297 |
| DQ267102 | 0.233696583 | 2.35E-12 | 0.000522858 | intron (NR_028261, intron 19 of 20) | 3235 |
| 9430019J16Rik | 0.21101941 | 2.60E-42 | 2.43E-05 | Intergenic | 195290 |
| Kirrel3 | 0.272017308 | 3.02E-12 | 0.000557909 | intron (NM_001190912, intron 1 of 15) | 170375 |
| Fam114a2 | 0.139494021 | 4.75E-12 | 0.019396695 | Intergenic | 53338 |
| Kcnj2 | 0.205938065 | 5.00E-12 | 0.000657188 | Intergenic | 600467 |
| Olfr459 | -0.11293978 | 5.28E-12 | 0.033527869 | Intergenic | -2900 |
| Sfrp5 | -0.298219214 | 5.32E-12 | 0.002887168 | exon (NM_018780, exon 1 of 3) | 372 |
| Olfr30 | 0.290345266 | 6.57E-12 | 0.000673376 | promoter-TSS (NM_146878) | 52 |
| Etnppl | -0.159871628 | 7.14E-12 | 0.001064161 | intron (NM_001163587, intron 2 of 13) | 256 |
| Foxp2 | 0.26071497 | 9.47E-12 | 0.000653267 | intron (NM_212435, intron 1 of 16) | 8004 |
| Hoxb8 | -0.122249039 | 9.99E-12 | 0.012924037 | 5' UTR (NM_010461, exon 1 of 2) | 906 |
| Tekt1 | 0.148963107 | 1.34E-41 | 7.82E-07 | promoter-TSS (NM_001282007) | -10 |
| Cngb3 | 0.173015711 | 1.15E-11 | 0.002178464 | Intergenic | -158468 |
| Snord12 | 0.226935011 | 1.17E-11 | 0.001604784 | 3' UTR (NM_008420, exon 2 of 2) | 37595 |
| Sema3a | 0.190717326 | 1.28E-11 | 0.001753977 | Intergenic | -353050 |
| 4833412C05Rik | -0.262126393 | 1.39E-11 | 0.003612794 | Intergenic | -46130 |
| Mctp2 | 0.169300179 | 1.44E-11 | 0.004023186 | Intergenic | 936964 |
| Gm19689 | 0.251166758 | 1.57E-11 | 0.000710971 | Intergenic | -42517 |
| Arhgap44 | 0.118476203 | 1.65E-11 | 0.009739727 | intron (NM_146386, intron 5 of 12) | -46147 |
| Nlgn1 | 0.108528696 | 2.31E-11 | 0.00799772 | intron (NM_001163387, intron 3 of 5) | 607543 |
| Il11ra1 | 0.105272538 | 2.32E-37 | 6.14E-06 | intron (NM_001163401, intron 5 of 12) | 2873 |
| Sgk1 | -0.232290305 | 2.57E-11 | 0.000920629 | 3' UTR (NM_001161849, exon 12 of 12) | 4195 |
| Ces5a | 0.190482117 | 2.99E-11 | 0.000907517 | Intergenic | 98647 |
| Uaca | -0.184547288 | 3.03E-11 | 0.000915893 | intron (NM_028283, intron 1 of 18) | 5137 |
| Nr2f2 | -0.228242662 | 3.37E-11 | 0.001221565 | intron (NM_009697, intron 2 of 2) | 2841 |
| Mir299a | 0.157184673 | 3.99E-11 | 0.004132553 | promoter-TSS (NR_029650) | -222 |
| Rab17 | 0.204451977 | 4.63E-11 | 0.003881313 | promoter-TSS (NM_008998) | -750 |
| Mir3969 | 0.242344001 | 5.14E-11 | 0.002108396 | promoter-TSS (NR_039550) | -385 |
| 1700029J03Rik | 0.101297653 | 5.33E-11 | 0.001920897 | Intergenic | -30956 |

| | | | | | |
|---|---|---|---|---|---|
| Gm10248 | 0.223236807 | 9.86E-37 | 4.90E-08 | intron (NM_001253369, intron 9 of 29) | -426474 |
| Dsc3 | -0.206960083 | 5.37E-11 | 0.009842669 | promoter-TSS (NM_007882) | -51 |
| Kcne4 | -0.163294172 | 6.10E-11 | 0.001323708 | promoter-TSS (NM_021342) | -94 |
| Slc6a5 | -0.182858734 | 7.21E-11 | 0.000101538 | promoter-TSS (NM_148931) | -670 |
| C730036E19Rik | 0.178966815 | 7.25E-11 | 0.001396951 | Intergenic | -30151 |
| Zscan10 | 0.233211667 | 7.50E-11 | 0.00654492 | exon (NM_001033425, exon 6 of 6) | 9319 |
| DQ267100 | 0.178669877 | 7.81E-11 | 0.005301065 | promoter-TSS (NR_046304) | -96 |
| Mir137 | -0.375837779 | 7.97E-11 | 0.002022591 | Intergenic | -2543 |
| Vmn1r43 | 0.267013348 | 8.52E-11 | 0.001520524 | promoter-TSS (NM_053220) | -248 |
| Cfap58 | -0.18477255 | 8.70E-11 | 0.001066943 | Intergenic | 110567 |
| Gm1527 | 0.194168024 | 1.10E-10 | 0.001057338 | Intergenic | -39543 |
| Gm6498 | 0.255895885 | 1.31E-10 | 0.001140812 | TTS (NR_003630) | 30633 |
| Kcnj2 | 0.201147236 | 1.56E-10 | 0.002854219 | Intergenic | 497483 |
| Rora | 0.247593056 | 1.99E-10 | 0.002640245 | intron (NM_013646, intron 2 of 10) | -12323 |
| BC006965 | 0.214135323 | 2.05E-10 | 0.001672763 | Intergenic | 603898 |
| Mir370 | 0.217267624 | 2.13E-10 | 0.005344312 | intron (NR_028261, intron 4 of 20) | 2214 |
| Kcnma1 | 0.165805797 | 2.14E-10 | 0.001668036 | intron (NM_001253369, intron 3 of 29) | 376805 |
| Tnfsf12 | -0.15282659 | 2.63E-10 | 0.012595978 | 5' UTR (NM_001159503, exon 1 of 13) | 297 |
| 4933409K07Rik | 0.134346038 | 3.18E-10 | 0.002033532 | intron (NR_033506, intron 1 of 3) | 43602 |
| Parp11 | -0.112597832 | 3.23E-10 | 0.003178943 | promoter-TSS (NM_181402) | -162 |
| Ric8 | 0.121669756 | 4.30E-10 | 0.006257808 | exon (NM_053194, exon 3 of 10) | 1077 |
| Obfc1 | -0.177502796 | 4.32E-10 | 0.012324338 | promoter-TSS (NM_175360) | -553 |
| Scly | 0.225798556 | 4.38E-10 | 0.003818376 | intron (NM_016717, intron 1 of 11) | 1308 |
| 4930532M18Rik | 0.209034575 | 4.45E-10 | 0.004043813 | promoter-TSS (NR_108050) | 92 |
| Tbx5 | -0.109100274 | 4.58E-10 | 0.015126965 | Intergenic | -2870 |
| Gm12159 | 0.17348663 | 4.75E-10 | 0.006984042 | Intergenic | -163641 |
| Sctr | -0.123149594 | 4.97E-10 | 0.002081526 | promoter-TSS (NM_001012322) | 26 |
| Fam19a1 | -0.143524932 | 5.05E-10 | 0.003628179 | promoter-TSS (NM_182808) | -15 |
| 4930504O13Rik | 0.152654402 | 5.34E-10 | 0.001648253 | intron (NM_207527, intron 1 of 3) | 324 |
| Ddx3y | 0.153803312 | 5.52E-10 | 0.021337496 | intron (NM_148943, intron 41 of 46) | -22091 |
| 7630403G23Rik | 0.2614195 | 6.45E-10 | 0.000964737 | Intergenic | -181951 |
| Phactr1 | 0.263091008 | 6.48E-10 | 0.00197237 | intron (NM_001005740, intron 1 of 12) | 174435 |
| Spink14 | 0.223045988 | 6.89E-10 | 0.004551081 | intron (NM_001039218, intron 1 of 5) | 329 |
| Vmn1r216 | 0.176758871 | 7.05E-10 | 0.001018245 | Intergenic | -2075 |
| Fpr-rs6 | 0.162636289 | 7.51E-10 | 0.002852597 | exon (NM_177316, exon 1 of 1) | 418 |
| Mpped1 | 0.16426885 | 8.48E-10 | 0.003776044 | 5' UTR (NM_172610, exon 2 of 7) | 11962 |
| 9430019J16Rik | 0.331209682 | 2.81E-28 | 3.21E-05 | Intergenic | -20558 |
| 1700094M24Rik | 0.167751184 | 9.02E-10 | 0.002011398 | Intergenic | -49720 |
| Larp1 | 0.195407299 | 9.65E-10 | 0.005567026 | intron (NM_028451, intron 1 of 18) | 22626 |
| Acta2 | -0.142835018 | 9.75E-10 | 0.02639753 | intron (NM_007392, intron 1 of 8) | 229 |
| Zfp423 | 0.173566012 | 9.87E-10 | 0.00953492 | exon (NM_033327, exon 4 of 8) | 178894 |
| D830026I12Rik | 0.158813742 | 1.01E-09 | 0.009184247 | promoter-TSS (NR_102304) | 146 |
| Mir3095 | 0.270201688 | 1.08E-09 | 0.001216155 | exon (NM_010944, exon 14 of 14) | 68040 |
| Gm13375 | -0.338088406 | 1.14E-09 | 0.001891468 | Intergenic | 68511 |
| Klra8 | 0.272513143 | 1.34E-09 | 0.005553952 | intron (NM_053151, intron 6 of 7) | 12424 |
| Rnf7 | -0.151774394 | 1.39E-09 | 0.004080065 | Intergenic | -52542 |

| | | | | | |
|---|---|---|---|---|---|
| Dlgap1 | 0.238207092 | 1.43E-09 | 0.004352573 | promoter-TSS (NM_001128180) | -611 |
| 4930581F22Rik | 0.17698972 | 1.47E-09 | 0.003381746 | intron (NM_001190914, intron 11 of 16) | -100172 |
| Nr2f2 | 0.270580796 | 1.56E-09 | 0.001971874 | Intergenic | 270779 |
| Mir542 | 0.143490488 | 1.59E-09 | 0.003370535 | promoter-TSS (NR_030264) | -12 |
| Cdkn2a | 0.29068074 | 1.62E-09 | 0.004232597 | Intergenic | 9106 |
| Olfr1151 | 0.204671407 | 1.63E-09 | 0.054922531 | exon (NM_146638, exon 1 of 1) | 359 |
| Slc2a5 | -0.13132513 | 1.64E-09 | 0.001312907 | promoter-TSS (NM_019741) | -23 |
| 4930452N14Rik | 0.195283567 | 1.66E-09 | 0.005461747 | promoter-TSS (NR_040629) | -34 |
| Krt1 | -0.195885674 | 1.67E-09 | 0.007348729 | promoter-TSS (NM_008473) | -77 |
| Col11a2 | -0.235462336 | 1.67E-09 | 0.029865085 | promoter-TSS (NM_009926) | -447 |
| Alox12b | -0.194048994 | 1.75E-09 | 0.005383946 | exon (NM_009659, exon 15 of 15) | 12517 |
| Etl4 | -0.194354412 | 1.80E-09 | 0.016750898 | intron (NM_001177631, intron 5 of 19) | 203282 |
| Cdh9 | 0.1933401 | 1.85E-09 | 0.007653388 | Intergenic | -1716037 |
| 4933432K03Rik | 0.140438343 | 1.93E-09 | 0.003279882 | exon (NM_001081327, exon 2 of 2) | 89234 |
| Sall3 | 0.147314296 | 2.03E-09 | 0.00220716 | Intergenic | -467336 |
| Mir744 | 0.151090012 | 2.10E-09 | 0.002150138 | Intergenic | 84574 |
| Spire2 | 0.155617771 | 2.22E-09 | 0.002647579 | intron (NM_172287, intron 1 of 14) | 1961 |
| Epn2 | -0.19300152 | 2.46E-09 | 0.014920471 | Intergenic | -21574 |
| Mir148a | -0.162818201 | 5.77E-26 | 5.24E-06 | Intergenic | 96930 |
| Prdm5 | 0.19714204 | 2.55E-09 | 0.002757997 | intron (NM_027547, intron 1 of 14) | 1322 |
| Mir882 | 0.188351347 | 2.62E-09 | 0.005035397 | Intergenic | -1310 |
| Arhgap28 | 0.205532075 | 2.73E-09 | 0.002906467 | Intergenic | -110261 |
| Ntrk2 | 0.201813296 | 2.75E-09 | 0.005679651 | intron (NM_001282961, intron 5 of 17) | 36411 |
| 2310011J03Rik | -0.15637916 | 2.83E-09 | 0.012833999 | exon (NM_011789, exon 14 of 14) | 5340 |
| Kcnj2 | 0.20512225 | 2.84E-09 | 0.003287024 | Intergenic | 502850 |
| 4932415M13Rik | 0.141822165 | 2.90E-09 | 0.002350784 | Intergenic | 58767 |
| Slc1a3 | 0.185588717 | 3.01E-09 | 0.002729209 | intron (NM_148938, intron 3 of 9) | 28509 |
| Gm21586 | 0.15979853 | 3.10E-09 | 0.008269825 | intron (NR_033506, intron 1 of 3) | 7115 |
| Kcnj2 | 0.175330795 | 1.23E-25 | 5.84E-06 | Intergenic | 582107 |
| Ext1 | 0.110945522 | 3.33E-09 | 0.004340673 | exon (NM_010162, exon 1 of 11) | 1187 |
| Olfr1259 | 0.190807604 | 3.33E-09 | 0.010213097 | Intergenic | -4572 |
| Ptgdr2 | -0.191665654 | 3.35E-09 | 0.005987415 | TTS (NM_023731) | 3580 |
| Medag | -0.288935224 | 3.92E-09 | 0.003428953 | 5' UTR (NM_027519, exon 1 of 5) | 169 |
| 4921531P14Rik | 0.138542263 | 4.12E-09 | 0.022942988 | Intergenic | -37592 |
| Wt1 | 0.139724126 | 4.30E-09 | 0.003091349 | exon (NM_144783, exon 7 of 10) | 40327 |
| Shisa6 | -0.142278669 | 4.54E-09 | 0.097718148 | exon (NM_001034874, exon 1 of 5) | 672 |
| Gm17751 | 0.245133322 | 4.59E-09 | 0.002607505 | Intergenic | -470395 |
| Met | -0.142527411 | 5.19E-09 | 0.002657038 | exon (NM_008591, exon 13 of 22) | 83090 |
| 3110009F21Rik | 0.242491298 | 1.56E-25 | 6.61E-06 | Intergenic | 72083 |
| Rab31 | 0.220505895 | 5.29E-09 | 0.009105974 | intron (NM_133685, intron 1 of 6) | 49118 |
| Lbh | -0.221023725 | 5.36E-09 | 0.005820152 | intron (NM_029999, intron 1 of 2) | 1249 |
| Kcns3 | -0.186170236 | 5.50E-09 | 0.008785343 | promoter-TSS (NM_173417) | 30 |
| C1qtnf2 | 0.164378666 | 6.05E-09 | 0.006003363 | exon (NM_026979, exon 4 of 4) | 16758 |
| Dnajc4 | 0.151557704 | 6.14E-09 | 0.008739768 | intron (NM_026675, intron 4 of 5) | -1284 |
| Ush1g | -0.194114876 | 6.42E-09 | 0.010896114 | exon (NM_176847, exon 2 of 3) | 3339 |
| Tmem79 | 0.303376341 | 6.45E-09 | 0.004470031 | exon (NM_024246, exon 2 of 4) | 1241 |

| | | | | | |
|---|---|---|---|---|---|
| Hoxb1 | 0.138579535 | 8.47E-09 | 0.006282816 | promoter-TSS (NM_008266) | -193 |
| Mir1188 | 0.164794543 | 8.51E-09 | 0.005426502 | promoter-TSS (NR_035419) | 79 |
| Mctp2 | 0.125061548 | 2.77E-25 | 0.000607055 | Intergenic | 922615 |
| Rgmb | -0.169964405 | 9.93E-09 | 0.004596893 | intron (NM_178615, intron 1 of 2) | 1090 |
| Ddi1 | 0.178976194 | 1.05E-08 | 0.003678022 | intron (NM_027924, intron 1 of 6) | -3671 |
| Hoxa2 | -0.293321096 | 1.20E-08 | 0.007566935 | intron (NM_010452, intron 3 of 4) | -10967 |
| Map3k7 | 0.230455391 | 1.22E-08 | 0.003703437 | Intergenic | 197711 |
| Mir5130 | 0.186204489 | 1.24E-08 | 0.003484336 | 3' UTR (NM_177715, exon 1 of 1) | 2574 |
| Prkrir | -0.267491267 | 1.29E-08 | 0.003338757 | Intergenic | -10163 |
| Ntsr2 | -0.122738092 | 1.34E-08 | 0.008189017 | exon (NM_008747, exon 1 of 4) | 265 |
| BC051665 | 0.213554407 | 1.35E-08 | 0.003616445 | intron (NM_199148, intron 1 of 7) | 431 |
| Arhgef15 | -0.302865307 | 1.36E-08 | 0.00652782 | exon (NM_177566, exon 13 of 16) | 9743 |
| Cdkn2a | 0.310332539 | 1.61E-24 | 7.39E-06 | exon (NM_001040654, exon 2 of 3) | 5538 |
| Zfp474 | 0.176697443 | 1.39E-08 | 0.004076864 | promoter-TSS (NM_025749) | 52 |
| Pomc | -0.121380557 | 1.47E-08 | 0.00674499 | promoter-TSS (NM_001278584) | -52 |
| Mir8104 | -0.29078826 | 1.51E-08 | 0.01742033 | intron (NM_176919, intron 4 of 8) | 135893 |
| Mir370 | 0.112613238 | 1.55E-08 | 0.021932465 | promoter-TSS (NR_029918) | -110 |
| Vmn1r44 | 0.276734998 | 1.94E-08 | 0.003550686 | TTS (NM_053227) | 1185 |
| Kifc1 | -0.229230902 | 2.15E-08 | 0.004541229 | TTS (NM_001195298) | 15958 |
| Slc2a3 | -0.165196104 | 2.15E-08 | 0.007656332 | intron (NM_011401, intron 1 of 9) | 518 |
| Cdk1 | 0.151077939 | 2.18E-08 | 0.003615123 | intron (NM_007659, intron 1 of 7) | 2313 |
| 2010111I01Rik | -0.252047915 | 2.19E-08 | 0.003885133 | intron (NR_110520, intron 2 of 5) | 5898 |
| Nr2f2 | 0.162004015 | 2.36E-08 | 0.010271085 | Intergenic | -504268 |
| Dbx1 | -0.31485662 | 2.44E-08 | 0.005995385 | promoter-TSS (NM_001005232) | -18 |
| Fstl5 | -0.209785298 | 2.46E-08 | 0.004178139 | exon (NM_001253719, exon 10 of 16) | 517905 |
| Hs6st2 | 0.233632225 | 2.51E-08 | 0.004069092 | intron (NM_001077202, intron 2 of 4) | 199992 |
| Slco1a5 | 0.177652099 | 2.56E-08 | 0.005317763 | 5' UTR (NM_001267707, exon 1 of 15) | 105 |
| Gm6588 | 0.170259108 | 2.80E-08 | 0.007853125 | intron (NM_001253917, intron 1 of 15) | 60930 |
| C530044C16Rik | 0.240076553 | 2.94E-23 | 1.02E-05 | Intergenic | 181447 |
| Prom1 | 0.167341158 | 3.02E-08 | 0.016619441 | promoter-TSS (NM_001163577) | -169 |
| Gm21586 | 0.112823154 | 3.59E-08 | 0.028629382 | intron (NR_033506, intron 1 of 3) | 9480 |
| Gm3893 | 0.106356229 | 3.77E-08 | 0.020432977 | intron (NR_033506, intron 1 of 3) | 6773 |
| Vmn1r213 | 0.169169466 | 4.54E-08 | 0.004991665 | Intergenic | -3195 |
| Palm2 | 0.202495574 | 4.61E-23 | 6.01E-05 | intron (NM_172868, intron 3 of 5) | 120086 |
| Pcdha7 | -0.167804056 | 4.36E-87 | 1.81E-10 | exon (NM_009957, exon 1 of 4) | 599 |
| B130024G19Rik | -0.105453814 | 2.13E-22 | 0.000330168 | promoter-TSS (NR_045850) | -296 |
| Nr2f2 | 0.297430147 | 6.62E-22 | 8.68E-06 | Intergenic | -749634 |
| Lpar1 | 0.232606511 | 6.98E-21 | 1.79E-05 | intron (NM_010336, intron 1 of 3) | 5261 |
| Mctp2 | 0.31055144 | 8.32E-21 | 2.22E-05 | Intergenic | 618435 |
| Gpatch2l | 0.210655666 | 1.47E-20 | 1.52E-05 | exon (NM_027405, exon 2 of 10) | 2464 |
| Sema3a | 0.211783865 | 1.47E-20 | 4.55E-05 | Intergenic | -456475 |
| Vamp2 | 0.121400948 | 1.81E-20 | 0.073052116 | intron (NM_009497, intron 1 of 4) | 139 |
| C330011F03Rik | -0.267460407 | 8.62E-20 | 2.75E-05 | intron (NR_037250, intron 3 of 4) | -86575 |
| BC006965 | 0.134029598 | 1.02E-19 | 0.001808062 | Intergenic | 298407 |
| 4930441J16Rik | 0.350178747 | 2.69E-19 | 1.97E-05 | intron (NR_040501, intron 1 of 4) | 248 |
| Lpar1 | 0.209436143 | 5.61E-19 | 3.60E-05 | intron (NM_010336, intron 2 of 3) | 6480 |

| | | | | | |
|---|---|---|---|---|---|
| Kcnma1 | 0.201215783 | 8.81E-19 | 4.33E-05 | intron (NM_001253374, intron 1 of 28) | 353776 |
| Cyb5d1 | -0.194718985 | 1.91E-18 | 0.004094833 | TTS (NM_001045525) | 2429 |
| Tgfb2 | 0.160470635 | 1.02E-17 | 0.002756052 | Intergenic | 147132 |
| Pitx1 | -0.15650164 | 1.30E-17 | 7.12E-05 | intron (NM_011097, intron 1 of 2) | 2728 |
| Elovl6 | 0.288581731 | 2.04E-17 | 0.000291618 | intron (NM_130450, intron 2 of 4) | 58527 |
| Rhox4a | 0.100421969 | 2.24E-17 | 0.000566343 | exon (NM_001039688, exon 2 of 4) | 734 |
| Ces1a | 0.278374757 | 2.76E-17 | 7.26E-05 | exon (NM_001013764, exon 1 of 14) | 131 |
| Rnf182 | 0.151830443 | 3.17E-17 | 0.007434033 | Intergenic | -11590 |
| 9430019J16Rik | 0.279892733 | 2.40E-80 | 1.42E-15 | Intergenic | 141196 |
| Rab38 | 0.313238581 | 3.24E-17 | 0.000111163 | intron (NM_028238, intron 2 of 2) | 48429 |
| Mir487b | 0.113008359 | 5.12E-17 | 0.0019343 | promoter-TSS (NR_030271) | 28 |
| Ces1g | 0.129212091 | 7.58E-17 | 0.000901036 | Intergenic | -90227 |
| Mir154 | 0.160987555 | 9.25E-17 | 2.68E-05 | promoter-TSS (NR_030437) | 76 |
| 4933409K07Rik | -0.116478411 | 1.20E-16 | 0.000164484 | non-coding (NR_033123, exon 2 of 5) | 396 |
| Atp5g3 | 0.287052204 | 2.06E-16 | 0.000112284 | Intergenic | -191590 |
| Olfr648 | 0.220396627 | 2.70E-16 | 8.53E-05 | promoter-TSS (NM_146751) | 34 |
| Sim2 | -0.201089191 | 3.25E-16 | 0.000189801 | Intergenic | -18819 |
| Mup20 | 0.202198968 | 3.85E-16 | 0.00063668 | Intergenic | -1152 |
| Nr2f2 | 0.22717336 | 6.76E-16 | 0.000151133 | Intergenic | -86553 |
| Klhdc9 | -0.333580567 | 1.07E-15 | 0.000102753 | promoter-TSS (NM_001033039) | -70 |
| Nr2f2 | 0.210549237 | 1.44E-15 | 0.000903431 | Intergenic | -850202 |
| Specc1 | -0.300475936 | 1.60E-15 | 0.053910347 | intron (NM_001281818, intron 1 of 14) | 58434 |
| Kirrel3 | 0.25923697 | 2.19E-15 | 4.54E-05 | intron (NM_001190912, intron 1 of 15) | 250636 |
| Utrn | 0.210202113 | 3.57E-15 | 0.000392063 | intron (NM_011682, intron 1 of 73) | 34836 |
| Mir582 | 0.365287962 | 5.18E-15 | 0.00012455 | promoter-TSS (NR_030644) | -104 |
| Rimklb | 0.278626806 | 5.53E-15 | 0.000200578 | Intergenic | 35805 |
| 9130227L01Rik | 0.132008828 | 7.69E-15 | 0.000187841 | Intergenic | 87751 |
| Hdgfrp3 | -0.128668526 | 9.88E-15 | 0.002009588 | intron (NM_013886, intron 1 of 5) | 681 |
| Wscd2 | 0.170800943 | 1.05E-14 | 0.000272579 | 3' UTR (NM_177292, exon 8 of 8) | 38873 |
| Mir432 | 0.125172087 | 1.07E-14 | 0.036984236 | promoter-TSS (NR_029550) | -63 |
| Lgi1 | -0.208406464 | 1.10E-14 | 0.000222746 | promoter-TSS (NM_020278) | -185 |
| 9430019J16Rik | 0.169970136 | 1.18E-14 | 0.001428811 | TTS (NR_040635) | 8199 |
| Mir1197 | 0.117735733 | 1.99E-14 | 0.00314263 | promoter-TSS (NR_030421) | 83 |
| Gata3 | 0.123845383 | 2.20E-14 | 0.00026601 | exon (NM_008091, exon 2 of 6) | 1191 |
| Akap7 | -0.175516753 | 2.54E-14 | 0.000294445 | intron (NM_018747, intron 7 of 7) | 98720 |
| Palm2 | 0.263643861 | 2.68E-14 | 0.000319235 | intron (NM_172868, intron 3 of 5) | 112063 |
| Mir148a | 0.178041104 | 3.99E-14 | 0.00044698 | Intergenic | 217808 |
| Gm3376 | 0.17238265 | 5.68E-14 | 0.004712634 | Intergenic | 64297 |
| Ccl27a | 0.118519023 | 7.66E-14 | 0.000298068 | Intergenic | -41270 |
| Gm10248 | 0.164860672 | 1.07E-13 | 0.011740703 | intron (NM_001253373, intron 13 of 27) | -399743 |
| Mir148a | 0.185484265 | 1.15E-13 | 0.001914531 | Intergenic | 165186 |
| AU023762 | 0.291718419 | 1.31E-13 | 0.000257545 | intron (NR_040760, intron 2 of 4) | 75666 |
| 1700094M24Rik | 0.209485337 | 1.34E-13 | 0.000295285 | Intergenic | 17356 |
| Gm3893 | 0.1012766 | 1.13E-44 | 7.14E-07 | Intergenic | -45226 |
| Gm10248 | 0.192966369 | 1.55E-13 | 0.00246442 | intron (NM_001253373, intron 13 of 27) | -392041 |
| Zfp287 | 0.115447951 | 1.80E-13 | 0.01763477 | Intergenic | -21425 |

| | | | | | |
|---|---|---|---|---|---|
| Rora | 0.188225404 | 1.93E-13 | 0.005574425 | intron (NM_013646, intron 2 of 10) | -30206 |
| Alox5ap | -0.180816539 | 1.94E-13 | 0.002439402 | intron (NM_009663, intron 4 of 4) | 22417 |
| Acsl6 | 0.122575939 | 1.99E-13 | 0.000990822 | intron (NM_001033599, intron 1 of 20) | 5069 |
| C530044C16Rik | 0.132118167 | 2.07E-13 | 0.018991563 | Intergenic | 201715 |
| Gm12159 | 0.17068806 | 2.21E-13 | 0.00293059 | Intergenic | -303640 |
| 2900052N01Rik | 0.183916602 | 2.25E-13 | 0.00258785 | Intergenic | -82490 |
| Dzip1l | -0.100392371 | 2.39E-13 | 0.002873754 | promoter-TSS (NM_028258) | -79 |

Table S7: DNA Methylation DMR DKO_EP vs DKO

| gene_name | maxbetafc | minfdr | Stouffer | Annotation | Distance to TSS |
|---|---|---|---|---|---|
| C530044C16Rik | -0.289256706 | 1.07E-33 | 0.534545104 | Intergenic | 149507 |
| Pcdha7 | -0.170591504 | 4.74E-15 | 1 | exon (NM_009957, exon 1 of 4) | 807 |

Table S7: DNA Methylation DMR DKO_EE vs DKO Top500

| gene_name | meanbetafc | minfdr | Stouffer | Annotation | Distance to TSS |
|---|---|---|---|---|---|
| Nup93 | -0.483262263 | 2.64E-59 | 1.78E-13 | intron (NM_172410, intron 4 of 22) | 52405 |
| Dlg2 | 0.215305377 | 5.68E-24 | 0.0000972 | intron (NM_011807, intron 2 of 22) | -350902 |
| Slc22a18 | -0.21770453 | 0.000000026 | 0.00092227 | promoter-TSS (NM_001042760) | -30 |
| Dlx6 | 0.17053343 | 2.64E-08 | 0.000973972 | intron (NR_015388, intron 1 of 1) | -2808 |
| Rnf149 | -0.134195354 | 2.65E-08 | 0.038749397 | Intergenic | -31537 |
| Scamp2 | -0.163891586 | 2.75E-08 | 0.000963609 | intron (NM_022813, intron 1 of 8) | 1562 |
| Glt1d1 | -0.136364304 | 2.76E-08 | 0.000549639 | intron (NM_177005, intron 8 of 11) | 53562 |
| Ank2 | 0.271850333 | 2.78E-08 | 0.000398479 | intron (NM_178655, intron 1 of 24) | 430 |
| Svs2 | 0.184060615 | 2.82E-08 | 0.001248072 | promoter-TSS (NM_017390) | -29 |
| Tnfsf13b | 0.153302047 | 2.84E-08 | 0.001286252 | Intergenic | 57954 |
| Rora | 0.147924276 | 2.87E-08 | 0.000839443 | intron (NM_013646, intron 2 of 10) | -65541 |
| 1600002D24Rik | -0.238461662 | 2.87E-08 | 0.001919329 | promoter-TSS (NR_040484) | -10 |
| Paupar | 0.152301237 | 2.88E-08 | 0.000699029 | promoter-TSS (NR_117095) | -85 |
| Dmbx1 | -0.426232196 | 2.97E-08 | 0.001033528 | intron (NM_001025567, intron 1 of 4) | 3599 |
| Bnc2 | 0.169362302 | 2.99E-08 | 0.000893896 | intron (NM_172870, intron 2 of 5) | 192443 |
| Scn5a | 0.14387319 | 3.03E-08 | 0.003145075 | intron (NM_009134, intron 18 of 27) | -52734 |
| Slc16a11 | -0.182588382 | 3.15E-08 | 0.005219651 | TTS (NM_172371) | 1937 |
| Lamtor3 | 0.115944624 | 3.16E-08 | 0.000703565 | intron (NM_001033155, intron 3 of 7) | -18486 |
| Gm15679 | -0.324830906 | 7.01E-24 | 0.000000208 | Intergenic | -1049021 |
| Cacnb4 | -0.242328981 | 3.19E-08 | 0.000607178 | intron (NM_001285426, intron 2 of 13) | 46547 |
| Ddx6 | -0.248499152 | 3.23E-08 | 0.000594468 | Intergenic | -22189 |
| Myh8 | 0.193105906 | 3.32E-08 | 0.006099605 | intron (NM_177369, intron 1 of 39) | 132 |
| Ddx43 | 0.110589471 | 3.35E-08 | 0.001668758 | promoter-TSS (NM_001191044) | -110 |
| Cmtm7 | -0.18262976 | 3.36E-08 | 0.000987194 | intron (NM_133978, intron 2 of 4) | 19829 |
| Ctf1 | -0.121529641 | 3.37E-08 | 0.022883185 | intron (NM_001301283, intron 2 of 2) | 2768 |
| Paupar | 0.185469622 | 3.38E-08 | 0.00024809 | intron (NR_002867, intron 3 of 3) | 105971 |
| Trim63 | -0.135017726 | 3.39E-08 | 0.00123923 | exon (NM_001039048, exon 1 of 9) | 104 |
| Slc41a2 | -0.216723855 | 3.43E-08 | 0.000739896 | intron (NM_177388, intron 1 of 10) | 2425 |
| Cd244 | 0.196153242 | 3.52E-08 | 0.001071975 | intron (NM_018729, intron 1 of 8) | 7870 |
| Glrx5 | -0.18765416 | 3.54E-08 | 0.000748818 | intron (NM_028419, intron 1 of 1) | 6699 |
| Brinp3 | -0.322952756 | 3.58E-08 | 0.000434097 | Intergenic | 974350 |
| Tcf4 | -0.316033295 | 0.000000036 | 0.000553469 | Intergenic | -148529 |
| Pcdh11x | -0.256867156 | 0.000000036 | 0.001219078 | intron (NR_073451, intron 1 of 7) | 4231 |
| Dedd | -0.102348841 | 0.000000036 | 0.001578001 | intron (NM_011615, intron 1 of 4) | 1837 |
| Cntn4 | 0.129685901 | 3.62E-08 | 0.00070309 | promoter-TSS (NM_001109749) | -8 |
| Crybg3 | -0.160972818 | 3.62E-08 | 0.000703657 | Intergenic | -13781 |
| Nfix | 0.180987579 | 3.62E-08 | 0.000709381 | promoter-TSS (NM_001081981) | -290 |
| Sipa1l2 | -0.120931602 | 3.66E-08 | 0.028338369 | Intergenic | -34043 |
| Lrrc8b | -0.280768334 | 0.000000037 | 0.000306247 | intron (NM_001033550, intron 1 of 4) | 2048 |
| Hsfy2 | 0.283442617 | 0.000000037 | 0.000607416 | Intergenic | 214991 |
| 8430423G03Rik | -0.106709318 | 3.73E-08 | 0.004198036 | Intergenic | 19744 |
| 1700025C18Rik | 0.115768099 | 3.75E-08 | 0.001112689 | intron (NM_174988, intron 11 of 11) | -24106 |
| 4932438H23Rik | 0.138168336 | 3.77E-08 | 0.001453429 | Intergenic | -71460 |
| 4930529K09Rik | -0.210579334 | 3.79E-08 | 0.000711108 | Intergenic | -477326 |
| Sh3bp5 | -0.272955759 | 3.86E-08 | 0.000619445 | intron (NM_011894, intron 3 of 8) | 34908 |
| Ces5a | 0.169032649 | 0.000000039 | 0.000637591 | Intergenic | 98647 |

| | | | | | |
|---|---|---|---|---|---|
| Olfr530 | 0.270318317 | 1.5E-23 | 0.00000138 | Intergenic | -2440 |
| Gsc | 0.181610707 | 0.000000039 | 0.000807476 | Intergenic | -2996 |
| Ppme1 | -0.128040895 | 4.01E-08 | 0.000638879 | intron (NM_028292, intron 1 of 13) | 15138 |
| Prss23 | -0.174337019 | 4.01E-08 | 0.00072587 | intron (NM_029614, intron 1 of 1) | 5586 |
| 3110099E03Rik | -0.138495524 | 4.05E-08 | 0.00072605 | Intergenic | -15597 |
| Mir467a-4 | 0.183848498 | 0.000000041 | 0.000621486 | promoter-TSS (NR_037254) | -537 |
| Sfswap | -0.145636834 | 4.14E-08 | 0.010437372 | intron (NM_172276, intron 1 of 17) | 1294 |
| Asb1 | -0.166902829 | 4.16E-08 | 0.000242333 | Intergenic | 110762 |
| Rora | 0.261228849 | 2.59E-23 | 0.0000169 | intron (NM_013646, intron 2 of 10) | -30206 |
| Sostdc1 | -0.188348416 | 4.16E-08 | 0.000945899 | promoter-TSS (NM_025312) | -62 |
| Fabp5 | -0.232128626 | 4.16E-08 | 0.000947258 | Intergenic | 11011 |
| Krt26 | 0.167921673 | 4.29E-08 | 0.000237302 | promoter-TSS (NM_001033397) | -67 |
| B3gat2 | -0.192087204 | 4.29E-08 | 0.001012251 | intron (NM_172124, intron 1 of 3) | 14811 |
| Mir8105 | -0.118120372 | 4.63E-08 | 0.029166521 | intron (NR_104290, intron 4 of 7) | -23421 |
| Myo5c | -0.1604212 | 4.66E-08 | 0.007205518 | intron (NM_001081322, intron 1 of 40) | 2233 |
| Etv6 | -0.217889977 | 4.77E-08 | 0.000663332 | intron (NM_007961, intron 1 of 7) | 53129 |
| Mir653 | 0.198628661 | 0.000000048 | 0.000679786 | promoter-TSS (NR_030612) | -225 |
| Ifnar2 | 0.315636429 | 2.61E-23 | 0.000000227 | Intergenic | -47127 |
| 4933427D06Rik | -0.181906619 | 4.81E-08 | 0.156493498 | Intergenic | -61148 |
| Pla2g2a | -0.151711905 | 4.83E-08 | 0.001911909 | promoter-TSS (NR_002926) | 21 |
| Mctp2 | 0.100260022 | 4.95E-08 | 0.077451389 | Intergenic | -330875 |
| Creb5 | 0.110955613 | 4.96E-08 | 0.11027696 | Intergenic | -102718 |
| Cntnap5a | -0.276782336 | 5.06E-08 | 0.000674017 | Intergenic | -1720364 |
| Ehf | -0.243033564 | 5.08E-08 | 0.00060093 | intron (NM_007914, intron 1 of 8) | 7906 |
| Olfr1377 | 0.134817157 | 5.08E-08 | 0.021522799 | Intergenic | -2617 |
| Ccnd3 | -0.151871035 | 5.25E-08 | 0.001525779 | intron (NM_001081636, intron 1 of 6) | 26593 |
| Foxo1 | -0.157431869 | 5.27E-08 | 0.002059462 | intron (NM_019739, intron 1 of 2) | 29456 |
| Slc43a1 | -0.269939671 | 5.28E-08 | 0.001752644 | promoter-TSS (NM_001081349) | -592 |
| Pknox1 | -0.123077182 | 3.55E-23 | 0.000842598 | Intergenic | -6858 |
| 4930519D14Rik | 0.142492707 | 5.34E-08 | 0.007877431 | promoter-TSS (NR_045975) | -137 |
| Ralgps2 | -0.181098009 | 5.52E-08 | 0.005646693 | Intergenic | -17343 |
| Wasf3 | -0.117905742 | 5.52E-08 | 0.006500241 | intron (NM_145155, intron 1 of 9) | 30297 |
| Btg1 | -0.168861743 | 5.73E-08 | 0.000704622 | Intergenic | 74770 |
| Stim2 | -0.177457549 | 5.77E-08 | 0.002284255 | Intergenic | -21718 |
| D16Ertd472e | -0.147455552 | 0.000000058 | 0.002503636 | intron (NM_001252438, intron 2 of 4) | 2657 |
| A730018C14Rik | 0.150647457 | 5.85E-08 | 0.000973561 | Intergenic | 17061 |
| Nrbp2 | -0.144914604 | 0.000000059 | 0.00103687 | promoter-TSS (NM_144847) | -110 |
| Gm15679 | -0.391632057 | 4.84E-23 | 0.000000242 | Intergenic | -715240 |
| Fsbp | -0.127947966 | 5.98E-08 | 0.001047834 | exon (NM_001256142, exon 1 of 2) | 229 |
| 6030466F02Rik | 0.124854466 | 6.06E-08 | 0.004243021 | Intergenic | -32037 |
| Neurod2 | -0.132575488 | 0.000000062 | 0.000837388 | Intergenic | 14154 |
| Psmd7 | 0.190489077 | 6.35E-08 | 0.001419935 | Intergenic | -480425 |
| Gm19782 | -0.204505273 | 6.58E-08 | 0.001809451 | intron (NR_045071, intron 3 of 5) | 87592 |
| Foxg1 | 0.158116554 | 6.92E-08 | 0.002738265 | promoter-TSS (NM_008241) | -700 |
| Rimklb | 0.449413367 | 3.32E-56 | 9.25E-11 | Intergenic | 35805 |
| Drd3 | -0.145336278 | 6.94E-08 | 0.005436544 | Intergenic | -8268 |
| Spred2 | -0.206633109 | 6.96E-08 | 0.000774502 | exon (NM_033523, exon 2 of 6) | 73762 |

| | | | | | |
|---|---|---|---|---|---|
| Plec | -0.110086099 | 7.24E-08 | 0.000909808 | intron (NM_201393, intron 18 of 31) | 7822 |
| 9330158H04Rik | -0.158202576 | 7.28E-08 | 0.017603724 | intron (NM_203491, intron 2 of 2) | -1147 |
| Ntrk2 | 0.169532611 | 7.32E-08 | 0.00972883 | intron (NM_001282961, intron 5 of 17) | 36411 |
| Nfatc2 | -0.184004121 | 7.69E-08 | 0.005345476 | intron (NM_001291169, intron 5 of 10) | 60249 |
| Mir7657 | -0.123690893 | 1.07E-22 | 0.000329446 | intron (NM_013867, intron 5 of 11) | 30510 |
| Wee2 | -0.413237599 | 7.87E-08 | 0.000788395 | promoter-TSS (NM_201370) | 13 |
| Npas3 | 0.221059072 | 7.97E-08 | 0.001317998 | intron (NM_013780, intron 3 of 11) | 368115 |
| Slc38a4 | 0.163543219 | 0.00000008 | 0.012671249 | intron (NM_027052, intron 1 of 15) | 6138 |
| Sorbs2 | 0.155861779 | 8.01E-08 | 0.001613197 | promoter-TSS (NM_172752) | 74 |
| Gm14634 | -0.197388056 | 8.02E-08 | 0.000196288 | Intergenic | 61705 |
| Bpifb6 | 0.165385463 | 8.03E-08 | 0.001028015 | promoter-TSS (NM_199303) | -84 |
| Gm16702 | 0.10369771 | 8.06E-08 | 0.000894844 | intron (NM_001161832, intron 5 of 10) | -7506 |
| 1700057G04Rik | -0.154077658 | 0.000000081 | 0.010615638 | Intergenic | -4212 |
| Stard13 | -0.21432856 | 8.18E-08 | 0.002415057 | Intergenic | -57316 |
| E2f6 | -0.274724711 | 1.23E-22 | 5.69E-08 | Intergenic | 40037 |
| Hs6st2 | 0.25961127 | 8.35E-08 | 0.000568601 | intron (NM_001077202, intron 2 of 4) | 22056 |
| Olfr1378 | 0.102523259 | 8.43E-08 | 0.003228485 | promoter-TSS (NM_146910) | -428 |
| Myh2 | 0.127879272 | 8.44E-08 | 0.002246762 | promoter-TSS (NM_001039545) | 29 |
| Vmn1r45 | 0.200221645 | 8.52E-08 | 0.000842174 | promoter-TSS (NM_011684) | -623 |
| Eva1c | -0.162935057 | 8.66E-08 | 0.001124112 | intron (NM_027627, intron 1 of 6) | 18360 |
| Engase | -0.171429802 | 8.66E-08 | 0.001282582 | intron (NM_001285436, intron 4 of 13) | 30347 |
| Lrig1 | -0.23480567 | 1.38E-22 | 0.000000448 | intron (NM_008377, intron 1 of 18) | 29043 |
| Mtx2 | 0.230091201 | 8.72E-08 | 0.003280183 | Intergenic | 174708 |
| Zdhhc1 | -0.15560914 | 8.98E-08 | 0.001672408 | Intergenic | -3144 |
| Hivep2 | -0.116460676 | 0.000000092 | 0.009365617 | intron (NM_010437, intron 1 of 8) | 98816 |
| 9430019J16Rik | 0.32856837 | 9.37E-08 | 0.002328311 | Intergenic | 309131 |
| A230077H06Rik | -0.222327816 | 9.42E-08 | 0.000865365 | intron (NR_040329, intron 1 of 4) | 22034 |
| Atg4c | 0.203777753 | 9.52E-08 | 0.004482484 | Intergenic | 191296 |
| Neurog2 | 0.100053299 | 9.52E-08 | 0.031237573 | exon (NM_027808, exon 6 of 14) | 53126 |
| Mir7219 | -0.174808935 | 9.57E-08 | 0.001678579 | intron (NM_172631, intron 4 of 5) | 18614 |
| Klk4 | 0.208180032 | 1.58E-22 | 0.00000108 | promoter-TSS (NM_019928) | -154 |
| Vezt | -0.18467116 | 9.67E-08 | 0.00166546 | intron (NM_172538, intron 1 of 11) | 15012 |
| Suclg2 | -0.114819273 | 9.79E-08 | 0.000163071 | intron (NM_011507, intron 1 of 10) | 9067 |
| Bglap | 0.159928864 | 9.96E-08 | 0.001313268 | 3' UTR (NM_025928, exon 5 of 5) | -9897 |
| Gtdc1 | -0.116277166 | 0.000000103 | 0.002078965 | exon (NM_172662, exon 4 of 11) | 105215 |
| Slfn5 | 0.142918864 | 0.000000103 | 0.034956281 | intron (NM_183201, intron 1 of 5) | 1121 |
| Nr4a1 | 0.241949939 | 0.000000104 | 0.001277718 | Intergenic | -6341 |
| Sdcbp | -0.149192681 | 0.000000104 | 0.004604244 | intron (NM_016807, intron 1 of 8) | 5563 |
| Cstad | -0.138289123 | 0.000000108 | 0.002140242 | Intergenic | -59859 |
| Abca16 | 0.286556229 | 1.6E-22 | 0.00000285 | promoter-TSS (NM_001278944) | -119 |
| Crnde | 0.12894596 | 0.000000109 | 0.000870768 | Intergenic | 183624 |
| Olfr525 | 0.20069458 | 0.000000112 | 0.001228892 | Intergenic | -2314 |
| Slc38a11 | -0.199573185 | 0.000000112 | 0.001884218 | promoter-TSS (NM_177074) | -25 |
| Cdkl2 | -0.129760104 | 0.000000112 | 0.010179437 | intron (NM_177270, intron 2 of 12) | 4737 |
| 1810011H11Rik | 0.279092519 | 0.000000113 | 0.023774691 | Intergenic | -1447 |
| Tex21 | -0.217188961 | 0.000000116 | 0.001831329 | intron (NM_019784, intron 3 of 9) | 17005 |
| Iglon5 | 0.194779614 | 0.00000012 | 0.000779538 | intron (NM_001164518, intron 3 of 7) | 11564 |

| | | | | | |
|---|---|---|---|---|---|
| Eif4g2 | 0.185350358 | 0.000000121 | 0.000759411 | intron (NM_001040131, intron 3 of 20) | 2409 |
| Lrig1 | -0.294654311 | 0.000000121 | 0.006896806 | intron (NM_008377, intron 1 of 18) | 6994 |
| Casd1 | -0.368427536 | 2.55E-22 | 0.000000199 | intron (NM_145398, intron 1 of 17) | 1178 |
| Magi1 | -0.239329774 | 0.000000126 | 0.001486265 | intron (NM_001286785, intron 1 of 24) | 100410 |
| Olfr112 | -0.281838355 | 0.000000127 | 0.00100131 | promoter-TSS (NM_001013575) | -546 |
| 1600014C23Rik | 0.126030514 | 0.000000127 | 0.007331361 | Intergenic | -138406 |
| Phc2 | -0.168095578 | 0.00000013 | 0.002099501 | promoter-TSS (NM_018774) | 90 |
| Zfp354a | 0.141666059 | 0.000000132 | 0.001485487 | Intergenic | -10408 |
| Entpd8 | -0.153235712 | 0.000000133 | 0.001601643 | promoter-TSS (NM_028093) | -13 |
| BC006965 | 0.147512627 | 3.1E-22 | 0.000110438 | Intergenic | 298407 |
| Il21 | -0.221088158 | 0.000000133 | 0.001684102 | intron (NM_001291041, intron 2 of 3) | 2864 |
| D1Pas1 | 0.208529863 | 0.000000135 | 0.001147465 | Intergenic | -40452 |
| Fgr | -0.173688043 | 0.000000139 | 0.000892305 | Intergenic | -13509 |
| Slc5a4a | 0.214129055 | 0.000000139 | 0.003239606 | promoter-TSS (NM_133184) | 22 |
| Msx2 | -0.129960984 | 0.00000014 | 0.000644452 | Intergenic | -190626 |
| Nfe2l2 | -0.221599665 | 0.000000147 | 0.003175238 | intron (NM_010902, intron 1 of 4) | 2620 |
| Eid3 | -0.178471287 | 0.000000148 | 0.001390734 | exon (NM_025499, exon 1 of 1) | 321 |
| Pitx1 | -0.103324392 | 0.000000149 | 0.000960476 | Intergenic | -43646 |
| Fam83c | -0.146976366 | 0.000000149 | 0.000962741 | promoter-TSS (NM_027788) | 66 |
| 1810053B23Rik | 0.141314927 | 0.000000149 | 0.001955735 | promoter-TSS (NR_040489) | 8 |
| Milr1 | -0.20621442 | 0.00000015 | 0.001970879 | Intergenic | -13728 |
| Ifnk | -0.362342336 | 0.00000015 | 0.002807298 | intron (NM_178061, intron 2 of 3) | -87099 |
| Rapsn | -0.122423727 | 0.000000152 | 0.001257058 | promoter-TSS (NM_009023) | -35 |
| Camk2a | -0.126234479 | 0.000000154 | 0.001411593 | intron (NM_177407, intron 1 of 17) | 288 |
| Tmem233 | 0.111216042 | 0.000000154 | 0.006522689 | promoter-TSS (NM_001101546) | -283 |
| Rprd1b | -0.123649791 | 0.000000155 | 0.001160973 | intron (NM_001291136, intron 1 of 5) | 6711 |
| Zfp366 | -0.293590268 | 3.72E-22 | 7.75E-08 | intron (NM_001004149, intron 1 of 4) | 11564 |
| Sgce | 0.186640268 | 0.000000159 | 0.00099181 | intron (NM_011360, intron 1 of 10) | 22590 |
| Lrrk2 | 0.253071435 | 0.000000161 | 0.001762231 | Intergenic | -1125 |
| Slc35f6 | -0.128084093 | 0.000000165 | 0.002538876 | intron (NM_175675, intron 1 of 5) | 1690 |
| Vmn1r54 | 0.185236815 | 0.000000165 | 0.004323097 | Intergenic | -22261 |
| Enpp6 | -0.110182391 | 0.000000165 | 0.019428844 | Intergenic | -38408 |
| Hs6st2 | 0.282726548 | 0.000000166 | 0.001086969 | intron (NM_001077202, intron 2 of 4) | 179382 |
| Vldlr | 0.122407984 | 0.000000166 | 0.002289625 | Intergenic | -25096 |
| Crip2 | -0.258162584 | 0.000000168 | 0.001271677 | intron (NM_024223, intron 1 of 6) | 2470 |
| LOC101056149 | -0.158667262 | 0.000000168 | 0.00480199 | Intergenic | 17369 |
| Tgfb2 | 0.2599262 | 2.97E-54 | 2.2E-10 | Intergenic | 147132 |
| Gm12295 | 0.273119859 | 3.88E-22 | 0.000000366 | Intergenic | -77943 |
| Reln | 0.350824171 | 0.000000169 | 0.001020045 | Intergenic | -58642 |
| Wwc2 | -0.192282309 | 0.000000169 | 0.001351651 | intron (NM_133791, intron 1 of 22) | 58382 |
| Zp3 | -0.122472548 | 0.000000169 | 0.001713887 | promoter-TSS (NM_011776) | 51 |
| Pds5b | -0.116206491 | 0.000000171 | 0.001211149 | intron (NM_175310, intron 1 of 34) | 2420 |
| C030007H22Rik | -0.141989941 | 0.000000174 | 0.001083579 | promoter-TSS (NR_040482) | 53 |
| Zp2 | 0.153399548 | 0.000000177 | 0.001026011 | Intergenic | -10860 |
| Phospho1 | 0.167904135 | 0.000000184 | 0.009261532 | exon (NM_153104, exon 3 of 3) | 6057 |
| Ano5 | 0.128653993 | 0.000000185 | 0.001769674 | promoter-TSS (NM_177694) | -37 |
| Gm2694 | -0.180339394 | 0.000000186 | 0.001072378 | intron (NR_125722, intron 1 of 3) | 28145 |

| | | | | | |
|---|---|---|---|---|---|
| Ccdc68 | -0.234310351 | 0.000000187 | 0.002213263 | Intergenic | -131083 |
| Arap1 | -0.198891025 | 0.000000188 | 0.001065895 | intron (NM_001040112, intron 1 of 30) | -2847 |
| Mir669m-1 | -0.169408321 | 0.00000019 | 0.004318393 | intron (NR_035474, intron 3 of 4) | 398179 |
| Kcnh3 | -0.202644855 | 0.000000193 | 0.001435488 | exon (NM_010601, exon 2 of 15) | 1455 |
| Ablim1 | -0.196136608 | 6.54E-22 | 0.000146833 | intron (NM_001290813, intron 1 of 20) | -2747 |
| Smim1 | 0.127082635 | 0.000000199 | 0.00331452 | intron (NM_001163721, intron 2 of 3) | 1456 |
| 4930503H13Rik | -0.173457461 | 0.000000202 | 0.000727497 | Intergenic | -138669 |
| Gpc6 | -0.176625114 | 0.000000207 | 0.003133479 | intron (NM_175500, intron 7 of 7) | -720179 |
| Acvr1 | 0.193317956 | 0.000000211 | 0.001153574 | intron (NM_007394, intron 7 of 11) | 96925 |
| Kit | -0.12577114 | 0.000000213 | 0.002443355 | Intergenic | 168816 |
| Slitrk5 | -0.153841988 | 0.000000215 | 0.004731003 | exon (NM_198865, exon 2 of 2) | 5617 |
| Hmx3 | 0.132123058 | 7.41E-22 | 0.000000158 | promoter-TSS (NM_008257) | -222 |
| Cistr-act | 0.189364458 | 0.000000218 | 0.00081103 | Intergenic | -50563 |
| Wbscr17 | -0.12960027 | 0.00000022 | 0.001491302 | Intergenic | -107833 |
| Srrm4 | -0.186877475 | 0.00000022 | 0.001522635 | Intergenic | -238499 |
| Tmem54 | -0.115498436 | 0.00000022 | 0.162958682 | promoter-TSS (NM_133881) | -43 |
| 4930441J16Rik | 0.227508859 | 0.000000223 | 0.001794964 | Intergenic | 177580 |
| Msx1 | 0.207759384 | 0.000000224 | 0.002031786 | Intergenic | -77655 |
| 1600002D24Rik | -0.239234896 | 0.000000226 | 0.001572845 | Intergenic | 90440 |
| Mir1904 | -0.216987694 | 0.000000231 | 0.001905372 | promoter-TSS (NR_035442) | 57 |
| Cbln1 | -0.15817904 | 0.000000233 | 0.001542666 | Intergenic | 12173 |
| Crmp1 | -0.139186894 | 0.000000233 | 0.044589337 | intron (NM_001136058, intron 1 of 13) | 642 |
| Dmbx1 | -0.131256739 | 7.53E-22 | 0.00405094 | intron (NM_001025567, intron 1 of 4) | 15912 |
| Bcl2l11 | -0.174420605 | 0.000000243 | 0.001228843 | intron (NM_207680, intron 2 of 3) | 9873 |
| Il34 | -0.247461429 | 0.000000244 | 0.001483459 | intron (NM_029646, intron 1 of 7) | 206 |
| Pcdh19 | 0.180468497 | 0.000000244 | 0.002377633 | exon (NM_001105246, exon 1 of 5) | 3282 |
| Olfm4 | -0.25839703 | 0.000000245 | 0.003822031 | exon (NM_001030294, exon 1 of 7) | 140 |
| Kcnf1 | -0.301766078 | 0.000000249 | 0.001900659 | Intergenic | 78950 |
| Dab2 | 0.197468504 | 0.000000249 | 0.003924727 | Intergenic | -19520 |
| Mir6404 | 0.193738783 | 0.00000025 | 0.002005747 | intron (NR_037302, intron 2 of 5) | 98215 |
| Abhd15 | 0.192442291 | 0.000000252 | 0.002184258 | 3' UTR (NM_026185, exon 2 of 2) | 3767 |
| Shfm1 | 0.114040067 | 0.000000255 | 0.005494543 | Intergenic | -105890 |
| Ica1 | -0.183968771 | 0.000000257 | 0.000549936 | intron (NM_010492, intron 1 of 13) | 8373 |
| Kcnma1 | 0.196550894 | 1.15E-21 | 0.00000215 | intron (NM_001253374, intron 1 of 28) | 353776 |
| Sln | 0.183945344 | 0.000000258 | 0.001251521 | promoter-TSS (NM_025540) | -307 |
| Ttc39b | -0.158470445 | 0.00000026 | 0.003390702 | intron (NM_027238, intron 3 of 19) | 58314 |
| 4921513I03Rik | 0.126736141 | 0.000000263 | 0.001610435 | intron (NR_038000, intron 1 of 2) | 162 |
| Gpr35 | -0.228983083 | 0.000000273 | 0.001183131 | promoter-TSS (NM_022320) | -8 |
| Wisp3 | -0.253026806 | 0.000000275 | 0.001778026 | Intergenic | -9438 |
| Zdhhc5 | -0.142472288 | 0.000000275 | 0.011220289 | intron (NM_144887, intron 1 of 11) | 1663 |
| Zfp536 | 0.297647664 | 0.000000282 | 0.001577927 | Intergenic | 448282 |
| Bmp5 | 0.165489195 | 0.000000283 | 0.00154692 | promoter-TSS (NM_007555) | 2 |
| Isg20 | -0.200040117 | 1.31E-21 | 0.000931902 | intron (NM_020583, intron 3 of 3) | 5252 |
| Sp8 | -0.268006304 | 0.000000283 | 0.002138428 | promoter-TSS (NM_177082) | 36 |
| Pxdn | 0.148123525 | 0.000000286 | 0.023451096 | promoter-TSS (NM_181395) | -367 |
| AI506816 | -0.140933762 | 0.000000287 | 0.002211156 | Intergenic | 36091 |
| 2310008N11Rik | 0.107016381 | 0.000000291 | 0.001474349 | Intergenic | 244682 |

| | | | | | |
|---|---|---|---|---|---|
| Mir8091 | 0.188595056 | 0.000000291 | 0.006651539 | intron (NM_015748, intron 4 of 36) | 71933 |
| Camk2d | -0.128062681 | 0.000000294 | 0.003651649 | Intergenic | -1166 |
| Slpi | 0.100566505 | 0.000000295 | 0.011589933 | promoter-TSS (NM_011414) | 57 |
| Alcam | -0.125174001 | 0.000000299 | 0.003645418 | intron (NM_009655, intron 1 of 15) | 8591 |
| Mir874 | -0.24352294 | 1.33E-21 | 0.000000228 | promoter-TSS (NR_030544) | -61 |
| Prrc2b | 0.127920855 | 0.000000302 | 0.002852643 | intron (NM_001159634, intron 1 of 31) | 22391 |
| Wwox | -0.104638056 | 0.000000302 | 0.141179051 | intron (NM_019573, intron 5 of 8) | 218546 |
| Cadps | -0.176787096 | 0.000000303 | 0.001693349 | Intergenic | -207736 |
| 1700024P04Rik | -0.12413903 | 0.000000304 | 0.003053755 | promoter-TSS (NM_027064) | -63 |
| Dytn | 0.150497469 | 0.000000308 | 0.002186159 | Intergenic | 71019 |
| Ly86 | -0.150231818 | 0.000000309 | 0.002640291 | Intergenic | 89749 |
| Wscd2 | 0.171868003 | 0.000000323 | 0.000528456 | Intergenic | 42778 |
| Calcr | 0.16126152 | 0.000000323 | 0.0027487 | intron (NM_007588, intron 3 of 15) | 10674 |
| Tspan5 | -0.229992918 | 0.000000324 | 0.003260401 | intron (NR_121195, intron 1 of 7) | 82165 |
| Zfpm2 | 0.260779867 | 0.000000329 | 0.003913694 | intron (NM_011766, intron 1 of 7) | 3447 |
| Sipa1l2 | -0.11779161 | 0.000000332 | 0.000579511 | Intergenic | -29671 |
| Serpina1f | 0.131215965 | 0.000000341 | 0.008249899 | Intergenic | 18426 |
| Krt5 | 0.144186093 | 1.73E-21 | 0.000000122 | intron (NM_027011, intron 5 of 8) | 2953 |
| Edil3 | -0.158301218 | 0.00000035 | 0.006353426 | intron (NM_010103, intron 3 of 9) | 251346 |
| Tgfb3 | -0.181203245 | 0.000000352 | 0.001722148 | intron (NM_009368, intron 1 of 6) | 4665 |
| Vwa8 | -0.138244966 | 0.000000354 | 0.00147769 | Intergenic | -47749 |
| Vmn1r54 | 0.159210492 | 0.000000354 | 0.001669372 | promoter-TSS (NM_053224) | 76 |
| 1700017J07Rik | -0.204158641 | 0.000000359 | 0.001300513 | Intergenic | 226335 |
| Acsbg1 | -0.188108981 | 0.000000365 | 0.003567676 | intron (NM_053178, intron 1 of 13) | 18688 |
| Mir6541 | -0.168756118 | 0.000000373 | 0.00764187 | intron (NM_175546, intron 2 of 11) | 27701 |
| Ephx3 | -0.155389991 | 0.000000382 | 0.057754018 | exon (NM_001033163, exon 1 of 7) | 220 |
| Vax1 | 0.13023174 | 3.58E-52 | 1.85E-08 | TTS (NM_009501) | 4090 |
| Zfp281 | -0.279004324 | 2.05E-21 | 0.00000112 | Intergenic | -19418 |
| C130080G10Rik | -0.168914433 | 0.000000391 | 0.001539811 | intron (NR_028422, intron 1 of 2) | 1468 |
| Mylip | -0.192167533 | 0.000000393 | 0.002202935 | Intergenic | -115150 |
| Sec24b | -0.239683974 | 0.000000404 | 0.001422673 | Intergenic | -12443 |
| Taar2 | -0.106516119 | 0.000000409 | 0.001348725 | TTS (NM_001007266) | 2944 |
| Gm6634 | 0.201937145 | 0.000000413 | 0.001423586 | Intergenic | -239395 |
| St6galnac5 | -0.166198883 | 0.000000413 | 0.002219432 | intron (NM_012028, intron 2 of 4) | 38992 |
| Pcnx | -0.22848592 | 0.000000413 | 0.002349 | intron (NM_018814, intron 6 of 35) | 67710 |
| Cped1 | 0.281886481 | 0.000000418 | 0.002770914 | 5' UTR (NM_001081351, exon 1 of 22) | 839 |
| Slc35f2 | 0.17321037 | 0.000000426 | 0.001936981 | intron (NM_028060, intron 1 of 7) | 21982 |
| Ano6 | -0.181325368 | 0.000000427 | 0.001422507 | promoter-TSS (NM_175344) | -364 |
| Mypn | -0.329041711 | 0.000000434 | 0.002148804 | intron (NM_182992, intron 2 of 19) | 26813 |
| Psmd7 | 0.147026578 | 0.00000044 | 0.009356224 | Intergenic | -360385 |
| 4933427I22Rik | -0.168577092 | 0.000000446 | 0.001830588 | promoter-TSS (NR_045160) | -63 |
| Gpr35 | -0.157049536 | 0.000000447 | 0.001231656 | Intergenic | 10508 |
| Magel2 | 0.246458122 | 2.82E-21 | 0.000000196 | 5' UTR (NM_013779, exon 1 of 1) | 150 |
| Ptpn20 | 0.172673243 | 0.000000448 | 0.007538163 | intron (NM_008978, intron 10 of 11) | 43868 |
| Cd93 | 0.155087647 | 0.000000454 | 0.002035588 | promoter-TSS (NM_010740) | -108 |
| Tbr1 | -0.226123379 | 0.000000454 | 0.004947646 | 3' UTR (NM_009322, exon 6 of 6) | 9191 |
| Tmem132d | 0.269974149 | 0.000000462 | 0.001971801 | Intergenic | -5897 |

| | | | | | |
|---|---|---|---|---|---|
| Vmn1r47 | 0.196973018 | 0.000000463 | 0.001344312 | Intergenic | -6228 |
| Fam46a | -0.173716861 | 0.000000463 | 0.037115637 | Intergenic | -21078 |
| Ccl27a | 0.106232253 | 0.000000469 | 0.002129715 | Intergenic | -54459 |
| D17Ertd648e | -0.157268502 | 2.82E-21 | 0.019655516 | intron (NM_016694, intron 6 of 11) | 269830 |
| Frmd5 | -0.140032019 | 0.000000503 | 0.001827862 | intron (NM_172673, intron 1 of 14) | 88165 |
| Arhgef3 | -0.11920849 | 0.000000505 | 0.083191808 | promoter-TSS (NM_001289688) | -872 |
| Bcan | 0.140049674 | 0.000000512 | 0.004352214 | exon (NM_001109758, exon 8 of 8) | 7301 |
| Foxd3 | 0.119159867 | 0.000000516 | 0.005458469 | 3' UTR (NM_010425, exon 1 of 1) | 1762 |
| Lurap1 | -0.142229033 | 0.000000519 | 0.022096051 | intron (NM_026547, intron 1 of 1) | 3260 |
| Ptchd1 | -0.231801208 | 0.000000539 | 0.002159335 | Intergenic | -132496 |
| En2 | -0.247637764 | 0.000000544 | 0.00061349 | 3' UTR (NM_010134, exon 2 of 2) | 5109 |
| Rbpms | -0.165382945 | 0.00000055 | 0.001542614 | intron (NM_001286168, intron 1 of 7) | 47454 |
| Rora | 0.326931446 | 2.86E-21 | 0.00000607 | intron (NM_013646, intron 2 of 10) | -12323 |
| Dlg2 | 0.23515645 | 0.000000554 | 0.005500709 | Intergenic | -96692 |
| Zfp536 | 0.177312632 | 0.00000057 | 0.001489784 | Intergenic | 303062 |
| Avpr2 | -0.143375267 | 0.00000057 | 0.00247253 | exon (NM_001276299, exon 1 of 3) | 159 |
| Dedd | -0.147871289 | 0.000000573 | 0.006958403 | Intergenic | -2927 |
| Plac1 | 0.166295617 | 0.000000574 | 0.008434412 | promoter-TSS (NM_019538) | -167 |
| Mir344d-2 | 0.162720907 | 0.000000575 | 0.014440748 | promoter-TSS (NR_037216) | 30 |
| Bbox1 | -0.216635986 | 0.000000576 | 0.005794328 | Intergenic | 125546 |
| 4933433F19Rik | -0.147189189 | 0.000000583 | 0.001815521 | Intergenic | -54754 |
| Tcf4 | -0.243662039 | 0.000000589 | 0.002246819 | intron (NM_001083967, intron 5 of 19) | 153473 |
| Catsper3 | -0.126017226 | 0.0000006 | 0.004302587 | exon (NM_001252487, exon 2 of 8) | 1841 |
| Mettl11b | 0.140729405 | 0.000000611 | 0.005911094 | Intergenic | 117672 |
| Hpse2 | 0.233040511 | 0.000000612 | 0.001621589 | promoter-TSS (NM_001081257) | -391 |
| 1700060C16Rik | -0.264336216 | 0.000000631 | 0.002779929 | Intergenic | 115780 |
| Mir1907 | -0.168253507 | 0.000000638 | 0.01240489 | TTS (NR_035444) | 509 |
| Gfra3 | -0.140633374 | 0.000000642 | 0.001611256 | promoter-TSS (NM_010280) | -74 |
| Il17rd | -0.24542963 | 0.000000651 | 0.001879598 | intron (NM_134437, intron 1 of 12) | 35894 |
| C530044C16Rik | 0.256711452 | 0.000000665 | 0.002812354 | Intergenic | 208483 |
| Lin28a | 0.18339191 | 0.000000665 | 0.003112693 | intron (NM_145833, intron 2 of 3) | 5550 |
| Gm12159 | 0.207055321 | 0.000000669 | 0.012900519 | intron (NM_007897, intron 8 of 15) | 215923 |
| 3010033K07Rik | 0.281276645 | 0.000000678 | 0.001994001 | Intergenic | 129486 |
| Sorcs3 | -0.220941742 | 0.000000696 | 0.002401363 | Intergenic | 884582 |
| Cmc1 | 0.11758592 | 0.000000698 | 0.014957449 | Intergenic | -115978 |
| Tmem171 | 0.20338954 | 0.000000699 | 0.004565218 | promoter-TSS (NM_001025606) | -131 |
| Lrig1 | -0.263156679 | 4.17E-21 | 0.000000581 | intron (NM_008377, intron 1 of 18) | 14256 |
| Ski | -0.127762953 | 0.000000702 | 0.001464266 | intron (NM_011385, intron 1 of 6) | 26103 |
| Mir7219 | -0.228948157 | 0.000000704 | 0.004843225 | intron (NM_172631, intron 3 of 5) | 29836 |
| Shc2 | -0.108171402 | 0.00000071 | 0.001805906 | 3' UTR (NM_001033473, exon 4 of 4) | -1866 |
| Mir7680 | 0.130832785 | 0.000000712 | 0.002502086 | non-coding (NR_046162, exon 5 of 5) | 7334 |
| Bcan | 0.15725076 | 0.000000722 | 0.002065822 | intron (NM_007529, intron 6 of 13) | 4949 |
| Arhgef12 | -0.155325277 | 0.000000731 | 0.009033741 | intron (NM_027144, intron 1 of 40) | 3321 |
| D730001G18Rik | -0.188560068 | 0.000000732 | 0.011832294 | Intergenic | -5425 |
| Proser2 | -0.147668825 | 0.000000734 | 0.004951302 | intron (NM_144883, intron 1 of 3) | 11454 |
| 1700110K17Rik | -0.22872865 | 5.71E-21 | 0.000000134 | intron (NM_172768, intron 2 of 19) | 48404 |
| Ets1 | 0.156218591 | 0.000000734 | 0.012322779 | Intergenic | 384295 |

| | | | | | |
|---|---|---|---|---|---|
| Colgalt2 | -0.14775968 | 0.000000739 | 0.001564293 | intron (NM_177756, intron 1 of 11) | 61917 |
| 5430421F17Rik | -0.286207326 | 0.00000074 | 0.002195892 | intron (NR_040352, intron 1 of 2) | 147 |
| Tmem220 | -0.149522869 | 0.000000749 | 0.009231804 | intron (NM_177392, intron 5 of 6) | 6175 |
| Slc35f2 | -0.177965794 | 0.000000755 | 0.006077275 | promoter-TSS (NM_028060) | -567 |
| Gm7325 | -0.118089552 | 0.000000758 | 0.002206276 | promoter-TSS (NM_001177470) | 14 |
| Degs2 | -0.174780977 | 0.000000759 | 0.005386666 | intron (NM_027299, intron 1 of 2) | 2235 |
| Mmd | 0.212457853 | 0.000000764 | 0.001664786 | Intergenic | -105871 |
| Zbtb38 | -0.217320717 | 8.47E-21 | 0.000000112 | intron (NM_175537, intron 2 of 2) | 35230 |
| Smyd3 | -0.1921755 | 0.000000768 | 0.003369927 | intron (NM_027188, intron 8 of 11) | 457895 |
| Thap1 | 0.16611675 | 0.000000769 | 0.002010687 | Intergenic | 84460 |
| 2810032G03Rik | -0.167967088 | 0.000000786 | 0.006019892 | Intergenic | 498743 |
| Mgat5 | -0.14046535 | 0.000000792 | 0.046708397 | exon (NM_145128, exon 3 of 18) | 101875 |
| Asph | -0.161280138 | 0.000000805 | 0.002294416 | intron (NM_001290367, intron 1 of 22) | 3367 |
| 9430076C15Rik | 0.156733048 | 0.000000811 | 0.003844297 | intron (NR_015553, intron 1 of 4) | 1367 |
| Nedd4l | -0.20985668 | 4.03E-52 | 8.98E-13 | intron (NM_031881, intron 1 of 29) | 1157 |
| Fam163a | -0.236863602 | 0.000000816 | 0.002673514 | Intergenic | -17628 |
| Ccdc85c | -0.15331571 | 0.000000824 | 0.014923345 | intron (NM_001159910, intron 1 of 5) | 4452 |
| Mink1 | -0.102207788 | 0.000000828 | 0.054646186 | intron (NM_176893, intron 1 of 31) | 14797 |
| Zfp536 | 0.173930122 | 0.00000084 | 0.004091062 | exon (NM_172385, exon 4 of 6) | 201679 |
| Susd2 | -0.137518373 | 0.000000844 | 0.001684432 | Intergenic | 11318 |
| Gm6498 | 0.209355273 | 0.000000851 | 0.001934792 | TTS (NR_003630) | 30633 |
| 9430019J16Rik | 0.143979916 | 1.03E-20 | 0.038902532 | TTS (NR_040635) | 8199 |
| Olfr120 | -0.300323518 | 0.000000874 | 0.000823445 | promoter-TSS (NM_146631) | -53 |
| Svs3b | 0.170507609 | 0.000000875 | 0.006199387 | promoter-TSS (NM_173377) | -849 |
| Pantr1 | 0.131405295 | 0.00000089 | 0.001777778 | Intergenic | 203190 |
| 4933433F19Rik | -0.285304929 | 0.00000089 | 0.002219363 | Intergenic | -125117 |
| Kcnc1 | 0.153238628 | 0.000000895 | 0.002289942 | promoter-TSS (NM_008421) | -364 |
| Irx3 | 0.175163603 | 0.000000897 | 0.001911372 | Intergenic | -63741 |
| Trpc7 | 0.114895367 | 0.000000898 | 0.012784782 | exon (NM_012035, exon 2 of 12) | 7669 |
| App | -0.22112854 | 0.000000903 | 0.00244543 | intron (NM_001198826, intron 1 of 16) | 45860 |
| Rmdn1 | -0.134274978 | 0.000000921 | 0.04152763 | intron (NM_025476, intron 1 of 9) | 1596 |
| Dlk1 | 0.140322501 | 0.00000094 | 0.008928312 | exon (NM_010052, exon 1 of 5) | 211 |
| Ano6 | -0.233129729 | 0.000000941 | 0.011917785 | intron (NM_175344, intron 1 of 19) | 39644 |
| Vmn2r98 | -0.250271726 | 0.000000944 | 0.001345458 | intron (NM_001104550, intron 1 of 5) | 1604 |
| Scn5a | 0.133780782 | 0.000000953 | 0.002011525 | promoter-TSS (NM_001253860) | -466 |
| Prkd3 | -0.187172775 | 0.000000958 | 0.002154715 | intron (NM_029239, intron 1 of 18) | 2224 |
| Pced1b | 0.156569645 | 0.000000965 | 0.010254091 | 3' UTR (NM_178114, exon 1 of 1) | -2472 |
| Comt | -0.287077503 | 1.92E-20 | 0.000000182 | intron (NM_007744, intron 1 of 4) | -2877 |
| Scn10a | 0.167703594 | 0.00000097 | 0.001827722 | exon (NM_001205321, exon 2 of 28) | 24819 |
| Fam122a | -0.129980968 | 0.00000097 | 0.004133759 | intron (NM_008846, intron 1 of 14) | -14060 |
| Vstm4 | 0.122258453 | 0.00000097 | 0.07297956 | promoter-TSS (NM_178791) | -220 |
| Pvrl1 | -0.214458076 | 0.000000998 | 0.0028064 | intron (NM_021424, intron 1 of 5) | 2150 |
| Uck2 | -0.136472695 | 0.00000101 | 0.002669174 | intron (NM_030724, intron 2 of 6) | 48239 |
| 4933407E24Rik | 0.15320774 | 0.00000102 | 0.001938863 | Intergenic | 41369 |
| Arsg | -0.168071895 | 0.00000102 | 0.003056588 | intron (NM_001166177, intron 1 of 4) | 2920 |
| Rxfp3 | 0.206771914 | 0.00000102 | 0.007908363 | 3' UTR (NM_178717, exon 1 of 1) | 2406 |
| Wdr11 | -0.175454528 | 0.00000103 | 0.004113298 | Intergenic | 152987 |

| | | | | | |
|---|---|---|---|---|---|
| Mir3964 | -0.229923099 | 2.08E-20 | 0.00000111 | Intergenic | 418235 |
| 2810032G03Rik | -0.114555482 | 0.00000103 | 0.010254632 | intron (NR_015579, intron 1 of 1) | 35030 |
| F13a1 | 0.170141541 | 0.00000104 | 0.001916778 | intron (NM_028784, intron 11 of 14) | 143596 |
| Treml4 | 0.140985044 | 0.00000105 | 0.002236105 | intron (NM_172623, intron 4 of 5) | 8192 |
| Plxna1 | -0.109933383 | 0.00000106 | 0.002033076 | Intergenic | 53566 |
| Smad4 | -0.146664315 | 0.00000107 | 0.002091102 | intron (NM_008540, intron 1 of 11) | 3033 |
| Sdc4 | -0.13819676 | 0.00000111 | 0.047240524 | intron (NM_011521, intron 1 of 4) | 7017 |
| Cftr | -0.436688265 | 2.39E-20 | 0.000000788 | intron (NM_021050, intron 1 of 26) | 10172 |
| Snx10 | -0.125908008 | 0.00000112 | 0.0031816 | intron (NM_001127349, intron 1 of 6) | 727 |
| Tnks | -0.1225705 | 0.00000112 | 0.004230048 | Intergenic | -154809 |
| 9430019J16Rik | 0.266113375 | 0.00000112 | 0.012992012 | Intergenic | 235064 |
| 1700108F19Rik | -0.204365299 | 0.00000116 | 0.00584308 | Intergenic | 18591 |
| Prok2 | -0.130134063 | 0.00000116 | 0.142091488 | Intergenic | -16542 |
| Vipr2 | -0.133647245 | 0.00000117 | 0.002038067 | Intergenic | 74532 |
| Tecrl | -0.19248637 | 0.00000117 | 0.003251284 | intron (NM_153801, intron 5 of 11) | 55604 |
| Thsd4 | 0.123189563 | 0.00000118 | 0.001935507 | intron (NR_040464, intron 2 of 3) | -16313 |
| Prdm12 | 0.120568528 | 3.24E-20 | 0.00000147 | intron (NM_001123362, intron 1 of 4) | 1156 |
| Kcnmb1 | -0.117716227 | 0.0000012 | 0.001975783 | 5' UTR (NM_031169, exon 1 of 4) | 217 |
| 1700023F02Rik | -0.258119157 | 0.0000012 | 0.002154096 | Intergenic | -130045 |
| Rara | -0.141067647 | 0.00000121 | 0.002902551 | Intergenic | -1117 |
| Pus7l | 0.18170806 | 0.00000122 | 0.002830763 | Intergenic | 65287 |
| Adamts9 | -0.125819133 | 0.00000123 | 0.002290853 | promoter-TSS (NM_175314) | 2 |
| Bmp2k | -0.144590566 | 0.00000123 | 0.015685304 | Intergenic | -75879 |
| Gjb6 | 0.128058111 | 0.00000123 | 0.019206594 | 5' UTR (NM_001271663, exon 1 of 3) | 371 |
| Ces1g | 0.138493919 | 3.82E-20 | 0.0000264 | Intergenic | -90227 |
| Mir378b | 0.121704734 | 0.00000127 | 0.003215615 | intron (NM_054043, intron 6 of 13) | 129641 |
| Pcdh17 | -0.222686253 | 0.0000013 | 0.002933108 | Intergenic | -18458 |
| Nuak1 | -0.128355444 | 0.0000013 | 0.011393465 | intron (NM_001004363, intron 2 of 6) | 35044 |
| Ccdc60 | 0.190596569 | 0.00000131 | 0.003816794 | intron (NM_177759, intron 2 of 13) | 79792 |
| Klk6 | 0.148201181 | 0.00000131 | 0.00549026 | intron (NM_001164696, intron 4 of 6) | 1758 |
| Psma6 | -0.196243529 | 0.00000132 | 0.002139886 | intron (NR_035430, intron 1 of 3) | -10619 |
| Dppa3 | 0.150877494 | 0.00000132 | 0.003861503 | exon (NM_139218, exon 2 of 4) | 2285 |
| Gpatch2l | -0.139752968 | 0.00000133 | 0.002343552 | exon (NM_027405, exon 2 of 10) | 2464 |
| Tspan5 | -0.380385382 | 0.00000133 | 0.003524739 | intron (NR_121195, intron 1 of 7) | 35997 |
| Gm1821 | -0.122044812 | 3.87E-20 | 0.001895626 | Intergenic | -139529 |
| Syt2 | -0.155224734 | 0.00000135 | 0.003051844 | intron (NM_009307, intron 1 of 9) | 5446 |
| Dmr | -0.154573393 | 0.00000135 | 0.009321391 | intron (NM_025833, intron 1 of 13) | -15990 |
| Arg1 | -0.112567462 | 0.00000135 | 0.191954881 | Intergenic | -20699 |
| Zfp217 | -0.137680438 | 0.0000014 | 0.002425128 | Intergenic | -93572 |
| Fhdc1 | -0.246647096 | 0.00000142 | 0.008735693 | intron (NM_001033301, intron 4 of 11) | 15413 |
| Mir6937 | -0.196201593 | 0.00000143 | 0.005768017 | TTS (NR_105902) | 123 |
| Hoxb1 | 0.190405352 | 0.00000143 | 0.03787319 | Intergenic | 41858 |
| Pcdh11x | 0.184392985 | 4.28E-20 | 0.00000052 | promoter-TSS (NM_001271809) | 46 |
| 4930465M20Rik | 0.227061557 | 0.00000144 | 0.002069541 | Intergenic | 210432 |
| Efhc1 | -0.181306088 | 0.00000145 | 0.008516634 | exon (NM_028829, exon 3 of 3) | -16234 |
| Pitx1 | -0.221589848 | 0.00000146 | 0.003090378 | promoter-TSS (NM_011097) | -365 |
| Cryba2 | -0.120138866 | 0.00000147 | 0.244075447 | promoter-TSS (NM_021541) | 83 |

| | | | | | |
|---|---|---|---|---|---|
| Hoxc12 | 0.106448693 | 0.00000148 | 0.048759836 | exon (NM_010463, exon 1 of 2) | 197 |
| Nckap1l | 0.203388543 | 0.0000015 | 0.009403092 | promoter-TSS (NM_153505) | 24 |
| Usp26 | 0.127690965 | 0.00000153 | 0.00770824 | exon (NM_031388, exon 9 of 9) | 45238 |
| 4931408C20Rik | -0.240227743 | 0.00000155 | 0.001924476 | exon (NM_001033764, exon 2 of 2) | 4671 |
| Mir351 | 0.215559526 | 1.03E-51 | 9.41E-12 | promoter-TSS (NR_029776) | -116 |
| Moap1 | -0.250602174 | 4.69E-20 | 0.00000141 | Intergenic | 13528 |
| Olfr118 | -0.244322716 | 0.00000156 | 0.002509867 | Intergenic | -3547 |
| Gm13032 | 0.253962689 | 0.00000158 | 0.005338392 | exon (NM_011059, exon 16 of 16) | 3724 |
| Rbpms | -0.16012115 | 0.00000159 | 0.003443721 | intron (NM_001286168, intron 1 of 7) | 44124 |
| Asb9 | -0.211800676 | 0.00000159 | 0.005675401 | Intergenic | 162658 |
| Tbc1d24 | -0.212158741 | 0.0000016 | 0.002564597 | intron (NM_001163848, intron 1 of 9) | 2672 |
| Olfr121 | -0.156036703 | 0.0000016 | 0.003780003 | intron (NM_146629, intron 1 of 1) | 817 |
| Msantd4 | -0.240339711 | 0.00000161 | 0.018027155 | exon (NM_019691, exon 14 of 17) | 72532 |
| Zfp292 | -0.163568223 | 0.00000162 | 0.002228199 | intron (NM_013889, intron 1 of 7) | 13441 |
| Nav1 | -0.142119841 | 0.00000163 | 0.035733351 | exon (NM_173437, exon 1 of 29) | 354 |
| 2700069I18Rik | -0.178026029 | 0.00000165 | 0.004816281 | intron (NM_030708, intron 3 of 10) | -53344 |
| Pabpc6 | 0.144239611 | 0.00000166 | 0.007275204 | Intergenic | 248246 |
| Tmsb4x | -0.162414788 | 0.00000167 | 0.205698598 | Intergenic | -18307 |
| Gm6644 | -0.238456707 | 0.0000017 | 0.002496737 | intron (NR_037965, intron 1 of 9) | 1113 |
| Btd | 0.215409333 | 0.00000171 | 0.001876299 | TTS (NM_025295) | 27778 |
| Lcn4 | 0.147526545 | 0.00000173 | 0.005027494 | Intergenic | -51773 |
| 9130227L01Rik | 0.111648804 | 0.00000174 | 0.002675763 | Intergenic | 219852 |
| 2210010C04Rik | -0.205133238 | 0.00000175 | 0.003102188 | Intergenic | -22390 |
| Pea15b | -0.10057276 | 0.00000175 | 0.088812984 | Intergenic | 46442 |
| Cacnb2 | -0.239048028 | 5.46E-20 | 0.00000155 | promoter-TSS (NM_001252533) | 10 |
| Psmc1 | -0.18031669 | 0.00000178 | 0.00412917 | Intergenic | -27804 |
| Bcar3 | -0.162096758 | 0.0000018 | 0.003339361 | intron (NM_013867, intron 3 of 11) | 39169 |
| Pdxk | -0.10864693 | 0.00000183 | 0.020972941 | promoter-TSS (NM_172134) | -309 |
| Magi2 | -0.108681472 | 0.00000184 | 0.006481087 | intron (NM_001170746, intron 9 of 22) | 486516 |
| Sp8 | -0.158125911 | 0.00000184 | 0.008369153 | Intergenic | -3273 |
| Nr1h4 | -0.206045157 | 0.00000185 | 0.003230412 | promoter-TSS (NM_001163504) | 26 |
| Ap1ar | -0.3112527 | 5.96E-20 | 0.000000893 | Intergenic | -24093 |
| Msantd3 | -0.2129827 | 0.00000192 | 0.00092928 | intron (NM_028137, intron 1 of 2) | 1775 |
| D830026I12Rik | 0.128831285 | 0.00000195 | 0.049120659 | promoter-TSS (NR_102304) | 146 |
| 4933433F19Rik | -0.291836049 | 0.00000196 | 0.002534524 | Intergenic | -286803 |
| Bnc2 | 0.142294956 | 0.00000196 | 0.135673953 | intron (NM_172870, intron 2 of 5) | 129079 |
| Pgpep1l | 0.192634019 | 0.00000204 | 0.013243124 | intron (NM_010513, intron 2 of 20) | 115448 |
| Mir692-3 | 0.155575936 | 0.00000206 | 0.007717418 | intron (NM_001081097, intron 1 of 15) | -1052 |
| Trpv3 | -0.188881655 | 9.28E-20 | 0.000111795 | promoter-TSS (NM_145099) | 10 |
| Wipf1 | -0.267148536 | 0.0000021 | 0.004467638 | Intergenic | -11571 |
| Kcnip4 | 0.135597595 | 0.00000212 | 0.003180891 | Intergenic | -239209 |
| Prrxl1 | 0.151102374 | 0.00000213 | 0.002224643 | Intergenic | -4025 |
| Degs2 | -0.204796757 | 0.00000215 | 0.003065073 | exon (NM_027299, exon 3 of 3) | 11819 |
| Gm15679 | -0.291083294 | 0.00000221 | 0.000134164 | Intergenic | -735118 |
| Acta2 | -0.14182263 | 0.00000221 | 0.006388301 | intron (NM_007392, intron 1 of 8) | 229 |
| Slc25a51 | -0.141770557 | 0.00000222 | 0.002514466 | intron (NM_001033306, intron 2 of 5) | -55793 |
| Dctd | 0.117160433 | 0.00000222 | 0.006740548 | Intergenic | 97248 |

| | | | | | |
|---|---|---|---|---|---|
| Ces1g | 0.190524423 | 0.00000224 | 0.004651743 | promoter-TSS (NM_021456) | -189 |
| Bdh1 | -0.163294799 | 0.00000224 | 0.005090576 | Intergenic | 79692 |
| Scx | 0.134638158 | 0.00000225 | 0.005559008 | intron (NM_013481, intron 3 of 15) | 8461 |
| Pxdn | 0.156540719 | 0.00000227 | 0.004632476 | exon (NM_001093778, exon 21 of 25) | -42903 |
| Wfdc6a | 0.101260836 | 0.00000228 | 0.008530371 | promoter-TSS (NM_001177848) | 6 |
| Fancb | -0.178360981 | 0.0000023 | 0.003200303 | 3' UTR (NM_183427, exon 9 of 9) | 148609 |
| Aicda | 0.240093484 | 0.00000232 | 0.00525934 | promoter-TSS (NM_009645) | 84 |
| Ywhaz | -0.124995563 | 0.00000232 | 0.034803545 | Intergenic | -45748 |
| Rtn4 | -0.145164348 | 0.00000236 | 0.00303364 | exon (NM_024226, exon 1 of 7) | 293 |
| Epb4.1l4a | -0.193256393 | 0.00000236 | 0.005770013 | intron (NM_013512, intron 2 of 23) | 57621 |
| Nrxn3 | -0.118926996 | 0.00000239 | 0.001166093 | 5' UTR (NM_001198587, exon 1 of 20) | 553 |
| Tbca | 0.169064822 | 0.00000242 | 0.002573959 | Intergenic | -141599 |
| C87198 | 0.138208664 | 0.00000247 | 0.021461302 | Intergenic | -1454 |
| Naip2 | -0.122275745 | 0.00000248 | 0.005386338 | intron (NM_001126182, intron 1 of 15) | 1680 |
| Igsf5 | -0.195658215 | 0.0000025 | 0.002895269 | Intergenic | -7260 |
| Nxt1 | 0.157692022 | 0.00000252 | 0.002846663 | Intergenic | -100375 |
| 4933430N04Rik | 0.187705128 | 0.00000254 | 0.00238275 | Intergenic | 137993 |
| En1 | 0.138488432 | 1.21E-19 | 5.93E-08 | 5' UTR (NM_010133, exon 1 of 2) | 450 |
| 4933402C06Rik | -0.230702227 | 0.00000256 | 0.002741693 | Intergenic | -482997 |
| Mir6949 | 0.127840944 | 0.00000257 | 0.083212707 | promoter-TSS (NR_105914) | -29 |
| Grin2b | 0.275579084 | 0.00000258 | 0.011192585 | 5' UTR (NM_008171, exon 3 of 15) | 1827 |
| Lnx1 | -0.199764018 | 0.00000262 | 0.004200827 | 5' UTR (NM_001159580, exon 1 of 10) | 156 |
| Cyp2c29 | -0.302789359 | 0.00000263 | 0.004199748 | promoter-TSS (NM_007815) | 50 |
| Ttc34 | 0.173018175 | 0.00000263 | 0.010882416 | Intergenic | -75956 |
| Mcpt4 | 0.171462717 | 0.00000265 | 0.001247499 | intron (NM_010779, intron 1 of 4) | 476 |
| Pde7b | -0.286437124 | 0.00000265 | 0.003111593 | intron (NM_013875, intron 1 of 12) | 66974 |
| Slitrk4 | 0.161370257 | 0.0000027 | 0.045358775 | intron (NM_178740, intron 1 of 2) | 1444 |
| Ccrn4l | -0.260984698 | 1.35E-19 | 0.000000746 | Intergenic | -16531 |
| Ifna11 | 0.197064832 | 0.00000274 | 0.001645249 | promoter-TSS (NM_008333) | -95 |
| Cfap58 | -0.1437714 | 0.00000274 | 0.002693769 | Intergenic | 110567 |

Table S8: AR Cluster 1 Genes Top500

| geneSymbol | annotation | distanceToTSS |
|---|---|---|
| Fkbp5 | Promoter (2-3kb) | -2972 |
| H2-DMa | Promoter (<=1kb) | 254 |
| Per1 | Promoter (2-3kb) | 2087 |
| H2-DMa | Promoter (<=1kb) | 200 |
| Per1 | Promoter (1-2kb) | 2000 |
| H2-DMa | Promoter (1-2kb) | 1424 |
| H2-DMa | Promoter (<=1kb) | 0 |
| Brd2 | Promoter (1-2kb) | 1316 |
| 1700016A09Rik | Promoter (<=1kb) | 22 |
| Per1 | Promoter (1-2kb) | 1749 |
| 1700016A09Rik | Promoter (<=1kb) | 0 |
| 1700016A09Rik | Promoter (<=1kb) | 188 |
| Brd2 | Promoter (1-2kb) | 1179 |
| Irf2bp2 | Promoter (<=1kb) | -190 |
| 1700016A09Rik | Promoter (<=1kb) | 189 |
| Per1 | Promoter (1-2kb) | 1619 |
| Per1 | Intron (ENSMUST00000101004.8/18626, intron 1 of 17) | 3145 |
| Irf2bp2 | Promoter (<=1kb) | -342 |
| 1700016A09Rik | Promoter (<=1kb) | -23 |
| H2-DMa | Promoter (<=1kb) | -384 |
| Per1 | Promoter (1-2kb) | 1543 |
| Per1 | Intron (ENSMUST00000101004.8/18626, intron 1 of 17) | 3464 |
| Midn | Promoter (1-2kb) | 1707 |
| Midn | Promoter (<=1kb) | 811 |
| Per1 | 5' UTR | 3552 |
| H2-DMa | Promoter (<=1kb) | -426 |
| Midn | Promoter (1-2kb) | 1098 |
| Irf2bp2 | Promoter (<=1kb) | -695 |
| Midn | Promoter (<=1kb) | 602 |
| Irf2bp2 | Promoter (<=1kb) | -889 |
| Midn | Promoter (<=1kb) | 188 |
| Irf2bp2 | Promoter (<=1kb) | -953 |
| Midn | Promoter (<=1kb) | 0 |
| Per1 | 5' UTR | 3756 |
| Rhof | Exon (ENSMUST00000192791.2/ENSMUST00000192791.2, exon 1 of 1) | -3525 |
| Setd1b | Intron (ENSMUST00000160227.7/ENSMUST00000160227.7, intron 1 of 2) | -3489 |
| Tet3 | Promoter (2-3kb) | 2896 |
| Midn | Promoter (<=1kb) | 450 |
| Neat1 | Intron (ENSMUST00000187697.1/ENSMUST00000187697.1, intron 1 of 1) | -3767 |
| Midn | Promoter (<=1kb) | 396 |
| Irf2bp2 | Promoter (1-2kb) | -1419 |
| Tet3 | Promoter (2-3kb) | 2708 |

| | | |
|---|---|---|
| Ptma | Promoter (<=1kb) | 46 |
| Cdk2ap1 | 5' UTR | 10607 |
| Gm15417 | Promoter (<=1kb) | 971 |
| Neat1 | Promoter (1-2kb) | -1184 |
| Neat1 | Promoter (1-2kb) | -1267 |
| Neat1 | Promoter (1-2kb) | -1442 |
| Trim8 | Promoter (<=1kb) | 945 |
| H2-DMa | Promoter (<=1kb) | -885 |
| H2-DMa | Promoter (<=1kb) | -942 |
| Per1 | Intron (ENSMUST00000101004.8/18626, intron 1 of 17) | 3816 |
| Irf2bp2 | Promoter (1-2kb) | -1517 |
| Cbx8 | Distal Intergenic | -15401 |
| Klf13 | Promoter (1-2kb) | 1080 |
| Ptma | Promoter (<=1kb) | -371 |
| Klf13 | Promoter (1-2kb) | 1558 |
| Klf13 | Promoter (1-2kb) | 1543 |
| Setd1b | Exon (ENSMUST00000195992.1/ENSMUST00000195992.1, exon 1 of 1) | -3230 |
| Klf13 | Promoter (<=1kb) | 896 |
| Ptma | Promoter (<=1kb) | -125 |
| H2-DMa | Promoter (1-2kb) | -1096 |
| Ptma | Promoter (<=1kb) | -60 |
| Per1 | Intron (ENSMUST00000101004.8/18626, intron 1 of 17) | 3981 |
| Tet3 | Promoter (2-3kb) | 2527 |
| Klf13 | Promoter (1-2kb) | 1275 |
| Klf13 | Promoter (1-2kb) | 1265 |
| Irf2bp2 | Promoter (1-2kb) | -1699 |
| Fkbp5 | Promoter (<=1kb) | -875 |
| Midn | Promoter (<=1kb) | -240 |
| Setd1b | Promoter (2-3kb) | -2514 |
| Zbtb7b | Promoter (<=1kb) | 416 |
| Trib1 | Promoter (1-2kb) | 1146 |
| Fkbp5 | Promoter (<=1kb) | -848 |
| Ppp1r9b | Promoter (1-2kb) | 1247 |
| Setd1b | Exon (ENSMUST00000192791.2/ENSMUST00000192791.2, exon 1 of 1) | -3950 |
| Tet3 | Promoter (2-3kb) | 2451 |
| Zbtb7b | Promoter (<=1kb) | 765 |
| Cdk2ap1 | Intron (ENSMUST00000031341.10/13445, intron 1 of 3) | 10392 |
| Trim8 | Promoter (1-2kb) | 1935 |
| Id1 | Distal Intergenic | 3316 |
| Ubtf | Promoter (<=1kb) | 330 |
| 0610009L18Rik | Promoter (2-3kb) | 2014 |
| Ppp1r9b | Promoter (<=1kb) | 679 |
| Tet3 | Promoter (1-2kb) | 1775 |

| | | |
|---|---|---|
| Vwf | Intron (ENSMUST00000100941.8/22371, intron 8 of 10) | 42437 |
| Id3 | Promoter (<=1kb) | -571 |
| Cic | Intron (ENSMUST00000169266.7/71722, intron 2 of 20) | 13543 |
| 1700016A09Rik | Promoter (1-2kb) | 1731 |
| Vwf | Exon (ENSMUST00000100941.8/22371, exon 8 of 11) | 41345 |
| Klf13 | Promoter (<=1kb) | 631 |
| Neat1 | 3' UTR | -4595 |
| Rasd1 | 3' UTR | 3108 |
| Rhof | Promoter (2-3kb) | -2867 |
| Fkbp5 | Promoter (<=1kb) | -591 |
| Irf2bp2 | Promoter (<=1kb) | 636 |
| Id1 | Distal Intergenic | 3246 |
| Cdk2ap1 | Intron (ENSMUST00000031341.10/13445, intron 1 of 3) | 10275 |
| Cbx8 | Distal Intergenic | -13448 |
| Ntn5 | Intron (ENSMUST00000040636.8/56546, intron 3 of 3) | -3712 |
| Trib1 | Promoter (<=1kb) | 992 |
| Neat1 | Promoter (<=1kb) | -478 |
| 0610009L18Rik | Promoter (1-2kb) | 1837 |
| Ptma | Promoter (1-2kb) | -1042 |
| Neat1 | 3' UTR | -4693 |
| Id1 | Promoter (2-3kb) | 2909 |
| Id1 | Distal Intergenic | 3003 |
| Ntn5 | Intron (ENSMUST00000040636.8/56546, intron 3 of 3) | -4027 |
| Vwf | Intron (ENSMUST00000100941.8/22371, intron 8 of 10) | 42923 |
| Ntn5 | Intron (ENSMUST00000040636.8/56546, intron 3 of 3) | -3753 |
| Cdk2ap1 | 5' UTR | 11495 |
| Vwf | Exon (ENSMUST00000100941.8/22371, exon 9 of 11) | 43348 |
| Pcgf2 | Promoter (<=1kb) | -438 |
| Id1 | Promoter (2-3kb) | 2770 |
| 0610009L18Rik | Promoter (1-2kb) | 1591 |
| Dalrd3 | Promoter (<=1kb) | -543 |
| Ntn5 | Intron (ENSMUST00000040636.8/56546, intron 3 of 3) | -3322 |
| Neat1 | Promoter (<=1kb) | -72 |
| Actb | Promoter (<=1kb) | 653 |
| Mir7687 | Promoter (<=1kb) | 65 |
| Id3 | Promoter (<=1kb) | -51 |
| Per1 | Promoter (<=1kb) | 382 |
| Hnrnpl | Promoter (2-3kb) | 2473 |
| Tnfrsf25 | 5' UTR | 15108 |
| Fosl2 | Promoter (<=1kb) | -491 |
| Zbtb16 | Intron (ENSMUST00000093852.4/235320, intron 3 of 6) | 112842 |
| Ier5l | Promoter (1-2kb) | 1159 |
| Zbtb16 | Intron (ENSMUST00000093852.4/235320, intron 3 of 6) | 112281 |

| | | |
|---|---|---|
| NA | Exon (ENSMUST00000147959.1/665635, exon 2 of 2) | 18856 |
| Zbtb16 | Intron (ENSMUST00000093852.4/235320, intron 3 of 6) | 112795 |
| Neat1 | 3' UTR | -4950 |
| Fosl2 | Promoter (<=1kb) | -181 |
| Trib1 | Promoter (<=1kb) | 916 |
| Neat1 | Promoter (<=1kb) | -23 |
| Fasn | Promoter (<=1kb) | 674 |
| Fosl2 | Promoter (<=1kb) | -84 |
| Junb | Promoter (1-2kb) | 1456 |
| Fosl2 | Promoter (<=1kb) | 0 |
| Zmynd8 | Intron (ENSMUST00000109262.7/228880, intron 1 of 10) | 13827 |
| Tgif1 | Promoter (1-2kb) | 1605 |
| Trib1 | Promoter (<=1kb) | 0 |
| Fkbp5 | Promoter (1-2kb) | -1343 |
| Hnrnpl | Promoter (2-3kb) | 2085 |
| Trib1 | Promoter (<=1kb) | 275 |
| Tbx3 | Promoter (<=1kb) | -532 |
| Zmynd8 | Intron (ENSMUST00000109262.7/228880, intron 1 of 10) | 13968 |
| Zbtb7a | Promoter (<=1kb) | 762 |
| Tnrc18 | Distal Intergenic | -3334 |
| Ier5l | Promoter (<=1kb) | 861 |
| Maz | Promoter (1-2kb) | 1179 |
| Hnrnpl | Promoter (2-3kb) | 2248 |
| Ier5l | Promoter (<=1kb) | 935 |
| 0610009L18Rik | Promoter (1-2kb) | 1543 |
| Tnfrsf25 | 5' UTR | 15348 |
| NA | Intron (ENSMUST00000146702.1/ENSMUST00000146702.1, intron 2 of 3) | 21089 |
| Tnfrsf25 | Exon (ENSMUST00000150958.7/56226, exon 1 of 7) | 15509 |
| 0610009L18Rik | Promoter (<=1kb) | 867 |
| Zbtb7a | Promoter (<=1kb) | 998 |
| Id3 | Promoter (1-2kb) | 1060 |
| Zmiz1 | Promoter (<=1kb) | 0 |
| Mir1191b | 5' UTR | -14484 |
| Ptma | Promoter (1-2kb) | -1442 |
| Junb | Promoter (<=1kb) | 890 |
| Sox1ot | Promoter (<=1kb) | 0 |
| Tob1 | Distal Intergenic | -49803 |
| Ptma | Promoter (1-2kb) | 1492 |
| Midn | Promoter (1-2kb) | -1363 |
| Zbtb16 | Intron (ENSMUST00000093852.4/235320, intron 3 of 6) | 98290 |
| Rasd1 | 3' UTR | 3672 |
| Ier5l | Promoter (<=1kb) | 606 |
| 0610009L18Rik | Promoter (1-2kb) | 1156 |

| | | |
|---|---|---|
| Tnrc18 | Promoter (<=1kb) | -472 |
| Klf13 | Promoter (<=1kb) | 343 |
| Zbtb16 | Intron (ENSMUST00000093852.4/235320, intron 3 of 6) | 98426 |
| Tnfrsf25 | Intron (ENSMUST00000030785.14/56226, intron 10 of 15) | 14222 |
| Ier5l | Promoter (1-2kb) | 1702 |
| Zmynd8 | 5' UTR | 14089 |
| Id1 | Promoter (<=1kb) | 278 |
| Tex44 | Distal Intergenic | 14017 |
| Mir191 | Promoter (<=1kb) | 131 |
| Ier5l | Promoter (2-3kb) | 2139 |
| Per1 | Intron (ENSMUST00000101004.8/18626, intron 1 of 17) | 4593 |
| Trib1 | Promoter (<=1kb) | -11 |
| Fkbp5 | Promoter (<=1kb) | -281 |
| Dbp | Promoter (2-3kb) | 2861 |
| Zbtb7a | Promoter (1-2kb) | 1645 |
| Tet3 | Promoter (1-2kb) | 1375 |
| Id3 | Promoter (<=1kb) | 0 |
| Cic | Intron (ENSMUST00000169266.7/71722, intron 2 of 20) | 13998 |
| Mir1191b | Intron (ENSMUST00000221817.1/18029, intron 1 of 9) | -16935 |
| Fosl2 | Promoter (<=1kb) | 115 |
| Tex44 | Distal Intergenic | 14014 |
| Ier5l | Promoter (<=1kb) | 484 |
| Junb | Promoter (2-3kb) | 2045 |
| Zbtb7a | Promoter (<=1kb) | 733 |
| Zbtb16 | Intron (ENSMUST00000093852.4/235320, intron 3 of 6) | 112102 |
| Tob1 | Distal Intergenic | -50097 |
| Gm15417 | Promoter (<=1kb) | 175 |
| Fasn | Promoter (<=1kb) | 288 |
| Tet3 | Promoter (1-2kb) | 1305 |
| Hes1 | Promoter (<=1kb) | 900 |
| Zmiz1 | Promoter (1-2kb) | 1093 |
| Per1 | Promoter (<=1kb) | 0 |
| Fasn | Promoter (<=1kb) | 103 |
| Zmynd8 | Intron (ENSMUST00000109262.7/228880, intron 1 of 10) | 12838 |
| Ppp1r9b | Promoter (<=1kb) | 0 |
| Ier5l | Promoter (<=1kb) | 277 |
| Ppp1r9b | Promoter (<=1kb) | 134 |
| Tbx3 | Promoter (<=1kb) | -298 |
| Zfp36l2 | Promoter (<=1kb) | 0 |
| Id3 | Promoter (<=1kb) | 223 |
| Dbp | Promoter (1-2kb) | 1431 |
| Fkbp5 | Exon (ENSMUST00000180802.1/ENSMUST00000180802.1, exon 2 of 2) | -4857 |
| Traf4 | Promoter (1-2kb) | 1512 |

| | | |
|---|---|---|
| Tle3 | Promoter (<=1kb) | -674 |
| Actb | Promoter (<=1kb) | 341 |
| Mir1942 | Intron (ENSMUST00000054449.13/18810, intron 1 of 32) | 16980 |
| Id2 | Promoter (1-2kb) | -1048 |
| Neat1 | Promoter (1-2kb) | 1985 |
| Fkbp5 | Promoter (<=1kb) | -112 |
| Tet3 | Promoter (1-2kb) | 1181 |
| Id3 | Promoter (<=1kb) | 311 |
| Fasn | Promoter (<=1kb) | 13 |
| Tnrc18 | Promoter (<=1kb) | 0 |
| Ppp1r9b | Promoter (<=1kb) | -67 |
| Trib1 | Promoter (<=1kb) | -303 |
| Pcgf2 | Promoter (<=1kb) | 0 |
| Hoxa9 | Exon (ENSMUST00000125581.1/15395, exon 1 of 2) | -3604 |
| Mknk2 | Promoter (1-2kb) | 1328 |
| Tnrc18 | Promoter (<=1kb) | -17 |
| Cyb561 | Exon (ENSMUST00000195385.1/ENSMUST00000195385.1, exon 1 of 1) | 8408 |
| Neat1 | Promoter (1-2kb) | 1909 |
| Klf13 | Promoter (<=1kb) | 167 |
| Ucp2 | Promoter (2-3kb) | 2529 |
| Zfp36l2 | Promoter (<=1kb) | -261 |
| Ankrd13b | Promoter (1-2kb) | -1241 |
| Tob1 | Distal Intergenic | -50408 |
| Tob1 | Distal Intergenic | -50419 |
| Ucp2 | Promoter (2-3kb) | 2035 |
| Fkbp5 | Exon (ENSMUST00000181029.2/ENSMUST00000181029.2, exon 1 of 1) | -5275 |
| Tnrc18 | Promoter (<=1kb) | -121 |
| Fkbp5 | Exon (ENSMUST00000181029.2/ENSMUST00000181029.2, exon 1 of 1) | -5206 |
| Maz | Promoter (<=1kb) | -55 |
| Rasd1 | 3' UTR | 3703 |
| Git1 | Promoter (1-2kb) | -1461 |
| E130317F20Rik | Promoter (<=1kb) | 0 |
| Cyb561 | Exon (ENSMUST00000195385.1/ENSMUST00000195385.1, exon 1 of 1) | 8193 |
| Rasd1 | 3' UTR | 4035 |
| Klf10 | Promoter (<=1kb) | 482 |
| Ier5l | Promoter (<=1kb) | 99 |
| Actb | Promoter (<=1kb) | 0 |
| Actb | Promoter (<=1kb) | 9 |
| Rasd1 | Exon (ENSMUST00000081980.6/192191, exon 2 of 2) | 4239 |
| Tex44 | Distal Intergenic | 13822 |
| Ubtf | Promoter (1-2kb) | 1964 |
| Zbtb7a | Promoter (1-2kb) | 1435 |
| NA | Promoter (<=1kb) | 110 |

| | | |
|---|---|---|
| Fasn | Promoter (<=1kb) | -164 |
| Tex44 | Distal Intergenic | 16432 |
| Mir21a | Distal Intergenic | 4185 |
| Ncor2 | Promoter (<=1kb) | 0 |
| Cbx8 | Distal Intergenic | -12758 |
| Id3 | Promoter (<=1kb) | 670 |
| Id3 | Promoter (<=1kb) | 313 |
| Id2 | Promoter (1-2kb) | -1583 |
| Mknk2 | Promoter (1-2kb) | 1660 |
| 1700016A09Rik | 5' UTR | 48678 |
| Brd2 | Promoter (<=1kb) | 143 |
| Vwf | Intron (ENSMUST00000100941.8/22371, intron 7 of 10) | 39543 |
| Gm15417 | Promoter (<=1kb) | 11 |
| Tob1 | Distal Intergenic | -50858 |
| Tbx3 | Promoter (<=1kb) | 0 |
| Actb | Promoter (<=1kb) | -177 |
| Fasn | Promoter (<=1kb) | -186 |
| Id3 | Promoter (<=1kb) | 569 |
| Mknk2 | Promoter (<=1kb) | 835 |
| Tle3 | Promoter (1-2kb) | 1505 |
| Zbtb16 | Intron (ENSMUST00000093852.4/235320, intron 3 of 6) | 111829 |
| Cyb561 | 5' UTR | 9213 |
| Fkbp5 | Promoter (1-2kb) | 1222 |
| Klf13 | Promoter (2-3kb) | 2487 |
| Zbtb16 | Intron (ENSMUST00000093852.4/235320, intron 3 of 6) | 98775 |
| Cyb561 | Exon (ENSMUST00000195385.1/ENSMUST00000195385.1, exon 1 of 1) | 7970 |
| Sin3a | Promoter (1-2kb) | 1450 |
| Neat1 | Promoter (1-2kb) | 1734 |
| E130317F20Rik | Promoter (<=1kb) | 51 |
| Baz2a | Promoter (2-3kb) | 2521 |
| Hes1 | Promoter (<=1kb) | -979 |
| Fkbp5 | Exon (ENSMUST00000181029.2/ENSMUST00000181029.2, exon 1 of 1) | -5284 |
| 1700016A09Rik | Intron (ENSMUST00000127664.1/108148, intron 1 of 15) | 46665 |
| Cyb561 | Exon (ENSMUST00000195385.1/ENSMUST00000195385.1, exon 1 of 1) | 9381 |
| Dbp | Promoter (<=1kb) | 865 |
| Sox1ot | Promoter (<=1kb) | -316 |
| Socs3 | Distal Intergenic | 3545 |
| Klf10 | Promoter (<=1kb) | -584 |
| Tob1 | Distal Intergenic | -49336 |
| Zfp36l2 | Promoter (<=1kb) | -520 |
| Egr1 | Promoter (1-2kb) | 1775 |
| Mir1191b | Intron (ENSMUST00000117966.1/18029, intron 1 of 9) | -13424 |
| Atp1b1 | Promoter (<=1kb) | -453 |

| | | |
|---|---|---|
| Esyt3 | Promoter (1-2kb) | 1402 |
| Mknk2 | Promoter (1-2kb) | 1125 |
| Ier5l | Promoter (<=1kb) | 0 |
| Klf13 | Promoter (<=1kb) | 82 |
| 1700016A09Rik | Intron (ENSMUST00000127664.1/108148, intron 1 of 15) | 48096 |
| Phf12 | Promoter (<=1kb) | -417 |
| Pfn1 | Promoter (<=1kb) | 758 |
| Cnn2 | Promoter (1-2kb) | -1506 |
| Id3 | Distal Intergenic | 3144 |
| Atp1b1 | Promoter (1-2kb) | -1350 |
| Zbtb16 | Intron (ENSMUST00000093852.4/235320, intron 3 of 6) | 96741 |
| Slc25a23 | Promoter (1-2kb) | -1121 |
| Atp1b1 | Promoter (<=1kb) | 508 |
| Traf4 | Promoter (<=1kb) | 254 |
| Atp1b1 | Promoter (<=1kb) | -208 |
| Gm15417 | Promoter (<=1kb) | 0 |
| Zfp36l2 | Promoter (<=1kb) | -603 |
| Scd2 | Promoter (1-2kb) | 1375 |
| Def8 | Intron (ENSMUST00000127664.1/108148, intron 1 of 15) | 3937 |
| Cbx4 | Promoter (<=1kb) | 594 |
| Sh2b3 | Promoter (<=1kb) | 902 |
| Cyb561 | 5' UTR | 8933 |
| Egr1 | Promoter (2-3kb) | 2426 |
| Cyb561 | Exon (ENSMUST00000195385.1/ENSMUST00000195385.1, exon 1 of 1) | 8613 |
| Tnfrsf25 | Exon (ENSMUST00000150958.7/56226, exon 1 of 7) | 16177 |
| Atp1b1 | Promoter (<=1kb) | -607 |
| Arid1a | Promoter (2-3kb) | 2369 |
| Cyb561 | 5' UTR | 8730 |
| Kdm6b | Intron (ENSMUST00000094077.4/216850, intron 1 of 22) | 3920 |
| Cbarp | Promoter (1-2kb) | 1385 |
| Mir1191b | Intron (ENSMUST00000117966.1/18029, intron 1 of 9) | -13963 |
| Pfn1 | Promoter (<=1kb) | 110 |
| Fasn | Promoter (<=1kb) | -368 |
| Gramd1b | Intron (ENSMUST00000119373.8/235283, intron 1 of 20) | 72318 |
| Bhlhe41 | Promoter (<=1kb) | 0 |
| Cpn2 | Distal Intergenic | 80002 |
| Tle3 | Promoter (<=1kb) | 840 |
| Cic | 5' UTR | 14508 |
| Phf12 | Promoter (<=1kb) | -398 |
| Sh2b3 | Promoter (1-2kb) | 1085 |
| Pim3 | Distal Intergenic | -10073 |
| E130317F20Rik | Promoter (<=1kb) | -238 |
| Lfng | Exon (ENSMUST00000200626.4/16848, exon 1 of 6) | 4054 |

| | | |
|---|---|---|
| Ddit3 | Promoter (<=1kb) | -338 |
| Hes1 | Promoter (1-2kb) | -1994 |
| Baiap2 | Promoter (<=1kb) | 336 |
| Sh2b3 | Promoter (<=1kb) | 863 |
| Baiap2 | Promoter (<=1kb) | 488 |
| Baiap2 | Promoter (<=1kb) | 243 |
| Scd2 | Promoter (2-3kb) | 2175 |
| Dbp | Promoter (<=1kb) | 12 |
| Pafah1b3 | 5' UTR | 14467 |
| Cbx4 | Promoter (<=1kb) | 7 |
| Hes1 | Promoter (1-2kb) | -1002 |
| Brd2 | Promoter (<=1kb) | 72 |
| Baiap2 | Promoter (<=1kb) | 451 |
| Irf2bp2 | Promoter (2-3kb) | -2438 |
| Cic | 5' UTR | 14512 |
| Sox1ot | Promoter (1-2kb) | 1023 |
| Phf12 | Promoter (<=1kb) | -290 |
| A230083G16Rik | Promoter (<=1kb) | 840 |
| Jun | Promoter (<=1kb) | -288 |
| Pim3 | Promoter (<=1kb) | -334 |
| Zfp36l2 | Promoter (<=1kb) | -633 |
| Ptch1 | 5' UTR | 8074 |
| Tle3 | Promoter (<=1kb) | 154 |
| Zbtb7a | Intron (ENSMUST00000125261.1/16969, intron 1 of 1) | 3176 |
| Cbx4 | Promoter (<=1kb) | -352 |
| Klf10 | Promoter (<=1kb) | 59 |
| Tmem259 | Promoter (1-2kb) | -1225 |
| Egr1 | Promoter (<=1kb) | 0 |
| Plekhh3 | Promoter (<=1kb) | -353 |
| Atp1b1 | Promoter (<=1kb) | -684 |
| Pim3 | Promoter (<=1kb) | 0 |
| Hmga1b | Promoter (<=1kb) | 119 |
| Ddit3 | Promoter (<=1kb) | -316 |
| Tbx3 | Promoter (<=1kb) | 324 |
| Cbx4 | Promoter (<=1kb) | 329 |
| Zmiz1 | Promoter (<=1kb) | -591 |
| Jun | Promoter (<=1kb) | 0 |
| Mus81 | Promoter (<=1kb) | -485 |
| Cbx4 | Promoter (<=1kb) | 0 |
| Dalrd3 | Promoter (<=1kb) | 0 |
| Zfp36l2 | Promoter (<=1kb) | -835 |
| Pim3 | Promoter (<=1kb) | -374 |
| Klf16 | Promoter (<=1kb) | -93 |

| | | |
|---|---|---|
| Upf1 | Promoter (1-2kb) | 1748 |
| Fkbp5 | Exon (ENSMUST00000181029.2/ENSMUST00000181029.2, exon 1 of 1) | -5903 |
| Pim3 | Promoter (<=1kb) | -104 |
| Hs3st3b1 | Promoter (<=1kb) | 0 |
| Zfp36l2 | Promoter (<=1kb) | -805 |
| Mir1191b | Intron (ENSMUST00000117966.1/18029, intron 1 of 9) | -13801 |
| Pcgf2 | Promoter (<=1kb) | 326 |
| Vwf | Intron (ENSMUST00000100941.8/22371, intron 7 of 10) | 28456 |
| Baiap2 | Promoter (<=1kb) | 656 |
| Pim3 | Distal Intergenic | -10640 |
| Plekhh3 | Promoter (<=1kb) | -551 |
| Pim3 | Distal Intergenic | -11202 |
| Klf16 | Promoter (1-2kb) | -1864 |
| Sh2b3 | Promoter (<=1kb) | 0 |
| Kdm6b | Intron (ENSMUST00000094077.4/216850, intron 1 of 22) | 3207 |
| Traf4 | Promoter (<=1kb) | 148 |
| Vwf | Intron (ENSMUST00000100941.8/22371, intron 7 of 10) | 28634 |
| Lfng | Intron (ENSMUST00000031555.2/16848, intron 1 of 7) | 3017 |
| Tob1 | Distal Intergenic | -50986 |
| Vat1 | Promoter (<=1kb) | 109 |
| Ucp2 | Promoter (<=1kb) | 648 |
| Itga3 | Promoter (<=1kb) | 214 |
| Mir195a | Promoter (2-3kb) | 2234 |
| Vat1 | Promoter (<=1kb) | 0 |
| Tle3 | Intron (ENSMUST00000178113.7/21887, intron 4 of 19) | 3829 |
| E130317F20Rik | Promoter (<=1kb) | 59 |
| Pim3 | Distal Intergenic | -10514 |
| Plekhh3 | Promoter (<=1kb) | -589 |
| Fkbp5 | Exon (ENSMUST00000181029.2/ENSMUST00000181029.2, exon 1 of 1) | -5938 |
| Junb | Promoter (2-3kb) | 2555 |
| Ssbp4 | Promoter (<=1kb) | -357 |
| Pim3 | Distal Intergenic | -11643 |
| Rcc2 | Promoter (<=1kb) | 950 |
| Pim3 | Distal Intergenic | -11237 |
| Ddit3 | Promoter (<=1kb) | 0 |
| Klhl21 | Promoter (<=1kb) | 448 |
| Klf16 | Promoter (<=1kb) | 293 |
| Kdm6b | Intron (ENSMUST00000094077.4/216850, intron 1 of 22) | 4374 |
| Bcar1 | Promoter (<=1kb) | 136 |
| Pim3 | Distal Intergenic | -11394 |
| Mir1191b | Intron (ENSMUST00000117966.1/18029, intron 1 of 9) | -12939 |
| Zswim6 | Promoter (<=1kb) | -23 |
| Cdk2ap1 | Intron (ENSMUST00000031341.10/13445, intron 1 of 3) | 9689 |

| | | |
|---|---|---|
| Lmna | Promoter (<=1kb) | 121 |
| Ncor2 | Intron (ENSMUST00000111402.8/20602, intron 1 of 47) | 3034 |
| Klf13 | Promoter (<=1kb) | -34 |
| Pim3 | Distal Intergenic | -10138 |
| Git1 | Promoter (<=1kb) | 16 |
| Ubtf | Promoter (<=1kb) | -473 |
| Junb | Promoter (<=1kb) | -862 |
| Klf16 | Promoter (<=1kb) | 508 |
| Id1 | Distal Intergenic | 4817 |
| Klf16 | Promoter (<=1kb) | 623 |
| Rasd1 | Promoter (<=1kb) | 615 |
| Ddr1 | Promoter (1-2kb) | 1224 |
| Pim3 | Promoter (<=1kb) | 267 |
| Pim3 | Distal Intergenic | -11659 |
| Mbd6 | Promoter (<=1kb) | 0 |
| Vwf | Intron (ENSMUST00000100941.8/22371, intron 7 of 10) | 28257 |
| Klf16 | Promoter (<=1kb) | 85 |
| Mbd6 | Promoter (<=1kb) | -498 |
| Bhlhe41 | Promoter (1-2kb) | 1175 |
| Dusp4 | Promoter (<=1kb) | 0 |
| Brd2 | Promoter (<=1kb) | 0 |
| Mus81 | Promoter (<=1kb) | -291 |
| Pim3 | Distal Intergenic | -10353 |
| Sin3a | Promoter (1-2kb) | 1199 |
| Pisd | Exon (ENSMUST00000201980.4/320951, exon 2 of 6) | 85078 |
| Ntn5 | Distal Intergenic | -6535 |
| Vars | Promoter (1-2kb) | -1828 |
| Trim8 | Distal Intergenic | 28832 |
| Traf4 | Promoter (<=1kb) | 0 |
| Tbx3 | Promoter (1-2kb) | -1819 |
| Traf4 | Promoter (<=1kb) | 73 |
| Mir21a | Distal Intergenic | 4550 |
| Wasf2 | Distal Intergenic | -45081 |
| Maz | Promoter (2-3kb) | 2463 |
| Klf16 | Promoter (<=1kb) | 224 |
| Egr1 | Promoter (1-2kb) | 1037 |
| Phf12 | Promoter (<=1kb) | -70 |
| Chd3 | Promoter (1-2kb) | 1877 |
| Hes1 | Promoter (1-2kb) | -1234 |
| Mir5135 | Promoter (<=1kb) | 0 |
| Egr1 | Promoter (1-2kb) | 1624 |
| Pfn1 | Promoter (<=1kb) | -549 |
| Zswim6 | Promoter (<=1kb) | 134 |

| | | |
|---|---|---|
| Cnn2 | Promoter (1-2kb) | -1106 |
| Ubald1 | Promoter (<=1kb) | 618 |
| Fkbp5 | Intron (ENSMUST00000114792.7/14229, intron 1 of 13) | 4232 |
| Vat1 | Promoter (<=1kb) | 569 |
| Mrps18a | Intron (ENSMUST00000233284.1/ENSMUST00000233284.1, intron 1 of 2) | -36701 |
| Zfp36l2 | Promoter (<=1kb) | 246 |
| Cdkn1a | Intron (ENSMUST00000119901.8/12575, intron 1 of 2) | 4610 |
| Atp5d | Promoter (<=1kb) | 0 |
| Mus81 | Promoter (<=1kb) | -192 |
| Dus3l | Promoter (1-2kb) | -1409 |
| Klf16 | Promoter (1-2kb) | -1039 |
| Fbxl15 | Promoter (1-2kb) | 1185 |
| Adamtsl4 | Distal Intergenic | 13258 |
| Tns2 | Promoter (<=1kb) | 0 |
| Mex3b | Promoter (<=1kb) | 47 |
| Cbx8 | Distal Intergenic | -17337 |
| Ddit3 | Promoter (<=1kb) | 90 |
| Cdkn1a | Exon (ENSMUST00000233069.1/ENSMUST00000233069.1, exon 1 of 1) | 3453 |
| Adgrg1 | Promoter (1-2kb) | 1542 |
| Cyb561 | Intron (ENSMUST00000143251.7/13056, intron 1 of 4) | 6555 |
| Klf10 | Promoter (<=1kb) | 0 |
| Lfng | Promoter (1-2kb) | 1465 |
| Hnrnpl | Intron (ENSMUST00000174548.7/15388, intron 2 of 13) | 3935 |
| Klf10 | Promoter (<=1kb) | -177 |
| Mrps18a | Intron (ENSMUST00000233284.1/ENSMUST00000233284.1, intron 1 of 2) | -36597 |
| Cables2 | Promoter (<=1kb) | 0 |
| Garin4 | Intron (ENSMUST00000195117.1/11910, intron 1 of 3) | -18815 |

Table S8: AR Cluster 2 Genes Top500

| geneSymbol | annotation | distanceToTSS |
|---|---|---|
| Fgd5 | Intron (ENSMUST00000089334.8/232237, intron 2 of 20) | 18357 |
| Gli2 | Promoter (<=1kb) | 0 |
| Mir704 | Distal Intergenic | 57295 |
| Fam76a | Distal Intergenic | -18507 |
| Gjb3 | Intron (ENSMUST00000106091.8/14620, intron 1 of 1) | 3189 |
| Dynlt1b | Intron (ENSMUST00000180035.7/547127, intron 8 of 8) | 11803 |
| NA | Exon (ENSMUST00000215081.1/270210, exon 2 of 2) | -5719 |
| Zkscan17 | Intron (ENSMUST00000139093.7/268417, intron 1 of 2) | 8181 |
| Zfp513 | Promoter (1-2kb) | 1950 |
| NA | Distal Intergenic | -27124 |
| Clec4f | 3' UTR | 10951 |
| 2310016G11Rik | Promoter (1-2kb) | -1704 |
| Cmip | Intron (ENSMUST00000166750.8/74440, intron 1 of 20) | 8047 |
| Mir704 | Distal Intergenic | 57481 |
| Fosl2 | Exon (ENSMUST00000201110.1/ENSMUST00000201110.1, exon 1 of 1) | 3796 |
| Shisal1 | Intron (ENSMUST00000080751.8/72301, intron 3 of 4) | 34699 |
| D630045J12Rik | Intron (ENSMUST00000117556.7/330286, intron 1 of 19) | 49622 |
| Or1d2 | Intron (ENSMUST00000102521.8/380711, intron 5 of 24) | 72956 |
| Ifitm3 | Distal Intergenic | 6032 |
| 4930451I11Rik | Intron (ENSMUST00000205324.1/68952, intron 5 of 8) | 10708 |
| Col16a1 | Intron (ENSMUST00000044565.14/107581, intron 10 of 69) | 7862 |
| Fndc5 | Intron (ENSMUST00000102600.3/384061, intron 5 of 5) | 5284 |
| Piwil1 | Intron (ENSMUST00000199537.4/231760, intron 2 of 16) | 106328 |
| Anks6 | Intron (ENSMUST00000107747.7/75691, intron 1 of 16) | 7133 |
| 5830454E08Rik | Distal Intergenic | 111695 |
| Abtb1 | Intron (ENSMUST00000113585.8/23945, intron 7 of 7) | 17479 |
| 9130230L23Rik | Promoter (2-3kb) | 2730 |
| Spr | Intron (ENSMUST00000203386.1/ENSMUST00000203386.1, intron 1 of 2) | 15120 |
| Slc16a3 | 3' UTR | 14005 |
| Sdc4 | Intron (ENSMUST00000017153.3/20971, intron 1 of 4) | 11893 |
| Hdac11 | Distal Intergenic | 21122 |
| Dnajb8 | Promoter (<=1kb) | 355 |
| Scx | 3' UTR | -4546 |
| Slc25a33 | Distal Intergenic | -36359 |
| Mt1 | Distal Intergenic | 15620 |
| Ahdc1 | Intron (ENSMUST00000154646.7/230793, intron 2 of 3) | 11941 |
| Hmgcl | Downstream (<=300bp) | -5639 |
| NA | Promoter (<=1kb) | -90 |
| Cdk18 | Intron (ENSMUST00000027697.11/18557, intron 1 of 15) | 7987 |
| Asap3 | Distal Intergenic | -5952 |
| Dysf | 5' UTR | 10609 |
| Mir129-1 | Promoter (1-2kb) | 1453 |

| | | |
|---|---|---|
| Gimap3 | Promoter (1-2kb) | 1226 |
| Hrh1 | Distal Intergenic | 95243 |
| Hspb1 | Intron (ENSMUST00000013615.13/58212, intron 12 of 15) | -19827 |
| Rpusd3 | 3' UTR | 10081 |
| Mir6937 | Intron (ENSMUST00000168129.9/217449, intron 8 of 11) | 20886 |
| Iqgap2 | Intron (ENSMUST00000068603.7/544963, intron 1 of 35) | 3866 |
| Hrh1 | Distal Intergenic | 95124 |
| Map3k9 | Intron (ENSMUST00000035987.8/338372, intron 2 of 10) | 29659 |
| Cartpt | Distal Intergenic | 33814 |
| Smim38 | Distal Intergenic | 100193 |
| Gm10406 | Distal Intergenic | 135929 |
| Edf1 | Downstream (<=300bp) | 4396 |
| Sec13 | Intron (ENSMUST00000101045.9/11941, intron 4 of 21) | -80442 |
| Atp6v1e1 | Exon (ENSMUST00000203527.1/11973, exon 1 of 3) | 21270 |
| Tmem141 | Promoter (<=1kb) | -235 |
| Mir5122 | Intron (ENSMUST00000030669.7/20544, intron 1 of 11) | 8738 |
| Ube2v1 | Intron (ENSMUST00000125544.2/66589, intron 2 of 7) | 8852 |
| Stk40 | 5' UTR | 13487 |
| Apobec3 | Distal Intergenic | -11044 |
| 1700001K23Rik | Distal Intergenic | -12208 |
| Prrc2b | Intron (ENSMUST00000036691.13/227723, intron 1 of 29) | 19698 |
| Ptpn1 | Distal Intergenic | -87150 |
| Apobec3 | Distal Intergenic | -11158 |
| Wnk2 | Intron (ENSMUST00000162581.7/75607, intron 3 of 27) | 46041 |
| Ppard | Intron (ENSMUST00000002320.15/19015, intron 3 of 7) | 19735 |
| Ccm2l | 3' UTR | 14999 |
| Slc25a33 | Distal Intergenic | -36357 |
| Hrh1 | Distal Intergenic | 95051 |
| Sergef | Intron (ENSMUST00000033127.11/27414, intron 8 of 10) | 30192 |
| Gsdmd | Promoter (2-3kb) | 2350 |
| Asprv1 | Distal Intergenic | -30263 |
| 4930521O11Rik | Intron (ENSMUST00000181509.1/74715, intron 3 of 3) | 15793 |
| Tamm41 | Promoter (<=1kb) | 0 |
| Tbc1d16 | Intron (ENSMUST00000036113.3/207592, intron 3 of 11) | 21398 |
| Mmp24 | Promoter (<=1kb) | 154 |
| Trim67 | Exon (ENSMUST00000041106.8/330863, exon 3 of 11) | 23923 |
| Ncf2 | Promoter (1-2kb) | -1912 |
| Fbxl14 | Intron (ENSMUST00000178696.7/22419, intron 1 of 4) | -9243 |
| Gm29719 | Intron (ENSMUST00000232402.1/101055731, intron 2 of 10) | 8805 |
| Spsb1 | Intron (ENSMUST00000146532.1/ENSMUST00000146532.1, intron 1 of 2) | 50144 |
| 1700017B05Rik | Distal Intergenic | -13785 |
| Trim13 | Promoter (1-2kb) | 1015 |
| 9630028I04Rik | Distal Intergenic | 16737 |

| | | |
|---|---|---|
| Cemip2 | Intron (ENSMUST00000096194.8/83921, intron 12 of 24) | 38035 |
| Tacc2 | Intron (ENSMUST00000207918.1/57752, intron 1 of 6) | 14283 |
| Apobec3 | Distal Intergenic | -11197 |
| Cilp2 | Promoter (1-2kb) | -1071 |
| Mtrfr | Intron (ENSMUST00000111477.1/72650, intron 1 of 1) | 8940 |
| Plekha4 | Promoter (<=1kb) | 796 |
| Rtl6 | Promoter (<=1kb) | 106 |
| Rere | Distal Intergenic | -14542 |
| 4930521O11Rik | Intron (ENSMUST00000181509.1/74715, intron 3 of 3) | 15748 |
| Gtse1 | Promoter (<=1kb) | 3 |
| Mroh4 | Intron (ENSMUST00000042035.15/107831, intron 26 of 30) | 51984 |
| Serpinh1 | Intron (ENSMUST00000068973.10/17760, intron 1 of 3) | 41420 |
| Mir6546 | Promoter (<=1kb) | 121 |
| Rin3 | Intron (ENSMUST00000056950.13/217835, intron 1 of 9) | 16258 |
| Cenpm | Promoter (2-3kb) | -2407 |
| Wasf2 | Distal Intergenic | -49513 |
| 2010106C02Rik | Intron (ENSMUST00000097275.8/18754, intron 2 of 14) | -184528 |
| Mir6546 | Promoter (<=1kb) | 232 |
| Gpr153 | Intron (ENSMUST00000075363.9/70025, intron 6 of 8) | -31701 |
| Iffo1 | Exon (ENSMUST00000139071.7/320678, exon 3 of 7) | 5915 |
| Pigr | Intron (ENSMUST00000125056.7/ENSMUST00000125056.7, intron 1 of 3) | -5531 |
| Hdc | Intron (ENSMUST00000028838.4/15186, intron 10 of 11) | 21762 |
| Mir7044 | Intron (ENSMUST00000164960.3/70727, intron 1 of 12) | -16444 |
| Plekhg6 | Distal Intergenic | -41662 |
| Fam102b | Distal Intergenic | -5179 |
| Col8a2 | Promoter (<=1kb) | 0 |
| C1ql2 | Promoter (<=1kb) | 0 |
| Arf1 | Promoter (2-3kb) | -2120 |
| Tspan9 | Intron (ENSMUST00000112173.7/109246, intron 1 of 7) | 87204 |
| Rrm2 | Distal Intergenic | 16248 |
| Il27ra | Promoter (<=1kb) | 107 |
| Trim13 | Promoter (1-2kb) | 1128 |
| Mir675 | Distal Intergenic | 6301 |
| Shisal1 | Exon (ENSMUST00000080751.8/72301, exon 3 of 5) | 29899 |
| Trim13 | Promoter (<=1kb) | 931 |
| Mrln | Intron (ENSMUST00000147545.7/76551, intron 7 of 8) | -20824 |
| Cobll1 | Promoter (<=1kb) | -119 |
| Mir7077 | Intron (ENSMUST00000166750.8/74440, intron 1 of 20) | -96345 |
| B2m | Promoter (2-3kb) | 2542 |
| Acaa1b | Promoter (<=1kb) | 142 |
| Grin2c | Exon (ENSMUST00000003351.12/14813, exon 13 of 13) | 16818 |
| Rrad | Promoter (1-2kb) | 1664 |
| Ly6e | Distal Intergenic | -27833 |

| | | |
|---|---|---|
| Aprt | Intron (ENSMUST00000127664.1/108148, intron 1 of 15) | -16640 |
| Gm29678 | Distal Intergenic | 126523 |
| Mir704 | Distal Intergenic | 61805 |
| Zbtb16 | Intron (ENSMUST00000093852.4/235320, intron 2 of 6) | 48576 |
| Aqp11 | Promoter (1-2kb) | 1964 |
| Slc24a4 | Promoter (<=1kb) | 421 |
| Slc10a4 | Intron (ENSMUST00000101127.11/72313, intron 18 of 62) | 98511 |
| Sync | Intron (ENSMUST00000102599.3/68828, intron 2 of 4) | 11718 |
| Prok2 | Distal Intergenic | -235469 |
| Trim13 | Promoter (1-2kb) | 1160 |
| Gm21859 | Exon (ENSMUST00000178424.1/545180, exon 4 of 4) | 25434 |
| Gprin1 | Promoter (2-3kb) | -2936 |
| NA | Exon (ENSMUST00000167717.3/547127, exon 7 of 17) | 16370 |
| Fam98b | Distal Intergenic | -18164 |
| Cog1 | Intron (ENSMUST00000125890.7/69806, intron 1 of 5) | -8123 |
| Xpo5 | Exon (ENSMUST00000087031.6/72322, exon 4 of 32) | 5110 |
| Mxd4 | Exon (ENSMUST00000130435.4/231125, exon 1 of 5) | -44952 |
| 4930521O11Rik | Intron (ENSMUST00000181509.1/74715, intron 3 of 3) | 16137 |
| NA | Intron (ENSMUST00000188746.6/547127, intron 2 of 15) | 6750 |
| Chrna3 | Promoter (2-3kb) | 2387 |
| Sox1ot | Distal Intergenic | -39095 |
| 4930404H11Rik | Intron (ENSMUST00000222756.1/73815, intron 5 of 5) | 56114 |
| 4930521O11Rik | Intron (ENSMUST00000181509.1/74715, intron 3 of 3) | 16162 |
| Mat2a | Distal Intergenic | -38302 |
| D6Ertd527e | Intron (ENSMUST00000042025.11/69538, intron 17 of 17) | 49004 |
| Fcgr3 | Promoter (1-2kb) | -1111 |
| Copg1 | Exon (ENSMUST00000113607.9/54161, exon 9 of 24) | 6037 |
| Mir1231 | Intron (ENSMUST00000040599.14/215690, intron 3 of 29) | -55790 |
| Tmem74b | Promoter (<=1kb) | 730 |
| 4833438C02Rik | Exon (ENSMUST00000237330.1/18034, exon 13 of 22) | -3460 |
| Zfyve21 | Intron (ENSMUST00000021714.8/68520, intron 1 of 6) | 5961 |
| 4930521O11Rik | Intron (ENSMUST00000181509.1/74715, intron 3 of 3) | 15934 |
| Shisa5 | 3' UTR | -10713 |
| Nfkbia | Distal Intergenic | 49263 |
| Trh | Distal Intergenic | -25383 |
| Tnrc18 | Exon (ENSMUST00000198181.1/231861, exon 3 of 3) | 24650 |
| Grik4 | Promoter (2-3kb) | -2267 |
| Dynlt1b | Intron (ENSMUST00000180035.7/547127, intron 8 of 8) | 10016 |
| Fcgr3 | Promoter (1-2kb) | -1135 |
| Scarb1 | Intron (ENSMUST00000086075.12/20778, intron 1 of 12) | 10639 |
| Ucp3 | Intron (ENSMUST00000098259.10/277939, intron 31 of 31) | -6111 |
| Slc25a33 | Distal Intergenic | -36183 |
| Vwa7 | Exon (ENSMUST00000087315.13/22321, exon 12 of 30) | -4977 |

| | | |
|---|---|---|
| Tmem161a | 3' UTR | -5266 |
| Ncf2 | Distal Intergenic | -3931 |
| Fam20c | Intron (ENSMUST00000026972.7/80752, intron 3 of 9) | 20076 |
| 2210406O10Rik | Intron (ENSMUST00000149687.1/76710, intron 2 of 2) | 14599 |
| Atp6v0e2 | Distal Intergenic | 5492 |
| Ror2 | Intron (ENSMUST00000130235.7/26564, intron 1 of 8) | 66254 |
| Pla2g5 | Intron (ENSMUST00000102513.7/18784, intron 1 of 5) | 3601 |
| Kcnn3 | Distal Intergenic | -27139 |
| Mesp1 | Intron (ENSMUST00000035622.7/626359, intron 17 of 17) | 11255 |
| Anxa4 | Distal Intergenic | -13176 |
| Cilp | Promoter (2-3kb) | 2497 |
| Wdr11 | Distal Intergenic | 57201 |
| Gm11468 | Distal Intergenic | 86194 |
| Slc2a1 | Intron (ENSMUST00000030398.9/20525, intron 1 of 9) | 8914 |
| 2700046A07Rik | Distal Intergenic | -15752 |
| Tnni1 | Intron (ENSMUST00000132795.7/21952, intron 2 of 6) | 5158 |
| 2010106C02Rik | Intron (ENSMUST00000097275.8/18754, intron 2 of 14) | -131898 |
| Nipsnap1 | Distal Intergenic | -11626 |
| Tmbim1 | Exon (ENSMUST00000087225.5/56695, exon 1 of 9) | -27051 |
| Syt17 | Distal Intergenic | -11866 |
| Atg7 | Exon (ENSMUST00000203217.1/ENSMUST00000203217.1, exon 1 of 1) | 146563 |
| Amigo3 | Promoter (<=1kb) | 0 |
| Lmna | Exon (ENSMUST00000029699.12/16905, exon 7 of 12) | 18489 |
| Camta1 | Intron (ENSMUST00000169423.8/100072, intron 3 of 24) | 109117 |
| Prps1 | Intron (ENSMUST00000208130.1/245643, intron 14 of 14) | -26154 |
| F7 | Intron (ENSMUST00000095456.9/17207, intron 5 of 29) | -36588 |
| Tacc2 | Intron (ENSMUST00000207918.1/57752, intron 1 of 6) | 14308 |
| Foxp1 | Intron (ENSMUST00000113326.8/108655, intron 3 of 17) | 537458 |
| Zfp423 | Intron (ENSMUST00000109655.8/94187, intron 2 of 7) | 65677 |
| Ehd2 | Intron (ENSMUST00000098799.4/259300, intron 3 of 5) | 6427 |
| Wsb1 | Distal Intergenic | 30328 |
| Gadd45g | Distal Intergenic | 111778 |
| Sin3b | Intron (ENSMUST00000109950.4/20467, intron 5 of 6) | 10869 |
| Trim13 | Promoter (1-2kb) | 1217 |
| NA | Intron (ENSMUST00000223481.1/ENSMUST00000223481.1, intron 1 of 2) | 47513 |
| Ctsd | Distal Intergenic | -11346 |
| Nr5a1 | 5' UTR | 3599 |
| Gm3636 | Intron (ENSMUST00000181787.1/ENSMUST00000181787.1, intron 2 of 2) | -15883 |
| Ass1 | Intron (ENSMUST00000102840.4/11898, intron 5 of 15) | 13013 |
| Gm29678 | Distal Intergenic | -25916 |
| Scn4a | Promoter (1-2kb) | -1731 |
| Srl | Intron (ENSMUST00000005862.8/83383, intron 6 of 6) | -4911 |
| Gm1968 | Intron (ENSMUST00000181815.1/ENSMUST00000181815.1, intron 1 of 1) | 159142 |

| | | |
|---|---|---|
| Sipa1l2 | Distal Intergenic | 158914 |
| Ldlrap1 | Distal Intergenic | -7482 |
| Rpa1 | Distal Intergenic | 74743 |
| U2af1 | Promoter (2-3kb) | 2477 |
| Fcgr3 | Promoter (<=1kb) | -863 |
| Gadd45g | Distal Intergenic | 37340 |
| Pstpip1 | Intron (ENSMUST00000059206.7/19200, intron 3 of 14) | 28604 |
| Htr1d | Intron (ENSMUST00000088677.4/15552, intron 1 of 1) | 15659 |
| Parvaos | Distal Intergenic | -58000 |
| Clec2l | Promoter (<=1kb) | 103 |
| Amz1 | Intron (ENSMUST00000060918.10/231842, intron 4 of 7) | 21725 |
| Bcl2l11 | Intron (ENSMUST00000019281.13/12125, intron 3 of 4) | 27074 |
| Fbxl14 | Intron (ENSMUST00000178696.7/22419, intron 1 of 4) | -28416 |
| Tacc2 | Exon (ENSMUST00000207999.1/57752, exon 1 of 5) | 18911 |
| Slc25a33 | Distal Intergenic | -36173 |
| 9530026P05Rik | Promoter (1-2kb) | 1094 |
| Mapk9 | Intron (ENSMUST00000020629.4/14584, intron 16 of 18) | -14582 |
| Mir1249 | Intron (ENSMUST00000047144.12/223739, intron 1 of 9) | -10457 |
| Tacc2 | Exon (ENSMUST00000207999.1/57752, exon 1 of 5) | 18350 |
| Ddit4 | Distal Intergenic | -11929 |
| Ndufv3 | Distal Intergenic | 21752 |
| Glrp1 | Promoter (2-3kb) | 2446 |
| Kcnj4 | 5' UTR | 19272 |
| Trim62 | Distal Intergenic | 28578 |
| Spats1 | Promoter (<=1kb) | 174 |
| Pole4 | Exon (ENSMUST00000000642.10/15277, exon 9 of 18) | -33001 |
| Sod1 | Promoter (2-3kb) | -2666 |
| Gm1968 | Distal Intergenic | 100189 |
| Gpd1l | Distal Intergenic | 43144 |
| Epor | Promoter (<=1kb) | 801 |
| Pllp | Distal Intergenic | -8130 |
| Miga2 | Promoter (<=1kb) | -278 |
| Zfpm1 | Intron (ENSMUST00000127664.1/108148, intron 1 of 15) | 12331 |
| Cpsf3 | Intron (ENSMUST00000174530.1/211914, intron 6 of 9) | -118081 |
| Rab11a | Intron (ENSMUST00000093829.8/214058, intron 4 of 22) | 146765 |
| Rbfox2 | Distal Intergenic | -43653 |
| Map4k4 | Promoter (1-2kb) | 1893 |
| Nfil3 | Distal Intergenic | -115608 |
| Fmnl3 | Intron (ENSMUST00000081224.13/22379, intron 1 of 24) | 18432 |
| Fbxo47 | Exon (ENSMUST00000017561.14/72324, exon 11 of 14) | -48166 |
| Spats2 | Promoter (<=1kb) | 928 |
| Etv6 | Intron (ENSMUST00000081028.12/14011, intron 1 of 7) | 25155 |
| Fbln1 | Intron (ENSMUST00000057410.13/14114, intron 14 of 16) | 39844 |

| | | |
|---|---|---|
| Mroh5 | Distal Intergenic | -335497 |
| Mcat | 3' UTR | -11243 |
| Pygb | Distal Intergenic | -14008 |
| Epha8 | Promoter (<=1kb) | -197 |
| Padi2 | Promoter (<=1kb) | -180 |
| Mreg | Distal Intergenic | -24385 |
| Gm1968 | Distal Intergenic | 101176 |
| Vgll4 | Intron (ENSMUST00000147639.7/232334, intron 2 of 5) | 42720 |
| Cbfa2t3 | Intron (ENSMUST00000127664.1/108148, intron 1 of 15) | 33034 |
| Tacc2 | Intron (ENSMUST00000207918.1/57752, intron 1 of 6) | 18987 |
| Trpv4 | Promoter (<=1kb) | 8 |
| Lipg | Distal Intergenic | 53674 |
| 2810414N06Rik | 3' UTR | 33339 |
| Mlph | Intron (ENSMUST00000027528.6/171531, intron 1 of 15) | 3180 |
| Snrpd3 | Intron (ENSMUST00000145890.7/68778, intron 5 of 8) | -15482 |
| Pknox2 | Intron (ENSMUST00000177218.7/208076, intron 1 of 10) | 115737 |
| Cdk18 | Distal Intergenic | -4575 |
| Htatip2 | Promoter (<=1kb) | 71 |
| Rgs6 | Intron (ENSMUST00000166429.8/217692, intron 16 of 22) | -160476 |
| Catip | Intron (ENSMUST00000027370.12/56695, intron 2 of 9) | -20297 |
| 4930549C15Rik | Intron (ENSMUST00000020971.13/108089, intron 6 of 9) | -24404 |
| Rnf44 | Distal Intergenic | 17770 |
| 4930413E15Rik | Distal Intergenic | -119239 |
| Gm14317 | Distal Intergenic | -23876 |
| Slc25a33 | Distal Intergenic | -36077 |
| Cabp1 | Intron (ENSMUST00000112113.7/29867, intron 1 of 5) | 6868 |
| Mir21a | Distal Intergenic | 15138 |
| Cdhr3 | Intron (ENSMUST00000095774.2/68764, intron 18 of 19) | 56397 |
| Sema4b | Distal Intergenic | -12575 |
| Fchsd2 | Intron (ENSMUST00000208439.1/207278, intron 1 of 6) | 7959 |
| Eya2 | Intron (ENSMUST00000088132.12/14049, intron 1 of 15) | 33271 |
| 2900040C04Rik | Distal Intergenic | -28144 |
| Gm13420 | Intron (ENSMUST00000095087.10/107746, intron 5 of 25) | 60116 |
| NA | Promoter (2-3kb) | -2234 |
| Mir704 | Distal Intergenic | 58004 |
| Srl | Intron (ENSMUST00000005862.8/83383, intron 6 of 6) | -4776 |
| Cpt2 | Promoter (<=1kb) | 100 |
| 2310043O21Rik | Intron (ENSMUST00000185323.2/69679, intron 2 of 2) | 24517 |
| Phf19 | Exon (ENSMUST00000028232.6/74016, exon 6 of 15) | 8548 |
| Pebp1 | 3' UTR | 4707 |
| Map2k3os | Intron (ENSMUST00000123544.1/24082, intron 1 of 1) | 3504 |
| Nfkb1 | Intron (ENSMUST00000029812.13/18033, intron 6 of 24) | 57036 |
| 8430430B14Rik | Distal Intergenic | -34167 |

| | | |
|---|---|---|
| Mex3a | Exon (ENSMUST00000193595.1/ENSMUST00000193595.1, exon 1 of 1) | -5840 |
| Gm10578 | Exon (ENSMUST00000081510.3/17314, exon 3 of 5) | -148273 |
| Rab15 | Intron (ENSMUST00000125842.7/110606, intron 5 of 12) | -60947 |
| R3hcc1l | Promoter (2-3kb) | -2811 |
| 1700016A09Rik | Intron (ENSMUST00000127664.1/108148, intron 1 of 15) | 9493 |
| Tbx1 | 3' UTR | 42431 |
| Errfi1 | Distal Intergenic | -36253 |
| Pnpla1 | Intron (ENSMUST00000056866.7/433091, intron 1 of 9) | 16558 |
| Or6s1 | Promoter (2-3kb) | 2131 |
| Cdhr3 | Intron (ENSMUST00000095774.2/68764, intron 3 of 19) | 12046 |
| Arntl | Promoter (<=1kb) | 714 |
| Fads1 | Distal Intergenic | -39599 |
| Atg7 | Distal Intergenic | -84848 |
| D430041D05Rik | Promoter (1-2kb) | -1653 |
| Pdgfra | 5' UTR | 3852 |
| Mroh5 | Promoter (<=1kb) | 0 |
| Atg7 | Distal Intergenic | -84599 |
| Mxd4 | Intron (ENSMUST00000094868.9/231125, intron 8 of 12) | -13467 |
| Ndufa6 | Distal Intergenic | -8624 |
| Klrb1c | Intron (ENSMUST00000204394.2/17059, intron 3 of 5) | 23138 |
| Atg7 | Distal Intergenic | -84876 |
| Anp32a | Intron (ENSMUST00000048043.11/235431, intron 3 of 11) | 96372 |
| 1700020M21Rik | Distal Intergenic | 110608 |
| Abcg1 | Distal Intergenic | -39390 |
| Lmod3 | Intron (ENSMUST00000113355.8/232288, intron 8 of 23) | -93887 |
| Baiap2 | Distal Intergenic | -10452 |
| Psd2 | Distal Intergenic | -24873 |
| Tacc2 | Intron (ENSMUST00000207918.1/57752, intron 1 of 6) | 19014 |
| Plekhg6 | Distal Intergenic | -45835 |
| Gm5820 | Intron (ENSMUST00000148850.7/71302, intron 1 of 5) | -11131 |
| Plekhg6 | Distal Intergenic | -45999 |
| Sec13 | Intron (ENSMUST00000101045.9/11941, intron 3 of 21) | -102790 |
| Slc45a1 | Promoter (1-2kb) | -1036 |
| Mir7074 | Promoter (1-2kb) | 1946 |
| Krt78 | Distal Intergenic | -12512 |
| Scamp1 | 5' UTR | 111614 |
| Prr5 | Distal Intergenic | -29715 |
| Rcc2 | Distal Intergenic | -7420 |
| Zfp648 | Distal Intergenic | -21020 |
| Mturn | Intron (ENSMUST00000190641.6/68235, intron 2 of 2) | 17114 |
| Morrbid | Intron (ENSMUST00000225087.1/100043424, intron 1 of 6) | 97913 |
| C2cd4a | Promoter (<=1kb) | 0 |
| Pigr | Exon (ENSMUST00000122968.1/ENSMUST00000122968.1, exon 1 of 3) | -6081 |

| | | |
|---|---|---|
| Mir6546 | Promoter (<=1kb) | 386 |
| Rmi2 | Exon (ENSMUST00000233200.1/545180, exon 28 of 46) | 46822 |
| Pan3 | Promoter (1-2kb) | 1174 |
| Pim3 | 3' UTR | 28706 |
| Rab35 | Intron (ENSMUST00000031492.14/77407, intron 1 of 5) | 5167 |
| Rexo4 | Promoter (<=1kb) | -652 |
| Bmpr1b | Distal Intergenic | -57112 |
| Cc2d1a | Intron (ENSMUST00000117424.8/212139, intron 5 of 27) | 4918 |
| Eif3g | Promoter (<=1kb) | 94 |
| Lrp8 | Promoter (1-2kb) | 1764 |
| NA | Exon (ENSMUST00000186821.1/ENSMUST00000186821.1, exon 2 of 2) | 42040 |
| Il31 | Intron (ENSMUST00000196809.4/381741, intron 4 of 11) | -8962 |
| Gm20556 | Exon (ENSMUST00000167499.7/73167, exon 7 of 7) | -38495 |
| Zfp648 | Exon (ENSMUST00000190485.1/ENSMUST00000190485.1, exon 1 of 4) | -82945 |
| Polr3h | Promoter (<=1kb) | -339 |
| Foxk1 | Distal Intergenic | -174711 |
| Cbs | Intron (ENSMUST00000236853.1/12411, intron 7 of 16) | 13634 |
| Hectd1 | Exon (ENSMUST00000040583.6/320487, exon 34 of 36) | -51443 |
| Ccdc12 | Intron (ENSMUST00000019803.8/72654, intron 1 of 6) | 31524 |
| Trp53cor1 | Exon (ENSMUST00000233713.1/100504267, exon 1 of 2) | 6803 |
| Pknox2 | Intron (ENSMUST00000177218.7/208076, intron 1 of 10) | 99508 |
| Sgk2 | Distal Intergenic | -10763 |
| Tmem115 | Exon (ENSMUST00000010210.12/56808, exon 14 of 38) | -19126 |
| Tspan11 | Intron (ENSMUST00000112173.7/109246, intron 2 of 7) | 89108 |
| Paqr3 | Promoter (2-3kb) | 2606 |
| Gsdme | Distal Intergenic | -16584 |
| Tacc2 | Intron (ENSMUST00000207918.1/57752, intron 1 of 6) | 14798 |
| Acvr1b | Distal Intergenic | -8165 |
| Gimap1 | Exon (ENSMUST00000054368.6/16205, exon 3 of 3) | 3864 |
| Lhfpl2 | Intron (ENSMUST00000054274.7/218454, intron 1 of 3) | 16557 |
| Mark2 | Intron (ENSMUST00000032557.14/13728, intron 1 of 17) | 30138 |
| Zfp648 | Distal Intergenic | -21050 |
| Gm3636 | Intron (ENSMUST00000181787.1/ENSMUST00000181787.1, intron 2 of 2) | -16423 |
| Ppp2r5c | Exon (ENSMUST00000221791.1/ENSMUST00000221791.1, exon 3 of 3) | 32455 |
| Tacc2 | Intron (ENSMUST00000207918.1/57752, intron 1 of 6) | 14460 |
| Tead4 | Promoter (<=1kb) | -780 |
| Tcl1b2 | Distal Intergenic | -52276 |
| Ackr3 | Promoter (2-3kb) | 2883 |
| Gzf1 | Promoter (<=1kb) | 603 |
| Fcgr3 | Promoter (1-2kb) | -1160 |
| Mbtps1 | Intron (ENSMUST00000212601.1/56453, intron 1 of 21) | 9650 |
| Mir3108 | Intron (ENSMUST00000220518.1/11906, intron 7 of 15) | -171913 |
| Etv5 | Intron (ENSMUST00000079601.12/104156, intron 5 of 12) | 8296 |

| | | |
|---|---|---|
| Spink2 | Distal Intergenic | 45946 |
| Gm28729 | Intron (ENSMUST00000190704.2/102635744, intron 6 of 10) | 17388 |
| Lgals9 | Promoter (<=1kb) | 0 |
| Spring1 | 3' UTR | 17261 |
| Epb41 | Distal Intergenic | -4588 |
| 4930521O11Rik | Intron (ENSMUST00000181509.1/74715, intron 3 of 3) | 17030 |
| Stk11ip | Exon (ENSMUST00000027414.15/71728, exon 18 of 25) | 10916 |
| Nes | Distal Intergenic | -10102 |
| Pomgnt2 | Distal Intergenic | -17344 |
| Nxpe4 | Distal Intergenic | -298006 |
| Mlycd | Promoter (<=1kb) | -511 |
| Foxi1 | Promoter (<=1kb) | 505 |
| Anapc7 | Exon (ENSMUST00000122010.7/56317, exon 9 of 12) | 17816 |
| Mreg | Distal Intergenic | -23923 |
| Cib4 | Intron (ENSMUST00000065486.2/73259, intron 3 of 6) | 24153 |
| Atp2c2 | Exon (ENSMUST00000212203.1/ENSMUST00000212203.1, exon 1 of 1) | 15346 |
| Mir704 | Distal Intergenic | 61831 |
| Cidea | Intron (ENSMUST00000144078.7/338360, intron 1 of 3) | -18690 |
| Acvr1b | Distal Intergenic | -8102 |
| D730045A05Rik | Promoter (2-3kb) | -2725 |
| Septin3 | 5' UTR | 6346 |
| Atg7 | Distal Intergenic | -84138 |
| Clip4 | 3' UTR | 100911 |
| Rpl11 | Promoter (<=1kb) | -264 |
| Noct | Intron (ENSMUST00000189687.1/ENSMUST00000189687.1, intron 1 of 2) | -150400 |
| Mis18a | Distal Intergenic | 14496 |
| Anapc4 | Distal Intergenic | 70765 |
| Spi1 | Intron (ENSMUST00000169852.1/20375, intron 1 of 3) | 11239 |
| Snx29 | Intron (ENSMUST00000180792.7/74478, intron 13 of 20) | 133477 |
| Trio | Intron (ENSMUST00000090247.6/223435, intron 1 of 56) | 46988 |
| Hsd17b14 | Intron (ENSMUST00000107752.11/66065, intron 5 of 8) | 6178 |
| Tspear | Promoter (<=1kb) | 505 |
| Scarb1 | Intron (ENSMUST00000198586.1/ENSMUST00000198586.1, intron 1 of 3) | 77885 |
| Kif23 | Promoter (<=1kb) | 193 |
| Agap1 | Intron (ENSMUST00000027521.14/347722, intron 1 of 17) | 41583 |
| Spata13 | Intron (ENSMUST00000160973.7/219140, intron 2 of 12) | 59104 |
| 4931429L15Rik | 5' UTR | 5185 |
| Zfp827 | Promoter (<=1kb) | -216 |
| A2ml1 | Distal Intergenic | -13234 |
| Hykk | Promoter (<=1kb) | 0 |
| Myod1 | Distal Intergenic | 4730 |
| Vps4a | Exon (ENSMUST00000118457.1/ENSMUST00000118457.1, exon 1 of 1) | 6803 |
| NA | Intron (ENSMUST00000183659.7/78286, intron 2 of 39) | -137350 |

| | | |
|---|---|---|
| Zcchc24 | Promoter (<=1kb) | 0 |
| NA | Promoter (<=1kb) | -559 |
| Galnt16 | Promoter (<=1kb) | 95 |
| C030007H22Rik | Exon (ENSMUST00000190308.1/ENSMUST00000190308.1, exon 3 of 3) | 103153 |
| Alox12b | Intron (ENSMUST00000021268.8/23801, intron 14 of 14) | -9047 |
| Tbc1d22bos | Intron (ENSMUST00000048677.8/381085, intron 12 of 12) | -7288 |
| Mrps24 | Promoter (<=1kb) | -844 |
| Abcc4 | Distal Intergenic | -10115 |
| Get4 | Exon (ENSMUST00000058716.13/77053, exon 18 of 23) | -10491 |
| Urb1 | Promoter (<=1kb) | 25 |
| Zbtb49 | Intron (ENSMUST00000124939.7/75079, intron 4 of 5) | 22953 |
| Abcb9 | Intron (ENSMUST00000031354.10/56325, intron 1 of 11) | 3156 |
| Egr3 | Intron (ENSMUST00000022678.4/73523, intron 4 of 6) | -111660 |
| Sox1ot | Distal Intergenic | -60294 |
| Pik3ap1 | Intron (ENSMUST00000059672.8/83490, intron 2 of 16) | 36650 |
| Ubald2 | Distal Intergenic | -7656 |
| Mir1966 | Intron (ENSMUST00000194091.5/11604, intron 1 of 5) | 6874 |
| Fry | Intron (ENSMUST00000200960.3/320365, intron 2 of 4) | 147333 |
| Syt16 | Intron (ENSMUST00000221220.1/238266, intron 1 of 6) | 16144 |
| Sun2 | Intron (ENSMUST00000162392.7/223697, intron 7 of 16) | 7263 |
| Gadd45g | Distal Intergenic | 19576 |
| Plekha4 | Exon (ENSMUST00000211227.1/69217, exon 8 of 16) | 11850 |
| Adam11 | Exon (ENSMUST00000068150.6/11488, exon 15 of 26) | 12815 |
| Abcg1 | Intron (ENSMUST00000236391.1/11307, intron 2 of 5) | 26263 |
| Atg7 | Distal Intergenic | -84883 |
| Mir704 | Distal Intergenic | 58038 |
| Myo7a | Promoter (1-2kb) | -1007 |
| Arhgap23 | Intron (ENSMUST00000121799.7/58996, intron 6 of 23) | 34380 |
| Atg7 | Distal Intergenic | -84374 |
| Mir1981 | Exon (ENSMUST00000161359.7/67247, exon 1 of 5) | -4806 |
| Potefam3b | Distal Intergenic | -66694 |
| Adra2a | Distal Intergenic | 36515 |
| Smtnl1 | 3' UTR | 11197 |
| Atg7 | Distal Intergenic | -84149 |
| Srxn1 | Exon (ENSMUST00000064061.3/545474, exon 2 of 2) | -11773 |
| Tacc2 | Intron (ENSMUST00000207918.1/57752, intron 1 of 6) | 14772 |
| Slc29a4 | 5' UTR | 9398 |
| Adssl1 | Promoter (<=1kb) | 50 |
| 1700109G15Rik | Distal Intergenic | -29547 |
| Acvr1b | Distal Intergenic | -7967 |
| Cd59a | Promoter (<=1kb) | 0 |
| C530044C16Rik | Distal Intergenic | 235874 |
| Mir101b | Distal Intergenic | 12637 |

| | | |
|---|---|---|
| Ksr1 | Intron (ENSMUST00000018478.10/16706, intron 1 of 18) | 14455 |
| Galr2 | Intron (ENSMUST00000021147.13/53413, intron 18 of 19) | 8805 |
| Atad2b | Exon (ENSMUST00000020958.8/208439, exon 12 of 14) | 166871 |
| Coro2a | Exon (ENSMUST00000127304.1/107684, exon 2 of 2) | 26742 |
| Scrn1 | 3' UTR | 63309 |
| Pald1 | Exon (ENSMUST00000219025.1/ENSMUST00000219025.1, exon 1 of 1) | 23396 |
| Acsl5 | Intron (ENSMUST00000225551.1/433256, intron 1 of 4) | 16456 |
| Fgr | Promoter (<=1kb) | 184 |
| 4930442L01Rik | Promoter (<=1kb) | -326 |
| Sytl3 | Intron (ENSMUST00000097430.9/83672, intron 1 of 15) | 6373 |
| Ddr1 | Exon (ENSMUST00000173840.1/ENSMUST00000173840.1, exon 4 of 4) | -33219 |
| Dnmt3aos | Intron (ENSMUST00000172719.7/13435, intron 4 of 5) | -6422 |
| Potefam3b | Distal Intergenic | -65980 |
| Mir330 | Promoter (1-2kb) | -1808 |
| Xrcc6 | Intron (ENSMUST00000165777.7/14375, intron 1 of 4) | 3211 |
| Gm29678 | Distal Intergenic | 129250 |
| Mir3083 | Promoter (<=1kb) | -401 |
| Tbc1d9b | Intron (ENSMUST00000093138.12/76795, intron 5 of 20) | 16281 |
| Txnip | Distal Intergenic | -5731 |
| Srgap3 | Intron (ENSMUST00000088373.10/259302, intron 1 of 21) | 94438 |
| Slc25a33 | Distal Intergenic | -80052 |
| Rph3a | Distal Intergenic | -16524 |
| Ccbe1 | 5' UTR | 10790 |
| Cnot11 | Promoter (2-3kb) | 2930 |
| Glce | Distal Intergenic | -58005 |
| Clic4 | Distal Intergenic | -14014 |
| Smug1 | Distal Intergenic | -16698 |

Table S8: AR Cluster 3 Genes Top500

| geneSymbol | annotation | distanceToTSS |
|---|---|---|
| Setd5 | Promoter (<=1kb) | 0 |
| Apip | Distal Intergenic | 70666 |
| Setd5 | Promoter (<=1kb) | 41 |
| Mfsd11 | Promoter (<=1kb) | 0 |
| Slc38a2 | Promoter (<=1kb) | 224 |
| Apip | Distal Intergenic | 70701 |
| Apip | Distal Intergenic | 70525 |
| Asxl1 | Promoter (<=1kb) | 0 |
| Slc38a2 | Promoter (<=1kb) | 218 |
| Arfip1 | Promoter (<=1kb) | 73 |
| Mex3c | Promoter (<=1kb) | 0 |
| Setd5 | Promoter (<=1kb) | 56 |
| Mir5127 | Intron (ENSMUST00000236629.1/ENSMUST00000236629.1, intron 2 of 2) | 480194 |
| Zzz3 | Promoter (<=1kb) | 595 |
| NA | Promoter (<=1kb) | 0 |
| Atxn2 | Promoter (<=1kb) | 524 |
| NA | Promoter (<=1kb) | -33 |
| Gtf2a1 | Promoter (1-2kb) | 1253 |
| Atxn2 | Promoter (<=1kb) | 581 |
| Rap2a | Promoter (<=1kb) | -275 |
| Zzz3 | Promoter (<=1kb) | 558 |
| Magi1 | Promoter (<=1kb) | -116 |
| Mir5127 | Intron (ENSMUST00000236629.1/ENSMUST00000236629.1, intron 2 of 2) | 480483 |
| NA | Intron (ENSMUST00000085546.12/15289, intron 1 of 4) | 44742 |
| Creld2 | Distal Intergenic | 11864 |
| Hspa9 | Promoter (<=1kb) | -202 |
| Tpm1 | Distal Intergenic | 36553 |
| Lonrf1 | Promoter (<=1kb) | 539 |
| Hes1 | Distal Intergenic | 4784 |
| Mfsd11 | Promoter (<=1kb) | -13 |
| Tent4b | Promoter (<=1kb) | 204 |
| Grhl2 | Distal Intergenic | -40195 |
| 2310008N11Rik | Intron (ENSMUST00000210732.1/72316, intron 1 of 2) | 39888 |
| Eif5 | Promoter (<=1kb) | 557 |
| Hspa9 | Promoter (<=1kb) | -503 |
| Kcmf1 | Promoter (<=1kb) | 0 |
| Sh3glb1 | Promoter (<=1kb) | 300 |
| Rasl11b | Intron (ENSMUST00000072857.12/212986, intron 5 of 8) | 55138 |
| Grhl2 | Distal Intergenic | -39960 |
| Kcmf1 | Promoter (<=1kb) | 203 |
| Map3k1 | Promoter (<=1kb) | 460 |
| Mfsd11 | Promoter (<=1kb) | -180 |

| | | |
|---|---|---|
| Mex3c | Promoter (<=1kb) | -707 |
| Arl6ip4 | Intron (ENSMUST00000086123.10/19679, intron 2 of 23) | 32606 |
| Yy1 | Promoter (<=1kb) | -341 |
| Sbno1 | Promoter (<=1kb) | 235 |
| Rap2a | Promoter (<=1kb) | -483 |
| Klf9 | Intron (ENSMUST00000236056.1/ENSMUST00000236056.1, intron 1 of 1) | -83604 |
| Hectd1 | Promoter (<=1kb) | 0 |
| Hspa9 | Promoter (<=1kb) | -54 |
| Map3k1 | Promoter (<=1kb) | 110 |
| Rybp | Promoter (<=1kb) | 0 |
| Sbno1 | Promoter (<=1kb) | 238 |
| Fzd4 | Promoter (<=1kb) | 0 |
| Marcks | Promoter (1-2kb) | 1500 |
| Jarid2 | Promoter (<=1kb) | 426 |
| Hspa9 | Promoter (<=1kb) | -116 |
| Map3k1 | Promoter (<=1kb) | 165 |
| Rap2a | Promoter (<=1kb) | -513 |
| Mex3c | Promoter (<=1kb) | -400 |
| Tpm1 | Distal Intergenic | 36276 |
| Yy1 | Promoter (<=1kb) | -299 |
| Mir7651 | 5' UTR | 14084 |
| Mfsd11 | Promoter (<=1kb) | -231 |
| Slc45a1 | Distal Intergenic | -32943 |
| Rev1 | Promoter (<=1kb) | 0 |
| Slc45a1 | Distal Intergenic | -33330 |
| Rybp | Promoter (<=1kb) | 114 |
| Rybp | Promoter (<=1kb) | 96 |
| Acaa2 | Promoter (<=1kb) | 0 |
| Arl6ip4 | Intron (ENSMUST00000086123.10/19679, intron 2 of 23) | 32569 |
| Slc45a1 | Distal Intergenic | -32839 |
| Klf3 | Promoter (<=1kb) | 122 |
| Uvssa | Intron (ENSMUST00000200682.1/71101, intron 7 of 8) | 25608 |
| Fryl | Promoter (<=1kb) | 386 |
| Hnrnpdl | Promoter (<=1kb) | 691 |
| Uvssa | Intron (ENSMUST00000200682.1/71101, intron 7 of 8) | 25764 |
| Acaa2 | Promoter (<=1kb) | -104 |
| Tle1 | Promoter (<=1kb) | -449 |
| Yy1 | Promoter (<=1kb) | 0 |
| Uvssa | Intron (ENSMUST00000200682.1/71101, intron 7 of 8) | 25675 |
| Zzz3 | Promoter (<=1kb) | 323 |
| Rybp | Promoter (<=1kb) | 283 |
| Mir207 | Promoter (<=1kb) | -63 |
| Acaa2 | Promoter (<=1kb) | 136 |

| | | |
|---|---|---|
| Klf9 | Intron (ENSMUST00000236056.1/ENSMUST00000236056.1, intron 1 of 1) | -84048 |
| Hnrnpdl | Promoter (<=1kb) | 706 |
| Bcl2l1 | Intron (ENSMUST00000007803.11/12048, intron 2 of 2) | 11025 |
| Rasl11b | Intron (ENSMUST00000072857.12/212986, intron 5 of 8) | 54634 |
| Slc45a1 | Distal Intergenic | -33085 |
| Fxr2 | Promoter (<=1kb) | 0 |
| Sbno1 | Promoter (<=1kb) | 52 |
| Hspa9 | Promoter (<=1kb) | -601 |
| Slc45a1 | Distal Intergenic | -33252 |
| Maco1 | Promoter (<=1kb) | 85 |
| Mir7651 | 5' UTR | 13929 |
| Mir8112 | Promoter (<=1kb) | 0 |
| Slfnl1 | Exon (ENSMUST00000136801.7/29871, exon 1 of 5) | -14592 |
| Rybp | Promoter (<=1kb) | 291 |
| Cc2d1b | Promoter (<=1kb) | 0 |
| Klf3 | Promoter (<=1kb) | 0 |
| Cachd1 | Promoter (<=1kb) | 40 |
| Uvssa | Intron (ENSMUST00000200682.1/71101, intron 7 of 8) | 25937 |
| Dhx8 | Distal Intergenic | -31862 |
| Klf9 | Intron (ENSMUST00000236056.1/ENSMUST00000236056.1, intron 1 of 1) | -84434 |
| F3 | Promoter (<=1kb) | 0 |
| NA | 5' UTR | -112032 |
| Exosc8 | Promoter (<=1kb) | 234 |
| Dnah8 | 5' UTR | -48074 |
| Foxp1 | 5' UTR | 603996 |
| Fbxl17 | Promoter (<=1kb) | -420 |
| Klf9 | Intron (ENSMUST00000236056.1/ENSMUST00000236056.1, intron 1 of 1) | -84131 |
| Apbb2 | Promoter (<=1kb) | 425 |
| Uvssa | Intron (ENSMUST00000200682.1/71101, intron 7 of 8) | 26056 |
| Hspa8 | Promoter (<=1kb) | 0 |
| Cnbd2 | Promoter (<=1kb) | -69 |
| H4c1 | Promoter (<=1kb) | -20 |
| Bmp2k | Promoter (<=1kb) | -31 |
| Fbxo11 | Promoter (<=1kb) | 394 |
| Apbb2 | Promoter (<=1kb) | 138 |
| Apbb2 | Promoter (<=1kb) | 10 |
| Mbnl1 | Intron (ENSMUST00000195724.5/56758, intron 2 of 2) | 28919 |
| Erbin | Promoter (<=1kb) | -433 |
| Scamp1 | Promoter (<=1kb) | 0 |
| Rap2a | Promoter (<=1kb) | -547 |
| 9230116L04Rik | Promoter (<=1kb) | 0 |
| Phlpp1 | Promoter (<=1kb) | 0 |
| Apbb2 | Promoter (<=1kb) | 185 |

| | | |
|---|---|---|
| Apbb2 | Promoter (<=1kb) | 353 |
| Spen | Exon (ENSMUST00000049927.5/ENSMUST00000049927.5, exon 1 of 3) | -7703 |
| Apbb2 | Promoter (<=1kb) | 328 |
| Gm5122 | Exon (ENSMUST00000216170.1/ENSMUST00000216170.1, exon 1 of 7) | 15069 |
| Pdcd4 | 5' UTR | 10779 |
| Fbxl17 | Promoter (<=1kb) | -448 |
| Hes1 | Distal Intergenic | 5376 |
| Phlda1 | Promoter (<=1kb) | 13 |
| 4930519K11Rik | Intron (ENSMUST00000172099.8/16531, intron 1 of 27) | -209846 |
| Prr5l | Exon (ENSMUST00000127358.1/72446, exon 2 of 2) | 50585 |
| Eps8 | Distal Intergenic | 182838 |
| Srd5a3 | Promoter (<=1kb) | 115 |
| Cul4a | Promoter (<=1kb) | 191 |
| Srd5a3 | Promoter (<=1kb) | 0 |
| Dnah8 | 5' UTR | -48084 |
| Nr4a3 | Promoter (1-2kb) | -1525 |
| Srd5a3 | Promoter (<=1kb) | 1 |
| Lifr | Distal Intergenic | -50797 |
| Cap1 | Promoter (<=1kb) | 0 |
| Bmp2k | Promoter (<=1kb) | -54 |
| Rap2a | Promoter (<=1kb) | -975 |
| Dnah8 | Intron (ENSMUST00000168787.7/224671, intron 1 of 10) | -48240 |
| Hnrnpdl | Promoter (<=1kb) | 968 |
| Srd5a3 | Promoter (<=1kb) | 10 |
| Rpain | Promoter (<=1kb) | 3 |
| Skil | Promoter (<=1kb) | -281 |
| Phlda1 | Promoter (<=1kb) | 338 |
| Stt3b | Distal Intergenic | -22943 |
| Mbp | Intron (ENSMUST00000091789.10/17196, intron 1 of 3) | 17621 |
| Apoh | Promoter (<=1kb) | -2 |
| Mir7651 | 5' UTR | 13670 |
| Maco1 | Promoter (<=1kb) | 260 |
| Arhgap35 | Promoter (<=1kb) | 0 |
| Rasl11b | Intron (ENSMUST00000072857.12/212986, intron 5 of 8) | 54455 |
| Sumo2 | Promoter (<=1kb) | 0 |
| Phlda1 | Promoter (<=1kb) | 0 |
| Foxp1 | Intron (ENSMUST00000113326.8/108655, intron 4 of 17) | 603777 |
| Zbtb18 | Promoter (<=1kb) | 106 |
| Tmem135 | Promoter (<=1kb) | 0 |
| Zdhhc14 | Intron (ENSMUST00000089185.5/224454, intron 1 of 8) | 129722 |
| Zdhhc14 | Intron (ENSMUST00000089185.5/224454, intron 1 of 8) | 129539 |
| Ctr9 | Distal Intergenic | -4910 |
| Phlpp1 | Promoter (<=1kb) | 64 |

| | | |
|---|---|---|
| Hnrnpdl | Promoter (<=1kb) | 735 |
| Fgfbp1 | Exon (ENSMUST00000200632.1/ENSMUST00000200632.1, exon 1 of 1) | 33371 |
| Adipor1 | Promoter (<=1kb) | 28 |
| Brip1os | Promoter (<=1kb) | 24 |
| Aebp2 | Promoter (<=1kb) | 633 |
| Kpnb1 | Promoter (<=1kb) | -266 |
| Ctr9 | Distal Intergenic | -4889 |
| Scamp1 | Promoter (<=1kb) | 390 |
| Slc45a1 | Intron (ENSMUST00000105682.8/68703, intron 11 of 22) | 82805 |
| Skil | Promoter (<=1kb) | 0 |
| Sbno1 | Promoter (<=1kb) | 39 |
| Hnrnph1 | Promoter (<=1kb) | 491 |
| Bclaf1 | Promoter (<=1kb) | 500 |
| Zdhhc14 | Intron (ENSMUST00000089185.5/224454, intron 1 of 8) | 129873 |
| Exosc8 | Promoter (<=1kb) | 0 |
| Skil | Promoter (<=1kb) | -116 |
| Fgfbp1 | Exon (ENSMUST00000200632.1/ENSMUST00000200632.1, exon 1 of 1) | 33304 |
| Fgfbp1 | Exon (ENSMUST00000200632.1/ENSMUST00000200632.1, exon 1 of 1) | 33657 |
| Fth1 | Promoter (1-2kb) | 1689 |
| Gm5122 | Distal Intergenic | 14680 |
| H4c1 | Promoter (<=1kb) | 56 |
| Yy1 | Promoter (<=1kb) | 460 |
| Bcl2l14 | Exon (ENSMUST00000081028.12/14011, exon 7 of 8) | -133174 |
| Eif5 | Promoter (<=1kb) | 582 |
| Mir8112 | Promoter (<=1kb) | -17 |
| Fbxo11 | Promoter (<=1kb) | 0 |
| Plet1os | Promoter (<=1kb) | 0 |
| Ccne2 | Promoter (<=1kb) | -317 |
| Cep43 | Promoter (<=1kb) | 0 |
| Lifr | Distal Intergenic | -51101 |
| NA | Promoter (<=1kb) | 145 |
| Ppp2r5c | 5' UTR | 38299 |
| Klf3 | Promoter (<=1kb) | -103 |
| Dcaf11 | Promoter (<=1kb) | 236 |
| Tecpr2 | Promoter (<=1kb) | 61 |
| Tra2b | Promoter (<=1kb) | 423 |
| Sbno1 | Promoter (<=1kb) | 15 |
| Zfand3 | Promoter (<=1kb) | 0 |
| Rybp | Promoter (<=1kb) | 660 |
| Zfand3 | Promoter (<=1kb) | -78 |
| 2310008N11Rik | Intron (ENSMUST00000210732.1/72316, intron 1 of 2) | 39411 |
| Tulp4 | Promoter (<=1kb) | 570 |
| Skil | Promoter (<=1kb) | -429 |

| | | |
|---|---|---|
| Stat5b | Promoter (<=1kb) | 0 |
| Skil | Promoter (<=1kb) | 72 |
| Cdv3 | Promoter (<=1kb) | 494 |
| Ccdc92 | Promoter (<=1kb) | 0 |
| Spred1 | Promoter (<=1kb) | -67 |
| Cdv3 | Promoter (<=1kb) | 318 |
| Cldn14 | Distal Intergenic | -14941 |
| Tra2b | Promoter (<=1kb) | 28 |
| Cldn14 | Distal Intergenic | -15148 |
| Bmp2k | Promoter (<=1kb) | -197 |
| Dmxl2 | Promoter (<=1kb) | -207 |
| Tulp4 | Promoter (<=1kb) | 67 |
| Cldn14 | Distal Intergenic | -14961 |
| NA | Promoter (<=1kb) | 381 |
| Myl12b | Promoter (<=1kb) | -63 |
| Nudt3 | Promoter (<=1kb) | 0 |
| Insig1 | Promoter (<=1kb) | 0 |
| Tmem135 | Promoter (<=1kb) | 96 |
| Smco4 | Promoter (<=1kb) | 0 |
| Insig1 | Promoter (<=1kb) | 10 |
| Sumo2 | Promoter (<=1kb) | 178 |
| Zbtb18 | Promoter (<=1kb) | 126 |
| Azin1 | Promoter (<=1kb) | -44 |
| Cldn14 | Distal Intergenic | -14980 |
| Insig1 | Promoter (<=1kb) | 63 |
| Zcchc2 | Promoter (<=1kb) | 883 |
| Tecpr2 | Promoter (<=1kb) | 0 |
| 2610306M01Rik | Distal Intergenic | 16301 |
| Klf3 | Promoter (<=1kb) | -349 |
| Ubr5 | Promoter (<=1kb) | 329 |
| Maco1 | Promoter (<=1kb) | 305 |
| 1110020A21Rik | Promoter (<=1kb) | 0 |
| Gm5122 | Distal Intergenic | 14841 |
| Stag1 | Intron (ENSMUST00000124487.7/20842, intron 1 of 3) | 45134 |
| Rmi1 | Promoter (<=1kb) | 135 |
| Exosc8 | Promoter (<=1kb) | -1 |
| Mir8112 | Promoter (<=1kb) | -154 |
| Fabp2 | Intron (ENSMUST00000180557.5/74057, intron 5 of 6) | -20510 |
| Arhgef37 | Promoter (<=1kb) | 659 |
| Maco1 | Promoter (<=1kb) | 488 |
| Phlpp1 | Promoter (<=1kb) | 108 |
| Myom1 | Promoter (<=1kb) | 0 |
| Myo10 | Promoter (<=1kb) | 321 |

| | | |
|---|---|---|
| Ifnar1 | Promoter (<=1kb) | -42 |
| NA | Exon (ENSMUST00000136077.1/634731, exon 4 of 4) | 71909 |
| Gm5820 | Intron (ENSMUST00000148850.7/71302, intron 1 of 5) | -25245 |
| Cep43 | Promoter (<=1kb) | 299 |
| Usp3 | Promoter (<=1kb) | 73 |
| 3830406C13Rik | Intron (ENSMUST00000022264.12/19270, intron 5 of 29) | -224393 |
| Apbb2 | Promoter (<=1kb) | 0 |
| Cep43 | Promoter (<=1kb) | 137 |
| Cap1 | Promoter (<=1kb) | 304 |
| Fth1 | Promoter (1-2kb) | 1941 |
| Tbl1xr1 | Promoter (<=1kb) | -255 |
| Zfp541 | Distal Intergenic | -4251 |
| Nin | Promoter (1-2kb) | 1808 |
| Tra2b | Promoter (<=1kb) | 200 |
| Maco1 | Promoter (<=1kb) | 585 |
| Ddx31 | Promoter (<=1kb) | 0 |
| Spred2 | Distal Intergenic | -186683 |
| Bicral | Promoter (<=1kb) | 750 |
| Hmgxb3 | Promoter (<=1kb) | -114 |
| Lmo4 | Promoter (<=1kb) | 73 |
| Stt3b | Distal Intergenic | -22940 |
| Pank2 | Promoter (<=1kb) | 0 |
| Bmp2k | Promoter (<=1kb) | -260 |
| Ppm1a | 5' UTR | 3045 |
| Tsc22d2 | Promoter (<=1kb) | 194 |
| Cap1 | Promoter (<=1kb) | 166 |
| Bub1b | Distal Intergenic | -11076 |
| Spred1 | Promoter (<=1kb) | 0 |
| Cep43 | Promoter (<=1kb) | 158 |
| Ndufv2 | Promoter (<=1kb) | 0 |
| NA | Exon (ENSMUST00000136077.1/634731, exon 4 of 4) | 71770 |
| Rev1 | Promoter (<=1kb) | 597 |
| Cdv3 | Promoter (<=1kb) | 267 |
| Cep43 | Promoter (<=1kb) | 222 |
| Spock2 | Intron (ENSMUST00000050516.13/69090, intron 9 of 10) | -23942 |
| Fasn | Intron (ENSMUST00000056781.4/71276, intron 18 of 18) | -22706 |
| Dnajc11 | Distal Intergenic | -30695 |
| Zbtb18 | Promoter (<=1kb) | 291 |
| Gab1 | Intron (ENSMUST00000210676.1/14388, intron 1 of 10) | 27365 |
| Fbxo11 | Promoter (<=1kb) | 27 |
| Rev1 | Promoter (<=1kb) | 141 |
| H3f3b | 5' UTR | 3665 |
| Spen | Distal Intergenic | -7261 |

| | | |
|---|---|---|
| Rybp | Promoter (<=1kb) | 739 |
| A930033H14Rik | Promoter (<=1kb) | 0 |
| Snx1 | Promoter (2-3kb) | 2043 |
| Cnot6l | Promoter (1-2kb) | 1662 |
| Fbxo11 | Promoter (<=1kb) | 60 |
| Smad5 | Promoter (<=1kb) | 242 |
| Hnrnpd | Promoter (<=1kb) | -107 |
| Rev1 | Promoter (<=1kb) | 618 |
| Sinhcaf | Promoter (<=1kb) | 0 |
| Sumo2 | Promoter (<=1kb) | 464 |
| Zzz3 | Promoter (<=1kb) | 192 |
| Limk2 | Promoter (<=1kb) | 260 |
| Aqp11 | Distal Intergenic | -9992 |
| Atp13a3 | Distal Intergenic | -7145 |
| Cckar | Intron (ENSMUST00000201991.3/19664, intron 2 of 6) | 118309 |
| Ccne2 | Promoter (<=1kb) | -246 |
| Mertk | Intron (ENSMUST00000014505.4/17289, intron 4 of 18) | 47610 |
| Fgf1 | 5' UTR | -63557 |
| Ywhaz | Intron (ENSMUST00000022894.13/22631, intron 2 of 6) | 4183 |
| Nsd2 | Promoter (<=1kb) | 125 |
| Aqp11 | Distal Intergenic | -10572 |
| Lifr | Distal Intergenic | -51123 |
| NA | Promoter (<=1kb) | 0 |
| Arid5b | Intron (ENSMUST00000219238.1/71371, intron 4 of 9) | 141932 |
| Cdv3 | Promoter (<=1kb) | 511 |
| 4930483J18Rik | Promoter (<=1kb) | 35 |
| Ppp1r3b | Distal Intergenic | -81799 |
| Fam120a | Promoter (<=1kb) | 207 |
| Rev1 | Promoter (<=1kb) | 802 |
| Jak2 | Promoter (<=1kb) | -105 |
| Rev1 | Promoter (<=1kb) | 545 |
| Atp13a3 | Distal Intergenic | -6677 |
| Shh | Intron (ENSMUST00000180878.7/654796, intron 1 of 6) | -33979 |
| Atp13a3 | Distal Intergenic | -6722 |
| Smagp | Promoter (<=1kb) | 0 |
| Numa1 | 5' UTR | 35567 |
| Tmx1 | Distal Intergenic | 135760 |
| Ptpn2 | Promoter (<=1kb) | 415 |
| Rab21 | Promoter (<=1kb) | 72 |
| Ptpn2 | Promoter (<=1kb) | 442 |
| Cep43 | Promoter (<=1kb) | 416 |
| Gldc | Exon (ENSMUST00000025778.8/104174, exon 19 of 25) | 60211 |
| H4c1 | Promoter (<=1kb) | 89 |

| | | |
|---|---|---|
| Aox1 | Promoter (<=1kb) | 513 |
| Ndufv2 | Promoter (<=1kb) | 61 |
| Gm10406 | Intron (ENSMUST00000171906.1/ENSMUST00000171906.1, intron 1 of 6) | 137793 |
| NA | Promoter (<=1kb) | 759 |
| Gm21992 | Promoter (<=1kb) | 387 |
| Rprd1b | Promoter (<=1kb) | 0 |
| Cdyl2 | Promoter (<=1kb) | 0 |
| Ino80dos | Promoter (<=1kb) | -4 |
| Pals2 | Promoter (<=1kb) | -457 |
| Hey1 | Promoter (<=1kb) | -269 |
| Arid5b | 5' UTR | 142086 |
| Gldc | Exon (ENSMUST00000235189.1/104174, exon 19 of 19) | 60260 |
| Gm21992 | Promoter (<=1kb) | 347 |
| Tob1 | Promoter (<=1kb) | -604 |
| Bclaf1 | Promoter (<=1kb) | 0 |
| Ptpn2 | Promoter (<=1kb) | 457 |
| Sdf4 | Promoter (<=1kb) | 379 |
| Sf3b1 | Promoter (<=1kb) | 0 |
| Pan3 | Promoter (<=1kb) | -247 |
| Limk2 | Promoter (<=1kb) | 0 |
| Atp13a3 | Distal Intergenic | -7085 |
| Cc2d1b | Promoter (<=1kb) | -259 |
| Smim23 | Distal Intergenic | -14047 |
| Lbr | Promoter (<=1kb) | 707 |
| Sbf2 | Promoter (<=1kb) | 159 |
| Fhl2 | Distal Intergenic | -108880 |
| Rev1 | Promoter (<=1kb) | 216 |
| Bclaf1 | Promoter (<=1kb) | 27 |
| Ptpn2 | Promoter (<=1kb) | 224 |
| Agps | Promoter (<=1kb) | -143 |
| Mapk9 | Promoter (<=1kb) | 176 |
| Gldc | Exon (ENSMUST00000025778.8/104174, exon 19 of 25) | 59942 |
| Fgfbp1 | Distal Intergenic | 34222 |
| 9230116L04Rik | Promoter (<=1kb) | -168 |
| Myo10 | Promoter (<=1kb) | 319 |
| Itga6 | 5' UTR | 41181 |
| Srsf7 | Promoter (<=1kb) | 294 |
| Dhrs13 | Promoter (<=1kb) | -41 |
| Rev1 | Promoter (<=1kb) | 810 |
| Dolk | Promoter (<=1kb) | 0 |
| B9d1 | Promoter (<=1kb) | 6 |
| Ccne2 | Promoter (<=1kb) | -215 |
| Sf3b1 | Promoter (<=1kb) | 6 |

| | | |
|---|---|---|
| NA | Promoter (<=1kb) | 9 |
| Tmem131l | Promoter (<=1kb) | 208 |
| Gm10433 | Promoter (<=1kb) | -566 |
| Shh | Intron (ENSMUST00000180878.7/654796, intron 1 of 6) | -34148 |
| Cckar | Intron (ENSMUST00000201991.3/19664, intron 2 of 6) | 117618 |
| Hmgxb3 | Promoter (<=1kb) | -26 |
| Mapk9 | Promoter (<=1kb) | 128 |
| Ptpn2 | Promoter (<=1kb) | 599 |
| Zfp541 | Distal Intergenic | -4841 |
| Glt28d2 | Promoter (<=1kb) | -45 |
| Sde2 | Promoter (<=1kb) | 0 |
| Gm4792 | Promoter (1-2kb) | 1189 |
| Slc16a1 | Distal Intergenic | -27896 |
| a | Promoter (<=1kb) | 146 |
| Zfp664 | Promoter (<=1kb) | 118 |
| Azin1 | Promoter (<=1kb) | 0 |
| Dhx32 | Intron (ENSMUST00000065359.11/66930, intron 1 of 10) | -11995 |
| Aox1 | Promoter (<=1kb) | 638 |
| Slfnl1 | Intron (ENSMUST00000000087.12/29871, intron 12 of 16) | -15763 |
| Proser1 | Promoter (<=1kb) | 0 |
| Ctps | Promoter (<=1kb) | 3 |
| Gm21992 | Promoter (<=1kb) | 81 |
| Fbxl20 | Promoter (<=1kb) | 809 |
| Slc16a1 | Distal Intergenic | -27908 |
| Atp13a3 | Distal Intergenic | -6724 |
| Trappc8 | Promoter (<=1kb) | -472 |
| Itga6 | Intron (ENSMUST00000133124.7/16403, intron 3 of 5) | 41074 |
| Rock2 | Promoter (<=1kb) | -343 |
| Rev1 | Promoter (<=1kb) | 329 |
| Mbp | Intron (ENSMUST00000091789.10/17196, intron 1 of 3) | 17928 |
| Eif4g2 | Promoter (<=1kb) | 371 |
| Zfp648 | Distal Intergenic | -50874 |
| Fam220a | Promoter (<=1kb) | 0 |
| Hectd1 | Promoter (<=1kb) | 360 |
| Zfp541 | Distal Intergenic | -4735 |
| Rock2 | Promoter (<=1kb) | -329 |
| Hnrnpd | Promoter (<=1kb) | 0 |
| Nr2c2 | Promoter (<=1kb) | 580 |
| Itga6 | Intron (ENSMUST00000133124.7/16403, intron 3 of 5) | 41983 |
| Shh | Intron (ENSMUST00000180878.7/654796, intron 1 of 6) | -34263 |
| Ubtd1 | Promoter (<=1kb) | 153 |
| Rbm47 | Intron (ENSMUST00000201544.3/245945, intron 3 of 8) | 74342 |
| Memo1 | Promoter (<=1kb) | 0 |

| | | |
|---|---|---|
| Slc45a1 | Exon (ENSMUST00000142379.1/68703, exon 1 of 2) | 82127 |
| Efhc1 | 3' UTR | 48504 |
| Fhl2 | Distal Intergenic | -108230 |
| Zbed4 | Promoter (<=1kb) | -48 |
| Zfp541 | Distal Intergenic | -4356 |
| Rab15 | Exon (ENSMUST00000154743.1/110606, exon 1 of 6) | -64105 |
| Fnbp1l | Promoter (<=1kb) | 472 |
| Mrpl40 | Promoter (<=1kb) | 37 |
| Ptgfrn | Promoter (<=1kb) | -239 |
| Smad5 | Promoter (<=1kb) | -50 |
| Sft2d2 | Promoter (<=1kb) | 229 |
| Psmg2 | Promoter (<=1kb) | 0 |
| Myl12b | Promoter (<=1kb) | 0 |
| Crls1 | Promoter (<=1kb) | 201 |
| Zbtb10 | Promoter (<=1kb) | -180 |
| Fth1 | Promoter (2-3kb) | 2011 |
| Tmem131l | Promoter (<=1kb) | 22 |
| Azin1 | Promoter (<=1kb) | -515 |
| Ttl | Promoter (<=1kb) | 197 |
| Zbtb10 | Promoter (<=1kb) | -434 |
| Gldc | Intron (ENSMUST00000025778.8/104174, intron 18 of 24) | 59925 |
| Agps | Promoter (<=1kb) | -288 |
| Mapk9 | Promoter (<=1kb) | 15 |
| Zbtb10 | Promoter (<=1kb) | 38 |
| Ube2e3 | Promoter (1-2kb) | 1223 |
| Erc1 | Promoter (<=1kb) | 39 |
| Gldc | Intron (ENSMUST00000025778.8/104174, intron 18 of 24) | 59811 |
| Glis2 | Distal Intergenic | -7732 |
| Ndufv2 | Promoter (<=1kb) | 189 |
| Gmeb2 | Promoter (<=1kb) | 225 |
| Add1 | Promoter (<=1kb) | 0 |
| Gab2 | Promoter (<=1kb) | 274 |
| Tspan14 | Promoter (<=1kb) | 27 |
| 2900093K20Rik | Promoter (<=1kb) | -20 |
| Sbf2 | Promoter (<=1kb) | 0 |
| Cldn23 | Promoter (<=1kb) | 185 |
| Pan3 | Promoter (<=1kb) | -257 |
| Pik3cb | Promoter (<=1kb) | 467 |
| Tent5c | Promoter (<=1kb) | 460 |
| Srsf7 | Promoter (<=1kb) | 669 |
| Kcnk1 | Intron (ENSMUST00000046765.9/16525, intron 1 of 2) | 23548 |
| Ptpn2 | Promoter (<=1kb) | 217 |
| Fus | Promoter (<=1kb) | 145 |

| | | |
|---|---|---|
| Rpe | Exon (ENSMUST00000061620.16/329178, exon 44 of 65) | -54045 |
| Cdc42se1 | Promoter (<=1kb) | 73 |
| Stard9 | 3' UTR | 101738 |
| Crls1 | Promoter (<=1kb) | 160 |
| Cdca4 | Promoter (<=1kb) | -130 |
| Brd8 | Promoter (<=1kb) | 161 |
| Ppp4r1 | Promoter (<=1kb) | 81 |
| Nr2c2 | Promoter (<=1kb) | 982 |
| Fus | Promoter (<=1kb) | 127 |
| Rab6b | Exon (ENSMUST00000035148.12/24059, exon 10 of 14) | -34754 |
| Cebpd | Promoter (<=1kb) | 0 |
| Hdlbp | 5' UTR | 33291 |
| Sel1l3 | Intron (ENSMUST00000031090.7/231238, intron 9 of 23) | 44233 |
| Rprd2 | Promoter (<=1kb) | 0 |
| Cadps2 | Promoter (<=1kb) | -190 |
| Mbnl1 | 5' UTR | 28527 |
| Yy1 | Promoter (1-2kb) | 1590 |
| Eif4g2 | Promoter (<=1kb) | 553 |
| Kif20a | Promoter (<=1kb) | 45 |
| Hdlbp | 5' UTR | 33441 |
| Fam193b | Promoter (<=1kb) | 194 |
| Stt3b | Distal Intergenic | -22697 |
| Kif20a | Promoter (<=1kb) | 0 |
| Rpe | Intron (ENSMUST00000061620.16/329178, intron 44 of 64) | -53617 |
| A930033H14Rik | Promoter (<=1kb) | 48 |
| Stag1 | Intron (ENSMUST00000124487.7/20842, intron 1 of 3) | 45249 |
| Scd1 | Distal Intergenic | 18445 |

Table S8: AR Cluster 4 Genes Top500

| geneSymbol | annotation | distanceToTSS |
|---|---|---|
| 2310001H17Rik | Exon (ENSMUST00000181517.7/76432, exon 3 of 3) | 24063 |
| NA | 3' UTR | 16424 |
| NA | Intron (ENSMUST00000131021.1/ENSMUST00000131021.1, intron 2 of 2) | 16770 |
| NA | Intron (ENSMUST00000131021.1/ENSMUST00000131021.1, intron 2 of 2) | 16774 |
| 3110009F21Rik | Promoter (1-2kb) | 1231 |
| NA | 3' UTR | 17383 |
| Myof | Exon (ENSMUST00000172095.2/226101, exon 7 of 53) | 57494 |
| NA | Intron (ENSMUST00000131021.1/ENSMUST00000131021.1, intron 2 of 2) | 16783 |
| NA | 3' UTR | 17334 |
| Zfp217 | Distal Intergenic | 75671 |
| NA | Exon (ENSMUST00000131021.1/ENSMUST00000131021.1, exon 3 of 3) | 17326 |
| NA | 3' UTR | 17369 |
| Hs3st3b1 | Distal Intergenic | 359357 |
| Pira2 | Exon (ENSMUST00000153846.7/ENSMUST00000153846.7, exon 5 of 10) | 21917 |
| NA | Distal Intergenic | -420809 |
| Gpm6a | Distal Intergenic | 396273 |
| Hmgb4 | Intron (ENSMUST00000129619.7/329942, intron 5 of 12) | 8029 |
| Gpm6a | Distal Intergenic | 395921 |
| Gpm6a | Distal Intergenic | 395927 |
| Tmem117 | Intron (ENSMUST00000080141.5/320709, intron 3 of 7) | 152497 |
| Potefam3b | Intron (ENSMUST00000178995.2/ENSMUST00000178995.2, intron 4 of 4) | -32692 |
| NA | Exon (ENSMUST00000131021.1/ENSMUST00000131021.1, exon 1 of 3) | 16118 |
| Gpm6a | Distal Intergenic | 395929 |
| Gm30790 | Intron (ENSMUST00000208702.1/ENSMUST00000208702.1, intron 1 of 1) | 92314 |
| A530032D15Rik | Promoter (2-3kb) | 2256 |
| 6820431F20Rik | Distal Intergenic | 226223 |
| Potefam3b | Intron (ENSMUST00000178995.2/ENSMUST00000178995.2, intron 4 of 4) | -33419 |
| Agtr1a | Intron (ENSMUST00000222503.1/11607, intron 2 of 3) | 23456 |
| 6820431F20Rik | Distal Intergenic | 226244 |
| Wdr27 | Intron (ENSMUST00000232147.1/71682, intron 23 of 25) | 95756 |
| Asmt | Distal Intergenic | 62926 |
| 6820431F20Rik | Distal Intergenic | 226190 |
| NA | Distal Intergenic | -418916 |
| 6820431F20Rik | Distal Intergenic | -31689 |
| Gpm6a | Distal Intergenic | 391020 |
| 6820431F20Rik | Distal Intergenic | 225087 |
| 3110009F21Rik | Intron (ENSMUST00000222231.1/ENSMUST00000222231.1, intron 1 of 2) | 74853 |
| Gm5797 | Distal Intergenic | -17937 |
| 6820431F20Rik | Distal Intergenic | 213260 |
| Gpm6a | Distal Intergenic | 391191 |
| Gpm6a | Distal Intergenic | 417720 |
| 6820431F20Rik | Distal Intergenic | 213015 |

| | | |
|---|---|---|
| Gpm6a | Distal Intergenic | 391216 |
| Tmem18 | Distal Intergenic | -5408 |
| 6820431F20Rik | Distal Intergenic | 225719 |
| Gpm6a | Distal Intergenic | 391219 |
| 2610042L04Rik | Intron (ENSMUST00000166848.7/ENSMUST00000166848.7, intron 1 of 3) | 180526 |
| Vmn1r3 | Distal Intergenic | 6442 |
| 6820431F20Rik | Distal Intergenic | 225927 |
| 6820431F20Rik | Distal Intergenic | 225757 |
| Gpm6a | Distal Intergenic | 417469 |
| Snx19 | Distal Intergenic | -149757 |
| Gpm6a | Distal Intergenic | 417491 |
| Gpm6a | Distal Intergenic | 417523 |
| 6820431F20Rik | Distal Intergenic | 225086 |
| Mogat1 | Distal Intergenic | 63038 |
| Gpm6a | Distal Intergenic | 417718 |
| Mpp7 | Intron (ENSMUST00000115869.3/75739, intron 1 of 16) | 10159 |
| Gm6260 | Intron (ENSMUST00000192421.1/ENSMUST00000192421.1, intron 1 of 1) | 63355 |
| Mmp20 | Intron (ENSMUST00000034487.3/30800, intron 6 of 9) | 20421 |
| Gpm6a | Distal Intergenic | 417985 |
| 6820431F20Rik | Distal Intergenic | 225039 |
| Cacnb4 | Intron (ENSMUST00000178799.7/12298, intron 2 of 13) | 26868 |
| Apob | Distal Intergenic | -5650 |
| Potefam3b | Intron (ENSMUST00000178995.2/ENSMUST00000178995.2, intron 4 of 4) | -33355 |
| Gpm6a | Distal Intergenic | 408167 |
| Gpm6a | Distal Intergenic | 417256 |
| Mogat1 | Distal Intergenic | 63193 |
| Slc16a14 | Intron (ENSMUST00000159114.1/ENSMUST00000159114.1, intron 2 of 2) | -74824 |
| Gm10406 | Distal Intergenic | 102859 |
| Foxq1 | Distal Intergenic | -8218 |
| Nfkb1 | Distal Intergenic | -60303 |
| Gm11482 | Distal Intergenic | 30045 |
| Atg7 | Distal Intergenic | -25698 |
| Pard3 | Distal Intergenic | -4033 |
| Gpm6a | Distal Intergenic | 397811 |
| Gm10377 | Intron (ENSMUST00000165136.1/ENSMUST00000165136.1, intron 2 of 4) | -58811 |
| Snx19 | Distal Intergenic | -154501 |
| Gpm6a | Distal Intergenic | 397818 |
| Rbks | Intron (ENSMUST00000201352.3/107976, intron 3 of 11) | -77406 |
| Gpm6a | Distal Intergenic | 399865 |
| Gpm6a | Distal Intergenic | 397831 |
| 2610042L04Rik | Intron (ENSMUST00000166848.7/ENSMUST00000166848.7, intron 1 of 3) | 180333 |
| 6820431F20Rik | Distal Intergenic | 180005 |
| Gpm6a | Distal Intergenic | 418264 |

| | | |
|---|---|---|
| Pdxdc1 | Intron (ENSMUST00000023361.11/94184, intron 8 of 23) | 36957 |
| Myc | Distal Intergenic | -70116 |
| 6820431F20Rik | Distal Intergenic | 224642 |
| Gpm6a | Distal Intergenic | 417998 |
| Slc16a14 | Intron (ENSMUST00000159114.1/ENSMUST00000159114.1, intron 2 of 2) | -74619 |
| 4933406F09Rik | Intron (ENSMUST00000180508.7/68308, intron 2 of 2) | 73648 |
| Gpm6a | Distal Intergenic | 396223 |
| Gpm6a | Distal Intergenic | 399814 |
| Ippk | Distal Intergenic | -10997 |
| Gpm6a | Distal Intergenic | 396237 |
| Gpm6a | Distal Intergenic | 396253 |
| Mir6400 | Distal Intergenic | 27102 |
| Potefam3b | Distal Intergenic | -71599 |
| Gpm6a | Distal Intergenic | 419854 |
| Gpm6a | Distal Intergenic | 419873 |
| NA | Intron (ENSMUST00000101655.9/78887, intron 20 of 31) | 16101 |
| Taf1b | Distal Intergenic | -24841 |
| Gm3264 | Distal Intergenic | -35126 |
| Gpm6a | Distal Intergenic | 399809 |
| Gpm6a | Distal Intergenic | 419754 |
| 6820431F20Rik | Distal Intergenic | 279224 |
| A530032D15Rik | Intron (ENSMUST00000159114.1/ENSMUST00000159114.1, intron 2 of 2) | 73511 |
| Smok4a | Distal Intergenic | -70961 |
| Gpm6a | Distal Intergenic | 418300 |
| NA | Intron (ENSMUST00000101655.9/78887, intron 20 of 31) | 16095 |
| A530032D15Rik | Exon (ENSMUST00000186987.1/ENSMUST00000186987.1, exon 7 of 8) | 42042 |
| 2610005L07Rik | Distal Intergenic | -3818 |
| Gm3264 | Intron (ENSMUST00000180138.1/ENSMUST00000180138.1, intron 1 of 1) | 92231 |
| Rnf133 | Intron (ENSMUST00000166458.8/320405, intron 9 of 29) | 195834 |
| Fam3d | Distal Intergenic | -6942 |
| Bbs9 | Intron (ENSMUST00000039798.15/319845, intron 18 of 21) | 288271 |
| 6820431F20Rik | Distal Intergenic | 214035 |
| Gm3411 | Intron (ENSMUST00000168480.9/ENSMUST00000168480.9, intron 2 of 4) | 69424 |
| 6820431F20Rik | Distal Intergenic | 224621 |
| Gm3264 | Distal Intergenic | -35134 |
| Atg7 | Distal Intergenic | -24708 |
| Smok4a | Intron (ENSMUST00000137459.7/72578, intron 2 of 5) | 30335 |
| 2610005L07Rik | Distal Intergenic | -4116 |
| Cyp2b10 | Promoter (<=1kb) | -61 |
| Hbs1l | Distal Intergenic | -25488 |
| Cyp2b10 | Intron (ENSMUST00000072438.12/13088, intron 1 of 8) | 4379 |
| 6820431F20Rik | Distal Intergenic | 224614 |
| Mogat1 | Distal Intergenic | 63646 |

| | | |
|---|---|---|
| 6820431F20Rik | Distal Intergenic | 214055 |
| Gpm6a | Distal Intergenic | 418434 |
| Dlgap2 | Intron (ENSMUST00000152652.7/244310, intron 2 of 14) | 346415 |
| 4930442L01Rik | Intron (ENSMUST00000135031.7/59020, intron 1 of 2) | -8778 |
| Pdk4 | Distal Intergenic | 52804 |
| 6820431F20Rik | Distal Intergenic | 224594 |
| Gpm6a | Distal Intergenic | 418475 |
| 2610042L04Rik | Distal Intergenic | -3265 |
| Gpm6a | Distal Intergenic | 417213 |
| 2610042L04Rik | Intron (ENSMUST00000166848.7/ENSMUST00000166848.7, intron 1 of 3) | 178906 |
| 6820431F20Rik | Distal Intergenic | 212514 |
| C130026I21Rik | Intron (ENSMUST00000159114.1/ENSMUST00000159114.1, intron 2 of 2) | 27125 |
| Gpm6a | Distal Intergenic | 417206 |
| Gm3264 | Intron (ENSMUST00000180138.1/ENSMUST00000180138.1, intron 1 of 1) | 92007 |
| Aldh3a2 | Distal Intergenic | -27055 |
| Or2aj5 | Distal Intergenic | 5209 |
| 6820431F20Rik | Distal Intergenic | 226788 |
| Sp140 | Exon (ENSMUST00000161763.1/ENSMUST00000161763.1, exon 3 of 3) | 3962 |
| Gm3411 | Intron (ENSMUST00000168480.9/ENSMUST00000168480.9, intron 4 of 4) | 72003 |
| Atg7 | Distal Intergenic | -12724 |
| Gpm6a | Distal Intergenic | 417145 |
| Gpm6a | Distal Intergenic | 418514 |
| Slc16a14 | Distal Intergenic | -30751 |
| Zfp982 | Distal Intergenic | -142794 |
| 2610042L04Rik | Intron (ENSMUST00000166848.7/ENSMUST00000166848.7, intron 1 of 3) | 179224 |
| Gpm6a | Distal Intergenic | 408484 |
| NA | Intron (ENSMUST00000101655.9/78887, intron 20 of 31) | 15608 |
| Mogat1 | Distal Intergenic | 63479 |
| NA | Intron (ENSMUST00000101655.9/78887, intron 20 of 31) | 15566 |
| Gpm6a | Distal Intergenic | 408176 |
| Potefam3b | Distal Intergenic | 32951 |
| Gpm6a | Distal Intergenic | 399307 |
| 6820431F20Rik | Distal Intergenic | 226246 |
| Gm31763 | Intron (ENSMUST00000217788.1/102634101, intron 1 of 4) | 3156 |
| NA | Exon (ENSMUST00000231996.1/ENSMUST00000231996.1, exon 6 of 6) | -44567 |
| A530032D15Rik | Exon (ENSMUST00000161724.7/381287, exon 5 of 8) | 18696 |
| Rad18 | Distal Intergenic | -13915 |
| Atg7 | Distal Intergenic | -13252 |
| Gm3264 | Distal Intergenic | 81161 |
| Atg7 | Distal Intergenic | -62608 |
| Potefam3b | Distal Intergenic | 32950 |
| 2610005L07Rik | Distal Intergenic | -87631 |
| 2610005L07Rik | Distal Intergenic | -87628 |

| | | |
|---|---|---|
| 2610005L07Rik | Distal Intergenic | -87626 |
| 2610042L04Rik | Intron (ENSMUST00000166848.7/ENSMUST00000166848.7, intron 1 of 3) | 179246 |
| Gm2897 | Distal Intergenic | -30563 |
| Sycp3 | 3' UTR | -13359 |
| Potefam3b | Distal Intergenic | 33012 |
| Gm10413 | Distal Intergenic | -64024 |
| Atg7 | Distal Intergenic | -13908 |
| Mylk4 | Distal Intergenic | 191657 |
| Gpm6a | Distal Intergenic | 399494 |
| Potefam3b | Distal Intergenic | -30258 |
| A530032D15Rik | Intron (ENSMUST00000159114.1/ENSMUST00000159114.1, intron 2 of 2) | 17396 |
| Aass | Exon (ENSMUST00000096092.3/ENSMUST00000096092.3, exon 1 of 1) | -27044 |
| C130026I21Rik | Intron (ENSMUST00000161923.1/ENSMUST00000161923.1, intron 1 of 1) | -60178 |
| NA | Intron (ENSMUST00000101655.9/78887, intron 20 of 31) | 15448 |
| Gm3264 | Intron (ENSMUST00000166025.7/ENSMUST00000166025.7, intron 4 of 4) | -13627 |
| Nudt9 | Distal Intergenic | -11331 |
| Gpm6a | Distal Intergenic | 399533 |
| 4930521O11Rik | Promoter (<=1kb) | 0 |
| Btbd35f20 | Distal Intergenic | 222888 |
| Or10v5 | Intron (ENSMUST00000214887.1/ENSMUST00000214887.1, intron 1 of 3) | -6711 |
| Potefam3b | Distal Intergenic | 83673 |
| Tnc | Exon (ENSMUST00000030056.11/21923, exon 2 of 25) | 29839 |
| Potefam3b | Distal Intergenic | 33165 |
| Mir148a | Distal Intergenic | 124782 |
| NA | Intron (ENSMUST00000101655.9/78887, intron 21 of 31) | 11296 |
| Psg25 | Intron (ENSMUST00000094795.4/114868, intron 2 of 4) | 3441 |
| Lonrf1 | Distal Intergenic | -12400 |
| C130026I21Rik | Intron (ENSMUST00000161267.7/620078, intron 1 of 2) | 11656 |
| 6820431F20Rik | Distal Intergenic | 181893 |
| Gm10375 | Intron (ENSMUST00000166110.1/ENSMUST00000166110.1, intron 2 of 4) | 30978 |
| Mir1951 | Intron (ENSMUST00000166472.7/399568, intron 5 of 10) | 16353 |
| Slc16a14 | Intron (ENSMUST00000159114.1/ENSMUST00000159114.1, intron 2 of 2) | -74379 |
| Potefam3b | Distal Intergenic | 64332 |
| Sp110 | Intron (ENSMUST00000161449.1/ENSMUST00000161449.1, intron 1 of 1) | 75700 |
| Potefam3b | Distal Intergenic | 32651 |
| Gm3264 | Distal Intergenic | -37102 |
| Gpm6a | Distal Intergenic | 399267 |
| Potefam3b | Distal Intergenic | 33194 |
| Trpm1 | Distal Intergenic | -67901 |
| Potefam3b | Distal Intergenic | 83641 |
| Potefam3b | Distal Intergenic | -30536 |
| Gpm6a | Distal Intergenic | 408659 |
| 6820431F20Rik | Distal Intergenic | 212455 |

| | | |
|---|---|---|
| Potefam3b | Distal Intergenic | 64312 |
| Ulk4 | Intron (ENSMUST00000171923.7/209012, intron 17 of 36) | 52202 |
| Mrpl40 | Intron (ENSMUST00000004222.13/15260, intron 24 of 24) | -81540 |
| A530032D15Rik | Intron (ENSMUST00000159114.1/ENSMUST00000159114.1, intron 2 of 2) | 17404 |
| NA | Intron (ENSMUST00000101655.9/78887, intron 20 of 31) | 15628 |
| Pcp4 | Intron (ENSMUST00000056102.8/13508, intron 20 of 32) | 198816 |
| Gm3099 | Distal Intergenic | 16093 |
| 6820431F20Rik | Distal Intergenic | -6358 |
| Atp5g3 | Distal Intergenic | -173416 |
| C130026I21Rik | Intron (ENSMUST00000166777.3/ENSMUST00000166777.3, intron 1 of 4) | -51457 |
| Nt5e | Distal Intergenic | -96924 |
| Gm3696 | Distal Intergenic | -4352 |
| Mir5127 | Distal Intergenic | -169293 |
| Defb13 | Promoter (<=1kb) | 993 |
| Sp110 | Distal Intergenic | 143933 |
| Gm3411 | Intron (ENSMUST00000170104.2/100861615, intron 2 of 5) | 11700 |
| 6820431F20Rik | Distal Intergenic | 197557 |
| NA | Exon (ENSMUST00000092696.4/381715, exon 4 of 5) | 3498 |
| Atg7 | Distal Intergenic | -13885 |
| Potefam3b | Distal Intergenic | 64276 |
| Pik3c2g | Intron (ENSMUST00000187069.6/18705, intron 4 of 11) | 154394 |
| NA | Intron (ENSMUST00000178227.5/ENSMUST00000178227.5, intron 2 of 2) | 3722 |
| Bmpr1b | Intron (ENSMUST00000098568.7/12167, intron 2 of 11) | 104007 |
| A530032D15Rik | Promoter (2-3kb) | 2457 |
| Potefam3e | Distal Intergenic | 205457 |
| Potefam3b | Distal Intergenic | 83587 |
| Wdr7 | Distal Intergenic | -110376 |
| Gm3636 | Distal Intergenic | -106515 |
| Gm3411 | Intron (ENSMUST00000168480.9/ENSMUST00000168480.9, intron 2 of 4) | 69396 |
| Ctsl | Distal Intergenic | -17600 |
| Tcp10c | Intron (ENSMUST00000233884.1/100041352, intron 11 of 12) | 16435 |
| NA | Intron (ENSMUST00000178227.5/ENSMUST00000178227.5, intron 2 of 2) | 3624 |
| 1700091H14Rik | Distal Intergenic | -79372 |
| Dlgap2 | Intron (ENSMUST00000152652.7/244310, intron 2 of 14) | 193582 |
| Akp3 | Distal Intergenic | 14537 |
| Asmt | Distal Intergenic | 65888 |
| C130026I21Rik | Intron (ENSMUST00000161267.7/620078, intron 1 of 2) | 13892 |
| Atg7 | Distal Intergenic | -24716 |
| Gm3411 | Intron (ENSMUST00000181015.7/ENSMUST00000181015.7, intron 2 of 2) | -136339 |
| Gpm6a | Distal Intergenic | 408733 |
| Lpcat4 | Distal Intergenic | -20029 |
| Atg7 | Distal Intergenic | -49635 |
| Gpm6a | Distal Intergenic | 398884 |

| | | |
|---|---|---|
| 2610042L04Rik | Intron (ENSMUST00000166848.7/ENSMUST00000166848.7, intron 1 of 3) | 178896 |
| Trpm1 | Distal Intergenic | -78547 |
| Gm6578 | Distal Intergenic | -150823 |
| A530032D15Rik | Intron (ENSMUST00000159114.1/ENSMUST00000159114.1, intron 2 of 2) | 17567 |
| Atg7 | Distal Intergenic | -13452 |
| Gm3264 | Distal Intergenic | -37137 |
| Fosl2 | Intron (ENSMUST00000201352.3/107976, intron 7 of 11) | -146929 |
| 6820431F20Rik | Distal Intergenic | 227234 |
| Chd9 | Intron (ENSMUST00000048665.7/109151, intron 2 of 39) | 58096 |
| Gnb4 | Distal Intergenic | -9553 |
| Gm3411 | Intron (ENSMUST00000238283.1/ENSMUST00000238283.1, intron 2 of 3) | -47887 |
| Gm10033 | Distal Intergenic | -77103 |
| Mir1951 | Intron (ENSMUST00000166472.7/399568, intron 5 of 10) | 16478 |
| Gm25018 | Promoter (1-2kb) | -1737 |
| Gm3411 | Intron (ENSMUST00000170104.2/100861615, intron 2 of 5) | 11711 |
| Gm3264 | Distal Intergenic | -31909 |
| 6820431F20Rik | Distal Intergenic | 226761 |
| Atg7 | Distal Intergenic | -13646 |
| Sp140 | Intron (ENSMUST00000113385.2/434484, intron 2 of 3) | 4186 |
| Potefam3a | Distal Intergenic | 31607 |
| Potefam3e | Distal Intergenic | 205476 |
| 2610005L07Rik | Distal Intergenic | -87272 |
| Gm5796 | Distal Intergenic | -34793 |
| Serpina3n | 5' UTR | -19525 |
| 2610005L07Rik | Distal Intergenic | -87265 |
| 2610005L07Rik | Distal Intergenic | -87235 |
| 1700091H14Rik | Distal Intergenic | -79549 |
| 2610042L04Rik | Intron (ENSMUST00000166848.7/ENSMUST00000166848.7, intron 1 of 3) | 178830 |
| Gm3411 | Distal Intergenic | 62560 |
| Cttnbp2nl | Intron (ENSMUST00000077548.11/80281, intron 3 of 5) | 23917 |
| Gm2897 | Distal Intergenic | -33387 |
| 1700091H14Rik | Distal Intergenic | -79561 |
| Atg7 | Distal Intergenic | -20191 |
| Btbd35f20 | Distal Intergenic | 223952 |
| Gm3411 | Intron (ENSMUST00000168480.9/ENSMUST00000168480.9, intron 4 of 4) | 72510 |
| Arhgap28 | Promoter (<=1kb) | -30 |
| Gm25018 | Promoter (2-3kb) | -2880 |
| Gm3411 | Intron (ENSMUST00000168480.9/ENSMUST00000168480.9, intron 2 of 4) | 69352 |
| 6820431F20Rik | Distal Intergenic | 212185 |
| 6820431F20Rik | Distal Intergenic | 226745 |
| Gm21671 | Exon (ENSMUST00000196214.1/ENSMUST00000196214.1, exon 4 of 5) | 15472 |
| Ear10 | Distal Intergenic | -14334 |
| Gm3411 | Intron (ENSMUST00000181015.7/ENSMUST00000181015.7, intron 2 of 2) | -136094 |

| | | |
|---|---|---|
| Potefam3b | Distal Intergenic | 49908 |
| C130026I21Rik | Distal Intergenic | -74192 |
| Gm25018 | Promoter (2-3kb) | -2067 |
| 2610005L07Rik | Distal Intergenic | -87282 |
| Potefam3a | Distal Intergenic | 31678 |
| Rapgef5 | Distal Intergenic | -64660 |
| Trappc6b | Distal Intergenic | 23310 |
| 2610005L07Rik | Exon (ENSMUST00000139264.1/381598, exon 5 of 5) | 29220 |
| Gm25018 | Promoter (2-3kb) | -2867 |
| Gpm6a | Distal Intergenic | 159263 |
| Gm25018 | Promoter (2-3kb) | -2075 |
| Clic5 | Distal Intergenic | -22591 |
| Gm3411 | Distal Intergenic | 113908 |
| Potefam3a | Distal Intergenic | 31636 |
| 6820431F20Rik | Distal Intergenic | 227281 |
| Gpm6a | Distal Intergenic | 398146 |
| 2610005L07Rik | Intron (ENSMUST00000140111.7/381598, intron 5 of 5) | 29195 |
| Adgrl2 | Distal Intergenic | 574137 |
| Gm3411 | Intron (ENSMUST00000168480.9/ENSMUST00000168480.9, intron 4 of 4) | 71719 |
| Gm25018 | Promoter (2-3kb) | -2305 |
| 1700091H14Rik | Distal Intergenic | -79351 |
| C130026I21Rik | Distal Intergenic | -74140 |
| Ifnk | Intron (ENSMUST00000102975.9/214944, intron 2 of 3) | -104345 |
| Gm3264 | Intron (ENSMUST00000180138.1/ENSMUST00000180138.1, intron 1 of 1) | 91968 |
| C130026I21Rik | Distal Intergenic | -74158 |
| Mir694 | Promoter (1-2kb) | 1634 |
| Gm2897 | Distal Intergenic | -33438 |
| C130026I21Rik | Intron (ENSMUST00000166777.3/ENSMUST00000166777.3, intron 1 of 4) | -53725 |
| Setbp1 | Distal Intergenic | -308131 |
| Cmas | Distal Intergenic | -5660 |
| Nobox | Distal Intergenic | 13082 |
| Potefam3b | Distal Intergenic | -56160 |
| 2610005L07Rik | Intron (ENSMUST00000140111.7/381598, intron 5 of 5) | 28972 |
| Potefam3b | Distal Intergenic | 83369 |
| 6820431F20Rik | Distal Intergenic | 197301 |
| Trpm1 | Distal Intergenic | -66336 |
| C130026I21Rik | Distal Intergenic | -74353 |
| 6820431F20Rik | Distal Intergenic | 212183 |
| 1700091H14Rik | Distal Intergenic | -79304 |
| NA | Intron (ENSMUST00000101655.9/78887, intron 21 of 31) | 11328 |
| Tmem117 | Intron (ENSMUST00000080141.5/320709, intron 3 of 7) | 95083 |
| 2610042L04Rik | Distal Intergenic | -96779 |
| Mir5127 | Distal Intergenic | -169131 |

| | | |
|---|---|---|
| Potefam3a | Distal Intergenic | 31638 |
| Atg7 | Distal Intergenic | -50719 |
| Atg7 | Distal Intergenic | -50903 |
| Atg7 | Distal Intergenic | -50704 |
| 6820431F20Rik | Distal Intergenic | 212167 |
| Gm3411 | Intron (ENSMUST00000181015.7/ENSMUST00000181015.7, intron 2 of 2) | -147495 |
| Fer | Distal Intergenic | -99996 |
| Asmt | Distal Intergenic | 174012 |
| Atg7 | Distal Intergenic | -24915 |
| Gm3411 | Distal Intergenic | 113771 |
| Slc16a14 | Intron (ENSMUST00000159114.1/ENSMUST00000159114.1, intron 2 of 2) | -72227 |
| Nkx3-1 | Distal Intergenic | 30911 |
| 2610042L04Rik | Distal Intergenic | 258064 |
| Pop4 | Promoter (<=1kb) | 0 |
| Gm3411 | Intron (ENSMUST00000238283.1/ENSMUST00000238283.1, intron 2 of 3) | -47871 |
| Mir1951 | Intron (ENSMUST00000166472.7/399568, intron 5 of 10) | 16526 |
| 2610042L04Rik | Intron (ENSMUST00000166848.7/ENSMUST00000166848.7, intron 1 of 3) | 180233 |
| Gm3411 | Distal Intergenic | -187244 |
| Pvt1 | Distal Intergenic | -25890 |
| Cutal | Intron (ENSMUST00000028225.11/66998, intron 5 of 9) | -12646 |
| Lmntd1 | Intron (ENSMUST00000148739.7/74071, intron 1 of 10) | 18042 |
| Atg7 | Distal Intergenic | -52987 |
| Potefam3b | Distal Intergenic | 90897 |
| Mir28b | Intron (ENSMUST00000004497.10/16795, intron 6 of 14) | 16919 |
| Sh3gl3 | Intron (ENSMUST00000179318.7/20408, intron 1 of 8) | 39687 |
| Ptprd | Distal Intergenic | -301711 |
| Nek7 | Intron (ENSMUST00000186017.6/59125, intron 7 of 9) | 112959 |
| Atg7 | Distal Intergenic | -10811 |
| Atg7 | Distal Intergenic | -40388 |
| NA | Intron (ENSMUST00000101655.9/78887, intron 13 of 31) | 22730 |
| Efcab11 | Intron (ENSMUST00000221929.1/78767, intron 5 of 5) | 68811 |
| Txndc8 | Distal Intergenic | 25726 |
| Gm5797 | Intron (ENSMUST00000100886.3/545013, intron 4 of 4) | 7770 |
| Potefam3b | Distal Intergenic | 50123 |
| Mboat1 | Distal Intergenic | -26981 |
| Potefam3b | Distal Intergenic | 50205 |
| Sgip1 | Intron (ENSMUST00000106911.7/18578, intron 3 of 16) | -311940 |
| Prdm1 | Promoter (1-2kb) | 1356 |
| Edn1 | Distal Intergenic | -59915 |
| 6820431F20Rik | Distal Intergenic | 211424 |
| Gm5797 | Intron (ENSMUST00000100886.3/545013, intron 4 of 4) | 7737 |
| Btbd18 | Intron (ENSMUST00000099941.8/12388, intron 1 of 17) | -21239 |
| Prr15 | Intron (ENSMUST00000145111.7/100042056, intron 3 of 13) | 33307 |

| | | |
|---|---|---|
| Potefam3b | Distal Intergenic | 49932 |
| 6820431F20Rik | Distal Intergenic | 197242 |
| NA | Intron (ENSMUST00000170480.7/100040852, intron 4 of 6) | 15933 |
| Mir3106 | Intron (ENSMUST00000082104.6/94109, intron 3 of 69) | -625096 |
| 4933406F09Rik | Exon (ENSMUST00000165448.1/ENSMUST00000165448.1, exon 1 of 1) | 6901 |
| Gm3264 | Distal Intergenic | -37176 |
| Vmn1r41 | Distal Intergenic | -9358 |
| Lhfp | Intron (ENSMUST00000059562.13/108927, intron 2 of 3) | 151650 |
| Gm3411 | Distal Intergenic | 82187 |
| 6820431F20Rik | Distal Intergenic | 211458 |
| NA | 3' UTR | -209639 |
| Lmx1a | Intron (ENSMUST00000177351.2/ENSMUST00000177351.2, intron 1 of 2) | 360832 |
| Exoc2 | Promoter (<=1kb) | 156 |
| Ang5 | Distal Intergenic | 62935 |
| Gpm6a | Distal Intergenic | 408736 |
| Gm3264 | Distal Intergenic | -37386 |
| Gpr50 | Distal Intergenic | -18078 |
| Asmt | Promoter (<=1kb) | 927 |
| Apaf1 | Intron (ENSMUST00000183156.7/77531, intron 15 of 25) | 486252 |
| Asmt | Promoter (<=1kb) | 920 |
| Miga1 | Intron (ENSMUST00000073089.12/215708, intron 7 of 15) | 25568 |
| Gm3264 | Distal Intergenic | -37421 |
| Golph3 | Distal Intergenic | -35796 |
| Gm3264 | Distal Intergenic | -29030 |
| NA | Intron (ENSMUST00000170480.7/100040852, intron 4 of 6) | 15966 |
| Chd6 | Intron (ENSMUST00000109443.7/19281, intron 17 of 30) | -490081 |
| Potefam3e | Distal Intergenic | -29999 |
| Mir30a | Intron (ENSMUST00000191045.1/ENSMUST00000191045.1, intron 2 of 3) | -69528 |
| Gm3264 | Intron (ENSMUST00000166776.7/100041306, intron 2 of 6) | 3518 |
| NA | 3' UTR | -209687 |
| Sp140 | Intron (ENSMUST00000113385.2/434484, intron 2 of 3) | 8592 |
| Miga1 | Intron (ENSMUST00000073089.12/215708, intron 7 of 15) | 25407 |
| NA | 3' UTR | -209692 |
| Potefam3b | Distal Intergenic | 83327 |
| 6820431F20Rik | Distal Intergenic | 211495 |
| NA | Intron (ENSMUST00000101655.9/78887, intron 21 of 31) | 11594 |
| Cdkal1 | Distal Intergenic | -29684 |
| Slc16a14 | Intron (ENSMUST00000159114.1/ENSMUST00000159114.1, intron 2 of 2) | -74358 |
| 2610005L07Rik | Promoter (2-3kb) | 2486 |
| Atg7 | Distal Intergenic | -53527 |
| Gm3264 | Distal Intergenic | -29182 |
| 2610005L07Rik | Intron (ENSMUST00000140111.7/381598, intron 5 of 5) | 28955 |
| Dnah10 | Intron (ENSMUST00000037865.12/21871, intron 9 of 19) | -14559 |

| | | |
|---|---|---|
| Gm21671 | Intron (ENSMUST00000196214.1/ENSMUST00000196214.1, intron 3 of 4) | 15468 |
| Gm3636 | Intron (ENSMUST00000181787.1/ENSMUST00000181787.1, intron 2 of 2) | -40532 |
| Gm3411 | Exon (ENSMUST00000168480.9/ENSMUST00000168480.9, exon 4 of 5) | 70657 |
| Asmt | Distal Intergenic | 174036 |
| Raet1d | Intron (ENSMUST00000105522.8/667281, intron 1 of 5) | 20299 |
| Fosl2 | Intron (ENSMUST00000201352.3/107976, intron 7 of 11) | -146731 |
| Otulin | 3' UTR | 14167 |
| Cutal | Intron (ENSMUST00000028225.11/66998, intron 5 of 9) | -12690 |
| Fbxo22 | Intron (ENSMUST00000139261.7/83961, intron 2 of 7) | 50829 |
| Miga1 | Intron (ENSMUST00000073089.12/215708, intron 7 of 15) | 25381 |
| Dnah10 | Intron (ENSMUST00000037865.12/21871, intron 9 of 19) | -14576 |
| Hs3st3b1 | Distal Intergenic | 336524 |
| Potefam3b | Distal Intergenic | 63971 |
| Dnah10 | Intron (ENSMUST00000037865.12/21871, intron 9 of 19) | -14578 |
| 6820431F20Rik | Distal Intergenic | 211551 |
| Mesd | Distal Intergenic | 43198 |
| C130026I21Rik | Intron (ENSMUST00000166777.3/ENSMUST00000166777.3, intron 1 of 4) | -51259 |
| Asmt | Distal Intergenic | 153444 |
| D830030K20Rik | Intron (ENSMUST00000225885.1/320333, intron 3 of 4) | 18481 |
| Potefam3b | Distal Intergenic | 83308 |
| Gm22109 | Intron (ENSMUST00000127664.1/108148, intron 1 of 15) | -5661 |
| 2610042L04Rik | Distal Intergenic | 257754 |
| 6820431F20Rik | Distal Intergenic | 211738 |
| Gm3411 | Exon (ENSMUST00000170104.2/100861615, exon 5 of 6) | 22694 |
| Atg7 | Distal Intergenic | -53145 |
| Cth | Intron (ENSMUST00000118539.1/107869, intron 5 of 11) | 13236 |
| Atg7 | Distal Intergenic | -53325 |
| Gm3411 | Distal Intergenic | 103372 |
| Gm3264 | Intron (ENSMUST00000166025.7/ENSMUST00000166025.7, intron 1 of 4) | -16714 |
| Gnaz | Distal Intergenic | -46044 |
| NA | Intron (ENSMUST00000101655.9/78887, intron 21 of 31) | 12402 |
| Gm3264 | Intron (ENSMUST00000166025.7/ENSMUST00000166025.7, intron 4 of 4) | -11626 |
| Gm22109 | Exon (ENSMUST00000179476.1/ENSMUST00000179476.1, exon 1 of 1) | -36892 |
| Atg7 | Distal Intergenic | -55156 |
| Gm3411 | Distal Intergenic | -187069 |
| Atg7 | Distal Intergenic | -21356 |
| Trpm1 | Distal Intergenic | -78370 |
| Gm3411 | 3' UTR | 26138 |
| Gm22109 | Intron (ENSMUST00000127664.1/108148, intron 1 of 15) | -5477 |
| Cdk15 | Intron (ENSMUST00000160662.7/271697, intron 13 of 13) | 89575 |
| 6820431F20Rik | Distal Intergenic | 211922 |
| Gm3099 | Distal Intergenic | -47000 |
| Atg7 | Distal Intergenic | -39978 |

| | | |
|---|---|---|
| Vcl | Exon (ENSMUST00000226113.1/22330, exon 5 of 8) | 56147 |
| Gm25018 | Exon (ENSMUST00000178755.1/100861502, exon 1 of 1) | -4821 |
| Atg7 | Distal Intergenic | -51216 |
| Gm22109 | Intron (ENSMUST00000127664.1/108148, intron 1 of 15) | -5667 |
| Eapp | Distal Intergenic | -33228 |
| 4930578N18Rik | Distal Intergenic | 96177 |
| Gm22109 | Intron (ENSMUST00000127664.1/108148, intron 1 of 15) | -5441 |
| 1700091H14Rik | Distal Intergenic | -78839 |
| Gm10406 | Distal Intergenic | 38359 |
| Gm3636 | Distal Intergenic | 135824 |
| Trpm1 | Distal Intergenic | -77604 |
| Atg7 | Distal Intergenic | -62429 |
| Adgb | Distal Intergenic | 150897 |
| Trpm1 | Distal Intergenic | -67132 |
| Dusp6 | Intron (ENSMUST00000189035.1/ENSMUST00000189035.1, intron 1 of 2) | 76853 |
| 2610042L04Rik | Distal Intergenic | 257824 |
| 1700095A21Rik | Distal Intergenic | -55840 |
| Flnb | Intron (ENSMUST00000052678.8/286940, intron 3 of 45) | 52632 |
| Gm3411 | Intron (ENSMUST00000170104.2/100861615, intron 2 of 5) | 11734 |
| 2610005L07Rik | Intron (ENSMUST00000140111.7/381598, intron 5 of 5) | 28917 |
| Aggf1 | Promoter (<=1kb) | 521 |
| Zfp455 | Promoter (<=1kb) | 27 |
| Gm3411 | Exon (ENSMUST00000190582.6/ENSMUST00000190582.6, exon 2 of 5) | 162265 |
| Gm4956 | Distal Intergenic | -8305 |
| Asmt | Distal Intergenic | 174310 |
| Trpm1 | Distal Intergenic | -67144 |
| Osbpl1a | Intron (ENSMUST00000121808.7/64291, intron 1 of 13) | 87683 |

### Table S9: AR Cluster 1 Reactome Pathways Top50

| Term | Combined Score |
|---|---|
| Diseases of Signal Transduction by Growth Factor Receptors and Second Messengers | 42.8475917175466 |
| Signaling by NTRK1 (TRKA) | 60.852649835615615 |
| Signaling by NTRKs | 55.94898472495188 |
| Nuclear Events (Kinase and Transcription Factor Activation) | 72.81066050553694 |
| Signal Transduction | 21.713605876038166 |
| Axon Guidance | 23.7312215109261 |
| Signaling by Receptor Tyrosine Kinases | 23.2947908087251 |
| Nervous System Development | 22.76107962350115 |
| Signaling by ALK Fusions and Activated Point Mutants | 41.04595799644717 |
| Signaling by ALK in Cancer | 41.04595799644717 |
| Signaling by Rho GTPases | 21.103202239307816 |
| Signaling by Rho GTPases, Miro GTPases and RHOBTB3 | 20.476224295037312 |
| Chromatin Modifying Enzymes | 25.756546989718053 |
| Chromatin Organization | 25.756546989718053 |
| Circadian Clock | 39.290613266569714 |
| Cellular Responses to Stimuli | 17.309210015525938 |
| Signaling by TGFB Family Members | 25.164028308492366 |
| Cellular Responses to Stress | 16.824109845293044 |
| Interleukin-17 Signaling | 29.800131446702725 |
| Negative Regulation of MAPK Pathway | 39.38194929703357 |
| MAPK Family Signaling Cascades | 17.562834583063474 |
| Oncogenic MAPK Signaling | 25.773681673886706 |
| Membrane Trafficking | 14.448391333416264 |
| NGF-stimulated Transcription | 38.22303047816518 |
| MAP Kinase Activation | 28.318428962495403 |
| ESR-mediated Signaling | 18.277535701026917 |
| Intrinsic Pathway for Apoptosis | 30.12881651642594 |
| RHO GTPase Effectors | 15.660688482510597 |
| Transcriptional Regulation by NPAS4 | 37.20626848784401 |
| HSP90 Chaperone Cycle for Steroid Hormone Receptors (SHR) in the Presence of Ligand | 27.837168546317816 |
| RHO GTPase Cycle | 14.017644724751076 |
| Integration of Energy Metabolism | 20.41654972921055 |
| Signaling by BRAF and RAF1 Fusions | 24.642421502100994 |
| PKMTs Methylate Histone Lysines | 29.354978985687012 |
| Apoptosis | 16.726784688890067 |
| MAPK Targets Nuclear Events Mediated by MAP Kinases | 36.3993544840078 |
| Signaling by Nuclear Receptors | 14.17358811010543 |
| Eukaryotic Translation Elongation | 18.550225440368536 |
| Toll Like Receptor 3 (TLR3) Cascade | 17.343505957685416 |
| Oncogene Induced Senescence | 28.511419597998064 |
| Signaling by WNT | 12.877868570527381 |
| MyD88-independent TLR4 Cascade | 16.441564398219928 |

| | |
|---|---|
| TRIF (TICAM1)-mediated TLR4 Signaling | 16.441564398219928 |
| G Alpha (12 13) Signalling Events | 18.431403475894147 |
| MyD88 Cascade Initiated on Plasma Membrane | 16.994943030622213 |
| Toll Like Receptor 10 (TLR10) Cascade | 16.994943030622213 |
| Toll Like Receptor 5 (TLR5) Cascade | 16.994943030622213 |
| TGF-beta Receptor Signaling in EMT (Epithelial to Mesenchymal Transition) | 51.076691194941716 |
| TRAF6 Mediated Induction of NFkB and MAP Kinases Upon TLR7 8 or 9 Activation | 16.039716999230166 |
| Repression of WNT Target Genes | 56.81883746313349 |

Table S9: AR Cluster 2 Reactome Pathways Top50

| Term | Combined Score |
|---|---|
| RAC1 GTPase Cycle | 19.260338404447804 |
| RHO GTPase Cycle | 13.36556948037757 |
| Signaling by Rho GTPases | 10.149015552437966 |
| VEGFA-VEGFR2 Pathway | 16.66953517153804 |
| Signaling by Rho GTPases, Miro GTPases and RHOBTB3 | 9.542271945028116 |
| Signaling by VEGF | 12.593594591882825 |
| Synthesis of PA | 19.024513442407258 |
| Regulation of Localization of FOXO Transcription Factors | 38.1809118721766 |
| Binding of TCF LEF CTNNB1 to Target Gene Promoters | 51.70740327412844 |
| RUNX3 Regulates WNT Signaling | 51.70740327412844 |
| Acyl Chain Remodelling of PE | 19.33609008995468 |
| Intrinsic Pathway for Apoptosis | 13.998831479372104 |
| Signaling by Receptor Tyrosine Kinases | 7.069335608868391 |
| Signaling by BMP | 16.211956347184724 |
| NGF-independant TRKA Activation | 69.4278439912912 |
| Constitutive Signaling by NOTCH1 HD+PEST Domain Mutants | 11.987727417971227 |
| Constitutive Signaling by NOTCH1 PEST Domain Mutants | 11.987727417971227 |
| Signaling by NOTCH1 HD+PEST Domain Mutants in Cancer | 11.987727417971227 |
| Signaling by NOTCH1 PEST Domain Mutants in Cancer | 11.987727417971227 |
| Signaling by NOTCH1 in Cancer | 11.987727417971227 |
| Acyl Chain Remodelling of PC | 14.789982468734953 |
| NOTCH1 Intracellular Domain Regulates Transcription | 10.660715320617895 |
| Activation of TRKA Receptors | 40.41613832740221 |
| Acyl Chain Remodelling of PS | 14.971448283324225 |
| Transcriptional Regulation by RUNX3 | 8.315863340841119 |
| Acyl Chain Remodelling of PI | 15.746729123047897 |
| Activation of BH3-only Proteins | 11.192155837148366 |
| Signaling by NOTCH1 | 7.68368915146209 |
| Oncogenic MAPK Signaling | 7.37496068804798 |
| Signaling by NTRK1 (TRKA) | 6.496404322795111 |
| Nuclear Events (Kinase and Transcription Factor Activation) | 7.41953567709952 |
| PINK1-PRKN Mediated Mitophagy | 9.404689104510574 |
| NGF-stimulated Transcription | 8.665412485702259 |
| Chk1 Chk2(Cds1) Mediated Inactivation of Cyclin B Cdk1 Complex | 14.82898941521559 |
| WNT5A-dependent Internalization of FZD2, FZD5 and ROR2 | 14.82898941521559 |
| Glycosphingolipid Biosynthesis | 11.902794266378095 |
| Signaling by NTRKs | 5.838409362207258 |
| Signal Transduction | 3.9954762995019903 |
| Defective ST3GAL3 Causes MCT12 and EIEE15 | 19.226975207478883 |
| Inactivation of CDC42 and RAC1 | 19.226975207478883 |
| Vitamin C (Ascorbate) Metabolism | 19.226975207478883 |
| Clathrin-mediated Endocytosis | 5.357681088888031 |
| Diseases of Signal Transduction by Growth Factor Receptors and Second Messengers | 4.3877698576486655 |

| | |
|---|---|
| Apoptotic Factor-Mediated Response | 10.451639216310252 |
| Carnitine Shuttle | 12.361541862848469 |
| LGI-ADAM Interactions | 12.361541862848469 |
| Methylation | 12.361541862848469 |
| Repression of WNT Target Genes | 12.361541862848469 |
| CDC42 GTPase Cycle | 5.089484190838621 |
| Signaling by NOTCH3 | 6.828607395332858 |

### Table S9: AR Cluster 3 Reactome Pathways Top50

| Term | Combined Score |
|---|---|
| Signaling by Rho GTPases | 39.527516628914526 |
| Signaling by Rho GTPases, Miro GTPases and RHOBTB3 | 38.7193790966898 |
| RHO GTPase Cycle | 43.12724536914632 |
| Cell Cycle | 32.113253725577415 |
| Deubiquitination | 34.026881237768464 |
| Cell Cycle, Mitotic | 26.876910420178927 |
| M Phase | 27.201612765379107 |
| RHOA GTPase Cycle | 35.19811652188292 |
| Transcriptional Regulation by E2F6 | 71.80738558831669 |
| Deadenylation of mRNA | 91.78523639077926 |
| Post-translational Protein Modification | 17.18688673013308 |
| Membrane Trafficking | 19.369598273680506 |
| Chromatin Modifying Enzymes | 23.6965605343528 |
| Chromatin Organization | 23.6965605343528 |
| Cellular Responses to Stress | 17.37305563983505 |
| SUMOylation | 24.776065933030267 |
| Organelle Biogenesis and Maintenance | 20.874510733994203 |
| RAC1 GTPase Cycle | 23.60446108602172 |
| RAC2 GTPase Cycle | 30.837408813844615 |
| Cellular Responses to Stimuli | 15.79445084188004 |
| Cilium Assembly | 21.63052127553087 |
| M-decay Degradation of Maternal mRNAs by Maternally Stored Factors | 39.7877830392194 |
| Epigenetic Regulation by WDR5-containing Histone Modifying Complexes | 23.413960057560043 |
| Anchoring of the Basal Body to the Plasma Membrane | 26.211542795604355 |
| Metabolism of Proteins | 13.312513725889682 |
| Diseases of Signal Transduction by Growth Factor Receptors and Second Messengers | 16.351754425032897 |
| SUMO E3 Ligases SUMOylate Target Proteins | 20.565799370455426 |
| Cell Cycle Checkpoints | 17.813164277211197 |
| RHOC GTPase Cycle | 27.383371693995326 |
| RHOG GTPase Cycle | 27.383371693995326 |
| Epigenetic Regulation of Adipogenesis Genes by MLL3 and MLL4 Complexes | 22.80989108160996 |
| Epigenetic Regulation of Gene Expression by MLL3 and MLL4 Complexes | 22.80989108160996 |
| MLL4 and MLL3 Complexes Regulate Expression of PPARG Targets in Adipogenesis and Hepatic Steatosis | 22.80989108160996 |
| Ub-specific Processing Proteases | 18.385799758770364 |
| Loss of Nlp From Mitotic Centrosomes | 26.776130454064038 |
| Loss of Proteins Required for Interphase Microtubule Organization From the Centrosome | 26.776130454064038 |
| Antigen Processing Ubiquitination & Proteasome Degradation | 16.307739973567926 |
| RHOB GTPase Cycle | 25.196397065627764 |
| HATs Acetylate Histones | 20.88675208887487 |
| SUMOylation of Transcription Cofactors | 30.99414880136365 |
| Protein Ubiquitination | 23.105915666811583 |
| SUMOylation of DNA Damage Response and Repair Proteins | 23.105915666811583 |
| Regulation of PLK1 Activity at G2 M Transition | 21.353291940411438 |
| AURKA Activation by TPX2 | 22.350441479846225 |
| Signaling by TGFB Family Members | 16.625340007924724 |
| Class I MHC Mediated Antigen Processing & Presentation | 13.517698257607737 |
| Mitotic Prophase | 18.408922888481513 |
| Estrogen-dependent Gene Expression | 17.83088380906366 |
| CDC42 GTPase Cycle | 16.382733080262316 |

Table S9: AR Cluster 4 Reactome Pathways Top50

| Term | Combined Score |
|---|---|
| NOTCH1 Intracellular Domain Regulates Transcription | 28.592533692745214 |
| Signaling by NOTCH1 | 17.979778906343302 |
| Formation of Apoptosome | 49.77503608834571 |
| Regulation of the Apoptosome Activity | 49.77503608834571 |
| Heme Signaling | 16.692675043246663 |
| Cytochrome C-Mediated Apoptotic Response | 32.10414633294334 |
| PTK6 Regulates Proteins Involved in RNA Processing | 71.55929183201883 |
| Apoptotic Factor-Mediated Response | 19.8409471371314 |
| Activation of the AP-1 Family of Transcription Factors | 29.495945420448916 |
| Circadian Clock | 9.660500661644962 |
| Toll Like Receptor 9 (TLR9) Cascade | 8.096155189921172 |
| Neurexins and Neuroligins | 9.597015310738149 |
| Constitutive Signaling by NOTCH1 HD+PEST Domain Mutants | 9.110741783770871 |
| Constitutive Signaling by NOTCH1 PEST Domain Mutants | 9.110741783770871 |
| Signaling by NOTCH1 HD+PEST Domain Mutants in Cancer | 9.110741783770871 |
| Signaling by NOTCH1 PEST Domain Mutants in Cancer | 9.110741783770871 |
| Signaling by NOTCH1 in Cancer | 9.110741783770871 |
| Signaling by ALK | 11.681415277842751 |
| Specification of Primordial Germ Cells | 18.713384647074502 |
| HS-GAG Biosynthesis | 10.703997350728587 |
| Activation of RAC1 | 15.379392822929068 |
| TRAF6 Mediated Induction of NFkB and MAP Kinases Upon TLR7 8 or 9 Activation | 6.421897205078066 |
| Signaling by Non-Receptor Tyrosine Kinases | 7.49503780586092 |
| Signaling by PTK6 | 7.49503780586092 |
| MyD88 Dependent Cascade Initiated on Endosome | 6.203470837033598 |
| Regulation of Pyruvate Metabolism | 9.04271578406926 |
| MAP Kinase Activation | 7.072787063642087 |
| Protein-protein Interactions at Synapses | 6.157173365480445 |
| Interleukin-17 Signaling | 6.474808655501876 |
| Toll Like Receptor 7 8 (TLR7 8) Cascade | 5.406075688689598 |
| Prolactin Receptor Signaling | 10.838991174963645 |
| Regulation of Pyruvate Dehydrogenase (PDH) Complex | 10.838991174963645 |
| EGR2 and SOX10-mediated Initiation of Schwann Cell Myelination | 7.580244018205018 |
| NPAS4 Regulates Expression of Target Genes | 8.534985089200532 |
| HDR Through Single Strand Annealing (SSA) | 6.585286113509688 |
| MAP3K8 (TPL2)-dependent MAPK1 3 Activation | 9.249656699168739 |
| PI3K Events in ERBB2 Signaling | 9.249656699168739 |
| Toll-like Receptor Cascades | 4.16680101269826 |
| Acetylcholine Regulates Insulin Secretion | 11.634287814140869 |
| Bicarbonate Transporters | 11.634287814140869 |
| PI3K Events in ERBB4 Signaling | 11.634287814140869 |

| | |
|---|---|
| Defective B3GALTL Causes PpS | 6.104655091209944 |
| RHOB GTPase Cycle | 5.037708220437412 |
| MAPK Targets Nuclear Events Mediated by MAP Kinases | 6.324701086190873 |
| Transcriptional Regulation by MECP2 | 4.899492804849313 |
| O-glycosylation of TSR Domain-Containing Proteins | 5.66588736412853 |
| Synthesis of PA | 5.66588736412853 |
| Estrogen-dependent Nuclear Events Downstream of ESR-membrane Signaling | 6.811640771484876 |
| RUNX2 Regulates Osteoblast Differentiation | 6.811640771484876 |
| FOXO-mediated Transcription of Cell Cycle Genes | 7.961648111764821 |
